# Transcriptomic analyses in the gametophyte of *Dryopteris affinis*: apomixis and more

**DOI:** 10.1101/2024.03.15.585305

**Authors:** Sara Ojosnegros, José Manuel Alvarez, Valeria Gagliardini, Luis G. Quintanilla, Ueli Grossniklaus, Helena Fernández

## Abstract

The gametophyte of the fern *Dryopteris affinis* ssp. *affinis* represents a good model to explore the molecular basis of vegetative and reproductive development, as well as stress responses. Specifically, this fern reproduces asexually by apogamy, a peculiar case of apomixis whereby a sporophyte forms directly from a gametophytic cell without fertilization. Using an RNA-sequencing approach, we have previously annotated more than six thousand transcripts. Here, we selected one hundred of the inferred proteins that seemed particularly interesting for a detailed study of their potential functions, protein-protein interactions, and molecular phylogenies. As expected, a plethora of proteins associated with gametogenesis and embryogenesis in angiosperms, such as FERONIA (FER) and CHROMATING REMODELING 11 (CHR11) were identified, and more than a dozen candidates potentially involved in apomixis, such as ARGONAUTE4 (AGO4), AGO9, and AGO10, BABY BOOM (BBM), FASCIATED STEM4 (FAS4), FERTILIZATION-INDEPENDENT ENDOSPERM (FIE), and MATERNAL EFFECT EMBRYO ARREST29 (MEE29). In addition, proteins involved in the response to biotic and abiotic stresses were widely represented, as shown by the enrichment of heat-shock proteins. Using the String platform, studying interactomes revealed that most of the protein-protein interactions were predicted based on experimental, database, and text mining datasets, with MULTICOPY SUPPRESSOR OF IRA4 (MSI4) showing the highest number of 16 interactions. Lastly, some proteins were studied from a phylogenetic point of view, comparing the alignments with respect to more distantly or closely related plant groups, identifying AGO1 as the evolutionarily most similar to that other ferns and the most distant to the predicted common ancestor. This work sets the stage for future functional characterizations in relation to gametophyte development including apomictic reproduction.

## 1. Introduction

Land plants, i.e., vascular plants and bryophytes, have a biological cycle in which two generations alternate: the sporophyte, which forms spores by meiosis and is typically diploid, and the gametophyte, which forms gametes by meiosis and is usually haploid. In ferns, which is a monophyletic group within the vascular lineage (PPG I, 2016), gametophytes are photosynthetic individuals independent of sporophytes. In addition to sexual reproduction by fusion of gametes, ferns exhibit asexual reproduction by apomixis, which involves two processes: diplospory (generation of unreduced spores due to alteration of meiosis) and apogamy (formation of the embryo from somatic cells of the gametophyte) (Liu et al., 2012). Apomixis occurs when environmental conditions are unfavourable, e.g. lack of light or water (Grusz et al. 2021), or when gametes are non-functional. Apomixis is more prevalent in ferns than in any other group of land plants (Dyer et al., 2012). Specifically, it has been estimated that 10% of fern species have obligate apogamy (Liu et al., 2012). Because apomixis circumvents meiosis and fertilisation to form embryos, introducing this mode of asexual reproduction into sexual crops has long been one of the main challenges for plant breeders (Grossniklaus et al., 2001; Spillane et al., 2004). The molecular and developmental study of ferns may contribute to this purpose, as they are the lineage most closely related to seed plants.

Vegetative and reproductive development of ferns is underexplored compared with other plant groups. In this regard, the most studied aspects in ferns are: photomorphogenesis (Wada, 2007), spore germination (Salmi et al., 2005; Suo et al., 2015), cell polarity (Salmi and Bushart, 2010), cell wall composition (Eeckhout et al., 2014), reproduction (Kaźmierczak, 2010; Lopez and Renzaglia, 2014; Valledor et al., 2014; de Vries et al., 2016; Domzalska et al., 2017; Grossmann et al., 2017; Rivera et al., 2018; Wyder et al., 2020; Fernández et al., 2021; Ojosnegros et al., 2022; 2023), or adaptation to the environment (Gill et al., 2009; Thagela et al., 2016; Sareen et al., 2019). In particular, these studies show that ferns are able to survive atmospheric conditions with a higher amount of carbon dioxide than we have now, and to resist in situations of salinity, drought, or soil contamination with heavy metals (Rathinasabapathi, 2006; Wang et al., 2010). In addition, ferns are also very efficient at extracting pollutants from the environment (Dhir, 2018). Molecular studies on ferns are also scarce. In fact, they occupy one of the last positions in the race to decipher genome sequencing because of their high chromosome number and large genome size (Barker and Wolf, 2010). To our knowledge, the genomes of only four fern species are accessible, which are *Alsophila spinulosa, Azolla filiculoides*, *C. richardii*, and *Salvinia cucullata* (Aragón-Raygoza et al., 2022). Thanks to the advent of next-generation sequencing methodologies, more genomes are expected to be deciphered soon, as predicted by Kinosian and Wolf (2022).

The gametophyte of *Dryopteris affinis* (Lowe) Fraser-Jenk. ssp. *affinis*, (hereafter referred to as *D. affinis*) has been widely used as a model for understanding apomixis (Cordle et al. 2012; Rivera et al., 2018). This fern produces unreduced spores that give rise to diploid gametophytes. These gametophytes form male gametangia (antheridia) but not female gametangia (archegonia) (ref. específica de *D. affinis*).Thus, sexual reproduction is not feasible and *D. affinis* is obligately apogamous (Menéndez et al., 2006). For this asexual reproduction, the egg in the archegonium forms an embryo without fertilisation. Consequently, the resulting sporophyte is genetically identical to the gametophyte from which it originated, which in turn is identical to the sporophyte that formed the diplospore. In other words, the alternation of generations occurs without changes in the ploidy level or in the genetic composition of gametophytes and sporophytes. In addition to obligate apogamy, gametophytes of *D. affinis* have several advantages for the study of asexual reproduction, such as easy in vitro culture from spores, small (few mm^2^) size that allows large samples of individuals in a small culture space, and direct microscope observation of complete individuals since they are mostly constituted by a single layer of cells. (Rivera et al., 2018). In a recent study (Wyder et al., 2020), some of us performed a RNA analysis using next-generation sequencing (RNA-seq) of *D. affinis* gametophytes, which generated a wealthy transcriptome and showed differential gene expression between the two growth phases of the gametophyte: the initial one-dimensional and the subsequent two-dimensional. Here, we carry out an *in silico* study to analyse in more detail this transcriptome. Specifically, we address the genomic sequences corresponding to a set of around one hundred of proteins, which are described by function, as well as their interactions and phylogenetics. For ease of understanding and interpretation, the proteins are discussed by clustering them into three biological functions: vegetative development, reproductive development, and response to abiotic and biotic stresses.

## 2. Material and Methods

### 2.1. Plant Material and Growing Conditions

Fertile fronds from several sporophytes of *D. affinis* were collected in Turón forest (Asturias region, Spain, 43°12′10′′N−5°43′43′′W, 477 m a.s.l.). Fronds were carried to the lab, where spores were released from the sporangia, rinsed in water for 2 h and washed for 10 min with a NaCl (0.5%), and Tween 20 (0.1%) solution. Spores were then rinsed three times with distilled water, centrifuged at 1.300 g for 3 minutes between rinses, and cultured in 500 ml Erlenmeyer flasks containing 100 ml of Murashige and Skoog (MS) medium (Murashige and Skoog, 1962) supplemented with 2% sucrose (w/v) at pH 5.7.Then, gametophytes at two growth stages were collected for molecular analysis. On the one hand, gametophytes in the first growth stage (uni-dimensional filamentous gametophytes) were obtained by liquid culture of spores at high density, located in a rotating shaker (75 rpm) for 50 days. On the other hand, gametophytes at the following growth stages (two-dimensional: spatulate and heart-shaped gametophytes) were obtained by maintaining spores in Petri dishes with 25 ml of MS medium supplemented with 2% sucrose (w/v) and 0.7% agar at pH 5.7 for 65 days. All cultures were grown at 25 °C, and photosynthetically active radiation intensity 40 μmol m^-2^ s^-1^ with a photoperiod of 16 h light and 8 h dark.

### 2.2. RNA Extraction and Sequencing

The methods of RNA extraction and sequencing are summarized in Wyder et al. (2020). In brief, for RNA extraction, 100 mg of fresh plant material was weighed, frozen in liquid nitrogen and stored at -80 °C until use. Among 3-5 biological replicates of gametophytes of each growth stage were used for RNA-Seq. Gametophytes of each growth stages were homogenised with a Silamat S5 shaker (Ivoclar Vivadent, Schaan, Liechtenstein) twice for 10 s and 5 s, respectively. Total RNA was isolated using the SpectrumTM Plant Total RNA kit (Sigma-Aldrich, Buchs, Switzerland). After DNA removal with the TURBO DNA-free kit (LifeTechnologies, Carlsbad, USA), RNA quality was tested using the Bioanalyser Agilent RNA 6000 Pico Kit (Agilent Technologies, Waldbronn, Germany). Sequencing libraries were prepared with the TruSeq RNA Sample Prep Kit v2 and sequenced on the Illumina HiSeq 2000. Transcriptome Shotgun Assembly project is available in the European Nucleotide Archive (ENA) (http://www.ebi.ac.uk/ena) with the accession number PRJEB18522. The *de novo* transcriptome assembly in fasta format and the transcriptome annotation had been deposited in the Zenodo research data repository (www.zenodo.org) (https://doi.org/10.5281/zenodo.1040330). Highly similar contigs assembled by Trinity were further collapsed using Corset 0.93 (Davidson and Oshlack, 2014) with a distance threshold of 0.3, which resulted in 166,191 transcript clusters (with a minimum contig size of 201 nucleotides (nt)). BUSCO version 2.0.1 with the Embryophyta odb9 dataset to assess transcriptome integrity was used.

### 2.3. “In silico” Analysis and Protein Analysis Using the STRING Platform

An *in silico* analysis of the transcriptome previously obtained was carried out with the GENEIOUS PRIME software version 2023.2.1, by using Araport11 database, to increase the number of homologies or identities, and getting a more comprehensive mapping of the genomic background in the gametophyte of the apomictic fern *D. affinis*. Transcripts annotations with more than 450 bp and an E-value of less than 10^-20^ were selected. The identifiers of the genes were used as input for carrying out Gene Ontology (GO) pathways enrichment analysis, including the three categories: biological and molecular function, and cellular components; Kyoto Encyclopaedia of Genes and Genomes (KEGG), and also Smart protein domains, by using ShinyGO v0.741 platform. In addition, critical expressed pathways were analysed by String platform version 12.0, and a high threshold (0.700) was selected to infer possible function of transcripts and association between them. At this regard, protein-protein interactions can be of different channels: (a) experiments: proteins that have been shown to have chemical, physical, or genetic interactions in laboratory experiments; (b) data bases: protein interactions found in the same databases; (c) text mining: proteins mentioned in the same PubMed abstract or article from an internal selection of the STRING software; (d) co-expression: protein expression patterns are similar; (e) neighbourhood: protein-coding genes are close together in the genome; (f) gene fusion: at least in one organism orthologous protein-coding genes are fused into a single gene; (g) co-occurrence: proteins that have a similar phylogenetic distribution; and lastly, (h) homology: proteins that present a common ancestor and similar sequences.

### 2.4. Phylogenetic and Protein Domain Analysis

Phylogenetic analysis was carried out in some protein sequences using the NCBI Genome Workbench. For their construction, experimental databases, an expect threshold of 0.05, the BLOSUM62 matrix with gap costs of existence of 11 and extension of 1, the fast minimum evolution the minimum fast evolutionary tree method, a maximum sequence distance of 0.85, and the Grishin distance, were selected. In addition, only the first 50 sequences with the lowest E-values were chosen. To study protein domains, SMART software version 9.0 was used. Notice that a phylogenetic tree is a graphical representation of the evolutionary relationships between genes, proteins, or taxa. In it, each branch represents a sequence. The genetic distance or divergence between sequences can be measured by a horizontal scale bar, which indicates the number of substitutions that have occurred per site, e.g. a scale showing 0.1 means that there is a 10% difference between the two sequences studied.

## 3. Results

Our results group the transcripts obtained from one-dimensional and two-dimensional gametophytes, in order to obtain a more complete transcriptome of the gametophyte of the apogamous fern *D. affinis*. Then, the transcriptomic data were treated with GENIOUS PRIME software, and yielded 27,037 transcripts corresponding to 6,142 protein annotations.

### 3.1. Protein enrichment classifications

Based on the GO classification of biological function (**Fig. 1a**), most of the proteins are dedicated to those events that organise and regulate cellular functioning, such as biosynthesis and catabolism of proteins, lipids, phosphate-containing compounds, vitamins, nucleic acids, and less intricate molecules such as tetrapyrrole, a precursor of chlorophyll biosynthesis. The second group in the ranking of biological functions were those processes associated with localization and transport, cell division, and gene expression, including protein annotations related to meiosis, chromosome organization, non-coding RNA metabolism, homologous recombination, and post-transcriptional regulation of gene expression. In addition, among the biological functions, it is response to stress induced by environmental factors such as temperature or radiation, and the resulting DNA damage, as well as alterations in the signalling of growth regulators such as abscisic acid, stand out. The categories with the highest number of genes were nucleobase-containing compound metabolic process, response to abiotic stimulus, and development. It is noted that in organelle organisation and carboxylic acid metabolic process the fold enrichment or gene ratio, i.e. the number of genes present over the total number of genes in that category, was higher, and in development and response to abiotic stimuli, the fold enrichment was lower. Regarding molecular function (**Fig. 1b**), RNA binding and oxidoreductase and transferase activities were the most represented functions in our gametophytes, with the highest number of genes found, followed by others such as transporter and ATP-dependent activities. The fold enrichment value was highest in the ATPase-coupled transmembrane transporter and ATP hydrolysis activities, and was lower in the oxidoreductase and transferase categories. In terms of cellular components (**Fig. 1c**), disregarding the organelle envelope and non-membrane-bounded organelle categories, the gametophyte proteins were mostly located in three organelles: mitochondrion, chloroplast, and nucleus. It was observed that the fold enrichment or gen ratio was higher in the different parts of chloroplasts such as stroma, envelope, or thylakoid, and lower in mitochondrion. As for the protein domains obtained from the SMART software (**Fig. 1d**), the highest number corresponded to ATPases associated with a variety of cellular activities, and catalytic domains of serine/threonine protein kinases, followed by WD40 and leucine-rich repeats. Regarding fold enrichment, the highest values corresponded to minichromosome maintenance proteins and actins, while the lowest values corresponded to EF-hand calcium binding motifs and leucine rich repeats.

**Figure 1.**
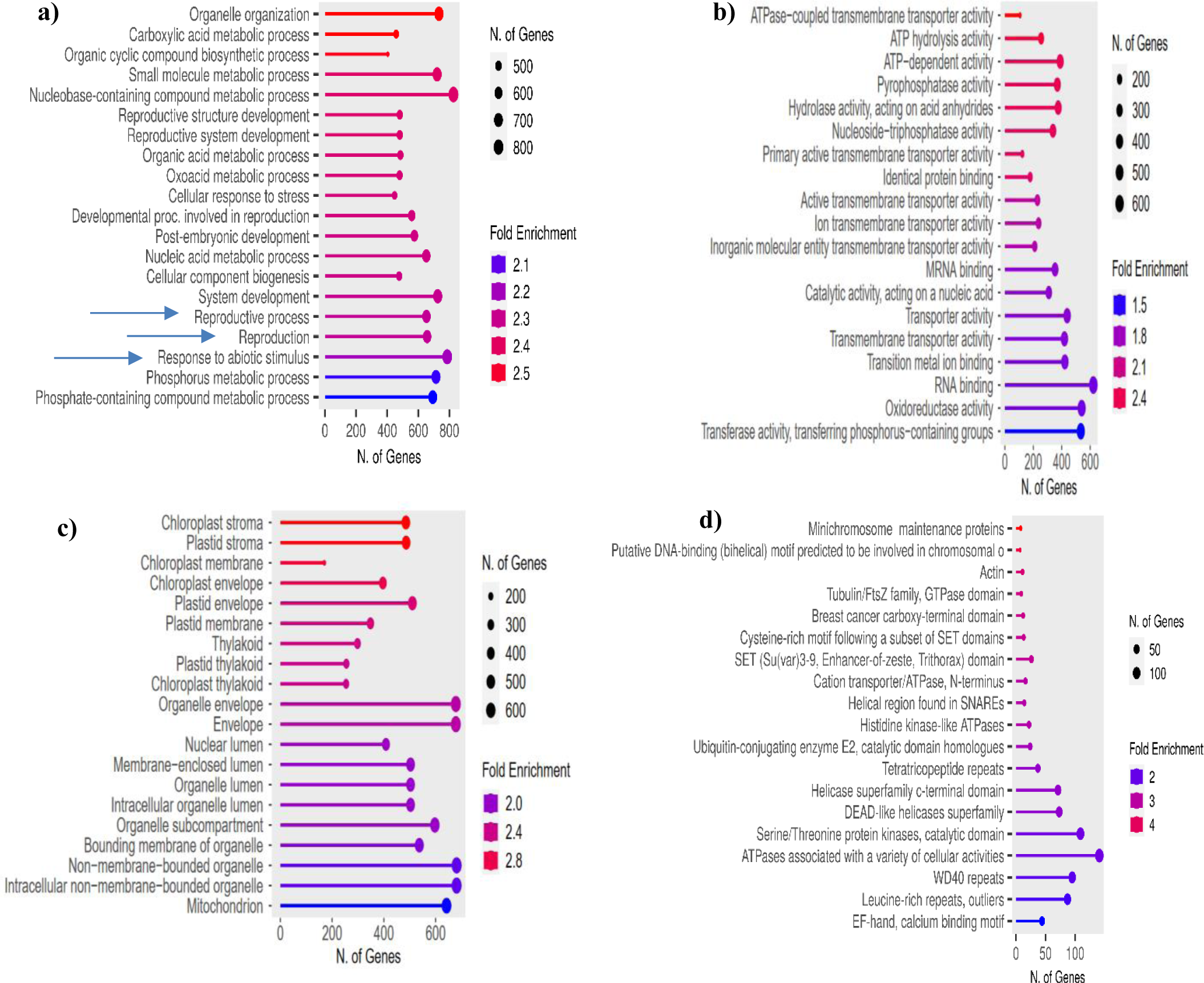
Classifications of proteins extracted from gametophytes of the fern *D. affinis* ssp*. affinis,* analysed with ShinyGO program. The size of the balls indicates the number of genes in each category. The fold enrichment is represented by a colour spectrum: blue shades indicate low values, while red shades indicate high values. **a)** GO biological function. Blue arrows indicate processes involving proteins explained in detail in the discussion section. **b)** GO molecular function. **c)** GO cellular component. **d)** SMART protein domains.

Likewise, the information of KEGG classification (**Fig. 2**) revealed that the three most abundant categories were related to processes of primary and secondary metabolism, highlighting the biosynthesis of secondary metabolites, and followed by carbon metabolism and biosynthesis of cofactors. Analysing all categories, the fold enrichment or gene ratio was highest in DNA replication, carbon fixation in photosynthetic organisms, and ascorbate and aldarate metabolism. In contrast, this value was lowest in the spliceosome and biosynthesis of secondary metabolites, as well as in glycolysis and gluconeogenesis.

**Figure 2.**
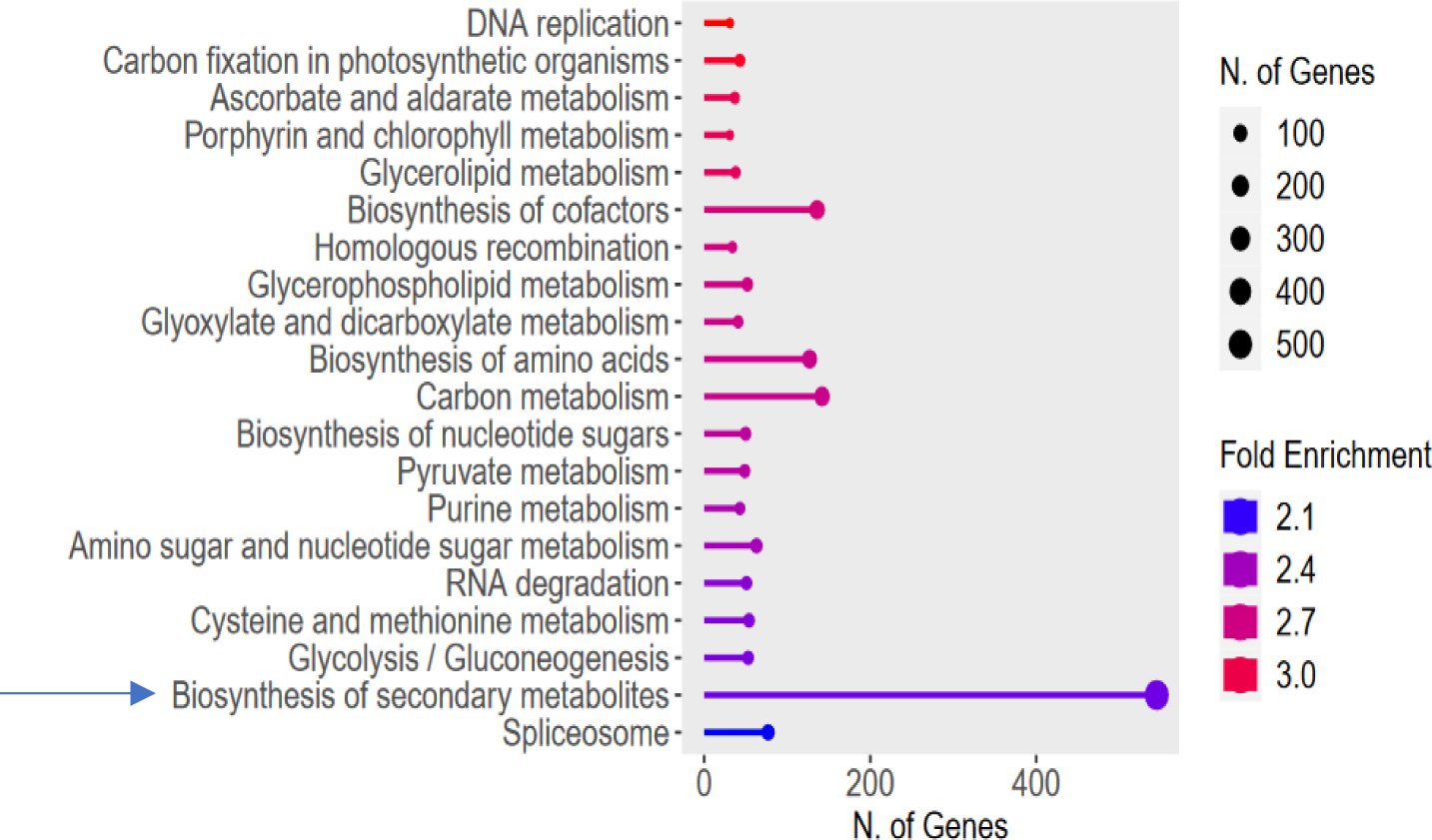
KEGG classification of proteins extracted from gametophytes of the fern *D. affinis* ssp. *affinis* analysed with ShinyGO program. The size of the balls indicates the number of genes in each category. The fold enrichment is represented by a colour spectrum: blue shades indicate low values, while red shades indicate high values. The blue arrow shows the large number of genes of biosynthesis of secondary metabolites found in gametophytes.

### 3.2. Proteins associated with apomixis

As expected, numerous proteins linked to reproduction, and specifically to apomixis were identified from the gametophytes (**Table 1**). Among them, there are members of the DEAD box helicase family, such as FASCIATED STEM 4 (FAS4), essential in male and female gametogenesis, MATERNAL EFFECT EMBRYO ARREST 29 (MEE29), needed in embryogenesis; SWITCH1 (*SW1*), involved in female gametogenesis; BRI1-ASSOCIATED RECEPTOR KINASE (BAK1), and the SERK family, required in somatic embryogenesis. Regarding the WD-40 repeat family, the protein SLOW WALKER1 (SWA1), related to megagametogenesis, and MULTICOPY SUPPRESSOR OF IRA1 (MSI1), whose mutants show parthenogenetic development, were found. Likewise, a member of the WD repeat ESC family was described: FERTILIZATION INDEPENDENT ENDOSPERM (FIE), whose loss of function mutants also are associated with parthenogenesis. BABY BOOM (BBM) is another protein related to apomixis, which promotes the reproductive processes of parthenogenesis, somatic embryogenesis and apogamy, and belongs to the AP2/ERF family. From the MADS-box family, the following two proteins were detected: AGAMOUS-LIKE 62 (AGL62) and 6 (AGL6), required in endosperm development and megasporogenesis, respectively. Three ARGONAUTE family proteins connected to apomixis were identified: ARGONAUTE 4 (AGO4), 9 (AGO9), and 10 (AGO10), all of them involved in gene silencing, and whose mutants exhibit apospory. Finally, it was detected SERRATE (SE), from the ARS2 family, and which is also important in gene silencing.

**Table 1.**
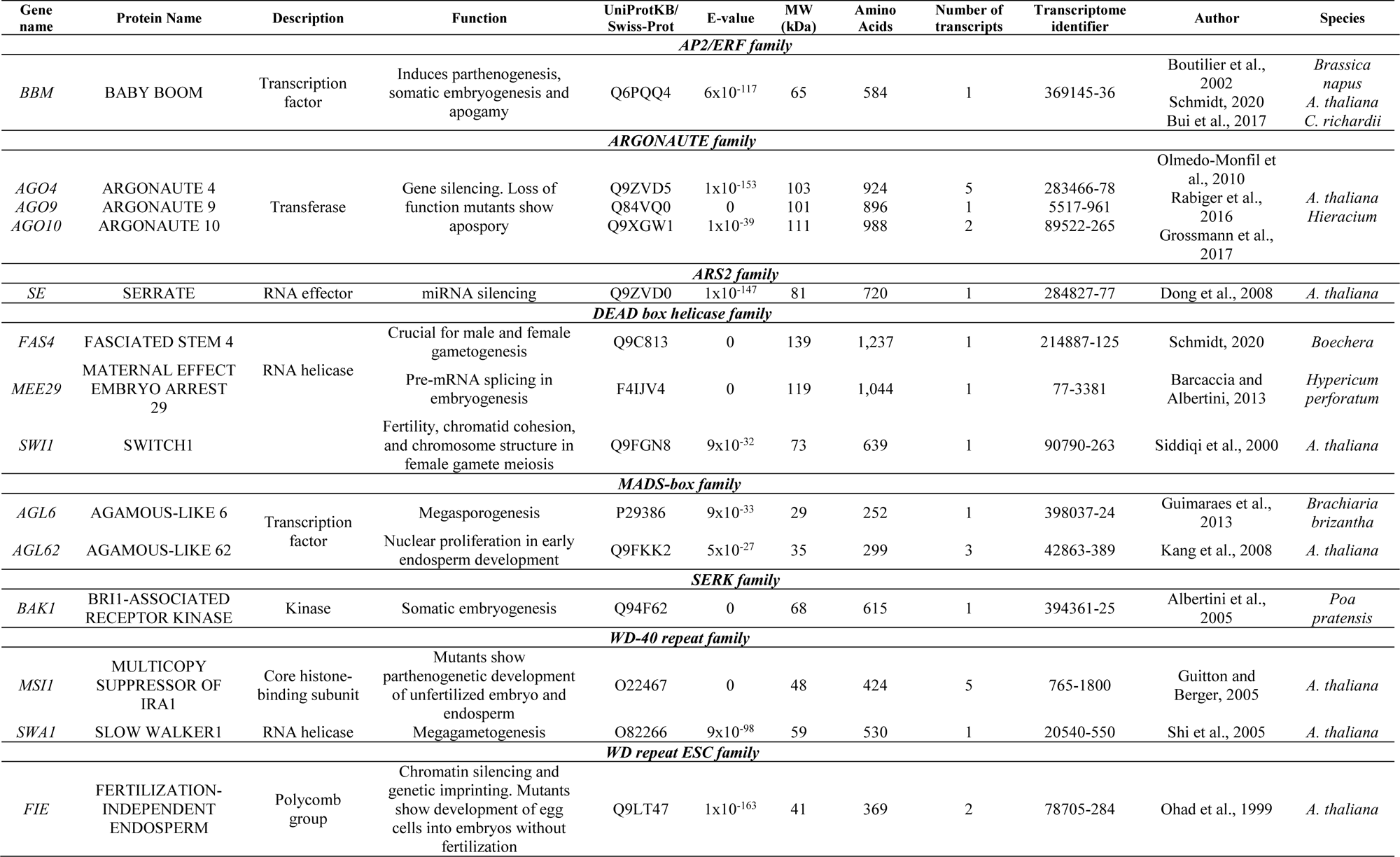
Proteins obtained from the gametophyte of the fern *Dryopteris affinis* ssp*. affinis* associated with apomixis.

Most of the proteins related to apomixis connect to relevant molecular processes operating behind gametogenesis or embryogenesis, such as gene silencing by RNA, splicing, chromatin remodelling, and methylation, among others. The large assortment of proteins related to these processes found in the *D. affinis* transcriptome can be observed in **Fig. 3**. This is the case of the ARGONAUTE family, represented here by five of the ten members that compose it in *A. thaliana*: ARGONAUTE 1 (AGO1), 4 (AGO4), 7 (AGO7), 9 (AGO9) and 10 (AGO10), associated with gene silencing during the transcription and post-transcription, as well as SERRATE (SE), implicated in control by gene silencing of the meristem activity and leaf polarity (**Fig. 3a**). The list includes also proteins involved in splicing, process otherwise reporting at least 33 entries linked to the maternal effect embryo arrest protein group (MEE), like MATERNAL EFFECT EMBRYO ARREST 29 (*MEE29*) (**Fig. 3b**). The protein cluster of embryo sac development arrest (EDA) is represented by 24 members in our dataset, signified here for three annotations: SLOW WALKER 1 (SWA1), 2 (SWA2) and 3 (SWA3), being some of them involved on megagametogenesis (**Fig. 3c**). It is striking the number of proteins found of certain groups such as helicases (**Fig. 3d**) and agamous-like, exemplified by AGAMOUS-LIKE 62 (AGL62) and 6 (AGL6) (**Fig. 3e**). Likewise, relevant genomic processes, such as chromatin remodelling, counts several proteins (**Fig. 3f**). That is the case also for the proteins belonging to the mediator complex, which regulates transcription, (**Fig. 3g**). Finally, proteins operating in methylation are copious, such as FERTILIZATION-INDEPENDENT ENDOSPERM (FIE), belonging to the Polycomb group, and which represses the development of embryo and endosperm without fertilization (**Fig. 3h**). Protein kinases were very abundant in the fern transcriptome, with a total of 345 (5.6% of all proteins obtained). The presence of 250 transcripts corresponding to embryo defective proteins (EMB) was also a remarkable result (**Fig. 4**).

**Figure 3.**
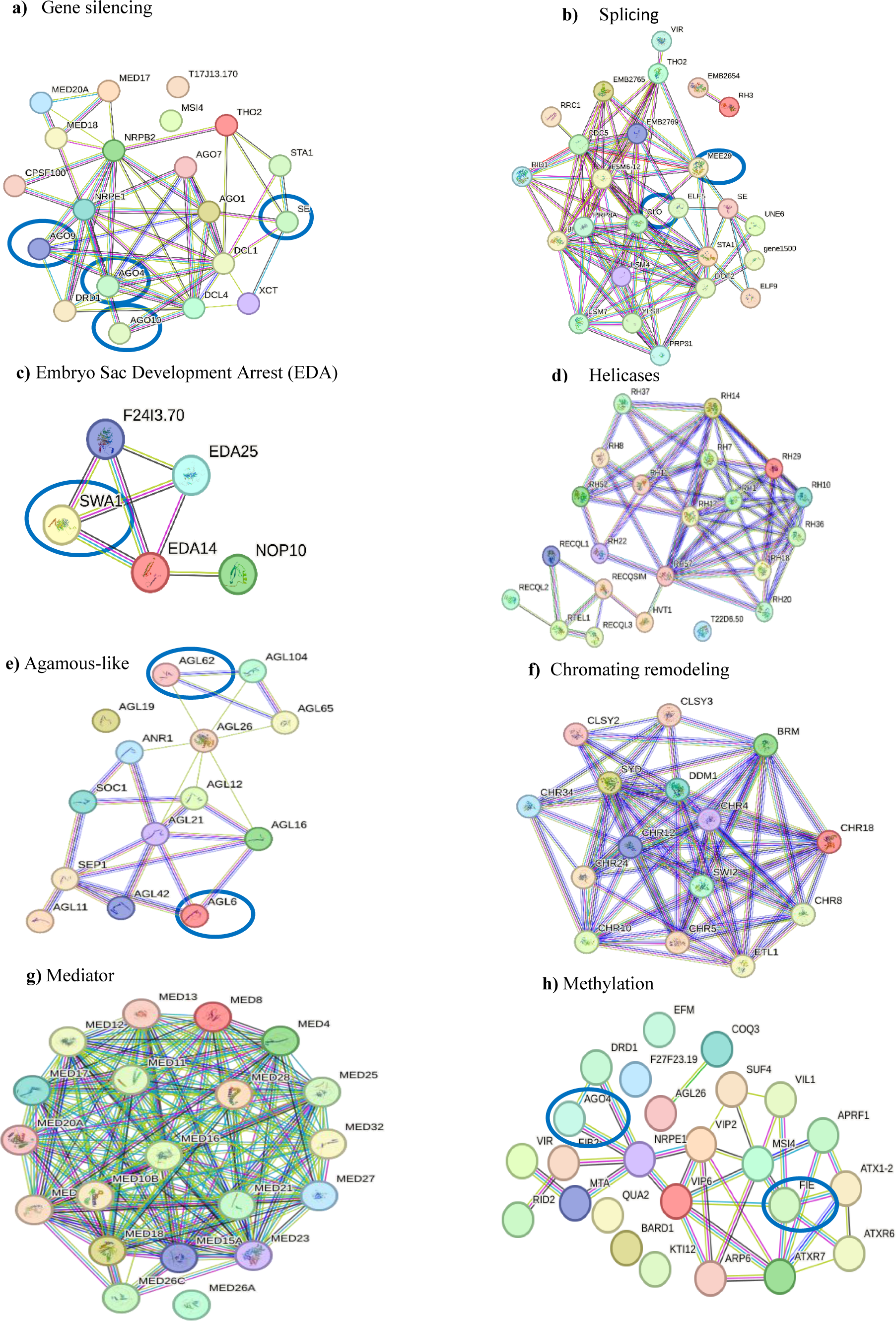
Networks formed by protein clusters interacting with apomictic candidate genes, marked with a blue circle, obtained from gametophytes of *Dryopteris affinis* ssp. *affinis*, provided by String program. Colour indicates the type of protein-protein interaction: pink lines refer to evidence from experiments, yellow lines from text mining, black lines from co-expression, and blue lines from databases.

**Figure 4.**
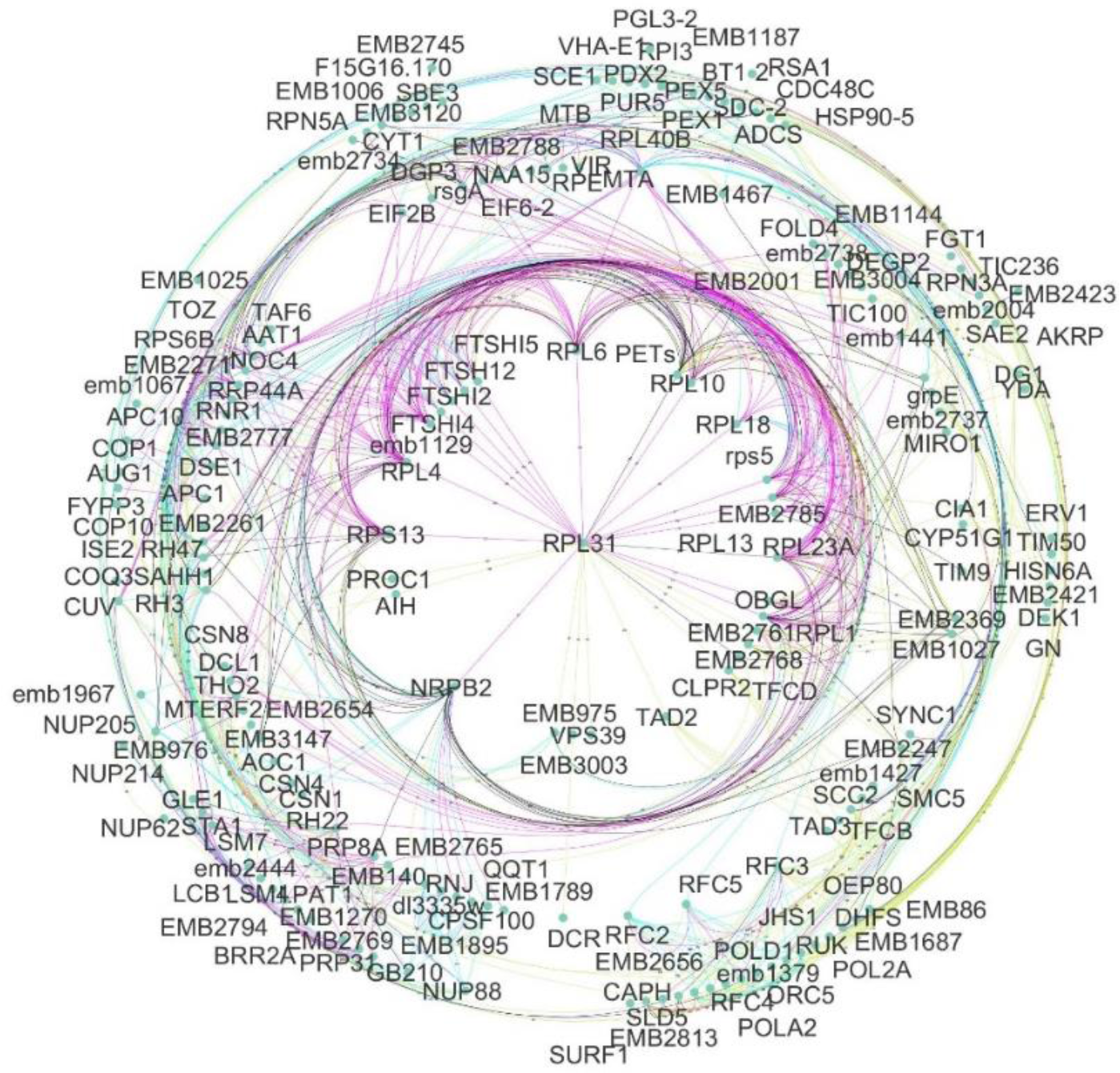
Embryo defective proteins annotated from gametophytes of *Dryopteris affinis* ssp. *affinis* provided by String program and edited with Cytoscape software. Colour indicates the type of protein-protein interaction: pink lines refer to evidence from experiments, yellow lines from text mining, black lines from co-expression, and blue lines from databases.

### 3.3. Proteins associated with vegetative and sexual development, and response to stress

A high number of proteins related to vegetative and reproductive development, as well as response to stress were identified in the *D. affinis* transcriptome (**Fig. 5**). Concerning the development of these anatomical structures, many orthologs were found, such as the protein HISTONE CHAPERONE HIRA (HIRA), implicated in gene silencing during leaf development, or ABERRANT LATERAL ROOT FORMATION 4 (ALF4), in the formation of roots independent of auxin (**Fig. 5a**), and others, such as SHOOT MERISTEMLESS (STM), which participates in shoot apical meristem formation during the embryogenesis (**Fig. 5b**). As with any other plant organisms, the gametophyte development demands the coordinated action of the main plant growth regulators, such as auxins, cytokinins, ethylene, brassinosteroids, jasmonic acid, abscisic acid, and gibberellins, and proteins involved in their metabolism, transport and signalling were found (**Fig. 5c**). Given the fact that the gametophyte is committed to cope with reproduction, it was expected to find plentiful proteins associated to gametogenesis or embryogenesis, some of them already mentioned to be associated to apomixis, and many others not involved like WUSCHEL RELATED HOMEOBOX 8 (WOX8).

**Figure 5.**
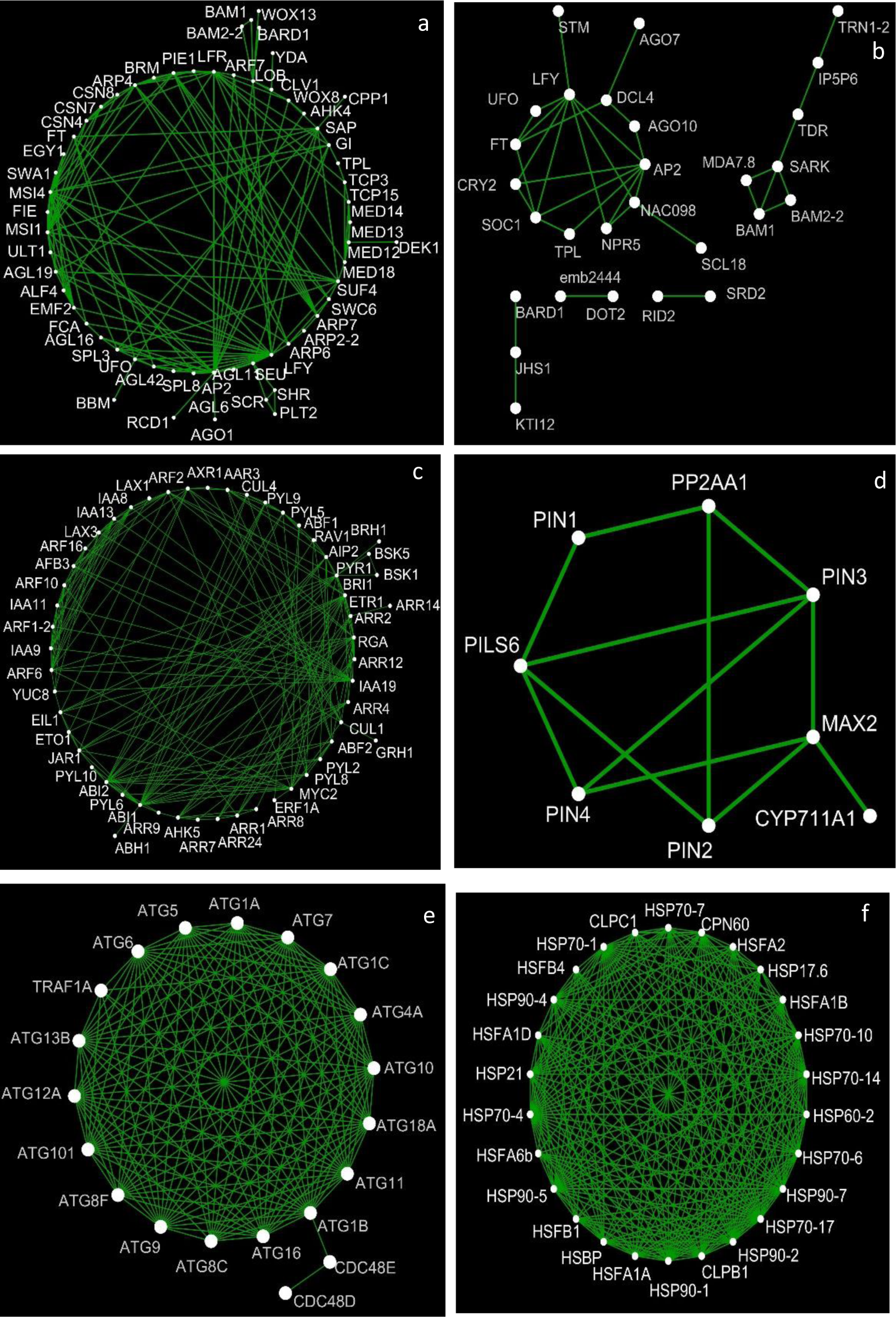
Networks of interactions between proteins obtained from the gametophyte of *D. affinis* ssp. *affinis* related to: **a)** vegetative and reproductive growth; **b)** meristem development; **c)** hormone signalling; **d)** auxin transport; **e)** autophagy, and **f)** heat shock protein family, provided by String program and edited with Cytoscape software.

Otherwise, although ferns do not form flowers, numerous proteins such as FLOWERING PROMOTING FACTOR 1 (FPF1), essential in the transition to flowering, were found, as well as proteins related to the circadian clock, like TIMEKEEPER LOCUS1 (STIPL1). The presence of numerous genes related to autophagy was also striking (**Fig. 5d**). Regarding abiotic stress, proteins coping with cold or heat were identified, such as ABA DEFICIENT 3 (ABA3), as well as several members of heat-shock proteins (**Fig. 5e**). Examples of proteins related to biotic stress were DICER-LIKE 4 (DCL4) and PRIORITY IN SWEET LIFE 4 (PSL4), intervening in defence against viruses and bacteria, respectively. All these proteins are a small sample that will be expanded later in the discussion section.

Altogether the proteins detailed in this work, involved in apomixis, vegetative development, sexual reproduction, and response to biotic and abiotic stress, formed a network composed of 120 nodes and 189 edges with a PPI enrichment p-value less than 1·10^-16^. This network had an average node degree of 3.15, an average local clustering coefficient of 0.446, and an expected number of edges of 81. The p-value obtained indicates that the network has significantly more interactions than expected, i.e., the proteins have more interactions with each other than would be expected for a random set of proteins of the same size and degree distribution extracted from the genome. Within this group, the proteins with the highest number of interactions are MULTICOPY SUPPRESSOR OF IRA 4 (MSI4) with 16 interactions, MULTICOPY SUPPRESSOR OF IRA 1 (MSI1) and SLOW WALKER1 (SWA1) with 12 interactions each, and FASCIATED STEM 4 (FAS4) with 11 interactions. Table 2 displays the scores for different protein interactions, emphasizing those with a total score higher than 0.9.

**Table 2.** Protein-protein interactions of the selected proteins from the gametophyte of *Dryopteris affinis* ssp. *affinis* with the highest total scores.

| Protein 1 | Protein 2 | Neighborhood Interaction | Gene Fusion Interaction | Co-expression Interaction | Experiments Interaction | Data Base Interaction | Text Mining Interaction | Total Score |
| --- | --- | --- | --- | --- | --- | --- | --- | --- |
| CUL4 | DET1 | 0 | 0 | 0.046 | 0.757 | 0 | 0.998 | 0.999 |
| FIE | MSI1 | 0 | 0 | 0.075 | 0.939 | 0.958 | 0.997 | 0.999 |
| CUL4 | DDB2 | 0 | 0 | 0 | 0.687 | 0.841 | 0.971 | 0.998 |
| DDB2 | DET1 | 0 | 0 | 0 | 0 | 0.900 | 0.913 | 0.990 |
| AGO1 | DCL4 | 0 | 0 | 0.063 | 0.410 | 0.313 | 0.975 | 0.989 |
| FIE | VIN3 | 0 | 0 | 0 | 0.050 | 0.900 | 0.859 | 0.985 |
| MSI1 | VIN3 | 0 | 0 | 0 | 0.058 | 0.900 | 0.862 | 0.985 |
| FAS4 | RH36 | 0 | 0 | 0.276 | 0.114 | 0.932 | 0.694 | 0.984 |
| FIE | MSI4 | 0 | 0 | 0.052 | 0.840 | 0.809 | 0.508 | 0.983 |
| AGO7 | DCL4 | 0 | 0 | 0.077 | 0.410 | 0.313 | 0.956 | 0.981 |
| CUL4 | MSI4 | 0 | 0 | 0.048 | 0.114 | 0.844 | 0.865 | 0.979 |
| MBD9 | TAF14B | 0 | 0 | 0 | 0 | 0.900 | 0.795 | 0.978 |
| CUL4 | MSI1 | 0 | 0 | 0.042 | 0.345 | 0.844 | 0.788 | 0.976 |
| RH36 | SWA1 | 0 | 0.005 | 0.349 | 0.062 | 0.808 | 0.808 | 0.974 |
| CDC5 | CLO | 0 | 0 | 0.128 | 0.937 | 0 | 0.538 | 0.972 |
| AGO4 | DCL4 | 0 | 0 | 0.058 | 0.410 | 0.313 | 0.934 | 0.971 |
| AGO10 | DCL4 | 0 | 0 | 0.056 | 0.410 | 0.313 | 0.931 | 0.970 |
| DET1 | HY5 | 0 | 0 | 0 | 0.280 | 0 | 0.951 | 0.963 |
| ALF4 | SWA1 | 0 | 0 | 0.500 | 0.776 | 0.560 | 0.224 | 0.956 |
| FAS4 | SWA1 | 0 | 0 | 0.285 | 0.901 | 0.350 | 0.083 | 0.952 |
| CERK | LCB2a | 0.107 | 0 | 0 | 0.062 | 0 | 0.931 | 0.937 |
| HOS1 | MSI4 | 0 | 0 | 0.050 | 0.510 | 0 | 0.874 | 0.936 |
| CDC5 | MEE29 | 0 | 0.009 | 0.081 | 0.889 | 0.087 | 0.362 | 0.932 |
| EDA25 | SWA1 | 0 | 0 | 0.388 | 0.056 | 0 | 0.892 | 0.932 |
| AGO9 | DCL4 | 0 | 0 | 0.052 | 0.410 | 0.313 | 0.837 | 0.929 |
| AGO1 | HST1 | 0 | 0 | 0.070 | 0.047 | 0.071 | 0.913 | 0.918 |
| SE | XCT | 0 | 0 | 0.069 | 0 | 0.911 | 0 | 0.913 |
| EDA25 | FAS4 | 0 | 0 | 0.911 | 0.049 | 0 | 0 | 0.912 |
| EIN2 | ETO1 | 0 | 0 | 0.065 | 0 | 0 | 0.909 | 0.911 |
| CUL4 | HY5 | 0 | 0 | 0 | 0 | 0 | 0.910 | 0.910 |
| ALF4 | FAS4 | 0 | 0 | 0.526 | 0.557 | 0.592 | 0.065 | 0.909 |

### 3.4. Protein-protein interactions

Regarding all the protein-protein interactions obtained from our dataset, only a few data on neighbourhood, gene fusion, co-occurrence and homology channels were observed. The proteins FATTY ACID AMIDE HYDROLASE (FAAH) and PYRIMIDINE 2 (PYD2) highlight in the neighbourhood interaction; CELL DIVISION CYCLE 5 (CDC5) and MATERNAL EFFECT EMBRYO ARREST 29 (MEE29) in gene fusion; and HASTY (HST) and SERRATE (SE) in co-occurrence; while HOMEOBOX PROTEIN KNOTTED-1-LIKE1 and SHOOT MERISTEMLESS (STM) in homology. Co-expression was low, with SLOW WALKER2 (SWA2) and FASCIATED STEM 4 (FAS4) having the strongest interaction. The highest scores came from the experiments, the database and the textmining channel. In experiments and database, FERTILIZATION-INDEPENDENT ENDOSPERM (FIE) and MULTICOPY SUPPRESSOR OF IRA 1 (MSI1) had the strongest interaction, while CULLIN 4 (CUL4) and DE-ETIOLATED 1 (*DET1*) showed the strongest interaction in text mining.

A molecular phylogenetic approach is included in this work, to explore the degree of genomic proximity existing between the proteins of our species and other land plants (**Fig. 6**). For that purpose, the phylogenetic trees of the amino acid sequences of some proteins were scrutinized. Specifically, the trees of the proteins ARGONAUTE 1 (AGO1) and WUS-INTERACTING PROTEIN 1 (TPL) from sexual reproduction, BABY BOOM (BBM) from apomixis, and DICER-LIKE 4 (DCL4) from response to biotic stress, are shown in the **Fig. 6a**, while the rest can be found in the **Supplementary Fig. 1**. Taking this value into account, in AGO1, the sequences of the species *Adiantum nelumboides*, *A. capillus-veneris*, and *C. richardii* are the closest to our fern sequence. The sequence of *D. affinis* originates from the same close common ancestor of the sequences of *A. nelumboides* and *A. capillus-veneris*, as they all have the same clade node. However, for BBM, the sequence of the *A. nelumboides* species is the most distant from our fern sequence. As for the third tree, the sequence of TPL in seed plants and in *Amborella trichopoda* are the closest to our fern sequence. Finally, in DCL4, there is more evolutionary distance between *D. affinis* sequence and the rest of plants studied, such as *Spirodela intermedia*. Comparing the fern sequences of the four trees, it is observed that the first protein, AGO1 is evolutionarily further to the common ancestor in each group of sequences, and has diverged more. From the phylogenetic trees shown in **Supplementary Fig. 1**, we want to highlight the proteins CULLIN 4 (CUL4) from photomorphogenesis, TIMEKEEPER LOCUS1 (STIPL1) from circadian clock, UBIQUITIN-SPECIFIC PROTEASE 26 (UBP26) from embryogenesis, and UNUSUAL FLORAL ORGANS (UFO) from flowering. In the CUL4 and STIPL1 proteins, our fern sequences are more separated from the rest of the sequences studied, such as those of the *A. trichopoda* species. The same occurs with the UBP26 protein, for which there is more evolutionary distance between our fern sequence and the rest of the species analysed. Lastly, as for the UFO protein, our *D. affinis* sequence is evolutionarily similar to the sequences of seed plants and bryophytes, showing more differences with those of other plants, such as *A. trichopoda*. Comparing the fern sequences of the eight trees, it is observed that the first protein, AGO1, is evolutionarily further to the common ancestor in each group of sequences, and has evolved more, showing 52% differences from the original sequence. Amino acid sequence alignments were also obtained with the same software and discussed later on (**Fig. 6b and Supplementary Fig. 2**).

**Figure 6.**
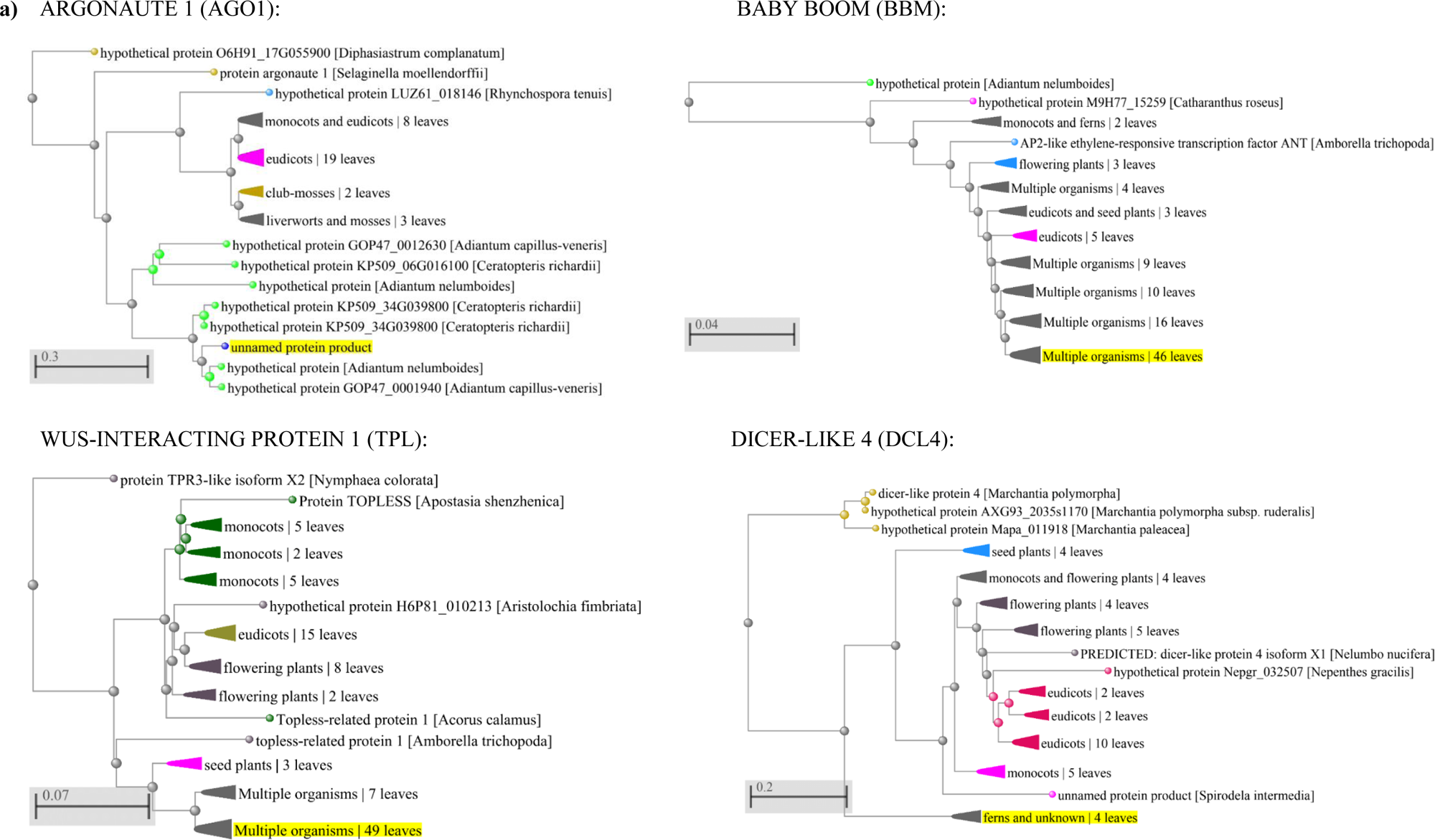
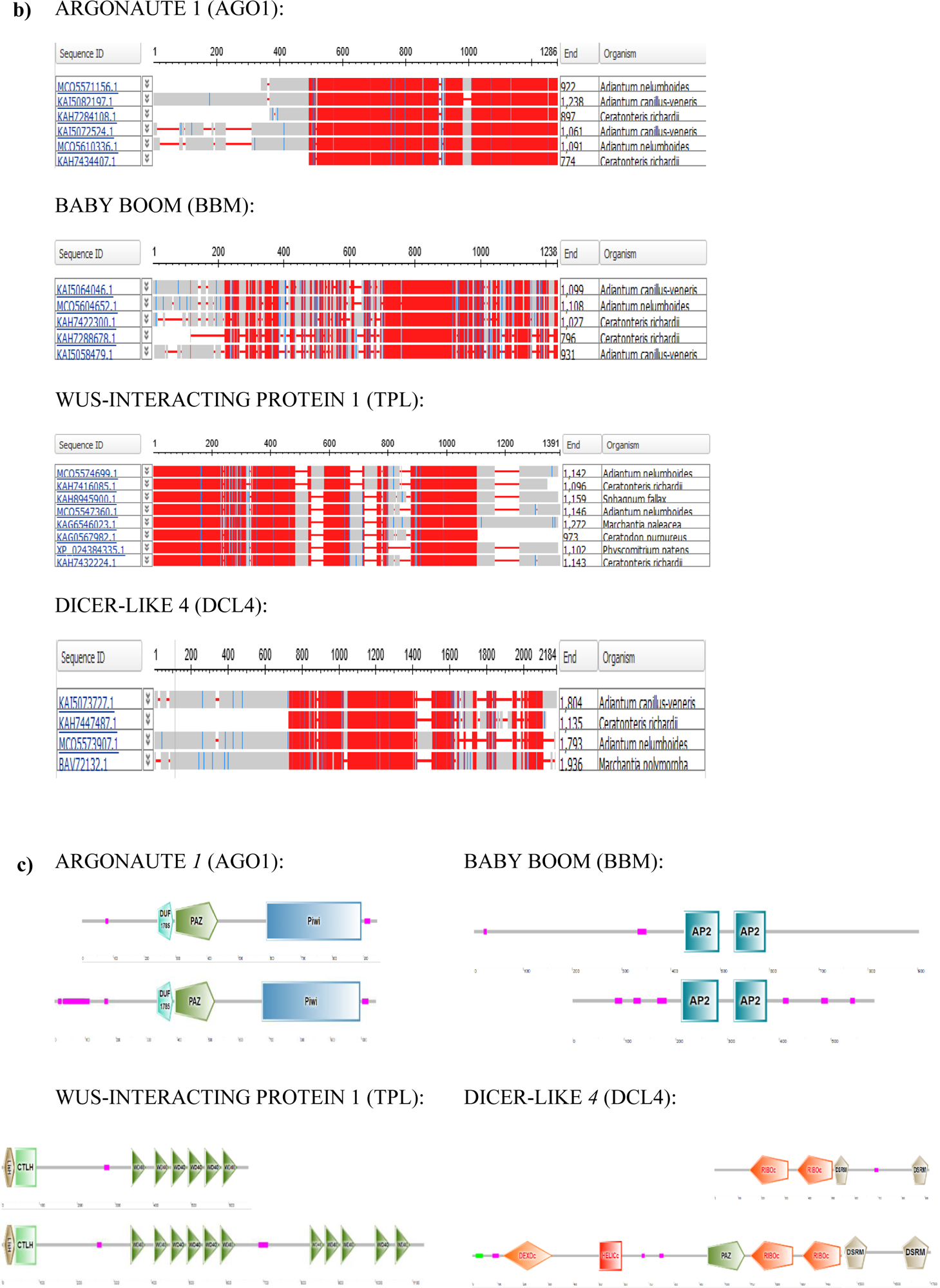
Phylogenetic analyses of proteins obtained from the gametophyte of the fern *D. affinis* ssp. *affinis*. **a)** Trees of amino acid sequences of selected proteins, performed by using NCBI Genome Workbench and NCBI BLASTP. *D. affinis* sequence is highlighted in yellow. Distance scale represents the number of differences between sequences (0.1=10%). **b)** Amino acid sequence alignments using the column-based method in NCBI Genome Workbench and NCBI BLASTP. Red indicates highly conserved columns and blue indicates less conserved columns. **c)** Domains of selected proteins from gametophytes of the fern *D. affinis* above and from *A. thaliana* below provided by SMART software.

Another aspect related to the evolution of protein sequences over time in different species is their protein domains. The domains of the proteins selected above for the phylogenetic study were analysed with SMART software (**Fig. 6c**) and the rest is shown in **Supplementary Fig. 3**. Comparing with those of *A. thaliana*, our data revealed that in the proteins AGO1 and BBM the same number and type of domains are conserved in both species: the DUF1785, Paz and Piwi domains in the first and the AP2 domain in the second. As for the other proteins, the sequences of TPL and DCL4 in *A. thaliana* have other types of domains and more repeats of some domains than those in the fern sequence. The first protein in both species has the LisH, CTLH and WD40 domains, while the second has in both species the RIBOc and DSRM domains, but in *A. thaliana* in addition there are the DEXDc, HELICc and Paz domains. The proteins CULLIN 4 (CUL4), TIMEKEEPER LOCUS1 (STIPL1), UBIQUITIN-SPECIFIC PROTEASE 26 (UBP26), and UNUSUAL FLORAL ORGANS (UFO) have the same number and type of domains in the two species studied: *D. affinis* and *A. thaliana*. CUL4 has a CULLIN and a Cullin Nedd8 domains, STIPL1 a G patch domain, and UBP26 a DUSP domain, while UFO has a FBOX domain.

## 4. Discussion

This work provides detailed information about the transcriptomic background yielded from gametophytes of the fern *D. affinis,* contributing to swell the list of information collected to date in non-model species. Our results open a new avenue for investigating apomixis in ferns and non-model species, as most previous reports have primarily focused on seed plants. The gametophyte of ferns is the phase of life cycle committed to cope with reproduction, and due to the lack of archegonia, *D. affinis* reproduces only by apogamy, i.e. it is an obligate apogamic fern, and as expected, numerous proteins linked to reproduction, and specifically to apomixis were identified. Therefore, this fern could serve as a valid model organism to deepen into this type of asexual reproduction. It provides an opportunity to speculate about the intricate molecular mechanisms that are still not well understood and to harness its potential in the future, particularly with economically valuable plants.

Likewise, the research performed enlarges our previous studies on this and other ferns, such as *Struthiopteris spicant* and *Dryopteris oreades*, focusing on the molecular basis of reproduction in the gametophyte of this group of plants (Valledor et al., 2014; Grossmann et al., 2017; Wyder et al., 2020; Fernandez et al., 2021; Ojosnegros et al., 2022; 2023). In this study, we specifically examine the transcriptome of the gametophyte, noting its ease of cultivation in a laboratory setting, which allows for the collection of sufficient material to conduct molecular analyses. According to a previous study (Wyder et al., 2020), a collection of 436,707 transcripts was obtained after assembled the RNA-seq reads, a higher number than in other species of ferns, such as *Adiantum flabellulatum*, with 354,228 transcripts (Cai et al., 2022); or *Vandenboschia speciosa*, with 36,430 (Bakkali et al., 2021), or 43,139 (Martín-Blazquez et al., 2023). All these studies contribute to enhancing our genomic understanding of unsequenced fern species, which have remained elusive until recently. This will enable us to gain a clearer picture of the evolution of land plant genomes in the future. Given that the *D. affinis* genome is not sequenced yet, the transcripts above mentioned were blasted (BLASTX) against the model species *A. thaliana*, being filtered those sequences that have an E-value less than 10^-20^. The obtained dataset revealed a total of 6,142 annotated proteins, which were analysed *in silico,* with the support of the GENEIOUS PRIME, ShinyGO, and STRING platforms. Approximately one hundred proteins were studied in more detail, and it represents the core of this discussion (sequences are shown in the **Supplementary Table 1)**. The compiled information has been organised and discussed turning around vegetative and reproductive development, and response to abiotic and biotic stress (**Table 3**), thus, showing the biological function of proteins, and envisaging a possible role in gametophyte development. Other studies about the interactions between proteins, and an evo-devo approach are discussed below.

**Table 3.**
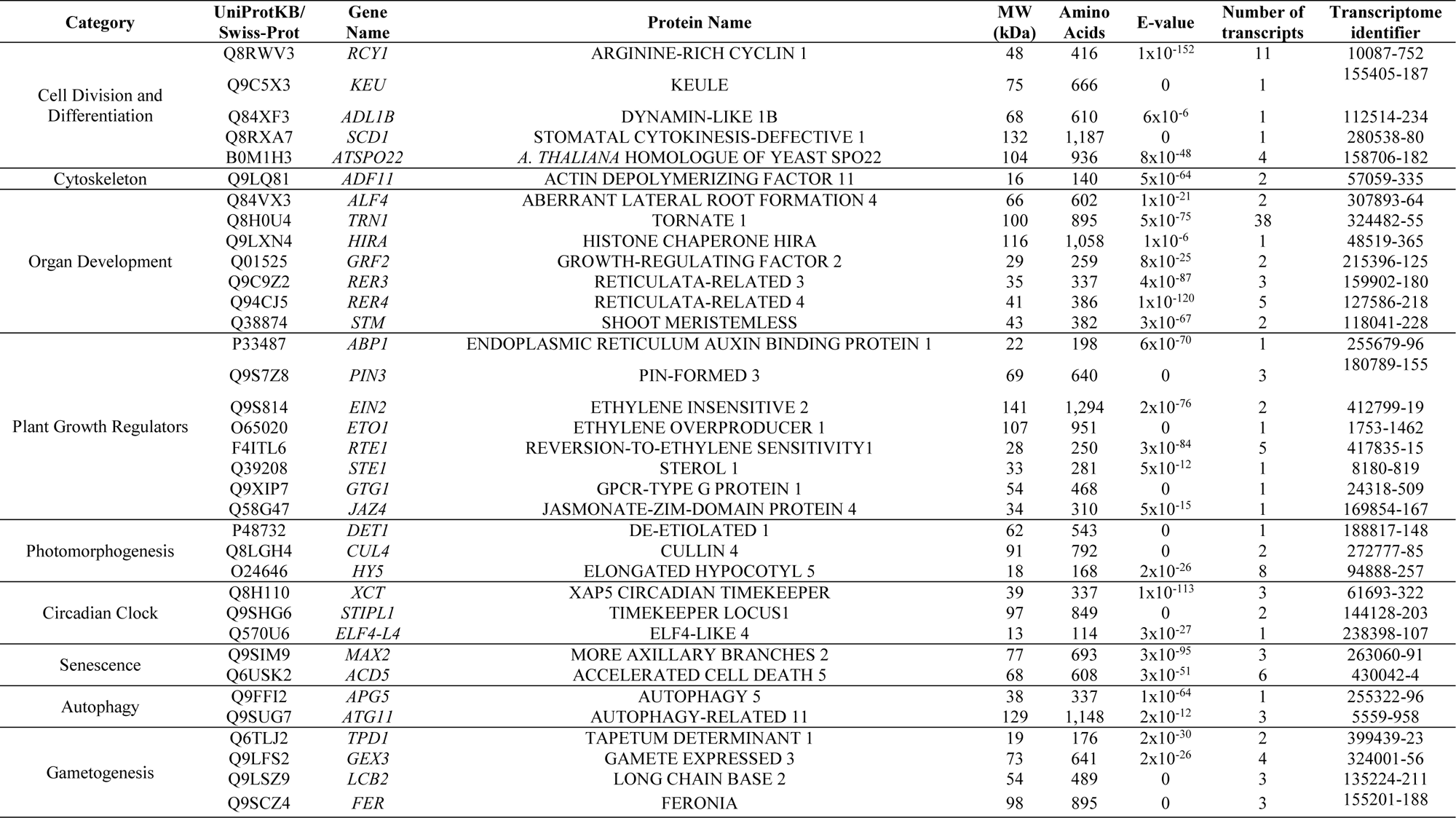
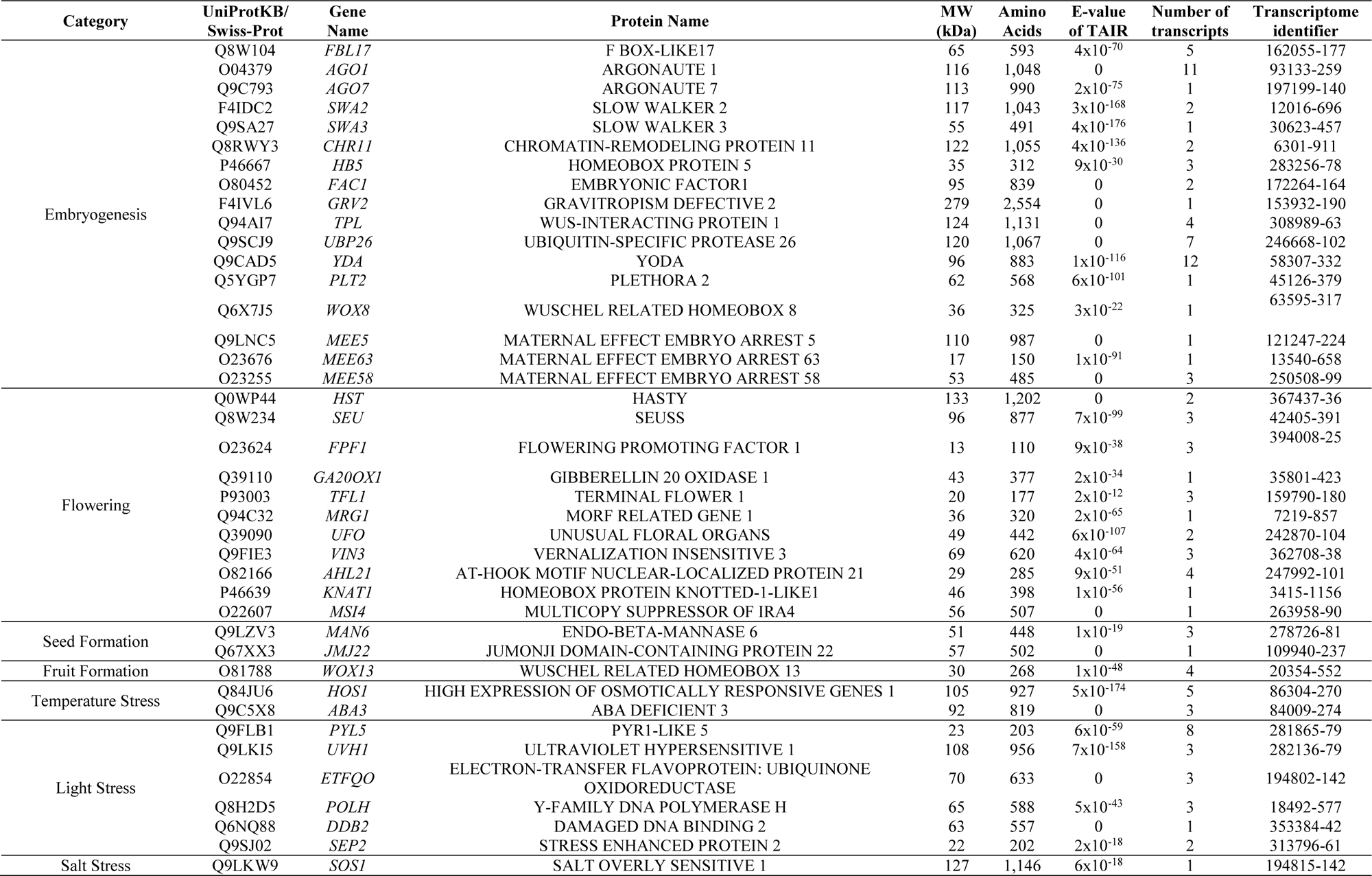
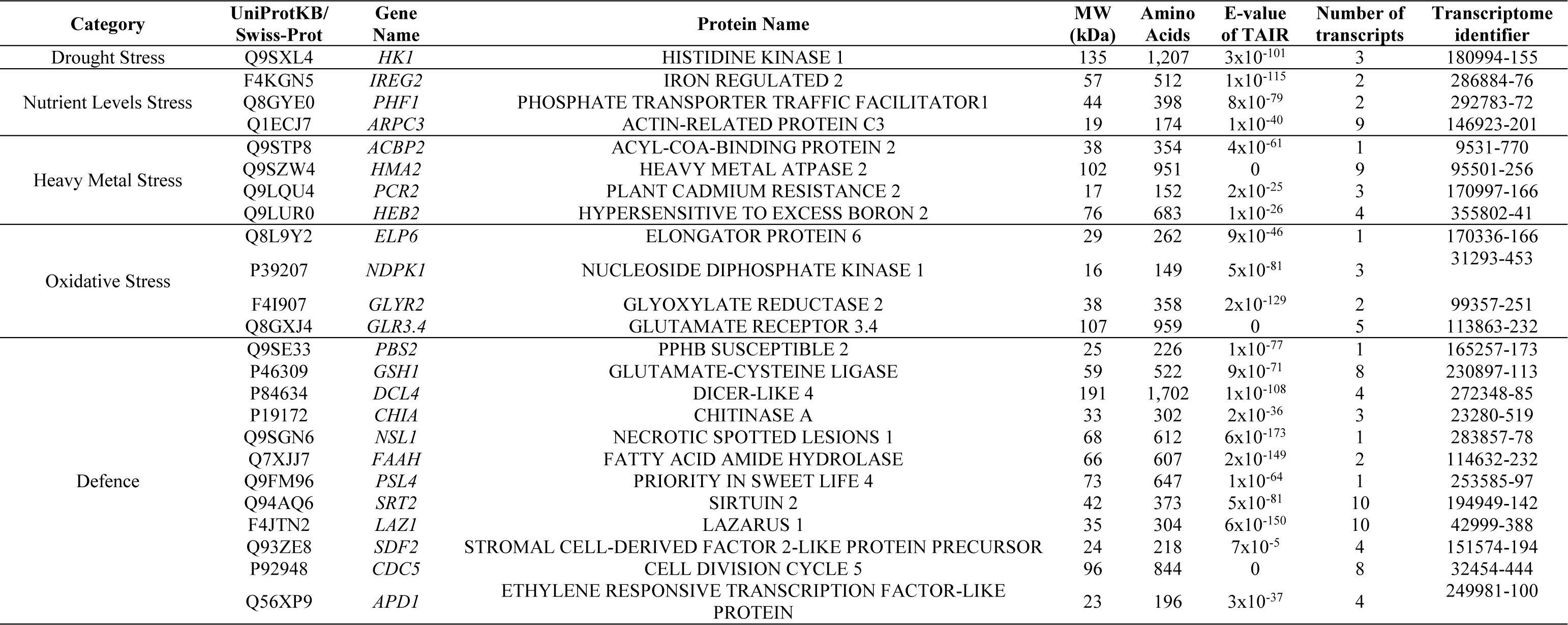
Selected proteins found in gametophytes of the fern *D. affinis* ssp. *affinis* associated to vegetative development, sexual reproduction, and response to stress.

### 4.1. Vegetative Development

In this work, many proteins linked to vegetative development were annotated, as many others coping with stress. Accordingly, the proteins mentioned in this section have been grouped into several clusters, depending on their function in processes associated to cell division and differentiation, cytoskeleton, even rudimentary organ development, plant growth regulators (metabolism, transport, or signalling), photomorphogenesis, circadian clock, senescence, and autophagy.

#### 4.1.1. Cell Division and Differentiation, and Cytoskeleton

Once spore germination occurs, the initial cell begins to divide actively, and differentiate the typical structures described above, each of them dealing with a function: photosynthesis, nutrients uptake or reproduction. As expected, among the proteins annotated in the gametophytes of our fern species, there are some related to cell division and subsequent differentiation, as is the case of several cyclins such as ARGININE-RICH CYCLIN 1 (RCY1), and other cell division-related proteins, including KEULE (KEU), DYNAMIN-LIKE 1B (ADL1B), STOMATAL CYTOKINESIS-DEFECTIVE 1 (SCD1), and ARABIDOPSIS THALIANA HOMOLOGUE OF YEAST SPO22 (ATSPO22). Likewise, actins and tubulins are essential components of the cell cytoskeleton that play an important role in division, and other processes like cytoplasmic streaming, organelle movement, cell shape determination, and extension growth. At this regard, the protein ACTIN DEPOLYMERIZING FACTOR 11 (ADF11) is one example identified in the gametophytes of our species of ferns.

#### 4.1.2. Organ Development, and Plant Growth Regulators (PGR)

Although fern gametophytes have no roots or leaves, but rhizoids and flat lobules, numerous proteins implicated in the development of these organs were annotated from the gametophyte of *D. affinis*, which otherwise, could gain sense in the sporophytic phase. This is the case of the proteins ABERRANT LATERAL ROOT FORMATION 4 (ALF4), required in the initiation of lateral root formation independent of auxin; and TORNADO 1 (TRN1), controlling the initial divisions of the root cap, the radial pattern, and the distinction between lateral root cap and epidermal cells. Ferns can contribute to fill out some missing gaps of knowledge in plant evolution. Schneider (2013) reminds that most fern roots branch via the lateral roots, which form endogenously from the endoderms or pericicle, resembling the lateral root formation of seed plants. In addition, the rhizoids of gametophytes, as those in bryophytes, have been suggested to precede the development of root hairs in angiosperms (Schneider 2013). Concerning leaf development, we identified HOMOLOG OF HISTONE CHAPERONE HIRA (HIRA), involved in gene silencing in this process, the transcriptional activator GROWTH-REGULATING FACTOR 2 (GRF2), which regulates leaves cell expansion in *A. thaliana*, or the proteins RETICULATA-RELATED 3 (RER3) and 4 (RER4). These proteins showed very low E-values in the *in silico* analyses performed, as the leaves of seed plants and ferns probably had initially the same common ancestor although they later evolved independently (Zumajo-Cardona et al., 2019). Likewise, meristems are another important structure in fern gametophytes, which show apical cell-based meristems and multicellular apical and marginal meristems (Wu et al. 2023). Fern sporophytes also show three types of meristems: root apical meristem, leaf apical meristem and shoot apical meristem (Schneider et al., 2013). Related to shoot apical meristem, we foundSHOOT MERISTEMLESS (STM), required for the maintenance of undifferentiated cells in *Arabidopsis* shoot and floral meristems (Endrizi et al., 1996).

Some annotated proteins from our transcriptome, which would hypothetically correspond to each anatomical part of a typical two-dimensional gametophyte, are depicted in **Fig. 7**, speculating about the existence of common genomic pathways in the gametophyte of ferns and other more complex phases of plant development, such as the sporophyte in both ferns and their closely related seed plants. Further analyses will be needed to decipher how genomic fingerprints evolve through the successive taxa and to understand the true functions displayed in them.

**Figure 7.**
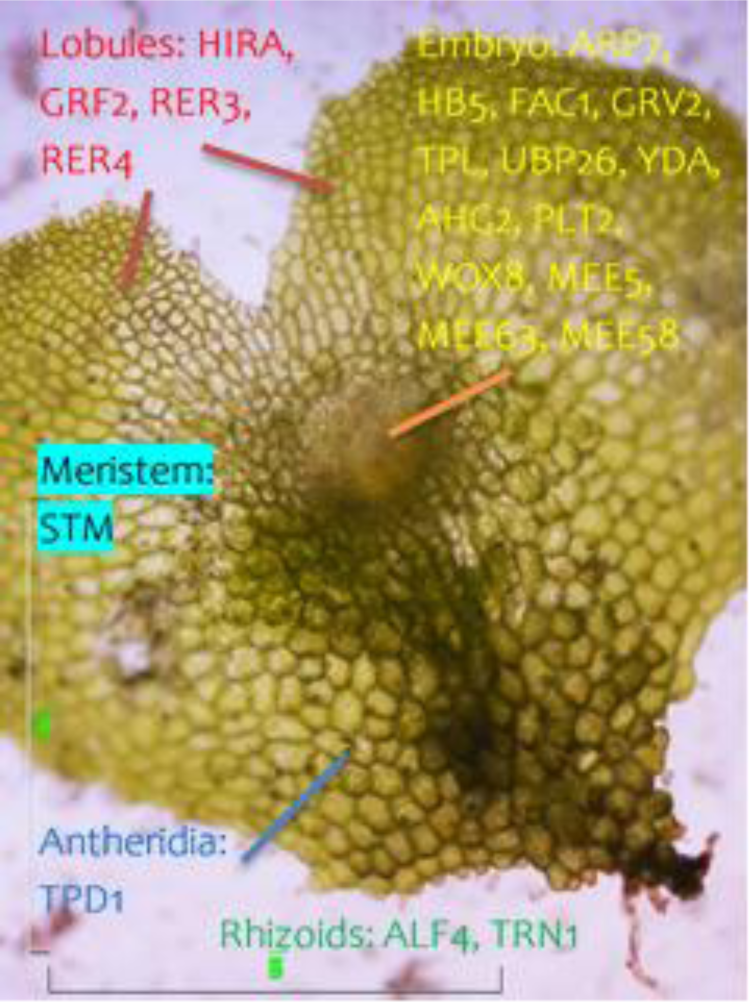
Light microscope image of a two-dimensional gametophyte of *D. affinis* ssp. a*ffinis* in which some proteins found in the transcriptome are linked to the corresponding anatomical part.

Our data also showed a high number of proteins implicated in th metabolism, transport or signalling of phytohormones, headed by auxins and cytokinins. *In vitro* culture of gametophytes of this species reflected their influence on both vegetative and apomictic development (Menéndez et al. 2006). In concrete, by chromatography and mass spectrometry analyses, it was reported increased levels at the dawn of the apogamic event. Regarding the auxin hormone, the receptor ENDOPLASMIC RETICULUM AUXIN BINDING PROTEIN 1 (ABP1) and others like PIN-FORMED 3 (PIN3), mediating its polar transport in *A. thaliana* were reported. Auxins and cytokinins are suggested to be involved in stablishing polarity in the syncytial gametophyte, in many plant species (Terceros et al., 2020). In ferns, in a study carried out in young sporophytes of the fern *C. richardii*, it was detected the expression of *PIN3* gene in the root tip, given the directional movement characteristic of auxin transport (Aragón-Raygoza et al., 2022). Recently, in gametophytes of the fern *C. richardii*, Whithers et al, (2023) demonstrated that correct auxin transport is involved in meristem positioning and regeneration, as they observed that upon destruction of a marginal meristem cell, cell proliferation ceased in this area, and an auxin synthesis gene was no longer expressed, but with the addition of auxins, such as indolacetic or 2, 4-dichlorophenoxyacetic acids, the meristem shifted position to the apex of the gametophytes. Ethylene is other important plant growth regulator that in *D. affinis* gametophytes is signified by several proteins, including ETHYLENE INSENSITIVE 2 (EIN2), ETHYLENE OVERPRODUCER 1 (ETO1), and REVERSION-TO-ETHYLENE SENSITIVITY1 (RTE1), all regulators of the ethylene-signalling pathway. In ferns, it is involved in the defence against biotic and abiotic stresses (Reynolds and John, 2004), and exhibits an apogamy-inducing function in *C. richardii* (Cordle et al., 2010). In the same fern species, other family of plant growth regulators, the brassinosteroids were documented to promote almost all fern growth and development processes (Zheng et al., 2022), being represented in our transcriptome list by many proteins, such as STEROL 1 (STE1). Regarding abscisic acid (ABA), which was assigned a vegetative growth inhibitory role in cultured *D. affinis* gametophytes (Menéndez et al., 2006), we found several associated proteins, such as the receptors GPCR-TYPE G PROTEIN 1 (GTG1) and PYR1-LIKE 5 (PYL5), the latter detected in the gametophyte of the fern *Alsophila spinulosa* (Hong et al. 2022). Finally, regarding jasmonate, which is used by ferns to detect and respond to infections caused by microbial pathogens (de Vries et al., 2018), we mention the presence in our transcriptome of the repressor JASMONATE-ZIM-DOMAIN PROTEIN 4 (JAZ4). All the mentioned phytohormones were detected in the gametophytes of *D. affinis* and its relative *D. oreades*, by means of chromatography and mass spectrometry analyses performed in our laboratory (Fernández et al., 2021).

#### 4.1.3. Photomorphogenesis and Circadian Clock

Apart from photosynthesis, light induces several morphogenic events in plants throughout their life cycle, from germination to senescence. Thus, through the quantity, quality, direction and periodicity of light, the development and growth of the plant is modified. Gametophytes, unlike sporophytes, can grow in low light conditions, probably because their morphology and physiology ensure efficient use of light (Johnson et al., 2000). Likewise, gametophytes are the first stage to cope with the environment, so how they respond to ambient light will determine the growth of the sporophyte, the next stage. Our results revealed the presence in the gametophytes of this fern of several proteins associated with photomorphogenesis such as DE-ETIOLATED 1 (DET1), CULLIN 4 (CUL4), or the transcription factor ELONGATED HYPOCOTYL 5 (HY5). As in many other organisms, cell plants contain an autonomous clock that connects the inside of the organism with environmental signals such as temperature or photoperiod to orchestrate plant development, allowing to synchronise inner processes with daily and seasonal changes and thus maintain a homeostatic balance. Circadian regulated physiological processes are extensive and in the gametophyte of *D. affinis* several proteins were reported such as XAP5 CIRCADIAN TIMEKEEPER (XCT), TIMEKEEPER LOCUS1 (STIPL1), and ELF4-LIKE 4 (ELF4-L4).

#### 4.1.4. Senescence and Autophagy

The gametophyte of ferns begins life and then dies like any other plant organ or individual. In particular, as fertilisation proceeds, the gametophyte may be consumed by the new sporophyte, which demands nutrients from the gametophyte to sustain its initial development. Thus, by degrading and utilising cellular components, autophagy helps the plant to respond to environmental and developmental signals while maintaining cellular homeostasis, as in many processes that take place in the plant, such as growth, degradation of leaf starch, regulation of lipid metabolism or development of anthers (Cheng et al., 2022). Our data report proteins related to senescence such as MORE AXILLARY BRANCHES 2 (MAX2) and ACCELERATED CELL DEATH 5 (ACD5), and a several proteins connected to autophagy like AUTOPHAGY 5 (APG5), and AUTOPHAGY-RELATED 11 (ATG11), being the last essential for plant nutrient recycling, proteolysis of chloroplast proteins in senescing leaves, degradation of damaged peroxisomes, etc. (Yoshimoto et al., 2014), and the second one, a scaffold of a complex dedicated to the delivery of autophagic vesicles to the vacuole (Li et al., 2014). Thus, *D. affinis* gametophyte seems to exhibit an important catabolic activity to remove unneeded constituents and to recycle intracellular nutrients, which can be associated either with its own development, and also with feeding the new sporophyte.

### 4.2. Reproductive Development: Sexual and Apomictic

The intervention of the genomic machinery in all the processes involved in reproductive development remains to be deciphered, especially in those species that have not yet been considered as good research models, such as most ferns. In this work, several genes involved in sexual and asexual reproduction that are expressed in the gametophyte of the fern *D. affinis* are reported.

#### 4.2.1. Gametogenesis and Embryogenesis

Gametogenesis comprises all those processes committed to produce male and female gametes. In ferns, these are antherozoids and egg cells, which occurs within the sexual organ antheridia and archegonia. Although the gametophyte of *D. affinis* is apomictic, male gametogenesis occurs as antheridia producing antherozoids have been reported (Menéndez et al. 2006). Therefore, it is not odd to have found a myriad of proteins related to male gametogenesis and fertility such as TAPETUM DETERMINANT 1 (TPD1), GAMETE EXPRESSED 3 (GEX3), LONG CHAIN BASE2 (LCB2); F BOX-LIKE17 (FBL17) or the receptor FERONIA (FER). Parallels can be drawn between anatomical parts of ferns and seed plants, and in a recent study with the fern *C. richardii*, TPD1 was shown to be essential for proper development of antheridia (Falls, 2022). In addition, in *A. capillus-veneris* it was showed that spore development in ferns is genetically very similar to pollen in seed plants (Fang et al., 2022). Aside, other proteins annotated in the gametophyte of *D. affinis* had been reported to be essential during megagametogenesis or female gametogenesis. In the gametophyte of *D. affinis*, the female gametogenesis is absent, occurring the apogamic event in the same location where archegonia development usually takes place. Therefore, we speculate about the importance of gene silencing to switch off the sexual pathways. Among them, there are two argonautes of five listed: ARGONAUTE 1 (AGO1) and 7 (AGO7), both involved in RNA-mediated post-transcriptional gene silencing (Cuperus et al., 2010), the three proteins SLOW WALKER 1 (SWA1), SLOW WALKER2 (SWA2), and 3 (SWA3), whose mutants result in the slow and delayed progression of mitosis during megagametogenesis in *Arabidopsis* (Li et al., 2009), or the protein CHROMATIN-REMODELING PROTEIN 11 (CHR11) (Huanca-Mamani et al., 2005). The female gametophyte of angiosperms is regarded as an ideal system to cope with gametogenesis because it gathers various critical pathways associated with cell proliferation and differentiation, determination of polarity, communication, etc. (Li et al., 2009). Nevertheless, its small size compared with those of gymnosperms or non-seed plants, like ferns, has represented a hindrance to carry out molecular analyses, being the free-living and larger gametophyte of ferns, an interesting alternative.

Given that *D. affinis* gametophyte is committed to form embryos, it was expected to find genes encoding proteins associated with embryogenesis, i.e. WUSCHEL RELATED HOMEOBOX 8 (WOX8), essential for several processes such as embryonic patterning, basal embryonic development after fertilisation, and embryonic cell division and proliferation (Wu et al., 2007). We also found the transcription factor HOMEOBOX PROTEIN 5 (HB5), the protein EMBRYONIC FACTOR 1 (FAC1), GRAVITROPISM DEFECTIVE 2 (GRV2), or WUS-INTERACTING PROTEIN 1 (TPL), a transcriptional repressor of root promoter genes in the upper half of the embryo during the transitional phase of embryogenesis (Wei et al., 2015). Following with other proteins exhibiting interesting roles for the embryo development it is YODA (YDA), regulating the first cell fate decisions in the zygote and the early embryo after the initial asymetric division (Lukowitz et al. 2004). In addition, a couple of proteins: UBIQUITIN-SPECIFIC PROTEASE 26 (UBP26), a heterochromatin silencing, and whose loss-of-function mutants show embryonic arrest (Sridhar et al., 2007); and PLETHORA 2 (PLT2), involved in the establishment of the stem cell niche in embryogenesis (Smith and Long, 2010), and which belongs to a large family of transcription factors known to regulate plant development. Our result agrees with other studies, such as those carried out by Youngstrom et al., (2019) and Aragón-Raygoza et al., (2022), who have detected the protein PLT2 in gametophytes and sporophytes of the fern *C. richardii*. Finally, quite a few proteins belonging to the DEAD box helicase family were found: MATERNAL EFFECT EMBRYO ARREST 5 (MEE5), 63 (MEE63), and 58 (MEE58), all important in embryogenesis.

#### 4.2.2. Flowering, Seed, and Fruit Formation

Although ferns do not form flowers, genes involved in flowering were annotated in gametophytes of *D. affinis*. These findings are even more remarkable, considering that the gametophyte of this ferns reproduces compulsory asexually. However, the fact that genes related to flowering, seeds, true leaves or roots are expressed in fern gametophytes, as Cordle et al., (2012) reported in *C. richardii*, and that their mutants cannot reproduce correctly by apogamy, suggests that these genes may have other roles, including asexual reproduction. Another possible hypothesis that would explain the presence of these genes in the fern genome and that they are not known to have any function, could be due to be silenced somwhow, supporting the hypothesis that asexual reproduction could have arisen by silencing sexual reproduction. Unfortunately, we are still far from elucidating the reason of these findings. Among this group, there are proteins involved in vegetative to reproductive phase transition as the transporter HASTY (HST), FLOWERING PROMOTING FACTOR 1 (FPF1), GIBBERELLIN 20 OXIDASE 1 (GA20OX1), the protein KNOTTED-LIKE FROM ARABIDOPSIS THALIANA (KNAT1), which in *A. thaliana* seems to be necessary to maintain cells in an undifferentiated meristematic state until the shoot apex undergoes the transition from vegetative to reproductive development (Kim et al., 2003); the protein GLYCINE-RICH PROTEIN 2B (GRP2B), a chaperone that binds nucleic acids to regulate transition to flowering and subsequent flower development; MULTICOPY SUPPRESSOR OF IRA 4 (MSI4), a core histone-binding subunit belonging to the WD-40 repeat family that in *A. thaliana* promotes flowering (Chowdhury et al., 2020), and AT-HOOK MOTIF NUCLEAR-LOCALIZED PROTEIN 21 (AHL21), a transcription factor that binds to the promoter of the FT gene regulating the flowering (Yun et al., 2012). Also, there are proteins which intervene in histone acetylation of flowering target genes, such as SEUSS (SEU), TERMINAL FLOWER 1 (TFL), and UNUSUAL FLORAL ORGANS (UFO). Other proteins detected involved in flowering time were MORF RELATED GENE 1 (MRG1), a chromatin remodelling factor that increases transcriptional levels of the genes FLC and FT (Bu et al., 2014)., It was also annotated VERNALIZATION INSENSITIVE 3 (VIN3), usually required during vernalization and cold treatment in seed plants.

Despite being a non-seed plant, *D. affinis* gametophyte has proteins associated with seed germination, such as ENDO-β-MANNASE 6 (MAN6) and JUMONJI DOMAIN-CONTAINING PROTEIN 22 (JMJ22). Continuing with the parallelism between fern and seed plant organs, these proteins in ferns may play a role in spore formation, as shown in *C. richardii* (Marchant et al., 2022). On the other hand, *D. affinis* is also fruitless, but a protein involved in fruit formation was found: WUSCHEL RELATED HOMEOBOX 13 (WOX13), a transcription factor that in *A. thaliana* promotes replum development and inhibits fruit dehiscence (Ikeuchi et al., 2022).

#### 4.2.3. Asexual reproduction

We firmly think that the gametophyte of apomictic ferns could throw some light to the understanding of the molecular bases underpinning apomixis, and opening the way for extending its agricultural applications, dreamed since long, by researchers from all the world. During the first studies carried out to elucidate the mechanisms that explain apomixis, Nogler (1984) demonstrated that this mode of reproduction is genetically controlled. Subsequently, Savidan (2001) noted that it can be affected by genetic modifiers or environmental conditions. In recent years, epigenetic mechanisms, such as DNA methylation or gene silencing, have been considered to be involved in apomixis, supporting this type of reproduction and inhibiting genes for sexual reproduction (Grimanelli, 2012). This hypothesis is receiving increasing support, as mutations in some epigenetic regulators promote apomixis in sexual plants (Hernández-Lagana et al., 2016). Likewise, RNA helicases, proteins related to the cellular machinery, since they are responsible for unwinding the secondary structures of RNAs, and transcription factors, necessary in gene expression, could have a possible role in apomixis. By whatever mechanism, the asexual reproduction has emerged in plants because of mutations in sexual reproduction genes in order to prevent infertility (Xu et al., 2022). Therefore, considering all this knowledge, we have focused on studying proteins associated with these processes in the transcriptome of *D. affinis* gametophyte.

Starting with methylation, it can be linked to apomixis, as discussed above. This is also reflected in some studies such as that of Podio et al., (2014), in which a reduction in methylation resulted in a decrease in parthenogenesis in *Paspalum* ssp. and, in addition, the apomixis determining region (ASGR) shows a higher level of cytokine methylation. Based on all these results, Schmidt (2020) proposed the hypothesis that apomixis would have overlapped sexual reproduction through epigenetic control mechanisms. Related to this, the list of our transcriptome includes a couple of signified proteins, which, by methylating the genome, prevent the transcription of genes that are not necessary at certain stages of the plant’s reproductive development. These proteins are part of a complex called FERTILIZATION INDEPENDENT SEED (FIS), an epigenetic regulatory Polycomb Repressive Complex 2 (PRC2): FERTILIZATION-INDEPENDENT ENDOSPERM (FIE) and MULTICOPY SUPPRESSOR OF IRA1 (MSI1). Both proteins repress the development of an unfertilized embryo and endosperm in *A. thaliana,* and their mutants can initiate parthenogenesis (Ohad et al., 1999; Guitton and Berger, 2005; Weinhofer et al., 2010).

Gene silencing is another mechanism that can be implicated in apomixis, so a study of silencing proteins may help to understand the mechanisms that explain apogamy in ferns (Barcaccia and Albertini, 2013). In our case, some of the silencing gene proteins found in the transcriptome of *D. affinis* belong to the ARGONAUTE family: AGO1, 4, 7, 9 and 10, having been AGO1 and 7 mentioned above. All of them are involved in gene silencing, but while AGO4 is required during transcription, AGO9 and AGO10 are required after transcription. Further studies pointed in the direction of a possible relationship between gene silencing and apomixis (Olmedo-Monfil et al., 2010; Rabiger et al., 2016), as *A. thaliana* and *Hieracium* mutants for ARGONAUTE proteins exhibited apospory. Likewise, other reports have suggested a role of small interferring RNA (siRNA), in apomictic development, as it is promoted by a decrease in the RNA-dependent DNA methylation pathway in eggs (Grimanelli, 2012). On the other hand, by silencing AGO10 controls shoot apical meristem activity and the establishment of leaf polarity in *A. thaliana*, as does SERRATE (*SE*) (Dong et al., 2008), another protein reported in the gametophyte transcriptome. Both proteins could be candidates to intervene in apogamy, as suggested by Grossmann et al., (2017), since these proteins could participate in the meristem development of the incipient apogamous embryo or in other yet unknown functions in the switch between sexual and asexual reproduction.

Continuing with RNA helicases, it deserves to be mentioned the large number found of this sort of proteins in the transcriptome of *D. affinis*. According to STRING program analyses, they have many interactions between them, with RNA HELICASES 17 (RH17) and 57 (RH57) having the highest number. Some of these proteins are involved in gametogenesis, so it is also speculated that they have a possible role in apomixis, contributing to shaping the plasticity of plant reproductive development (Schmidt, 2020). In line with it, the protein FASCIATE STEM 4 (FAS4) was reported, which was found to be differentially regulated in the sexual and apomictic angiosperm *Boechera* (Schmidt, 2020). Three other RNA helicases mentioned above are SLOW WALKER1 (SWA1), 2 (SWA2), and 3 (SWA3), all needed in female gametogenesis. Specifically, SWA1 is essential in this process in *A. thaliana* for nuclear division and organization in mitosis (Shi et al., 2005), SWA2 is involved in the export of ribosomes (Liu et al., 2010), and SWA3 participates in ribosomal biogenesis, being this, together with RNA processing, fundamental for the progression of the cell cycle during the development of female gametophytes (Liu et al., 2010). It is likely that SWA3 belong to the same ribosome associated pathway that FAS4, controlling different aspects of mitosis in gametogenesis, since it was observed that mutants of *SWA3* gene show the same defects as mutants of the *FAS4* gene (Liu et al., 2010; Schmidt, 2020). However, future studies are needed to better understand the possible role of these proteins in apomixis. The last RNA helicase we want to highlight here is MATERNAL EFFECT EMBRYO ARREST 29 (MEE29), essential in pre-mRNA splicing during embryogenesis in *Hypericum perforatum* (Barcaccia and Albertini, 2013).

Other proteins associated with apomixis were transcription factors, such as BABY BOOM (*BBM*), that induces parthenogenesis and somatic embryogenesis in *Brassica napus* and *A. thaliana* (Boutilier et al., 2002; Schmidt, 2020), and apogamy in *C. richardii* (Bui et al., 2017); and the AGAMOUS-LIKE proteins, belonging to the MADS-box family, widely represented in our transcriptome. We remark here two members: AGAMOUS-LIKE 62 (AGL62) and 6 (AGL6), the primer promoting cellularization in early endosperm development in *A. thaliana* (Kang et al., 2008), while AGL6 is involved in megasporogenesis in the apomictic and sexual angiosperm *Brachiaria brizantha* (Guimaraes et al., 2013). A deeper study of how these transcription factors influence the genome of the *D. affinis* gametophyte would allow a better understanding of the mechanisms behind apomixis. Finally, other proteins with a possible role in asexual reproduction are SWITCH1 *(SWI1),* required in *A. thaliana* during meiosis of male and female gametes, and chromatin remodelling (Siddiqi et al., 2000); and BRI1-ASSOCIATED RECEPTOR KINASE *(BAK1),* intervening in somatic embryogenesis and embryo sac development in *Poa pratensis* (Albertini et al., 2005).

### 4.3. Response to Abiotic and Biotic Stress

The gametophyte generation is considered a critical exploratory stage for ferns, being, in most all cases, more stress tolerant than sporophytes, growing in areas where sporophytes cannot. Certainly, gametophytes, despite their delicate appearance, can be robust in response to abiotic stress (Krieg and Chambers 2022). Our data have revealed the expression of a large group of genes that allow the gametophyte to cope with environmental threats imposed by variations in temperature, light, salinity, water, nutrient levels, or biotic attack, some of them being commented next.

#### 4.3.1. Temperature and light stress

Temperature can stress plant development by heat, cold or freezing. Indeed, possible increasing temperatures, and varying precipitation patterns due to climate change, may exceed the resilience of some species, including ferns. As occurs with other stressors, studies about the effect of temperature on development and growth of fern gametophytes is scarce. Indeed, we have little information in general on fern heat tolerance. In this study, it was annotated the proteins HIGH EXPRESSION OF OSMOTICALLY RESPONSIVE GENES 1 (*HOS1*), and ABA DEFICIENT 3 (ABA3), involved in stress by low temperature. More space would be needed to detail the numerous heat shock proteins annotated from the transcriptome of *D. affinis*, revealing the rich machinery present in the gametophyte to deal with any threaten environment. Likewise, in the response to light stress, while sporophytes can increase light capture by producing large fronds and growing higher, fern gametophytes must adapt their photosynthetic machinery to achieve a positive carbon balance in their prevailing light conditions. Aside, proteins induced in the absence of low light conditions can be regulated like the protein ELECTRON-TRANSFER FLAVOPROTEIN: UBIQUINONE OXIDOREDUCTASE (*ETFQO*). Other proteins aim to avoid the damage by ultraviolet light, such as ULTRAVIOLET HYPERSENSITIVE 1 (UVH1), Y-FAMILY DNA POLYMERASE H (POLH) and DAMAGED DNA BINDING 2 (DDB2). In addition, the protein STRESS ENHANCED PROTEIN 2 (SEP2) is involved in non-photochemical quenching, minimising the generation of oxidative molecules.

#### 4.3.2. Salt and water stress

Salinity is one of the most studied stress factors, affecting also prothallus development. Higher salinity of the substrate produces a significant decrease in gametophyte sizes, and affect the development of gametangia as well (Pangua et al., 2009). *D. affinis* gametophyte contains proteins to overcome salt stress including SALT OVERLY SENSITIVE 1 (*SOS1*), one of the best known. This protein is member of the salt overly sensitive signalling, to maintain ion homeostais. Regarding water stress, it must be mentioned that fern gametophytes requires wet environments for the success of sexual reproduction, allowing male gametes swimming until reaching the egg cell. Nevertheless, *D. affinis* gametophyte seems to be well equipped to fight against water scarcity and escape damage by producing proteins like the osmo-sensor of water stress and signalling component HISTIDINE KINASE 1 (HK1). This is only one example of the genomic arsenal the gametophyte of *D. affinis* has to overtake water scarcity. It is striking that the gametophyte, despite such a limited water storage capacity and rudimentary cuticle, has developed remarkable degrees of desiccation tolerance, preventing its survival to be compromised.

In order to first obtain only one-dimensional and not two-dimensional gametophytes, the spores of *D. affinis* were overcrowded, which caused nutrient stress, which might be consistent with the obtaining of proteins like IRON REGULATED 2 (IREG2), and PHOSPHATE TRANSPORTER TRAFFIC FACILITATOR1 (PHF1), involved in iron and phosphate scarcity, respectivel. ACTIN-RELATED PROTEIN C3 (ARPC3) was also reported, which was detected in gametophytes of the fern *Diplazium maximum* as a protein that upon exposure to extreme changes in the environment increases its presence in response to stress (Sareen et al., 2019).

#### 4.3.3 Heavy metal and oxidative stress

Another type of stress are heavy metals, which can be harmful for plant development. Some ferns have been proved to be good hyperaccumulators, being screened for gene candidates for the design of stress tolerant crops (Chen 2022). *D. affinis* gametophyte shows to be provided with proteins to respond to plumb like ACYL-COA-BINDING PROTEIN 2 (ACBP2); cadmium such as HEAVY METAL ATPASE 2 (HMA2) and PLANT CADMIUM RESISTANCE 2 (PCR2); and boron with the protein HYPERSENSITIVE TO EXCESS BORON 2 (HEB2). In relation to oxidative stress, we only mention here some examples of a long list: ELONGATOR PROTEIN 6 (ELP6), NUCLEOSIDE DIPHOSPHATE KINASE 1 (NDPK1), and GLYOXYLATE REDUCTASE 2 (*GLYR2*). Apart from all the above, there are other stress-related proteins, including GLUTAMATE RECEPTOR 3.4 (GLR3.4), required in rapid calcium-based transmission of environmental stress.

#### 4.3.4. Defence

It has been considered that bryophytes and ferns are rarely consumed by phytophagous insects in nature. It has been speculated that this is due to lack of flowers and fruits in these groups of plants, the accumulation of secondary metabolites, and the presence of genes of resistance (Markam et al. 2006). However, little is known about defence mechanisms against insects in ferns and bryophytes unlike what is documented in angiosperms. These authors reported that extract proteins might cause decrease in damage in leaf-disk assays to larval insect growth in several fern species. In the trancriptome of *D. affinis*, a long list of proteins involved in defence were annotated. Among them, we identified the proteins PPHB SUSCEPTIBLE 2 (*PBS2*), GLUTAMATE-CYSTEINE LIGASE (GSH1), the antiviral helicase DICER-LIKE 4 (DCL4), CHITINASE A (CHIA), NECROTIC SPOTTED LESIONS 1 (NSL1), FATTY ACID AMIDE HYDROLASE (FAAH), PRIORITY IN SWEET LIFE 4 (PSL4), STROMAL CELL-DERIVED FACTOR 2-LIKE PROTEIN PRECURSOR (SDF2), SIRTUIN 2 (SRT2), LAZARUS 1 (LAZ1) and CELL DIVISION CYCLE 5 (CDC5). They intervene in plant innate immunity, being induced in response against the pathogen-associated molecular patterns, or in programmed cell death through the hypersensitive response. Likewise, several ethylene responsive annotations were found in the fern gametophyte transcriptome, such as ETHYLENE RESPONSIVE TRANSCRIPTION FACTOR-LIKE PROTEIN (APD1), being necessary for the systemic acquired resistance. Ethylene responsive transcription factors represent a key regulatory centre in the response of plants to abiotic stress, as they are involved in signalling hormones, such as ethylene, jasmonate and abscisic acid, and in oxidation-reduction reactions (Muller and Munné-Bosch, 2015).

### 4.4. Protein-Protein Interactions

The proteins MULTICOPY SUPPRESSOR OF IRA 4 (MSI4) and 1 (MSI1), SLOW WALKER1 (SWA1), and FASCIATED STEM 4 (FAS4) had the most interactions, probably because they are involved in many processes of vegetative and reproductive development (Guitton and Berger, 2005; Shi et al., 2005; Chowdhury et al., 2020; Schmidt, 2020). In the software used (String), each interaction score expresses an approximate confidence on a scale of 0 to 1 that the association is true, given all available evidence (Szklarczyk et al., 2019). Accordingly, the strongest neighbourhood interaction in our dataset was between the proteins FATTY ACID AMIDE HYDROLASE (FAAH), required in defence, and PYRIMIDINE 2 (PYD2), involved in nitrogen recycling from nucleobases, which is explained by the proximity in the genome of both genes (Zrenner et al., 2009; Khan et al., 2017). In the case of gene fusion channel, the interaction was strongest between CELL DIVISION CYCLE 5 (CDC5), essential for plant innate immunity, and MATERNAL EFFECT EMBRYO ARREST 29 (MEE29), needed in embryogenesis, because at least in the fungus *Pneumocystis jirovecii* orthologous protein-coding genes are fused into an only gene (Barcaccia and Albertini, 2013). The strongest co-occurrence interaction was between HASTY (HST), involved in the vegetative to reproductive phase transition, and SERRATE (SE), which regulates leaf polarity, because their genome has evolved similarly over time (Dong et al., 2008). As for homology evidence, ARGONAUTE 1 (AGO1) and 7 (AGO7), involved in RNA-mediated post-transcriptional gene silencing, showed the highest values, as they present the same common ancestor and therefore, similar sequences (Cuperus et al., 2010). AGO1 and AGO7 are paralog proteins, i.e. both are found in the genome of the same species (Jacques et al., 2023). In terms of co-expression, the strongest interaction found was between SLOW WALKER2 (SWA2) and FASCIATED STEM 4 (FAS4), as their expression patterns are similar. Both are required for female gametogenesis and believed to be candidate genes for apomixis (Li et al., 2009; Schmidt, 2020). The next evidence studied were experiments, i.e. proteins that have shown to have chemical, physical, or genetic interactions in laboratory experiments, and databases, i.e. proteins found in the same databases. In our dataset, the strongest interactions in these channels were between FERTILIZATION-INDEPENDENT ENDOSPERM (FIE) and MULTICOPY SUPPRESSOR OF IRA 1 (MSI1). FIE and MSI represses the process of parthenogenesis, so both are considered to be apomixis-related proteins (Jullien et al., 2008; Fei et al., 2019). Finally, the proteins required in photomorphogenesis CULLIN 4 (CUL4) and DE-ETIOLATED 1 (DET1) showed the strongest text mining interaction, as they are mentioned in the same PubMed abstract or articles from an internal selection of the STRING software (Nassrallah et al., 2018).

### 4.5. Phylogenetic analyses

Evolutionarily, ferns diverged from gimnosperms and angioperms 400 million years ago. Thus, they constitute a phylogenetic clade that can provide a great deal of information; however, this plant group is markedly absent in most comparative studies (Plackett et al., 2015). In order to study which species have the most similar sequences to our fern proteins and to understand their genealogical relationships and evolutionary history over time, phylogenetic analyses were tackled. For this purpose, the pattern of branching in phylogenetic trees, corresponding to amino acid sequences of certain proteins of vegetative and reproductive development, and response to biotic and abiotic stress were obtained. These trees are constructed after making comparisons between multiple sequences, relating the degree of similarity with the evolutionary relationship between organisms. In the protein ARGONAUTE 1 (AGO1), the sequences of *A. nelumboides*, *A. capillus-veneris*, and *C. richardii* are the closest to our fern sequence, while in BABY BOOM (BBM), the sequence of *A. nelumboides* is the most distant. All these plants are ferns, as is our study species. In WUS-INTERACTING PROTEIN 1 (TPL), one of the closest species is *A. trichopoda*. Likewise, in DICER-LIKE 4 (DCL4), our sequence is further away from other studied species, such as *S. intermedia*, a monocot. As for the other proteins studied: CULLIN 4 (CUL4), TIMEKEEPER LOCUS1 (STIPL1), UBIQUITIN-SPECIFIC PROTEASE 26 (UBP26), and UNUSUAL FLORAL ORGANS (UFO), there is more evolutionary distance between the sequences of our fern and the rest of the species analysed, such as *A. trichopoda*. After comparing the fern sequences of the eight trees, the first protein studied, AGO1, showed 52 % differences from the original sequence, which means that it is evolutionarily further away from the common ancestor in each group of sequences and has evolved more. These comparisons between sequences are made after measuring the distance from the common ancestor in the tree, as this is a measure of the degree of divergence that two sequences have undergone, i.e. it provides an estimate of the time that has passed since two sequences ceased to be the same and each evolved in a different way. Thus, the evolutionary relationships between different amino acid sequences are inferred, studied, and evaluated.

As for the protein domain analyses, in AGO1 and BBM the same number and type of domains are conserved in *D. affinis* and *A. thaliana* species: the DUF1785, Paz and Piwi domains in the first, and the AP2 domain in the second. The Paz and Piwi domains are found in all ARGONAUTE proteins, with the Piwi domain having a role in epigenetic regulation of the transcriptome (De Storme and Geelen, 2013). The protein BBM belongs to the APETALA 2/ETHYLENE-RESPONSIVE ELEMENT BINDING FACTOR (AP2/ERF) family, which has three classes of genes depending on the number of AP2 domains present. BBM always has two AP2 domains (Bui et al., 2017). The protein TPL in *A. thaliana* has one LisH domain, one CTLH domain and eleven WD40 domains, five WD40 domains more than in *D. affinis*. These WD40 domains function as a platform for interaction with other proteins or DNA (Xu and Min, 2011). Regarding the fourth protein studied, DCL4, in *A. thaliana*, in addition to two RIBOc domains and two DSRM domains, has one DEXDc, one HELICc and one Paz domains, which are absent in *D. affinis*. The DEXDc and HELICc domains are responsible for recognising the RNA of the pathogenic virus in the plant’s defence against infection (Yao et al., 2022). The other proteins studied, CUL4, STIPL1, UBP26 and UFO, have the same number and type of domains in the two species studied: *D. affinis* and *A. thaliana*. CUL4 has a CULLIN domain and a Cullin Nedd8 domain, being the first a scaffold for ubiquitination and the second the site where the NEED8 protein binds for ubiquitination to occur (Rabut and Peter, 2008). The STIPL1 protein has a G patch domain, a glycine-rich domain involved in eukaryotic RNA processing (Bohnsack et al., 2021). As for the UBP26 and UFO proteins, the first shows a DUSP domain and the second a FBOX domain, both required for protein ubiquitination and subsequent degradation by the 26S proteasome (Elliott et al., 2011; Zhang et al., 2019).

In conclusion, an exhaustive study of the transcriptome obtained from the apomictic gametophyte of the fern *D. affinis* using RNA-seq has proven to be a very useful methodology to annotate thousands of genes, categorise them and discuss their role in plant development. Although the gametophyte is a very simple structure, and ferns lack complex reproductive organs like flowers, the large number of genes found to be associated with reproduction, including gametogenesis, embryogenesis, flowering and seed development, is striking, with some of them being possible candidates for apomixis. Among them, we reported some RNA helicases, such as FAS4, SWA1, and MEE29; other genes intervening in gene silencing, represented by AGO4, 9 and 10, and SE; other linked to methylation, such as FIE and MSI1; or the protein BBM and two AGAMOUS-LIKE proteins: AGL6 and 62. The protein-protein interactions are provided by String platform and came from experiments, databases and text mining channels, and were led by MULTICOPY SUPPRESSOR OF IRA 4 (MSI4) with 16, MULTICOPY SUPPRESSOR OF IRA 1 (MSI1) and SLOW WALKER1 (SWA1) with 12, and FASCIATED STEM 4 (FAS4) with 11. In addition, proteins associated with vegetative development, as well as with the response to biotic and abiotic stresses are also pointed to. The highest number of protein-protein interactions provided by the String platform came from experiments, databases and text mining channels, and were led by MSI4 with 16 interactions, MSI1 and SWA1 with 12, and FAS4 with 11. Regarding phylogenetic analysis, the protein AGO1 was the most evolutionarily similar to other ferns and the most distant to the common ancestor. All this knowledge provides novel information that open new research lines to the understanding of the molecular bases underpinning gametophyte development, including apomixis.

## Acknowledgments

This research was supported by the University of Zurich, the University Research Priority Program Evolution in Action, and the Program for Mobility of Excellence at Oviedo University. We would like to express our sincere gratitude to Dr. Stephan Wyder, for his great work with transcriptome performance. Also, thanking to Hanspeter Schöb (University of Zurich) for administrative and logistic support, and the Functional Genomics Center Zurich for sequencing and generous assistance with all the analyses.

## References

Albertini, E., Marconi, G., Reale, L., Barcaccia, G., Porceddu, A., Ferranti, F., and Falcinelli, M. (2005). SERK and APOSTART. Candidate genes for apomixis in *Poa pratensis*. Plant Physiol. 138, 2185–2199. DOI: 10.1104/pp.105.062059

Aragón-Raygoza, A., Herrera-Estrella, L., and Cruz-Ramírez, A. (2022) Transcriptional analysis of *Ceratopteris richardii* young sporophyte reveals conservation of stem cell factors in the root apical meristem. Front. Plant Sci. 13, 924660. DOI: 10.3389/fpls.2022.924660

Bakkali, M., Martín-Blázquez, R., Ruiz-Estévez, M., and Garrido-Ramos, M. A. (2021). *De novo* sporophyte transcriptome assembly and functional annotation in the endangered fern species *Vandenboschia speciosa* (Willd.) G. Kunkel. Genes 12(7), 1017. DOI: 10.3390/genes12071017

Barcaccia, G., and Albertini, E. (2013). Apomixis in plant reproduction: a novel perspective on an old dilemma. Plant Reprod. 26, 159–179. DOI: 10.1007/s00497-013-0222-y

Barker, M. S., and Wolf, P. G. (2010). Unfurling fern biology in the genomics age. Bioscience 60, 177–185. DOI: 10.155/bio.2010.60.3.4

Bohnsack, K. E., Ficner, R., Bohnsack, M. T., and Jonas, S. (2021). Regulation of DEAH-box RNA helicases by G-patch proteins. Biological Chemistry 402(5), 561–579. DOI: 10.1515/hsz-2020-0338

Boutilier, K., Offringa, R., Sharma, V. K., Kieft, H., Ouellet, T., Zhang, L., Hattori, J., Liu, C.-M., van Lammeren, A. A. M., Miki, B. L. A., Custers, J. B. M., and van Lookeren Campagne, M. M. (2002). Ectopic expression of BABY BOOM triggers a conversion from vegetative to embryonic growth. Plant Cell 14(8), 1737–1749. DOI: 10.1105/tpc.001941

Bu, Z., Yu, Y., Li, Z., Liu, Y., Jiang, W., Huang, Y., and Dong, A. W. (2014). Regulation of *Arabidopsis* flowering by the histone mark readers MRG1/2 via interaction with CONSTANS to modulate *FT* expression. PLOS Genetics. DOI: 10.1371/journal.pgen.1004617

Bui, L. T., Pandzic, D., Youngstrom, C. E., Wallace, S., Irish, E. E., Szovenyi, P., and Cheng, C.-L. (2017). A fern *AINTEGUMENTA* gene mirrors *BABY BOOM* in promoting apogamy in *Ceratopteris richardii*. The Plant J. 90(1), 122–132. DOI: 10.1111/tpj.13479

Cai, Z., Xie, Z., Huang, L., Wang, Z., Pan, M., Yu, X., Xu, S., and Luo, J. (2022). Full-length transcriptome analysis of *Adiantum flabellulatum* gametophyte. Peer J. 10, e13079. DOI: 10.7717/peerj.13079

Chen C-H. (2022). Unveiling novel genes in fern genomes for the design of stress tolerant crops. Crop Desig 1. DOI.org/10.1016/j.cropd.2022.100013

Cheng, S., Wang, Q., Manghwar, H., and Liu, F. (2022). Autophagy-mediated regulation of different meristems in plants. Int. J. Mol. Sci. 23(11), 6236. DOI: 10.3390/ijms23116236

Chowdhury, Z., Mohanty, D., Giri, M. K., Venables, B. J., Chaturvedi, R., Chao, A., Petros, R. A., and Shah, J. (2020). Dehydroabietinal promotes flowering time and plant defence in *Arabidopsis* via the autonomous pathway genes *FLOWERING LOCUS D*, *FVE*, and *RELATIVE OF EARLY FLOWERING 6*. J. Exp. Bot. 71(16), 4903–4913. DOI: 10.1093/jxb/eraa232

Cordle, A. R., Irish, E. E., and Cheng, C.-L. (2012). Gene expression associated with apogamy commitment in *Ceratopteris richardii*. Sex Plant Reprod. 25, 293–304. DOI: 10.1007/s00497-012-0198-z

Creux, N., and Harmer, S. (2019). Circadian rhythms in plants. Cold Spring Harb Perspect Biol 11, a034611. DOI: 10.1101/cshperspect.a034611

Cuperus, J. T., Carbonell, A., Fahlgren, N., Garcia-Ruiz, H., Burke, R. T., Takeda, A., Sullivan, C. M., Gilbert, S. D., Montgomery, T. A., and Carrington, J. C. (2010). Unique functionality of 22-nt miRNAs in triggering RDR6-dependent siRNA biogenesis from target transcripts in *Arabidopsis*. Nat. Struct. Mol. Biol. 17, 997–1003. DOI: 10.1038/nsmb.1866

Davidson, N. M., and Oshlack, A. (2014). Corset: enabling differential gene expression analysis for *de novo* assembled transcriptomes. Genome Biol. 15, 410. DOI: 10.1186/s13059-014-0410-6

De Storme, N., and Geelen, D. (2013). Sexual polyploidization in plants-cytological mechanisms and molecular regulation. New Phytologist 198(3), 670–684. DOI: 10.1111/nph.12184

Dhir, B. (2018). Role of ferns in environmental clean-up. In: Fernández, H. (Ed.) Current Advances in Fern Research, Springer International Publishing, Switzerland, 517–531. ISBN 978-3-319-75102-3

Domzalska, L., Kedracka-Krok, S., Jankowska, U., Grzyb, M., Sobczak, M., Rybczynski, J. J., and Mikuła, A. (2017). Proteomic analysis of stipe explants reveals differentially expressed proteins involved in early direct somatic embryogenesis of the tree fern *Cyathea delgadii* Sternb. Plant Sci. 258, 61–76. DOI: 10.1016/j.plantsci.2017.01.017

Dong, Z., Han, M.-H., and Fedoroff, N. (2008). The RNA-binding proteins HYL1 and SE promote accurate *in vitro* processing of pri-miRNA by DCL1. PNAS 105(29), 9970–9975. DOI: 10.1073/pnas.0803356105

Dyer, R. J., Savolainen, V., and Schneider, H. (2012). Apomixis and reticulation in the *Asplenium monanthes* fern complex. Ann. Bot. 110(8), 1515–1529. DOI: 10.1093/aob/mcs202

Eeckhout, S., Leroux, O., Willats, W. G. T., Popper, Z. A., and Viane, R. L. L. (2014). Comparative glycan profiling of *Ceratopteris richardii* C-Fern gametophytes and sporophytes links cell-wall composition to functional specialization. Ann. Bot. 114(6), 1295–1307. DOI: 10.1093/aob/mcu039

Elliott, P. R., Liu, H., Pastok, M. W., Grossmann, G. J., Rigden, D. J., Clague, M. J., Urbe, S., and Barsukov, I. L. (2011). Structural variability of the ubiquitin specific protease DUSP-UBL double domains. FEBS Letters 585(21), 3385–3390. DOI: 10.1016/j.febslet.2011.09.040

Endrizzi, K., Moussian, B., Haecker, A., Levin, J.Z., and Laux, T. (1996) The SHOOT MERISTEMLESS gene is required for maintenance of undifferentiated cells in *Arabidopsis* shoot and floral meristems and acts at a different regulatory level than the meristem genes WUSCHEL and ZWILLE. Plant J. 10(6):967–79. DOI: 10.1046/j.1365-313x.1996.10060967.x.

Falls, K. P. (2022). Roles of EMS1 and TPD1 in gametogenesis and sporogenesis in the fern *Ceratopteris richardii*. University of Iowa, United States of America. DOI: 10.17077/etd.005500

Fang, Y., Qin, X., Liao, Q., Du, R., Luo, X., Zhou, Q., Li, Z., Chen, H., Jin, W., Yuan, Y., Sun, P., Zhang, R., Zhang, J., Wang, L., Cheng, S., Yang, X., Yan, Y., Zhang, X., Zhang, Z., Bai, S., Van de Peer, Y., Lucas, W. J., Huang, S., and Yan, J. (2022). The genome of homosporous maidenhair fern sheds light on the euphyllophyte evolution and defences. Nature Plants 8, 1024–1037. DOI: 10.1038/s41477-022-01222-x

Fei, X., Shi, J., Liu, Y., Niu, J., and Wei, A. (2019). The steps from sexual reproduction to apomixis. Planta 249, 1715–1730. DOI: 10.1007/s00425-019-03113-6

Fernández, H., Grossmann, J., Gagliardini, V., Feito, I., Rivera, A., Rodríguez, L., Quintanilla, L. G., Quesada, V., Cañal, M. J., and Grossniklaus, U. (2021). Sexual and apogamous species of woodferns show different protein and phytohormone profiles. Front. Plant Sci. 12. DOI: 10.3389/fpls.2021.718932

Fernández, H., and Revilla, M. A. (2003). *In vitro* culture of ornamental ferns. PCTOC 73, 1–13. DOI: 10.1023/A:1022650701341

Gill, A. T., Farrant, J. M., and Rafudeen, M. S. (2009). Characterisation of the heat stable proteome of the desiccation tolerant form of the resurrection fern *Mohria caffrorum*. s. Afr. J. Bot. 75, 433. DOI: 10.1016/j.sajb.2009.02.143

Grimanelli, D. (2012). Epigenetic regulation of reproductive development and the emergence of apomixis in angiosperms. Curr. Opin. Plant Biol. 15, 57–62. DOI: 10.1016/j.pbi.2011.10.002

Grossmann, J., Fernández, H., Chaubey, P. M., Valdés, A. E., Gagliardini, V., Cañal, M. J., Russo, G., and Grossniklaus, U. (2017). Proteogenomic analysis greatly expands the identification of proteins related to reproduction in the apogamous fern *Dryopteris affinis* ssp. *affinis*. Front. Plant Sci. 8, 336. DOI: 10.3389/fpls.2017.00336

Grossniklaus, U., Nogler, G. A., and van Dijk, P. J. (2001). How to avoid sex: the genetic control of developmental aspects. The Plant Cell. 13, 1491–1497. DOI: 10.1105/tpc.13.7.1491

Grusz, A. L., M. D. Windham, K. T. Picard, K. M. Pryer, E. Schuettpelz, and C. H. Haufler. (2021). A drought-driven model for the evolution of obligate apomixis in ferns: evidence from pellaeids (Pteridaceae). Amer. J. Bot. 108(2): 263– 28. DOI:10.1002/ajb2.1611

Guimaraes, L. A., Dusi, D. M. de A., Masiero, S., Resentini, F., Gomes, A. C. M. M., Silveira, E. D., Florentino, L. H., Rodrigues, J. C. M., Colombo, L., and Carneiro, V. T. de C. (2013). *BbrizAGL6* is differentially expressed during embryo sac formation of apomictic and sexual *Brachiaria brizantha* plants. Plant Mol. Biol. Rep. DOI: 10.1007/s11105-013-0618-8

Guitton, A.-E-., and Berger, F. (2005). Loss of function of MULTICOPY SUPPRESSOR OF IRA1 produces nonviable parthenogenetic embryos in *Arabidopsis*. Curr. Biol. 15(8), 750–754. DOI: 10.1016/j.cub.2005.02.066

Hernández-Lagana, E., Rodríguez-Leal., D., Lúa, J., and Vielle-Calzada, J.-P. (2016). A multigenic network of ARGONAUTE4 clade members controls early megaspore formation in *Arabidopsis*. Genetics 204(3), 1045–1056. DOI: 10.1534/genetics.116.188151

Hong, Y., Wang, Z., Li, M., Su, Y., and Wang, T. (2022). First multi-organ full-length transcriptome of tree fern *Alsophila spinulosa* highlights the stress-resistant and light-adapted genes. Front. Plant Sci. 12, 784546. DOI: 10.3389/fgene.2021.784546

Huanca-Mamani, W., Garcia-Aguilar, M., Leon-Martinez, G., Grossniklaus, U., and Vielle-Calzada, J. P. (2005). CHR11, a chromatin-remodelling factor essential for nuclear proliferation during female gametogenesis in *Arabidopsis thaliana*. PNAS 102(47), 17231–17236. DOI: 10.1073/pnas.0508186102

Ikeuchi, M., Iwase, A., Ito, T., Tanaka, H., Favero, D. S., Kawamura, A., Sakamoto, S., Wakazaki, M., Tameshige, T., Fujii, H., Hashimoto, N., Suzuki, T., Hotta, K., Toyooka, K., Mitsuda, N., and Sugimoto, K. (2022). Wound-inducible WUSCHEL-RELATED HOMEOBOX 13 is required for callus growth and organ reconnection. Plant Physiology 188(1), 425–441. DOI: 10.1093/plphys/kiab510

Jacques, F., Bolivar, P., Pietras, K., and Hammarlund, E. U. (2023). Roadmap to the study of gene and protein phylogeny and evolution-A practical guide. PLOS ONE. DOI: 10.1371/journal.pone.0279597

Jin, J., Zhang, H., Kong, L., Gao, G., and Luo, J. (2014). PlantTFDB 3.0: a portal for the functional and evolutionary study of plant transcription factors. Nucleic Acids Res. 42, D1182–D1187. DOI: 10.1093/nar/gkt1016

Johnson, G. N., Rumsey, F. J., Headley, A. D., and Sheffield, E. (2000). Adaptations to extreme low light in the fern *Trichomanes speciosum*. New Phytologist 148, 423–431. DOI: 10.1046/j.1469-8137.2000.00772.x

Jullien, P. E., Mosquna, A., Ingouff, M., Sakata, T., Ohad, N., and Berger, F. (2008). Retinoblastoma and its binding partner MSI1 control imprinting in *Arabidopsis*. PLOS Biology. DOI: 10.1371/journal.pbio.0060194

Kang, I.-H., Steffen, J. G., Portereiko, M. F., Lloyd, A., and Drews, G. N. (2008). The AGL62 MADS domain protein regulates cellularization during endosperm development in *Arabidopsis*. The Plant Cell 20(3), 635–647. DOI: 10.1105/tpc.107.055137

Kaźmierczak, A (2010). Gibberellic acid and ethylene control male sex determination and development of *Anemia phyllitidis* gametophytes. In: Fernández, H., Kumar, A. and Revilla, M. A. (eds.) Working with Ferns. Issues and Applications. United States of America: Springer, 49–65. ISBN: 978-1-4419-7161-6

Khan, B. R., Faure, L., Chapman, K. D., and Blancaflor, E. B. (2017). A chemical genetic screen uncovers a small molecule enhancer of the N-acylethanolamine degrading enzyme, fatty acid amide hydrolase, in *Arabidopsis*. Sci. Rep. 7(41121). DOI: 10.1038/srep41121

Kim, J. Y., Yuan, Z., and Jackson, D. (2003). Developmental regulation and significance of KNOX protein trafficking in *Arabidopsis*. Development 130(18), 4351–4362. DOI: 10.1242/dev.00618

Kinosian, S. P., and Wolf, P. G. (2022). The natural history of model organisms: the biology of *C. richardii* as a tool to understand plant evolution. elife 11, 1–12. DOI: 10.7554/eLife.75019

Krieg, C.P., and Chambers, S.M. (2022). The ecology and physiology of fern gametophytes: A methodological synthesis. Appl Plant Sci. 2022, 10(2):e11464. DOI: 10.1002/aps3.11464.

Li, F., Chung, T., and Vierstra, R. D. (2014). AUTOPHAGY-RELATED11 plays a critical role in general autophagy- and senescence-induced mitophagy in *Arabidopsis*. The Plant Cell 26(2), 788–807. DOI: 10.1105/tpc.113.120014

Li, N., Yuan, L., Liu, N., Shi, D., Li, X., Tang, Z., Liu, J., Sundaresan, V., and Yang, W.-C. (2009). *SLOW WALKER2*, a NOC1/MAK21 homologue, is essential for coordinated cell cycle progression during female gametophyte development in *Arabidopsis*. Plant Physiol. 151(3), 1486–1497. DOI: 10.1104/pp.109.142414

Liu, H.-M., Dyer, R. J., Guo, Z.-Y., Meng, Z., Li, J.-H., and Schneider, H. (2012). The evolutionary dynamics of apomixis in ferns: a case study from polystichoid ferns. J. Bot. 1–11. DOI: 10.1155/2012/510478

Liu, M., Shi, D.-Q., Yuan, L., Liu, J., and Yang, W.-C. (2010). SLOW WALKER3, encoding a putative DEAD-box RNA helicase, is essential for female gametogenesis in *Arabidopsis*. J. Integr. Plant Biol. 52(9), 817–828. DOI: 10.1111/j.1744-7909.2010.00972.x

López, R. A., and Renzaglia, K. S. (2014). Multiflagellated sperm cells of *Ceratopteris richardii*are bathed in arabinogalactan proteins throughout development. Amer. J. Bot. 101(12), 2052–2061. DOI: 10.3732/ajb.1400424

Lukowitz, W., Roeder, A., Parmenter, D., and Somerville, C. (2004). A MAPKK kinase gene regulates extra-embryonic cell fate in *Arabidopsis*. Cell. 116(1):109–19. DOI: 10.1016/s0092-8674(03)01067-5.

Markham, K., Chalk, T. and C. Neal Stewart Jr. (2006). Evaluation of fern and moss protein-based defenses against phytophagous insects. Int. J. Plant Sci. 167 (1), 111–117.

Marchant, D. B., Chen, G., Cai, S., Chen, F., Schafran, P., Jenkins, J., Shu, S., Plott, C., Webber, J., Lovell, J. T., He, G., Sandor, L., Williams, M., Rajasekar, S., Healey, A., Barry, K., Zhang, Y., Sessa, E., Dhakal, R. R., Wolf, P. G., Harkess, A., Li, F.-W., Rossner, C., Becker, A., Gramzow, L., Xue, D., Wu, Y., Tong, T., Wang, Y., Dai, F., Hua, S., Wang, H., Xu, S., Xu, F., Duan, H., Theiben, G., McKain, M. R., Li, Z., McKibben, M. T. W., Barker, M. S., Schmitz, R. J., Stevenson, D. W., Zumajo-Cardona, C., Ambrose, B. A., Leebens-Mack, J. H., Grimwood, J., Schmutz, J., Soltis, P. S., Soltis, D. E., and Chen, Z.-H. (2022). Dynamic genome evolution in a model fern. Nature Plants 8, 1038–1051. DOI: 10.1038/s41477-022-01226-7

Martín-Blázquez, R., Bakkali, M., Ruiz-Estévez, M., Garrido-Ramos, M.A. (2023) Comparison between the gametophyte and the sporophyte transcriptomes of the endangered fern *Vandenboschia speciosa*. Genes 2023, 14, 166. DOI: 10.3390/genes14010166

McCarthy, F. M., Gresham, C. R., Buza, T. J., Chouvarine, P., Pillai, L. R., Kumar, R., Ozkan, S., Wang, H., Manda, P., Arick, T., Bridges, S. M., and Burgess, S. C. (2011). AgBase: supporting functional modeling in agricultural organisms. Nucleic Acids Res. 39, D497–D506. DOI: 10.1093/nar/gkq1115

Menéndez, V., Villacorta, N. F., Revilla, M. A., Gotor, V., Bernard, P., and Fernández, H. (2006). Exogenous and endogenous growth regulators on apogamy in *Dryopteris affinis* (Lowe) Fraser-Jenkins ssp. *affinis*. Plant Cell Rep. 25, 85–91. DOI: 10.1007/s00299-005-0041-1

Mercier, R., Vezon, D., Bullier, E., Motamayor, J. C., Sellier, A., Lefevre, F., Pelletier, G., and Horlow, C. (2001). SWITCH1 (SWI1): a novel protein required for the establishment of sister chromatid cohesion and for bivalent formation at meiosis. Genes Dev. 15, 1859–1871. DOI: 10.1101/gad.203201

Muller, M., and Munné-Bosch, S. (2015). Ethylene response factors: a key regulatory hub in hormone and stress signalling. Plant Physiol. 169(1), 32–41. DOI: 10.1104/pp.15.00677

Murashige, T., and Skoog, F. (1962). A revised medium for rapid growth and bioassays with tobacco tissue cultures. Plant Physiol. 15, 473–497. DOI: 10.1111/j.1399-3054.1962.tb08052.x

Nassrallah, A., Rougee, M., Bourbousse, C., Drevensek, S., Fonseca, S., Iniesto, E., Ait-Mohamed, O., Deton-Cabanillas, A.-F., Zabulon, G., Ahmed, I., Stroebel, D., Masson, V., Lombard, B., Eeckhout, D., Gevaert, K., Loew, D., Genovesio, A., Breyton, C., De Jaeger, G., Bowler, C., Rubio, V., and Barneche, F. (2018). DET1-mediated degradation of a SAGA-like deubiquitination module controls HS2Bub homeostasis. eLife 7, e37892. DOI: 10.7554/eLife.37892

Nogler G. (1984). Gametophytic apomixis. In: Johri B (ed) Embryology of angiosperms. Springer Verlag, Germany, 475–518.

Ohad, N., Yadegari, R., Margossian, L., Hannon, M., Michaeli, D., Harada, J. J., Goldberg, R. B., and Fischer, R. L. (1999). Mutations in *FIE*, a WD polycomb group gene, allow endosperm development without fertilization. Plant Cell 11, 407–416. DOI: 10.1105/tpc.11.3.407

Olmedo-Monfil, V., Durán-Figueroa, N., Arteaga-Vázquez, M., Demesa-Arévalo, E., Autran, D., Grimanelli, D., Slotkin, R. K., Martienssen, R. A., and Vielle-Calzada, J.-P. (2010). Control of female gamete formation by a small RNA pathway in *Arabidopsis*. Nature 464, 628–632. DOI: 10.1038/nature08828

Ojosnegros, S., Alvarez, J. M., Grossmann, J., Gagliardini, V., Quintanilla, L. G., Grossniklaus, U., and Fernández, H. (2022). The shared proteome of the apomictic fern *Dryopteris affinis* ssp. *affinis* and its sexual relative *Dryopteris oreades*. Int. J. Mol. Sci. 23(22), 14027. DOI: 10.3390/ijms232214027

Ojosnegros, S., Alvarez, J. M., Grossmann, J., Gagliardini, V., Quintanilla, L. G., Grossniklaus, U., and Fernández, H. (2023). Proteome and interactome linked to metabolism, genetic information processing, and abiotic stress in gametophytes of two woodferns. Int. J. Mol.Sci. 24(15), 12429. DOI: 10.3390/ijms241512429

Pangua, E., Belmonte, R. and Pajarón, S. (2009). Germination and reproductive biology in salty conditions of *Asplenium marinum* (Aspleniaceae), a European coastal fern. Flora - Morphology, Distribution, Functional Ecology of Plants. 204, (9), 673–684. ISSN 0367-2530. DOI: org/10.1016/j.flora.2008.09.007.

Plackett, A. R. G., Di Stilio, V. S., and Langdale, J. A. (2015). Ferns: the missing link in shoot evolution and development. Front. Plant. Sci. 6, 972. DOI: 10.3389/fpls.2015.00972

Podio, M., Cáceres, M. E., Samoluk, S. S., Seijo, J. G., Pessino, S. C., Ortiz, J. P. A., and Pupilli, F. (2014). A methylation status analysis of the apomixis-specific region in *Paspalum* spp. suggests an epigenetic control of parthenogenesis. J. Exp.Bot. 65(22), 6411–6424. DOI: 10.1093/jxb/eru354

PPGI (2016). A community-derived classification for extant lycophytes and ferns. 54 (6), 563–603. DOI: 10.1111/jse.12229

Rabiger, D. S., Taylor, J. M., Spriggs, A., Hand, M. L., Henderson, S. T., Johnson, S. D., Oelkers, K., Hrmova, M., Saito, K., Suzuki, G., Mukai, Y., Carroll, B. J., and Koltunow, A. M. G. (2016). Generation of an integrated *Hieracium* genomic and transcriptomic resource enables exploration of small RNA pathways during apomixis initiation. BMC Biol. 14(86). DOI: 10.1186/s12915-016-0311-0

Rabut, G., and Peter, M. (2008). Function and regulation of protein neddylation. “Protein modifications: beyond the usual suspects’ review series”. EMBO Rep. 9(10), 969–976. DOI: 10.1038/embor.2008.183

Rathinasabapathi, B. (2006). Ferns represent an untapped biodiversity for improving crops for environmental stress tolerance. New Phytol. 172(3), 385–390. DOI: 10.1111/j.1469-8137.2006.01889.x

Raven, P. H., Evert, R. F., and Eichhorn, S. E. (2005). Biol. Plants. United States of America: W. H. Freeman and Company.

Ravi, M., Marimuthu, M. P. A., and Siddiqi, I. (2008). Gamete formation without meiosis in *Arabidopsis*. Nature 451, 1121–1124. DOI: 10.1038/nature06557

Rivera, A., Cañal, M. J., Grossniklauss, U., and Fernández, H. (2018). The gametophyte of fern: born to reproducing. In: Fernández, H. (ed.) Current Advances in Fern Research, United States of America: Springer International Publishing, 3–19. ISBN 978-3-319-75102-3

Salmi, M. L., and Bushart, T. J. (2010). Cellular, molecular, and genetic changes during the development of *Ceratopteris richardii* gametophytes. In: Fernández, H., Kumar, A., and Revilla, M. A. (eds.) Working with Ferns. Issues and Applications. United States of America: Springer, 11–24. ISBN 978-1-4419-7161-6

Salmi, M. L., Bushart, T. J., Stout, S. C., and Roux, S. J. (2005). Profile and analysis of gene expression changes during early development in germinating spores of *Ceratopteris richardii*. Plant Physiol. 138(3), 1734–1745. DOI: 10.1104/pp.105.062851

Sareen, B., Thapa, P., Joshi, R., and Bhattacharya, A. (2019). Proteome analysis of the gametophytes of a Western Himalayan fern *Diplazium maximum* reveals their adaptive responses to changes in their micro-environment. Front. Plant Sci. 10, 1623. DOI: 10.3389/fpls.2019.01623

Savidan, Y., Carman, J. G., and Dresselhaus, T. (eds). (2001) The flowering of apomixis: from mechanisms to genetic engineering. CIMMYT, IRD, European Comission DG VI (FAIR), Mexico.

Schmidt, A. (2020). Controlling apomixis: shared features and distinct characteristics of gene regulation. Genes 11, 329. DOI: 10.3390/genes11030329

Schneider, H. (2013). Evolutionary morphology of ferns (Monilophytes). In: The Evolution of Plant Form. Ann. Plant Rev. Vol. 45. Ed. Barbra A. Ambrose and Michael Perugganan. Clackwell Publishing Ltd. DOI: 10.1002./9781118305881.ch4

Shi, D.-Q., Liu, J., Xiang, Y.-H., Ye, D., Sundaresan, V., and Yang, W. C. (2005). SLOW WALKER1, essential for gametogenesis in *Arabidopsis*, encodes a WD40 protein involved in 18S ribosomal RNA biogenesis. Plant Cell 17, 2340–2354. DOI: 10.1105/tpc.105.033563

Siddiqi, I., Ganesh, G., Grossniklaus, U., and Subbiah, V. (2000). The *dyad* gene is required for progression through female meiosis in *Arabidopsis*. Development 127, 197–207. DOI: 10.1242/dev.127.1.197

Smith, Z., and Long, J. (2010). Control of *Arabidopsis* apical-basal embryo polarity by antagonistic transcription factors. Nature 464, 423–426. DOI: 10.1038/nature08843

Spillane, C., Curtis, M. D., and Grossniklaus, U. (2004) Apomixis technology development-virgin births in farmers’fields? Nat. Biotechnol. 22, 687–691. DOI: 10.1038/nbt976

Sridhar, V. V., Kapoor, A., Zhang, K., Zhu, J., Zhou, T., Hasegawa, P. M., Bressan, R. A., and Zhu, J.-K. (2007). Control of DNA methylation and heterochromatic silencing by histone H2B deubiquitination. Nature 447, 735–738. DOI: 10.1038/nature05864

Suo, J., Zhao, Q., Zhang, Z., Chen, S., Cao, J., Liu, G., Wei, X., Wang, T., Yang, C., and Dai, S. (2015). Cytological and proteomic analyses of *Osmunda cinnamomea* germinating spores reveal characteristics of fern spore germination and rhizoid tip growth. Mol. Cell. Proteomics 14, 2510–2534. DOI: 10.1074/mcp.M114.047225

Szklarczyk, D., Gable, A. L., Lyon, D., Junge, A., Wyder, S., Huerta-Cepas, J., Simonovic, M., Doncheva, N. T., Morris, J. H., Bork, P., Jensen, L. J., and Mering, C. V. (2019). STRING v11: protein-protein association networks with increased coverage, supporting functional discovery in genome-wide experimental datasets. Nucleic Acids Res. 8(47), D607–D613. DOI: 10.1093/nar/gky113

Terceros, G. C., Resentini, F., Cucinotta, M., Manrique, S., Colombo, L., and Mendes, M. A. (2020). The importance of cytokinins during reproductive development in *Arabidopsis* and beyond. Int. J. Mol. Sci. 21(21), 8161. DOI: 10.3390/ijms21218161

Thagela, P., Yadav, R. K., Dahuja, A., Singh, P. K., and Abraham, G. (2016). Physiological and proteomic changes in *Azolla microphylla* roots upon exposure to salinity. Indian J. Biot. 15, 101–106.

Valledor, L., Menéndez, V., Canal, M. J., Revilla, A., and Fernández, H. (2014). Proteomic approaches to sexual development mediated by antheridiogen in the fern *Blechnum spicant* L. Proteomics 14(17-18), 2061–2071. DOI: 10.1002/pmic.201300166

Vries, J. de, Fischer, A. M., Roettger, M., Rommel, S., Schluepmann, H., Bräutigam, A, Carlsbecker, A., and Gould, S. B. (2016). Cytokinin induced promotion of root meristem size in the fern *Azolla* supports a shoot-like origin of euphyllophyte roots. New Phytol. 209(2), 705–720. DOI: 10.1111/nph.13630

Vries, S. de, Vries, J. de, Teschke, H., Dahlen, J. von, Rosa, L., and Gould, S. B. (2018). Jasmonic and salicylic acid response in the fern *Azolla filiculoides* and its cyanobiont. Plant Cell Environ. 41(11), 2530–2548. DOI: 10.1111/pce.13131

Wada, M. (2007). The fern as a model system to study photomorphogenesis. J. Plant Res. 120(1), 3–16. DOI: 10.1007/s10265-006-0064-x

Wang, X., Chen, S., Zhang, H., Shi, L., Cao, F., Guo, L., Xie, Y., Wang, T., Yan, X., and Dai, S. (2010). Desiccation tolerance mechanism in resurrection fern-ally *Selaginella tamariscina* revealed by physiological and proteomic analysis. J. Proteome Res. 9, 6561–6577. DOI: 10.1021/pr100767k

Wei, B., Zhang, J., Pang, C., Yu, H., Guo, D., Jiang, H., Ding, M., Chen, Z., Tao, Q., Gu, H., Qu, L.-J., and Qin, G. (2015). The molecular mechanism of SPOROCYTELESS/NOZZLE in controlling *Arabidopsis* ovule development. Cell Res. 25, 121–134. DOI: 10.1038/cr.2014.145

Weinhofer, I., Hehenberger, E., Roszak, P., Hennig, L., and Kohler, C. (2010). H3K27me3 profiling of the endosperm implies exclusion of Polycomb group protein targeting by DNA methylation. PLOS Genetics. DOI: 10.1371/journal.pgen.1001152

Withers, K.A., Kvamme, A., Youngstrom, C.E., Yarvis, R.M., Orpano, R., Simons, G.P., Irish, E.E., and Cheng, C.-L. (2023). Auxin involvement in *Ceratopteris* gametophyte meristem regeneration. Int. J. Mol. Sci. 24, 15832. DOI: 10.3390/ijms242115832

Wu, X., Chory, J., and Weigel, D. (2007). Combinations of *WOX* activities regulate tissue proliferation during *Arabidopsis* embryonic development. Developmental Biol. 309(2), 306–316. DOI: 10.1016/j.ydbio.2007.07.019

Wu, X.; Liu, X.; Zhang, S.; and Zhou, Y. (2023) Cell Division and meristem dynamics in fern gametophytes. Plants, 12, 209. DOI: 10.3390/plants12010209

Wyder, S., Rivera, A., Valdés, A. E., Cañal, M. J., Gagliardini, V., Fernández, H., and Grossniklaus, U. (2020). Differential gene expression profiling of one- and two-dimensional apogamous gametophytes of the fern *Dryopteris affinis* ssp. *affinis*. Plant Physiol. Biochem. 148, 302–311. DOI: 10.1016/j.plaphy.2020.01.021

Xu, C., and Min, J. (2011). Structure and function of WD40 domain proteins. Protein Cell 2(3), 202–214. DOI: 10.1007/s13238-011-1018-1

Xu, Y., Jia, H., Tan, C., Wu, X., Deng, X., and Xu, Q. (2022). Apomixis: genetic basis and controlling genes. Hort. Res.9, uhac150. DOI: 10.1093/hr/uhac150

Yao, S., Chan, J., Xu, Y., Wu, S., and Zhang, L. (2022). Divergences of the *RLR* gene families across Lophotrochozoans: domain grafting, exon-intron structure, expression, and positive selection. Int. J. Mol. Sci. 23(3415). DOI: 10.3390/ijms23073415

Yoshimoto, K., Shibata, M., Kondo, M., Oikawa, K., Sato, M., Toyooka, K., Shirasu, K., Nishimura, M., and Ohsumi, Y. (2014). Organ-specific quality control of plant peroxisomes is mediated by autophagy. J. Cell Sci. 127(6), 1161–1168. DOI: 10.1242/jcs.139709

Youngstrom, C. E., Geadelmann, L. F., Irish, E. E., and Cheng, C.-L. (2019). A fern *WUSCHEL-RELATED HOMEOBOX* gene functions in both gametophyte and sporophyte generations. BMC Plant Biol. 19, 1–13. DOI: 10.1186/s12870-019-1991-8

Yun, J., Kim, Y. S., Jung, J. H., Seo, P. J., and Park, C. M. (2012). The AT-hook motif-containing protein AHL22 regulates flowering initiation by modifying FLOWERING LOCUS T chromatin in *Arabidopsis*. Plant Biol. 287(19), 15307–15316. DOI: 10.1074/jbc.M111.318477

Zhang, X., Gonzalez-Carranza, Z. H., Zhang, S., Miao, Y., Liu, C.-J., and Roberts, J. A. (2019). F-Box proteins in plants. Annual Plant Reviews Online 2(1). DOI: 10.1002/9781119312994.apr0701

Zheng, B., Xing, K., Zhang, J., Liu, H., Ali, K., Li, W., Bai, Q., and Ren, H. (2022). Evolutionary analysis and functional identification of ancient brassinosteroid receptors in *Ceratopteris richardii*. Int. J. Mol. Sci. 23(12), 6795. DOI: 10.3390/ijms23126795

Zrenner, R., Riegler, H., Marquard, C. R., Lange, P. R., Geserick, C., Bartosz, C. E., Chen, C. T., and Slocum, R. D. (2009). A functional analysis of the pyrimidine catabolic pathway in *Arabidopsis*. New Phytologist 183(1), 117–132. DOI: 10.1111/j.1469-8137.2009.02843.x

Zumajo-Cardona, C., Vasco, A., and Ambrose, B. A. (2019). The evolution of the *KANADI* gene family and leaf development in lycophytes and ferns. Plants 8(9), 313. DOI: 10.3390/plants8090313

