## Supplementary Figure 1 for "Transcriptomic analyses in the gametophyte of *Dryopteris affinis*: apomixis and more"

CULLIN 4 (*CUL4*):

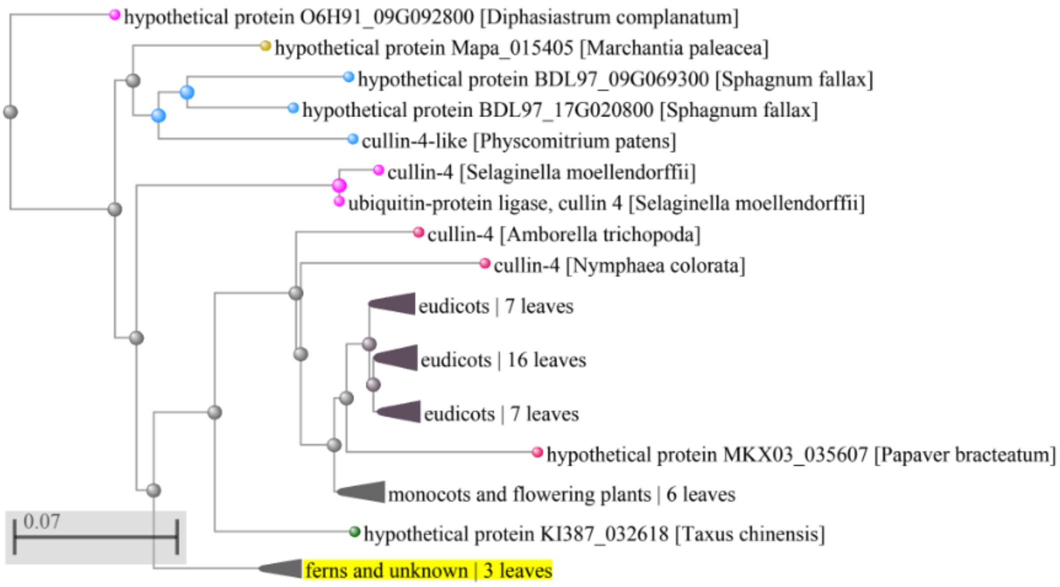


XAP5 CIRCADIAN TIMEKEEPER (*XCT*):

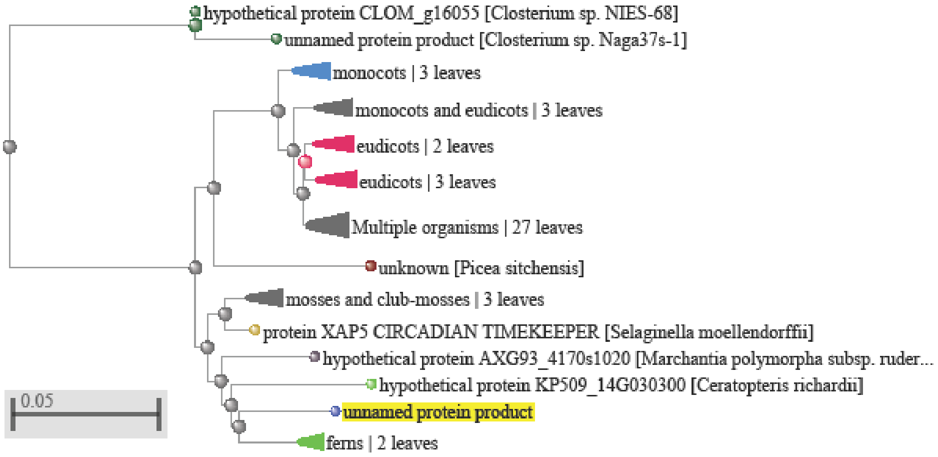


TIMEKEEPER LOCUS1 (*STIPL1*):

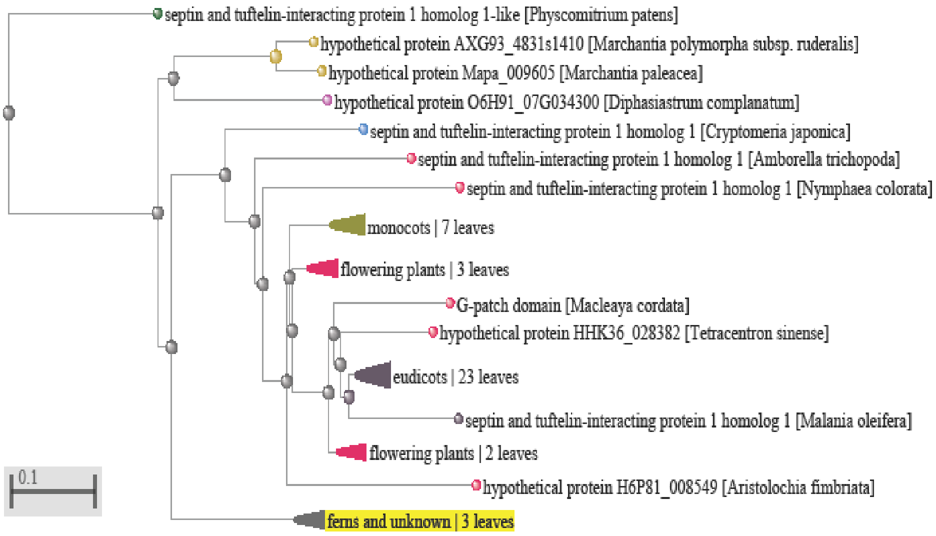

EMBRYONIC FACTOR1 (*FAC1*):

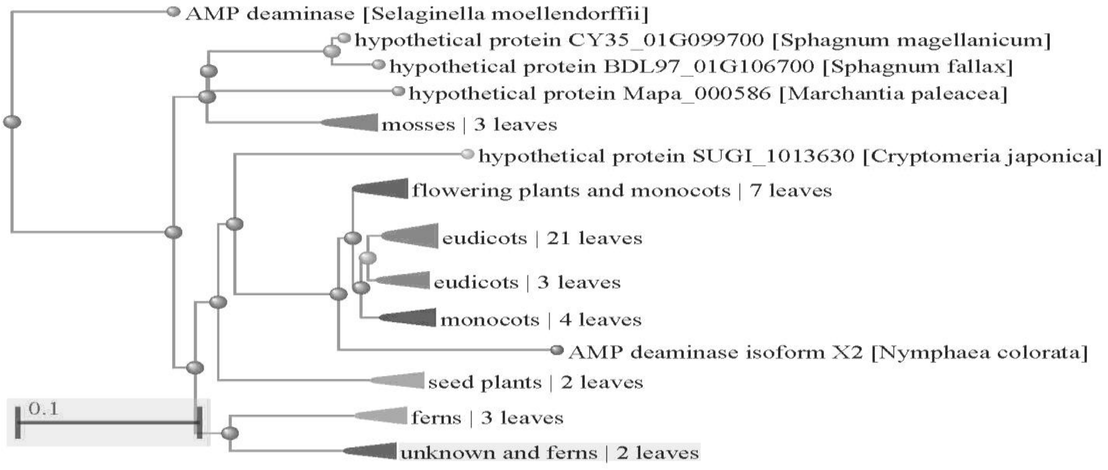


GRAVITROPISM DEFECTIVE 2 (*GRV2*):

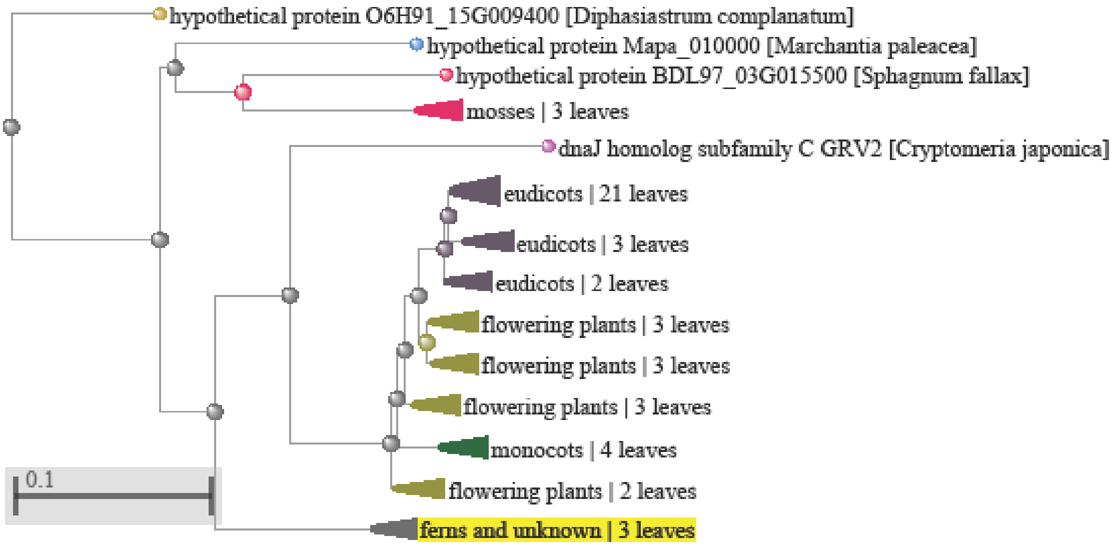


UBIQUITIN-SPECIFIC PROTEASE 26 (*UBP26*):

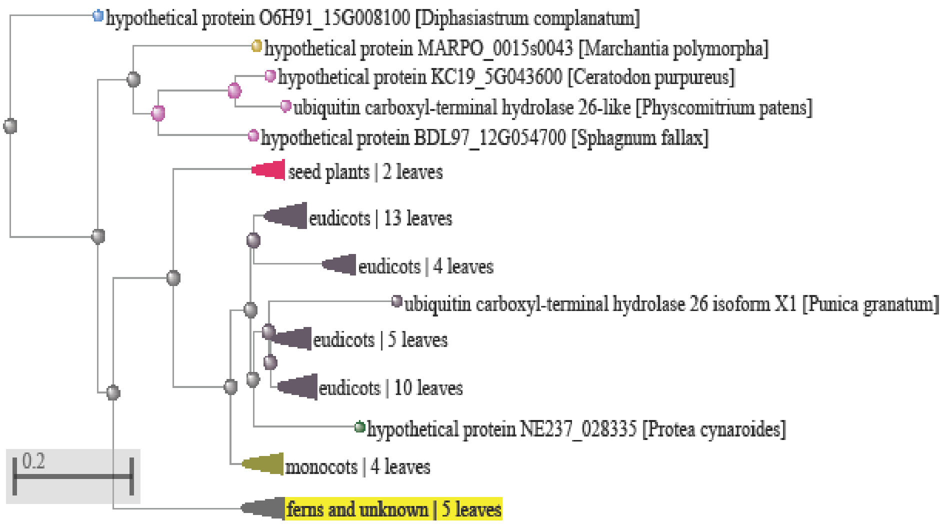

HASTY (*HST*):

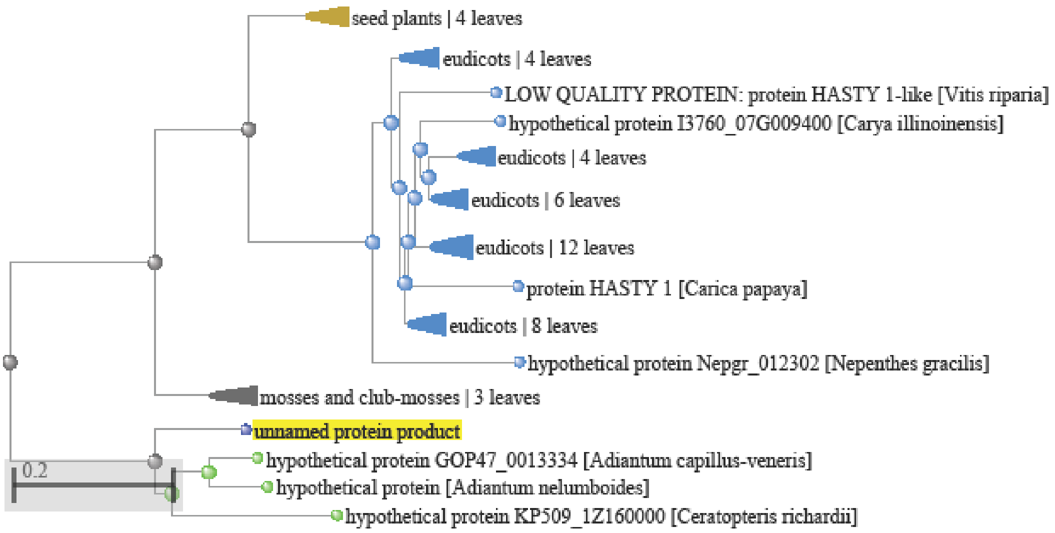


GLIOMAS 41 (*GAS41*):

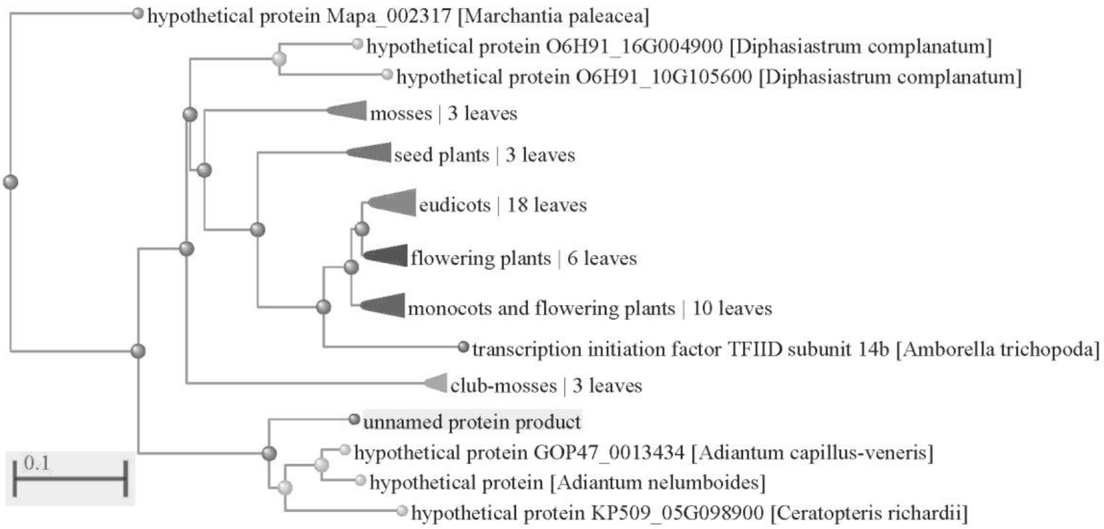


SEUSS (*SEU*):

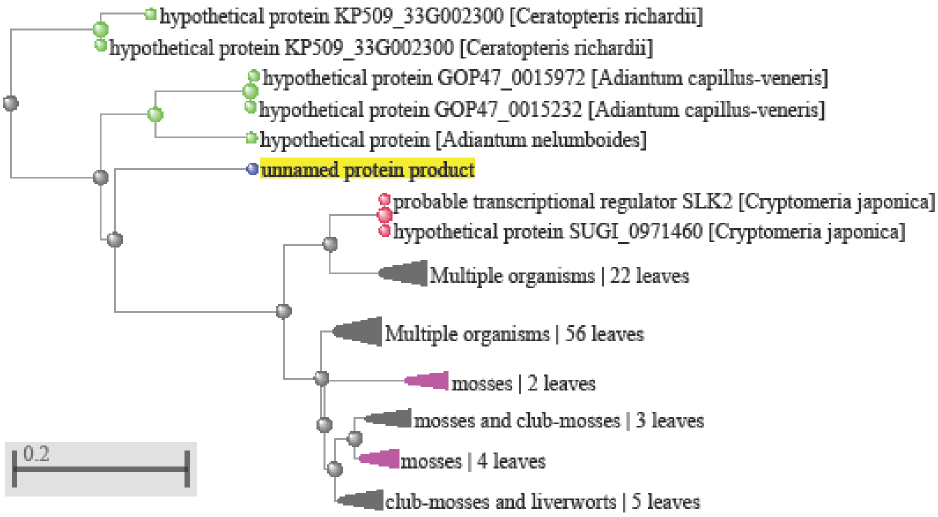

UNUSUAL FLORAL ORGANS (*UFO*):

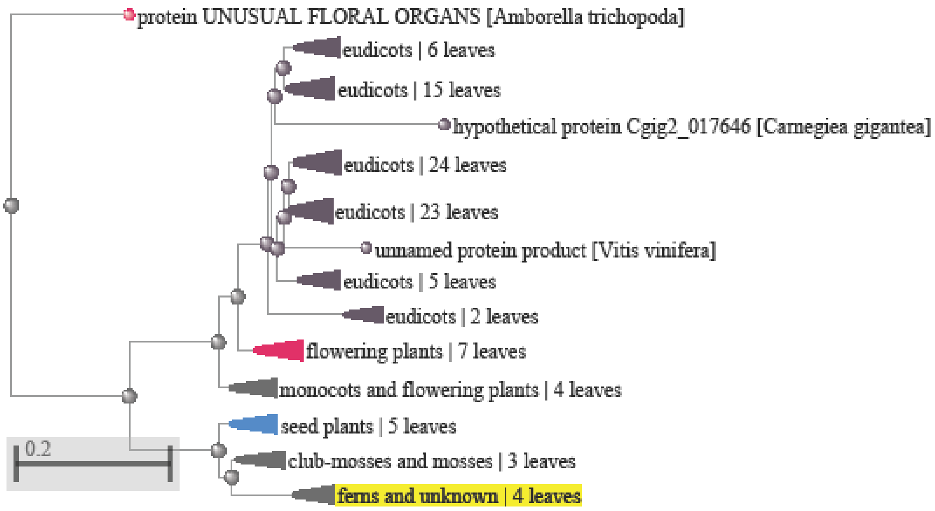


VERNALIZATION INSENSITIVE 3 (*VIN3*):

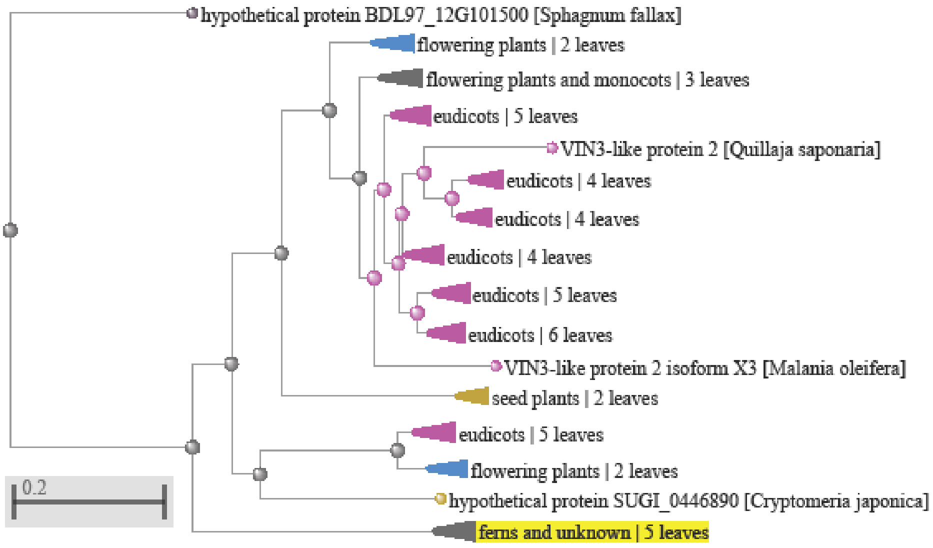


EXPRESSION OF OSMOTICALLY RESPONSIVE GENES 1 (*HOS1*):

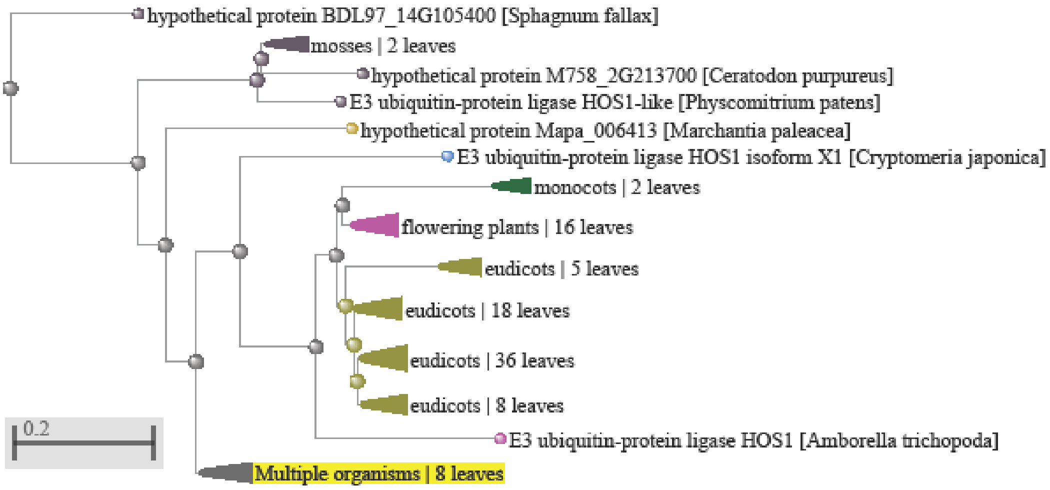

ULTRAVIOLET HYPERSENSITIVE 1 (*UVH1*):

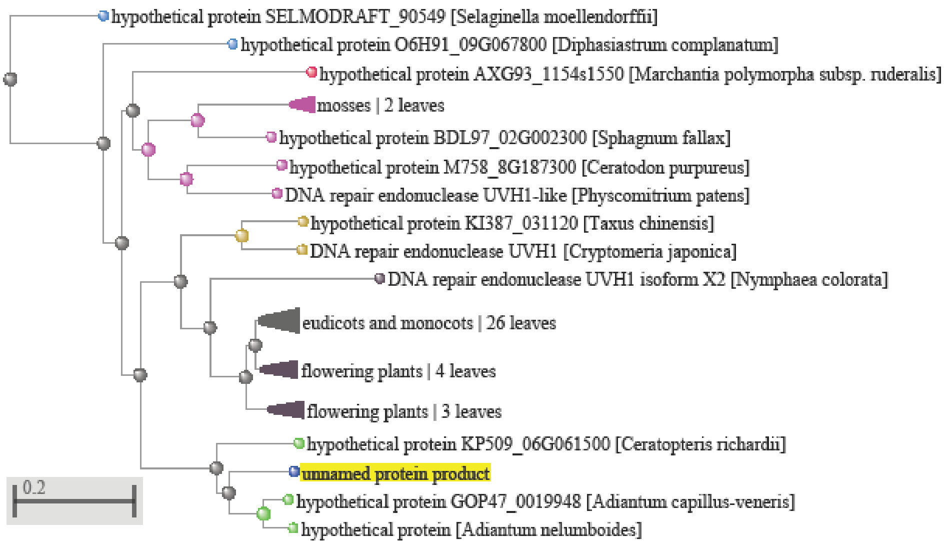


**Supplementary Figure 1**. Phylogenetic trees corresponding to proteins associated with vegetative and reproductive development, as well as with response to stress from the apogamous gametophyte of the fern *Dryopteris affinis* ssp. *affinis*. The yellow line indicates our sequence. Distance scale represents the number of differences between sequences (0.1=10%).
