## Supplementary Figure 2 for "Transcriptomic analyses in the gametophyte of *Dryopteris affinis*: apomixis and more"

CULLIN 4 (*CUL4*):

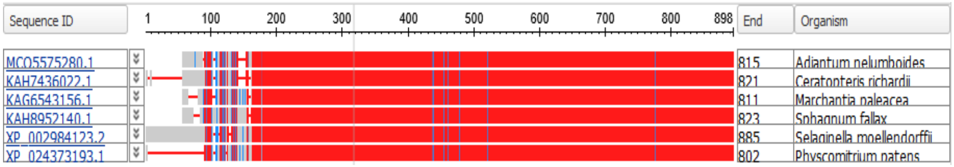


XAP5 CIRCADIAN TIMEKEEPER (*XCT*):

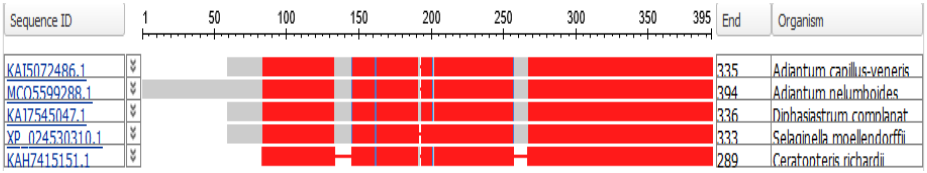


TIMEKEEPER LOCUS1 (*STIPL1*):

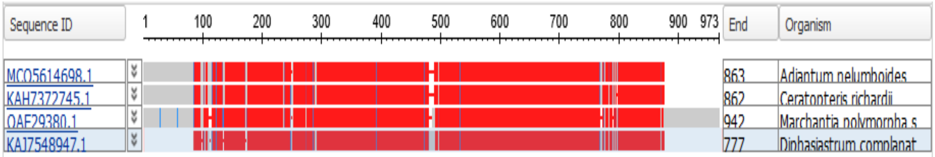


EMBRYONIC FACTOR1 (*FAC1*):

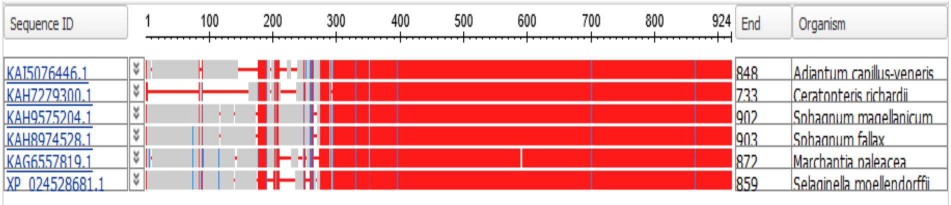


GRAVITROPISM DEFECTIVE 2 (*GRV2*):

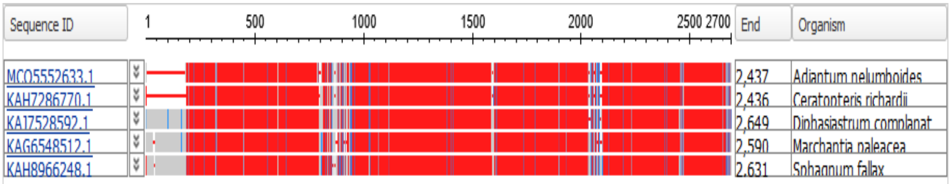


UBIQUITIN-SPECIFIC PROTEASE 26 (*UBP26*):

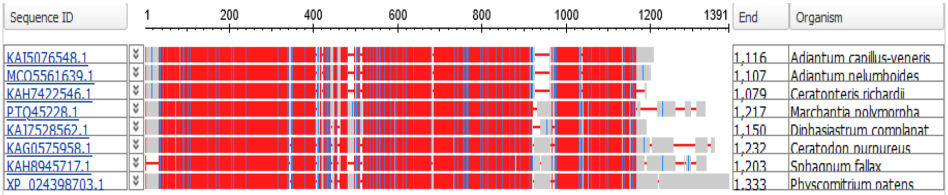


HASTY (*HST*):

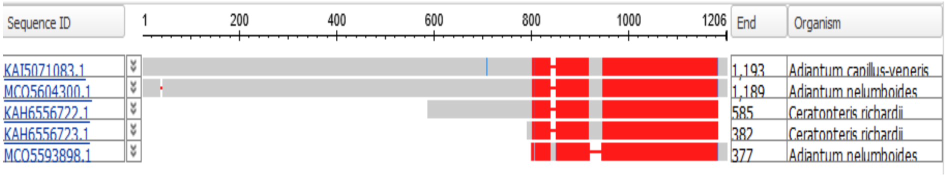

GLIOMAS 41 (*GAS41*):

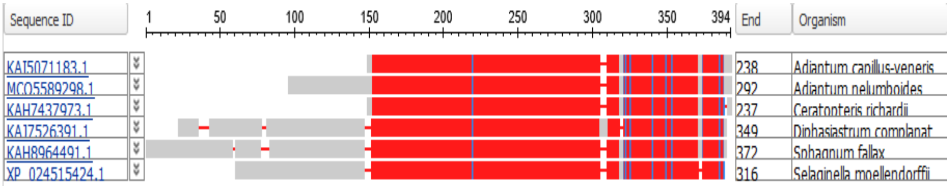


SEUSS (*SEU*):

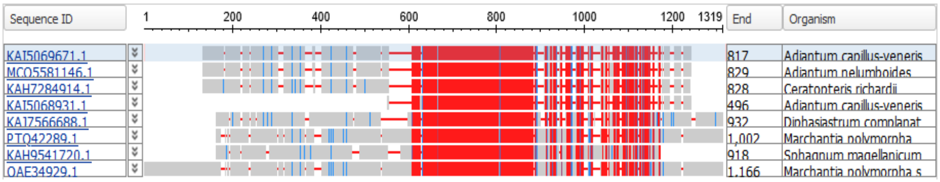


UNUSUAL FLORAL ORGANS (*UFO*):

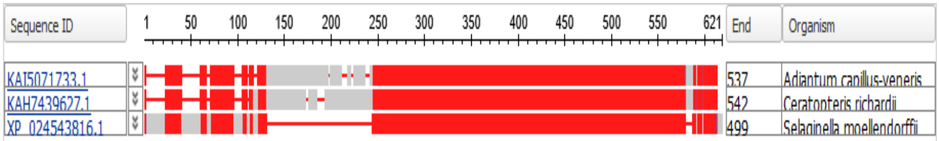


VERNALIZATION INSENSITIVE 3 (*VIN3*):

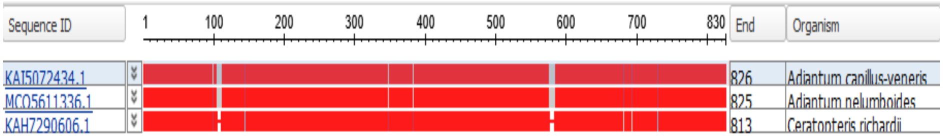


EXPRESSION OF OSMOTICALLY RESPONSIVE GENES 1 (*HOS1*):

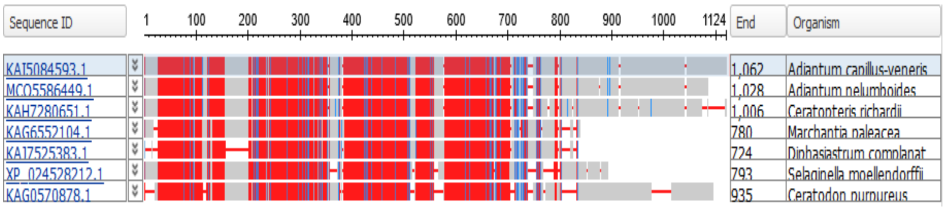


ULTRAVIOLET HYPERSENSITIVE 1 (*UVH1*):

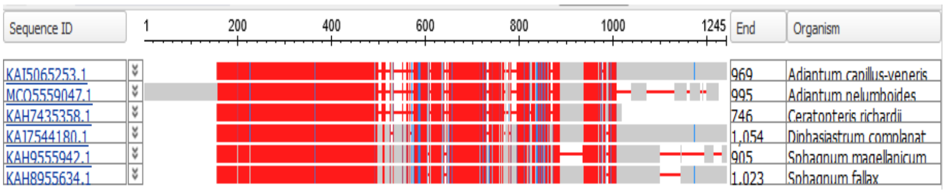


**Supplementary Figure 2**. Amino acid sequence alignments corresponding to proteins derived from the apogamous gametophyte of the fern *Dryopteris affinis* ssp. *affinis* associated with vegetative and reproductive development, as well as with response to stress. Red indicates highly conserved columns and blue indicates less conserved columns.
