## Supplementary Figure 3 for "Transcriptomic analyses in the gametophyte of *Dryopteris affinis*: apomixis and more"

CULLIN 4 (*CUL4*):

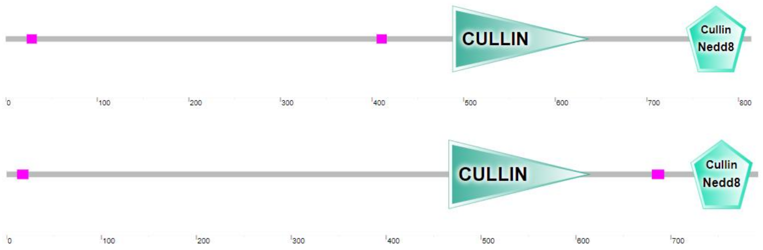


TIMEKEEPER LOCUS1 (*STIPL1*):

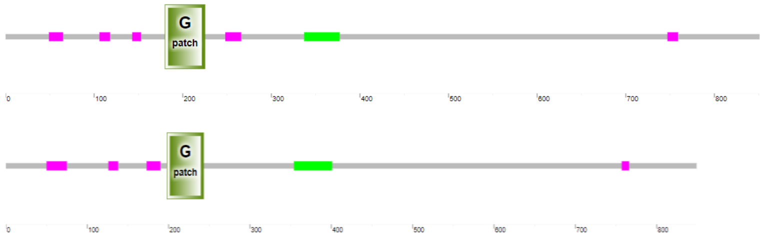


UBIQUITIN-SPECIFIC PROTEASE 26 (*UBP26*):

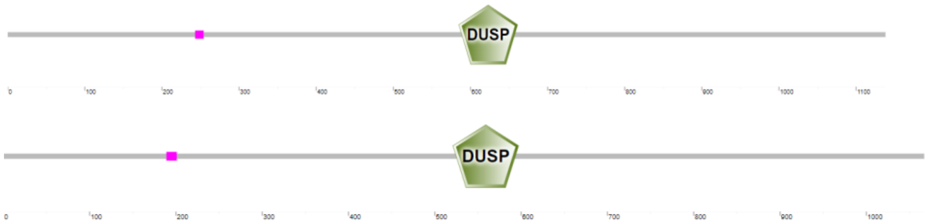


UNUSUAL FLORAL ORGANS (*UFO*):

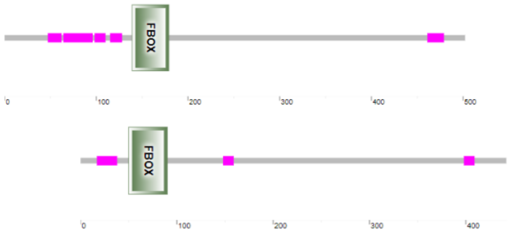


**Supplementary Figure 3**: Schematic representation of domains of proteins from gametophytes of the fern *Dryopteris affinis* above and from *Arabidopsis thaliana* below obtained with SMART version 9.0 software.
