## Supplementary Table 1 for "Transcriptomic analyses in the gametophyte of *Dryopteris affinis*: apomixis and more"

**Supplementary Table 1**. Nucleotide sequences of proteins found in gametophytes of the apogamous fern *Dryopteris affinis* ssp. *affinis*.

**ARGININE-RICH CYCLIN 1 (*RCY1*):**

TCTCTCTCTCTCTCTCTCTTCCTTCCAGCCCTAGCTAATGATTTACACCGCCATCGACACCTTCTACTTGTCGCAGGAGC
AGCTGCAGAATTCTCCCTCCCGCAAAGATGGCGTTGATGAGGAAACAGAAACTGTGCTTCGGCTCTATGGTTGCGAGCTCGTA
CAGGAGGGCGGCATACTTCTCAAGCTACCTCAAGCAGTCATGGCGACAGGCCAGGTGCTGCTTCATCGATTTTTCTGCAAGAA
GTCATTTGCCAGGTTCAATGTGAAGAGAGTAGCTGCAAGCTGTCTTTGGGTTGCAGCAAAACTTGAGGAGAGCCCAAAGAAG
GCCCAGCACGTCTTAAACGTGTTTAATAGAATGGACTGCCGTCGAGAAAACAAGCCTCTTGAACTGCTAGACCCTTTTTCTAA
GAGGTACGAAGAATTGAAGATTGATTTGATCAGGACAGAACGCCATCTTTTAAAAGAGATGGGTTTCATTTGTCATGTTGAGC
ATCCACACAAGTTTATTATCAATTACTTACTACAGCTTGAAACTCCACCAGAGTTGATGCAAGAAGCATGGAACTTAGCTAAT
GACAGCTTACGTACTACGCTTTCAGTTCGATTCAAAAGTGAAGTGGTGGCTTGTGGCGTTGTCTATGCTGCTGCGCGGAGGTT
CAATGTCAGTTTGCCTGAAAGTCCTCCTTGGTGGAAGGTATTTGATGCGGAGAAGGGTGATATTGAGGAAGTATGCAATGTCC
TTGCAAACCTCTACAAGAGGCCCAAAGCGAGCTATATTGAAGTGTCAAAAGATTCCAAGTCTTTTGTGTTAAGCAGTAGAGCT
TGGGAACCTTCTGCAAGTGTGAAGGATTCTCTCGGAGGTTCTGTAATTGCAAATGGAAGCAAACAGGGCCAGATGACCACAG
AAACACGAGAGTTTATGGTTAAAGTAGCCCTAGATAAACTGAAGAACACTTCTGCACAGGATTCAAGGAGACCTTCTGACGA
GGCCAGGGCTTCTTCTGCCAATGGGGAGCTTTCAGACGAGTTAGTCAAGGGAGCTGAGGAATTGGACGAGTTTACGCCTGAT
AGAAACAAGGAGTCTGCAGACACCGAACTCAGGATTAAAGAGAAGGACAAAGACAGGGCAAAGGCTCGCGAAAAGGGAAG
AGAAAAAGATAGGCAGCGAGAGTATGAGAAGATTAGGGACAAAGATCGAGGAAAGATTCGAGAAAGTGAGTATGAAAGAG
ATCGAGATCGAAAAGAGAGGGAGCATCGCAAGGAAAGGACAAAAGAATCAGGTTATGCTGACAAGAGTAGACATCATGGAT
CAAGCCGAGAAAGGGAGTATCATAGTTCATACAAGCTTTTCCGTGAAAAGGACCGACATCGGCATCATCCTTACTCATGAGA
AAGTCTACGAAGATTTAGATTTTACTTTCTTGTAGCTTAAAAGTTATCATCCAGTATGGAAGGCTTTTATTTGATGGTGCTGAT
CAGTGATTGTTTGGTCAGTAGCTTAAACAACTGTTTCTTACAGTTTCCCTTAGTGGAGACTTCCTTGTAGAGCTGACTTTTGCT
CGGTGAGCCGCACTGGATGATAAAAGAATGGAGAGTTGGGAAGGTTGCTGTTCAAGCTGATCTGAGGAGCTGCTTCATATAC
TTCACGCCCCTCGTAACAACACAGCACACTGTGCTCATGAGGGGACAATGTGGCAGGCCATTCTAGTTCTGATCTGGTGGTAA
GTGCGGATGGACCACCTTGTGCTGCAATCGAAGAATGCAGCAGGATGTAAGATCACATCAGACGGGTTTCCAGTAAGTGGAA
CTGCATGTTTTTATCAAATGCTTTATGGGTTCTGGAGCATGCCTCTATATAAAATAGTAAAGCAAAATGGGTTATTGTATACCT
TTTATGATTGTGGGTGTTTTCGCTATGTTTTTCTACTTTGGATGGGGAATACTTGAAGGCTTTATTATTGTTGGAAGTAGCAGG
AGCTGCTGGAATTTCATTAATACTGGATGCTCATAGTTGTCCTTGCAACTATTTCAAGCATACCTGCCATGCGAGAGCATGGA
GTCTTTCATTTAAAGATCCAAAGCTAGAGCAATGACCACTTCACATAGATCGCAAGGATAGAATTTATAACTTTTGATGCATG
AAAGTGGAAACAATATAACTCTGTATTTACCGCTCCATCATCATCTGATTGTATGGTAGGTCAGCAAGCACCTAGCCATTAAT
GTTATCTAGCAATTTAGCCAATGCAGGCATTCTTGATTTTGGCGACCTTCAGTACACAACTTTTTGTTTGCATGCACAGTCGAG
GGAGTTCTTGAGGGTCAAATTATCATCAGACGAATGACTCTCTAGATGGTCTATATTTGGAGTCGCAGAGACGCACATAATTG
ACTTGGATCTCCTGTTGGGGTTCTGGCATTTTGTTGAGTATAAATTGGTGTACAACCTTAAATATTAGATATTTCTGTCATGGT
TTCTTGTTTAAGGGGTACTATTGCTTGTGGTGTTTAAGCGAGTTATCATCAGTGGCTTTGTAGCTTAAATTGGTGCTAGGGGAG
GTTGTACAATTTGACAAGAAAGAAAGAAGCCCACAGAAACCTGCCTCCTTGGTTTGCATATTAAGAGTTGAGCAGGGTTGCT
CACTATTTAAGTTGAAGAGGTTATCCCTTTTTTTCAAACAACTAAATACCTGGCTAAGAAAAGTTTTCGAGTGTCTTATTGCTT
AATGTAACTTCTTATACATATAATGACTCGTAATGTTATCTGTACAATTGTGGGCCATAAGAAAAGCAGCACTTGTGAAGCAA
AACGTTTATTGGAGTTGAAAAGTGCCTCCTTGTCAATCAGATCATCTATATTGAATTGATTTATGATTTATCGATTGCTTTTTG
GTTTAATGAGATCTCAAATGGGGGATTTGTTGTGCTACCACCAAGGGAGTACCTAGTGATCGTAATGTGTGTAGATTACAAAT
TAGTAGGTCAAATAATCATTTTTCAGAGAGGAACTGGTCCAGCAGCTCAAAGCATTTCATACAAAGACTTCAGAGTCAGGATT
TGGTTATGTGAAATCGGTGAGTGAAATGACTTCATAGTCCTTCCTGTATATACCAACAAAGCTCAGTGCGACATGGCTTGTTT
TGAAAAGGCTGCTTGAGTGCCTAATTTTTTTTCTGTTTTAAGGGTTTTGCTTGTAATTATTCCATCCTTGAGGACAGTAGAGGT
ATTGCTTCACATTACCGGTAAATATTAGCTCTGCCCCTTTTCAGAAGGTTTTTGTTGCACCAAAAAATTTGTGAGATTTCAATG
CATAATAAGAATCTCATACCGTTTTTGTAATTATTGTTTTATTTTTCTAATGGAGAGGTTTCACCTTTGAGAATGTTGTATCAA
GAGGATTCATGTAATTCAATGTTGTGAGGTGTGAAACAAGTG

**KEULE (*KEU*):**

GCGAATTAGTGATCTCACTATTCATCTTGGCCGTGGCAGGCTCCTTGGTTTCATGGTCAATCCGTGAAACCTCCACTCG
TCTTCCGCCATTTTCTAAGAAGAGGAGAAGAAGACGTGCCGCCTTGCCCTCCTCTCTCGGAAGCTTCCTCTTCGTCTTCTTCCT
CTGCTCGAGCCCTCCTGCATCGCCCCTTTCGGAACTTCATTCTTCAATCCCTAATCCTGCAGAGCTTTGCAATACACAGATATT
TCTTAGTTTTCACGCAAAAGGAGATCAAGTTGTCTTCTGTAAGGTTTCCGCAACCCTAATCAACCATAGGAAGGGTCTGGTTC
TTCAAACTCTTGATTATCAGGTGCTCCAATCAAACTTTCTACTCTACATTTCATCAAGCAGAGGTTGGATGCAAAACATACTTG
TAACTCTGCAGAGCATCCAGAGATCATAAAAACCTAACATTATATTGCTTTTGCACTACGACACGCGGCTGCTTTATCAAAGT
GAAGAAGTTGCTACACTTGTTATCTACGAGGACGCCATAGCCATAGCCCTGCACGACATCTTCAACCTTTTGCCTTCACCATG
TCAGGATCGGAATCTTCTTTGCATTCTTGGAATTCTGGCAGCTATAAATCATTCCGCCAAGTTACTCGAGAACGTCTTCTTCAT
GAAATGCTCAGGCTGGCTAAAGGAAAGGATTCTTCGGATTGGAAGGTACTTATCATGGACGAGGTTACAGTGAAAGTGATGT
CATATTCTTGCAAGATGGCAGACATAACAGATGAAGGCATTTCATTGGTGGAGGACTTAAATAAGCGGAGACAACCACTTCC
AGCATTGGAAGCAGTATACTTCATTCAACCTAAAAGAGAAAGTGTTCAGAAGCTCAAGATGGACTTGTCTGGAAAAACACCT
CTTTATAAAAAGGCCCATGTGTTTTTCAGTTCGCCCATAGCTAGAGATCTGGTACAGGCCATCAAAAGTGATGCGCTGGTTTT
TTCCCGGCTTGCTACTCTTAGAGAGATGAATTTGGAATATCTCACGATTGATTCTCAGGCATTCTCAACTGATAATAGTAGAG
CTTTGGAGCAGTTATTTGGAGAGCATAGTGAAGGTAGTCGTGACTATGATGCTTGTCTGAGCACAATTGCCGCTCGACTCAGC
ACTGTATTTGCATCTCTCAAGGAATTTCCAATTATCAGGTATCGGGCTGCAAGGTCGGCAGCAGTAGATTCCACTGCTGCAAC
AACTTTTCGAGATCTTGTACCGACCAAGCTGGCAGCAGCTGTCTGGGACCGTCTTCAAGGATATAGAAAGTTGCCAGGTTTTC
CACAAGAAGAAAGTTGTGAACTTTTGATTGTTGATCGATCAATAGACCCGATTGCTCCTATTATACATGAGTGGACCTATGAT
GCAATGTGCCATGATTTGCTGAATATGGAAGGCAATAAGTATGTGTATGAGATCACAACCAGTGCTGGCAGGCCAGACAGAA
AAGAGGTTCTTCTTGAAGAGCATGATCCTGTTTGGGTTGAGTTACGCCATCTACATATTGCTGCAGTTAATAAAAAGCTTGAT
GAGAAGATGACAAATCTGAAGGAGAAGAACAAGGCTGCTCAAATACGACTTGGCTCCAGGGAAGGGCAGGAACTCACCACA
CGTGACATGCAAAAGATGGTACAGGCGCTACCTCAGTTTCGCGATCAGATTGAAAAGCTGAGTCTGCACATTGATATAGCAA
CAAAGTTGAATGCGAAGATTGGAGAGCTTGCCTTACGAGACATCGGCACTTTGGAGCAAGAGTTGGTGTTTGGAGATGCTAG
CAGCAAGGAGCTTATAAACATACTAACATCGAATCAGAATCTCACCAAGGAAAACAAGTTGCGCTTGCTTATGATATATGCT
GCTTCACACCCAGAAAAGCTTGATGCAGCAAAAAGGCTTCAATGGATGAAGCTCGCAAGGCTTACTGAGGAGGATATGAATC
CTGTGGTCAACATGGAGTATATGGGTGTGTCTGTTTCAAAGAAGCAGTCTGGTGGTTTCTCCCTTAAATTTGGTACTCGCAAA
AACAAGCGAGGTTTGAGGAAGGAGAAGAACCAAGATGAAGAGATTTATGATTTGTCTCGATTTTGTCCAATGATCCAGGAAA

TAATAGAGGATTTATCGAAGGGTGAGCTATCCAAAGAAGAATACCCTTATGTGACGGAACCACCAGCTGCAGTTTTGCACTCT
ATGTCAGGGGACAGTCTCTCCGTAAGGTCAACCACAACAAATTCTAAACCAGCACAGTCCATGAGAACCTCAAAGCCTGGTT
CTACTTGGGCCACCAGGTCCAAGCCTTCTGATGATGGACAGTCTAGTGATTCGGCGTTAAGGCATGCATACAGTGATCCCAAG
ATAACTGGAAAGCGCATTTTTATTTTTGTGGTTGGTGGCGCTACTCGTTCTGAGTTGCGCGCAGTGCACAGACTTACTGGGCA
GCTAAGGAGAGAGGTTGTACTGGGAACTACCAGTCTGGATGAACCTCATGAATTTCTAACGAAATTGAAAGTGCTTAGCGTT
CTGGACGAGGTTTCGCTCGACGATCTTGAGATATAATATACTATCAACAGTTCTATCTGATACTGGGTTCAGGAAAGCTGCGG
CAAGGTGTCTGCATCAGATGGTGTACAAAAGTTTTCTAAATCACAGGAATGCTCACTTTTGCGGTCCTCATCATTTACCAGTTA
TGCAATGGGCTCACACCAAGGGGAACTGTTACCTAAATGCACTGAACATTCTTAAAAAATCAGCTCGTTGTGGTTGCATGAGC
AATTCCCTGGAGACAATTCATAAGTTAGAAGCGTTCGTAACCTAGTTAGCGAGCATATTGGGGCTGTATTTTGTCTTCTGAAC
ACCTAGAATGGTTTGGGAAGTAAAATTGTATTTATTCTTGAGAGAGATGTGTATTGCCATTTCTGTATTACGTGGCTATTTTGA
ACTGGAGTGATGGTAAGGGCACAATTATTTTGTAAAGCTTAGGGATCATGAGTGCCCCATAGTAATCCTCTCTTGTATTTTTA
AATAAATCAAGTTACGTTTCTAACAAGGAATGGGATGGTGATTTTTCTGTTGTGCACCAAAGGCCAACTTGGTGTGTAGGAGA
TTAGAGATGGTGCACAATTTTGTCA

**DYNAMIN-LIKE 1B (*ADL1B*):**

CCCTTGTCTTGCAACTGCATAAAATTGGTGAGGGATCAGAGGATGCAGAGTTTATGCATCTGCCCAAGCGGAGATTTA
CATATTTTGGATAAGAAATTGTGGAATTCCTATTTTCATTCCCTTAATTGAGGTGGTTGCAGTTTGTAGCTCAAGTACTGGGAG
ATTAATAGAAGTTGCAGAGTATCGCTCAGGTGAAGGCGGGGGGTTAAATCAATAGCTTTAGCTGGTACAGGTGGTGGTTCAA
CAGAAACAGGAGGTCATCAGGTTCAATAAGTTTTGGTGGTTCTGGAAGCTGAGGATGTTGTAAATGCATTATTAGAATGGCTA
GAGCTTCGAATTGGGCGTGGAGGCAAGGAATATTTCACGCGCTTTAAACGCCACGAGAGACCCACTTTTTTTATCCACCACTG
TAACTTCAACCGAAAGCTCTTTAAGTGGTAACGAAGAGCGATCATACATACAAAGTGTTGACAGATTGCCAAGAAGCTTGAA
GCACTCTGGGAGCTTGAGGCAGTCTGGCAGCTTCAGGCAGTCTAGTAGAGATGGTTTTGTTACAAGGAGGAGGAGCGATGCA
GATATGTGGCAATGACACATTGATGATGGAGCTTTAGAGCATTTTACAGCCCGAAGGAGTGATG

**STOMATAL CYTOKINESIS-DEFECTIVE 1 (*SCD1*):**

CTGGTTCTCACTTGAGAGAGGAGAGAGAGAGTTACAGAAGTCGATACGGTGGCTGGATCAAGGCAAACAATACAGAT
GAACACGTCCATAGCTTGTGAGCAGGATCAAGGGATACATCCGATCTTACATTGATCTGTGGGTCGTCTTGCAAATCACAGCA
ATGGGTTGGGCAGCTCGGGCCATTGGGTACTTCGTAGTCTGTGGCTTGGGCCCACAAGTGAAGACCATGGAGGGCAAGTCTG
GCTTCCAAGGATTCGAAGATACTTACATGCCATCTCTTTTGGATCAGTTTCCTCCGACACCTTCCTCCGGCGTATCCGAATTGC
CTCCTCAGCTACCTCAGTGTGCATTGCCTACAGGGGTGGAGTTCTACTCATCTGGTCTGACCAGTGGAGATGAAACCAGCCAG
CCTCACAGCTATCCCATCATTTTGACAGACGGGGATGGCGCAAAAATTTATGTAAGCTGTGTTGCTTTTCGAGATCCAGTTGA
TGACGATGTCGCTCAAGCTTATCGAATTCCACCAAATTCTTATGTAGCTAAATGCATCTGTATAGTTTCACATTCACCTTATTT
TCATGTCTGCCGAGAGGCAGTAGAAGAGCTACATCGTCTCTGCTTCTCAAGTTCAGGCTGCAGCAAACCTTTGTGGGATATAA
TTTCGCATATAGTGTTAAATGTACCCCTGCCAAGTCCAGGTGGGGGCATTGTTCTTTTTGCAATGGAAAATCATTTGTTATCGA
TAGAAGGTCCACCAAAGGATGGTCTTCCACATGCAGACCTCTCATTTCAACCGTTGATCCAGTGCTTGGATGTAGATAACGTA
ATTCGTTTGTTCACGGCAGTTTTTCTCGAGAGGCACGTTTTGCTACGGGCTTGCAAGTACTCTCTTTTGACAGTTGTGGCTGAA
GCTATTTGTCATTTGCTGTATCCAATCAAGTGGCAGCATGTTTTCATTCCAGTCCTCTTCTGTGGAGGAGTGGATTACCTTGAC
GCTCCAACTCCTTATCTGATGGGCCTTCATTCTAGCATTGACATATCCTACCTTAAGCTTGATACGGTGGCTGTGGTTGATCTT
GATCGCAACACTATCACACTTTCAGAAGAGATGCCTTCTATCCCAGAGCCGGAGTTGGGACAACTTAGGGCAGATCTGGTGG
AATTTTTATATCCCAATCTTGCTGATATGGATCGAGTGCAAGACGGTGCTGGGGTTCTTAGAGATTCTCGCATTCTAAGAGGA
AACAAATCTTGGAGTGCAAGTCATGACTCTGAATTAAGGTTAATTTTTTTGAAATTCTATGCTACAATTCTATCTGGTTACCGT
GATTTTGTGGTGTATTGCAACGCTAGCAGCCAGGAGACCACAGAGGAAAACTCTTTCAACAAAGAGGCTTTTTTGAGGAAGC
GCTCTAGAATTACCCGCCTTCCTCCGGACCCTACGATGGATCAGTTTTTAAGCTCCATCGGTTTTCTTGAATTTATACAAAAAG
GCTACGGATCTTCAAGTACAGGGCAGAATCTTATGGATAAGTTGCAGCAGGCAATGGCTAGAGGTCAGAATCCTTCTGTTGTT
CTTCCTGCTCCAGCCGCCGAGCCCAACATACTGACCATTTCAGATCATTCAAGTCAAGAAATAGCATCAAGAGCGAAGTATCG
ATATGAAAGATTTCCAGCAAATGTCAGAACTCATGAGCAAGAGGAGGAAAGAAAGAGAATTCTAGAGGCTGCAAGTACAGC
TTTGGACCATCCAAAGCGACACAGCTCAAGCCCGGGGCTTTTGCAAACAGGGAAAATGGATGGCTTATCTCCACTTGAGAGG
GCTGCTGAAAGAGAACGAATGGTTCTTGACATCAAGGTCAAACTACAAGGGCTGTGGCAACGCCTCCTTGTTCTTGGCTCAGC
AGAAGACCCGCTTGCTTCTTTTGAATATGGCACAATTCTAGCACTGATCGAGTCAGATGCAGAAGGAATAGGTGGAAGCGGG
TTCATTGAGTGCATCGGTGAACACATCAATTCGGGTTGGAGTTGTGGTTTGACTGAAGAGCAGTTTATTGCTGTGAAAGAACT
TATTAAAACCACCATTAGCCGTGCGATATCTCGGAATGACATGGTGACTGTGCGTGCAGCACTTGAGTTTTCTGCTGTAGTTT
ACAGAAAAGATGTGGGCTCTGTCTCTGATTATGTCCAGCGCCACCTTGGTGTTCTTCCTATTTGGGATGAATTCAGGTTTTGGG
ATGGCTACTTTGATAAACTTTTGGATAGTTTTGCAGACAAGTCTGGAAATTTTGCTACTTTGGTTTCAGAGCAACTTATCATCG
TTGCACAGCACATGGCGGGCTTGGGTCTGCCAGATTCTGAAGCATGGTTTATTCTAGAGGCTATTGCAAGGAAGCATAATTTG
GGACCAAAGCAGATGATCAAGCTTAGGGGCTTGCTTGCTTACATGCAACAAGTTAGGGTTGGTTACTGGGGTTTTACTCCTAT
ACGATCCCAAACTGCCTATCCTGGTGGAACAAACCAAGCTCAATCTCATGCGCTAACAGATGAAGCACAACAACCAGTTGAA
GCTGCAGGCGCGACACGCGGCTGGGTACATTCCATGTTTAGCAGAGATAAAGCTGCATTTGCTGCTGGTAGACGTCGTGGTAC
ATCAGATAGCAGTGTGTTAGCTGCCAATGACAATATTCAGAACATACAACAACCGAAGAATGTGGACGGTTCAAATACTGCC
CAAAGAAGAGGCTCTTCTAGTGTTAGGACACTCAGAGGGCATAAGGCTGCAGTAACAGCATTGCATGCTGCAACAAAAGCAG
AATGTGGGGACAATGGTGTGGACAGTGATGAAGTCGGCCATTTTATCAGTGGTTCTGCTGATTGCACGGTTAAAGTTTGGAAT
CCATCTCAGCGAGGATCAGAGCTGCGAGCAACTTTGAAAGGACACACTATGGCTGTTCGTGCGATTGCGTCAGACAGAGTTC
GTATTGTTACTGGCTCCGATGACCAACGTGTGCTTGTTTGGGACAAGGCTACAGGAAAACAGCTTGCAGAGTTAAAAGCTCAC
AATGACAAGATTAGTTGTGTGCGAATGCTATGGGAGCCTTGTGTTTTAACAGCCTCACATGATGGTACGGCCAAGATGTGGGA
TGTACGAGCTGATACATGTGTCGCAACAGTCGGGCGAAGTGCAAGTGCTATTCTGTGCGCGGACTATGATTATGAAACAAGG
GTCTTAGCAGCAGCTGGAATGGATGGGGTCTGCAATATCTGGGATACACAAGCTGCAAAGCAAAGACACAAGTTTCTTGGAC
ACAGCAAATGGATCAGGTGTCTAAGAATGTCAGGAGACACCGTTGTCACAGGAAGTGATGATTGGACTGTCAGAGTATGGTC
TGTTTCCAATGGGGCTTGTGATTCAGTTTTGACATGTCATTCTGGGGCAATTACATCAGTTGAGTACTGTGCAGCGGATAAAG
GCTTAGTTACAGGCTCGGCTGATGGCATGGTGCGGATGTGGGAGAGGGATGAAGGGAAATTGAGATGCACTAAGAATGTCGG
AATCCATTCAGCATCAATTCTTTCTCTGCGGGCGGCTGACAAATGGCTGGCAATTGGGGCGGCAGATAACTCGATGTCCCTGT
TTCAGAAGGCAGAACAAAGAGGGGGGACAGCTGGTTGGCAACTATTCAGAACACCGCAGCGAATGGCAGCTGTGGTAAGAT
GTGTAGCATCGGATCCGGAGAAAGGAAGGATATGTAGTGGAGCTAGAAACGGTTTAATCCGACTCTGGGATGCTGCAATCAC
TGTTTAGTCTTGAGTGCAGACCTTGAGGTTGGATTCTTTTGTTAGGTCCAGATCGGTTGCACCTGCAAGTGAGAGAAAGAAGC
TTAGTACAGATCATTGAATGCAAGAGCATCCACATCCAATTGGAGCTGCTGTGGCCTTGCTAGGGATTCAATCTGTCTTATGT
ACATTAACTCCTTTGTGAAGCATTTTTGCTACCAAGGCTTTAAAGCAGCTCATGTTTTTATAGTTATTTTCAAACCTCTCTCTCT
CTCTCTCTTTTTCTTATCTCTCTTTCTCTCTTCTCTTGTCCATTTGTAATAAAGAAAGTTGGTATAAAAGAGAGCCTGTAGTATA
AGTTACTTACTCTCTACTCGCAGCACCATGAGTACTTCCAAAATGGACTGCGAAAATGGTAATGTTTCTGCTTTACTTGAATCT

TAAGGCAATTTTTAATTCTTATTCACATTTTTAACAGAAAATAGCTCTCGCGCCACTTCTCAAACTGTTTCTGAGGAGGAAGGA
GAGGAAACGCCCCTTGTTGCATCACCAAGTGACCAAAGGCATGTGCAAGAGAGTTTTCCCTCTCCTTCAAAATTTGATGGACA
ATTTGTTTTAATCTGCAATGAAAATGAAAGTGAGTGTACGAAGCGAAAAGTCAGTCGCGTCAGCTGTTTCCTTTGAAAGGAAA
TAAAGGTCTGTATATTTGTTAAGTCAGTACTCTGTGTGGTTTTAAATTGTTTCTCTAAACTATGTGCACATTTATTCTTTGTTTT
AAAGCTCGATGAGGAAAGTTCTAGCGGACAGAAATCTGTGATACCTTTTGCTACACTGACATCCTCATCTTTGCCTTTGCTCA
CTATCTCACAAATTTCAAATACCAATCTCCGTTTTGCTTTTGAGGGCAAAATTATTAATGTGTATGGGACTTTTCAAACTGCTA
AAGACTGTTCAAAGTATCTCAAAGCTGAACTTGCTGATAGAGAAGGTTGTAGCACTATCACTCTTGTCATAACTGAAGCGTTC
ATTCAAAAGTATCTTGAGAAAATAAAAGAAAATGAATTCATTAGGGTCGATAAAGGTACATGTGCCTTTGCTCTTAGTGTAGA
TGCGACTACCTTCATTGTCAAGGCTGAGCCGTTCCATGTATGCTTAATGTTTATGCCTGAGATAAAAATTCGCAACGTCCTTGC
CAAAGCAAAAAATGAGCCTTTCTTGAAAGGGACCATTGCTTTTGTTGTCATTCAAGTAGATAAAGTCAAAAAAAGTAAAGGT
ATTGAAACTTTTGAACAATTGACTGTGGCAGACGGCTCTACTTCAATGGATTTGGCAACGGTATTTTTCTAACTCAACCATTCA
AAAAATCTGAAGTGTTTTTTTAGCTCACATTTGTAAACAAGCGCAAGAATATCTATCGTGAGCTCGTAGAACAGTATCAGAAA
GGATCTATTGTTATGTATGTTGCCCACAATATTGGTGTATGGAAAGGCCAACCGAACACACTGTCTGTTTTTTATCTTATTTAA
CTCCTGTGATGGGTTCAAAGTGTAAGGCAGAGCTAGAGACAGCCTTTCGTGTCCAAATTTCTGAAATGGGCTATAGCAAACA
GGTGGTCTTAGTTCAAATGCGTTTCTTCTTTTCTCTTATTCTGTTTTTTTTTCTAACTTTTCAATTTATGCAAGTTTTTGGCTCTCT
GGAGATTACAATTTTAATGCCGTCTTCATTGTGACAAAGTGTGCTGTCTGCAAACAGCCGAACATCAACAAGCTTCGCGAAGG
AGGTTTCTGATTGACTGCGGCAAGCCGGTTGAGCTTTACGAAGTGCCTTTTGAAGCAAGGATTATCGCTCCTTCCGGACAAGC
AACCGAAGTTACATTGGTGGGTGACGATGTCGACAAAGTGCTTGGGCTTGAAGATTCACTTGTTAACCTTTACCATAAGAATG
GTATTCTCACCCGTGTCTTGCTGTCTGAAATTTCAAAAGTGGGAACCTTTCTGATCGATAACCAGGGTTCTGTTATTGGCTTCA
ATGCTCCTCCTCCATTACCATAACTATCAACCATGTATATTTTGTGTCATGTCTCCTTAGAAGCCTTGTAGTTATGAAGATGTTT
GTCGTACACTATAACCCTTGTATCTATGAAGTCTGTTGTATACTATAATCCTTGTACCTATGAAGATTGCTCTACACTATAATC
CTTGGTCTTTTTTGTAACTTTCCCAATGGACATTTTGCAAATATATGATGTTTTGTCCAAAAAAA

***A. THALIANA* HOMOLOGUE OF YEAST SPO22 (*ATSPO22*):**

TTTTTCGTTAAGCAGGTCACAGTCGCAAATTGCTGATCTTCCAGGTAATCTGCTCATCTATTTCCATCAAAGCAGGAAT
TTTTCTGCAGTAGATCTCGTCATTTTCCTTTTGTAACCGAGCACTAGGAGCTTCCTGCAGAAGAAAGTGCATTCTATTGAGCTT
CTGATCAAACAGCCTCAGGAGCCACGTTTACTTGAATCCATCGTAGCAGCTATTCCAATCTGAGGAACGATGCAAATTTCTGA
GGTTTCTAGTGAACAATCTGCAGCCAGTGATATAAGCAATGATCAATCCTGCTTGCCGTTACAAAATTTGGAGAGTCTGGTTG
TGGAGGTGGAGCAAATGGCTCCTATGATTAGTGCGGAAACTGTGCAGAAGCTTAGAAACCTCATTCCTACCTTCCCCTCTCGA
TTGAATGAGGACCAGAAATTTCGGCTATGGAAACTTACCTACCGCATCTGGAACACCTGCATTGAGATCAACAACAGCCTGC
AACCAGGCCAAGATGTTGACGAAGAGCACGCTAAGCTTCGCCATGTTGCCAGTGATCTTCTTGTCATTGCAGGTAGTGTCAGT
TCCGTACATACAAGCCTTTCAAAGACGGCTATGTTTTTCTTCAAAACAGGAACAATCTGGCACAAGCTTGAGAACCATGCCAT
GGCTGCTGCTTGCTTTGAGAAAGCTACTGAATTTAGTAACAGAGCCAAAGATCATAAGACAGCTGCTATGTCGCAGGCACAA
ACAGAAGAGGAAGAAAAGTTCTTGTTCGATTTGTTTGTGGCACGGGCAAAAACAGCATGGGAGCTTCAGCAGAAAGCTTTGG
TTCGAAGCTTGTTGAGCAGCGCAAGGGGGATGTTAAAGCTGGCATCTTCTTCTTCATCTTCAACGCTGATGTACGAGGAGCTT
GCAGAAGAGTACCTGCACTATGGTAAGGTGCTTCTTGCGAGGCAGGATAAGCCATCACAAGCAGAGTCTATCCAGTTTTTAG
AGCTGGCCTTTGAGGTTTGCTCAGATGGAGCTTCGGCTGTTGCTGCTGCTAGCGGGGATGGCGGTGATGATGAAGACATGCTT
GGGAAAACAGACAGGGAAACCCTTGCCTTGAGCAATCTGAAGTTCAAGGTTTTACGATACCTTGCTGCTGGCCATCTGCAGA
ATGACAATTTTGAAAGCGTTCTGAAATGTGTTGATATTCTGAAGGCAGGGCCAGACCATGCAAGCACACCATTCTTAGCCTTA
AAGGCCTACGTGGGGCTGAATCGTTTTGAAGGGGCAGAGAAGGAGGCCTACATTCTGGTAGCACAGAGATCTACGCCAGTCG
ATGTATGTGCGGCTGCAATAGATGTGCTGATTGAGGGCTCATGTAGCTCGGCAACTCAGATTAGTATCAAGATTCTGCAAGAT
GCTGTGAAGAAGGCCTTTGCCATCGCACATGCCAGGTTTCCATCCAACAAAGAGCTCACTGTAAGGGTGTTGGATAAGCTTTT
GACTCAGCCCCGTGACCAGTTCTCCAAAAAGAGGGTTGAAACTGCCCTTGCCATTGCTGTGGACGAAAAAGTCATCAACAGC
ATAGTTGAAATTTCAAGCAGAAGAAAATCTGCAGCAAACATTATCAACGAGTTGGAGGACGAGTCATCCACTGGAACTGCTC
AGCAATGCATCTACACCTTGCTATGGAACAGGTGCAGAGTAATTCTCACTTCCTGTTTCTATCTTTCTTCTCAAATTTATATCC
ATTCAAATCAATACTGGCTGCAATCAAGATCATTCTGGACTATGAAACAACCTGGCCTCGCTTTGCATGAAGTTTTTCTGCAA
TTCTCTGCAGTCTCACCACAGAAACCAAATTTAGTTGCTGTGTTTTTTTTTTGAGGCATTTCATGTATCTCCTCCGTCGTCGTGG
TCTTAATTACAAGGCCAGAGTTCAAGGAGGTCCGATTTTCTTTTGCAAAATAATGCAGTCTGGCCTTTGCCTTCGTATGTTAGA
ACTTGGATTGTGAGATGCATGGCTAGGGCTGTAGTTTTTGATAGCTTGACCGGCGTGTTGCGGTAGCCTTCAGGGCCCTGATG
TCCACGAGTTACTCTATTCTTTGAACAGCTTGAGAGATAACTGATTTCTAAATTTCAAAACTGCAAGTCTCTAGTACTTGAGCT
GGATTTTCAGCATCAACGTAGTTGCATTCTATCAACATTACACTCACGGATCTGTCAGAATCTGCTGCAGTGCACAAGAATTT
TTTGTAGCCAAGGAGTATGCAACTTGCATTCAGTTGTTTGAAGCATCTCTTCTGTACCTCTCTCCAAAAGACAACAGTTACACC
CAGCACAGAGCCAAGAATCTTCGTATCCTCTGCATTTGCCATCTTGCCCTTGACCAGTATGATCGAGCCTCAGAGTATATTAA
TGAGGCAAACAAGCTGGAACCTAGCATTGCTGGAGTATTTTTAAAGGTTGAATGTGCTCATCTTCTACATCAATTTCAATCAT
TCGAGTGTCTTTCTTGAAGCTGCCAAAATTGTGCTGTTCCTTGGAATTTTGACCTTGGGCCTGCTTGAGGCAATGGAGAGCTAT
CTTGAATGTTGGTAATTAGGATAAGCATTGAGCAAAGTCATTTTAAACGGACCATTTCTTGAATTTTGTTTTCCAGTTCAAAGT
GGCTCTTCAGACAAATGATGAAGAAGGGGCAGCCAAGCAGGTTGAAGAAATGTTTCGGTGCCCAGATTTTGAGCCAGATTAT
CTGACGTTGGCCTCCCATGAGGCCATATCGTGCAAATGTATTGGCGTGGCCATTGCTGCACTGTCACGCTTGCTGGAGCTGTC
ATGTTCCTCTGCAAGCTCTAATACTGGAGACAATAAAGAGGCAGTATTAATCAGAAACATTATTGCACTCTCTTCTCAAGACT
CCAGCAATTCAAGCACTTCTCTTAAATATTTCAAGCTTGCTCGATCTCGTCTGGCTGAGCTGGGACCTGAGGGTTTCTTGGGGT
GTGGGAGTATGGAGGAAAAAGAAGCTCAGTGGTATGCAGGCTCTGCATGGAACCAGGGCTTAGATACAGTCAAAACTCAAG
AATGGGGCTTGTGTAGAGAATTCTTTCAGTGTGCTGCTGATTTCTACGAAGTTCTCCCTGACACCCTTGAGAACCTTGAGATG
AGGAGAGCATCTATGCTGCTCGCTGTAGGTTCAATCCTTGCAGGAAACAGAAAGGATGCAGGTCTGAAAGAGGCTTCCATAT
ATCTTGAAAAAAGCCGTAAGGTAAAGTCTTTGAGTACACTTGTCAACTTGACATGTAAACTACTGTCCATCATTCTTTGAAGA
AAATTTGATGGATACAAACCAATAACCTCTAAACCAGGACCTAAATGATCTCTATTAGGAAGGTTAAGCCATTAAAAGAGTC
TAAAATTTGGTTACTGAATCTGTTTTGCGCCTCAGATACATGAGATTGTAAGGTCGAAGAGCATTTCAGGTGTTAACAATGAT
ACCACTGAGATTTATCTGAATCTACTCTCCTTCGAAGTGAAAAGGAGGATGAAGGATCATAAAGCTCAGCTGGACATCATTTG
CCATTGCAGCACACTGCCAGGTTTTCAGCCAAAGCTCTTTTTCTCAATGGGATCAAGCGGAGGAGAAGGCCAGGATGTTGAG
GTCTCCATGGCTGCATTTGAGGCTTGCCTCAACTCAGCCTTGGCTTGTCCAGATCCTGATTATGCTTTGGTTGCAGCTACCATA
AGAAAAATGATAACACTTGCAGACACATTGAGACGAGACAGCCCTAAAGCAATGAGCTTGTATAAGCAGGCACACCAGATA
CTGCTTGGCCTGGGCATCGGAGTGTACCCAAAAGAAGAGGTTCAGTGGTTGGTTTCGACTGCTTGGAACAGAGCATCTCTCCA
TGTCAAATTTAATAGATACTCAATTGCAGAAAAGTGGATGAACTTGGCCCTGGACATGCTTCAGCATGCACCCTCAATGGAAA
ATGTTAGGTCTGTCATGCTGGAGAACCTTAGTTCCGTGCTGAAGGGAAAGGCTGATGAGATGGATGAATGAGCTGGACAGAA
TTAATGCATACGAGTTGAAAAGGATTCCCACAGTTTCTATTTGCGGGTCTCTTTCTAAATTTCTGCAATATCTCTTAAGATCAA
AAGGATTCTACAGCTTCTAAGTTCTAATAGTACTGCCTTATAGATATTCTTATGGTGAAGGAAATGAAATATGGGCCACTCTT

TGTCTTTTGGTGTCTCTAGCTGATGCCTGCATTGTCTTATCAGATGACAGCTTTTGCTTGAAATGGAAAATAATGTTTGGTCGA
AGTTGATTGGACAATTATACCTTCTGTGAAAGAGAAGCTTTGGACCACTGTATTTCTCAGGCTTATGGAAGCATTTTTTTAATA
CCCACTTTTCATTTTTTGATACGCTCAAATTCGTCAACGATAGGGCTCAAAT

**ACTIN DEPOLYMERIZING FACTOR 11 (*ADF11*):**

GAGAGAGAGAGAGAGAGAGAGAGAAGTCGGACAGAGAGAGAGAGAGAGAGCTTCCTGAAGTCTGTGCACTACGTCGCG
TTGCCATGGCTAACGCTGCTTCTGGGATTGCTGTTGATGATGAGTGCAAGCTCAAGTTTTTGGAGCTTAAAAGCAAGAGAACA
CACCGCTACATTGTGTACAAGATTGATGACACATTGAATAAGATCGTGGTTGAGAAGCTTGGGGGACCTGAGGAAAGCTACG
AGGCCTTTACTGGCTCTCTTCCTGAGGGAGATTGCAGATATGCCATCTACGACTATGACTTTGTTACAGAGGACAACTGCCAG
AAGAGCAAGATCTTTTTTGTAGCGTGGTCTCCTGATGTGGCTCGTGTGAGGACTAAGATGCTGTATGCTAGCTCCAAGGATCG
TTTCAAGCGGGAGTTGGATGTGACCCATGAAATTCAGGCTACGGATTCCACTGAAATTGACATAACAAATATTCAAGAAAAG
GTTAACTAAGGAGCTCTGCAAGTTACACAAGAATTTATAATCCCTATGCAGTGTTCTGGTGTGTTTGTGTATAGTTGTATTTTG
CTCGTTGCATCAACTCTTTGCATTAGGTTACTTAATTATGTATTGGGCCATATACTCGGGCTTTCACCTTACATTATTGCCAAGT
TTGGACCTCACCAGGGTGATATTTCTTTTCAAGGAGTGGTTTTGATCTTCTACCATTTTGCTGCTCTTGTTCTCTCTTTTGCAAA
TTTCACGGCCTAATGAGGTGACTAGGCTCTACCCTACAGTTGGTTCAAGGTTGTGCCGTTTATCGGTGATCAGGCTGTACGCG
ATTGGCGACTGAATTATTAATGTATTTAATACTACTCTCTGTCTCTCTCCCCAATGCATTTTGGTCTTTGTGCTGAGGGGAAAA
GTGTTCATCATAGTGGCAACAAATGCCAATGTGTGTAGTTACTGTCTTTGAGGTGGGCTTTCAAAGCGGGGAGAGAAGTGTAT
TTTGTCTTGATTGTGGTAGCTTGGCTTTCAAGAGATGCAAAGCCTGTAAGATGTGTCTAATGTGGTTT

**ABERRANT LATERAL ROOT FORMATION 4 (*ALF4*):**

AGAGTGAAAACAAAAGAACAAGATGGACAACAAACTCTTAAATTCTAGTCTGTGATTACTTTTGGGGCATGTCAAGA
TCGCTAACATTCAACTTATTCAACTGTAAGACTGATGATGAGGATTTGGATGTCTTATTTTTTTCGAGATCATTCTATTAAAAA
ACACCTTAAGGGCAAGGTCCAAAGTGGATGTCTTCTTCACATGGACTTGAAGGAACATTGGCAACGAGGACGCTAAGAATGA
GGGCGTAGGGCATGAGCAAATCTTGGGAAGAGGGTCATGCACGTAACTTTTTTCCAACACTTGGGGGAGCCGGCTGCATAAG
GTAAAATACGTGCCTTACCAGCTGCTGCCAGTTCAGGAGATTACAGGTCCCAAGCCCCCCCTCCACATAGGCAAGCGAACTTG
ATGGCCCTGAATTCTAGATAATTATGGCCCACACCCTCTCCAGCAGCTATAGATGCTGCTGTGAAGGACCTATGCGGAGGGGC
TCTAGACATGCAAGCTCACCAACCTTGACGTATGACGTGATGTTTACAACATGGCTGGGGCATAACAGTTTAGAAAAAGCTTG
AAGTGTTGCCTGACTAGCATTCCTCTGTTTTGGCACATCGTACTCCATTTCAGTGCATCATTTGCCGAGAGGTTAGCACTTGTC
GTCAGGTCATGTGATCCATGCCCTTCACCATACCGAGTAATATTCCAAGTTTATGCAGACTGGCAATGACTAGCTGATGTTGG
TAGCCTAACATACAGTAAGCTCAAGGAGCTGGTATTAGCCAGAATCTTCTTTCAGTGCTACCTCCGCCAACTCAAGGCATCGG
TATAGCACCCCTTCCAGATTTGATATAGATAGAAGAACATTGGCAGTAACTTCAGAATCATCAGCAGAGCAATCCCTTTTAAA
GCTTTCAAGAAAAGAACGAAGAGGCTGGAACCATTTTGAACAAGCTTCTTGAAGTCTTGCTTTGAACAAGACCCCTGTATAGT
TGGACTTACCTGTACAGAGCAATGACTTACAACTTTATTTTCATGGTTTTTGCAAGTTGCGAAAAAAACAGTGAACAGTAACC
TGTTGTTTCCTTGATCAGAAGAAACCGATACAGGTTTAACACTGACAGGACAGCATCTATCTGATTTGGGAGGTCAGGAGGGC
CCCCAACAGGTGGATGCAGAACAAAATCAACAAGCTCCAGCACTTGCTCACTTGCGAAAGGTGAATTTAGAAGTGAACTAGT
TCCTTGGACTGTGCTTTTAGATGCTAATGCCTTGTCCGCTTCCTCTTTGACAACATAAATAAGAAGAGATACCATTGAAGAAT
GCTCGCACTCGATGATGAGTTGCTTCAAAGCTTTAAATCTTTCGATTGGAGGAAGTACTTTTCTGACAATAATCACCAAGACA
CTATATGCACGTTGCCTTGTCCTTGACTGTGGAGAAAAAAATCACCACATTTTGAACCACCTGCAGAAGTTCTTTCAGCTTGAG
AAACTCGTTGGGCTCCAGAAACTCTAGTTGGATGTGAAGAGGGTTATCTACAGAAGGCGTTAAGCCCCACTCCAGAACGATG
GTCAACAAATCAATGCCAATCTCCACTTTTTTCCATGTTTTTTCTTTTGAGGAGAGGAGGACCAAGACATTAGACAATGCACTT
ACAAATGCATGCCTTGAGCTCCGCACTCCATCAACTACAGATATTACCTCTTGATTATCTATTGCTGTTTCACCAAAACAGAAC
CTCCACCATACTGCAAGAGCTGCCCCTTGTCTGGTGTCTAAAATGTTACCGCCCTTGTTGACAAGATCTTCACTCATGGAATCT
ATGAAACTTTCAAAAGGGCTTCCTGCAATCATATCTAAATAAGACAAGCCACAGGTGGGCATTAGCTTAGCAAGTTCCACCA
GTACAGATTGTACAGCATCTGAATCATTCAACTGCTCAAATTCAGAGAATCTGGCCAGAAGTAAAAGAATAAATACACCCAC
TGCTTCATGTAGACTTTGTCTTTGTTCTTTTACCTCTTCTTGTGCACCAATGTCAAGTAAAACCATAATTGTTTGAACCAATTTA
TCAAGGATCATCCATGGCTCCCAATTGGTATCAATTGCAAGGCCAGCTATTCTCAAGAGGCCTGGCAAAGACTCCTTAAAGTA
TTCAACCCTTCGTCTTCTTAGGCGAGTGAAAATTTTGGGTAGGCCATCTAGGATGACTGGGCAAAGTGCCAGAGTTTCTGGCT
CAGAATACAAATGCAATGCCTGAAAACATGTGAAGCACATTTTCAGAGCAAACAAAAACAGCAAGTGGAACAAAAATGCAA
AGCTACCTCCATGTAGGCTGCAAACATGTCGCGTGGGTCACAGCTTTCCAACAAAGTTTTCAGGATCAACTCTGCAATCCCTC
CACATGATCCATGTGGAATGATGGCAAACTTGACAAGAATAGATGGCATATCGATTCCAACGATCTCCAGTAAATCCTGACC
AGTGGGACTGGATGGGTCGAGATGAGCCTTGACCTCCGCTAACAAGGTCTCCATATCATCTGACGGCTTGCCACCTCGTTCAT
CTAATCTCTGCGCAGCAACTGACACCCTACGTACGAGTTCCTTGCCCGCTGCTTCCAATTCCATCTCCACCAATTTTTCTTCAA
AACATGGCATTATATTTTGCCATTTTC

**TORNADO 1 (*TRN1*):**

AGGAACTCCCTTAAAATACAATGCTGCAATTTTTGCTCAACTTGCAAGGAATTTGAGAAATGCAGAGCTAGCAGAGTT
GCTCCAACCTGAACCCAAATATAATATCAAACCCAAACCCCCTCCTAAATGTAGCAAATCCAAACCCATTCTTAAACTCAACC
TAAATGTTTTTGGAAGGTTGGACTCAGAGGAAGAAGTTAAACTCAAACCCCCTCCTAAACGTAGCAAACCCAAACCCATTCTT
AACCGTGCCGATGACTTTGACATGGTTAAACCAACAGCAGTTAGACTTTTTCTTTGTGGCCTTCCTTATTCAGGAAAAACCAC
CCTTTCTCAGTCTTTTGTGGCATATAAATCGCAAATGGTTGGTACTTCAGTTCGAAGGTTTCACAGGGCTCATATAAACCGGTT
TAGGGTTAAAGGCCGCGTGCCAGAACGGACACGTGGCATAGAAATAACCACTCTGGAGGAGGATGATTTGAAGGTTTCGCTT
TGGGATTTGGCCGGTCAGTCAGAGTACTATACCTTTCATGACTACATGTTCCCCAATCTTAGCACTCGGGCAACACCAGCTTTT
TTCTTATTTGTTTGGAATCCCATCGAGGTTGATGAGCGAGGTCGTCATAAGACAGATGATAATGGAGCTCTTGTTGAAAAATC
TATTGAGAAGTTTGAGAAGGAGTTCAAGTACTGGCTTCGATTCATTGCATCCAAAACTCTAAGGTCAGAAAAGTTCAAGCCTA
GGTTGATAATTGTCATAACAAGAAAAGATTGTAATTTGAGCCACATGAAAGGATGGACTCATAGCACCCTTCATAATCTTCAG
GATACTTTTGGAGACATCATCCAATTTGACATGAATGGACTTTTTGAAATAGATGCCAGGGATCCTATGGAGCTAAAGGGATT
AGCTAAATATTTCTTTGATTGTGCAAGAGATATTCTGGATTCAGCACCAAAGGTGCATGCGGCATGCAAGGACGCAAGGTCA
ATGATTGCAACACTAGTGGACACTAGAAGCATCGCTCCTTTGATTACAAGAGATAAATTTTTGTCTCTGTGTGAAAAGCATCT
TTGTATCAACTCAGAAAAGGTGCAAGAAGCAATCGCTCTCTCTCTCAATGATTCTGGAGATGTAATTTATTTCGCATCACTTCA
GTTTGTGGTGGTAGATACACAATGGTTTTGTCGTCAAGTCATGGGTGACCTGATGTCCAAGAGCATTCAGACTCAATCCAATG
GCAACAGAAATCCTCAAGGTATCTTTTCGGTAAAGTATCTAGAGGATTTGTTGGAGGCACAAGTGAACTCAGCAGCATGTAG
AAAGTCTTCTCGAATCAGGCACGCATTCAAGGCCTATAGAAGAAAACTGGTGACACATGCAAAGGTGGTGGGAGAAGACTTA
GTAAAGCTGCTAGTGGAGTTGCAACTAGCTTGTCGTCTGAATGAAGATGAAGGGAACTTGAATGTTTTCATACCTGCAAGTCT
CTCTCAAAAGAATTATGCTTATGCACCACACTTTTCCTTAAAGCGAGAAGGCATCAAGGCATTCCTTGGAAGACGGCTAAAAT
GTAAAGACTCAAAACGCACTTTCCTCACACCCGGTGTCTTCCCGATAATACAGGTTGTTTTTATAGCACCATCAAAACAGCC
AACAAAAGAGAATCTGCTGTTACACTTGAAGAGGATCTCATTTCATATGCTCAAGAAGGTAATGGTGTAATAATAGAACTTTT
TAAGCAGCAGGGAGAAGAACATTTCATAGATATTTTAGTCTGCTCTCATCTTGAGCTAGCTCATACAATCAAGTGGGTTCATA

AGAACGTAATCACAATCATAGAAAGAGTGTGCGCAGAACCGAAGGGAATCCAAGGTGTGGAGTTAAAACATGAAGTGATTC
GACCAGAGTGCGTGGAATACATTGACCGAATAGAGAAAAGAACACAGTCGGTTTCTGTGACAGATTTGAGAGAGAGGTTGG
AGAAACATTTGCGAGAAGAAGGATTTGGGGCAGATGGTTTCAATAATGTGCTTCACACTTGGATACCTGGAGGAAATTCACA
TCGTGTTGATGAATTGTTGGGAGAGGAGGCGACCCTACAGATTGTTGAGACTTATTGGCGAGGTTTAAGTGTGGCTGCTGCTG
ATGCATTAGGTGTTGTGCCGAAGGACATAGGTACTAGTAAAATAGAGAATAAGATGGATACTAGCAATCTACAGGAGCAGCA
GGCAGGCAAAGATAATGTTGGAAAGGGGCACCTTTACGATGGCGAGGCAATTTACAATAAACTGGATGAAGTACTTCTGGAG
GTGAAGAAGCTGCGCAAGGAGCAGCAGGCAGGCAATGATGAAGTACTTCTGGAGATGAAGAAGCTGCGCAAGGAAATGCAA
AATCATGGAAATAAAATTGACAAGTTGCGAAAGGCTGTGATTCCAAAGCTGGAGGATCTGTTTACATTTTTCAAGGAGGGGG
AGGCAAAGCTGCCAAGAATGCTTGTGGTAAGCAAAGGGCAAAGTGGGTTGGTGAGCATGGTGGCAACTGCAATGGTACGTA
AGCTTAGTAGCTGGCGTCTAGAGCTTTTGTGCGAGTGTGAGAGTGGACCCCACTCTGTAAAAGGGTACCCTGGAATTGACATT
GCAACCCTAGAGGATGGACTTCTCAAGCGTGGCCTTCCTTACATAAACACATTCTTGAAAGTGGCCAATGTGGCTCTCAAAGT
GGGCGCTCATGTGACTCTCGGTCTTGGGAACGTGGTACCAGATTTTCGACATGTGCTTGCATTGATAGAAGACACTAATGAAT
TTCCTGGCTTGCCAACAATTTTCGACCCTGCTGATCCTTTGGGTACCACCTCTTTCGCATCCAGAGAAAATTTTCAAAAAGGCC
AACAATGGCTTCTTCAAGTCCTTTCCAATCGAAATTATCTCTCTGATATCCAGATAAAGGAAGTCTTTGGACTTACTCGTATTC
GGTACCCTAGCCACAAAATTGCATGGGTGTGCAAGGAACATTCCCATAAAGGTCAACTCTGCCCAGCATGGCTTTGATTTGTC
AAGTTGTGGTCCATTTTCTTATCTGCGAAGTCTAGTAACAAAAACTCACTGGGGGCAATAACCTCTGTTTGTAAAAGTTCAGT
CACTGGAGGAAATTTGGCTCTTCTGTATTCCAAGCGTGCGAAATTATGCATGATCAATATTTGTAATTTAATAAGTTGTATTTG
AATGATCAAACACAACAGTGTACCCAAATGGAAGTAATTTGGCTCCTGTATCCCCAGCGTGCGAATTTATATATG

**HOMOLOGUE OF HISTONE CHAPERONE HIRA (*HIRA*):**

ACCATCATACTTGGCATAGCCCTACTTCCAGCTTTAGTGTATACCTGTAAAGCTCCATCCTGGCAACCTGCAGCAGTTA
CAAGAAATACCACTGGCCTTTCCCGTCAATTTATCTCGCCACTTGACATCCCCCACCCTGAGATCAGAATCAGTCTTTTGGGG
ATTCATGCTGGCTGATCCACCACTGAGACTCGAGAGGCCTTGCTTCAAAGCATATTGGCAATCCCTGCTGATCGGCCTCTTGC
TTGCTGGTGGAAACAATTATGGAAAGGACACCTTTCCAAATTACCAGCAGCATCAGTCGAGCAACAGAGTTGGTTGCTGTGA
CAACCTGATGATTGCCAGCTGTGTGGCCACCACCTTGGG

**GROWTH REGULATING FACTOR 2 (*GRF2*):**

CCCACGATGCATGTGCCTCTCACAATACTTCTGATCGGGCACTACTGCCCTCCCGCACCTCCACTTCTTACCATCCGTC
CTCCGGCATCTCCCTGGCTGTGGATCAGCCAAACCCAAAAAATAACTACCTGCATTGCTGCCAAAATTAACGGGCACATTCTC
CACTTTACAGCCCATGGGTAGCAGCAATGCTGATGGTGGTGTGTTGCATGCAGCCACATGCTTGAGTACGTACGCCTGCTGTC
TTAACTCTGCCCATTGATCTGGTGTTAAAAAAGCCTGTGTTGGCCCTAAGTAAGGCAAGCTCCCGTGATGAGATCTTGCAGCA
GATGCAGCTTGGGCAGCCGGAATTCTGCAATCTTCAAGCGCAGTAGAAGCATAAGATCTGTTCTGAGAAAGCAAAGCGGAGC
TATGGAGACCATCGGCACCACGGATCGCCCCCACCATGTTATGTGGGTACTGCAGAGAGCTTGCACCTAGGGGGTTTGCTATG
GCAGATGCGTGATACTGATGATGACGAGGAGGCTCCATGCGCGCAAGCTTGAATGGCCTTGTATCTTCTGTAGTGTCGTGAG

**RETICULATA RELATED 3 (*RER3*):**

GTGGGGTTGGAGGTGGGCGCGATGATAATCATGCAGATAATGAAGGCCAAGGTAATTCTGATGATGAAGGAGGCCTC
TTTGGTGCCCTGTTAAAGGGCTGGAATGAAAGGGTAAATGCAGATCCACAGTTTCCGTTCAAGGTTTTGATGGAACAAATTGT
GGGTGTGGGTGCGTCTGTCATCGGAGACATGGCTTGCCGTCCCAATTTTGGCTTGAATGAGCTCGATTTTGTGTTCTCCACCCT
TGTTGTTGGGTCCATCTTGAATTTTTCCCTCATGTACATGCTCGCCCCTACGTCTCTGGCTGCATCTACAGCAAGCACACTGCC
TTTCATCTTCTCCACATGCCCATCGGGGCACATGTTTGAACAAGGGGCATACTCTGTTCTTGACAGGTTTGGTACATTTGTGTA
TAAAGGAGCTGTTTTTGCTGCTGTTGGCTTTGGGGCAGGTCTGTTCGGGACATTCCTATCAAATTGTCTCATTGGTCTCAGAAA
GAAGATGGATCCGAGCTTTGAGCAGCAGAATAAGGCGCCTCCAACTCTCTTGAATGCGTCTACATGGGCTATCCACATGGGG
CTGAGTAGTAACCTGAGATATCAAGTTATCAATGGAATGGAATTTGCAATGGAAAACGTCTTATCCCCACCTGTATTTAAGGG
GTCGGTTCTCTGTTTACGTGGATTCAACAATGTGTTGGGAGGCTATTCTTTTGTGACATTGGCAAGGTTGACGGGATCTCAAA
AGAAAGCAGAGGCAAAAGAAGTTCCTTCTCTTGCTTCTAAAGCACCTACAAATGAATATGAGGAGCCCAAATCGTCTAATAC
AGATCCCTTATCTATTGAAAAAGTAGTAACCACAGAAGGGAATAGCACTAACATTGGAGAGGGTAGCGCTAAAGTGGGGGAT
GCGACCTCACAAAAAGGGAATAGCACTAACATTGGAGAGGGTAGCGCTAAAGTGGGGGATGCTACCTCTTAGCGGGGCCCCC
AGTTTTGAATTCGCCAATGCAGTATAAAAAAAAAAAAATTCCCTCTGTGCAAGCAGATCCCTAACCTTTTGCAGCAATCAGTTA
GCTGTTTTGTGCTTCATGCTTGGTCTTTTTGTAGGTGTAGCAGAATGTTGATCAATGTAGTTTCTCTTTTCATTTTCTGTTGTGA
GCAACCATGATACACACGTTGAGAGTTTGAGCTGAGAGCTTGAGTAATCTGCTTCTAAGTATCCCTTTGTAAAAGTTTGATGT
CTATTAATCATTAATCAGAGGAGTAATGAAAAGTCTCACATTTCTAGCTGTTGACCGTGTCTTTGCTACAATTGTAGTGGATTT
CTTATTGGCGTCGAAAGATCGAGTCAAGACTTGAGCCACAGAAGAAGGCTGAGTTGCTACTGCCCGTGCAAGAAGGCGTCTC
TGTGTTTACCTTGGCCATCCACGTGGGGCTCGGAAGCAATTGTATTTATCTTGTAATGCAATGCAAGTGACTGAGTTGTTGCTT
CTCCTGTTTGCTGGCATGGGTTGGGAAGCAAAGGTGAGGAGTCTAACATGACAATCTAGCAACAGACAGGAGACATTAGCTT
TAAATTCAGCCGTTTAACTCATCGAAAAGAGGATCTTTTGAAGGAAGAGCCCATCAGTTTTCATTTTTTAACAATACATCTGTT
TTGAGACCAGGTTCCTACCCCCAGAAGCAAAACTCATCGATCTGCTCTTTAGGTCTTCTTCTCAACACTCTAACCGATTCTTTC
TTACGAGTTAGCAGTCCAAATTGGCTTGGAATCCTGCATACCAAGTATCATTGGTCAAACCTTCGAGTCAAAGTGATTACCTA
CAATACGGACCGGGTCCAGGAAACATTCTTTATATGCATGCATTTGCATAGTTCTTGTGTATTGCTTGCAAGCAGTTGGCAAA
GGTGCCACATTGAAGCTAGCTATCCTTCAGGTTTGTTTAATTCTGTTGTTGCTACTGTCAATAACATGTTCCTTGGTTGTCTTGG
ATTGGGTGTTGTATTAAGGTATTTTATATGAGCAGGAGGATTTGCAGCTTTG

**RETICULATA RELATED 4 (*RER4*):**

TACCCAAGCCAAGTTGCTCGTGTTCAGTAAGTATAGATTTCTTGCAAATAGAACATAAACACAATCTTGAATAGAAAG
ACACTAGATACATCTTTTGGCATGATCAATTTATTCATAAATGAGATCGTAGTCCGATGAAAGACATGTCTAGGCAGAAATCC
TCAGCCAAATGCATTGAAGGCACATCCGGAAAGCTTAAAAGGATAAAAAGCAAACACATTGCAAAACAATGCTTCATCGATC
CACCATGATGCGGGGGGGGGAACGTACTGCCTAAGGTCTCATGTATGCTGAAAGCATAAGGTCTCATGTATGATGAAAGCA
GACTGCAAGATATGCTGTACATGAAACAGTCTCATAAAGATAGACCCAGTCTGAGCAGACCCAATCTGGCCAAAGAACTGAG
GACTCAATTTGAGTGACACCAGTTACAATGCTACAGCACTAAATTGATAAATGGTGTAAACATCCCTGCCGCTAACATTTTGT
ATGAGTCTGTAATAAATGAAGCTGAAAACTACTTGCTAGAAGAGTTATTCCATCCATGAGAACAAATTGTTTGTGGCTGGCTA
TCCATGATTTGTATGTAGTAATGGTTGAATCTACAAAAAGCTTTCTCTGAACTAAAAAAGTGTAAGCTAAATGGATGCTTCTT
GATAGAGTGGGCATTGATATTTGTCAAACAAATACTAAGCTGTGGTCTCAGAATCTTCTTTGATCTTCTGAACACCCACCCAC
CGAGCATAGTCCACCCACATCAATGACCCCAAGAAGGTATTGCCAGTTCGAACAATAAAGCATAGAGCACTCAGCGCTAGCC
TGTGATTCTCTAGCATAGGTTGAAGAAACCGCTGCTCCACTACACCAGCCAAGAATTGATACCTCAAATTGCTGGATACAGCT
GCATAAGCACCATAAGCAACACTCATGGACAAAATAGGCACATCCTCTGTACTGCCAGCGTAGTTCTTGTTGAGAGCTTTCCT
AAGATAAATCAATACATTTGTTGCAGAGGTACCCACAAGTGACGCAGAAGTAGCCACACAGAAAAGCTTCCCACCATTTCGT
ACTATGGCTCCACATCTTTGTAGCAGACTGTAAGAAGTTCCTGCCAAAGCAACCTGGAAAGCATTATCGGGACAACCTCGAA
AGAGGTTTGCAAAAAGTCCAGACTGCTTGGAAACTAAAGAGCCGAGAGGGACGGTTGGTGCAGGAAGAAATACCAACATAA

AGTCAGCAATAATTGCCATCATCACGTCAGCAATAACAAAATCCAGTTCTTTGGAAAAGTTCTCCCTTCGGCGCTCCAGTTCA
GCTGCCGTCTTTGTGATGGTACCAACCCCACATTCAATTGCCACTTTGGTCATGAAGAGGTCATCAGCCAGCAGCCTTTCTTTC
ATGCCTCCAAATTGAAGCAGCCATCGAAAAAATGGAACCTTCTCGAGCTCGAAATAGCGGCGTACAATAGATCCAGGAATTT
TACCCTGTTCTACAGCTCTCGCAAGATCTGCGGGCAGATCACTAAGTGCCTTGCCCAGGCTTGCAAGCACAAGCAATGCTTCT
GATTTATTGCTTGAATGTCCTCCCGGCTGCGACTCCCCATCGCCACCACTGCCTCCTCCCCCACTGCTGCCTGCTCCTGCACCA
TCATCTCCCCCTCCGCCACTTTCAAGTACGGCAGCTTCAAAGGACGCAAGCAAAGGCAGGTTGGGAGTGGACCTGCTGCTAA
CAAACCCAAAAACTCTTGCAACAGGGAGCAGGGGGGAGAAGTTTGACAAGGAAAATTTCTCTGCAGGTAACTGGTGATGGG
CGAAGGAGGCAATGTAGGAGGGCGCATGAGCTTGGAGCCTGCAAGTGGGAAATGCAGAAGGAGCGCTGTGAGAGAAGGAA
GAGCTTACAGAAGAATGAAGATTCATCTGTGCGAAGGCCATCTGAGAGGTTTAGGGCTTTCTGCGAGAAGACGCAGGACTCG
CAGAAGGGCCTGCAAGAAGAACGCAGCAGATCCTTTCCTTTTCCCCGTTCCAGCAGCCGCTTTTCCTATGAATATTTCCATTAT
TTACTACCTTTGTCCCTTCGTGTAAGGTGG

**SHOOT MERISTEMLESS (*STM*):**

AATTACCTGATTGCATGTAAACTTGACTCAAAAAAGTTGCCTCCAGAGCTTATTGATCGACCTCAACACACACACACA
AAGAAAAAAGCAAAACACATGTATACATAGACGCATACACACACATACTCACTGGCACACAGCGCTCTAAAGTCTCAAAAGT
CATACATACACATCCACAGCCATGCACATGCACATGTCTCCAGTAACATTGGTAATGTTTGAGCTGCAGAGATAAGGGTAAG
CAAGAAAAAATCTATGTGTGTAGTGACACTATGGACCTGAGCGCAGCTGATCATCAGCTCTTGTACAACAGCGCCATGGCCG
GGGCCGACGCAAGTCATCCTTCTTTTGGGTCCATGATGGCTCTCATGACTGCAGAGCAGGATCTTGAAGGAGCTAGAAAGTCT
CAGGTTCATCATTCTCCTCATTTTATATACTACAGTGATCCTTACCAAGCTGCAGTTTTTCACACGGACCAATCCGGCATGCCC
GATCAGCTCTCGACATACGACCTCATGATTGCTGCAGTGGACGACGCACATGCGGCATCTGTACGACAGCTTGATCATGGAG
GAAGAAGAGAAGGAGAAGAAGAAGATCAGAAAGTGCAGTTGCAGCAGAGCGCGGCGACGACGGTGAACAAAATCCACCCC
AAATCGGAGGTCGTTCCGTGCGGGAACATATCCGTGTACTCACAAAACTCTTCTCACAGCAATGAAAGGAGAGGTTTTTCTGC
AGGTTTGAGCTCTGGGGCATGCAGAAAGGAGCCAGCAGAGCGCGGCGACGACGGTGAACAAAATCCACCCCAAATCGGAGG
TCGTTCCGTGCGGGAACATATCCGTGTACTCACAAAACTCTTCTCACAGCAATGAAAGGAGAGGTTTTTCTGCAGGTTTGAGC
TCTGGGGCATGCAGAAAGGAGCAGTCGGAGAGTTTATGTAGTACGAGCATTACAGATGATTCAATCAAAGACAAGATAAGA
GCTCATCCTGAGTATCACAAGCTGGTGACGGCATACATAAACTGCCGCAAGGTGGGTGCGCCGCCAGATGTCGCGAGGCGGT
TGGAGGAGCTGAGCAAAGAGTACGATAACCCGCATCTGCTCAGCGCCATCGTTTGCAAAGCTGCTGCAGCTCCGGATCCTGA
GCTCGATCATTTTATGGAGACCTATTGCCATGCTCTAAACAAGTATGAGGAGGAGTTGTCTAAACCCTTCAATGAAGCCATGG
CCTTCCTGCAAAAGGTGGAGCTCCAAATAAGCCATGTCAGCAGAGCGGAATTCCGCGTACCTCTGCAAGGTGATGGGTCATC
TAGGCTTGGCATGGAGGACACAGAGCAAGACATCGAAGATGAGGAAGAAGAAGATGAAGGTGCCGGAGGCTGTGGCGAGGT
CGATTTTGAGATGGATGCTTCCATGGACCCCCACACTGAAGACAAGCAATTGAAAGAGCAGCTTCTTCGAAAGTATAGAGGC
TACATCAGTACCCTCAAACACGAATTCATGAAAAAGAAGAAGAAGGGCAAGCTCCCCAAGGATGCTAGACAGCAACTCCTTG
ACTGGTGGAATGAGCACTACAAGTGGCCCTACCCAACGGAAACGGAGAAGGGAACGCTAGCAGAGACGACGGGGCTCGATC
AGAAACAAATTAACAATTGGTTTATAAACCAGCGCAAGAGGCACTGGAAGCCGTCCGATCAAGATATGCGCTACGTGATGGT
GAACGGCCAGGATCCTCCTATGCAGCACGACCACCCGCCGGACCAACTCAATTAGACCATGCACGTGCAGCACAACCTTCTC
AGCAACTCTCTCATAAAAAGAACCAACGTGTACCTTTTTACTCCTAATTTTGTAAATGTTGAATGAGCTTCAATATATAAATTT
TATTTCTTGCAAGCTCCGAATTTAA

**ENDOPLASMIC RETICULUM AUXIN BINDING PROTEIN 1 (*ABP1*):**

GAACACTCTTGCAACAATTTTGCCATTATAAAAAGAAAATAGAAGAGAAAAAAAATCACCCAACTGATATCCACAAC
ATTCTTTCTCAACTAAATTACAATTAGAGCTCCAAAGATTTTTTTCTTGAACGTGTGCAAATCTATAGTGCATGGCTACTTTTA
CACTTACAAGTATTGGGTGTTTTACATACAGTAAGGACATTAAGAAACAATTTTTGTAGATACAATGGTTTGTTACAATTCAT
CAAACATATAATCTGACTCAAAATTCTTTATCTCATTTTCCAAACACATGGCATCCCACGCGTATGGAAACTTTAGTACTGCTG
CAGTATGTGGAGTAAACCAATCCTTGTAAACAAACGCTTTGATTGGAGGCCGTGATATGATCACAAGGACTTGTAAGTCTTCT
TCCACTTGTGTATTTTTTACCTGATGTACATGATTGATAGGAATTGTGAATGTAGTATTTGAAGAAATCATGAATTCGTTAGGT
TGTCCCGGAATCTTCTTGCCACTATCAGGTTCCATATATAGAGAGCCTTTTCCCTTCAATGTTACAAAAACTTCTTCACATGAA
TGTCGGTGGATTGGAGTCCCAGCACCAGGAGCAAATGTTTGAAGCCAGACTTCAACCTCTTGCATGCCATGGTGTGTTGCCCC
AGCAATAGTTATATGTGACACACCCGGCCGTCCAAAACTGTCTTCATACATCTCATTTATTTGACTTACAACCGGAAGTTTTGA
GTGTGAGCAGTCGAGAGGTGGGCGGCCGCTCATGCTCCGAACAATGAAGCTCAGCAGAAGCCATAGCAGAACCCCTCCCATC
GAGAGAGAGAGAGAGAGA

**PIN-FORMED 3 (*PIN3*):**

TTTTTTTAAGTAAAGGTGTTGATCTGCATTTAATGTGGATGCTTGTACATTCAGAAGCATATCATATACAATGAACAAT
TATGTTTAGATGTCCATGGTTAGTGGCTAACACACACCAAATGCCCCATCATAAAGGATGACTTCAATTTCCCCGACGGGGTA
AAGACGTGCTACATGACTCCCAAGGAGCGCAAACTGTCATCCTATTAAGAGTAATTGGTGCAAGCCAAGAGCCACATCATAT
AATGTTGTCAACACTCTTGAGTTAGTACTTGCTAGCCTCCATTTGCACAAAATCACCCCTACGCAATAACCGGCCGTCTTTACC
TTCAAACCCATGGTTGTTTTCAACTGTGACTATTTGGGCCCTTGACAAGCATGGTTGCTTCCAATTGTACCTATTTTTCGTATG
AGGCCGAATGTAAGCCCCGCCGCTCCGCTGACACTATCCAAAATTCCTAAGTAGTGGAAGCAAGACCAAGGGTGGGCTCTTT
TCCCCAAAAATTAGCTCAGTTGGTTGAACCTAACTTTACATAATCTGATTTTACTACTGATGTTTTTTACTAAAGTCCCAACAG
GACATAGTAAATTAGTGTTGTTGGTAACGCAACGAGCATGCCAAATATGACTGCTGTGCTGAGTATATCAGGGTGCAAGTCGT
ACTCTCTCGCAAAAACGAATGGAACGATCCCTTGAGGTAGTGCTGCCTGAACAATGGCTGCATGCAAATCGACACCTCGCAA
GCCAACTACAATTGAGGACGCCGCCATGACAGCGGGCCCCGATAAGAATCGAACTGCCATTGCAAAAGCAGCGAGTGTCGTT
CCACACGCTATTAGTCGAGGCTGAAGTGCCATGAACAAACCTAAACTAAACATAGCCATCCCAAGGCCAGCATCTGATAGTA
AGCTTATTGAGCCCGCAAGCATTTTTGGCATAGCGACATTCCACTTGAATGCAATTAGAGACCAAATTATGCCAAGTAAACTC
GAGTATGTGTTTGGATTTTGAAGAAGCTTCCGCCCAACCATTCGAAGAATGAGCTTAGCCAACACAGGGGCTGGAGGCATTTC
TTTGGGCTTACCAAAACCTTGCAGCTTCGGAGAATGTGCTCCTGCTGAGGACTTGGATATGCTTTGCCTATGTTTATCTGCACA
CCCACCTGTCTCCCCAGGAAATTTAGCTCTGCTACCAAAACTGAAGTCCTCACCTACATATGGAGCAGTTGGAGGGACCGGAT
TAAGTGGCTCATGAAGATCTCCATTTGGCGCTTGCTCGTTGTGAGGTTGCACATGCATTCGCACTTCTTTGTGGTCATGTGCAG
GCTTAGTGAGGTCAGTTGCGTTTACATCGTTGCCTCCAAAGACATGCAAGCCCCCTTCCGACATGGGGGAAGCTGTGGAGCTC
CACACAAACATGTGAAGCTCCTTGCCATCATCTTCAAGCCTTGGAGAGCCTCCAGGGTCTAGTACCTTCTTGGAGATCTGTGA
TGCTTTGGGCGAAAAGCTCGTGGACTTCCCTACTGAGTATCCAGGCCAGCTGCCTGTATTGACGTGCCCAGGACGATTCTGTC
CTATCCCATACGGTTCGTTTGGCATGCCATTTGCACCATAGTTTTTGGTGCCATAGCCATGGCTCAGGGTAGAAAAGCGAAGC
GGATGCATGCTTCTTGAACCTGGGTGTGCATCCTTATTGCTGTCCTCATGCACATTTGATGCCCTCGGGGTCGGAGCCTGCGAG
GACTGCCCAGAATGAACATCAGAAGCCGTGAAAGAGTGCCGCGGACTCACTGGCCTTGTTGACATGACTGAATGCAAATCTG
GATTGTTCATTCTATACTCTCGAGGAGTATTGTTTCGAGATGAGTAGACAGAGTACACCTCTACTCCAGTCAAGTTTGAAGGC
CTTGGCGTAATGGCCTTTGATGAGGGCATTGATGTCATTCCTATAGACCTTCGAGGGGATGCAAACATGTTCAAAGAAGGCGG
GCGGGACGAAGTAGACTTGCGTATAGTCACCCTTATCTTTCCATCCTCCCCTATGTCAGCTTGTGTTTGAAGAGGCTCCCGGCC

ATCTAGAGAAACTACATCCGACTCCACTTTAAACTCGACAATAGAAGCTCCAACACCGGGAAACTGTTCCATGACTAGATTCG
TGGCAGCCCTGTACTCAAACAAAAAAAGTAGGAGAGTGTACCATATGATGCATTGAAGCACAACAAGCTGCACCATAAGACC
TCCATAGAACATCCCGTACATGGCTTCCAGTAGGGGAATTCCCATGACTAGAGTGTTTGGGAGAGTAGCGACGGCAAAAAGG
GTGATCATCCAATCCAAGCTGCCATTCGGCGAGAACTTGACCCATGCAGCTAGCACAATCAAAACAGCTAACTTCGAAAGCG
TGTCTGCTAGAATGAAGGGGCCATCCATCTTGTAAGGATTATTTGTGGAGATGAAATGAAAGGATAGCAAGGGCACAGCAAA
CACAGAGACAAAGCGGTTGATACCCGAACATTGAACAGGCGTAAAGATTTTCCACCATCTCACTGACCCATATGCTAAAATC
ATCGCCACGTACAGGGGCACCATCGCACAAAGCACTGCGTAAAGGTCTTTGAGGGAAATCATGTTTGAATGCTCGAAAATTG
TGGGCCAAATTTCAGATTTTACAGTGGGGGGGATCAAATCACATATATATATGGATGTCAGACATCGATGAGGAGCCCATAA
GGTATGCAGAGATGCTCTTCTTTAGAGCTTATACATGTGTGTAAACGAAAATTACAACAAAGAGGCTGCATCAGCAAGCTAA
CGCTAAACATGCTGAAATGTAAAGCATGAAAGCATGAAAAAAAAACTGACAAAAATGCTATAAATACTTGTTAGGGAGATGT
CCAAAGCTCACATTCAGGGAACTCTTGCACCCAGAAAATGCAAGTAAAACAATTCAAACGATAGAAAATTTAAGCTACCTCA
CTCAAAAAATAGGCTACCAGTGGCTCAAATCCACCAATAATCTGTGCAGCCAAGCGCAGACAGCGACAAATTCCCTACGCGT
GAAGAGACGCAGCACAAACATGGAAACACTCGAAAAGCTTGTTAGCTTAAGCTACACTGCAGAAAAAAATCAGCTGGCCAA
ATGCTTATTTATCCCGCGAGCAAAAAAATCTGGGAGCTTCGAGCTCTTGTGCAGAAAATAAGAGCATCCTGAGGAGCTACAA
GGGCTGAAATCGGTGGAGAACCATTTTCTTCTTACCCTAGAATTGGGCAAAGCAGCCCCAATTGGGAAAGAGGAGGAGGGGA
GGAAAGGGGCAGACGGGAGAGAAAAATGAAGAGTATTTGCCCAATGCAGAAAGGGAGGTTGAGGGTCTTTGAAGGAAAGG
GAGAAGAAGAGAGGGAGAAGAAAAGAGGGGCATGCCCATGCCAATGCCATAGCAGCTGTGGTTGCGTGTAGGGGGGCGCCATG
TGCAAGGAGGCAGGTTAAAAGTAGAAGCAAAAAAGAGGCGGGGCTAGAAAGGGAGAGTGTGCAGAGTACCTTCGATGCTAG
CTCTACCTTAGCTTGCAACTCACAAACACACACACACACACAGCTAACCTGTTACCCAATGGTGGGTTTGAAGTAAAACCCAA
GCAAGGAGTAGCTGTGAAGCAAGCAAGGAGGATGACGATGAAGCAGGAGGACGAAAGGGGGTTCATCGCAATCGCTCGTAG
AATTAGCCTGCTGCCGCGCCGTGTATTTTATACACGCGAGAGAGTGTCGATGGACCCACAA

**ETHYLENE INSENSITIVE 2 (*EIN2*):**

CACACACACTCACTCACACACACACAAGTACCACACTCACGCACGCACCTACACACGTACTCACACGCAGGTGGGTG
GCTTTTTGGAGACACACATGAAGAGATTTACTGGATATGACAGCAGAGAACAAGAAGAAGAGATTATGAAGGAGACTTTGCA
AAAGCGGTGACCAGGAGGTTTGCAAATTGCCATGGAAGCTTTGTGGGCACCTGTGAACTCTTTCTGTCTTATCTTAGAAAGGA
GGATTTCATGCTCTGTCACTTTGAAGATGCGCCTGCAGGCCATGGGTTTTGGATAAATTCAGCTCTCTTCTGCATTGGCTACCA
TCTCACGAAACCCTAACCCTAATTCTAGCCCTAGTACAAGCATAGCTAGATAAGCTTTTTTGGGTCCTTTGAAGAATATCTTTC
TGGGTCCTTGAACAAATTGGGTCCTCTGAACAAATGTAACAAGGCAGGGAACTTTAATTTGGAATAAAAAAGCCTACTCCTG
ACCTAAAAGTAGTTTCCCCCACCCCCCCCTCCCTGCTGCTCATTCAATGGCTTGGAGTACCTCTATATCATGCTTGGGTCCTGC
TTTTTTAGTGTATGTGGGATGTATGGACCCTGGCAAATGGGCTACTGCTTTAGAGGGCGGCTCTCGGTTCGGATTGGAGCTTG
TGTGGGTGATGATTGCTGCCTGTGTTTTTGCTGCTTTTTTACAGTCCCTTTCATCCCGGCTCGGCCTCGCAACTGGACGGAACC
TTGCCCAGATTTGCAGTGATGAGTATCCAAGATACCTATCTGCGCTATTGTGGTTTCAGTGTGAAATTTCAGTTCTGATTTTGG
ACCTTACCATGGTGTTGGGAATTGCGATGACTTTGAATGCGCTGCTTGGCATTTCAATGGTTGCTAGTGTGGTTGTTACAGCAG
TTGATGCGATAGTTTTTCTTCTGATTCTTCCTCATGTGGGACTATACAAGGCAGAGATTTTTACTGCAAGCATCATGGGTGTTG
TTCTTTTATGTTTTAGTGTGGAGGCTTTATACAGCAATGCAGCCCTCACACCTGGCATTTTCCAAGGATTGATCCCGAAGTTGA
AGGGAGACATGCTGTACATAGCTGCTGGCATTTTTGGTGCAAACATTATACCTTGCAATTTTTATCTTCATTCAGCACTCGTGC
AAGATGAGAAGCATACAGTGAGCTATGAGCCTGGTGTGTTACTGCATCATAGCATTGTAGATATTTTGACTGGCTCAGGTGTT
ACCCTGCTCATCAATGTAGCAGTGCTAAGCGCAGCAGCAGCAGCGTTTCACAGCTCTGGTCATGTGGTTATTACCCTTCAAGA
TGCCCAAGCACTCATGGAACAGATATTAAACAGTTCAGTGGCACCTGCAGCTTTCGGATTTTCCCTGCTTTGTGCTGGTCAGCT
TTCAAGTTTTAGTCTGACAAAAGCAGGGCAAGTGGCAACAGAAGGATTACTGGGTTTTAAGTTCCCTCTTTCTTTTCATAGAG
TTTTTACCAGACTGGCCGCGGCATTGACAGCTACTTGTCTGCTATGGAGTCTCGGGAGTGAAGGGACATATCAAGCCCTAATT
TTTTCTCAGGTTATCCTTTCATTGCAGCTTCCATTTGCCATAGTTCCCCTTCTTAAGGCCACCTCGTCTCAGGCTGTCATGGGCA
GTTTGAAAAACTCCTTGCTTGTGGAGTCGCTTTTATGGGTCTGCACAGGATTCTTATATGTAATGAATGTTTGGGTGGCTTTTG
ACATGTTCTTTGGTGAAAGCGAAGAGTACGTGGGATCAGGAAACTGGAATTTGATAAGGGACTTGGTTGATTCGAGCACAAG
CTGGAGAGAAGTGATCCGTATCCTCTGTCTCATTACTGTGGTTGGTGTGACAACAATATCTACATTGCTTATTATGCGGATGAT
TTCTGCACCATTCAAAGTACAGGACACAAGGCTTCTAGGTGGACCTGTAAGCAGTTCTGTGGAGTCTACTGTCTCAGGACATG
CAGAGGGTTCTTACAAAAGTGACACATTAGAAGAGAAACTGACTCTACCTGCATATAGTCCCCTAGATAACTACTCCTTCACC
GAAGAAATCGATACTAAATATCATGAGGCTGTTAGGGATAGCGAGTTTTTACTGCCTTGTGAGCTCATCAATGAGCTGCTACC
GGAAGGGGGTGCAATTTCTTGTGCCAAAAGTCTAGAGGTGGAACAAGGTGAAATACTTGATGCATCTGTGGCTCAGCAGGTA
GTAACGGAAAAGGAGACTCACATGGAAGAAATGCTAGAAGCTACTACAACATTCTCAACAGTGCAGGAAAGCAAGGAGCAC
TGTCTGGAGCCCCTGCAAAAGGATCCGGATCAACAAAAGCAACCCCAGTCAACTACATCTACTGATAATTCCTCCTTTGTTGA
ATCGTCCTCAGTAAACTCTTTGTCTCGCGATAGTTTTTTGTCATCATGGGCAGTTTCTCCTCCTGAGTTAACAACAGCAGAGAC
AAACAATGCACCTGCCACAAAAGTTTCTGGAGGTGGTGAAGCTGAAAATTTGAAAAGGGTTATTGAAGATGTTGATCTTGAG
CTACTTGACAAGGATGATTATGATATTGACGGATGGGAAACCTTAGATCAAGATGATGCTCTTCAGGATTCCATATCAGTTTT
AGGTGGCAGTGTCAATTCATTAACATTTGAAGAGCCAGAATCGGGGAGAAGCTTTAGTAGGAGTGATGCAAGCGAGGGCAGC
TGTGGTGGTAGTGGGAGTGGCAGCTTGTCAAGGTTGTCAGGGTTAGGCAGATCAGCACGGAGGCAATTTGCTGCCTTGCTTGA
TGAGTTCTGGAAAAAGCTGTATGATTTGCATGGACAACCAGTTTCCCAAACGCATGGAAAATCACGAGGGGCTAGCACCAAG
ATCTTACAGTCTGATACAGCAGATCACCAGCCAGTATCAGCTTACAAGGAGGCATTTGGTCCAAATTCCACCTGGTATACTTC
TAAGCTTGCACAGAAATCTTGGCAGAAGAATGTGAAAGGAGATATGGCAGGTCAAACTGATGCATATCTTTACCCTCCTTATC
TAAGCTCCACATCAGACTCTTTGGTGGCAATGACCCCTTTCGTCAAAAGCATTCTTCACTTGATGTGATTGAAAAGCGATATT
CAAGCTTGAGATATCCATCATTTCGGGATGGGCTTGATAACCAGCCTGCTACCATTCATGGGTACAAAATGGCTTCATATGCA
GGAAGGAATGGAAGCTTCCCAACAGGTGTGGACACTAGCTCTTTTCTCATAGAAAGGCAGCGTCAGTTGATGCAGCGGTCTT
CAGATGAGCAGTCTTTTTCTGCCAGTTCCTTATATGGGCTTCCAGCATCGGCTTGCTCCGAGCTAGATACCAGCATGTCGACCA
GTTCTATTCAATCATTCCTGTCATCACCGTACGCGAAGCAGTCATTAGGTAGACAGATTGATAGCTTTTCCCTTCCAAGCAAAT
CTTATCCCATTCAAGCAGACCGGCCTGACCCATACTCAAGATCTCAATGGACTCCATATACAGGGAGAGATTCTGCAACTTGG
GATCCTTTAGTATATAGAGCTACTGGAGAGTTAGACATTCCTCACTATGATCAGATAGCTCTGCAAAGAGACAAAGACTCTTT
GAACTCCTCTTTTGGACATCGAACTGGTGAAAGAAATTACTTTGGCACTGGTTCAAACAATAGCAATCATTTTAAGCCTCAGA
TTGATAGAGGTCCACTACCGTTTGACGAAATTTCTCCATCTCCGTCCCACAAAGATGCCTTTTCTATTCAATCAGCCCCTCAGA
ACCAATCCCTGTGGGCCACTCAACCATTTGAACAGCTTTTTGGGTCCGTTCGCTCCCCAATAGGGAGTGGCAGGGCAGCCACA
CGGAGTGCTACAAACAGTTCTTTTGGCCAAAATGGAAGCCATGCATCTGTCAAAAATTTGCAGACAACATTTGCAGGAAATG
ATCATGAGATTGAGGTCTTGAACATGCTACGCTCGTGTATACGCAAATTGCTGAAGCTTGATGGCTCAGAATGGCTTTTTCGA
CTTGAGAGTGGCTCAGATGAAGATTTGATTGGTACAGTTGCAACTAGAGAAAAGGTTTTACTCGACGCAGATGCCAGGGAGC
TACACAAGCTTCACACAGGCAATGAGCAGATTTGGGGTTCCAGCTCTCAAAGATCAGTTGCAAGTAGTAATAATGACTTGGC
CTTCTTAACACTTGCTACCTCTGGTGTACCACATTGTGGTGAGGATTGTGTATGGAATGCAGAACTTTTGATAAGCTTTGGTGT

GTGGTGCATTCATAGAGTTTTGGAACTGGCACTGATGGAAAGTAGACCGGAACTTTGGGGAAAATACACTTATGTGCTAAAT
CGTCTTCAGGGTGTCTTGGATCCGGCATTTTCAAAGCCGCGAATTGTGTCAACGTCATGTATATGTGTGATATCCGAGGCTTCC
GATGGGCAAGGAAGAGGTTTAAGTAAGCAAGGAAGTCTCAAGGACGGTGGGATAGGAGACATGCTAATAAGCAGGTCTCTT
TCTGGTGGTTTTCCAGGGAATTCTTATCAAGGGTATCCACAGTCTTGGCCTTGGGGACGCAACTCGAGCGCAGGCAAAGGGA
AAGGGGCCAGCTCATCAATTTTCTTGGAAATCATTAAGGATGTAGAAACAGCAATAGGTACTCGCAAGGGCAGAACTGGAAC
TGCTGCAGGTGATGTAGCATTTCCCAAAGGTAAAGAGAACCTAGCTTCTGTTTTGAAGCGTTACAAACGTCGTCTCGGAAACA
AAGGCCCTGGAACACCAGCAAATGGCAGCAGTAACAATTCTGGAATTCGAAGGGCACCAGCATCAAGCCCAAATATCTTCAT
GTAGCATCTGACACCAGTCTCGGGCTATCCATCCCACAGACATCTAATACGCTTGCTGACAACCCTCGCTTTAGCTTGATCAT
GTCATTATGGGCGTCGTGAAAACTCAACAGAGTTAGAGCAGTTCATGTGTGTCATTATATGGGTGTTTTAAAGGAATGGAATT
ATAGCAATTCATGTGTGTTCCACCTCTTCATGGATGTTAAGCCCATACAGTTGGGCTTGGTCATACAGGAGGACTCGGTGCAA
AAAGGGCACCAGCTTGTTGGCGATGAAGATACACAGCGTTCTCTATCCTCAAACCGAGGATAGTAGTAATTGTTCATATTACC
AAAGGAAGCTTGGGGATAATTGGGGCTAGTCTCGATACGCATGCTTTGAAACATCAACGATGGATCACACCAGGGGCTAAAG
CTTCAATATGTCAGAACCACTTCGGCCTAAAGAATAAGGCATGAGCAGTGGCATGTTTGACGCAAGTGGGCAGTGCTGTGTTT
GTGAGCCAAAATTTTTCTGGATTTCGCTTAGCTGCATCGGTGAAGAAAATTTATCCTGA

**ETHYLENE OVERPRODUCER 1 (*ETO1*):**

TCAAAAGAAAAAAGTCGAATAGATGGGTCAGATAACATGTGATTGTTTTGCTCAATTTTCTTACATGAATGCTAGATT
CCAGCAGCTTGAAAAGAAACAACTGTTTGGATGCCACATTCAACATTAAGATCATGTATAGCTAATTCAACAAGGGCCGAGG
AGCATGTTTCCTGAGTGTTTCATCGCTTGACCACAAAGACAACATTGTAACAATGCCACCTGGAGATATGGAGGTGGACACG
GGAAGATTTTGACAACCTTGATGATTGCAAGACCCTCCCAAATTACATTTCATTACAGAGAAGCAATCTTGATTTCTCCCCCTT
AGTGCTGTTAGTTATAGTTACAACATCACGTCAAACTAGCATACAGATGATGGTACACCAGGATTTACTCAAAAGTGGACCAC
TAATGATCCTTAATCACAGCTTTTGAAGAATTGGCTGAGGAATACAAATTGTCCTCATATGCAATCACAAAGCTGACAGATAA
AATCAAGGCTCCCTGCTAGAGACGCGGTTATGAAGCTGCATTGTTTCCACATGGCTTGGATCCACTGACAAAGCTGCTCGACA
ATCACGTAAAGCACTGCTTGCATCTCCAATGCACTCGTGAAAGGCTGCCCGCAAATGAAGTAAATGCAAGTCAGCTTTAAAG
GCTATAGCTTTTGACAATTCATCAATGGCTTCCTTCTCTTTATGACTGTCCATTAGAACTGCAGCCCTGTATCTGTAAGGGTAT
GTTCGCAAGGGATCCAACTGGGTGACCATGTTCAGATCTGCCAAAGATTTTGTGCGTTCACAGTATTCAGAGCGTTTTTCGAA
GGCAGACGCACTACTTTTTGCCTTCTCAATCAGCTTTGTCATCTCTTCATAGGCAGAAGGTCGGTCATTCTTCAAATAATACAC
TCGAGCAAGGCCCTGGTGTGCCCGTGTATGACGAATCTCAAGAGCACTAAGATAAGAATCAGCTGCAAGGTCTAGCTTGTCA
CAGTCGACATAAACACTACCTAAGTTATTCAAAGCCTGCCCCTTGCGAAGCCCATCTGATGGGCATTTTAGTGCTTCCTTCAG
AAGATCTACCACCTCTTTAGACACATCAGCGTCCCGATTAGTATCTGCTAAAGCATAAGCCTTCAAGAAGAAGGCTTCAAATG
AGCGTTGTATATCTATAGACTTCTGTGCTTTGACAATCGCTTCCTTGCAATGCCCTGTATCATAAAGAATCCATCCCTCATATA
CTAGCCGTTCATGCTCTGTGGGAGCATTCTCTCGAGCAAGTTGAAGACTCTGCATTGCAGACTTCGGACAGTTCAGCCGCAAG
AGTAGGAGAGACTGTCTGAAAAAGAGAAGAGCTTTTGAAGGGTCTGACTCAAGCATTTGATGAACAACAGCAAGTGAACCG
ATGTCATCTACAGAAGACCATCGGTCATACAATTGCATCCAGCAATCTGCAACATTCCACTGCTCCACATGAGGGGATAAAA
GATCTAGCAACTTGGAGGCCCTTACCCTACCTGAGTACAGAGTATACTTTGGGTTGAGGGTGAGTAATGCACGGACATCCCTT
ACAGCCCCTGCATAGTCCTGAGCTGCCAGACAAAAATATACTCTCAATTCGAGACAATCAGTGGTGACCTTGAACCCTAGGAT
GCGATTAATTTCCAATATAGCCTCATCAACTTTAGAATCATCCATAAGCCTTGCCGCTCTATACTTGTAGGGGTAAGTTAATGT
TGGATCAAGTTCTGTTGCCTTGTTCAAGTCATCCATCTTCTCTTCTTCATTACAGTACAGAGACCTCTCAACATACATCCACCC
AGTAGTAGGCTTTTGACTGCACAATTTCGTGCTAACTGTGGCAAGAGCAGCATGTCTATGTGCCAGCTCATATTTGGCACGCG
CAAGACCAGCCAGAGAGTACCGGTGACCCTCTTCAACAGCTGCTTCAAACAGTTGATGGGCATCCGACAAGGATTTCCTTTCC
AGCATTGCACACCCGAGTTGATGCAATGCCAATGCTTTCTGTCTGGAAGTTGTCGCCGATTCTCGTACGTGCTCAAGAAACAT
GATGCACAAGTCCGATCCTGCATCATCTTCCATAGCTACTTGGCTCAGCAAACTGTAGAAAGCAAATGCGGAATGCGTCCTCA
TCATAAACATCCTCATATCAGCTCTGCTTAACAAGGATGCCACTTGTGGATTTTGCAGTATGCCTGGAAGACCTTGCAAAAGA
ACCTCCAAGCATGACGCAACTAGTAATGGTGCACTCTCCTCCAAGCCATACTCCATCAGCATAACTGCATCCTGCCTAGTATG
GACTAAATTTGCAAGTGCGCTGTCACAAGCCTCTTTGATGTGCTCGCAACAGAACCTGTTAGCAAATGTAAGCACTTCCAATA
GTCCATCAGGCTCCAGGGGAGGCAAGCTACCTGTTTTACTAAACTCGTCGACAGATCGCATACTTTCCACTGAAACCCCACTT
TGTGAAAAGTCTATCCGCTTCAATTTAGACTCTGTGAAGCATCCATTTAGCATAGTATCAAAAGGGACAGACAGGGAAGCAA
TCCTTTGGCGCCCACATATTACTTCTTCACTACCTATGGAAAATGTGATGATTTCATCCAAACTGCCAACACTCAAGGTATCCG
CATCCGCGCAATCACTACTATCCCTGCTCAATTCTGAATCTGCAGTACACCCACCCCCATCCGAGTGAGCAGTTCCTGCATGCT
TGTCTAATATGGACGACCTGCTACTCAGGTCCTCCCCCTTTTTCTCCAACTGCAACCAAGCTTTGTACACGAGCTTTGCATGAG
GCGTACTTGCAAGATTAATGGCTGCTCCCAGACTTCGACGAACCAGTTTATTCTCACCAAGCCCTTTGAAAATGAACGCCTGC
AATAAACATAAACTGGCCTTATCGTGTTCTGATGCATTCTCTATCTCCTCGTGCAATTGTGCTAGCGTTTCCACATAGTCTAGA
GGCTTGAAATGTGATTCGACTGGGGGATCCGCAAGCTTGGACGCCAAAGAGCTGCGCGGTCGTTCTGACATGCTGTCCCACTG
CTTTTTGCATTTCCGTCGCTCCAATTGTACCCAAGAATCTGGTGATTTATCCAAGGGAGATAGCGCATGAACCTGGCTGCCCTT
ACATGAATCTGCCAGCCAAAGACCTCTCATCTGGCTGAAGAAGGCCAAAGGAAGCTACTTTCACAAAGTTCCAAGAGATTTG
CAATTCCGAAACACGTTATCCTCTGCTTCCATGACTTCTGCATCTCTGTGTGTGGATTCAACAAATTGAAGAGAAATCGGCCTC
TGATTCATGGCCGCAAGCTCCCTCAATCAACCATGGAGATACATTGAGGGTTTGACGCAGGAGCGAGAGGGTTGCTTGCGAC
TGCAAGAAGCTTATGCACAGATAGACGAGTGGAAAGACAATGAGCCACTCTCCCTCTCTCTCTAGAACAGGTGGGCAGAGAA
CGCATTTCCTCTCATCACTCATAAACTCTCTCTCTCTCGCTCATGGGGGGTGGGGTACGCTTCGATGTGTGTTAACCTTTTTAGT
CCGTCTGGTAAAG

**REVERSION-TO-ETHYLENE SENSITIVITY1 (*RTE1*):**

AAAGCCCAGACATTCTTTCACTAGGGTTTAAATTGCTCCTGCTTTTGTCTCCAAAGGCCTTGAGGCTTGACCTTTTGTA
ATGTTGATCCTACTCATGTCGTGCATATGTACCTGCTCAGCTTACATGAGGAATATCATGTGGCATTTATGACCGACCTCGAA
GAGGCCGTGCCGACCTTCCCGTTAATCAATGGGCATAGCTGCATAGATGATGTAAAATTTGAGGAGGAGGAGATGTCATTAA
AAACGAAAGCAGGGAATGCCGTGGTTGTTGAGGCCGTGCAAAAAGGCGATTGTGTCCGTGTGGGAAGCAAAGCCGCCAAAT
TAGAGATTGTGCAACCAACAAGGGGTAGAAGCAGCAAGGTCAGCCATCACTCCAATTCTTAGGAAAGCTTTATTGACGAGCT
GGAGCCACAAGTTGCAAGACCAAGGGCAAGCCGAGTCATGGCTGAGCTCCATGATGAGGAGGCTTTGATGCATTCAAGTGAA
TCTTCCAAGCCAGGCATATCAATAGACCCTTCCTGCTCTAGATTCCCCCATTGTATAGTTTGGACTCCATTGCCAATAGTGTCC
TGGTTAGCTCCATTTGTTGGCCATGTGGGTGTTTGTCGTGAAGATGGTATAATATTGGATTTTGCGGGCTCTTATCTCGTGAAT
GTTGACAATCTTGCATTTGGTCCGACAGCAAGATATGCTCACCTCGACGGGCAACAGTGCTGTTTTCCTCCTCATCTGTATGGT
CACACATGCGAATCCAGATTTGAGCATGCAGAACTCGGCACAGCAACCTCGTGGGATGATGGACTCCGCAGTTGTATGCAGA
CTTTCCAGCATAAATGCTTTAACCTGTTTACTTGTAATTGCCACTGCTTTGTCGCCAGTTTTCTGAACAGGATTGCGTACCAAA
AGTCAATCAAGTGGAACATGATCGACATTGTATCATTGGTCTTGTTGAGAGGCCAGTGGATAAGCAAATGGTCTGTAGCAAG
GTCTTTGGTTCCCTTCACAACAGTGATGTGCCTTGGCTTATTGGTAGCTGGCTGGCCCTTTTTTGCTGGTTGGGCTGTCTTTGAC
TTTCTCCTCATCGCTTGGTTTTTGTTTGGTACTTACATTAAAAGAGAACTCATTGAGTCCTAGTTTTAGCTTGAAAGGTCATGT

ACTTTCCAGATTCAAGTGATAGAAGAGCTTTTCTAGGTTCTCTCTCTTACAGTGTACTTGTATGGATGTATACATGTACTGTGT
ATGTTTATGTTTG

**STEROL 1 (*STE1*):**

CAAGTGATAAAATTTAACTGAATTATCAAACCAAATGTCCTTGGTGATTTTTTTTTCATACAACTCTCTGAACAAGCTG
GAAAACATTGAGTAAAATAAACTTGAAGTACTTCAAAGACACTATTAGTCAGCAAGTAAAGTAACACCTTTCGCATCAGTAA
AAAATCTTATCTAACATTACAACGCTGTTTCCAGCTAAAATAATTTCACTACAAGCTGCTCGCATGACTGTTTCATCCATCTAG
GAGGCCTTCGACTTTTCAGGTTGAAAAGGGTCTCGCAGGGTTCCAAACATCCAATCCATCCAGATTGTGTAGTGTCCATAATT
ATGTCGGTATGTGGTATGATGGATTGTATGATAACCCGCACCCATTATTGGCCAAACTTTTCCATGAATGCAGTCATGGATAT
TTGTTGTCCATATGCTTTCCACAAACAATGAAAGTTCATGAGTAAAAAAATGTGTGGGCACAAGAAATATGAAGATGACATG
GGGGGATGCCTGTAGTATTCCATCCAATGGGTTGAACGCTAAACCTGCAAATGGTGACAATGTGTTCTCCTTATTGTAGATGT
GGTGCGTTGCATGCAACCATTTGTAGAGCGGTTTCACATCGTGTAAACCCCGATGCATCCAGTAAATCCCAAATTCCACAAGA
GCAAGGTACAAGAAGGTCAACAATATGTACCTCGGAACTCCAACATCACCTATGTTAAGATAGCATCTCGTCCACCCTCTTTC
AATCATGTACTCAGATAGTGTGGGTAAGGAAACATAAATAGGCATGCCTTTCATGCTCACCCAAATCTGTAACAAAATTGGCT
CTTTGGACGGTACACCACCTTTGGGAAAATAGACATGTCGTTTCAAATAATAGATGTAGAGGCACCAGAGACCACCGGTGGT
GAAGTAAATCAAAACACCTGCAAAATAATTGCGGAGCCATGTTTGCGCAGTATGTGGAATTTTGGATCTAAATTGCGGAGAT
ATAAGGAAACCCAGAACCAGGTCGTTGTACCAAGATGTCTCTTCCACAAAGTATTGTAAGAAGGGGCTTGATGTGGGCATTG
AAGAGGATGTCTCTTCAGTAGCATTCACTGCGACGGAGGGGAACGCAAGGGCGACATGGCAAGCTGTGAAGAGGATAAATA
AGGCAACAAAATGCAAGTGTGCCATACCCCAAAAGCTTATGCAGCAAGAGCCGTTGTGTGGAGGAGGCACTCTGCAATAACC
ATAGCCAAAGAGAAGCGCGCCCAAGAACAAAGGGAGCCTTCAACAATCTGTGTGAGGGGTAAAAAGAGCGATTTTGATGTG
TGCGCGTGTTGAAAGAGAGAGAAGATGCGAT

**GPCR-TYPE G PROTEIN 1 (*GTG1*):**

AAATTGCCAGAAAGACCAAGACGGACTAATTTGAGCATTTTAAGCCATGATAAAAGATGTGTGTTTGTAGTATTGGTC
CTCTTTTAAAGACCAATAAGAAACTACGAACGATTAAATAGAGATAAGATTAGTACAACCCAAGCTCCTCAATCTGAATGAT
GTTTACAATTCCTAAACACAACTTCATATTGATACATGTCAACTGCACCTTCGATCAATAAGACAGTTGAACAACCCTATCTA
AAGGCTTACAGGCACTCACAACTTGCACCTTATTTTCTTTTGATATACATCCAATTCAAAGGGTACTTTATGACAAGTTAAAAT
TTACATTCGTGATTCTTGAACAGCTTATAGAAAACTACAATGCATCACAGATGATGCTCTTTCACGATCAAAAACGGATGAA
GGTTCCATACAACAAACGCCAATGATACGATACTGATGTGGATGAGCTTGAGATTTGGACAGATGGAAGGGGTATCCATGGA
TGTATTTTGCAGCTAAACACTAGTACATAACGACTAGTACATAACGACCTGCTCCTTGGAACTTTCTGTACTTTTGAGACAAGT
AATTTGCTTCTGCTATTTTATATTACCATTAGTTCATATTCAATAGGATGCTTGAGATTTGAAGGGACAAGAGGGTTGTCCATG
ATCAACAGTATATGCCAACCTGCTCCTTGAAGCTTTCTGTACAATTAATTTGCATCTTGTGTTTTATATAGCCAAAGGTTCATC
AGGCTTCATTATTCAATAGGATGCTTGTCAGACTGACGGGAAGTATACTGCACCGCGATAACTAACAATGAGACAAATGAGC
TTGCCACAAAAATAGCATCAAACCACCTATGATAGAAGTTGAACTGTATATCTCCACCTAAAACATCTGTGATGATAAGCCTA
TACTCATTCGCAAGGCTCTTCCTTATTAGCAGGATTGAGGAGATGAAGTACATCCCCATTACCTCCGACAAAAGCAACACAAC
ATTGGTTGATGAACCACTCCCACCTCCCGATAACACATAGAAGGACTTCATGAGATTCTGGAGAAAGCCTCTGACAGACATTG
TGATCAGCATTCCAATGAAAACAAGCGATACATACTGAGACCAAAGAGCAACATTTATGCCAATATGGAAAAATTTCAAGAA
GAAGCTTAGTGTCTGCGTAACTGGATCTACTGAATTTGACTCCTTGAATATTACACTTTGCATTGACTTCAACATCTTATAGAT
GCAATAAATAGAGAAGACGTAGCCGACTAGATTCTTCATGTGTCCTCTCCAGGTTCTTGAATAAGCTGCTGCTTCCTTTGCCTG
CCGCAATTCATAAACTTCCAAAAACAACTGCCTAGAGAGCTCTTCCAATCCATTAATTTCTGCTTCCATGCTCTTGATATCTTG
TTCCCGTTCATCTTCTTGGACGGTTCGCACCACCGTGCCAATCAATCTCTTGAAAATGGATCTTGCCTCTAGATGCTTTTGCGT
ACCTTGCAAGCGCTCCATTTCCATGCGAGAGAGAATTATTTTCTTTTTCTTTGCAATGCATATGTCCATGGCTTGCAAAAGTCG
CCTTTCAAGGGCAACAACATCCACCTCTTCTATTTCCCTGATAAACAATGACAAATAGCTATATGGAAGATTAACAGCTCCAA
ATCCAGATAGTATTGCCATGACTGTCACACCAATAACTCCAACTCTGCTAAGCAACTGCGGCATTGTGAAAAAACCTTTGTCT
GGCGATGGCATTGGGAAATGGATACCCATTCGCCAAAATGCGTAAAGTAAGGCGATCAAAAAAACAACAGCTCCAGCAGCT
GCTCGTTGAAATCCCAATCCTGTGTTACGAAGCGAAAGATAACAGTGGTAGTATGGTAGAACAAAGACGAGCAGTGTGATCA
AACAAAAGAGATCCATCTTCCAGTTCATCCACCTTGCACTTTTGGAAAGAAGAGGAATGATTTCAAATAAGACGAGCTGGAG
AAGGTTGAAGGAGAAAGCGAAGACAATGCTGAAGAGGATCTGCACAAGAGCGCTTCTCTCCTCGTACTCCTTGTAGAGCTTC
CTGTTGAGGAACCACAGGCCGCTCCATGCCAACCCTGCTAGCGATCCTGCAATCACCAACCCCTCCAACGCCACGTACGCCAT
CGCCATCCTCAGACCCTCATGTGAGAGAGAGAGAGAGAGAG

**JASMONATE-ZIM-DOMAIN PROTEIN 4 (*JAZ4*):**

CTGTACGTACGTAGCCCAAAAGCTTAAAGGGCCTTCAGGAATCACATTCTCGGAGCCTCTCAGATTCTCTCCCTCTTTC
CACAGCGCTTAAGCTGCTATCCTTTTGGATCCCTAGCTCTCTGCACAGCATTTGGGCTTTCTTCAGCGCCTGCTCTAGCTCTCA
TTTGGGTTTTCTTCATCAGCTCTAGCTTTCATTTTCAGGCCTCACTTTTTTGGCGAAGTCTCCATCCTTTCACTGCATGAATTTT
GTCTCTCTTGCATTGCCCCCTGCACAATTAGGGTTTTCTTTACAAAACCTGCAGAGTGTTCGCTTTGCTGCTTCCAAGCTGATG
AGATGGGCCGATCAGATCAAGAGAATGCCGGGGAAAAAGATTTCATGCGGTTTGGCAGCGACACTGCAAGCTGCGCCGAGG
CAAGAAAACATATGTATGCAAGCAGCCACGTAGTACGCTCGGAAAGTAAGCACATGTGCATGAACAACCACATGTATAATGT
GGATCCTCATGGGAGCAGCCAGCATTATGCAGACGCTATTGCCGTTGTGAAGTCCAATATTCTGGATGCTGTTCAACGGCCTA
CCCCTTCCCGGTGGTTGTACAACAACAACCTCACAAACAGCCTGGCAAATGGGAGCTTGGCCTTTGCTGACATGGTGCGCTCT
GATTGGAGTAATACTGGTGGCAGGCAACAGTTTTTTTCTTTAAATAGCTCACGAGAGCAAAAAGTAGTGGAGAAGGTCAATG
ATGCCTACCAAATCATGCATGCCAGGGGCCCTGACAATGCTGTGTTGAATAGCACTGCCTTGAATTGCATTTTGCCAGCCAAC
TCCTTACCAGGGTTGGGAATCAAGGGGTTCCCTTTTTTAGCTCCGCACCAAATTCCAACGGCAAGTGGTCAAGCACTTACTCC
TGCTTTGTTAGTGGGTGGGGGTCCCTCTACAAGAATGCTACCAGCTGACTCTAGCATCAAACCAAGCAGCACACAGCTCACTA
TATTTTATGCAGGCAAAGTTAATGTCTACAACGATGTGTCTTCAATCAAGGCCCAAGAGATAATGCTGATGGCGGCGGCTGGT
AATTCTTTGCCATTAAACATAGATATGTGTTGTGGGACATCTATTAGAATGGTAACACCACCTGTCCCAGGGTCTCTCAAAAA
TGCAGACCCTCCGGAATTGTGTCAGATAGACCCTTCTACGTGCCAGTCTGATCAGTTGGCAAGGGATCAGGTTGGTATAGGCC
CCTTACCTGCCCCTATACTTCAGGCATCTCAAGCAGAAGAAAGTACTCAAAGCACATATCTCGACGATGAGCCTGCCTTTCCA
CGAGCTTTACCTCATGCAAGAAAGGCATCCTTAGCCCGATTCCTTGAGAGGCGAAAAGAAAGGGTGCTGTCAAAATCTCCTT
ATCCGGTGGAAAAAACTTACCTTTGAGGAAGTGACCTTAATGAAATCAATTGTCCTTCCCATATCATTGTGGGGATTTATATA
GCTCGTGTACAGCAAGCGATTGCAGGAGCATTGTTTTTAACCACATCATCATGTGGATTTATAGAGCTCTCAGTTCTTGTACA
TCAAGAGATTGCAGGAGCCATTATTTTTTCCTCTTTTCATTATTTGTCAGTAGACGGGGCTAGTTCTACATAATATGCAGCTCA
CCCCGAACTGACTTCTATGCATGTTTTATCACAAATCTTAAACAAATATAAAGCCGATGTAGAAAGGCTCCATCCGTTGAGGC
CTTACTTTATGTGCTAAGTGATTGTTGATGCGTGACATTGTTATGATGTGGAGTAACAATTTGACTCTTTTGTTTTGCAAATTG
ATGCCAAG

**DE-ETIOLATED 1 (*DET1*):**

CACAAACTCGTCGAAATTAAGCTGTTAGGCACTGCTCCCTAAAGACTTTTTCCACCACCACCACCAACAAATCTGGCA

GTCTAAAGCTTATCCGACTTAACAATTTGTTAATCATGGATCCAAATTACTTCTTACATAAGGCTACTCTAAATCTGTCCTAGT
TTACAAATCAAACATTTTGTGTAGTGTTTCATAGGTATACCCCCATGTGCACATGGGTAAAAAAGGGCGGTAACAAACCGAA
GGATAAAAACCATTGCTATCTTCGCATGTGAAAATTTACAACAGCGGGCTGCATATATGCTTGCTGGACAGTTATGGCAAAAG
GAAATATAGGATGAAAAATAAATGAAGCTACTTTCTTCGCACGCCCATCACCAGTTCCAAGCTCAAGGCCTGGATTAATTTTA
AATTTGAGGACATTTGGCTTGCGCCTGGAAATAAACTTTATAGGGTGCTCTGTGCAAGGCTTGTATTGATCGGTTGCAGAAAT
AAGCTTTTCATCAAAGTGGAACAAAGACTGATCAAAGTATGGTGAAGGACTGAGTGATTGTGCATTGCATGGAAGAGAGGCA
AGGGTTCTCTTAATAACCTGGGCAAAGCTATTTGGCTTTGTGTTATTGCAAGCTGACTTCTGCTTCCTGAATTGCTCTCTTGCA
AACTTGTTATTTGAGCAGGAAGAGATAAATTGCATGTACAAAGGATAAGATGGTGCAACTCGAAAATGGTCAAAGTATTGCT
CCAACAGCAACAAAAACTCTTCAGAAGAGTTCGGAAAAAACCCAAGTATTTTGGTTGATTCAAAGTTGTAAATGACAAAGAA
AGCTATTTGATTGGATGCATCTGAGTTTTGTAAAACCACGCCATCCACACTACCAAACTTGATGAGCAAGTGGTACCTGTCCA
AAAATTGTACCTTCCACATCACCAAGTCTGCATAATGTTGGAAATGGAAGTAAAATCTTTTGAGATGTTGAGCTTTCGCAACA
GGATCAGAATCTTGACTCTCAACGCTTCGGAAAATGTAAGACAGCATCCGTTGCTTCAGGCCACCCAGCATGTGATCACTTTG
CATATTGGCACCCATGTCATCAAAAGAGTGAGTTTGTGAAGTGGAGGTCTCTTGACTTGGTCCTGCCTGGGAAGTCATCTCAT
GATTTGAACCTGTTGAACTTGACCTCGTATTCTCACCGAAAATTGTGTTCTTGTGCCCATCAATATGAGTATCACCACAGTTGC
TTCTGCTTGAAGGACCTCGTAATGCCCTGGAACACCCTTGTGCATGGGAGCTAATTATCAGTTCATCATCTTCTCTGCAGAATG
GTCCAATAGTTTGAACATCAACAAACATCCCTGCCCTTCGCACTTGCAGTATGTGAATGGACTGATAGCGCACAGATAGAATA
GAAAGGAAGTCGTCATACATGAATATGCCAGCATTATGGGCCAAGTGGATGAAATCATCGTAGAACTTCCTTTCATCCATAAT
CGTACCATCTGCTAGCCTCACAAGAGACAGCGTGATTTTCTTCATGGAAGGGATACCAGGAATAGCTCCAGGCCTTGCAGGA
GCATCAGACTCGAGAGCAGTTGCAGTGGCAAATATACCATAACAATTGCAATCTGTTGCCAGGAAAAAATCTTTACATATGA
GCTCGTTTCCAGATGCAAGACTTAAAGAATACAACTGGGAGAAATGGCTCTCAAACTTGTTTCCCTTGATGGGGAATTCTTCC
TCTATATCACAGTTTACTCCTTTACTGCAGTACGTCAAGCAAATGTGTCGATAAACAACCAGGTTCTGATAATTCCGACTGAA
ACAGATCAAAAACTGTCCATCATTTGAAAATTTTCGGAAATGATAATCTGGGCACTCCACGTCATACACTGTATAATTTGGAA
CAATGTTTTCATACAGCACTCTTGCTCTGTGAATGTGAGTATTGGGTGCAGCAGCAGAGATCTGGCGGTCGAAAAGCCGGTGC
AAAATGTTCTCGCTTTTCCTCCACATCCTTATACTGTCGTATCTCGTGGCCACAGCTGCTTCTCTGTGACAACACGCTTGTGGA
CGCCTTCGTCCATTGTTAAACTGCTGAACTTCGAGTGACTCCTAGTTCTCCATAAAACTTATGTAC

**CULLIN 4 (*CUL4*):**

TTTTTACGAAATTTGGATCTAATTTAGCACAAAAATCTTCAGAAAGTAGTCTGGAAGTCCAACTTGCTAAGAAATATG
GAAGTAACAGGTGATACAAGCTTTGGAGTCATGCCTTGACTACACTGAAAGAAATCACCGTACACATGGAAGACATTTTCTA
GATGATGACGCATTGTGCTTCTGAAGGAGCTTTAGAATTTACACTTTTCAATTCCCTTGCACTTTGCTGATGATTGTGGTTTTCT
AGAAGGCTGGCCAAAGATGGATTAATAGATCTCTTTGGCTGTTCCACGCGGTTGGCTAAAAAGCTGTACAATATACAATAGT
GCAATGGAGTACTGAAGTTCGTAAAATGCTATGCTCCAGCAAGTCATCATGCTGATTGGTATCATTTACAATTAGAAACACTG
CTAAGAAATAAAAGAGTATCTAGATCCCTGTTGAGACATCACATAAATACTTGTTATGTATGAACCCACATTTAGGTATAAAA
AGGTCAAAAAAGCTAAGGGCAATTTGTCAGCATCAGAATACTGAGATCATACAAGAGAAAGTGCATTTACAACTCATGGAGC
AAATTTCCTTGCAAAGCATTGGCATATACGACGCGTTTGCTCTCTGAGGATAAACATCTGATAATCTGTTATGCCTTGCCAGTA
GAAACCTTATAATCCAATGTTCTCCACAATTGAGACTAACCATGATATAGTCAACGGCCTGGTCCCAACAAGCTGTAAGTCAA
GCAAGGTAATTGTAGATTTGGGGATTGCTTTTGTCCCTCTCAAGATACTCTCTGTCGATCAAACTTTCTATTCTTTTCTTCAGAT
CAGCTGGCTTTATGGGAAATTTGAGCTGTTGGAAGAGCTCAGTTATTAACAGTGTATGGCTAAGAATCTTTCTTGTCTTCATGA
TCCTTACAATAGCAGCATCTATCTGATATTGCCTATCCTGAAAGACTCTTTCAGTTGTGCTGGTATTTTCCTCCACAGTCTCCTT
TAACTGAATTGCGTTGACTTTGATCCGGAAGAGAGGGGCAACAAAGTCCACATTGAAAGAAAAGCAGTCATCATCTTCAACA
TCTCTTCCTTTGGGTTCCTTAGAGAGCACTCTCACTTTTCCACAAGCCAATGACTGAAGAGTTCTACGGAGCTCTTTGTCCTCG
ATGGCAGTGGTATCCTTGATTTCTTGAAAGGTGAGCCTGTCTGAACTGTTGAAAAGCATCAGCACAACCGTCTGAAATAAGGA
AACAGAAAGCTCCTTTCTGCCTTTTGGATAATCGGCTTTCAAAACACAGTGGCCAAGTGAGTTTTGCCACATCAACCTCCGGC
CACTATGCTTGCTGAGGTAGAAATCCTTGAAAATGTCTTGATAAACATTCAGCTCATGAGGTAGCCGGACTTCCATAGGCGGG
TAAGTAGGCCAATACCCTGTTGTCAATACATGCACACTCATTTCAATGCCTGAAGGCAGCTTCGTTCTGGCCTGTGAAGACTG
TTTAAAAGACTCGTTAATCTCCTTAGAAAGCTCTATGTCCTTAAACATGCCTTCCAGTTTATTCGTAAATTGGCTTCCACATTC
TGTTTTGAGCTTTGAAATCATTGATTTCTCTGCGTCGATTGATGCACTTTTTCCCAGAAGAAGTCTCTTTGCAAGGTCCTTTTTA
TAGAAAGCTTCAAAAACATCCTTTCCCTGTATAAACCGAAACAAAATCAGCACTCTGTCCAGAGTGTTTTCCAGCTCCTCTTC
AGATGTGCCTTTGTTGCCGGCCCTCAGCTTACCATCTATAAATTTTGCAATTAACTCTGCTGGGCGATTCTGCCGCAAGTTGAT
AAGGTGCTCAAAAGCATCCTTTATAGCATTTGCAAATAGCTCATTTTCAAAAAAGCTTTTTTCCCACACCACATCAAGTCGAG
CTTTGAAGTCCAAAAGCCACAATACCATGTCTTTGTCTTTCTCATCATCCATAACAACATTGTGTCCAGAGGCCTTGATGTAGA
CACTAAGTGACTGCTTCAGCTGTTCCAATGCGCCCACTTTTGAGAACAAACTGTACATCCTCTGAAGGTCTTTGACACGATTC
GCATCCATCAGCATGCTGAAACCCTTGTCTAAAACGGCTGAAGTGTGTCTCTCAAGCAGCTGCTTTTCTGCAGCTGCAACCAA
GGGCTTGCGGGTGCTTGCATCCAAGTAAAGAAGGCAGCGATCATGCTCTTCATGTAACCTGACCTCAACATGCTTGAGGTAAT
CAGGCACATCAGTTTGCTGCATAAATCTAGTGCCTTCTGCCCCATAGAAGTCTGTCGTACAATTAAGGAAAGGCTTTTCAAAA
CTTTCCACATAAACACCTAAAGCACTGAACATGCGAAGCAGGTGTTTCAGAAGAGTTCGGTCCACTGTCTCTCCCATCCGTTC
TTTTTCTATCAACCTCAATAAACCAGTTACAGTTTTATGTTCTACTTCTGGACATGATCCCAAGTGTTTGCGGAATAGCTGCAA
CCCCATATCCCATAAAGACCTTGCATTTGAGGTCTGTATGACATATGTTCGGTCCAAATACAGCGCTATGCTACGAATCATCA
ACATCTGGTCACAGTGTTCCTGCCAGCACCTCTCCACCAGTGACAGGAAGACTACTGCATCTGGGCTCTGGCCTACCAGAGAC
TGCAATTTGGCTGCAATATGCATCTCACATTCTTCTTGTAGACGACTGTAAAGATTGCCTGCCATTTTGTGCAAGCATAAGTCC
TCCACAGCCCGATACAATTCCTCAAGGCTGCAGGAAACTGGTTGCTTTAAATGGATAGCAGTGACAGCCTCCTTTATTTTGGC
CCAAGTAAGCTCCTCAAAGTTGGTGGGAAGTTTTGGCTTGTCTTTGAAGGGCTTGATGACTAATCGCCTGGGCTGAGGAGGGG
TAGCTTTCTTCCGGGACAAGTTTGCCGCAGTACCGGCAGCGATATGGGATCCAGGGAGGCCCTCATCCTTGATATCCATGAGC
ATATCATCAGGGCCGTGGTCACCTGTGGAGTTGAAGAATCCCAAGCCGTTGGATCTAGGGTCCAGGTTCTCCTGTTTTGGCTT
CTTGCTACTAGAGATCATGCCGATGCCCTCACTGCTCCCGTTCACACAGCTGCTACTTGTTTGGCTTGCCGGCCGCTTTGAATG
CGACATCAACCGCTCATGCATTGCACCAACACCTGGCCCTAGACTGCATACCCACCGCCGCTCGCTGTTGCATCTCTCTCTTTC
TCTCTCTGAGGCGAGCGTACCCTTCTCTGTAAGCGTACGCTCAGTTTAACCCATGGAATTCCGACACCTTAGCAGAATCCAGT
AAACTATGTCTGGTAAGGCAAATTGATGGAAACAATAACACAGCTCGACGATGGTGAGACCAAGCTCGCTGCTAAGAAAAAC
TCAGCAGCAATCCCTCTGTATGTTAAAATG

**ELONGATED HYPOCOTYL 5 (*HY5*):**

TCTCTCTCTCTCTCTCTCTCACATCACTGTTCAGATACGTATAGCAGAAAAGCCCTCGCCTTTGTCAAGGTTTTGCACA
GCTTGAACTGTAAGTTTCCATGGCGGCTTCGGAGCTTCGTCATTCCTCTTCTGTCTCGTAGTTTAGGGTTAACTTGTTTCTTTGT
CTTCTTCTTTTCTTTTTACGTCACATGATCCGGCAATGTCGGTTAAGAGCTTCGGTTTCCTGTCTCTGGAGTAAAGCTATGGCG
AAATTCGAGCCGTCGATAGATCTTTTTTTCTTTAACTCCGTCGTTTGCTGAATCAGTCGTGTTTCGGATATATTGTATTGGTTGC
TCTGCTCTGGAGCCGTAGCGTCTTCTGCGTGCACATCCTTGCTCATTTCTGCAAATCTCCATCGCCGAATTTCCTGTTTCTCC

AGGCTAGCACAAGCGTTCTTTGAAGCAGGCTGTTTCAAATATGTAAGCAGAGCTGCTAGTGATGGTGGACTCAAGCTCTTGGT
TTAGGACCATGGCGGATTGAAAGCTCTCACAGGAATAGGTCTCAGGGGTAGTAGAGAGAGTGCAGTGGCCTGTGCAGCTTTT
GGAATACGAGAATGCAGGCTTCAACGTCAGCCATGAGCTCCCCCCATGGTAGTGAGGAAAAGCCGGCTCCCTTCGAAATCAT
GGCAGCTTCCATGGATCAGCTCAGCAAAGACGGTATGCATGCCTTTTATTCTGTAGAATCGATTCTCTGTATGAGAGAGATCT
TCTCCTTAGAGTGTGTGTGAAGAGAAAGTGTGTGTGTACACGTATTGTCTCCTGTGTTGTGAAAGATGTGCAATTTTGGCGTTT
TCAGCGTAACTTCTTGTTCGGACACGCTCTGTGCGTAAGTAGTTGTTGTTGTGTGAATATTGTCTCTACTCTCTGCATTGTGAG
CGTGCACGTGAATGCACACATGTTTGTGTGGTCCACTGTGTGTGCTTTGTGTTCGATTGTGTGTAGTGCATACCTCTTTGTAAA
AATCTCAGCGTGAGTGCATCTGTGTATTGCGAATGCTTTTTTGCGTTTGGATTATCACAACCCGTGCAGTCTTTTCCAGTGAGA
AGTTATCTTTTGTTTTAGTCATTTCAGTTGTCTGGTGCTGACAAACATGAATCTACACCAATTTGCCAAGGTATTATCTTCCCTT
CTCCCGCCAGTTCGTCTCATTGTGATTTGGGAAATATGTGCTGCAGATATCTTTTGAATGTGTGTGTACACGGGCATTTGTCTT
TGTTTGCATCAATGGGTACACAGAACCCACTCTGCATATGGGGACGGATGTCTGAGCAAGCATTGGTGTGCACAATTATTGAG
TTCACATGTCTAACACAAAGTTAAGTGCTGGTAATGATGTGTATGAACAAGTTAGTGAAGTAATGGGGTAGAAGCTAGTCTTG
GAAGCCCAAGTCAAGTGATGGTTTTGATGTTTAAAGTGTGTATACATACAGGTCTAATTCACGTGGTAAACCCTTTTCATAAT
GTGCTTTGTTGTGTCAGGCGATGATAGTGATTCGGATGTGAGGAAGGTGCCAGAGATTGTTGGGAGGGCTTCAGCATCAGTA
GGTGGATCTGGGAAGCATCAAGGAGGCAGGAAGAGGAATGCTTTACCTGCCGAGAAAGAGCACAAGAGGATGAAGAGATTA
CTGAGGAATAGAGTGTCTGCACAACAAGCAAGAGAGAGGAAGAAAGCCTACATGAGTGACTTGGAAACCAGAACAAAGGAG
CTTGAGCAGAGAAATGCTGAGCTGGAGGAGAAGGTGTCTACTTTGCAACAAGAGAACTCCATGCTACGGAAGATAATCAAGA
GCACGGCTATCAGAACTGGCTCGGCCGGTGACAAATGAGGAAACAAACAATTTTAACTTAGGACATACTGTTTTATTTAGAGT
CCAGTTATACCCTCGTCAAAGACCATAATGTGGCTTATAGCCCATTCCCTCAATGGATAATTGCCTCTTATTTTACCTTTTTTA
GGGTGTGATCCCTTAATCTTCAGGCCTTTTTCCAAGGGCCCTAGGACCAAAACATAGTTTCATGGAATATCGGCTTTCCATCCT
TACACTTATTTTTGAGTTTCTGGGAAAGCAGACTGGCTAGGCATGTTTCATGGAAAATATTGGCTTACACTTCTTTTGAGTTTC
TGGGAAAGCATACTGGCTAGAGATGTGGATAAACTATTTGCACATCCTTGTAAATGTAGTTTCTCAAAATTGCACAAAGCTTA
TTGCTTCTTTCCTTCTCTGCTTTGGATAAATAGTTTTCTCTCCTTTGTCTTGGGA

**XAP5 CIRCADIAN TIMEKEEPER (*XCT*):**

GGAGGAGCAGAGTGGGCGAGGGCAAGATGTCGGGCATGGGCGATGGATATGTAGGCACAGCGCAGGATGCAGCTCG
CATCCGGCGGCTGGAGAAGCAGCGCGAGGTGGAGCGCAAGAGGATCCAGGAGCTCAAGAACAAGTCCTCTGGTGGCCAGAC
TGGCCTCCTTCAGTTTGGCTCCGGCACCTCTGAGCTTCTAGAGACTGCTTTCAAAAAAGAAACTGTGGGACTCGTCACAAGGG
AGCAGTATGTTGAGAAGCGTGTTAACTTGCGGAGCAAAATCGAGGAGGAGGAGAAAGAAAAGCAACAAAAGCTTGCGCAAG
AGTGTACAAGCTGCAATTCAATATTTGTAAGTTTGTATGTATTCAATGGCATCTTTTGTGTTTGCACAAGCACAAGCTTACATA
CACACATGATTATCTTTTGATGAATGGCCTGCTGGACGTACGCCTTGATGGATTTTTGCATATAATAGATGCACCTGAATGCA
CAGAAAGATACCCATATAGGTAAACATAACTTATGTGCAGGTTGATGCGTAATTTATGTGTGTGGCGAGTGTGCGGCATGCAT
AGAATGTTTTGTGATGTTGTTGGAGATTACACTCATTTTTATCTTTCAGAAAATGAGGAACCCAAAAGGGGAAAACTCGGTAA
GTTTGGTAAGGATCCAGCTGTTGAGACTAGCTTTTTGCCAGACAGGGAGCGCGAAGCAGAAGAGCAAGCTGAGCGTGAACGA
TTGCGGAAACAATGGCAGAGGGAGCAAGAGCGCATCAAAAATGAGCCACTAGAAATTACTTACAGTTATTGGGATGGAGCT
GGTCATCGGCGTGTAATTCAGGTTCGTAAGGGAGATGCAATTGCAGAGTTCCTCCGGGCGGTCCAACAGCAACTAGCCTCAG
AGTTCCGGGAAATCAGAACGGCTTCAGTGGAAAATTTATTATATATCAAGGAAGATTTAATAATTCCACATCAACATTCGTTT
TATGAGCTGATAATCAATAAAGCAAGGGGCAAGAGTGGGCCGCTCTTTCATTTTGATGTGCATGAAGACATCAGAACAATTG
CGGATGCGACTATAGAGAAAGATGAGTCACATGCAGGGAAGGTTGTAGAAAGGCATTGGTACGAGAAGAACAAACATATTT
TTCCAGCATCCAGATGGGAGATTTATGATCCCACTAAGAAATGGGAGCGTTACACAATACACGGTGACTGAGGCACAATCAA
AGATTAGAAGCACACAAATCAGCAAGATGTTTTAGGGATGACCTCTTTCTAAACTTGTTATGTCAGAGATTTAACATCGTGTT
ACATGAGAACCTCAGCATTCTAACAAGAAATTTTGTGATGTATGTGGAAACATTTGGGATCATGAATTATTGGCGGTACATCT
TGTACAGTTTTAGAAGTAATTTTGTTTTGATTAAAAAAAA

**SPLICEOSOMAL TIMEKEEPER LOCUS1 (*STIPL1*):**

GACACACTCGATATTTGAATATCGCATCACTTCAATGCGATTAAAAAACGATTATAATACTACAGCGTGATATTTGAG
TAAGCCTTTGAATCCTACCAAGTGAACAGGTTTTACATCCCACCCGGTTTCCAGTAATAAGTTGCCGACATGTCTAAAATTGC
AGAAGCACAGCATCAATACTACAATAACCTCAATTAAGGAACAGTGCAATCCATTACCTACAAAACTATGAGTGGGATTCAA
CATTTCTCTGCTCAACCTTAGAAGAGATACAAAAGAACACAAGCATCGTCCGTCGCACTTCCAAGGCAGAACCCCCCAAAAA
CCAAATAAAACAATCAGAACAACAAACCTTTTTAAACCTGCCCTGATTTTGAGCTAATATACCCAAAAATACACATGAAGAT
GGATCAAAACGCTTCTCTGTTCACCACATTCGATAAGGACTCCGAACACACTCCAAAGTATCCTTTGTTGCTTGTTCAAGAAA
CGAAACAGCAGTAACATAAGAAAAGGTAACAAAGACAGTCTGAGCATGTCAATTCATAATTGAATTTGACATATGACATTGT
ATTGCATTCTTTGCAGACCACATCTCAGCATAAATAGGACCACTGTTCCTTGATAATGTACAAGCTTGGCAGCCTACTTCCATC
GGCTAGACTGAGCCCGATGCATTTCTAGTAACTGTTCCAGTGAAACTGCCAACCATCTGTCCCCTGATTGCGCATACAACCTT
TGCTGTGCATTGTCCATGCAAACATTTGTAGATCCAAAACCATAGACTTGAAAACCTTCATGTAAGCGCCCAATTTTGGGCAG
AAATTGCACATCGTTCTGCTCGGCAAATGCCTCTACAACCTCTTTCAGACTCATTTCCACCGAGCCATTATCGCGTGCAACATT
ACCAAAGACGGAAGAATGCTGCTGCGTGTATGCAGCTGCGGCTGCTGCTTGCTGTTGATTCTCAAACTGTCTTTTTTCTGTGAC
ACGTAGATAGCTTACATTCTCCCTTGCCCCCGGTTGTACAACAGGCATACCCTCCACAGCTTGATCCATCATGTCAAGAGCAC
CTTGCAGATGCCACCTCACCCTTTCATTTGCCAGCAATTCAGGAGTAAGTAGGCTTTTCCAACCCAGATACCATTGTGTTACTT
CTTCAAAATCGGGTGTTGAGCAAAGCCAATTATAAAGGACTTGGCGCCACTTCATAAAGAAACCTGCTTCCAATAAGGTAAC
CATATGATGTATGGGAACTGCCGCACCCCACGCCATCACCCAGCGAAAAGGCTCAAGCTGCTGTGCTTGAGGGTTTATGATA
AGTTCTTGCAAGGCAACCATGAGTTTGGGAAGAATAGACTTCACTAACAGAAGCTCCCAGCTGCCTGGATCAAAAACAGTTT
GCCAAGGGGATAATAATGCGTACGCTGAACTATCACTGGCATGCCACGCTCGTAATGCACTCTCCAGCTTGTAGCGAATAGG
ATGGTAAAGAGGCTCCATTCGTTGCCCCAGTAAAGGCAACCATGGATGAAGCCATGCATGAATAGGAACTGTTTCTTGCCGA
GGATCCCATTGATCCACTGCTAGGCTAAGCTTTGGCATTATTAAATGCTCTAGAATGCTGTGAAGCACAGATTTTGGAAGCAA
ATTCTCCCATATCTCCAGAAATCTTAACATGGGTTCAGGATCACGAGGTTCCCAAATGTTAACCGCTGCAAACCGAAGAGCAG
GAAACACAGCCTCCATTACCAACTGGGCATATGGCGAATCCCCATAGCTGTTTGCATCTGGAAAGATTGAATAATCTATGGGA
TCGTCTCCTTGAAGCAAGCATTTCCAGAGACCTAATATGTCTGTCCCATGAAGAGGTTGAAGCAATGGTTCCCAGCCATGAAA
CACTTTTACCATTAAAGGGAGCCCGAATGATAAAGCTATGGCTGAAAGATTATGCAGCTTGTATTCTTCTGCATGATCTCTCT
GCAAGGCGCTGAACGCACCTGCAACTGCATCTAAATTCATACCTCCACTCGCAACCTTTTGCTGCACATCCCCAATTGCCCTC
GTAATAAGTTCCAGGGTACCTATCTGCTTCTTTTGCCTATCAGCTTCACCTTGTAGCCGCACCCGCTCCTTTTCAAGAAGCACC
GCTGTGTCTTTCTCCTGCCTTAACTTCTTATCAAACGTCTGTATATCAGCCTCCACAAGATCCACAATCAATCTAACATTATGC
TGAAGCTCTGGCATGGGGATATCATCTTCAATATCTGTTTGCTCAGCATTCAAATGCTCGAGATTCGTGAGAACCCGGACCTG
CGGCCCTCTCATATCCAAAATGGTTTGTACAGGCTCAATACCCTGCTCACTCTTCAGAACAAGCAAATCATCTGCTGTTTGGA
AATCCTTCTTCTTGCCTTTAAATTTTTTGAGCCAGAGTTTCTCTTTAGGCTTAGTTTCCTTGTTTTTAGCTTCTGCTTCTTCAGGT
CCTGGTAGAGCAGAGACCGCCGCATGCACCTCTCTGTAGTCATTGAAGCCCATACCCATGTTCTTGGGTCGCAATTTTGCTTC

AATTGGGACAGCTATACCCTGCTCATTTTTACCAAGCCCCCCACCTTTGTAGCCCATTTTTTCAAGAAGCCTCATACCTATACC
TTTTGTGTGCTTCTCAAACTCAGGCTTATCAGCAGCTGCCGTACTTGCAGAGAGACCCCCCTTGCTCTTCGACGTGGTCGAAAG
CCTCACCTTTTCTTTCTCTCTTTCTTTCTCACGCCTCTCTGCGCCTTCCTGGATGCGCTTCCCAAAAGCTGAAGGAAAAAGGGTC
TCTTCTTCATCTGCTTGCTCGGGCACAGAACCTGCAAACCCTAATCCAGCAGATCCCAAGCCCGCGTTGGGAGGTTGATCTGC
AACATCCTCTTCTACTTTATCTATTTCCTCAGAGGGTTTAACAGTACCGGTAGAGACAAAACTCACAAACTTGGAAAGGTCGC
CTTTCTTGATGGTGTCGCCACGGCGCCTGCGTTTGGAGGAGGATTCATCAGCATCAGAGTCGCTTTCGTCGAAGACACCATAT
AGAGCTTCCTCACGGGTCTGCGAGCGCATCTCCCTGCGCTTCTTGTTGTAGTAGAATTCGCCCCCGATCCATTGCCCACCCGTG
TAGTCATTCTCCATGTTGAAACGCTCCATGTGCTGGTACTCATCCATGGCTACGGCACTCTGTGTGCAGCTTTGAGAAAGCTTC
TGTGCAGCGTCGGTTTACGTGATTGCTATGTCGGCTGGTTATGGCGAGTGAGCTATCGACTGTGGTGGCATAAAATTCTCTCG
CGAGGGAGATGGACAGAATCAAGGTT

**ELF4-LIKE 4 (*ELF4-L4*):**

AACCAATCCAAGTTTACTCAAATCCATTCTGCAAATACATGATGGAAACATTTTTTCAGGTAAAAGAAGGGTCCCACT
GTGCAAAACACGGTGCACACGGCCCAAAGACTTGCAAGAGAAACAAGAGAAGATCAGCTGAAGCTCCACAAGCATTCACTC
CTCCTCACATGTTTTACTGTATTTATCACATGCATTTGAGGATGAAAACATGACCGCTTTTGGGAGACGCTGTTGCTGTTGCGC
ACAGATGCAAAGCAAGTGGCATTTTCCTCACTGGACGACAATGATGAAAAGTATGGGATTTGCCTCCCTCTCTCAGTCTCCCT
CTAGAGCTGCAAAATGAAGCGCCAGATCCTTGACCTTGCTCAAGAGACACCAAAATTGAGCACATGTAACTGGTCAAGTAAG
CCGGGGCTTCTTCTGCGCCTCATGCGAATGACTGTTAAAACCGCCTCTATCCATCTTCACCATGCGATTTGCTTCCCCTTTTCCT
GCAGATTCCTCCATTATTGATTGTACAAACTCTGTTGATAGGTTTGCATAGACCTTCACAACACGGGCAATATTAGCATTGAG
CTCCCTTATAAGTGAGACATTCCGTGCGAGGTTCTCCGGTACTTTACTTTCTTGATTCTGGTTGATCTCTATGATAAGGATGCG
ATTCTGATCAAGGATTGTTTGCACCTGGGAGAAGCTCTTCTGAAACATCTCCCAAACCTTGCAATCCATGTTATTCCCATCTTT
CGAACATTCTGTATTATTGCCCCCATTCATCAGGGCGCAAAAACAGAAAGCTTCGATAAGTGATATGTATCTGCCTTTGATGA
GTGGAAGTTTTCCAAAAAGCAGCTAAGCGCACATTAGCTGCAGCTACCGAGAAGAGGAATGTATGGACTTCATTTGGACGTG
TCGTAACAAAAAGCTTTAACAAGCTGTTTGATGTGTTTAAGCTTCACAACTGTGGCTAAACCGACGAACATGTGCGTTGTATG
TGTAATTCGCATGTGTGGGGATTACAGAAATTAGCTTTATTACTCTTTGAGGCTTGTCAAAATTGTTGTGTGTCCGCGCACAAT
CGACAAAAGAGTGTCAAGAGCTATTGTGCAGATACTACTGCCTGTGAGAACTCCCAGGGGCCAAAGTGGGGTACTGCGATGG
CAAGAAGCGCTGTCGGTGTAACAGAGAAAACTAAAGAGAGAGGG

**MORE AXILLARY BRANCHES 2 (*MAX2*):**

GAGTGTGTGTATGAGAGACACTATTTGCAGCTATTCATTGAACTTAAACCGATGAACCCAGATCAACAATTGGGCTTC
CTGACCATTTCTTACAGCAAGCTCTTCAACAATTGGCATACACCAGCAAATCAACAATTGGGCTTCCCTTCTCCAAGCTCTGTA
GAACATGACCATTGCTATTCTGCCTGGAATTGTCCAGGTCAGCGCTCACTCATAGATTGTGGGAGCAAGAGGAAGAACTACA
AGGCAGGAAGTCTAAGAGCTTGAAAAGACACTAAATGGTAGCATTTTGCCACAAAGGGTAGGAATGGAGCCTCCGGCTCTTG
AAGTCCCTCCTTCTCCCCCCCGATGGTCTCGCATTCCTTTTGACATATTTGAAAATGTGCAAGGAAGAGAGAAGAGGGTTCGA
GTTGCGGCACCAATGTCAACGACGTCCATCAAAGATATTCCTGAATCAATCTTGGCCATCATATTTTCTAGCATCTCAGAGAC
TCGCAGCAGAAACTCTTTTTCTCTGGTTTGCATGGAGTGGTACAGCTTGGAGAGGGCAACAAGGACAAAACTTTGTCTCAGGG
GCTGTGTAAGTGATTTGTATCTCATTCCCAGATCATTTCAAGCAGTTGAGTGCCTTGATCTCTCGCATTGCTCTCCGTGGGGAT
ACTCCTTGTTTCAATCCATCCCAGAAAGAGCTGTCCTAGTTGGACAGGTCTTAAAGCAAGCTTTTCCAAGAGTGAAAGACATG
ACGATTTATGTTCGGGATGTGAGAGATATTGATATGGTGGCCTGTTTTTGGCCAGATTTGCAGTCTTTGAAGCTTGTTAGGTGG
CATCAGAGGCCCACTCAACCAGAAGAGGCCAGTGGCCTTGGCATGGAGTTTTGCTTCTTAATGCAGTCATGCAGATTCCTTAA
AAACCTTGACTTGTCTGATTTCTACTGCTGGACTGAGGACATACCTCCTGCTTTGATGGTGGAGCCATCTGTGTCAAGCAAGCT
TCACAGCCTCAATCTTCTTAAAGTCTCAGTAGAGGGTTTCAAGGCCTCTGAATTGGAAGCCATTTCAACTGCTTGCACAAGCT
TGGAGGAACTACGAATAGTCCTTGTGTTTGATCCGCGCTTTTTGGATTTTGTTGATGATGCTGTACTTCTGCACCTTGGAGCTA
GCTGTAAGAAATTGAGAGTGCTACATCTCATAGACAATGCCGCATTCAATGATGCGCGTGGTGATCCGGAGGATGCCTTTGAT
GAGCAGGATGCACAGATTGGAAGGCAGGGCCTTGAAGGTGTTTTTCAAGCTCTTCCTTTGCTTGAAGATCTTGCCATTGTGCT
TTCACAAAATGTGAGGGATGCTAGACCCATTTTGGAGGTACTTGCTGGTTCTTGTAAGAGGTTGACATCCCTATGCCTTGGCC
ATTTTCATGGTTTGTGTAGTGGGCCGCAGCCTAATGGGGTTGCCATGTGTCATGGGCTCACAAACTTAGTCATCAAGAGCTGT
GCAGATCTTGACGACATTGGTCTGGTTGCAATAGGATCTGGTTGCACAGGGCTACGAAAACTAGGCCTTAACTCTTGCAGAGA
AATATCCAGAGCTGGGTTGCGCAGCTTAACAAGACAACTTAGTCACAGTCTTGTGGATGTTCAGGTTATTTGCTGCTTACAAA
TGGATGCAGTGGACGCTCTTTGGGCCTTGCAACCAATCCAAGGCACCATTGTCAACTTGCATGTCGACTTTCTGCTAAATCCA
GATTTTGTGGCAAGGGAGGCAGAAGATGCTAAAAAAAAGCAGCCATTACCTCCTTCCTCCAGTTGTTTAAGTCTCAAGAAGA
CATGTAATGAAAGTGTAGGATCTTCTGGCACAAGGAGCAACAGTTCACAGCTGAGCAGCTCTGGCAGAGATCGTATCAGTGA
TGGACTTCAAAAGAGTCATTCTATTGAGAGTTTGGATCTCAATGTTGCCGTAGTTGGCCAAGAAAGTGACACAGGATCCAACT
CTCTTTGGCCCCTAGCAGACGGGTATGGATTTCCTACACAGAGTAGCAGTAGCTCAGGGGGGCTTGCAAGCGATTGCCTGACT
GCAGCCGAGCAGGACACCAGGGGTTCCACTTCCTTGTGTGATGCCAGTGTGATTACTGATGACCTCTCAAGCCACTATTCAAA
ACGCCTCCCAGGACTTGAAGTAAGCTCTGAGTGTGGCGCTTCTGAATTCTCTCTGGCATTTCGTGAAGGAGAATGGAGCAAGC
TAAGATCTTTGTCTTTATGGATCCCTATAGGTGAGCTGTTTTCTCCATTACCCATCATGGGTCTGCAGTTTTGTCCATCTTTACA
AGAAGCTCGAATCAAGGTTGAAGGTGATTGTCGAGCATGTCCCAAGCCCAAAGTTCGCTTATCTGGTATAAGCACTTTTGCAT
CGTACCCAGCATTGTCAAAGCTCACGTTAGATTGCAGTGAAATAGTAGGGTATGCTCTCAGTGCTCCCCTAGGTCATATGGAA
CTCAGTTTATGGGAGAGATGGTATCTGCAGGGTATAGAACACGTGAAGATCTTAGAGCTTGATTACTGGCCGCCTCAGGACA
GGGACGCGAATAGGAGAGGGCTCTCTTTACCATCCGCTGGCCTTATATCTCAGTGTCCAACATTGAGGAAGCTTATCCTTCAT
GGCACAATTAATGAGCATCTGCTGAGGATGTTTCTAAAGGTTCCAAACCTGCGGGATGTGCAGCTGCGGCTTGATTATTATCC
AGCTCCTGAGGTGGAATTGAACACAGAGACACGTGCAGATGCTTGTAGAAGGTTTGAAGCGTGTCTCGCTGAACGTGGGTTT
CTAGACTAACTATAGGATCTTACAGGAGTCACTAACATCATGACACAACTTATTTTACTAAGGGTAAGGGCCTTCAGCTTACT
TCCTTTTCTGATTAATATGATGGCTGTCTGAAGAAAGTTATTGACAACCATTAGGTCTTGCTAGTGAATTTGACTTGGTTGTAC
AATGTTCATCTTTTGCAGGGACACTTGCAGGTTTATGGTGGGGGGCATCACCATGGACTGTTTCAGCAGATTTTAGGTAGTTTT
TTGGATGGGATCTCAAAATATCTTAACGAGTCCCTATTTTGTATTGTGCATGAGATGTACAAATCCACATTTTTGCATCATGGT
GGTAACAGCTCTCTTGGAGGTCTAAAGCCATTTTGATGTTGTGACTGTACCAGAGACTAACGAGCTTGGGGTCAACATTTCTG
GCTTTTCAAACATGTTCGCAGTATTAGCCATATATA

**ACCELERATED CELL DEATH 5 (*ACD5*):**

AAAAAGGCTGCAGGTGGACTCATTGATAGAAGCAGCTGCACTTGTTGTCTCTGCGGCCTTTTTGTTCAAGAGAGGGCG
TCGATCTGCAGAGGCTGGTAACAATGTTACTGCACTATTGCTTACAGATTTTCCATGATTTTGAGGTTTTGGTGGGAGTTTTGC
CCTATGACGCCGCAGCAATAGGCCATTTAGGACTTCATTGAAGAGGCCATCACCACCCACAATAATAACTCCATCTAGTCTTG
CCAAGTCATCATCACTAGTGCTGTTTATGACCTCAAATGCATGTCTGGTCCTCTGTGTCATTAATACCTTTGTGTTAATTTTTGC
CCTTTTGAAGAGAGAGATGAGACTTCTTTCCAAGTATTTGAAGCATTTTTCTTGCCACTATGTGGATTGATTATTACCATGAGATT
TCTGGGTCGTTTGGTGTCTTTGTCTAATAAGGACTGTATTTGATTCATCCATCCCTCGCAAACACCTTGATCAGTATGATAAAA

GACAAATGTTTGAGGTATCCATGCTCCCTTCTGGCTGGAAGACTTTTGATAAGCATGCACTACAAAGCGATGCACCTCTTTTA
AAATTTCTCCAGAAAATGTCTCTGGTGATTGTCCTCCTACTGTATCAGCAATAATCGGTTCGCTAGATTCAACAGCATATATTT
GTGACATTGCCAACTGATTAACCTCATCAATGTTCTTAATGGGGACCCAGCAACATACGGTGTCAATATCCAGCTTATCTTCA
AGAGCCTGCCAAGCCAGACAATTGGATGATAGAATAACAGAGACTGGGCCCAGAATGTCCAAATGAAAATTGGCTTCCAGCA
GGATATCTTCAGAGTTTGCAGCAAAGTTTGCAGCCCCATCAAAGCGCCTAAACTTTTGAAATTTACCCTGACCATCCCACAAA
CTAGCACCAAACCTCAATCTCTTTGTACCATCTTCTATGTGCGATGACCCAACCGACACATCCACTTGCTCCCCCATCCAGAAA
TTTCTCTTTGTGAGAATCTGGACTGCTTGTCTGCTAATGCAAAAGCTGCTGGGAGCAGACACGCGCATGGAAAACCACTTTTA
CATAAGAGCGAGAGTGCACGCGTGTGTGTGCGCGCGTGTATTGTTTTGGGCGTTTCAACCTTTGAAAATTTCTACATGAACGT
TGATTTCGTG

**AUTOPHAGY 5 (*APG5*):**

AAAAAATCTCTCGAACAGCGCAACTCCAAGGTGATCAGCAGAGTCGATGATGCAAATTGTCTCTTATTTGAGCTTTTA
CTTCAAAACAGTCTTTGCTGCTCTATTCTAAACTACAGGATTGCTGAAATGTGATAAATGAGTTCCATGTAACAATCTGGCGG
GACAAAAGATATGGTTATTGAAGAACGCAAGATTGTATGGAAGGGTGGGATACCTTTGCAGCTGCAGCTCCATAACTCAGAG
GTTACAACAGTCCCGCCACCACCTCCTTCATTGATACTTGCACCACGTAATGGATATCTACCACTTCTTGTACCAGTTTTGAAG
CCACAATTTCAAAGTGCTTTGCCAATTGGACAAGATACAGGCTGGTTTGAGTATGAAGGCCTTCCTTTAAAATGGCATGTTCC
TACTGGCGTGCTTTTTGATCTTTTGTGCTTTGAACCAATCAGGCCCTGGAACCTTACGGTGCACTTTAGAGGTTACCCGTCTGA
ATTGCTCCCTTACGAAGGAGAAGATGTAATGAAATGGAGTTTCATCAATTCCCTAAAAGAGGCATCATATGTGATGTATGGA
AGCACAAAGCATGTAATGAATCTATCTCAGCCCGACCAATTGGATCTCTGGCGATCTGTTGTTCAAGGTGACCTTGAAAGTTA
TGATCGCATCTGTTTCAGATTGGCACCTAGTTTGGGAACGAACTCGAGTGCCGCAAAAAATTTGGGTGATTCAGAAGGTTCCT
CTAGTAGACGACAACAAGGTAATCGATCATGTGGAGGGCCAAATCATGGCTGCGGCTTGTATTTGGATGTTCTGAATCAACTT
GGGAACTTCTTTGGGCAGCGGCGGAGATGGCGGCTCTCAAAATACCGTTCCGCCTTTATGTAAGAACAATAGAAGTGAGCGA
TGGGGATTTGCCGGAGGGATCCCCACTAAAGAGCTGGGATGACATTACATACATATCAAGACCAGTAGATGTATGGAGAGAA
GATGGTGTAGTACTCACACTTTGGGAAGCCCTGCAAAAAATTACACCACAGCTGTTTGCATATGCATCGGAGCTGCCTAGTAG
AGACCAAAGCCTGCCTGCCCCTTTAATCTGGGAAAGTGACTCAATCAGCACTACAGCAAACTCAGAAAATGATTACTCAGGT
TATCCAAACATCAAACCAATAAATTCTTCTCGTAGGCGGCCACAAGCTCATTATTTGAACGGAATCATTCGTATACAAGGTAT
AGAGCCTAGTCCAGATCTGCCTATAGATTGGATAGCTCAGAATCTGAGCGGACCAGATCATTTTGTACATGTTTGTATACTGA
TAGCAACTAATGAAAACAAAAAGAGTACATACTAAGGAATTAACATACATGAAGCCCATCTTCACTTGGAGAAAGAAGCTTC
AAGGTGGAAGAGGCAAAGTTTTGATTGCGGGGTTTAAAGTGCAAGCAAGTATTTAGTACATTTCAGAGTCTGCCTAGAGGGT
ATGCACAATGGAATTTCTGAAGAATTCCGCATCAATTAAGAAGCTGCATAGAGAATGGGCAAGAGAGGTACGCCTCTAAAAT
ACATTTGCAAGATGCAACATTGGGTTTGCATCTGAGCAAGAACTTAATGTAAAGCTCATAATTAGAAAGCAGCGCACTTTCAA
TAATAGGTGTTCCTTTTGCTTGTAAACCTTCTTGACCTTGTAACAATGGTAGCTGTGATCAATACAATGAACATGTCTTCTCCA
ACAAAATGTCATTGCAATTCTACTTTCTGGTAGAAGCGGTGCAAATCGGAGTATCGGCATGGGTTACGACAACTCTGATTTTA
GATTTACTACAGTAGCTCCATTGCGCTTAGTAGCTTGCCATTGTTGTAAGTAGCCTAGTAAACAGTTGAATGATAGTCAAGGC
TGCAATTATCTTGAAGCTTATAGACCTTTTCTTGGAATTTCAAATCACTCTGAATAAGCCATGTATGAATTGAATCTAGAGAG
GTAATGGAAGTAGCAAGCTTTTATCTTAAAA

**AUTOPHAGY-RELATED 11 (*ATG11*):**

GCCAAAACGAGCAGAGGAAGCACGACAGTCCAGAATAGAGTCTCTTAATATCAGAACGCGCTCAGTATCGCAAGATA
GAAGCTCAACTCTAATCAAGCGTAAGAGAGGTTTTTTCTGCATGTTCTTTCTTGGTTGCAGTACAATGGGAAATGTAGTACAC
TAGTTTAAAAGATGAGGTAACATGCTGCACATGTTGGAGAGAGAAGTGGACTTAGTTGAAAATTATGATGTCCTTGTGAAGC
TAGTGTGTATAACATCACATCATGGATGGAATTGTAATACACCTTGTTGTCACTTACCCTTTCTCAGTAGTCTTTTTGAGTTTG
ATTTCTACTAACATCAGAGTTTTTGCGCTGCAGTGAGGATGTGAATACTGTAAAGAAGCTGATTGATGAATGTGTGAGCAATA
ACCTTGCTTTATCAATGCGACCACATGATGCCGTGTCTGCATTTGGCCCAATGTATCATGTGCATGTAAAT

**TAPETUM DETERMINANT 1 (*TPD1*):**

AGGGGGGGGGGGAGGGGGAGGGGGATAGGCAAATACTGGAGAGGACTAATTGCTGCTAATGGGGCCATGAGTATGT
GTTTGTAGTGAAGTGTGAGCTTTGTATCATCTAGGCACCTAGATACAACATCTCAAAAGAACAGAAGGCTAGAACTAGATGC
GACATTGGAGAGGCAAATTCGGTGACAAAATTGGACCCAAAAAAAGAACTCGCAGGAAGCAAGCAGTTTTTTGGGAAAGGG
GAGGATAAACAGAGTTCGAGGGAGGAAAAGACAAGAAAGATTTCAGCAGAACTTAGCAGATTTAAGCTTCAAGGGGTACAT
GAAGGAGTTAGCATATTGAAAACGGATGATGTCCCCATATTTGAGGGGCTTTCCATTGTTAACAAGGCAGTCATCAAAGGCA
ATGCGTCTAAAGAGCGCGGGGTTCACTAAGGGCGCTGAGGCAAACCAGCCACATGAGAGGTGTATTTCTGATGGCGCACATC
CTGCTAAGCATGTGTTCACTATCTGAACCACAAACTGTGGGATCCCCTCTGATCCATCGAGCCCCTGCGATATGCTTATATCAC
TAGCTGAACACCCATTTAATCTTCGCCGCTTAGAGTACTTATGGGGAGGAGTTGTTGCAATGTCGGCATTCAGAGTGTTGTAA
CTGCGCCGCTGATCACAAGATTCTGTATTTTGGGTAAGTTTCAACTGTGGTGTTTGATCATCTGCACCACTGCTAGAGCTCCAG
CCTCTAAGCCTAGACTTTATCATTTTCAAGCTCTCAGAAGGTTTTTGTAAGGGAGCCCCTATGTAATTTCTGCTACAGTTATTG
ATTGATACCGAAGCTGTCGATGTAGCTGTTAGTGATCCTTGTCGCGAGGAATGCTTATCTGAGGATGCTTGTAAAACGTTCAC
AGCTGAAAAACCTATAGCCCACGCGAAGCCAGCAAACGGTAAGAGAAGGAAGAGAAACCAACGAATGAGAAGTGAAGGGC
GACAACAACGCTGCAGCATATTGATGCTGATGCTGATCACGCCAGAAGCTGCCCAAGAGAGCTAAGCTCATCATCAACGTGC
GCTTCTCGCTGGGAGACAACGGGGAGGAAGGAAGAAGAGGAACGAGGAATGCGACATGGAGAGAGAGAGAGAGAGA

**GAMETE EXPRESSED 3 (*GEX3*):**

GGAAACAGATTCTGTGCTTGGAGGGAAAAGCTCTGCTGACTTCTGTTCCAAGCTGACAATAGAATCATTTGATGCACT
AGTTGAATACTGATCAATTCCCTCAAAAAATTCATTAACGGGAGAAAATGATACAGTTTTTTTAGTGGAGCTTCGAGCTACGG
TTGTGCTTTTATCATACAAGGGGTTACTGTACTTCTCGGTGATACTTTGGCCGAGAACTGCACTGGAGTCTTCCTTGCTGTTTT
GATAATGAGAGGCCTGCATTGCACTTGATGATCGCTGCAAGCTTAAACTACGTTCATGGACTCGTGGCGTTATCCATTGAGGA
AGAGTCAGTGTGAGAGGAGCTGTGGGGCAATCATCATATAACGAAAGAATGTGCATTTCCTCAGTGAAATCCTCGTCTTCCG
GTGCACCGAGTAAAGCGCGGAAAGCAGTATCGAGCAACACAGTCCAAAACCCACCTCGTGAATTGGTAGATGTGAAATCCAG
GGAAGCAATTGCATCCTCTACCAGAAGCTGCTCTTGACGAAAATGATCACCAAGCAATGTGTCTGATGGAAATAATGCCAAG
TGTCTTTCTATGAGATGAAGCCCTTCAAGGGTGCCCTTTAACTGTTCGAGAATTTTAGAAGATCTAGGTTGAACCTCAAGTTCT
CCCTGCAGATGCTTGATCGTTTGTTTGTACATTCTTTGTCTCTTTAAAAGCAGCCCCTGACGCTTCCGAAACCTCATCTGCTTCA
GCAACAGACAAAGTCTGTGCTTGTGCAATGCTTGCTTTTTCCTCCAAACACAATATATGAGCACGCTCAAGAGTGACAATAAC
AGAAAGATGAGGAGAATAGAAACAGAGAAGGCAATGATTGGCCCATCATTACCTGGGTGTTGCACTCTTGTGACTGTAAACA
AGAACTGTGAACAGCTGACTTCATGAAGTTTTTGTTCGGTTTTGCAGAAGCTCCTACTTGAAAGATTTGCAATTGATGTAAGC
GCAATCAGCCAATCAGGGTCAATACTGTATGAGCTCAAGTCCCTCTTTACTAAATCTTCAGGAAATTGGTCTGTTCAAAAAAC
CAGGGTACAATGTATTCCACATGATCTGTCCACTTGATGGAACAACCATGTATAAGTTCATGCCCAAATATTGTGTGAAGGAC
AAAGCTTTGGTTCCATCCATTTGCTCCAATATAGTTTTCGACTTGACTCTAGTTTGAGCAACAAATAAAGCTTCTTTTGAGCAA
TCAAGAATAGGTGAAATTTGTATGCTGACTGGAGCCGAGTTGTCCGCTGCCAGATATTTGCTCACATATGCTCCATCTGGGGA

AATGACATAAAGAAATCCATCCAGAGAGCCAATGGATACAAAACCTAGGAAGTCGACCACTGGGATGCAACTTGTTTCACTG
AGAGGTCCAACCTTTGCTTGCCAAACCAGATTCCCCAACTCCGTCGATAAAGCATACACCATATCATATTTGGCCATGGCTGC
ATAAATGCGCCCATTACTTCCAGTAGTTATTTTGACTGATGATGGTACATCAAGACTGAACGTGTAACGCCACCTCATAAACG
GTTGCCAGGTTGCATACGAAAAAAGCCAGCCGTCTGTACGAGCGACATAGATGCTTCCATCACATTGGTCCACCGCCAATGAT
GGAAGAAGGGTGCAATTTGTCATATTATGAGCACAAGGTCCCATAGACTCTGGAGACTTCATCACGCTATTGCTGAACTGGAC
AAGCGTGCTCCACAATGGTGTCCCATCCAACATAAATGCGTGAAGGCCTGTACCCGTGTTAATGTAAAGGACAGAGCTCCAA
GATGTCACCGCAAGGCCCGTTATGGAAGAAAACTTGCCAATAGCTTTCAAGCTTATGGCTGGAACCATACTAAAAAAATCAG
CTGCAGAAGCTTGTGTTGCTTGCGATGCTGGTGGAGTCACAACCACTACAGTCTGCTCTGCTGCCACATAAACCTTTCCATCA

**LONG CHAIN BASE2 (*LCB2*):**

TGTCATTTACTTTGAATTATAGAAGGCGAGCAATCAAGTTGACATTTCAATGAGCGAGAGTATTTAAATTGGGAAAGG
ACACATGGCGGGATATGGTTGGGTAGCGAATCAGAAAGTGAACGTCCCCTGAGACCATAATTATGGGAAATGGAGAAACGG
GAATGGGAAATGGAACAACAATTTTGAGAAAAATCGAAACGGGGATATGCCATATCAGCAAGAAATTGTAAAATTTTGGGA
AGGAAGGAGAGAAAAAATTGAGAGAAAGGAAGGTGATGTATCGAATTTTAAGGTTAATGTCAAGATTTTGCTCTATATTACTA
AGATATTGAATGAAGCAATATGTGAGGAACAATTAATGCGATGTTATTGTTTCAATTTGAGATCGGAAATGGACTGAAACAG
CCGAAACGGGAAATGGAGTTTCCAGTAACTATGCCTCAAACTACACCCATGCAGTTTGCACAGCACCCAGCTTGAGCCTTCTG
TAGCTCATCTGATCTGGAAACACCTTGCAATGAAAACCCAGAATAGTAGCAATAATCTGGGCCTTCTGGACCAGATTAGGAA
GCATACATAATGGTTTTGCCCCTTATTTGGGCATGGAAGGAATTAAAATGTTGTTTGAGTGTTTTATGGCATTGTTTAACTCCA
CTCCATCTCCTCAGTCGAACAGAGAAGACACGGACTTGCATAAATCTGGGTTCATACAATTATCTTGGATTTGCTGCTTCTGAT
GAATATTGTACACCTCGAGTAATTCAGTCGTTGGGAAAGTTTGCTCAGAGCACATGCAGCAGCCGTGTGGATGGTGGAAACA
CAGTGCTGCATGAGGAGCTTGAAAATTTGATTGCAAGATTTGTCGGAAAGCCTGCTGCAATGGTGTACGGTATGGGTTTTGCG
ACCAACTCTACGACTCTGCCATCCTTGATAGGAAGGGGAGGTCTGATTATAAGTGATGCTTTAAACCACTCATCAATAGTCGC
AGGAGCACGTGCTTCAGGTGCCAAAGTGGCAGTCTTCAGCCATAACACACCAGAGCATTTGGAGGAGGTTTTGCGGGAAGCG
ATTGCCGAAGGGCAGCCAAGAACTCACAGACCTTGGAAGAAGATAATTGTTGTGATCGAGGGCATTTACAGTATGGAAGGAG
AGCTCTGCAAACTTAGTGAGATAGTGGCAGTCTGCAAGAAGTACAAGGTATACATCTACCTTGATGAAGCTCATAGTATAGG
AGCCATTGGCAAAACAGGCCGAGGTATTTGTGAGCTTGCTGGCGTCAATCCTGATGATGTCGACATAATGATGGGCACCTTTA
CGAAGTCTTTTGGATCTTGTGGAGGTTACATTGCTGGCTCAAAGGATCTTATAAAGTATTTGAAGTTTGTTAGTCCTGGGCATC
TGTACGCAACGGCAATGTCTCCACCAGCAGTTGAGCAGGTCATCTCGGCTCTGAAGGTCATCCTTGGTGAAGATGGTACAAAT
CGAGGGGCTAAGAAGCTAGCACAAATAAGGGACAACAGCAACTTTTTTCGAGAGGAGCTTAGAAAAATGGGCTGCGAGGTT
ATCGGCGATCCGGACTCACCCGTGATGCCAATAATGCTGTACAATCCTGCCAAAATCCCAGCTTTTTCAAGAGAGTGCTTCAA
GCGGAATGTCGCAGTTGTAACCGTTGCGTTTCCTGCTACCCCATTGCTGCTAGCAAGGGCTCGCATTTGCATATCCGCAGCTC
ATACAAGAGAGGATTTACAACGAGCACTTGAGGTAATAAATGAAGTAAGTGACATGGTGCACTTGAAATACTTTCCCTTTCTT
AAAGATGCTGAGGAAGAGCAGCAGAAAAAGAAGCTGCAATGATACCTTGTTTGAAATAGTAGACAATTGGGCTGAAAGTAT
CTTGAGATGATAACAGCTAGCTCAGCTTTGGAAAGCCAATAGAAGTGCTTGGTCATTTTTTTCCCATTTAATGTATCAGTATCA
AGCAGGGTCCTCCCTCTATTTAATCACACTTCTTGACCATTCCAGAAATTTGGAGGAGAGTATTCTTTTTGTGACTTTAAGACC
TAATGAATGTTTTTTTACCATGATTCATTGTGCATTTTGTATTCGGTTGTGAGTCGACGCTTTTTCTGAATGATAGTTGGTGTCT
ATTTTGG

**FERONIA (*FER*):**

AAGTGGGGCGTTCAAGCAAACCATAGCCATAATTTATAAACTACAAGACTTCTTCTTCTCTTTGTTTCGTTCATATCAT
TGATTGCATCAGTATCAGTACATATCTCCTTTTCCCCCATTGATACGCTCCTTTGCGCTTTCTGCTTTCTGTCAGAGGCTCGTAG
ATACAATTCGCAAGTGCCAAGAAGCATGGGCGATCTCCTCATGCATCCAAACATTCGGGAATTTTTGGTAATTTGCCCATCAG
AGCTTTTTTTGCTTTGAGAGATGGTGTTTTGAAGGTGCCCAACCCTTCGCAGCCTGAACAATTAGGGTTTTCTTCATGGGTCTC
CTTCATTTCTCTTTCTTCAATCACTGGTAAGCTTTCTTTCCGAAGCTCATTAGCTTTAAAGTCTCAGACATGTGTCTGCGATTCG
CCTGCAAAATGAAAAGCGGTGCAAATTGCTCATTGCACAGTTGTATTTCTTGATCTTTCAGTAGTCAATGATAAGCTTCGACT
GCCAAGCTTTCCATTAAAGCTTCACAACTTCAAGGTCACACAAATGGGTGTTTTTGAATAATGAAGCAAATCCTAGAAGTTTA
CAAGCTGTAAAGAGCTGAATTCCCTTTTTTGTTTCTTCGTCCTGAAGCTTTTACCAAGCACCAAATCAGTTTTTTTTATTTGAGC
CAAGCATTGAATATCTCATGGTTTTCTGATGGCTTCAACAATCCTAACACGCTTGTCTTGAATGGGCTTTTCAAGATCTCTCTT
GTGGAGTGAGTTATCATCTTTCTGGTGCTCCTGAGGAGCTTTTCTCGGTGTTGGGGTTCCATCAAGTTCAAAGGCTTCTAGGTC
TTCTTGCTTTGTGGCGGATTTGCGTCCAAGCTTTGCTTTTGAAGGCTTGACTGATTTTTGTCATGCCAAGCTATGCCCAAGATA
GCTCCTTACCCCAGTAACATGCTTGCAATAGTCATATCAATTTAACACCATCAGTCATCAGATTACTTAATATTTGGGCTGATA
TCTTTCAGTTTCACGTCAAAGCCTTTAATGGGGGTCATTACTTTACTTCTCCATATTGTTTCATTTCTGTTTCTGCATATTACAT
GCCTCTTTGTTCTTGTACAAGGGCTATCCTACACCCCTAAGGATAACTACCTTATCGCCTGTGGGAGCTCCCTTGATAGCGCTA
CAGATTCTGACGGTAGAACATGGGTGGGTGATGAGTCTCCTTCTGCTTCACCATTTTTGGCTGGAAATCCTGTATCTGTGGCA
GCTTCTACTAGTGTTCAAAACCCATATCTCCCAAGCACAGTTCCTTTTCTTGGCGCTCGCATTATCACATCTCCAGTTTTGTATT
CGTTCCCTGTTACGCCAGGGCGCCATTGGTTGCGCTTGTATTTCTATCCTTTTGCCTATGAAAATTATGATCCTGATTTGGCAA
TTGTCAGCGTGAATGTAGATAATTATGCTCTGGTTGCCAATATGAGCATTACTAGAGAGATGAATGCTCTGAACTATGGGTAC
ATTCAGAAAGAGTTTTCCATCAATGTGACCTCCAAGTCATTGTTGGTGAGCTTCATTCCAGGCAAAGCAAATCATGCATATGC
TATGGTGAATGGAATAGAGGTGATTTCAATGCCGGATAACATGTTTGAAGACAGCATATATAATGTTGATCTTTCAGCAGACA
TACCCATTGCCATGAGTGCGACGGCTCTAGAAACTATGTATCGGGTGAATGTGGGCGGACAAGCAGTTAGTGCAGCAACCGA
TGAGCTTTCTCGAACTTGGGGCACCGACATTGGTTTTATCCCCACAGCAGCAACAGGAGTTGCAATGAATACAGAACACCATA
TACAATACGCAACCGCCAACAACTACACTGCCCCTCAAGTTGTCTATCAAACAGCCCGGACTATGACAAATGATGACCATGTT
AATTTAAATTTTAACCTCACATGGCGCTTCTCTGTCGATCCTGCATTCACCTACTACGTTAGGCTTCATTTCTGTGAGTTTGTT
ACCCATTGACCAACATGAGGGTTTTTGACATCTTCATCAACTCAGAGGTCGCCGAACTGTCCTTTGACGTCAATGCACAGGCA
TCTGTTTTGACTGGCAATACAGGAAGTGGTGAGTACAAAGCTGTTGTGATGGATTTTGCTACTTCAATTGCATCCAAGAGTGC
ATGGAATGCCAGTGATATAATACTTGCACTTCATCCTAACAGTACGAGTGCCCCAACAAATTTCAATTCCATATTGAATGGCT
TGGAAATTTTTAAACTAAACGATTCTTCTGGAAATTTGCAAGGCCCTCCTCCCCCTCTTGTTGCAGTTCCTAGTGGTACTAGCA
CAGCAAGGCAATCTTCGCCTTCTCATGGAGGGCTTAAACAACCAATCATCGGTGGGGTGGCAGGGGCTGCAGCTGCAGTTGC
ATTACTGTTGGTAGGAGGCTTTCTATGCATGCATAGGCGCAAGAAAAGCGCCAAGAACTCCTCACAAGCTTGGCTGCCTTTAC
CTCTCTATGGAGGTACGTCTCGGTCACTTATCAGTAAAGGGTCTACTGCATCTCCAAAGAGTGCAGCGGGTAGCTATGCATCG
TCGGCACCTTCTAACCTGAGTCGCCATTTCACATTCGAAGAGATATCCTCCATGACTAACAATTTTGATGAGGCCAGGGTCCT
GGGTGTGGGGGGCTTTGGAAAAGTCTATGAAGGAGTGATTGAGGATGGAACGAAGGTTGCAGTGAAGCGAGGCAACTCTAC
TTCAGAACAAGGTGTCATGGAGTTTCAGACAGAGATTGAGCTACTCTCTAAGCTTCGCCATCGCCATCTTGTCTCGCTCATTG
GCTATTGTGAAGAACACAACGAGATGATTCTGGTATATGATTGCATGGCAAACGGTCCTCTGAGAGGTCACCTTTATGGTACT
GATCTTCCTAAATTATCATGGAAACAAAGGCTTGAGATATGCATTGGGGCTGCTCGTGGTCTTCATTATCTTCATACAGGGTCT
GCACAAGGCATCATACACAGAGATGTGAAAACAACCAACATACTCTTGGATGAAAATCTCTTAGCTAAGGTATCAGACTTCG

GGCTTTCCAAGACAGGCCCTACTCTTGATCATACACATGTGAGCACAGCTGTAAAAGGAAGCTTTGGGTATCTGGATCCAGAG
TACTTTCGAAGACAACAGCTTACAGATAAGTCAGATGTATACTCTTTTGGGGTAGTCTTGATGGAGGCACTGTGTGCGAGGCC
AGTTATCAACCCCTCTTTACCAAGAGAGCAAATAAATTTAGCTGAATGGGCAATGAAGTGGCAGAAGAAGGGGATGCTGGAT
CAGATTATTGACCCTTATTTAGTTGGAAGAATAAGCAGAGAATCATTGAACAAGTTTGCGGAGACAGCTGAAAGGTGTCTAC
AGGATCGCGGCTCTGAGAGGCCAACTATGGGGGATGTTCTGTGGAACTTGGAATATGCCTTGCAGCTCCACGAATCTTCTGTG
GAGAAAGCATTGGATGAGAGCAAGCCTCGAATAGTGGATTTGCCGATTCGAGCATTGGATACAAGTGAACTCGATGATAGCT
CTGAGGTGAAGATAAAAAACAGTAATGAGAGCTACAGTAGCCCAAGGACTCAGAACTTGCTAAGCGAAGACTCGGATGATA
CTTCCATCAGTGCAGTCTTCTCTCAGCTGGTTAACCCTCAAGGACGCTAGACTGCTTTTGATCTTCTCCCCATGAAGAGATTGT
AAAGGTAAGCTGGGCCAGTGTATTTTTGTACAAAATAATTCCTTTTGGTTTTTGGCTGTTGATAGGTGCACTGTTTAGTGTTCA
AGAAGACTAGCCAAACCCTTTGTTGTGTACACTATTGTTGGCTGTTAAAGGTTTGCCAGTTTTTAAGGAAATATGCCAAAAAG
CTTACATTTGTACACTAGCATTATTGGATGCTGCTAAAGGCACAATTTCCTATCATTCTTGCCGGATCTAGATGCTACTGTGTT
AAGGAAAATTGTGTTGCATCTTAGCTTTGGTGAGAGTTGTCATGCGCAAGTCTAGGGGCTAGTTTGAATTTAGGACTTGACAT
ATTGAGGGAATTGGGTGTGAGCTGTTTGCATTCTCCATCTCTTTGTTTCTAGCGGGTTGTTGTTTGTGCCATAGATGGCTGATT
GTAATAGAAATACAACTATGTAAGCAAAGAATTTCCTTTTACAGATGTCATGTTATCCATGGAATCAGTTAGAAGTGCTTGAT
ACTTAGCTTTTATCCATTGCCTCAAAGCTGTTCCTTGGTAGGTTTTGCTTTTA

**F BOX-LIKE17 (*FBL17*):**

CATACACACATTCAAAACAAATCCCTCCATGCTAGCCTAGCCACTGTGATGTAGGAAAATGTTTGCATCCATTGTTTA
CACCACGCATGTGCTACAATTTGCCCCCACTACTAGTGATTTCCATCAAGGAAGTTCATTAGAATTGACTGTCTACTAATTCCA
ACATTCCATTCACGGATTTGATTAAATGATTACCTCTTGGTCTTACAAGAAGGGAGCATTCATGCTTGTTACGTGCATGGGAT
GACAAAGCTATTGATCTCAGTGTAATGAGAAGTTAACAATGTTAAATGTGTTGATTAGAAAGGCATTCATTGCAAACCTCTCA
AAGCTAGTTGGAGCTTGTAAGAAAAGATTATGTAAATGATTTTGAGTTGAGGAGATGAGACTGGCAAATGTGCCTTTCAGGA
CATCTCAAGTATTGCTTGTTTGCCGTATACAAATACTTTTACTATCATCGTAGTGCTTGGACACGCTTCGAGCCATCTGCTTGT
CCCCTCCAGATTGTAGTCGGATCAAAGTGAGCCAGCTCCAGCTATAACAAAAAAAGAAAAAGTTCCATTAAACCAAGTAACT
ACAATCTGAATTGACTAAACCATCTTAAACCAAGTATCTTGCCTCCTTTTAGCTGGTATCAACAATTTAACAGAAAGAAAGCT
TTTTCTGATATCATCCATCCCTACAGATGCCTACCTGACCGTGAAGTTCGTGTACCAGCTCTTCTTCGCACTTTCCCACATAGA
CATCTTTTAAGGATGGGCATAACAATTTCAGTCCACCCGGTTGAAGTTTTGAACAACCCGTCAAATTGAGGTCCACAAGACTT
GGGCAGTCCAAAACTAGCACCTTAAGTCCAGTGCAGCCCCAAAAACTCAGCTTCTGTAGTTGCTGATGCTGAAGCCGCAACTC
CCCACGTCCAGGTTTAGCTTCTAAATGATTAGGCTCAATAAAACATGTACTGTCAGCATTGTCTGCCCATGTGTAGGTCCTCTT
ATCAGTAATGACCTTCCCACAATCTAGAAGCTGCAGAAGAGGAAGCTTAGAAACCACCATCTGCAAACCAACTTGAGATACT
CTTGGACAGAGAGCAATCAAAAGCTTGGATAGACTTTTTGAGAACGACTTGCAAATCAGTTCAAGTGCCATGTCAGTGATGCT
TGACCCACTCAGATCAAGCAACTCTAGATTCTCACAGTTAGCCACAATTGCATCCACGGAACAGTCTGTAATGCCCAATCCAA
ATGTGAGGGAAAGCATCCGTAACGACTTTAAGCCAGGAGTGAAAAGACCATGGACAACACGATCTTGTAGCCAGGGAGAAG
CAACATGAAGGCGGGTCAGACCTCTTGCAGTATTTCCCAAATTTTTCATAATAGCTGCTAGGCTGTCTGATTTCACGTGGTCCT
GCTCCAATGGATCCACATCTTCACGAGGCAGGAAGTCTAAGCACAGCTCTGATAACTCTGGACAGTCCAGATACATATATTTG
AGGTGATGGCATCCAGTCAACCAAAGCGTTGCAAGGCGTGGACTGCTTAGCTCCACTTCAATCACACGATCACAACCCTCAA
ACTTAACAGATGTCAAAGATCTGCAGTAGCGGAATAGGTCTCTTAACCCTTCACCGGATATTGTGTTTACACCACCAGGTTTC
AATGATACTTCCAGAGATGAAAGCTGGTAGCGACTGAAAGTAGAAATGCGCTTGAGCAGTTGATCTCCCAGATGGCTTGTCA
CGTCCAGGGTGAGCTGCTGAAGATTCGAACACTTCCGCAGGATGAATGGGAGGAAATCTGCGTCGCCTTCTACCCTGTTAGA
ATGAGACGCCCTGATCGAGACGATTGTTTTCCGCAAACCTGCCCAGACATACTCCGCACATTGACGCCACCGCCTACATACAC
TGCTAGCAGCTACAAGATCTTGCGGGAAGAGACGGGATAAGATCTGCACCAGACAAGCAAGTGGCATCGCCTCACAGTCCTT
TGCTTCATTCACTTCCAGATCGTAACTGCTGTTTAAGCCAAAACAGCGAAGACCAGCATGAGGACACCTGCCTGCAAATTCAT
CAGCGACCAGCAAAGCAGTGTGTGACCATGAATCCGCGAAATACTTATTGACAAAGTGAACCGCGAGGATATTCCGATCCGA
GAAGCAAGGCTGGGGAAACGAGGGCAGCTTCAAGTACATCAAGTACAAGGTAGGTTTTCACGGGGACATAAAGAACAATGG
GGTTCTACAACAGCAAGAAAATGACGTTTTTCTATCACATGAGAGGTCACAAACGACGGCACAGAAGAACCAACGGGACTAG
AGAGTAGCCTCTACACAGACAAAAGAAGAATTTTGATAGGGAGGGCACCTTAAAGAGTAAGCACATAGAGGATTAAGCTTTA
CTCTTATATCATACAGGCGGATTAGAGGAAATCGAGGGCACTTCTGCGAAATGAGCAGACAATGCATGTTCCCAACCGACAA
AATCACGAGTTGAAGAACAAAAGAATCAAGCCTTTAGGATATCAGAGC

**ARGONAUTE 1 (*AGO1*):**

CCTGTTTCCCCCTTCCCTTCAAGATCAGCTTATCTTTTTTGGGTTTCTTGCGTCTCGCCAATCGCAGTTTTGTGTATCGC
CTGTTGCGCGCACACATTGTGCCGCTTCAATATCATCGGCTACCCTCGTTTTCTTTTTCTGTTTTGTCTGTTTCTCTGCATCATG
CCTCCTCGTAGAGGTCAAGGGCGTGGTCCTAGCGCCCCTGCCCGAGTTCCTGAATTTGAGGAGGATAGAGAAAGCAGCTTTC
GTGGTGGTGAAGTGTGGAGGCAGCAGCAGCTGCATCCTCAACAGGAGCATCGACAGCAATACCCCCATGTGCAAGCGCATCG
TGGTGGAGGCCGTGCTGGAGGGCGCGGTTTTTGGCAGCAACAGCCTTTTCTTGAAGAAAGAGGCGATCAAGGTTTGTCTGGA
TGGAGAACAAGTAGTGGTGGAAGAGCAGGTCAGGATCCTGCACGTGCACCGTCTCAGGATTGGCGTGCAGGTGCAGGGGTGC
TTCCTAGGACGGTGCGAATTGTAGCTCCCGAACCTGTGGGAAGCTTTGAGTATGGTGGAAGGGGGGAAGATCAAGGAGGTGG
AGGAGGGAGCCTTACACATGGTGGACGTAGAGGGGAAGATCAAGGAGGTGTTGGTGCTTTTATGTATGGTGGTAGAAGAGG
GGAAGATCAAGGTGGTGGGGGTGGCATTGGCTATGGGGGTAGAAGAGGTGAAGATCAAGGTATTGGCGGAAGAAGAGAAGA
GTATGGAGGGGGAGGGGGCACCAGGTATGGCGGGAGAAGAGGAGATGATCAAGGAGGGGGTGCCAGTTTTGGCGGGAGGA
GGAGTGAAGATCAAGGAGGGGGGGAAGGCCCTGGCTTTTCAATCAAAAGAGGGGATGATCAGGGAGGACCTGGCTATGGTG
CGAGAAGAGTTGAAGATCGGGGAGGAGGGGGGCCCATCTTTGGCGGGAGAAGAGGGGAAGATTATGGAAGGGGAGCCCGA
AGAAGTGGAGGTGCAAGAGGAGGGGGACGATTCATCGACTTGGAAGGCAGAGGTCTCGAGGAATCTGAAGCTGGAGGTAGC
AGCATTGCGAGCGGGAGCTACAGCGGACGTGGAGGTGCAAGGAGAGGCGGTCGCTCATTCCAAGCACCATCTCAGGGTTTCA
CGTTCTTCTCCCTCTTCTGTTGTTAGTTTTATCATGGTTACATGTATTCTAGAATTTGTATGTGTAGGCAGCAATGGCCCGGAGT
TCTGTAAGGACTTGCATTTGGCGACCGTATTGCAATGCCAGGGGTTAAATATTAGTGTGTCTTGTAGTGCTTAGAGAACTTCC
CCATCTAATCCTGATGATGTTAGCCCTCACAAAGGAGGTTCTCTTTTCTGAAGCAACTCGAGGAAATAGGTGTTTAGTGTTGA
TCGCGATAGTTGTGCATGAGGTAATTCTGGCAATTTTGGGTTTTCCTGGTAAGGATGTAAGCAAAACACTCGCTCGACCAAAA
CCTTATTCAGGGTGGAATATGTCTGATCTTCGCAGCCATTAGCTTGGTCGAGTATGTGCTGTTTTGCAACTAACACATGAGTTT
TGAAGCAGCCTGGTTGTATCTTTAATCTGCCTTCCCGTTTGTTTGTTGCATCTTGCATTCACCTGTAGCTTGTCGATCCTATAAC
AGAAGCAATTTGTTAGATTGTTATACATGCTTTTTTTTCCTGGCATTTATGTAGAGGTGGCTATTAGTCTAACTGACAATGGCA
GCAGAACTACTGTAGATGTGTGGGAAAAAGGGCATTTTCATCCCGTGCTGTGTAGGTGTTGTGTAATTTGCTAAAGTAGGGAG
CCTAGCATCACGAAATCACACATACGCACTCGAGAATAGGGGACTCAATCAATGCACACAAGTTTAAGTAGTAGTCCAAACA
CTGAACAAGCTTTCAAATGCAAAATCTTCCCTCCATGTCAGATCTGAGTCGGTTGGCCAAATGGTGGAGTCAGTTTTGATGTG
TGCCTGTACACCATGCATTCTTCTAATCATGATTGACTTGGTGTTTTAATTCTTTTCATTTGATCAAGATGCTATTCAAAATTTT
GATATCTGTGGTTGTAAAAATAGCTAATTAGTTTTTTTGGCTCATGGCTGTAGAAAGTAGTTGGTTGGCTTTGATTAATTTATT

GGCTCAAGCAGGATCGTGGGCTTTATCAGCACCACCGTTGGGCGAGCTCCTTCCAGGTTTCTCTGGCTTGGGGATACATGAGG
AACCATCCCAACCTGGTTCTACATCTGCAGTAGAGGAAGCGTTGCCCCAAAATGTGGATGAAGGGGAGACGTCTGCCATTGT
GCCACAGGAGCTTCCATCCGCATCGGGGTCCTCTTCACGATCATCTTTGAAGCTGATTGCTCCCAAAAGGCCACTTCCTGGAA
GCACAGGGCTAGCTATCAAAGTTCGGGCGAATCATTTCAAGGTAACCTTCCATCCTGGTGACAATATCTATCAATATGATGTG
GATATCAGCCCGAAAATCTCATCCAAGATTGTTGCTCGCACTCTTGAACGCCAGCTAGTTGAGATGTATAGTTCTGATTTCAA
TGGCAAACTTCCAGTTTATGACGGCAGCAAAAGTATGTACACTCATGGTCCCCTTCCATTTGAGCATTGTGAGTTTGTTGTCAC
TCTTTCAGAAGAGAGAGGAACTACTGGCAGGGAGAAGAAATTCACTGTAGCAATCCGTCATGTCTCTACCCTGAACAGGAAA
AACTTGGAAGATTTTATGAAAGGCAAGCAGGTTCCAACTCCACAAGAGCATCTCCAGGCCCTTGATGTGTTACTGAGAGAGC
ACCCTGCATTACATTTCATAGCAGTTAGCAGGTCTTTTTTCAAAAGCGAGTTAGGTAGTGCTCAGTTGGAGGGAGGGCTTGTT
GCATTGAACGGTTTTTATCAAAGTCTTCGACCTACAGAGAGTGGATTGCAACTCAACATTGATTTGTCCACAACAGCATTTCA
TGCTAGCATTCCTATAATTGATTTTCTCAGGCAGCAGTTAAGGAATTTCGACCCACGGTATCGGCTAACAGATGTGGTCCGTG
TGAAGGTCAAGAGAGCACTTGCTCGCTTGAAGGTGCAAGTTATTCACAGGCAAACACCTCGGAGATACAGGATATCTGGCCT
TTCCACAAGCCCGACGAAAGATTTGAAGTTTCCCATCGAAGGTGGGGAGGAAATGCGGGTTGTGGATTATTTCAAGCTGACG
TACAATTATGTGATCGAGTTTCCAGAACTCCCTTGTTTACAAGTTCAAGCCAACAAGCCTAGCTACCTGCCAATGGAGGTTTG
TGTGATCTGTGATGGCCAGAAATATGGTGGCAAGCTCAATGACAGGCAAACCACGAGATTAAGAGGGCTGGCTTGTGTCCTG
CCTAAAGTGAGGGAAGCTAAAATTAGGAGCATTATGAACAACGATGATGGTCCAGGCAGAGGTCCGCATGTAAGGAATTTCG
GTGTGCAAGTAGCCTCTGAGATGACCTTGGTGAATGCTCGCCTTCTGCCTCCCCCTAAGCTCAGGTATGGAGATCAAGGGAAA
GTTAAGGAGATAGTTCCCACTGATGGTGCATGGAATCTGTTAAACTCCTGTGTCGTGGAGGGTGGGAATGTTGCTTATTGGGC
TCTAATCAGCTTTGATCAGACTGTGAATGACTATATTGCTGGCAATTTTATATCCAGCCTAAGCAAGCGGTGCAATGACCTTG
GTATTCAGATGGCTGAACAGACAGTAATCCCTCCAGTCTTGCGCAGGTGGGAGGACCTTGAGACCCCACGACTCGAGAAGAA
TCTGAGATTTGTCTATGACCAAGCTTCTCAAGTCATTCACAGTAGTGAAGGGCAGGGAATACGTCTGCAATTGCTTGTTTGTG
TAATGGCTGATAAACACCCAGCTTATGGTGAGCTGAAGAGGATTTGTGAAACCCAGATAGGTATTGTAACTCAATGTTGTCTG
TCAAGGCATGTAAAACAATGCAAGTCTCAATATTTGGCGAACTTGGCTCTGAAGGTGAATGCTAAGGCTGGTGGAAGAAATG
TTACATTGGCACTTGAGCTTCCCAAGATGTGCCCTGTTTTCAATAGACCGACAATTATATTTGGCGCAGATGTCACACATCCAT
CTCCAGGTGATGATACAGGACCCTCCATTGCAGCTGTTGTGGCCAACATAGACTGGCCTTCCGCCAACAGATACATAGCACGC
GTGCGTGCACAAACTCACAGAGAGGAGATAATTGAGTACTTGAGGGAGATGGTGCAGGAGCTTTGGCATGAGTTTTGTGAAA
AGACAAAGAGTCGGCCGGATCGCGTGATAATGTTTAGAGATGGTGTAAGTGAAGGTCAATTTGATGAAGTCCTGCAGAGGGA
GGTGGCCGCATTAAAGGATGCTTTTATAGAGGTGGGTGGCCCAGATTACAAGCCTTTAATCACATGGGCGGTGGTGCAAAAG
AGGCATCATACCAGGCTATTTCCTGCCGATGACAAATGCAAGGATAAGAACAACAATATCTTACCAGGCACCGTGGTAGACT
CAACCATCACGCACCCAAGGGAGTTTGATTTTTTTCTTTGCAGTCATGCTGGGATACAGGGCACAAGCAGGCCGACACATTAT
CATGTGCTTTGGGATGAGAACAATTTTAAGTCTGATGATCTGCAGGGTCTGGTTTACAATCTATGCTATACTTATGCTCGGTGC
ACACGTTCGGTTTCTGTTGTTCCCCCGGCTTATTATGCACATCTTGCAGCATATAGGGCTCGCCTTTACCTGGACTCTCTAGGT
GGTTCCGATACTTCTGCTTCGTTACGGGGTCAGCGCTCAGGTACTGGATCAAGTGGTGGAGGTGGCAGCTCTAGAGCTGCTGC
TCCTGCTGTGCGTCATCTTCCTCGTGTGCAACGCAATGTGCAAGAAGTTATGTACTTCTGTTGAGAAGTCTGACAATCATCATT
GACTTGTTTAATTTTTGAATGACTTGCATACACAATTTTGTCTGGTCATGTGATCGGTGCTTAGTTTCTGGACAACATTAATTA
CTCAGTTGTTGCCATGTGGCCAGCCTAATGCAGGTACCTGAACCTAAATCTTTGAGGAGCAGAATCTTCTGCATTTTGCTTTTG
TGTGAAGACTTTTACCGGGTCTTAAGTGTTTTCTCTTTCTTGCAGTTGAGATTCAATTTTGAGGGTGTGTAACTTACACCCTTG
GTTCAGCTACTGGATGCTCGACATGATTAGTATGCATGCAAATTCCACATTGTGAAGTGTTTGTCTGATACACCCTTGGTTCAG
GTACTGGATGCTCGAAATGATTAGTATGCATGCAAATTTCACATTGTGAAGT

**ARGONAUTE 7 (*AGO7*):**

GCTGATCATATCTACCAGTATGATGTGAAGATGGTAAACTCACGGGTGCGACAAGTCCCTGACAAGCCTTTTAAACCT
CTTCCGAAAAAGATAAGCAGAAGGATTAAAGAACAGCTTATTAAGTCTTATCCTGATGTATTTGGGAACATAAATCCAGTGTT
TGATGGCAGCCAGAATTTCTTCACAACAAGCATGCTTCCAATTACTGAGCTGAAAAGCTTCGACGTGACCATCGTTGATGATT
GCAATCACCAGCCGATAACGTACTCTGTGACGGTGCAATTAAGCGCGAGCTTCGCAATTGCGAGTGTGCAGAATTACATCTTT
GGGCAAGTTGTCGAGATGCCTCAGGCTGTGCTACAAGCTTTGGATGTGGCAATGAGGGAAACATTGTTGGGTCATTACACAG
AACAGGGTAAATCATTCTACGCATCAAGCACAAACAAACAAAAATACAGTGGATTTCAGACTGATTACATGGAAATATGGGA
TGGTTTCTTTCAAAGCTTGAGGATCACAGAGCAGGGTCTAGCTCTGAATTTGAAAAAATCGCATGCAGCATTCTTTGCAGATA
ATAAACTCCATGACAAAATTACAGTTGCAAATTATATCTTCTATTTGAACCAATGCCGTCCTATTGATTTAGAGGGCTTAAAG
AATGAATTGAAAGGGGTGAAGGTGCTAGTTCACATGGAAAGGGATGCTGGACAGGAACCAATTAAGAGCCGGATGTACAAG
ATAGAAGCCATCAGGAATAATGCATATGAGGAGAAGTTTATTAAAAATGGGAGAGAGGTGTCTGTGGCACAGCATTTTGAAG
AAACTGGGAGAACACTCCATTTTCCAGAAAGTCCGTGCGTGAAGGTGCTCCACTTGCGCAACATACTCTTCCCCTCTGAGCTT
TGTGTTATCGAGTACCCCCAGCACATTTCCAAGACGCTATGCTTTGCACCGCACCGCAAACATATGCTCGATGTAGCTGCTGC
CCTCGACCCCACAGAACATTGCAAACAGATTCAAGACCTCATGCCAGAACCTCCCACAGCTGCATCGCCCTTACCCTATGTTG
GCCCCCCGCGCCAAATTTGGGGTGTTGTCACAGTACACCAAAAGGGTTTGAGATGCAGGTGTCCTGTTCGATGACGGAGTTAG
AGGCGAGGCTAATGCGAGCACCCAAGCTGAAGCAAATGCCAAGCAGGCCAGGGATGCAACCACTACCAATAGTAATAGAAG
ACAAAGGATCATGGAGTCTAAAATCAATAAAAGTAGCAGAAGCAGCAGAAGACATCACTAAATGGCTGCTAATCACCTTTTC
TAACGGTCCAAATCAAAAGAATTCCCATTTTTCTAATTTCCCAAGGCAGCTCTCTACAAGATGCTTCCAACTTGGCTTTAAATT
AGCAGAAGCCCCTGCCCTTAGAGCTTCTGAATGTTGCATCCCTTTCTCTTACCTCTCCAACTATGAGGTCTTGAAGACCAAGAT
CTCAGATTTTCTAAAAGCCGCAAAGTCGGTGCGTCCGCTTCCGCACAGTACTAAGCTTCTTGTTTGCATGTTCCCTTGTAAATC
CCCCTCCGGTTATGGCTTCCTCAAACACATCTGCGACTATCAACTTGGCGTCGCCTCTCAATGCTGCTCCTTGATAAAATGCAC
CCAACCTGTACCTGCCCAGCTCGATCAGTACCTCGGCAACCTTGCTCTCAAAATCAATGTCAAACTCGGCGGTGTTACCTCCC
TCTTGTACGAGCCACCATTCCCCGAACAAATGAAGTCGATCCTCCTCCTCGGAGCCGATGTATCGCATGCGGGCAGCATTGAT
GACCCAGCAGTAGCTGCTGTCGTCGGCAGCACCGACTGTAATGCAGTGAAGTACATCGCCCGCATCTGCTTGCAAAAGGCTA
AGATGGAAATCATTCAGGATTTCAAATCGATGGCAGAACACATGTTGAAATTTTTTTCGAGACGGAATGGTGCAGCGCCGGG
GGGAGTGATCATGTTCCGAGATGGAGTGAGCGAGGGACAGTTTGAGGAGGTATTAAACAAGGAGGTGAGGGAGCTAAGGGC
AGCGATAGACAGCGTGTACTCGAACCTGAGAGAGGAAGCGAAGCCAAAGATCACATGGGTGGTGGTGCAAAAGAGGAACAA
CACAAGGCTATTTCCTGCTCAAAAGGAAATAAATGATGGGAATGGCAATGTAAAACCAGGGACAGTGGTAGATCACACCATA
ACTCATCCTTTTAACTTTGATTTCTACCTGTATAGTCATGCAGGAATCCAAAAGAAAGGCACCAGCAAGCCCACCCACTACCA
TGTCCTCTTTGATGAGAACCGCTTCACATCTGAGGCCTTGCAAGAAATGGTTTACAAATTGTGCTACACCTACGCCCGTTGCA
CTCGCTCTCTTTCCATTGCGCCGCCCGCTTACTACGCTCATCTCGCCGCCGCTCGCGGCTGGCTCCTCTACGACGCTCGCCGAA
AGATGAAGACTCCACTACCCTCCGACCGACCGTTTTGTATCACTGAAGCCATCCCCGAACCCTTCCAAGAATCCATGTTCTTCT
GTTGACAGCTTATGGAGTTCGTATTCAGCAGATCCATGTTTTGGAGTCCCG

**SLOW WALKER 2 (*SWA2*):**

GTTAGGAAAATTTTAAATTACAAAATGAAGCAGCAAACTTAGTTTCTGCTGCCATGGCGAACAATAAGGAGGACTTG

AAGAGCAATAAGGAGGACTTGAAGAGCATACAGGCTGACGTCGCCTCCTTCGTTTCCGGCCTCGGCTTCTCTACAGCTTCTGT
CAATGGAGATGGCTTCTATGATGGCGACTTCAGAAAGAAAGGTCGCATCTCTGATAAGAAGCCCCAGAAGGCGCCCCTTAAT
GCACAGAAAAGCTCGTCGGAGCTCAAACAGGCCAACAAGGAGAGTAAAATTCTCAAGAACAAATCAATCACCCCAGAACTT
CATCAATCGCCCATGAAAAGCAATGGAAAGCTTGAAAAAACCAGCTTAAAACAGTCTTCTTCAAAGAGTGAGCTTACAGAGG
AACCCAAGTTTCAACAGGGAAAGGGAAGCAGAGAACTTGACAAGGATGCATTGAAGAAGAATAAGAAGAAGCAGAAGAAG
GAGGAGGAGCAGGAGGAGAAAGCAGAGAAGTTTTCTTCTGAGAAGAAGCATAAAGCAAGAGCTTCTGTGAGTCTTGCTATTA
AGGAAAAACAAGAAAAGAAAGCTAATGTTTTTGGCAGTGCTGATTCTAAGAAGAAACGGAAAGCAGAGAATATGGACAGCA
ACGCAATTGCCCCTGCTAAAAAGCAGAAAGAAGAAGGGGCTCAAGGCTTCGTTGAGAAGGATCAAAATCTCTTAGCAAAGAT
GGACTCTGCAAGAGACGGTTCAAAGACGCTTTCAAAAATATGGTCACTTTTGAAGTCTGGACGGTGGTTTGAAGCATCATCAT
CATTATATTCAAGACAGAAAAGTAATGGAACTCTTGAGGGGGGAGACAAGAAGAGTGGTAAACTTTGTAATGGCATAGAGG
GGAGTGCATTGGTAGCGAAAGCAAGAAGGAAAGGTGACGAGTTGATGGAAAGGGCTGCAGCAGAGTACGAGAAATCTCGTG
GGAAGGACAGTGATATGCGGTGGCTTATGGTAGCTCGTAAAACAGGGACAACCGCCGACAAAGTTGCAGCTTTCACCGTTCT
TTTGCAGGACAATGCAATTGCCAACCTCAAGTCATTAGATGCTCTTCTAGGGATGGTCACTTCAAAAGGAGGAAAGCGACAT
GCTGCAATGGGCATAGATGCACTAAAGGAGCTATTTGTGGCAAGTTTGTTGCCCGACCGGAAGCTTAAGTATTTGACGCAGC
AGCCTCTACAAGTGCAGCCAGGCACACATGAAGCAGAAGAGCTATTATTCTTGTGGTATTGGGAGGATTGCTTGAAACAAAG
GTATGAACAGTTTGTTGGTTCATTAGAGGAGGCAACAAAAGATAACCTTCCTTTTTTAAAAGAGAAAGCTGTGAAGACACTTT
ATGAGCTTCTCAAAAGCAAGCCTGAGCAAGAAAGGAAATTACTCTCTGCCCTTGTGAACAAACTTGGGGATCCTGAAAGAAA
AGTTGCTTCAAATGCTGGCTACTATCTGTCTTGCCTTCTAACTGCTCACCCAAACATGAAGATGGTTGTGGTGCAAGAGGTGG
ATTTCTTTCTTTTCAGACCCCATTTGGGATTGCGGTCAAGATATCAAGCGGTTGTCTTCTTGAACCAGATTGTGCTGAGCAACA
AAGGGGATGGACCGAAACTTGCCAAAAGACTGATCGATATCTATTTTTCGCTCTTCAAGGTCATAACAGCGGGCGAACCCAG
TAATAACCAAGAAAAGGGTAAAGGCAATGCCAATGGGACACAACAAAGAAATAAGAAAAGGAATGAGGATGAGGACGATG
CGGTTGTTGAAATCGATTCTCGTTTACTATCGGCACTTTTAACTGGAGTCAACAGGGCTTTTCCCTTTGTTTCAGCTGAGGATG
TGGACACTCTTATAGAAGAGCATACTCCTGTTCTCTTTCGTTTGGTGCATTCGAAAAGCTTCAATGTCGGTGTGCAAGCTCTTA
TCCTCCTGTACCAGCTTCTTACAAAAAACAACGCTGTTAGTGATCGCCTTTATCGGGCTCTCTATGGGGTTCTTTTATCTCCTG
GTCTGTCCAAGTCTTCCAAGATGGAGATGTTTCTTGGTCTAGTATTCAAGGCCACTAAATCAGATATCAATTCAAAAAGAATG
GCTGCGTTTTCGAAGCGTTTGATGCAGTTGGCCGCCGTGACTTAAAGCTCTCTTTTGAGAAGTTGGACCGACTCTTTCATTCTT
GAAGAATAGTATCAACTGAATGCAAGGATCTCCCTTGTCGCCTTGCAAGGGGCTCCCCAATTTGCGTGTGGATGTCTTCTACT
GTTGTCAGAGATTCTCAAGTCAAGACCGACTCTTTGGCATGCAGTTACTCAGCCTGAAGATGCAGATGAAGAGATTGAGCATT
TTGAAGACTTTAAAGAAGAAGATACAACAAAGGATGATACAGTTCGTCCCGATCAGCTGGATAGTAGTGCCATTCAGAGTTC
AAGTGAAGCTATTGATAATGGAGAGGTGGCTATAGCGCTTGCAGGATGGCCTAAGGAAGGGCAATATAATCCTCGCCATCGG
GATCCTTCCTTCTGTAATGCAGATCAGGCTTGCTGGTGGGAACTAACAGCACTAGCATCCCACGCACATCCATCTGTTGCAGC
AATGGCCAAAACACTGCTCTCTGGTGCAAGTATTCTTTATGCTGGGGATCCGCTAAGGGACCTCACTTTCAGTGTTTTTCTTGA
CCGTTTTACAGAGAAAAAACCTAAACCACGCCGCAAAAGTGAAGCTTGGCATGGATCCTCTTTGTCCGCGCCAGCCAGAAAG
GCTTCACTAATGACGACGCTGCCCATTGGAGATGAGTTCTTGAAATTATCGGAAAAGGAGGTAGCACCAGAGGATGTCGTGT
TTCATAAATTTTACACATCCAAGGTAGATAGGAAACAGAAGTCCAGCAAGGCAGAAAAGAGGAAGGAAGATGCTGTTTCCAT
AACTGGAGAAGATGATCTGCTGGGTGATAATAGTGATGATGATGAAATCGAGGATCTGCTAGACCAGGATGAGGGGGTGGA
GATGGGCTTGGATGATCCCTCTGAAGATGAGAATGAATCAGAGGGAGAATGGACTTACCATGAACCAGATGGTGGGTTTGAA
AATGCAAAACATAACAGCTCTGATGACGTGGATTTTGAAGATATGAATAGGGCAAGCTCGGATGAAGATATACTCTCTGTGG
ACCTGAATGAGCATACTTTTGAGGAAGGCATGAATGAAGTGAAAGAATCTCAATCACTTGTCAATAAAAATTCCAAAAAGCA
CCCAAAAAAGCAGTCAACTTCTAGTAAAAGGGAAAAGAAACAGAAGGAGAAAAGTGCTTTTAATAAGGACGGCTCTAGTAA
ATCTCCTTTTGCCGATCTTGATGAGTATTCTCACCTTTTGGTGGATGAAAGACCTGAAAAGGGCAAGATAGATGCAGTACCAA
GCATGAGGAAACGAAAGAAAGCTAAGTAATAGTAGTATCCCGAGACTCCAATTCTCCTGTGTGTGTGTTTGTTGCAGGGTGTT
CCCTCCTCCTAGTTATCCTAATGACCTGCTATGCAGCGTGTGTTTGTGCTTGTATGCATGTACAGTTTTTTTTTTTT

**SLOW WALKER 3 (*SWA3*):**

GGAGCAGGAGCAGGAGCAGGAGCCAGCCCTCTTCTCCACGAAGGCCAGGCTGCAGAAAAGCTCCACCAGCAGCGAC
AAGAAGAAGAGCGGGCATCAGCTCAACAAGCCTCTCCTCGCTCCCTCTGATGATGATTTGGCGAGCTTCTGTGACCTGGGTCT
CCCCGAGTGGCTTGCTAACACCTGCCTTCACATGGGAATCAAACGCCCAACTCCTGTGCAGAGGAGCTGCATCCCCCACATCC
TCAATGGCCTTGATGTGCTTGGCCTCGCCCAAACAGGTAGTGGCAAAACTGCAGCTTTCGCCCTCCCCATCCTCTACCGACTC
GCTCAACGCCTCTATGGCGTTTTTGCCTTGGTCCTCACGCCCACCCGAGAGCTCGCCTTCCAGCTTTCTGACCAGTTCAAAGCC
CTTGGCTCTGCTATCCACCTGCGTTGCTCTGTAATCATTGGTGGTCTCGGCATGATTGAGCAGGCCAAAGCTCTGATGCAGCG
GCCACATGTTGTGATTGCTACTCCTGGACGCATTGCAGACCTCCTTAGGAATGACCCAGGCATAGCATCTGTGTTCAAGAGCC
TTAAGTTCCTCGTCCTGGATGAGGCAGATCGCATTCTTGATGTTGGGTTTGAGGATGAGCTGGGGACAGTTTTGCAGAACCTG
CCTTCTCAACGACAGACCCTATTGTTTTCAGCTACAATGACCCCTGAACTTAAGGCCTTACACGAGTTGTCTGGGGACAATGC
TTACTTCTATGAGGCATATCAAGGCTTGCAGACAGTTGAATCACTACAACAACAGTACATTTTTATGCCTTTGGATGTTAAGG
ATGTGTATCTTTCATATGTCTTGTCTATTTTGGAAGACAAGAGCATTCGGTCTGTCATTATATTTGCTTCCACATGCAGGACTT
GCCATCTTTTGAGTTTAATGCTGGATGAGCTTGGAATTCATACTGTAGCATTGCATTCTATGAAGTCACAACCTCAACGGCTG
GCTTCTTTGAATCGTTTCAAATCGGGGCAAGCTCCTGTTTTAATTGCTACAGATGTTGCAAGTCGAGGCTTGGATATCCCAACT
GTTGATCTAGTCATCAATTACGATGTTCCCAGGTTCACTCGCGACTATGTCCATCGTGTAGGTCGTACAGCACGTGCAGGCAG
AGGTGGATTGAGTATCACTTTAGTGTCACAGTATGATGTGGATCTTGTGCATGAGATCGAGTCACTTATTGGAAAGCAGCTAG
ATGAGTATGAGCTAGAGGAGGAGGATGTGTTAAAGGGCATTACAAAGGTTTTCAAAGCCAAACGCCTGGCTTCACAACGAAT
GCAAGATAGTGGGTTCGAGGATACGGTCAAGGATCGAAAAGAGAAGAAACGAAAGGTGCTCAAGGAGAAACTTGGTTTGCA
AAAGAGGAAGAAAAGGAAACAAAGTGAAGCAGCAAGATGACTTTACGTGTGTACCATTGCTGTGCCATAGGATTAGCTAGA
AGGTTGAGGATTTAAATATAGGAAACCAAAAAGAATTTGATTTGAGTGGTTCAACAGGCTGGACATTTCAGTAATTAGATGA
ACTGGCTATGATGAATTTGTATAATTTTTTCGTTGTATTTCTCAGTGGAGAAATGTGGATGGCATTGTTCAAGGGAAAAAAAT
CACTGTCCTTAGCATGCTATTATTTTTCTAATGTTTCTATACAATCTTTGTCTCCACTAGTTGCATCTTAA

**CHROMATIN REMODELING 11 (*CHR11*):**

TCGCTCGAAGAGATGTGTTCAAATTCAACCCAAAAATATTACTAGTATTTTCTTTAAACTTCACCAGTCAAAATTCTGT
AGTGCCCTTACAAGTGGGGGAGAACGCCACAATTTCAAGGAAGTCCCATCGGATTTTTTATGGCTCAAAAAATACTGTCGAA
CCAACTTTTCAATTGTGCTCATCTCACCAAGGGTTAAGTTAATTTCTTTTGCCGAGGTCAACAATGTACAGCTACATGTCAAAT
CCCTTCCCAATCTGAGGCATGCACATCACAACCATCCAACGCTGAAACACGATACCCCTGCACTTTCACGGAGGTCATTGACC
TTTCATCGAACTAGAACGATTTATGTACCGACTTCAAGGTATCAATTGAAGTGCATATCGAAGGAGTGCGCTTATTTCAATAA
GGATGCAACAGGCAACTCCAGGAACTAGATTTGGATTGAAAATGAGGTTGCATAATTATCCTTCAACAGTAAGGTGCCTGAG
AGGTTATCCCCTATTCTTGAAATAACTTACAAAGCATCCATCTGAGATTCCAAACGTACCTCCACTTAACAATAAAATGCTGT
GGCCACGTATTGAAGCTTGATAGATGCTGTTTGTTAGCGTAAATCCTGCTTCAACATATAAGCTGCTTACAAAACTTGAATGC

AATAAACTACAGCAGCAATTTCTGACTGATTAAAAACATAGTGATTGACACTGCAATCAAATACTATGTTGATCCAAATACTA
TGTGCCCGCAAAGCACAGGAGCGTAATTTCTAGGAGGGCTTGCTGGAGATGTCAAAGTATTCAGCACAAGATTGTAAAATGC
AAGTCGATGTAAGCAACATACTTCTAATAGATCAAAAAGGGCAAGTGACATTAGTAAGATTACTGCATATAGCCCAAACATG
GCATGGCTACCCCATCCACCAAAAGAAAATCTTTGAATTTGATTTCCCCTCTTCCCCTTGGATTAATACTTCCAAGCAAAAGAT
GGGTTTTGACCATACATTCCTTCCATCAATGGCACACCACGAATTTGTGTAGGAGAGCCATTAATGATCCAAAAAATTACTTT
TCCACAATTTTACTAATCTTTGGGGACCTCCATTCGTTCTGGACTAGTCCGCTTAAACCACACTGCGACCATCTATTGCCCCCT
TCATGGACCACCAAAATGCATTCACAAGCGTCCAAGGGTAGCAGAAATGTAGCCAAATCTCCGGATCTTCATCAGATTCACTC
GAAGCTTGCGAAGCAATTTAAAGAGCGATGAACCCCAGAATTTGCTGGACTAAACCTGGAAAACACAGGCAGCACATTTCAG
CTTGCCTTTCCTCTCGTGGCTTCGGTTCCTCCAACATCATACCAGAGTTGAACTGAGCTTCATTCCACTCATCTCAAAACTTAT
CCTAATCAATGATTACCACCTCCTTAGCATTGATACACCTTGTTCCATTGAGTCCATTCAGGGTTTCCAGTGAAGGCTTCATCT
CACGATCATCCCATGTGACTTCTGATTTGTGTACGTCCTCCAGGGACTGGCTAAACTGCTGCACTGTCTTCACTAAGAGCTTGT
CATCCTGTAAATCCTCGATGCAAGGCTTAGATTCATGAACATGAGATAGATGCACGCTTCCCTCCGGCTTGTGAGATTCCACA
GACGATAGTTTGGAGGTGGGCTTTATGTCTTTCGTCTCCAATGTATCCAAATCTGGCATGTCACCTGCCTGTGATCCAACAGCT
GGGAACTCAAAGTTCCCATTCACTTGATGTTTTGCTAAATAAACGCAGTAACTACAGCAGTGTATGAAATAAGCCTGTGTAGG
ATGAATTTGCCCAAGATTCTCAAAACGTTCACTCGTCCCTATGAGTTCCGCCTCTGCTGGAAGGTCATCTTCACAATACGCATT
AGGACACATTTCACACCGAAACAAGAGACCACCAACAGTGTGAGCCTTCCTCGCACAAATTCCACATGTATGATGAGGACAA
AACCATTGCAACTTTGAAACCTGGGCTAGTTCAGCCTTGGAATAGCCAAGGCACTCTGAGTGGTATGCAGCTGGACACCCATC
ACAACAAATTAACAAACCCCCGTCTTTGCACAACAGACAAATATCATCATGTTCATAATCAATGCCAGCCCTTTGAGCTCTCT
TTTTCTGAGTCTTGAATTCCTGGCTATCATTCAAGTATTCCCGTCCAAAAACACTTGGTTCACCTTCCTCCAAGTCGTACAGAT
TTTGACGACGAATAGCGAACCCATCCACTTCGACAGTTGTGGAAACTCGCTTGCGCTTACTAGCTTCTATCCATTCCTCCCCAA
TACCCTTGTTTGACGTTGGTGTATAGTCAACACCCTCAAAGACTCTAATGGCTGTAAATGGATCCATCACTACTGACTCTGTCT
GCTGAATTTTTGAGCACTCCTGCTTTCCTATTTCTCCTTCCATGGTTGATGGGGAGCGACTTATTATCTTGTCCAGATCTTCGTC
GGTCAACTCCTTGGTGTTTTCTGAACTTATGAAGGTCTTTGCTCCAAAGGTGATCGTAGCCAGCAATTCAAACAAGTTACCAG
GTCCATTGTCCTCACTCTGACTCATACCACCATTGTTGACAGCATGATCAAGAAAGAGCTTTTTCTCAGACCGTTGTTGAATTC
TCTCTTCCACTGTACCAGCTGTAATGAGTCGATACACATGCACAGGCTTTGTTTGCCCAATTCTGTGAACCCTAGCCATGGCTT
GCAAGTCCACCTGCGGATTCCAATCACTGTCAAAAAGTATGACCGTATCTGCAGTCTGAAGGTTAATCCCAAGACCACCAGCC
CTTGTAGACAGCAGAAAGATGAATATGGGACTAGAAGGCTTATTGAACATATGCATGTCAATAGTACGCCTTACGCGATTGG
TGGAGCCATCCAATCGTCTGAAATCAAAGCCACGGTATTGACAATAATCTTCAAGAAGGTTCAGCATCGTTGTAAATTGAGA
AAACAGCACCACTCTATGTCCGGTGCTTTGAAGCTTCAACAACAGACGATCCAACAACTGCATTTTTCCTGAGCTTGTAATAA
TGTCGTCGCCTACACAGCTTGGATCTTCCTCAGCTCCTGGAAATAAATACGGATGATTTGCACATTTTCGAAGCTGCATTAGC
AAGCTCTGCAATTTTTTCCATGCATCCCCGTCCTTGCTTTTTTTGAAAAGTGCTGCTGATTCTGCCTCTAATTGAGATAAAAGT
CTGCTATTTTTCAGAAGCAAGCGCCTATACCAAAAATGTTGCATTTTTGACAGAGGACTATAGATCTTCAATTCTGTTCGGGG
GGGCAATGACTTCTCAACATCCACTTTAAGTCGCCTCAAACTCAAGTTCTTGAACAGCAAGTGCGCTTTGTCCAGAGTCTTAA
AGTCACATTTGGATTGAACCATACTGTAAATGTTATCGAACAACTTTGGATCGGTAAAGATATCTGGAATAATCTGGGTAAAA
AGGGCCCAAAGCTCATGCAGATTGTTCTGCAAAGGAGTTCCAGTGAGCAGCACAAGGCCCTGGCGATAGATGTGGCGAACTG
CATCTGAGATCAATGCATTTTCATTTTTTACCTTATGTCCCTCGTCCAAAACTAAATATCGCCAATGAATTTTTGAGCGAAGAA
TCATCTTCATGTTTTGAGAAGCCACCATCTCATAAGTTGTCACAATAACATCAAAAGATGATGAGTTATTGAGCACCTCCCTG
CGAAACCGTTCTCTCTCTGCCTTGTCTGATGAATGCAGCCGAATAACTCGAAGTTCCGGGCACCAATGCTGAAATTCTTTCATC
CAAGATGATAAAACTGAAAGAGGACATACTACCAAGTGTGGACCTCCCACGTGCCTCTCAAACTTGAGGTACCCCAAAAATG
CTATTGTTTGGAGAGTCTTTCCGAGCCCCATTTCATCTGCCAGAATAGCGCTGAGGCCATTGTCAAACATGGATACTAAAAAT
GAAAGACCTTCTAGTTGATAGTCTCGCATTTCTCCATTAACAATACACTTAGGTTGCCTTGTAATCTGTGTTTTTGGCATTGGT
TGTTTGAAAGGAAGCTTCTTGACAGATTCCACCAGCTTGTTCTTCACATCAGATGTAATGAACGGCTCAAGTACATCAAGTAT
GCTAAACAAAAATACATTGGCTTTATTTCTGGCAACTATCATGTCTTCACTTATCTTCTGATTGTTGCTTCCCCATGCATTCTCA
TGTACCCTTTTCACAATTTGGTCAGCATCCTTCTTGCTTTGATTGCTGATTTTTCTTTTCTTCTCTGAGACCACTTGAACCTCTTC
CTCTTGAGCTGCCCTATACACGCTAGGCTCACCTTCTTCTAATGTGTAGGTGTTCAGCCGCAGAACCGGTTGCCCGTTGACGTA
GGTTATCCTGGACTTGCGAGAGCGGTCACTTCGTCTAGGTTCATCCATTGAGGTTTTTGTACAATCATCCATCACACTATCACC
AACACCGGGCATGTGGGCGGGAAGAAAGAGGGCGTGCAGAGGAGAAGCGGAGGGCAGCACACCTGCGCGCTCTTTGCTTCT
GAAAGGCCCCCCGGGCGGTGTAGGTGTGTGTATGCGCGCGCGCGCGT

**ACTIN-RELATED PROTEIN 7 (*ARP7*):**

TAAATTGAGCAACTAGTCCTCTCACTTCTCATTAGAACTAGAGTGTTATTGATGGAAGATCTGTTCATAAAATTATGTT
GAAATGGTGGTTTTTTTTGCCACATGTTTCCCTTACAGCATAGTCAATCTCACAAATAATCAATGCAGCACATGAGTGAGACG
TCCAACAAACACTCCGTACAATTTCATGGCGCATTTGCTCCACTACCTTTTCTTTAAAATTGGTCTAGCTGTTGCTTTGAATCA
GAGAGCTTTTGATACACATGCGCTGATCGCAAATGAGAACTGAGACAAACTAACTTTAATAACTGCTCTTTGAACCCCATGTT
ACTCAAGGGACCTCATCTCAGTAAATGTCAAAAGATAACTATCTCCTCATTTAAAATTACTATATCCATACTTGAGCTTAGCA
AACCTTTGTTGAGCAGCAAAAGCTTGGATGCAGCAGCCTTCCTTACAATCTGAAGTGTGAATCCGCAATCGTGCCATGCATAG
TCCTGCTATCAATAGCACTTCTTGTGAACAATGCTTGGACCCAGCTCATCATACTCAGATTTTGTTATATGCTGGTTCTGAGGA
AAAACGACCTTCGCAAGAATTGCGCCTCCCATCCATGCAGAATAGCGTAAAGTGTTTTCTGGCATATATTCAGGAGGCT

**HOMEOBOX PROTEIN 5 (*HB5*):**

TCTCTCTCTCTCTCTCTCTCTTGTGTGTGTGTGTGTATGTCTCTTTGTGCGCTCCACCTATGAAAACCATCTTCACACAT
CCTCTTTTGTAAGCTCTTGTTTTCTAATGAACCTCAGTTGCAGCCTCCTCTTTCCATAGCCAGTTTCTCCTTCTCCTCTGTCATTT
CTTCTTCTCCCCTTGTGTGTCTGCAGAGCTGAAGTCCTCTCCATGGCTCTCTGTTTGGAGATCCCCATTGGCTTCAAGATGGCG
AAGCAGGTTATGCATCTCCATACCTTCCTCTCCAGATATCTGCACCATGTCCCCCTCTTTGCTATCTTCCTTATGTGATAGCCTC
TCCTCTGTGCAGGTTTCTCTCTCCTCCATAGCCCTCACCTTCCTCATCTCTCTGCGCCTCATTTCTCTCGCTTTTCTCCTCTTCTC
TTTTCTCTCTGTACATTCGGTAGGAGCATACTTGCATAAGTTAACAGAACAATTGATTGAGGGGCCATGGAATGGAGCGCTGC
ATCTGGTTCTATGAAGGCGCTTCACGATGACCGCGATGATACTCTGCTAGGGCTTGTCTCTCCTGGGCACTCCTCAATAACTCA
AGAACACAATCGCAGAGGAAGTAAGAAGCGATCTCACCACTACGCAGTAAAAGATGAAATTGATGAGGATGATTTGAGCGA
TGAGCTCTCAAACCAGGCCGAGAAGAAGCGCCGGCTCACTGTTGAGCAAGTGAAGTTTCTTGAGATGAGCTTCAACAAGGAT
CTTAAGCTTGAGCCTGAGAGGAAAGCCCTACTTGCGAAGCAGTTGGGCATACGTCCTCGCCAAGTTGCAATTTGGTTTCAGAA
TAGGCGAGCCCGTTGGAAGAATAAACAGCTTGAGCAGGACTATGAAACCTTGAAGGCCAAATGTGACTCTATTTTGAAAGAG
AAAGAGATTATATTGCGGGAACATGAGGCAGTTATGGGTGAGAAAAAACGACTCCAGGCTCAGGTTGTTCGTCTGATGAGTT
TGCTTGAAAGCCCAGGGGATAATGGCGTGCATGATGTAATGGGAGTGGGCTCGGAGGGTAAGTCAGAGCTTACGTCTCCTAC
CAAATCAATCGATGACCAAAGCGATATCCAAGATTCGGGGCAAAGCTATCCAGTATCAGAGGGAGTGGCTGGGGTCGACATG
ATTGGTCCATCACATGTAGAAATTGATCCTGTGGGAGCTTGGACTATGAATGTGAAGATGGAAAGCGAATTTACGCCTTCTTT
AGCTGCTGATGATGTTTTCGGGAACATTCCGCCGCTCTTCCATCAAATTGCGGCTAATGCAATCTATCTAGAGGATGCATTCTT

CTATAACTGTGAGGATCACTTTACTGGGTTGGCATATTATGGCTGATGACAGTAAGCCCTTTAGTACTATACTATTATGCCTAG
ATTAAGCATTATGCATATGAGAAGTAAAGCATTATGCATATGAGAAGTAGATATTTAGAATTATTCAGCCTATGGTCAACCCT
TCAATTTTTGTCAGCACTTCTTGCAGGTTATGAATAAATGCGTGGTAGGTTGACATGATCAAATTCATGAAGATCACCTGCCA
CCATTTCCTCTATGACAATCTCTTACTGGTTAAGGCTATCCTGTAGAAAATATTGTACAATAGATGTAAATATTATTTCGGCTG
ACACAAGCTGGTTCTAATTAAGCAGGAAACAGTCTTGTTTCAGCAATATGCAAAATGAGAACAGGTCTTTGTATGTTCGCTGT
GGCGCATTCTTGTTAGATTTAGTTGTTGTTGAAGCTGACATGAGATATTGAATGTGACATGAAACTATTTGATTGTGCTCATGA
TATCAATACGAAATGATATCACAACTCTTTGGGGGAGAAGATTGCTAGGCAGAGGTTTGTTGTTAAGGTTGTTCGGCTTTTGT
GTAACATGGATGTGTAAAGTTTTTCACATAAGTCTTCAGTACTGAACAGCACTATGATGCTTGTAAGTTTATTACAG

**EMBRYONIC FACTOR1 (*FAC1*):**

GAAAAGACAAGCACTCACAACCTACCAAAAGGAGGAGACCGTCACCCACCTTAGAAGGTGACCGCTTTTTGAACAGA
AGTGCGCCAGTTTTCTTAAGTTTTTAATTCCTTTAGTCTAGAAGATGGAGAGTCGAGAGTAGCTAGAGCTCACTCGCTACCTC
GCACAGATTGAACATTGCCATTCTCCTCACTTTGAGGTTGCCATCAAACGAGCGATAGTTTATATCTACTAGCTTCGTTCACCT
GCAAAAGAGCTTATCAAACAAAGGCTATGGATGCATCTACAAACTGGGACACTTTGGTCAGATTCCTAGCAACAGCTATCGT
TGGTGCGGCGCTATTTGCTGTGCCCGCAGTTTTGTACCACAACCGGGTTGTGTATGAGCTTAAACACCAAAAAAAGCACCACG
GAGCTGACGACACACGTCGCCATCAGGGTCCATTCAAAGCTTCACATCGTCGGGTTCGAAGCCGCTCATCAAGCAGAGAGAG
AAGTTCTGCATCTCTCCGTGAGGTTGTCATGCTACATGAAAAAGATGGCCAAAAACCGCACCGGACCTCTAAGTGGCCTGAG
CAAGATAATAGTGTGGATAGTAAACCTAGCTCGCAGTTGCCTTTCAATACACTGAGCTCAATACCGCCTGGGCTTCCACGGGT
GCAAACACAACGGGAAGGTGCGAACAATCATCTGAGTCCAACCAGACGATTTTGTCCTTTATCAAGTACTGTACTTAGGCCTA
ACACACCAAAATCTCCAGTTGGTGCCAGTAGCTTCTCGGACGAGGAAGACATTGCAAATGACCAGGATGAGCAGGCCTTCGA
ATACGAAGAAGTGAATGAAGAGTCTCTTACCTCTAATGCAACCACGGTGAAAAAGGAAGAATGTGCAAAACCTTCTGCTGTG
TTGGCCATAGACCAGTCAAATGGATCTGGTGAGCGAACAGGAAATGTTGTAAGTAGATCACACAGCATTCCTGGAGAGCTTC
ATGGCATTCATGCACCGGATCCGGTGGCAGCTGATATCCTCAGGAAAGAGCCAGAGCAAGAGACTTATGTACGTCTACAAAT
TGGGCCAATTGAGGCTCCATCTACAGAGGAGGAGGAGGTATGTAGAATGATGCAAGAGTGTCTTGCTTTGCGGCAAAAATAT
GTTTTCAGAGAGAGAGCTACACCTTGGGACAAAGAGCTGATAACAGATCCCAGTACTCCAAAGCGAAATCCAGATCCTTTCT
ACTACGAACAAGAGCCAGCTTCAAAGCATGTTTTCCAAATGATTGATGGGGTGGTTCATGTCTATGCCAATGCAGAAGCGGA
GCAAGATCTTTTCCCTGTATTTTGTGCAACAGAGTTTTTCACTGATATGCATCGAATTCTGAAAATAATATCACTTGGGAATGT
TCGCACACTTTGCCATCACCGGCTTCGTTTGCTAGAGCAGAAGTTTAGTCTACACCTGATGCTCAATGCTGATCGAGAGTTCTT
GGCTCAAAAGAGTGCTCCACATCGAGATTTTTATAATGTTCGGAAAGTTGATACTCATGTACATCATTCCTCATGCATGAACC
AGAAACATCTTCTACGATTCATCAAGTCTAAGCTGAGAAAGGAACCTGATGAGGTTGTAATTTTCCGGGATGGAAAGTACCTT
ACACTCAGGGAAGTCTTTGAAAGTCTGGATCTAACTGGGTATGACCTGAACGTGGACCTTCTGGACGTTCATGCGGATAAAA
ATACATTCCATCGCTTTGACAAATTCAATTTGAAGTATAATCCATGTGGTCAAAGCAGGCTAAGGGAAATCTTTTTGAAGCAG
GATAATCTGATCCAAGGGCGTTATTTGGCTGAAGTAACTAAACAAGTTTTTCAAGATTTGGAACAAAGCAAATATCAGATGG
CTGAATACAGAATATCAGTTTATGGCAGAAAACAAAGCGAATGGGACCAACTAGCTAGCTGGTTTGTCAACAATGAGCTTTA
CAGTGAGAATGTAGTGTGGCTTATTCAGCTTCCTCGTTTATACAACGTTTACAAGGAAATGGGAATAGCTCATTCATTTCAGA
ATATGCTGGACAATATTTTTATACCTTTATTTGAAGTGACCATTGATCCTGGATCTCACCCTCAGCTGCATGTTTTCCTTAAAC
AGGTGGTTGGTTTTGACTTGGTAGATGATGAGAGCAAGCCAGAACGTCGGCCCACAAAGCATATGCCATCTCCTCTCGCTTGG
GATATTATTTTCAACCCTGCATTCTCATATTATGCATACTACATCTATGCAAATCTTTACACCCTAAACAAGCTTCGGGAAACC
AAAGGTATGCCGATCATAAAATTTCGTCCTCATTCTGGTGAGGCTGGTGAGCAAGACCATTTGGCTGCCACCTTCTTACTTTCT
CACAATATAGCACATGGTAACAATTTGCGCAAGTCGCCTGCTTTGCAATACTTGTACTACTTGGCTCAGATTGGGTTGTGCAT
GTCTCCTTTAAGCAACAATTCCCTATTTCTAGATTACCACAGAAATCCCTTTCCAATGTTCTTTGCTAGGGGCCTGAATGTATC
TCTTTCCACTGATGATCCTCTTCAGATACATTTAACAAAAGAACCTCTTGTAGAGGAGTACAGCATAGCTGCCCAGGTGTGGA
AGCTTAGTCCTTGTGACTTGTGTGAGATTGCTCGCAATTCAGTGTATCAATCAGGTTTCACACATGCTTTGAAGTCACATTGGG
CTGGAAGCAACTACTACAAGCAAGGCCCTGAAGGGAATGATGTGCACAAGACTAATGTTCCGAATGTTCGGGTGGAGTTTCG
GCATGAGGTGCTGAAGAATGAGTTACAGTATGTGTATCTTGGAGCAGCGCCAATTTCAGATGTGATACAACCATAGGGAGAA
ACCCTTTTTACGATATCTTGGTTTCAGAACTTGAAGGTTGTGGCATCCAATTTTGATATGCCTATGTTGGTGCTTGTAGTCTGTT
TAAAACTTCTCGTCTGCAGAACTTGATCATGAAAAATGGCCAATTTTTATGTTTCATGGCCTTACGTGTGTTAAGATGAGAAA
AAGTAAGCATTCATTTGTTCTAACTGGTTCTCCATGATGTTTAGAGAGAGAGAAAGAGAGTGGATAAAAAAAAGTCCTTATG
CAACCTCTCTTTTATGCAATTGGGAAGTGGTTGCATCTTCTCTTTAGTGTATTCTCTCAGATTGGAAAGCATACATTGGTGTAT
GGCTTTTCCACACCTCTTGCTATTTCAAGAATCCCGTAAAGCCTGCTCACTCTCTTCAATAAGGATAATGAAAGTGGATGATGT
TGTCAATAACGTACGCTATATTTTGACATTTTAGTGGAA

**GRAVITROPISM DEFECTIVE 2 (*GRV2*):**

TTTTGAGGAAAATAAATCAATTCACCTTTCATAAATCAATGTGTCAAAACTGTTACTTATGATTTACCATTTACAAAAT
GTCAAGCTTTACAAGCAGAACTGTGAGGGAAATTGAAGCCAACACCTACAACATGAAACTCATTTCCCAAATTTGGGAGATA
TATAACATTATTTACTGATACAAAAAGAGCTGGCCATTGCTGCCAGCTACCCAAACAAAATGCAAAAAGCACAGTTTGGGTC
ATTTATTGTCTTTTCAACAAGGATAATTAGCCATTTGCAATTACATATTTACAAGATGGATGATACACGATCGCAGAAAAAAT
CATTATCACAACCTAAAATGCACATATACCTTCCAAACAGGCATATATTCAACCTTAGAGGCTACCAATCTCTTCATCCGATT
CCTCTGCAACAGCCTCCTGGAAAGGGCCATTGGTTATGGTCGAGCGAAGTAATGGCCGCGGGGTCGGAGCAGGAAGTGCATA
CGTCAAAGCCCCAGATGAACCCTCAATAAGGCCTGCAACCCCAGCTGCAGCAGACTGCGCATTTGTTGGAAGGAAGAGATCG
TGTCTTTGATCCTTGTAAGCACTCCAAACATCTGATGCACTCAAAATGTCTTGAACTTTAGCTGAATGAGCTCCCTCACTGGCA
AAAGCATGCAAGACCTCAACCGCCAAAACTCTTCCAACAGAGGCTTCAGACTCGTTCCATTTCATCTGTGCACAGAGTCCATT
TGTACCACCTGATCGCCAATCAAGCAGTCCCAGAAGCACCTGCACTAACCCCACCTTCAACCCCTGTGCAACCAGAGCATCTC
TTGCCCGATTTCCTGCAACCACTACTCTTTTCAAGGTCTCTAGAGCAAGCACACTGCCACCTTGCCATCCAATGGCTTTCATCA
ACAGTGGCACTACCTGAGGTGAACCTACTCCGGTGGTTGCCATGGCTTCAGCACAAGCAGTGCTAGCAGCCAGCTGATGCAA
GACTCGCAAACAGCTTAACCTCACCCTTTCTTGGGGGATTTGACTTCCTGATATTTCTCCAGCATCCTCTGTAGTCAAGCACTC
CTCTGCACGATCTCCCATACGTCCTTTTGCCCCTGACGACATTGTTTCTCTCCTGCTTTCATTTGCAGTTGATGCAACGAGCTTG
GGAATATACCCTAAATGACCCACATGATCAGCTAATGTAGGATGCACTCTCAGAAGAGAGACTAGGGCTGCTGACAAGAGCA
ATGGGAGCTCAGGGTCTGCAAGAAGAGGTCCTTTTGCATGAGTTGCTGCAACTGCAGAGACATACTGGTCAAGTAGGCCCTC
AAGAAATCGTTTTGGATTTCGCAGAGGAAACTTTGGGTCTTTAAGGAACAGTCTGACATAAACACCCCCGACCTGGGGTTCAT
CACGCATCTCCTGTTGGCCAGAACCTTGCTCAGGCAAGTCCCATTCAAGAACACTTCCCTTCACTTGCTCGCGATAAAGATTA
GATGCCATGGTGGCGACCTGTGCACTGAGAGACGCTGCCATAGCAGGACTCCAGACAAGCTCAGGGGTTTCTGTAGTCTGTTC
CAGGGCTGCAACTACCGCATCTCCAGGTCCATCCCGAATGGCAGAGACGAGCCCATCGGGAAGATACCTGGCAAGAGTTATT
GCCACCCTAGGACCATGCATTGGCTGCCCAACAAGCTTACCCAAAAGAGATGCAGCAGCTGCTCGTTGTTGAAGGGGAATCT
CTTCTTGCATGGGCAGCAAAAGTTGAAGTATATAAACAAATCCTCCATGTTTTGCAGTAACCCAAGCCAGCTCTGAAGTACTT
GCCAACGCAGAAAGAACCCTTAAAGCCCCTGGCCGACATACCGGTGCACTATGTAAAAGTTTGAGTAGAAGCAGCAGGCTTG

ATCGATCAGCAACCATAGCCTCAACACTTGGGGCGTGCGTTGTCAAAACAGAGAGAATACTCAAACACAACTGCGGCACTTC
CCCACTTGAAGCTTTTGTGTCCACGAGGCATTCAAAAAGTGGGATCAGATGCTCTTTGGAGCTAAAAGTTGAAGCCAGGCTTG
GGGTGGCAGTCAGCAAATTCTGAAGTGCTGTTAGCCCAGTTGTGAGATCCCTTAAAACGATCTCTTCGTATGACAACTTATCA
TCAACTAGCTTGCTGTGTGCGTCTTTTCTTGACAGCTCTACTTCAGAACTTTCATCACCAGTATCAGCCTCTGACGATTCCTCA
GATGAAAGTTCTGACTCATGGACCGGATCTAACTCTGCTGTCAGCTTTGCCAGATAAATTTTTTTCCTTTCATCTACCAACTTC
GAAATAAATTGCAATAAAGAGTCGCAAAAGAGCTCCGGCTGACTGAGCTCATAATCAGGCTGCTCATTGTAGACTCGTAGGT
ATACTTCACCTACATGCAACTCTTTGGCCAGGGATTTATATTTAAATGACTCTGCTTCTTTCATTTCATATGAACCATCTTGAC
ATTGAGAGACACGCTGTTTATCCACAAACTTCAAAAGCTCGCCCCTTGTTGAAGAATTCCAGATAATCTCTGGCGATTCCAAA
TTGGCATTAAGGTTGCAAAGAAGCTCCTTCAAAAGCTCTGTGGTCAGCATGGCTGCTAGTTTGGGAGTTAGCAGAGCTTTCAA
GGCATTAGCTGCAGTCTCATTATGAGGTGTCGAGTATTCGTCTGTCAGGGATCCACGCAACCGACTGAAAGCACGTGCTGCAA
GAAAAGCATGTGTATTTTTGGCAATTTGCACTCTGCTCCCTACACCATGGGACTCACTTGACCCATTTTCTTCAGCCGTTGAAT
CGTACTGCAAAAGAAGTGGAAGTAGGTACCAAAGTACCCCAGCCTTCAGCATGCAGTTTTGCAGCTCTTCTGATACAGCCAC
ACTACCCATAGTGTGAAGGGCAGCTTCAACAGCTACAGGAGCATGCTCTAGCTCAGAGCAATGCACAACATCTTCTATAAAT
CCAGGACATCGAAGCATTTCTTTCCATGCACTCCTGAACTTGCTCAAGCCGGCAAACGTGCGCATGACATTTGTCACAATTTTT
GCTGCAGGATCTGTGCCAGGGGTGGTACGCTGTACTATGTACATGCATCTCGACAGCAACGTGGCCAGTAAAGACACCCCGC
CATCCCGTACTAGCTCCTCACCATTCAAAGACGACGATACACATGTTAGCCATACGAGCTCTGAAGCTGCAACAAGTAGTGG
AGCCCTTTCTGTAGAAAGAAAATTGTTATCATCTTCATCAACTGTGATACAGTTGAAGAGCATTGAATACCCAGCATATTTGA
ATGGCTCTAAAACATTCCCATATCTTCTGTACAAAATGCATTGGCCTTTCAATAGGAGTAGTAGTCTCCAAGGCTGAGGCCCT
TGGAGTCCTTGTAAGGTGGCCTGCAAACACTCATAAGCTTTTTGAACAGCAACAAACTTCTCTCTTCCTTCTGGGTTTTTGTCT
GGATGATACTTCATGGCAAGCTTTCGGTATTGCCTCTTCAATGTTTCTTCATCAACACGGCCAGCCAGTCGATCTCCAGATTTT
CCATCATCCACAACATGCTTAGCATCTTCATTATGCATGTCTTCTTCTAACGATATCTCCAATATTTTGCACGCTTGCTCCTCAG
AAAGATCCATAGGTCTTCGTGTGAGCTCCTCGCGCCACATGGTAAGTAAGGACTGGAGAAACTCCACATGCTCTACAATGGG
CCATTCAGGGAACCTGATTTCATCACACAGATTTCGAAGATAATAACGATGGCACCACATCTCATCCTGAAGCTCTGGATAAG
TGACTGGAGGTATAGGTGCATAGTCATATAAAGAATGGCAATGTTGGGACAACTTCTGAGTAAAATCTCCAAGATGCTTAAG
AACCTGATCAATGAGTCTCTCTGCCCTCATCTTGTGTGTCCATATTATTTCCGGAGTGTCTGAATCTGCAACCATTGCAGAAGC
AAAGGATGCAGATCCAGATCTCTCAAGCACGTAAAGAAGACTCTCTGGCAAAAGTCCACCAAGTATGCTCCGCTTTGCTAAC
GGCAGTGATGAAGATAAAGCTGCCTCAGCACCTCCATGAAAGGCTTGGTGGGTATGAGTAGCTGCAAATAGGTTTGCCGCTG
AATTCAGATTTGACCCAGCATAGGTTAACATGAAGTAGAAAGCACCCGTGTTATAAAGTCGGGCCATCGCTTTTGGATTCCTT
GTTACCACAGCCTTCAAAAGAGATGCTGCACCCTCCACAATAACTGGCTCCCCTGTCAAGATGGCCTGTGCAACATGAGGAA
GACAGCGAGAGCTGGACAAGATACGCTTCACCCTTGGAGTTGGAGTCACAAGTTCACCAGCATCATCAAGGTCTGAATGCAC
ATTTACCATGCTGTGCAAAATTGACAAAGCAACCTCCCCAACCTGTGCTGGTGTGAGCACCGGAACTCTAGAAGCTAAAGCC
CACCGTAATTCACGGATATCACTGAGCTTTCTCCACTCCAGCATCCCTGAAGCCCAACATTTGGTATCCCAATCAATTTCCTGC
TTTGACCAAGCCCTCCTGATGCCATCCTTCTCCATCGGGCCAACTTGCACTCCATCCTTGCTCATGTAATGCCATTCTTTCTGTG
GCTCCACAAAAGCTGTAGCTGCTATAAGATTTGACTGCAAAGGAATTGAGGTTCTTTCTGCCGTTTCGTGCGCTGTAGTCAAA
AGATCTACAGCTAGTACACAACCTCCTACAGACACGCATGCCTCTGCATTAGAAGGCACCTTCATGAGCACCTTCAACAGCAG
TAAAAGTCGATGTCGAAGCGCTCTATCACTTGTCCGATCCAAAAGAACTGTTATATGGGCAGTTCCATCAAAAGGGCCTATAA
CAGCATAATGCTGCTCATAGACGATTGTCATTGCTCTGGCACATAACTCCCTTACTGCAGATCCACCTCCTCCTCCAAACCCAT
CAAGCTTGCCCACATCACACCAATCCTCTAATGTACCCATTTCATCCCAATTTGCTCCATCCACTTGAAGTCCCATATCTGCAT
CACATAAGAAGCGGTGATATAGTGCGCTGAAAAAAGCAGCTGGATCCCTAAGAGGGAAATCCTGAGCCCGGCCACCCAGTCC
CCCACCATCAAGTAAAAGCCTGAGATAATATTGTCCAACACAGACTTCCTTTGAAAGGCTTGGGTACCTGACCCCAAACTCCA
GGTAGTTCCAAGAGAGTTGACTAGAGACCAGTTCTCCTTGTTTACTGTCACTAGTCCCATATCCCAAAATATTGACATCTTCTG
TGCGTTCCTTCTCCACATCTAAATGGTGAACCTCAGTTTTCAATGCTTCTTTAAGCTCTTGTCGAGTGCGCTCATTCCAAATTA
GATCTGCCCGACTATGATCAAGTCCAAACTCTCGCCAAAATTCTGGCCAGTTACAGAGAAGTCTGCCTGATCCAACAGGTGTG
CTTTCCATCACCACTTGGGCAGGGGCTGGAGCACTAGAAGAGAACGGTAGACCCATGCTAGACTCACTTGCAAATGCATCAA
TAGAGTTTGGATTAGAGAAAGCAGTAAAATCTGAAGATAAGGGTGGAGTAGCCCCATCAGCTGTAACATTGTTCTCGAGATA
AGATGGCCCTCCAGCTGGACTTGATGTTGGCATATTCATAACTGTGACACTATTGACTTGTATCTGTGCAGTTGGTGAAACGA
CTCCATTCTCAAATGCAGAACTGGAAAGAGGATTTGTCTCTTTCCGATTCGCAACTTCTGCTGCATTTTTTGCACGAAGCAACC
TACCCCTTCTCCGCTGAAGAATGCGTCTCTGTCTTCGTTTCGCAGAAAAACTAACTTGAGTTTCCGCCTCTGGCACATCTTCCT
CGTCCTCACAAGGAAGTGTATGCAAATATGCCACTAGACCAGGAGGAATTACTCGAGAAAGAAGCTCTAATGCTGGCTGGTA
GGAATCTGCCCAGAGGGCTACAAGCTGCCTACTGACCTCACGCCTTTCACCAGGCAGTAGAAAAAAAGCATGTTCAAGATGT
CTGAGCAGAGCACCATCACGCAAAGCAGCATCTCTCATGGGCTCTGCAGCAATTGCATCCTCCTCTGCAATTGTCCGCATAAT
CACCGCCACAGCCTCCCTTACACTTTCAGCAGGATGTCCAAACAAAGCAAATAATCTCCTCCGTAGTCCTGCTACTTGTCGTA
ACATCTCTACAAACATAGAGTGGTCAGTTGTCTCGCTCTGAGGTTCACAAAGCATTGCCTCCAGCACTTCTACAACGGCCATA
GATAACAAAGGTGACACAGACAAAGGCTTGAGTCTATTTACTAAGATAGTTACATAACTTTGGTTCCCAAAAAGGGAAGACT
TTGCATGCATCAAAGTTGCATGGCTCTCCCCCTTTGAATTAAGGACGCAACTGCTCTCTCCTGGGCCACCTCCAATAAGCACA
GCAATCAAACTAGCACCTTCTGCTGCTACCCCCTCAGATCCACTCCGCAGTAAGCCCATAATTCGTTGTACAGCAATGGGCAT
AGAGATAACAACTGAAAATGCAGACGTAGAACCAAGAAGTCGCCTAAGGCAAGCAATGAGACCCATTAATGTTGCAGATGC
TTTCGGTGATGGAGCAGGTGGTGGAGGTGCTTCTGGAGGCAAATTTGGTGGAGCTGGAAGCATTGATAATAAGGCCATGAGT
GTAACATCTGGTACCTCAGAGCTGGCTTGCAGTCCACTGTAGGGAACACATGCATTGAATTCCCTGACCCTGCGCCAGAACTT
TGCTTTTGATCCAGGTAGGGACCCACCTTCAGCAACAGCATCTTTCGCAGCTGTTGCAAGGTGTTTCAAATGTAGTACCTCAC
CTTCAATATCTGCAAAGAGACGGTGATGAGAAGAGCTTGCAGGTTGAGGCAGTGAAGGAACTTTGCCACATGGTGGATCAAG
CCGATGACCTGGAAGGGTCAGTCTTGGCAAAACTGGCACAGGCCTTTGCCCCTCCATCTGAACAACATCTTGTACTGCAGCCA
AGAGACTGTCACGAGATGTGCTGGAATAAACATGAATTGGGCAGCCATCATTGAACTCCAAAGCAAAGAGCTGTGGTTCTTC
TACAAAGCGGACTAGAGCTGCGACTGAAGATAAAGGCCGTACAATGACTGCCTCGTAATTATCTGGCCTCCTTTCAACCAGTG
ATAGCTTTGTCAACACAAGCTGTCTTGGTACAGAATCACCATCCTCCCCAATTCCGCCTTTCCCTCCTTGCCCCAGTCCTGCAC
CTATATTGTGCACAGTCCCATGCCCTGCAGATCGAAGTCTTATAACCGACCATTCTCCATACCTTGTCTCCTCAGCTCCAACTG
CCACCATAGCTCTTTGCTTCAGAAATTCCATTGCACCAATTGTTTGTGAGCTATCAATCATCAAAGTCAGGCCCATCTGTGTCT
TTGCAACATTTGTCAACTTTGAAAGGATAGCTGAATTTGACACACCAGCGGCTGCTATAAACGCTTTGCATTTCCTGCCATAA
AGAGGGCAAAAAACAAACCCACCTCCTTCTACGCCTTTTCTTCCATAAGCATCAGCAAGAAGAATAATACCAGGTGAATCCA
CATCCCGAAAATCGAGACACCAGTGCAAATCTCCTGTTTTTGGCTGTCTCACTTCCATACCGACAGCTGTAATGCGTAAAGTA
AATGGGACCCAATTGGAGGTTCTCCGCCGCAAGTGAAGTACCAAGAATTCTGCCACAGCTGAGGTCGAAGGCTGATGCCCTA
AACTTCTGCTGCCTTTCAGCCTATGGAGCTCTGATAAAATTGCAACGCGATACCTGGAAGAAAATTTGATGGCTTTGAACTTT
CCACGGCCATCAGTTCTGACGTTGATGATAAACTCTTGTGCGTGCTGAAGCTGGGGATCATCCCTTCCAATGACAGGTGCTGC
CGTTTCGAAATCGGTCAGGACATCGTATGAATTTGTGACGCTAAGTGTTGCAGGATCAAGGGTTATGATAGAAGCCTGAGAT

ATGCAGAGGATCCGCTTGTATTTGCCACGCCAGGAGTGCTTGATGACCATATAACGGGCAAGGTACTCGGGTTCTTCTTGTAC
ATGAGCATCTTGTGCAGGCGCGTTCTCGGATGGGGATGACATGGCTGGGCGATTGTTGACATAATCCATGGTCTGAAGGAGG
TGGAGAGGGTGCAATGGGGGGCGCGCATTCGAGCGGAGAAGCCACCAGCTCCACGCCGCTTCCTTGCGGCCCCCCGACTGGC
TTGCCCCCAGCTCTTCCTGCTGCTTTAAAACGGCGCTTTGCTTGCGCGCGGCGCTACCAATTTCTGCAGATGGCAACGAGCTCA
GATCGCGCTATGGCTTCCCTTGAATTCACCTGCAGCTCTCTGCACACACACTCCACGCTATTGCACATGAAAACTGTCCCGTA
GGGGAGAGAGAGAGAAAGAGAGAGAG

**TOPLESS (*TPL*):**

GCTAGAATCTTTATCCCATTGTCATTTGTAGTCACAGCTAGAAGAGAACCGTCCTTGTTGAACCAAAGCCGGGGACTA
GCCTGCAATGAGGATGTGCATACTGCATTACACCCTTACTCAAGGGAAAGGCAAAAACACCACAGAACCTAGCACACCCTAT
GTGTTCCTTTGTTCAATAGTAATCTTACAGGCAATCCACCATCTGCATCGATGGTGGTAAGCAAATTAGTGTTATCCACATCCC
AAAACTTAATTTGGAATTCATCGCCAGCAGCAAGAAAATGATTCCTGGTAGTATCAAACTGCACTACTCCCAAAGAACGCTTT
CTGAAACCTGAATAGGTCCTCTTTATTGCTCCCTCACTTTCATTCCATTCCACTAGATAGGATTCACCTTCTTTGCTTGTCCCAC
ATGAGAAAAGCCTGGTTCCATCAGCACTGTAAGCCATGGTAGTACACCATTGGCCTGGAGCGTCATAGTCTACCCTTGATCCC
AAAAGATCATACAACCAAGCCTTGATTTTCCCATCAATTGCTGTCGAGAAAATAAACTGAATGCTTTCTTTATGATGAGGGCA
TACTGAGTATACAGGAGCCTCATGACCCTCAAAAGTGTATTGTTTGCGGCCATTAACAGCATCCCAAACCTTAATAGTTTTGT
CATCTCCACAAGTGATAATGCAGAGTTGCTTGTTAGGATGAGAAAACGCAAGGTCATTTACACCACCAATATGGGCATCAAT
CTCAAGGTGTTGACGCAAATCATTTCCTCCCGCACATGAGTAAATATGGACCATGTGTTTGGAAAATGCAACCCCAAGTAGAG
TACCGTCCGGACTCCATACACATCGATTGACGGAAACAGCAGGGTCTTTGACAAGCGCAGCCTGCATGGGCATGGTGCATAC
ACTAAGGTCCCAGACTTTAAATGTTCTATGTGCTAGCCTTTCCTTTGTTCCAACATCCCAGATAGCAATATCACCAACATTTGT
TCCAACAAGCAGTATGGTCTGCTGAATTGGATGAAAATCCATGCTCATTACACATGATCCTTGATTCAAAATTCGAGCCACCA
TCTTTGGCAATTCGTCAGGAGAGTACATGGGAGAGGGATGATTTGGGGCAGCTCCAACAGGGTAGGTCACCTCATCCACAGC
TTGCCCTCCAGAGCGAACCCTCTTCATAATGTGTTCTGAATCTGCAGATTGGTAATCAAGAGCAGAATTATTTGCTGGTGGGG
TCCTAGGTCGTTTCAGAAGGGCAGCAGCTCCTGGGTTGGAAGGGGCCGCAAGTGCAACGGACCCAGCTGCAACAGATGGGTG
GGGTGCTGGCGGGTTTGAGTTTGCCATCCAACCAGCTATTGAATTTGCAGTCGGAGCTGACGCAGGGGGAAACGGTCCATGT
GCCAAGGGAGGAAAGGCTCCCTGTTTGGGGAGCCCACCAACCAGCGAATTGCTAGCAGCAGGGGGAGCACGGGCTCCATTTT
GAGGTCCACAAGAGTGATCTACAAACAATGTCTTAATATCCGGATTGGGTCGAGGGTTCTTGCACAGCTGATGTTGCCAATTC
AGACTCTGGTTGATGAGAGTTCTTAATCGTGAAGCCTTCAGGGCAGGGAACACTAATTTGTCACGAAAAAGAGGATTGGCTT
CAATAAGCTTTTTAAGTTCTAGTAGCATAATGCTCCTTGCAGACTTGGTATCCCCATACTTTGACAGCTGCTCATTCTCCCTAA
AATTCTCTAGGGTCAAAAGTTGTGTGATCTCCTTATACAATTCCTCGTTGAAAGAGGAAAAAACCTTAAGATCCTTGACAAGT
ATATCCACAGCTTTAGCTCGATCTTGTTTGTCCAAAGCTTCAAGGTACTTTTGCTTTCGGATCTCAAAGAATATCTTCATTGAG
TAGCGGTTATCATCCACTTTGGTGAAGCCTGACAAGTACCTTTCTACTTCTTCCCATTCCCCTGCCTGGACCTGATCCTCAAAA
TACTTCATGTTGAAGAAGAATCCGGACTCCTGCTCAAGCTTGTGTACAGTCTCCTTGAACTTCTCCTCATCCAGGAACTGCAG
AATCAGGAAGACGAGCTCGCGGCTCAGCGACGACATCGTGCAACAATGTCCGCTACGGATCTTAACAGCAACGAATCAAAAG
ATGGGGGGTCAATACAATAAACAGAGCGGCAGGACCCATGTTGCTTGAGGGCCCCTTAAAAGGGCTTATGTTGTGTATTTGG
GTTCCACTCTTTCATGGCCAAGCTCCAAAGCATAATGGATTTGTGGCTATGGCAGTCAAGGTATAATTCAGCACAAGGTAAAA
GGAGAGATTGGCTATGGTCTTAATCTGCACAGAGGTAACACACGAGGATTCACTCAAAGACAAATGCGTAATTTGTGAAGGC
AACAAGGGAGAAGGGCGGAGTTAGAAGATTTGGGAAACAAAAAAAAAACAGAGAAGTTAGCTATGGCGGGGAATTGCGCA
GGGATAAGAGAGGACGGGAAGAAAGGCTTTAACAAAGAGGAAGCGAACAGGAGTGTAAGAAAGCGTAGCTGGCAGGCAGT
AGTTAGCTATGGCGGGATGCTGCAGAGGGGGGGAAGCAGAGGAGAGAGAGAGAGAGAGAG

**UBIQUITIN-SPECIFIC PROTEASE 26 (*UBP26*):**

TGAACATGTACAAGAAAAATTAACAGTCAAAAGTTCTTTAGATGCTGCCACAGAAACCGAGGGGCAGTTTTTCTCCTA
TACCCCATAATACTCTAATGCACTTACATTTCACTCAAGTGCTGCATTGGCAAAAGATGGACCAACAAGATCAGGCAATCTGC
TTAGCAGAGTGCCCCTAAAGCCTCCTTCCATCGAAACTCTGTCAGTGTCTTGCACATAAAATTCCTCTGCTATATCCCGGTTTT
CATGCAGCCCCGTGTCAGTCACCCAGAGATGGGCTCCAGGTAAAATGTTCAGGTCAGACAAAGTGGCTGAATCATCAGTAAG
CACTTTCTTACAAAAATGTAGACATTGATTTTCTTTCACAATTGAAAAAGATTCCCAAATCAGAAGCCTGAGCTGATAAACCG
ATGTAGCTCCAGAAACCTTGAGAGCAATCCGTTTGTTTGATGAAGTGGTCCCTCGTCGAGCTCGCTTAGATACGCGTCGCTCG
CCTTTCAAGGGCTCAAGAAGTGCTTTCGGGGGCTCATCTCCATCCACAAAATCCACAAAGATTTCTTCATCAACATATTGAAG
CTTCTTTATAAGTTCTGTACTCTCCCTTTCCTCGATACAAACATGACACACCTCTGGGTTTGTGAGGAGAGTCGGAATTTCAGT
ATCATTTTTACATCCTTCAGATACTTCCATTCCCACAGCAGCTTCTGTGGAGCTAGTGCTATTGCCATCTTGCATCGCTGGCAT
CACAACAATAGCTTTAATGAATTGGGAGTTATTCACATTCCATCGCCCACAAAGATCTATCCAGTTATCTTCCAATACAATTGT
GAACATGTCATCATTTGGATTACCCTGCACAAGCTCACCACGCCGATTTCTCTGCAGTTGTGGGGGTTTGTAAAGAAGACCCT
TATGCTTCTCACAAAGTAGATCTCGTAAGCTATTCTCAAGCCTGAGGGGTTCTTCAGCCTTCTGCGATTTCTTGCCACTGCCTC
CCAAATAAGAACGCCATTGATGCAGCCACATTGAAGGCACAAGATAATATGTAGACCCAGGCACTATTGGAATGCTCCCTCC
AGAAAACAGAGCTTCTTGTTTTTGTCTTTCTTCAATCTTTGTAGCCCTGAGATCTTGCTGTTGAGATGCAACTTCCATCATTTCT
GCATCACAGATGGAGCAAGTAGGAGTATCTACTGGAAAGCTCATGCATCCTTCACCCAAATCTTCTTGAACTTGTTGAGCTAC
TTGAAAAAAATAGTCCCAAACCTCTTCAAGTACTGTCTGACGCTTTGCACCTGTAAAGTTATCAGGAAGTAAAGCCTTGTGAG
GGCATGTTATAGCAGCAGTTGGAGATGCATCAACTTCTGTCGGGGCTTCTGCTGCTTTGCGTCGCAACCAATTTTGTAGCCAA
GCTCTAGATACAAAAAAATGTCTTCCAGCTCCATTGGTAGCATCAGCATCTTCAAGCAGCTTTTTTATTTTCAATCTCTCATTT
TTGAAGGAGTTGACTGATGCAATATTCTTGGCATTCTCCATGACGCACTCAACACAGCAATCGTATGCAGACAGCTCTGGTCC
ACCCCCATACTGAGATTGTAAGTTGGCCCAAGCTCCTTCAGATATACGCTTCATTGCACAAACATTGGATGGAGGTACTTTGC
CATGTTCACATAGCAACTCGCTGTTGTCAATGGGAAAAGGTTTCAGCTCATCTGCCCATGATCGAAGCCAACTGCTAGATATC
CAAAAGTATCCATTATCTTGATTTAGAGCTGGCAATTGTCCTGAAATTGTTTTTATCTCTTTCTGCCTATCTGCTCTGTCCGAAA
TCTCTTTATCTAGCTTTAGCTTGTACTCCCGGCATTTGTCTAACAGATCTTTATTTTGAGCATCTATCTTGCAGCGCAACTCGTC
AGGCAGTTGAAAAGAGTCATCACTGGTGGATGCCGATGTTGCACCAAGTGGTTCTCTCAGGTTGTAGATCAACATATATGCAT
CAGCAGAGGTCAGACAAATCTCAAACTGATCAGGTGCTTCCTCGCGTGTTTCAGTGTCTGATGCCAGCATGGAATCTGGTGTG
GCACTGCAAGATGCATCCTGATAGCTAGAAGCCTTTTCACTGCTGACATTAATTTCCTTTTTTGCTTCTTTTTTAAGAGCTTTTC
CAGGGACTTCCCCACAGGGATGAGAACCCAAGCTTGTTACAAGCTCATCATCAAACTCCCACCACTCTCCTTTGACATCATTC
TTGATGTTTGCAACATAATGGCCACTATTTGTTGTGCTTCCTTTGTGAAATAAAATACAAGAAAGATCATATAGCAGAGATTT
TTCAGCACGTGGTCCACTTGATCCATCAATGCTAAGCCTGGAGCTCATGTCAAGCACTTGTGGAAAGCTAAATTTTGAGGTCA
CCTTCTTCTTCATTGCAGTCTTGGCATCAAATACAAAGCGCTTGAGCTGAAAATTTAGCACTGGTGGAAGAGACCTCAATTTG
GTGAAGTGAGTTGCATCCACTCTTGCATTGCAAGCCTCGCATAAGAATTGATTTTCCCCAACCAGCTGCTCCACACTGAGGTA
ATCATCAAGGCTTCCATCCAAACTTGCCAACCCTTTCACATTGAGCTCTAGTTCATAAAAGTCCACAACTTTCTTTGAAGCGGG
AGACTCCTGCCCGCAACTTGAACACCTAGTCACATGGGACAGGGTTCCTCGAAATACATCTTGCACGATTGTCTTCACAACAG
CATGCCTAGAAAAATTCAACAAGCCTTCTAGCGAAGACAAAAGAAGCTTCAAAAATTCCTGTCCATCCTGCTGTATCGAGTTA

TTCAGCTCAAGCACCTCTGCAAATGGAGCAGAATCCACAGCTTTCTTTACCCCAAAGCGCAGCTCACCAAAAAGAAGTGCAA
GCTTCTGAAGTACAGGCTGACTTTCAAGTAGTTCTGGTTCAGCAGCAAAGAATCCCCGCATAAACGGCTTGATTCTATACAAG
CACTGCAAAACGCTGTTTACATAGCAAGTAGCCCCTAAATTTGTTAAGCCTGAAGGTGAGTCTATGGAGGGGTGAATCAGTTC
TTGAGGGTTAGGACCGAGCATTGCCTCTGTATCAGATACTTTTGCCATACCGTTCCACTTTTACGCGATCCTCCAGAAGGAAT
TAGTGCACAGAAACAATTTGGAGAGTCCTTGTTATTCACACGGCAACCTTCACAAGGTGGCTTATTGATCCGAAAAAGCTCCA
AAATGTCCTCATGGCTAACCTCATTCTTCTCATCAATTTGTCTGAGAATGGATGTTTCAGGATCTTCAGCAGTATTGCGCTGCC
TCCGGGTTTTGTGTCGGCTGCCTGCCCCGCTCATACTGTGCATGTGCAGCTTAGGATGAGCAGCTTCAATGGCGAGCTTCTGA
ATTGCAAACAAGTGCAGGCTTGCAAAGCAGAAATGGCAACTCCTAGCTTCTTCCCCAAAATGGACTCGAGTTTGTGTGTGTAT
CAAAGCAAAGCAGAAATGGTGTGTATGCATGCAATCATGTGTGTTTAAGGTTTTGCCTTTCAACAACAATTTGGTGATGTTTG
AAACCTCAAACGCGCATCAATCTGAGCAGAAAACCTGTAAGCTTTCTTTGTTTGTAAAGAAAGGCGCCATGTAAAT

**YODA (*YDA*):**

TTTTTTTGACTCAAAACACAATTTTAAAAAGAGTTGTATCGTTCGGATATTTATCACCGCCAACAATATCCAATACAG
GTTACAAAAGTTCACATTTGCAAACTTTGTTATATGGTTCTGTCAGCAGCCCTTCTACAAAGGTTGCATTAACAAACTTTGATA
TACAATTGTTCAGAATCATGTCTTCATTGAGGCCCTTAAGTGGGATTTTACTGCATTACTGTGACCAGTCATCAAGACTGAAAT
GGGCAAAATGCTAAAATACAGCCGTGCTATGAATGCTGCTTTGCTCTTGAATCCGATATGTTAAGGGGACGGAGATGCATTGC
TGCAAAAAAAAAGAATGCTTTAAGACTCCTCGCCAACATCGAGCAGTAAGTGGAGTTCGCCGGTACGCACACAAATACTCAAA
ACTGCACACAATCAGCTTTCCAAGCCTCCACCATTACACATAAGTTAATCTGAACCACATGCTCATGAGAGCAAGGTAAATAT
CATGAGACCACCGAAGCCCTCCAGGCTTCCAAAAACTAGCCTGACAAGTTACTAGCAGCCTCGTAGAGTAACTCAGACGCCT
TCTGTGCGTCTGCTCGAGAAGTTTGCAAGTTTGCTTCCCAACTCCATGTGTTACTCGGAGTTCTGCAACAATGATTGAAAGTTG
CAGGTTCCCCTATCTCTGCCGCTCTTTGATCGTGACCATGGGCTTCATCAATTACAAAGTGATAGCTTCAAATCAGATAGCAC
ACTATAGATTCACTTCATGACTGCCTGTACGACTGATTGCCGTTGAACCTGCGGGCGATGTCGGCCCATAATACCTTGGACTTT
GCTGTGGCGATCTTTCACGAAGCACCTGAGTTACTCTCTCATCCATCCTTTGCAGAGCACCAGTACGCTTCTGCATATAGTCGT
CCCTTTCTCTAGGGGGCCGGGCAGCCGCTGGCATATAAAACCCATTGTTACCTGGACCATAAATCTGTGGCATTTGCCTATTCT
CTGCAATTATACCCCATTGAGCATCAGGTCCATTCACTGTAACATGTAGAGAGGCCGGGATACCATGCCGGACGAGCGGCCTT
CCTGCCCTTCTATTGGCCAAATTTGAGTGATTTCCAGGTGTTGTATACAAGTCAACACCAGTGGTCGACAAACCTCCCGGAGC
ACCTGTTGGGGGGGTTGCAAATCCAGGAAGAAGACCAGCTGACCGTGTTGGAATCAACCCATATGATGACCGTGGAGAAACA
GGAGCACTCACAGAAGGGAACATGCCTGGTCGGGCTGGTACATGTGATTCCCTCGTGTTTGAGAACTGAGTTCCTCCTGCTCT
TGAAAACCTTCTTGAACTCCCTTCATTATCTAGCCGCATACTTCTCTCTCCAACATCCAGGAAATCAGGAATGTTCAAGGCATG
GGCTAACATTTCAGGACCATCATTTGGGATTTGTACAAAAGGATGATCCAGTAAGAAGGCTGCAGAAGGACGGTCGGCTGGA
TTCCTTTGCAAACAATATCTGATGAACGACTGTCCTTCTGGTGAGAGGCCATCTGGAATGGGAGGAGTCTCTGCTCTGGTGAC
TTTAAACATGGCTGCTACCCCTTCATACTCGCTCCATGGAGGCTTCCCTGTGACCATCTCAATGACAGTACATCCAAGGCTCCA
TATGTCAACGGCAAACTCATGTCCAGTATTCTTCTGCATGATGACCTCTGGAGCCATCCAGTAAGGGCTTCCTTTAAAAGACA
AAGGGACTCCTTGCTCCTTTATATGCTTTGCCATGCCGAAGTCAGCTAATTTTACTGTTCCATCCTGATCCACAAGTATGTTTG
CCCCTTTTATATCCCTGTGAACGGTATGCTTGGTGTGCAAAAAATGTAACCCCTGCAAAATCTGTCGAGTATACCGCCTTATG
ACTGGCTCTTCAAACCTTTTAAATTCTTGATAAAGCTTGTAGATAGACCCCCCTGAAACAAGTTCCAAGTATATGTACAAATTT
CCACCCACCATCTCACTCCCTTTATAATGGACAATGTTCTGATGTCGTAAAATACTTAAAAGCATTATCTCCTGCGCAAGCTGC
TTTGCGGATTCTTTTGACTTAGGATCATCCCTCACAAACTCAACCTCCTTTATGGCACAAAATTCCCCTGTTTCACTATGAATT
CCTTTATACACTTTGCCAAAAGACCCGCTACCTAGCAAACACCCTTTCAGCCATTTTCCAGCTGATGCCCCAGCCTCTGCTTTG
CCAGGACTCCGAGGTGGTACTCCAGAAGAGCGTGGTGAAAGTGGTGTTGAAGGTGCAGTACCCCCACCTGACGTTGAAGGAA
GTGGCAGTGGATGTCCATCACTCCAAGATGATGGATCATAAGCTGTACTATTGGACAGAGGAGGATGTCGAGGAGATACAGC
CCCACTTTGAGCTCTTGGACTTAAATCTGAGCTTGGCCTTCTTTGCTGCAATGGAAAGTCACTTGATATTGAATTATGCCCTGA
ATTATGGCCAGAGGTGGGACTAGAGACCTGTTGAAGATGGTTCGATGCAGGGTTCCAAAGCCAAGGCTGCTGCTCTTGATGA
CCCATCCCCCGAGGGCTAAGCACTGGGCTGGAGAATGTACTAGTCGAAGCACTATTCTGAACTCCATTGACATGTATGTGGAT
GTGAAGATCGCCTCCTTCACCCTGACCGTTTCTGAATGCCAACCTAGGAGAAGCAATAGTACCAGCAGGGCTTCCCGGTTGCG
TCCCATCAGCCCGTGCAGTAGATTGAATAGGGAAATGTGAAGTAAACCTTTGAGAGTCTTCAAATGGTGAGGAGTGCTGTAG
CCTCAAAGCTTCCCCTTCAGAGTGCCTCCTGTAGGAAGTAGAGAGCTCATGACCAATGCCGCTTTCTGATGAGCTATCGCTTG
ATTCCGAATCCGTGGGATATACGGAAGCCTCCCTGCTATCATTGATTCGTGACTGCAGCGGAGTTCGAGCACAAGGTTGGATG
GTTGGCTGTGCTACGCATGCATTCTGCCTTGGGGGGAGAGGCAAAGGTTTTGGGATATCTGTGCATCTAAAAGCTGAATTGGA
TGGAGATGGTGATTGTGAAACCCATGAAAATCCAGTGGCAACCTCTGCAGAGTTGTCACCATTGAATAGCGAGTTCCTTGACC
TATCAACGCTATCATCATCCTCCTGGCGCCGATTACCTCGATTAATCCCGGATCTCGCTAGCCTGGGGGGATTCTGGATTGATG
AATCCTGCTCTGACCTGCAACTTGACCTCTCTCTCCCACTCAGAGAGGAATCCTTCACACTATCGGTGCTTGCTTTCTTCCTTGT
GTCCTTTGCTCGTCTGAAGAACGACGGAAACATCGGCATTTGCAAAGCGGTCTCAGCACTACATGTGCACACTGACGACCTAG
AAACCTGAGAAGGATACCGGACCAGTGAAAAGCTGGGTTCAAAGAAAACCCTGATGCAGGAGGATTTTGAAGAAGCCTCAG
ATCGACTAAGTGAAACCTCTGAGTGAAGTCCTAGCTCCAGCACTGCCAGGTTGAGCACTCTGATAAACTCTTCACTATGTCAA
AAGCCGTGCTGGTAATGACCAACCGAGTTAAGAGGAAGCGCGCTGTGTATTTCGATGTTTGTGAATGTAGAACAGACTAGCA
TTTTCGAGATCCAGCTGCATTTTGGGCACATTCCCTGTATTGTAGCTCCAACTCAACGATAAGATCATTGCTGTGGAGGAAGG
AGAAGATAGCGTATGAGAATGGCAGCTCTGTGTGGAGGACCACTACAAGCTTGAAAGGCAGTTCGTGAGTTACCTATCCCTG
CTCTCATGGGTCAACAATGCACTTTTCCGCAGTCTCTTGAACCACAAAGATGATGAAAGCTTGAAGGAACGTCTGTGTCAAAA
TTGCCCTGCCTCAAGCTCAGCACGCTTAGATCTTCTCCCAAAATTTAGTGCTAAAATGGCATAACCTGCTACACTCCTTGTAGA
ATCTGTGAAATCTTAACATTGGGTCTGAGAATCCATTGAAAATTTCCCGCAGCCTTTGCGTTGGCTAAGACTGGATTTTGTCTG
CTATGAGTGCACGGCAAAGCCTGCTATACTTCTGGAGAAACCTGAGAAACCCTAAAAATTGAGTATGAGAATCTAAGGAGAC
CGTGACAAGGGATGAGTGTAGGCAGCTGTAATGTGGCCCAGCTGAAAAGAGCAAAAGGGCAAGGGGAGGGTAATAATCAGG
CGAATTTTCCAGCACAGACAAGGTGGGCATCAAAGAGTGCAAATCACCAGTAATGCGGGAGAAAGCAAAAAAGCAGGAAAG
CAAAAAAGAAAAGCTGCAGCCACAAGGCGAAATAGAGGGGAAGGGCATGCAGGGAAGGGGAAGCCCCTCAAGGCTCAGAT
GGGTTTCTTCTTCAATCTTCACCTCCTGGAAAGTGCAATGCTCATGACGATGGTGGCCTCCTCTTCTCCCCCTCCTCCAGCTT
GCACACTCTCCCGAGGGTCCAAGGCAAGCCTCACTTCACTACTAGTGCGCCCGTGTGTGTGTGTGTGTGT

**ABA-HYPERSENSITIVE GERMINATON 2 (*AHG2*):**

ACGACTGTTTGAAGCGTCAACTCTCGATTTGGCTTTGCAGCTCCACTAGAAGTGGACATTATAATGGCAAAGCTTTAC
ATGTTTATACCGAGGCGTTAAAAAGGTTAGCTACTAAAAATTAGTAAGGAAATGAAATCAGGGGAAAAAATCAGCACAACAT
ACAAAAGTCAATCAGCGATCAGAAGTTGAAAACTTTTCACATGACATGAAAGTCTCTAATGTTCAGTTTTTTAGGATGTTG
TATCAGTAGGAAAAGAGGGCAGACAATGAGCTTCTTTGTATAAGCTGACCTGATTTTGCATCTATGTAAGCTGTGGGTTCCCG
CACACACCGCTACCTGATTTGGTTTCCCACCACAACTGTTTCACAGTGCCCTCAAGTGCGCCGGCCATGTGCTGCAATGCTCCT
TTTGGATACCTGGAGTATCTTGTGTCATGTTCCTAGCTAGGAAGGCATGGAGGCAGATGATGTGATACATCAGGTACCTTTAC
CCCTCAGCACAACAAACTGCTTTCATTACAGGAATCTGATCCAAATTCTTAGACCCACCCCCTGGCTCTTTCAACTTTCCTGTG

CATAAATCCAGTATCCCTGGTCCAAAATTCAATAAATTTGTGAACTCTTCAACTTGAGAAGCTGTGGGAGTGTCATGGGCAGT
TGAAAGAAGACCCTTGACATTAACATTCAGTTGATGGCAGATTTGGGCAAACACACAACCAGTCATGAAGGCATCATAACCG
GCCTCATGTAAGAGGTTATTTTCTCTGTCTTGGTACCTCTTAAACGCAGGCATGATGTCTATATGAGCTTTACCAGAAAACTGG
TTGGTAGCTCCCGCAAACCTTGCAGTTCCAACCCCAATACTCCTTGATTTGCACAGATAAGAATAAACAGCATTCAATGAGGT
ATGTTTCCTTTTCCAGGACGATTTAAGGCTTGGCACAAACTTCATAAGATATTTGGTATCAATAATGGTTGGAAAATGAGCTA
AAAGAGAAGCTGAAAACTCTTCTGCAGAAGAAGGAAGCGGCGAGAGAAATTTATCATGTATGTGGGCTAAATCAAGAGCAC
AGTTATGTCCAATAAGAGGTTTTTGGGATGACACCATGGCATCAACAATGTGCCGAAAACCGATTGCCTCGTTGATCTTAGCT
TGTTCACATCCTTGTTCTAGCTCACTCTTTATAATTTCAAAATATTCTTTGGACTGGGTGAAGAAAATCTTGATATCGTGACTT
GCAGCAGAAACACAATCACTCCTTACAGTAAAAACAAGGTCCTTAAAATGTTTGTCAAGCACCAGTTGTGCAAGACAAACTT
GGTGGTGATTCTCCAGCTGCAAAGTCAAAGAAGGGCACTTGCTCAATACACCACTATGTGGCACCTTAAAATCTTTAATCGGA
CATCTATTCAACAGCTTGAAGTTAAAAATATCTGCCGCTAGGTCACTGCCATGGGAATTCACATCCAAACTGCAGGTACTTAT
CTTGCTGTGCCACTTCTCTAAATCCACCCGTAGTCGTTCAGCAAACACTACATCTGACGTGTTGTTTAAACTTCTACTACATCC
ACAACTCTTCAGTCTAAAAGGAACATGTTCATTGGCCAGCTGTACCCTTGCTTCAGATTCTTGTGCCTTAGACAAATATGATAT
ACCATCCTGTATGCACAAGTTGAAGTCAAAACCATGAGCTGCTAAAAAATCCAAAGCAGCAGCCTCGCAAGAAAACGTTGCA
GCAGGTAGACCAAGAGAAAGCTCATTTCGAGGGAACACATAAAAATTGTACGGGTAAGCGAGGAACCGGTTGCAGGAAGGG
TCAAAATGGAAGAGGGCGAGGCCGACCTGGAAGACAGCGAACTTCTCAGCGAGGTGCTTGATTCTGTGATATGACCGGAGGA
GGGGAGATGGGGGGGCTGCATGCGAGCTTTCAGGGGGCACACTGTGAATGCCACTAAGCTCCAAGTCGAAGGCAACAAAAT
CTGCAGCGCCCAAATGCGCCTTGAAGCTGTCAAGCGAACTCCAAAAATTCTGCCTTGATATGTTGCCCTCCTCTGCTTGTACTT
GCCATCTCCCTTTCTGTTCTTCTAGTTGATGGGTGCTGTTCCCCTGCTGTTTTGCTTGTTGATGGGAGCTAAAGGCTCGCGCGCT
TCCAATAGTCCTCCTCGCACTCCCCCGCCTCAGCCAATTCATTCTTTGAAATTCATACCTGCCTCCTGACTGCCCTTTGCTATTT
ACTACTACTAGCTGGCAATTTACCACGGGGAAACATGGTGTT

**PLETHORA 2 (*PLT2*):**

ACACACACACACACACAATCTTAGAACAAGGCAACGAATGTTCTTACTGTAAGAGCATGCTTTAATCATGGATGGGG
CTGCGATAAAATCTGTCGATGAAATGTCCGCTACAAGTAGCGATGTCTCCAATTGCACAGGAGATGAAAGAGAAGCTTTCAT
GAGATCTTATGGTGAAGAGATAAACCAAGGTCACCCAATGCTTGAGGACTTTCTTGGGAGCAAAACCAAAGCACACCAGGTG
TTTGACAATGGCAGTTCTATTAATACGCATGCATTGCGCGGTGTAAATGAGGGAGGCTTCCTGCCGTCTTCCCTCAAGCACTC
TGTGCTAAGTGTTTACAAAGACTCGAAGACGATAAACGATGGATTACAAGGGAGCTCTGCTATTGATAGTGGCAGACTGAAG
AGGGCTTTAGGTTGCTTCAAGGATAGTGTAGCAAATGAGGGTATGCATCAACAGTATCAATACAATGGGGCAACTTTTGCTA
ACCCCACTTTATTGACATTAACACAACCAATTACTGGTCTTGATTCAACTAATCATCCATTTTCATACTACAAAGCGAGAGATC
GCGGCGCAAATGAACTTCAAAGCATATTCCAATATTCTGCTACTGATAGCTCTGAAAGCACACTACAGCAAAGCAACCCTGG
TTTGAGCCTAGAAGACTGCAATTTGGGGAGGGCCATCCTCTACCCTAATTGGAGAACTAATAACATTGCAGCAAATCTACATG
ATATGGGGAATTTAAGAGAAAACTACTCACATTCTTATAATAGAGAGGGGGAATGCATGGCAAATGGGGATTGTAATATTGG
AGGCAATAGGCCTAATACTAATGGGTGCTTACAAGGAGGTTTTCCTAATAATTTTAATTTTAACAAGTTGCAGGGTACTTCAA
TAGCCCCTAGTGCTACTCTGGGGCATTCCCTCTCACTCTCTCACTTCCCATTCAATCACCAATCGGCTCTCTCTCTCTCTCGAGA
CGCTGCTAATTCTGCATTAAAGAGTAGTAATATCATGATCAACCCTCAATGTGCTGATCAATATAAGGTTGATTACAGCAATA
TACAACCTGCAGAACCAATGAAGCTTTCCATGAGAAATTGCAGTAGTCACTTTGATGCAGCAGGAACAAAGAGAGCAGATCA
TCTGCTATGGCATCATCTTCCTCAGCAAGAAGGCTTTGCAGCATGCGCTGCAAATAGTGAAAGCATAGAAAGCACTGTTAATA
TGAAGGCAGTTAGTGATCAAGATGATGATCCATGTAAGGGTAAGACTGTAGGACCTGCTGCTGCTGCAGGTTTAATGGAAAG
CTCCATTCCAAACAGTAATGATGATCACTTGCAGCTTCAGCTAGTTCATTCAAGCGCTGAGCAGAAGATATTGCAGCATAATA
ATAGTGCTTGCAGTCTTGATGCTGGTAATGCGGAGTCCACTTCTCGCAAATCAATCGAAACCTTTGGCCAGCGCACTTCTGTCT
ATCGAGGGGTGACCAAACACCGTTGGACAGGAAGATTTGAAGCCCATCTTTGGGATAACAGCTGCAGAAGAGAGGGCCAAA
CCAGAAAAGGAAGGCAAGTCTATCTGGGTGGGTATGACCTGGAAGAGAAGGCGGCTAGAGCATACGACCTTGCAGCTCTAA
AATATTGGGGTCCAACAACAACCATCAATTTTCCGCTTGAGAACTATGAAAAGGAGATGGAGGAAATGAAAAGTATGACCAA
ACAAGAGTTTGTTGCATCTCTAAGAAGGAAAAGTAGCGGGTTTTCAAGAGGTGCTTCCATTTACCGTGGTGTAACAAGGCATC
ATCAGCACGGACGGTGGCAAGCTCGGATTGGAAGGGTGGCTGGGAACAAAGACTTATATCTAGGCACATTTAGTACACAAGA
AGAGGCTGCTGAAGCATACGACATAGCCGCCATAAAATTTAGAGGATTGAATGCAGTGACAAATTTTGACATGAATAAGTAT
GACATTGAGAAAATATGCTCCAACAATTTGCTGCCTATACAATCAATGAGAAGGGTAAAAGAAAATGTGGGGGCAGTTGATT
TAATGCAGGGGGAGGATGCAAAAGGCAGTAATGCCATTGGATTAAGTGCTCATTTAGCATATGGAAATGGAAGCTTTGATAT
TGCAGCAAGGGCAGGGAGTATACAAGAGTGGCAGTTAATTAATCAGCTCGCTCAGCCAGCTAATCCATTATACCATGATAAT
TCCTTGTTGATGTGGAGCAAAATGGATCAAGAGAGAGCAAAATATGCACTCCACACAGGCTTGCTCTTCCATCACAGTAATA
ACTTAAAGTTCCTTGAGGGAACTTATAGTTCTCTAATGCAAGGAGCTACAAACTTTCCTCATGTGGAAAGCTCTCCTCACTCTA
TGCAGACTCAAGGTGCGCCTGATTTGGAAAGGCCTCCTTTTATGGGTGCGACACAATCGTGCAGCAGATATAATTTTGGGGGT
GGCCAGAGCCACTTGGCAGAAGAACCCTCACATGTTTGCGAACAAGTACAAGCTCCTGAGTGTGTAAGTTTGCAGTTGTAAG
AGCATATTCCCACGCTCTTTCATTTTAGTATGTCTGACATTAAAAATACTCCACATACAGGATCATTGCAATTTTGCACTCTTA
CTTTAAAGCCCATTTTGCTGTTTTAGGCTAAGTGTCACATTTTCCATGTTGGTGCAAAGTTACTTGAAGCCTCTACATGTTTAT
G

**WUSCHEL RELATED HOMEOBOX 8 (*WOX8*):**

CACACACACACACATATATATATACACAGATACCTGCAAGCACGCCATCGCCATACGCTTCCGTTCATTCTTAGCAGC
CGCAGACCTCATCTCCCCTGACTCAAATTCAAGCAAGCTCTAGTTTTCAGCGGCATCAAAAAGTCAGCGGCACAGTCGCAGCC
ATTAGATCCAGCCAGTAGGGGACTACAGCCGTTGGATCGTGTCTAAGGAGTTTAATTAGCGTGGTGCGTACGAGGGGGACTT
GTTTGACGCTGCAGCTGGGCGGTGCTGGCGCTGCAGAAGCTCTCTCTGCTCTCACTTTCAAATACAGAGATTCATGTAGTAGT
ACTCCCACCACCATCTCTGCAGAAAACGCAAGCTCCTTCAAGCACCCATTTTTTCATCCTCATCATTGTCTGCTGCAGCTTTCT
TCTTATAATTATTCGACCCGTGATCATTATATGGCGGCTCCTAACTCAGAGAGCGCATTGAATTTAAACCCAAGGAATCAATC
TATACAAGCCGCAGTCAATGCAGGATATCTAAGTAGAGAAGAGGCGCAGCTGCTAATTATGCAAGATCCCGGTAATTATGCC
ACACATGCGCTACCGTTACAAGGGCTGTCCATGAATAATATGCAGGCGCAGTTCTATAATGAGCAGATCCTGCGTTCTATGAG
TGGGTCGCACTTCACGGGAATAGAACCCACAGAAAATCAAGAGCTGATCAACGTAATGCAAGCACAGCAGAGCAGCTCGAC
ATCTAATGCAGGGTTGCAAAACCTATTAAGCATGGAGAACGTTGGCAGATCGGTTTCAAATGCGCAGGCTGCGCCAGTGCGT
TACGAGGAGACATGCACACCAAGGCCAAGGTGGAGTCCCACGCCAGAGCAGATTGAGATTCTGGAGGCACTTTTCAACTCCG
GAACAAGGACCCCGTCACGCGATATGATAGCGGACATCGCGGTGCAGCTGCAAAAATATGGCAACATCGCAGAAGCGAATG
TCTTTTACTGGTTTCAGAACCGCAAAGCGCGAGCCAAGAGGAAGCAGCTGCAGCCAGCGGCATCAAGGTCATCAAGCAGCTG
CCCCTAACTTCTAATGGCAGCTCAGGCTATCAGATATTATATAATCCTTTTATTTTCTTAGTTGTCCTGCTTGTCCTAACCATAA
GGCTAACTAACTAGACTTAGCCTAGTACCGGTTACCGGGATGTCCGCTTTGCCTCGTACGCGGCCAAGCGGACCGATAACTAG
CAAAATCCAAGAAACTAGTCCATCTCTTGTCTATTATATCACCTTCTCTTCTCTTTTATCCAAACTAAAATTTTGGTTCTTGCTT
CAACAAGACTGTTTATGTCCTCCACCTCAATACATTCTAACATAGCAATGTGGATCACATTTTGTACCCCCAAGTCTTAAAAA

AAAAATGACCAATCTTTTTATAGCATATATGTTTTTTGA

**MATERNAL EFFECT EMBRYO ARREST 5 (*MEE5*):**

GCGGTTTTGAACCCACGAACATTTCTATTTTCAAAATCAAAACCCAGCGCGACTGTGCAGACGAGAAGAGGAGAAGA
GGCTTGCAATCATGGACGACACACTGTATGACGAGTTCGGGAACTACATAGGCCCGGAGCTCGACTCAGACGAGCAGGATTC
CGAAGCTGATGATGAAGACGACGGAGAAGAGGAGGAGGAAGCAGAAGAAGCTGATGGGCACAGCCGCAGCCGCTTTCCGG
ACGGCAACTCCGGTGATGGCGCCGACAACGGCTACGGTGCCGACCTTGCTCTCGATGCCATGGACCTTGACACCAGTAACGG
CATTGTACTTGCAGAGGACCGCAAATACTACCCCACTGCTATGGAAGTCTACGGCGAGTCTGTTGAGGCCCTTGTCATGGATG
AAGATGCCCAGCCCCTTGAACAACCCATCATCAAACCTGTCAAAACCAAGAAATTTGAAGTGGGTTTCAAAGAGGGTAGCAT
GCTGTCGCATGTTTCCACTGAATTCCTTCTTGTCCTCATGTCGAACCCTGTGCTCGTGCGCAATGTTGCCCTTGTTGGGCACCTT
CACCATGGAAAAACCCTCTTCATGGACATGCTTGTCGAGCAGACGCACGAGATGAAGACTCTCGATCCTAACAGCGAGAAGC
ACCTCAGGTACACGGACACACGCATCGATGAACAAGAGCGGCAGATCTCCATCAAGGCTGTGCCCATGTCCTTGGTTCTCGA
AGACAGCAACAAGAAGTCTTTTCTTTGCAATATCATGGACACACCCGGGCATGTGAATTTTTCAGACGAAATGACTGCTGCGC
TGAGGCTTGCAGATGGTGCTGTACTTATTGTAGATGCTGTCGAAGGTGTCATGGTAAACACAGAGAGGGCAATTAAACATGC
TATACAGGACAGAATTCCCATTGTGGTGGTTGTAAATAAGGTTGACAGGCTAATTACTGAGCTAAAGTTGCCACCTACGGATG
CATACCATAAGCTGCGGCATACAATTGAGGAAGTCAATCAATCGATTGCCAGTTTCTCTACAGGGGCAGATGATCCTCAGGTT
GTTGATCCGATTCTCGGAAATGTGTGCTTTGCAAGCGCAACTGCAGGATGGTCTTTTACTCTCTTATCATTTGCCAAGTTGTAT
GTCAAGCTGCACGGCATCCCATTTGATGTAGAACAATTTGCTTCTAGACTCTGGGGGGATTGGTATTTTCATCCTGACAGAAG
CTTTAAAAAGAAAGCTCCTCCGGGTGGAGAGAGAACATTTGTCCAGTTCATTCTTGAGCCGCTTTACAAGCTGTATAGTCAAG
TGATTGGAGAACACAGGAAAAATGTGGAAACAACCTTGGCTGAATTAGGGGTCACACTGAGCAATGCAGCGTACAAGTTGA
ATGTGAAGCCTTTGTTAAAACTTGCTTGCAGCTCGGTATTTGGTTCAGCCACAGGATTCACTGACATGTTGGTTCGTCACATAC
CGTCTGCGAAAAAAGCAGCTATCAACAAGGTCAATTACAGCTACACAGGTCCTCAAGATACAGCTCTTGCCGATTCTATGCG
AGCTTGTGATTCGAAGGGGCCGCTGATGGTCAATGTCACAAAGCTTTATCCTAAATCCGACTGTAGCGTGTTTGATTCTTTTGG
AAGAGTTCTGAGTGGAACCCTTCGTACAGGCCAGACTGTAAGGGTTCTAGGTGAGGGTTACTCTCCAGATGACGAAGAAGAT
ATGGCGGTTAAAGAAGTCACAAAACTGTGGGTTTATCAGGCAAGATACCGCATACCTATAAGTCAAGCTCCAGCTGGTTCAT
GGGTTCTTATTGAAGGAGTGGATGCCTCTATAACAAAGACAGCAACGCTGTGCCCGGAATTTTCTGATGAAGATGTTTATATT
TTTCGGCCGTTAAAGTTCAATACCTTGTCAGTAGTGAAAACAGCCACAGAACCACTTAATCCTAGTGAGCTGCCAAAGATGGT
AGAAGGCCTCCGAAAGATCAGCAAAAGCTATCCGTTGGCTATAACAAAAGTTGAGGAATCAGGAGAGCACACCATTCTTGGT
ACTGGAGAGATATACCTGGACTCTATAATGAAAGATCTCAGAGAGCTTTACTCTGAAGTGGAAGTTAAGGTGGCAGATCCTG
TAGTATCTTTTTGCGAAACGGTCGTAGAAACATCGTCACTGAAATGTTTTGCAGAGACTCCAAACAAGAAGAATAAGATTAC
AATGATTGCCGAACCTTTAGAGAAGGGCCTTGCTGAGGACATAGAGAACGGTGTGGTCAGTATAGATTGGCCTCGTAAAAAG
TTGGGGGATTTTTTCCAGGCAAGGTATGACTGGGACGTACTTGCTGCTCGTTCAATATGGGCGTTTGGTCCAGACAAGCAGGG
TCCAAACATTCTTCTGGACGACACTTTACCCAGTGAAGTTGACAAAGGACTTCTTAATGCAGTAAAAGACTCCATTGTCCAAG
GATTTCAGTGGGGTGCTCGTGAAGGCCCTCTTTGCGATGAACCCATCCGAAATGTAAAATTCAAGATTTTGGACGCAACTATT
GCTCAAGAGCCTCTTCATCGCGGAGGTGGCCAGATAATTCCTACGTCTAGAAGAGTTGCATATTCTGCATTTCTCATGGCGAC
CCCTCGGCTAATGGAGCCATTCTATTACGTGGAGATACAAACACCAACTGATTGTCTTACAGCGATTTATACAGTGCTTTCTA
AACGCCGTGGGCATCCAACTGCAGATGTTCCCAAGCCTGGAACACCTGCGTATATTGTGAAGGCATATGTACCAGTAATAGA
GTCTTTTGGGTTTGAGACTGACTTGCGGTATCATACACAAGGCCAGGCTTTTTGCTTGTCTGTTTTTGACCATTGGTCAATAGT
GCCTGGAGATCCTTTGGATAAGTCTGTCACCTTACGACCTCTGGAACCTGCTCCTGCACAGCACCTGGCTCGAGAGTTCATGG
TCAAAACTAGACGCAGAAAGGGAATGAGCGAGGATGTAAGCATCAATAAGTTTTTTGATGATCCGATGCTTCTTGAGCTCGC
TCGACAGGATGCTGATCTTCAGCAGATTCTTTGATTTCCTCGGGAGGCTAGTATTGGCATCTGCACCCTTACCTATATGGCACT
TAAATTTGGACCATGTCTATCCCAGCAAACCACACGGGCAAGTGAGGGCCGGCACAGACAGCGTGCTTGTTGTCAAACATGA
ATGCAAAAAGTCCTCACAAATCAGTACCGTAGAGCTTGTGCAGTTAACCACAGAGTTAGTGGCTTTTTGACATCAGGCTGATA
TTACAGTAGTGGCCATTTTGGAAGAGGCATGCCTTTTTGAGAAGTGGTACATTCAGAGTCTTCTTGACAAGTGCACAAGCCGA
GATGTGAGTGTATACAAATTTTTTTAGAGGGTATCCACACAAGTTTACAAGTGATGATAGGCTGAATTGCATTGATTGCAAGG
CACTAGTCAGCACTCATGACTGCTTCACTTTAGCCGGCTTAACCATATTAAGAAGTTGGGCAAGTTTGTCCACTTGAGCGTTTG
CAGACTTAACCAGAGGGCTTTTGGCTCTCAGTGTGCGCGAGCATCGATTCCTCTGCAGCTTGCAATGATAGGAGCCTTGCTGG
AATCATTTTCGTAATCTTAGGCTTTGTAGTTCACATGAAAGGGGTCTGAAGCTAGTCAAGGGCTCAAACCATAAAGAGCTCCA
ATGTAAACGATTATAATGAGCATCTTCTCGTTTATCCTGCTTAGCTTCTCATTAATCTAGATCTAAACCTCAAACTGTTTCTCT
GGAAGCTGAATTGTTAATTCAGCATCTTTGCACCAATCTCCTGTGAGGTGGTGAAGTTCTTATACTTCCAAGTTTTACTTAAGT
TTAGGAAGTTCTAGCCCCTGAATGTGTAATTTAATTGGCAGATGTATCGTCTAAATATAAATGGTGTGTTATGTATCTGTATAT
GTAACTTAAAAACAGGCATTCTGGGATTGCATCCGAACAGATTCTAGCTTGATGGCCACAAGTCAGGAAGCGGTCTG

**MATERNAL EFFECT EMBRYO ARREST 63 (*MEE63*):**

TCATGTACATATCTATGTATGTAAATATAATATATAGATATAAATATATATGTGTTAGGTGCTGTTGGTTTAAAGTATA
GGGCTAAATTCGTTACAAGCGCGGAGCCATGGCGGACTCGGATTTGGGAGAATTCTATGTACGCTACTACGTGGGGCACAAG
GGCAAGTTCGGGCACGAGTTCCTGGAGTTCGAGTTCCGCCCTGATGGCAAGCTGCGCTACGCCAACAACTCCAACTACAAGA
ACGACACCATGATCCGCAAAGAAGTCTTCCTCACGCAGGCTGCCCTACGTGAGTGCCGCCGCATCATTGCCGACAGTGAGATT
TTGAAAGAGGATGACAATAACTGGCCAGAGCCCGATAGAGTCGGCTCTCAAGAACTCGAAATTGTGATGGGAAATGAGCAC
ATCTCTTTTACGACCGCAAAAAATTGGCTCCCTTCTTGATGTTCAAAGTAGCAAGGACCCAGAGGGACTTCGCATCTTCTACTAT
TTGGTGCAGGACTTGAAATGTTTTGTATTCTCCCTCATTGGCTTGCATTTTAAAATAAAGCCAATCTAGATGTTTTTTGGAAGA
GCACCTACATGGTTTCAAATAGCAGCTATATGGAAAAGATTTTATCTTTTGCAGTGGCTGTGAGCACCTTTTTATATCATTGGT
GTGAGCATTTAGGCAGAGTCTTGCAACTGGCTTCATAGGAACTTGCTTTCCATGTCAATAGTTTTTGGGCTCCTTTTTATCGTT
ATAAACTCTCGCTTCATGACAAGAGACACGACTTTTATATTGCTGGCAGGATTATATGCAGATTATTGTCACAAAGCTCGTTA
TGTGAGTAATACAGAGCCACAATTCCGGGTGTGTCAACATAGTTAACAAATTATTTTGAGGTCAGATTGGTGTAATTTGAGTG
CATATGAGAGCAATAGGTTACTATCTGTTAGTTTTTTTGCATCTATTTGATATATTGTGATTTCCAGTCAAGGTTGGTTTAAAT
CAGGGCTGGCATTTTGGGTTTGTTTTAATGCCTTG

**MATERNAL EFFECT EMBRYO ARREST 58 (*MEE58*):**

GTGAAGAGGCTTGTAGGGAAGGACAAGCAAGATCTTTAAAATGGTTTTTCACCAGCAGAACCCTACGCCGGTGTTTCT
TGCTCGCCATAGCCAGGAAGCCCGTAGCGACTGACGAGGAAGACGCACACAGCCCTCCGTTTCGAGCTCGGCGTCCTTCATTT
CTGCGCCTATCTTGCTAGCAGAGATGGAGCTCTCCGTGCAGAAAGGTGCTAAGGGTTTTGAATACAAGGTGAAAGACTTGTC
GCAAGCTGATTTTGGCCGTCTGGAAATCGAGCTCGCTGAGGTTGAGATGCCTGGCTTGATGTCTTGCCGCACCGAGTTTGGTC
CGTCGCAAGCGTTGAAGGGAGCACGCATCACCGGCTCTCTTCACATGACAATTCAGACTGCTGTGTTGATCGAAACCTTGACT
GCCCTTGGTGCTGAAGTACGCTGGTGCTCTTGCAACATCTTCTCTACCCAGGACCATGCAGCTGCTGCTATTGCGCGTGATAGT
GCCTCGGTCTTTGCTTGGAAGGGAGAAAATCTTCAGGAGTATTGGTGGTGTACCGAGCGGGCCTTGGATTGGGGACCTGGAG
GTGGCCCAGATCTCATTGTTGATGATGGCGGTGATGCTACTCTTCTCATCCATGAGGGAGTCAAGGCCGAGCAAGCCTATGCT

AAGGATGGTACCCTCCCCGACCCCACTTCCACTGATAACCCAGAGTTTCAGATCGTTCTCTCCATTATCCTTGATGGCCTTAAG
AAGGATTCCAAGAAGTACCACAAAATGAACGACAGATTGGTTGGTGTTTCTGAAGAAACCACCACTGGTGTGCACCGCCTTT
ACCAAATGCAGGCTAATGGGACTTTGCTCTTCCCTGCCATCAACGTGAATGATTCAGTGACCAAGAGCAAGTTTGACAACCTC
TATGGGTGCCGCCACTCTTTGCCTGATGGCCTCATGAGGGCCACTGATGTCATGATTGCTGGGAAGGTTGCTGTTGTTTGTGGC
TACGGTGATGTCGGCAAGGGTTGTGCTGCTGCCATGAAGGCTGCTGGTGCTCGCGTTGTTGTTACTGAGATTGACCCCATTTG
TGCCTTGCAAGCCCTGATGGAGGGACTTCCAGTCTTGAGGCTTGAAGATGTTATAGACACGGCTGATATCTTTGTGACTACAA
CTGGTAACAAGGATATCATCATGGTTAGCCACATGAAGAAAATGAAGAATAATGCTATTGTGTGCAACATTGGTCACTTTGAC
AATGAGATTGACATGCATGGCCTGGAGACGTACCCTGGTGTGAAGAAGATAACTATCAAGCCACAGACAGATCGGTGGCTTT
TCCCTGAAACGAAGACTGGCATCATTATTTGGCGGAGGGCAGGTTGATGAACTTGGGTTGTGCCACCGGACACCCCAGCTTT
GTCATGTCTTGCTCTTTCACCAATCAGGTGATT

**HASTY (*HST*):**

TGCAACCAATACATGACGCATCATATATGACCATAATGGGTAGTTCTTCCCATCCAATTTGTCCTCAGCGGCCAACAT
ACGAAATACATCATTGTTGTCTGACATGATTGCAGATTTGGTTTAAGAAATAGTACAGTTTTCCTGCCTCTCAGTTTCTTTTTC
CCTTGTGCTACTATATAAGTTGAGCCAAGGTGGTCGGTAGAGGCGCATTCATTTTATCTATGCTTTCAAATTCCTGTAACTACT
GCTGAGATGTAGCAGTGGTATTGATAAAAGTGCAAAAAAAAGGGTCTGTTTGGACTTGCATTAATACTGATGTACAAATAAA
TATAATGAGCACAAGCATGCTCTGATGGAGGTAGTTTTGAAGATCTTGGCTACAGTAAGTGGTACCTTGAGATGAAAAAACA
TCAAAAACATTTACATTCATTGGTAATGTAAATAGCATACAAATTGCATGCCGGGGAGGCAGGACTCGTATATAATGTCAAAT
AAACTGTGTGCATCCTCCTGTACAAAAAGGCTCTCCGGAAGAATAAAATGATGTGATAAGCACTCACAAAAATGCCGCTAGA
CCAATCTTACCTGCTTCCTCATTCTCAATAGCAGCTACATGGTGTGTAGGTCTTGGACAATTTGCAACATTCGAGATTGGATTT
GGAGACTTGGCTGCCATCAAAGCCTTGATATTACCACCACCAGCCTGCAAGAGAAGGCTCTTTATATACTGCCTATGTTCTTT
AGCACTGCAGGTTGTGGATAATGCATTTTCAAAAGCCAATAGAACCTCTCCTGTAATTGAAGGGGATGATTGTAGGATCTGCC
GTGGAGCTGCATGCTGCTGTGCCATTATCAAATATATAGTCCTAAAGAGATTGATCAGCTCTGTCTGCGCTGACGCATTTGAG
TCCATGGTAAGTGCCTCGATAATTGCTGTAAACAAATCAGTTGCCACATAGCTTTGAAGTTCCATATTGCCAGAAATGATAGC
TACTGTTACTGTGGCTCCACAAAAAACTAAGGCTTTATGTACTGCTTCAGTATCTGGCCAATTCAAAGCTTGTATGCATATTGA
TAGAGATGCCTTGGCTATACTATGATGCTGTAACACAAGTCTCATCATACAATCAGATCCAAGTACTGCAGAGCTATCAAACT
TCATTGGTTCCATTCGAGACACTAACAACTGATCTCCAGAAGGTAAATTTGGGTTGAGGGCAGGGGATGCCAATACTGCCAAT
AATGTGCAAGTTTCTCTCGAGAGATCCCTGAGCAGTTTTTCTTCTAGCACTTCTTCCTTCAGGCCAAGTCCTTCACTCTTGCAA
GCACCCTCAGAAAGCTTCACTGCTCCATTCTTTATCAGATTCATCCAGGAAAAGGATAATGCTGAGTCGCAGAAAGCAAGAG
GACCTGGTAGAATTTTGCCAAGCCATAATTCCCTCATATGCCAGGGACAAGACTTGACAAGAGGAATCACTACGAAATGCAG
CAGTAGCCGCACGTGCCTGTATTCCATTGCTTCAATATTTTCCAATAGAGCGATTTGAAAAGCATTACAGTCATGGCCAAAGA
ATATGTCTGTGAAGTGTGTAGCTGCCAGACCAAGTAGGCTGTAGCTGCTGTCACGAACACCCTTCAACCAATTCCGGACTTCA
TTTTCCTTTCCACCTATTTGACTATCTGTATGCATTTCCACATCATTAACAATAGGAGCATTTGCTTTTGAAGACTTTGTTCCAG
CTTCTCCAATGATGGATGCCTGCTCTGCGCTGCTCATTAGCAAAGCACCTTGTACCACACTAGGTAATGAATGCTTTATATTGG
GGGACCATAAAGCGTGTATGCATCGCAGCAACTGCAAAAGCGGTGTAATGATCCATTTTATATGATTTACTAAAGGATGTAG
AGGTTCTGTTGCAGTCGCTCCTTGACAGAAGAAATCGGGATGTTGAGTTGCACATCGTTTTACAACCCTCTCTGTAAAGACGA
CCGAATGGAAGATTGACCATAACTCTTCATTCGCATTATTGCCTGTTGGTTTTGTCAATAAAGCCACAAGATTTTCTGGGCTTA
CAAGAAACCGCTCTTGCAAGTTATTACTGCTCCACGTAGTTTGTATGGGAGTCAGAAAGCAGTCTAAAGCCTTTGCTTCACAT
TCTTTCCCTGCTGCTGAGCCTGCCACCAAAAAAGCCTCAGCAATCAAGTTATGCTCGCCTTGCTTCAGTAAACCCTGCTCTTGT
AGAGTAGATATGGTTTTCCACATGACCTCTAAATGAGCAGAAACCGTCTTGTCAGCTGCCTTTGCAATTCGCAAAAGGGATGT
ACAGACTTGCAGCCTGGAGCGTGAATTTTCAATCAGTGAGGAATCTTTTCCAAAGCTCGTTGAAACCACTGGAAGGCTTGCAA
GCAATTCAAAAATCTTATTTAAAACACTTAAAGTTGCAGCAGGTGAGCTTTTCAGGAAAGGGCCCATGGCATCCAACAGGCG
CCCATGAATTTCTACATAAGCAGAACCGCTCCACTTGGCCATCAGAAGAGTGCTTAGGATTCCTTCGAGTGGTGCCTGCAGAG
AAGGAAGCTCTGTGGATGGAAAGAGATTTTCTCCACCAAAAACTGCATAAATAACAGCCTCCAACAATTTTTGGGCACTGTC
AAGAACTCCCAGTGTCTCGTAAACGCGTTAAGACAGCAGTTTGAAAACTGGAAATGGAATTTAACGCCACATCACCATGGAT
ATATTTGAGCTTGCAGAACTACCACATGCCTCCTTTATTCTTTCGGAAATTTTGATGACAGCTATTAAAGGACGCCGTACTGCC
ACCAAACGAAGCAACTCCAACAAGCGGCCACGATAGTTTCCATAATCCTTCACATCAGAAATCTCTTCAATGTTTCTTGCTGA
ACCACCAGGAACCTGCGCTTTTGAGCTGTTTTGAAAATACCAGTTGAATGTATGATCTAAAAGGCTTGCATAAACATCGTTTG
AGATCAAGATCATAATGCCCTTCTTGTCCTTGAAAACTGGTCCTATAACACCCAGCTGTCCGGCAATTAACTCGTTTCCATGAC
TATATTCAGGATCAGCAGGTATGTCCCTTAAAACTCCCAGAATAACAAGGTCGAAAAATGAACACCAATTTTACTGTGCTGAT
AAAACCAGAGCATCTGCTGAAGATAAGCCGTAAGGTGATCTTGAAAACCCACAAAACATTGAAGGTTTTGTGATCCAAAGGC
AACCAATGCCTCAGTTAGATATTCTGCAAACTCTGAGTCATCCTCTTTATTTGCAGATGGTGCATTTTGTAAAGCTCTAGATGC
TTGAGACAAGATTTCTAGAACACGTCTTAAGGCAACATCAAATGCTGACGCACTTTCATCCAATGGCCTTTTTCTTGATGAAA
TTTGCTTCATAATTTCACCAGACCGTAGACGAAATTCCACAGTGTTCAGCAAAAATCCACAAGCTTCGATCAATCCACTCTCA
GCTATCTTGTCTACAGGTGCCCATTCCACGTAAGCCAGAACAGCATTCAGTGTTGCATTGACCACAGCTGCATGCTGCTTTGC
AATGTCCAGCTGATTTCTTTGCAATAAGGTCAGAGCTTCACCGAAATGACACTCTAGCATCTTATAAAACAAAGGGAATATCT
GTGGCAATGATTGTGTTAGGCTTTGAAGCAGTTGTCTACGCCTGTCTCCTTCCAAATCCTCGTTATAGACTGTTATGTCCTCAG
GAAGCCACCTCAACACCATGACGACCAACTCTGCGTATGCAGGGTTCACTGCTGCCAGCTCCATGAGTTTGAACAACAAACCT
TGCCACAAAGAAATATCTTCGTGTCTTACAACTTCAGCCACCAAAGCTGCAGTCTGGCTCTTCAAAAACCATTCATCCTGAGA
AGTTGCAATCTCAGGCACCAGCCCCAGCACCAAATCAGCCAGTTGTGCCCTAGTCTCCACCATTAGCTCTTCCCATCGCATTC
GCACTAGATATTGTAACAGCTTCAGCCCATAGTGACGAACTTCGGATGACAAGGAGCTTTGTTTAACTAGAGCAGATGCAAT
GGATCCAAGAATTTGAACATCTCCAGTTTTCAATGATTCAAGGAAGGTTGAGGCAGCAAGCCTTGCTTCAGGAGGCGAATTGT
AATCCAAGGCAGCATGAATTGCACGGGCTGTGATCTCCAACTGCGAGCTCTCCATGACCCACTTAAAAACCCAAATAAAAAC
GAGGGCTTTTTTCTTTTCTGCTAGGTTTGCAATGTGCTTGCTTGCTCTTCTTTCTTTAGCGTGGGAAGAAAGCACAAGAAAGGC
GGGCTCACTCTAATCCTTCACGATCGTTTCCCATGCAATGCCTGTAGTGTAGGCTTCTCATTTGCGTGCACTCAAAACTAAAAC
CCAG

**METHYL-CPG-BINDING DOMAIN 9 (*MBD9*):**

GAAGATAGAGAACAATATATTACTTGGAAAGGCAGATTTCATGATGTAGCACTTGTATAAAATGTGTCGTAAAATGC
AGGTTGCATTAGATTTAACTGAATACCTCACTTTTGCTAATGAGAACTACAAAATTTCTTACATAAAAGTAAAAGCCCTGAGC
AACTCCAGCAGAATCCTAGACTGCTGCTGCAATGCTGGAGATTAACACAAGGGCTGTTCATTGTGCTTAAGTAGGCAAAATTT
TACTTTACAGCATGCACATTTCCTTGGTTGATTTCTACAAAGCTCATAAGGCTACCAGCAGCACCCTTACAAATCCCAAAGAA
TCAAGGCCTTGGCAATCTTAAAACTGGGAACTACAATATGGTACAGAGTTCAAAGCAGATTGGCTCATCATTTCTTCACACTT
GCGCTGTAGTCTTCTTTTTCCTACCCGATTTGTTGGGCTTCTCCGAGTCTTCAGTATCTGGATCTACTGATGATCTTTTGTACTT
GATGGCTGCATCCAAAGAGTAAATTCTCAATGCCAATGCGCTCAGTGTATTGATCCCAACTCCAGCTGAGAGATGTGTCCAGT
TACACCAGGATTCCTTCAAGCAACTTGCTTTGACCATCTGTTCGAGCAATGTAACAGATTGCACTAACTCGAGTATAGAAACC

GCTGACTTGACAAAGGATCTCCAAGCTCGTCTGCGATTTGGTTCGCTTCTTGACGGCTCCATAATATCAACGGACAACAAAGA
TTCTATGTCAAGCAAGCTTGCTTTTAGTTTCTTGATAGCCCACAAATTTGTCAAAAGCGAAGGTTGAGATGGTGGGAGCAAGC
TCGACATCGTGTTTGCAAAGGAAATCTCTTCCAATGTTCTGGATCCTGACGCATTTTTCCCTTGATCATGATGCTCAGATGTAG
GTTCTTTACCAAAATGTGTGCTGGGCTTATCAGATAAACCATTATCAAAAACATAGACATCTGTAGGTCCATTTTCTTTGAGCC
CTGCATGTCCATCAATGCTTTGAATTGACTCAACAACAATCTGGATCTTCTCTACTTCCATCGAAGAACTGCAAAGCCCTGTTT
CAAGGGTTTTCATGTCAGACAGTACATCTAATGATAATGAATTCATGGTTGTCATGTGGAGGGCAGGATCCAGCAAGGGGGA
AACAGGAAGAGCAGGTATAAATCTAGGGATTTTGTCAGCAAGGCACCCGATTTGCAAGATTCTCACACGATTTGAGGTATGC
AAGGTAAACCTTTTTAATACAAGAGAGAGGTCTGGAGGATGAAATTCTTGGCCCTTGCTTTTTGCTTTCTTGGCCTTTGAAAG
AACCAAGGCTTGACCTTCATCTTCAAATCCCCTCCAGCTGCATGCCCCATAATTATGACCCTGAAGTTCTGCAACTGAGTCAT
AGGTATGATGACAACATGGACAATGGATTCTTGTTGTCCACACTAACTCCAAGCAATCACATCTGAATATCTTTCCCTCACCT
AAGGACTTCTTCTTTCTGCCCCGTTTACGAGAAGGTTCTTCACCAGGTGCTTCTGACTGAGATATTGATCCAAACTTTTTGGAA
AGAATGGTAGCAGCCTTTGTATTTACAGAGACCGGAACAGCAGATCTTGATTTGACTGTGTGCTGTTTATCTGGTTCTTTTTTA
TACTTAGAACTCCAAATCAAGCACCACTTTGTAAGAGCAGATTTTAGGCATCTTTCAGTAGCAATCGTGGGTTTTAACCACTT
AAGCAGCTGCTCAATACTTTCGTCATCACAATAAGCAGACCATTCATGGATGCCTGGGTGCGCAATAGCACTCATCTGAGCAT
TTCGAACATTTGAGCCAGTTGAAACCCTTGTGGCTTCAGATAAGCACGATGACTGGTTGCAATTAGCAGCATTTGTATTGTCA
TCTACTACAGAGCCTTTTTCAACTATTACCCAAGGAAATTTTCCTGCCCAGCCAAGGGCCCAATAGATCCTTCCCAGATCATCT
CTTCCCAGACAATCTCTGCGAGGGGCCATTTTCATCAGATCAATCTCTACTTTCTCAATCTCTGCCTCCAGTAACTCCAATGCT
GTCTTACCAGGTGTTGCTTGAACATTGGGTAAAGGAGAGGCTTCAAAATCACCACCACGATGCACATTTGAAGAGCTTAATG
GAATTGAACTCTTATCTATTTGACCTTCAAGTTTCCAACCTTGTAAAACACCACTGGATCCCTTCTGGTCATCTACAGCAGCTA
AATCTATTGAGCACCTGAAGGTGTTGACAGGTGTTGCATGGAGAGAATGATGAACTTCGTCAATGCTGCCTTGCATGCTTCCA
AGGCAATTGGCACTCGCCAATTCCTGCAGTGGGAAGTTTCCAATTTGAATATGTGTTACATTCTGTTCAGGATAGGTTACTCC
AACTTTGTTGAGATCCTCTGAATGATCCCTAGAAGCCTGAGCAGTTATCATGGGAGGCACATCTAAATCAGGATGCCTACTGC
TCATCTGTGCTTCTTGTATGGTTAGCCCTGAAGAGCCTTTAATCAGAATAAGATCCTGTTTTTGAGTGTCACCATACTTTTGAT
GCATTCCAAGAGCAGTCTCGGACTCACTTTGAAATAACTTATCAATTATTTTCCCATCCATAGTACCAACTGTAGAATCGAGTT
CATAACCACATGAATTCTCCGCTGGTATTCTGAAAATTTCTATTCCACTCTGGTTTCTAGTGTTTTCGTCCAACATGGGTGTGC
TAGATTTCAAACTCTTTCTGCCTCTTTTCTTTGGCAATGTTTCCTTGGATCCAGCTTCTGTGTTATCTGATACAGGAACATCAAC
CACCCTGTGGTTTCCTTTCATTGCAGCAGCCTGGCGTTCTATCAAGAGAACTCGTAACCGCTGTTGAAGTTCAGGTAACCCCTC
TATGCTTTGGTCAATGTGATCCCTGATGAAAGTTGATTCCAAGACTTTGTCACATAGATACTTTAACAATTGCACCCTCTCTGA
TCCACTTAAATGCCAGTAATCCTTGCCATCCATGCTCCTCGCCAGCGAAGACAAAATGCTTCCATCTTCCGACAGTCTGCTTTG
ATAGAACTTAAGAATTGTACTGGGTTCAGAAGCAACTGGACAGCTATTTGAACCTCCTTGCTCCAAAGCAACACAAGATGGA
CAATACCAATTCCCTTCTGGAATCATGGAAAGAGGGGGCTCTAAGCAGTAAATATGATATTCCGCATCGCATCCATCACACAA
TAGAGTGCTATCATCATCCTTGTCTATTCCACAAATCTTGCACACCCCTTCCTCCCAAGGAGCTTTTTGGAGTTCGTCATCTGA
CATCATTGATGAAGAGATATTGCTATGCTCCGTTGCAGCATCCTTTCCTTCTGTTACATTTACTTGGTTTGTTTCTTTGAGCTTA
GCAGCTTCCTCTTTACCATTCACCAGCTTAATCACCTTCTCCTCATACAAAGTTTCAAAGTGCTGCGACATTTCTTGTCCAACTT
TAATAAAATCTTCACGATCTCGAAAAACAACAGGGACATTTTGAAAAAGCTGCCGCATGTCTGCAGCATATGATTCATGTGA
ATCACCGTAGGATCCAGCAGCCAATCGAGAATCAATCATTCGGAAATCAAGAGGACGGGCAACTAATGGAAGGGGTACTTCA
ATGTTGTCATCCTCTTTTTCAAAATATCCCAATACTGTTCCTCCGAGAGAATTGCAGAAGAACCGCACATCATCTGGTGTGATA
AGTTGTCGAAGAATGACACGACATCGTTTCATTACAAGGTCAGGAGAGGCAACCATATACTCAACCTTCTTCTCCACTTTTGG
AGGACTTACATACTTTTCATTGGCAACTTTCTCCAAGACTGAGAGAACAGCTCTCTTCGTTGGTCCAGATGCATTTCCCTTGTA
AACCTCTTTACTGATTGACCATTCTAATGTTTCTCTGGCCCAGTCAGGAGGATTCATTTCAAGTGCGTCCCTTACACAATTTCG
GATTCTTGCACCCACATTTGTGGCAAGCTTTCGGACTGGTTCTAGAACAATGGCCCAAGGAGGCTTCCTAGAAATTCCTTCGG
AATGTGACTCATCGGAAGGCTTGCCATCTGAGCCTACCTTGCCTACGGGCTCAGAGTCAGACTTGTCATCTGCACCTGGCAGC
CAGCAGGAGAGTTGCTTTTCTGCAGCAGCAAGCAGCAGAGTATCAGCTTCAATGGCTGCCACTCCATCCAAAGCACCACAAA
GAACTCCACCATCTCCTTGTATGCACCGTAATATCTTCCTGCGCTCTTGAACACTTATTTCAAGACTCTCTGCACATGTATTTA
GTGTTAAAAGGCCAATAACATACCGCCTAGCTACTTCAGGCCAGGTCAGGTCATTCAATGGCATAAAGCTAGCTGACTGCTTC
TTCATCGTTGCTGCAGCTTCCCCCTCCTTCTTTTTACCACGCTTTGATTCAAGAGCCAAATCGGAACCAGCTCCCAATGCCATT
TGCTGAAGATCAGATAGGGCAATTTTTAAAAGAGCAATATTTGAGGCAGAAAATGCTACACTGCAGCCTGGAGAGCTAAAAG
TTTTCTTCTTTGTGAGCCTTTTACTCTGGAAGAAGCCTTGGTCCTGAACTTCTGAGTTGGCATGCTTCACAACATCCAAGTCAT
TCTGGAGAAAGCCTTTATCTTGACTGGCACGTTGGCTTGCCATATACATGTCCTTTTTGCTATGGCCTCTGTCAGTGTCCATCA
CTGAGGCATCAGCTACAATTTCTTCCAAGTCCCCCAAAGAAGGGGGCTCCTTCTGACCAAGAATGTCTGAGAACCGACACAG
AAACTCCCAAATCTGAAGCAAATCCCCAACAAAATTAGCAGATACTTTTTGACAAACCGGCAACCCAGTAGGTGGTGTGGGA
CGTAGCAATTTTAGCTTAGCCTTTTTGGCTTCCAGATACTTCTGTACAACTGGTATATTGAAGCCAAACCGATCCTGCCCCATC
CATTCATATAATTTTGAAAAAAAAAACTGTGCATCATGTAGTTGTCCTTGCGTCGCACCCTCTACGTTATTTGAAACATTGAGT
CCCAGAGCAGATAGAAGGGAGCTTTCCACATGCTTGCAGCCCAGCTGAAGTGAACCCTTTCCATACAATTCTGCTGCACGTTT
TACAACCTGTTCTGCAAGCAGTCTCCAAGCTCCCGAAGGCGAGGTGCTCTCCACAACAATCTCTCCAATTTCATCCCCTTCCCC
TAGATTCCTGACACAATTTTCAGGCTGATGATTAAGTTCCACGATACTCCTTTTTTCAACTACTGTAGACCCAGGGCAAGTAGC
CTCATTTAAATCCAGCTGTGAGTAATTCCACCCTTCATCTCTTGCTATAGATACACAAAGGTCTCCGTCAGTAAAGCTCGTAAG
GGTTCCTTGCAGCATGCTTAGCTCATCATCTTCATCATAAACATGGTTAAGGTACTTCGTCTCTATCTTCACAGCTTCTGGTCG
AGCTTCCACACCAGATTTGTCTGGTCTCTGTGAAACCAAGGCCTTTGGGAATCTTCCATCATCGCAAGAATCTAACGGAACAT
TACAGGACCTTCGTGTAACCCTGAAAATAGGAGGCAACCCTCCACCATCTACAACCTCTGAGATGCAAATAGACCCTGTTACT
CCATCATGCCAGGAGCAACGATAGCCAACAGGCCAGACAGAATTATCACTGTTATACCCAGGTTGCATATCTACGACACCCA
AAGCCTCCACTCTCAAGTCCATGCATTGCAAAGGCAGCTGATGCTCAAGGTCCCGAGCAGGGTCCTGCAACTCCAAAAAATT
ATCAGTCTCCATCTTTCTTCGCTTTATCAGTTCTGAAAACTCCCCATCGCTCTGTTGAGTTTCTTGTGAGACATCCCTTTTTCTT
CCTCTTGCGACAGTATCTTTTGAATTGATATCGTCTTTCCTCTGCTCCAGAACACCCAAAAACCTGGCTACTTCCAGTTTAGAG
CGGAATTTTTGTTTATCGGGAGAAAAGAACATAACATCCATGGCCTTGTCAGAATTGGCTCTTTTTTTGACGCTAACACTCCAC
TCCCCTTCTAAACTTCTACCTTGCTGTCCTAAAAACTCCTTCAATTTCTGCAAATGCTGTCTTTCCAGGTCTGTATCCTTTGATG
AAGCCTCCATATCATCATCTAGGTCTTTGCTCAATGCACTAGCGCTTGTGAAAGACGAATCCCTCTCCCCAGCAGAGATGGAC
TCCAATATATCTGCTCTTCCAGTGTCTGTTGGCACATCCTCCTTAGAACGACACAGCGTCTTCAACCTTGTCACACGTGTTTTC
GGTTCTTCCAAGGTGTTGTTGCTAAGAATGCTTGGCGGCATGCCCATATCTGCATCCCCATCCCCTTTCTTCTGCCTCTTGCACT
TTTCTTTTTGACTTCCGTCGGAAGGAAGCTCACTGTCAACCAAAAGTGGGCTTTCTGCAGCATCCATAACACCGTCATGAGGC
TGTGATGGGTGGATTTGCAAGCCATCAACAGTCGCAACTTTGGAACCTTGACCGGCAAATAAGCTGCCATTGTTAATTTGAGG
TGTGCCCTGAGATTCGTGGCTGCACGCAATTCTCCCTAGTGGCCAATGCCGTTTTCCCTGGTTTGTGCTGCATGCCAGGCAAAC
CCACTCATCTATGGCAGAGGCCTGGCTATCTCGAAGTTTCAGGCACTTCGAGTGGAAACTCAGCCCACAGACATCACAGATCA
ATGTCTTTTTACGAACTTCCGGCTTGCCACAGCAAGTGCAACTTACAAAAGCACCATTTTTAAGAGGGGTGCCCGCTTCTCCG

AGGAACTCCGCAGCTGGTCCAACATGGTTTGTGCGTAGTCGATACAAAATGGCAACCTGCATGGTGTCGAGGCCTGAGAGAC
TTGCAATGGGCAGTGTCTGTGTAGTTGAACTAGCAACTGGAAGAGTGTGTGAGGCCTGCATGGCTTCTGGGACATACTGGCCA
TACTCAACTTCTCCTGACGAATCTCCTTGCTGATTTAAATCAATCATCAGGAACCCTCTGCCACCCATGAAGACAACTCTGAG
AGAATAGAAGCTAAACCTGCAACGAGAGACAAGTTGCTCGCCTGTCTGGCCAGCGACGCTTATCTATGGAGCTTCCATGGTG
TACAAGAAGAAATAAACCCTAAAAGCAGATCTGCCAGTGCCTGTGTTGAAGGAGACGAGTTCAAAAGACTGCTTAGGGCGAT
GCCGCTCCTCCGCCCCGCCCCGCCCCCTGCTCG

**GLIOMAS 41 (*GAS41*):**

ATTGACGCTCTTTCAGCTCTTGAATTGATGAGCTGCACTTGCTTTCAGTTGTAAAGACTCCGTCTGAAGAAGGTTAACA
TCTCTTCTCAGCTTCATAGTCTGGCTCTGTACCTGTTGTCGAGCTGCTTTTAACCGTACTAACTCGTCTAAATCAGAATGCTTTA
AAAACCACTGCACTTGTGGATGGTCCTTTGTATCACCTCTTTTCTTCTCAATGGAAGTTTCAGCTGCAGGAGTCGGAGAAAGA
TCGCTGAGCAATCCATTGACTATCACAGAAGGGTGATTCCTGATGCGTTGAACAAAAGCTTCGGAAGGCTCAGAAAAAACGA
TTTCGTCATACGTCTCCACAACAACTGGCTTCTTCATAGAAGGTGGGCCCGCATCATCGTCTGGATACAATTTCAAGTGATGA
GACAGCTCAATGGGCTTTTCTGCTGCATCACCATGAAAGATGACCGTTATTCCAATCTCAAATTCACCCCATCCTACCTCGGTA
AGCTCAAATGGTGCTGTCTCCACCACCCTTGTAGGATTATTGAAACTTGGGTGAAGTGTGAAGATCACTCTCTTAATCACAGA
ACCAAGGTCCTCGTTGGATATGCTTCTTATGTAAACTGTCCATCTGTGAGAGCGGTACTCATCTGCCTTCTTGCCTAGCCAGTA
AGAAATCGTTCCGTAAGCAATCGGGCACGATATCTCAACATCTTTGACACGACGAACTGGAAGCTGCATGTCTTGGTAATGCT
TTTTGCTGAACGAGTCCCTGAAGCTTCGTGCAACAAATTTCTTTGCTTGAAATCAAGCATTCATAGCTCATTTGTAGGCGCCTG
TCTGCCCGCAATTGATGAAGGCAGCTTCTGACGTAGCACTATAAAGTTTAACAAAGAGGACGTAGGAACAATACTTTAAAGT
TTAACTTGGGCGTTAAGGTGATTCATTGCCAAAACCGGGTTCGAACCCTGTAGGC

**ENHANCER OF AG-4 1 (*HUA1*):**

GTGGCATTCCAGACTGGAAGGAGGTGTCAACAACTCCTACAACCAAAACACAGTCGTTGCCTGTGCGAGAGGGAGAG
CCTGACTGTTCGTTTTATATAAAAACTGGAAGCTGCAAGTTTGGCTCCAAATGCAAGTTTAATCATCCTGAAAACAAATCAAC
TGCTGTTACTACAAATCAAAGTAATAACGGCAATGTGAAGGCATCGGCAGCTTTAACTGATGCTACAGTTGATAATGGATAC
GAAGAAGGTGAAGAGGCTAAGGCTGCTGGCTACAATGCAAAAGGGCTTCCTCTAAGACCTGAGGAGACTGATTGCTCATTTT
ACTTGAAGACTGGAAGCTGCAAGTTTGGATCTGCATGCCGCTTTAATCACCCTGCAAGCGTGGCCATTCAACAAAGCCAGAC
GACAGTTGCTCCTGTGATGGGAGGCTTTACTGCACCATACGGAAGTTTCTCTATATTTCCAGGGGCTCTACCAGACTATGGGT
TTAGCATGCCAGGAGCGGATCTCGGATTGTCTAGTCTGGCTCCCCCTACGGTGTATCCACAACGGCCTGGTGAAACCACCTGC
TCGTTCTATGTGAAAACAGGTATTTGCAAGTTTGCAACCAGTTGTAAATTTCACCATCCTATTGACCGAAAGGAGCCCTCCAC
TAAGGTCACACTTGCCGGCTTTCCCAGACGAGAGGGGGAGCAAGCTTGTCCATTCTACATGAAGACAGGCACTTGTAAATAT
GCCCTAACGTGCAAGTTTGATCATCCTCCACCTGGTGAAGCAGCTGCCAAAGCAGTAGCTGAAGCTGGGAAGTCGGAACAGC
CTGAAGCTGGGAAGTTGGAACAGCCTGAAATAGGCGAGCTCACCCCCAATTCCAGAGTTGCTCGAGGCAAAGTTGTATGGAA
TGACTGAGGGAACAAAATAATGTTGCCTTGGGCTGTAGATGGTAGTGGTGTGCAGCTTCGTGTGATGTGGGTTGAGGTCCATG
TAACGTTTTGATTGTGAGTAGGATTATTTAGGCTTACCTGATTACGGTAATGTAGTGTTTTTACTAGAGTCGAGCATTTTAGCT
AGGCTAGGGATATCTACAATTTTTTTTTGGGTTCATTGTTTGCAGAATCTTGTTTTATTCCACATTAAGTTCACATCCGTTGTTT
TTCTTTTGTGGATGGAACTTGTAGTCTTGTTTATGAATTGAAGGAAAGAAGCCAAGGCCTTCTTTTATACAAAAGTTTGGGAG
GATGTAATCAAGTCTGCGCTTCACGTAAAACTCTGATGATAGTCGTCACATTCTCTAACAAAGTCAATTGTGATGGTATTTAT
GAACATCTTCAGAATGGTTCTTCAGAATGGTATTAATGAACACCTCCAGAATTGAAGTCTGGCAAAA

**SEUSS (*SEU*):**

CATACACACACACACACACACACGAGTCTTTCTCTATGAGTTTCTCTCACACACAACCCACAATCAAATTTGACGTTCT
CTCCCTCCCTCTCTCTCTCTGTGGGTTGTCTGCGCAGACCTTCACCTGCAAGCCCGAGGAAACGCTAACCCTAACCCTAATCCT
CACTTTTTCTTAGCTTATAATCAAAGAGTCCCTGCAGCTGAAGCTAAGCAAAACAAGAAAAATCAGACTTTACATTTATCCAT
CTGGATTAACCGGATTTAACGGGCAAGAGCACAGTCAAGCTGGCCTCAGAGCAACAAGCAGGGCCCGGCCACCAGCCAGCC
CTGCATCCTGGTGGCCCTCATCCAGTGGCAGGGACTCAAGTAAAACCAACTCCAACCTCCATGGCAGGTTCGCAAGGCGACA
GCCCCATCAAAGCAGATGGAACATCTTCGCACTCCGATCAATCATCTAACTTGTCTGATTGCTCCCAGGTTCTCCAATACTTTA
ATTGCTCCGATGGTCTGTCTGTATCAGCAAGTCTTATGCCCACCATGGGATTTGGAGAGGGAAGCCAAGCGCTAGTGAGTTCA
GTCAATCTCAGTGCCACAAGTGCCAGCTCTCTTACGACAGGGGGCAACTCTCTTATATCAGGTAACTCAACTTTACAGCAGAG
CATGAACATGAAGGCAGACTCCTTGAATTCTACAGGTGCAGCTTCCATGGGCTTGCCTACTTCACCTTTGTCTTTCTCATCTAG
CAATATTAGTTTGCCAGGTTCTTCAGGTTTTAGTATGACGTCCTTGTCTCAGTCAGGGGTGAGGTCTATCAACAGGCTAGATGC
GCCTCACGACCCTACTATACATCCAACCAAACTAATGAAATTGGACAATGGCTATCCACCCATGCTCAACAGTGAGGAGCCA
TCATCTAAGTTCGGGTGGCAATACAATGTGAATAACTTTGATGTTGCACGCCAAATGCTAGGTGCCAATGCGGCTGGGAGGTT
TGACCATCAATTCAAAGGTCAGGGTCTAACAGGAGCACGGCAAGATTTTCATAGTAACGAGCATCAGTTGGCTCAGTTGCAT
GCTTCTATGTCAGCTGCGAAGGCTATGCAAAAGGATTACTTGCAGCAGCAACCTGTACAAAACAGATTTCAGCGGCCAAACC
CTTTGCTATCAGCTCAACTTCATAGGCACGATGTCTTACAGCAACAGCCTCAACATATTCATCCTCAACTTCTACAGCAGCGCC
TTTTGCAGCAGGAGCACCTCTTGCGAATGCTGCCACCACAGCTCCAAAGGTCACATGTTTTGCAGCAGCAACATCAACTGCAG
CATCAGCAGCAGTTGCAACAGCTGCAACATCAAATGGCTGGAATGAACAAGCAACCAGATAAGGGTTTTTCGGGATCTTGTT
CCCGGCGCTTGATGCAGTACCTGCACTATCAGAGGATGCGCCCAGCAGACAACAATATTGGGTTTTGGCGAGAATTTACAGC
CGAATTTTTTGCCCCTTCTTCTAGAAAAAGGTGGTGCCTGTCCCAGTGCAGTACTGGTGGGCGGCATCAGCCAAATGGAATCT
TCCCACAGGAACTATGGTGTTGTGAAATATGTGGTTCAAGTCCTGGTCGTGGATTTGAAGTCAGTGTTGAGGTTATGCCTAGA
CTTTGCAAGTCCAAATTTGACAATGGGGAACTTGAGGAGCTGCTTTTTGTGGATCTTCCTCATGAATACAGACTTAGTTCAGG
ACTGATTGTTCTGGAGTATAACAAAGCTATTCAGGAGAGTGTTTATGAGCAACTTCGAGTTGTTTGGGAGGGTCAATTAAGAA
TTATTTTCACTCCAGAGCTGAAGATTCTTTCTTGGGATTTCTGTGCCAAGAGTCATGAGGAGCTCCTTCCTCGCCGAAGTATTG
TGCCTCAGGTGAATCAGCTAGTTACTCTGGCTTCTAGATATCAGGCTGCCATGGCACAGAGTAATGGAAGTGGGCTACCTGTT
CAAGATCTTCAAGCATTCTGCAAAGTGTTTGTTTCAAGTGCTCACCAACTGACCAACAGTTTGGAGCCACCCTGTGTGAATGA
GCTCGGGTTTACTAAGAGATTTGTTCGCTGCATACAGATAGCGGAAGTCGTGAATAGTATGAAGGATTTAATGGACTTCAGTC
GTGACCACAATTTGGGCCCTATTGAAAGTCTCGCTAAGTATCCAACTTCAAGGTCATCTGTAGAAGTCACAAATTTTGTCAGG
CAGAGTGAGTTGGCGGGGCTGGATCGACCTGGAGTAGAAGAGAATTCGACCTCCAGTTTCCTACAATCAAAGATTGATGATC
AAGGAAACCTGATAAGCAGGCCTGTACTTCAGCCACGGTTGTTGCAGCAGGGTACCAGCTCTTTTCCAAATTTTGCACATATT
GATTCATCCACAGTAGGGACACCAACTGCTGGAGGTGGCTCTGCAGCAAACTCCCTACAGGCCTCTCCTACCAACTCCCTAAC
TTCGTACCAGAGCTGTTTTTCTTCATTGACTAACAACTCTCAACAACCTATTACTTCTTATCAGAACCCTCTTACTGTCAATTCG
GGTAACTTGAATATACTAAACCAGCCTTCTCTTCATCAGCACATTCAATCTGGTGAGCAGATTGGCAGTGGTGCAGTGCAGCA
GTTTCTTCAAGGAATGATTAGTTCTCACCCAAGGGCAGGACAAGCTTTAGTCCAAGGAGCTTAGGGATGAACATTAGAAAT
ACAGAGGGAATGATAGGATCTGGCCTTTCTAGAAGCATATCAGGACTGAACAGTCCTGCTGGCAGCGGCAATGTCGGTGGTT
CAGGAGCCATTTCTGAAACTCGGCAGCAATGATCTTACATTTTCTGTGCGTTGATGGCCTAATTTGCCACAACTTTAGATCACG
ATACTTCTGTAATGTTTTTGCAGAGCTTGTTTTGATGCATGGCTTCATCATCAGGGATTTGGAAAATTGAGTTGACCATAACTA

ACAACAAAAAACAGAGTCACTAAATCAGCTTGATTTAAGGTTGTGCCTCTTTGGCAACAAAGCTAGAAGTGCAAACAATCAA
GAAAACAAGTATGGGGTCTTGGTATAGTAGCAAATGTACATTTTGGATGGCCATCCTCGGAAAGATAGCTTTGTCCAAAGAA
AGCTCCGTGCTGTATGGAGTTTTGTGAATGCTGTGGGCCATGGTCTTACTTTTTACTGAAGCTGCAAAATCAACATTGTGGCA
GCTTTTTGGACATGACATTCCACGTACGTAGTCGTTTGCTACCTGCCAAATAGTCAAATGGTTACTTACGTTGCAAATTGGCAG
GCTGGTAACTTGTATTTCTCATGGAAGATACGGTGGGTTTCCCAGTGGAAGTATCATAGCTGGTTTTGACCCATGTTGAATACT
TTGGCCGCCGCTTTGTATTTGTATCAAGGGTGTTAGCCACAGAGGCCATTTATGTTACCGAAAGGTGCAGCTTGCTTACTTTTT
TTTTGGCTGTGCCATTGTATGTTATGACTATTACTTTTGTAGTTTTTGGCTCAGATATTTGGAGTCTCCAAACAGTGTCTATAGT
GGCCGATCCACTCTTATTTGCATGTTAGACAGCGAAAGTTGAGTCTGTGTCGTTTGTAAAGAGGATTATGCTGTTTTTTGATCT
TTTGTACTTAATTCTTATGTTG

**FLOWERING PROMOTING FACTOR 1 (*FPF1*):**

TTAGTAGTAGACATCTTACCACATAGATCAGCATGCTATAGGTCGCGGTCCTCGAAGTTATAGGTCGCGGACCTCGAA
GTACTCGCGATTTTTGAGGACAATGTCGTACATGTGCATTGTTTTGAATTTGCGGAAGTCGGAGGGGAGGGAGATGAGGAGG
GAGGAGGAGTAGCAGCGATGGTATTGGAGCGAGTCATCGGAGGCGGCGGACGAATAGTATCGCTCCCAACCGAGGCCCTGG
AGCTTTTCCTCTAAATCTGAGTAGGAGGAAACAACTTCACTAGTGGGTAGATAAACTAAGGCTTTCCTTCTGCCCTGCTGCCA
CTCTTGGCCTCCATCAATAGGCTCCACCATGGGGTTTGAACAAGCTTTGCTACACCGTTCTTGAAAACCCATACACCTGCCAT
TGCCTCCTGCACTCTTGTGTGTGTAGCCCACCTGCTTTTGTTGCTTTTCGCTCTCTGAAGCCCACCTGCTTTTGTTGCTTTTTACA
CTTAGCCCACCTGCTTTTTGTTGATTTTTGCTCACAGCGCGCTTGTGTCACTTCTGTGTGCGTAGCCCACCTGAGTGTCATTTCT
CTGCTGCATTATGTTAGTGTGCGTGCCAGCCAGGAGATGATGTGAGCTGTACATAGAGCTGTGTAGTGTGAGGTAGGCA

**GIBBERELLIN 20 OXIDASE 1 (*GA20OX1*):**

TACCCGCCATGCCCGGACCCGAGCAAGATCTTGGGGCTCGGGCAACACACAGACCCGCAGAGTCTCACGCTGCTGCT
GCAAGATGAGGTGGGCGGTTTGCAGGTGCTCATGGAGGACGTCTGGGTCGCTGTCAAGCCTCGCCCCGATTGCTTTGTTGTCA
ACGTAGGAGACACTCTCGAGGCATGGAGCAATAGTAGACTTAGAAGTGGCATACATCGTGCTGTGGTCAACACTACTTTCCCT
CGTCTATCAATGGCTTATTTCCTTAGCCCTGCACTTAGCACTCTCATATTACCTCCACCTTCATTAGTGGACAAAGATGACCCT
CTAAAGTATAAGCCCTTCACGTGGCTTGACTTTCAAACTGAGCTTCTCAGACAAAGACGAGTAGTTGGTAAACAAGCGCTCAA
CAAGTTCTTCATTG

**TERMINAL FLOWER 1 (*TFL1*):**

TTTTCATCCAAGTTTTTAAATGAGCGTGCAGACTTGGTTCCTGCATACACATACACGATGGCAGGCCACGTTCGTTACC
TGTTTTTAAGTACTCAAGCACGCAGGCGGCAGAGGGCGAGCCATGGCGAGCGAGAAGCTATGGCTTCTGTGGTTGGTGATGG
GCGTTGGGGTGGTGGGAAGCAGGGCAGCAGCAATAGAAAAGGAAGAGAAGGGCTGTGTGATTCCGCCGTGGGTGGATGCCT
ATGCGGCGCCGCGTGTGAAAGTGTCCCTTTCCTTCCTCGCCACGCCCGTCCGCTCGGGCCAGCTTCTTCCCAGAAACCTTACCC
AGATTCGCCCTCATGTCTCCCTGCACGTGCCTCCTCTCCTTGCCCATAGTTTCCTTACCCTCGTCCTCGTCGACCCCGATGCCCC
CTCCCCTTCCTCCCCCTCCGTTTCCAATATCCTTCACTGGATTGTCTCCAACATCCCCTCTGCCCACCTTTCCGAACAAGAGATA
TGGGAGTTGGGTGTAGAGAATGTACCTTACAAAGGGCCTGCACCTCCCAATGGAACTCATCGTTATTATGCCTTGGTCTTTGA
GCAGAAAGACAAGATCCATGTCGAGCTTATTCAGAACAGAGCAAACTTCAGCGTTCGAGATTTCACCCGAAGATATGATTTG
TACTATCCGATTGGAGGGACTTTCTTCAGAGTTCGTGCCAATACGTAAAATTGGTGTCTTGCTGTATATTATCATGCATTGTGA
ATTGATTAAGCATTCTGATCAATACATTTTCTCAAATTCTAAACTCAATAAAGCTACACATTGACATGATATACGTGCTATGTG
CTTAACATGGTGAGTGTGCAAAAGTTCATAGGGAAACCAAAGATGAAGAAAATGAAAGCATTTTGGAATCCCGGGCATCATT
TGGATGCACACAGTACTACAGATGCAAAAAGT

**MORF RELATED GENE 1 (*MRG1*):**

CTATAATGACTCAAAAAAAAAAAATCTAGCATTGATTCATGAATTACCTTTGAAGTCATCTAAAGGATGGACCATGTAT
GAGACTTCTCTTAACAGCTCTCTCCAGCTTTCATAGTGATCTCACTAATGACTGATAAAGCTGGAGTGCTGTGTCAAAATGAG
CAAAATCACTTACTGCTCTGATTAGGCAATATGGTCTGGTTTATCTTGCAGCCAGTCACTTTTCTAGTACGTTATGGCCCACAT
AATTTGGAATTTTAATGTCACTTTTGGACTGTAATATTGCCAAGCATTTCTAAGAGCAAACATCTCATTGAGACCAGCATACA
TTTGCAGCAGCAGTTGCACTACTGCACACAGAGCAACGGGGAAGAAACACACATTTTGTCATCTTCCACAGGCTCAAATTGCC
GCCTGGATGAGTTATCTGCAATTGATGTAATTAAGCTGCTGCATACTTCTATGCTAGTTCGTGTCTATTCTATTGCTCGCTATG
CACCAGCACTCCAAACTAGTTTGCCTTCACTTCATTATCCTGCAATCGGCTTTTGCTTTCTTCACTCCAGGAGACCTTCCCAGT
AGCGTTGATAAATTACCAGCAAACACAGAAAAAGCACGGAGCATGCTTCAATTGCAGCCATGCTAGCGAAGAGAAACAAAA
GCCATTGAACACCATGCAGCTCATAGAAACCGATGCTGCTTCGGCTTGCATGTCACATCCTCTTGACCAAACCCATGCCCTTC
ACCATCCCTGCTAGGAGATTATAAGCAACCTCAAAACTTACCAATGTGAAAGAAGCACCTCAAGGGTGCCATCTGATACAAG
AGGCAATTAACATTTTTAAATGAGGCTTTGTCTACAAGCTTAAGGAGTGCTTTGAGTGAAACAAAACCCTATGTGAATGCACT
CGGGTATTTTTTCATAAAGACCTGCAATTCTCCACCATTGGAAGACACGTGGACCAGATAATCTGCAATAATTCCCTTAGGTT
GAAGTCTATAATATTCTGCGTCATTTCCAAAATGTGTAATTCTAGCAGCTAAGTAACCCTAAAAATTTGAGAGTATTACTTAC
ATTCTTACCTCCCCCAGGGGTGTTAGAAGCTAGTTTATATGGTACAGGCAAGTAGAGTACACAGAAGTGTTGTAGCAAATACA
AGAAATACAGTCCATTATAGGGAGACGTTTAATTAGAATTTCCTTCATTGGTTATGGTGGTGCACTAAACTTGGATGACAACC
CTAGCAGCCTGTGATCAACTTTGTAAGGAGCACTTGTTAAGAATGCCTCTCCTTTCCTTCATCTACCCCAATATCAACCTTTGG
TCCATCATAGGTGGACAAGAAGAAGACACTCTGGTTCTTCTGCAAATACTTGAGAAAATCTGCCAGCTTCTGTTGCAACTGTG
TGAGAGCATCATCCTCCATGTTGGTGTACACAAGCAATTCAGGCAGTTTCACAAATAGGCGCAAAAGATGCTCTGCTCCATAG
ACAGACGAGGGGGCAATAGTGCTATTTTCAGGTACATGCTCGAGATATTGCCCTCGCTCTGGTTTATACAGCAACATGGCAGG
TAGAGACTTGTCAAAGTATGTGCGTAAGCCATTCAAAATCTCCACCAAAGCATCACCAACAACGCCTTCCTGCTTTGTTTTGA
ATTCCAAATACTTTTTTAGAATGTCGTCGACAGTGGGGTGACGTGGAAGCTTTATCAGCTTGCCGAGTTGTGTTACAAACTCC
CAATCATCAACCAGCTGCTTCTTTAAAGTGCCAGGAAGGGGAATCTTCATCACTTGTTCTGGTTCATCAGTATCCCTATCCTCA
ACACCTGAATCTGCTTTGCGCTTTTTCCCCCTTGTCACTGCATTACAAATAACAAAGTAAACAGTGCTGTAAGCCATGATGAT
ATGTATCCCCATTGACACAACATCTAGCATGAGCTGTCCCTCCCTTATAAAGCTTGAAAAAAACAAAAACCTTTACACATAGA
ACATAACTTGCTATCTCTCCTAACCTACCAACTGGCAGACCTTAGACCCAATTCAGGTACAAACTCAATCAGAGCCACCCTTC
ACTGTAACAGTTTGCTTGCCGTGGAAAATCTCACAGGTCAACTACTCTCATTGGCTGGATTATTATCTCAGCACTCGGCACAC
AAAAGACACAAAGCGAGAAGTTTGTAACGATGAAATCTGTGAGAAGAAAAGGTGGACCGGCTGAGGCCAAATGCAAAGGAA
AAAGCAAACAGAAACAAGAAAAACATCAAAAAAAACTAGAATCATTTAAAGAAATGAACTTGATGAAGATTTGGCTGGCTA
GGCATTCCACCAACATAAGCTATCATGTAGATCAACATGCTCATTTTGTGTGTTTTTTGATTGGCTCAATACCCCTCAGCTTGA
TGGGTTGGCATATGTTAATGCAACCTTCGAGTCTTCGAGAAAAGTGGAGGACTAGATGGCATAGATCCTGTGAGTTGGCTTCC
AAAGCAAGTGGATCATCGACAGATTGTAATCGAGGCTGCAGATATCTCGCGTATTCTTCCACACACAACAGAGGATAATGCA
TATGATTTTATTCGCAAGGATATGATGAGGTTCACACGGCAGACTCGTTCAACTACAGCTCGTCGTCGTGGAGAGTCTGGTGG
TGGTACTAGTAGTTCTTAGCCTCGTAGAGGTCGTGGACGAGGTCGAGGAGGTAGCTCCAGACAGTGTATTGCAGGGCCTTCTA
GTCCTAGGCGACTTACACCCACTCCTTCAGAGTGTGAGAGTGATGACTCATCCCATTCAGAGGGAA

**UNUSUAL FLORAL ORGANS (*UFO*):**

TCTCTCTCTGTCTCTCTCTCTGCATGGAGTTCACATAGGGGCCGCGGGGGGTACGAAGAGTAGACGCTGACTGACCGG GAGACAAAAGAGAGCAAGCGAAAGAGAAATTGAGAGGGAGCGCCCGTGTTCTTCTGTGATGCTGGGACAGGTGGAGGACCC AACAGCTGCAACGCTGGAGTGGAGTAGCTTACTGCTCCGTTCCTTTTTGATGGAAGCGGGGCTCTCTATGGTAGAGATGGACG CCCCTTTCATGAGCTATTATCTGGATCACACCGGCGGCGCCCTTGTCGACACCGGTGGCGGTGGCTATGGCGCTGCTGCTCGC TTGGAGCAACAACAGCAACAGAAACTCCAGGATGAAGACGATGATGATGATGACATGCAACACCTCATGATGCAACCCCCTA AGAAGAAGCTCATGATGAATGCTGTCGCTGACAGCAGCTCCGTTGTGACTTCTGTTGATAACATAATGCAGGGAGGCTCTGGC TCTGGTGCCGACACAGCTGCTGCTGCGTCGCCCTGGCTCGACCCTCGCCTCTGGAGCAAGCTCCCTGAGAAGCTTGTGGAGAG GGTTGTCGCCAGCCTGCCCCTCCCAAGCTTCTTCCGGTCGCGGCTCGTTTGCAAGAGGTGGTACAGTCTCTTGTTTTCCGACAG CTTTCTTGAACTGTGCGCGAAGGTGCGCCCGGCCAGGCCCTGGTTCCTGCTCTTCCGGCGGGGGGTCTGGTCGGAGGCCTTTG TGTTCGATGCCGTGGGGAAGGCGTGGTTCCGCTTGGACCTCTCCTTTCTGCCCCCGCGGTTCACTGTGGTAGCGGCGGCGGGC GGCCTGCTCTGCTGCATTTCTGAGGCACGGGGCTGCAAGACAGTGCTTATCTGCAACCCCCTCACCCGTGTCTGTGTGCAGCT TCCCGCCGCCCTTAAGGAGCGCTTTGTTCCAACTGTGGGCCTTGTTGTCGACTCCATCACCAAGGCTTACCGTGTTGTTGTTGC AGGGGATGATCTCATCTCCCCCTTCGCCGTCAAAAACCTCACCACTGAAATGTATGACTCTCGCCTGCAGCAGTGGCGCATGA CTGCCCCCTTACCTCGCCTCTGCAATCTTGAGTCCGGCAAGACAACCTATGCTAATGGCTTCTTCTACTGCATGAATTACAGCC CTTTCAGCGTTTTGGCATATGATACTGAGCAGGGAATATGGAGCAAGATCCAGGCACCCATGAGACGCTTCCTTCGTACGCCC AATTTGGTGGAGTGCCGTGGCCGTCTGGTGCTGGTGGCAGCAGTGGAGAAAAATAAGTTGAATGTGCCGAAGAGCATCAGAA TTTGGGGGCTGCAGCACTCGCGCACAAGCTGGGTGGAGTTGGAGCGCATGCCACAAGGCTTATATGAGGATTTTATGCGTGT GTCAGGGCACAAAGCATTCCACTGTATTGGTCATGGTAACCTCATCCTTATCACACTCCCCGACTGCCCCGACATGCTGCTAT ATGACTTCTATGAAAAAGTGTGGCGCTGGGCGCCGCGCTGCCCTTTTGCAGGTCACCCACCGCCCCCTCATGCCGCAGCCTCT CATTCGCTTTCAACAGGACCTCAAGGATTTCATGCCTTTGCCTTTGATCCCAGGCTGGAAGCGTCTGTCTACTGATGATGAATT GACTGATGATGAAGATCAGTTGAGTGCTGTTGTTGATGATCAGGTTGGTGCTTCGCTACATTGCTTCTTATCGTTTCATTTCAA TTCTGCATCTCTAGGGTCAAAGAACTGGACTAGTGGCATTGCTCTCTTTTTTGGGACCTACTTCTCCTGTGTCTAAGACAAAAT TACTGTCTATTATGGGACATCTTGCTGCTTTTTGTCATGATGTTCGTGGGAAATTAAAAGACTCTTGTTTTTCATTGCTGAGTG ATCCTATCTCAAACCTGTGATCCAAGCTTCTGCCAATTTACCTGTCA

**VERNALIZATION INSENSITIVE 3 (*VIN3*):**

AGAGAGAGAGAGAGAGAGAGATGTTGGTGAAGAGTACCTACGCCCTTATGTGAGAGCCCTTAACGCATAAAATCACTCC AGATCTGTGGCGCATAACTAAATGTCACCGCCCAAAACGTCCCAATTGGAGCCCACTTCAGGAGCGTATTTTTTAGGAGCATA TAATCCAACGATGGGAGGCCTAAGCCTATCTGAAAGGCAGGAAATCTTGTATCAAGCTGCACAAGGGGCTGAGGGGGGCCGT GAAGCCCTCGAGTCATGGACACGGAAAGATTTACTGCAACTTATTTGCATTGAAATGGGCAAGGAGAGAAAGTATACAGGTG TTTCCAAAGTCAAGATGGTAGAGCATCTGCTAAAGCTGGTTTCTGCGAAGGAACTGAGAGAAGGGGCAGACCCTGATTTTAC TTCTCCAATATCTCAGTTTCCAGCACCACAGAGCTCATCTAGGCGGCAGCGAAAAGCAGGCCGTCCTGCCCGAGTTCCAGCAG CTATCCAAGTGCCGGCAGCTGCAAGCATATCAAAGCCAGAGCTTTCTTGGGTGTGTAGAAACACAGCATGCAGAGCCCAGCT GCCACAGGGTGTGAGTTTTTGCCAGCGTTGCTCTTGCTGTATTTGCAAGAAGTTTGATGATAACAAGGACCCTAGTCTTTGGA TAGTTTGCACACCTGAGCCACTCAATGTAGAAATGGACTGCAGGCTATCTTGTCACATAGAGTGTGCTCTCAGCAGTGACATG GCTGGGGTAGTGTTGGATGGACCAGATATTCTACTTGATGGAAGCTATCAGTGTTGTTCTTGTGGCAAAATAAGTGATGTGAT AGGTTGCTGGAAAAAGCAGCTATTGATTGCAAAGGATGCTAGAAGAGTCGATACTCTTTGCCAACGTCTCTCTTTGAGTTACA GGTTACTAAATGGCACCTACAAGCACAAGTCACTTCATCAGCTTGTAGAAAAAGCAATCCACAAGCTGGAAGCCGAAGTGGG CTCAATCACTGAAGGGTCTGCTAAGTTTGCTCGTGGTTTGGTGAATAGACTCTCTTCAAGTAGTGAGGTACTTGAATTGGTAA TTTTGGCCCTTGAAAAAGTAGATGTTCTTGATGAAGAGCCGGTTTCGAATCACAAAGCAGGAACAGAGACGGTAACTATCCA AGAGAAGAATGCCGGTATGTTGTGCACGATAGAATTTGATGATGTATCATCATCTTCTATTGTGTTGACTGTAAAGGGTGGTG GGGAGCTTGTGATTGGCTATCGGATTTGGCATCGCAAAGCATGTGATGCTAGCTTTGCAAACAATCCAACTTGCATAATCACT GCCAATCCTGGGAGGGCTCAGATTTCGAGCCTGCACGCTTGCACAGAGTATGCAATCTATATCGTTCCATTCTTTGAGAGAGG GATTGGAGAGCCAGCAGAAGCTCGATGCTTCACAAAAAGTGTGGAGTTGCAATTACCCAAGGAGGCATGTAAGGACGGGCA CCTTAACACTGGCTTGAATGTTAGTATTAAGGAGGATGTTTGCATTTCTGACAAGCTCGAATCCAATTTCAAGGTTCGTGAAC TTGGCAAGGTCTTGCATTCCGCCTGGGCTGAAGAGAAACAGTCTGCCCATGTTCGGAAAGGTATTTTTTGTGGTAAAGGTGGG GTTACAGGCTTGATTGATCGCAAGAACAATGAAGAGAAAAAGAATGAATCTACTTGTTTGGGATTGAACGTTCTCTCAAGCC CTGGAGCAGACTTGTCCCATGTGGACAACAGTACGTGTAAAATAAGCCCTGAAAAAGCACCTAGCAGCGAATTGAATGCCTC AGTAAATTTAGAAGCTGGTGTGGAAGAAAGTAGAGTAACTGTGGAGGTAGAGGTGGCTCCACATCCGATTAGAACCCTTTCG CGGAGAGATTCAAGTGTTTTATTACAAGAAGGTGAGGCTAAAGGCCGGCCTGCATGTGCTGAGGTTACAGATTTGGATGACA CAAGTGACCCTGGAAATTCTCACACTTCACAAAGGGAAGAAACGGAGGCCCTATCACAGGTTGTTTCGGATGGAATGCAGAA CAATGCAGCAAATGAACTGCACGATGTTTTAGACAATAGGCGTAGAATCGGAGATGGTGAGTCCAATGGGCATGGAGAAAG CTGGGCGATGCAAGTTCGAAGTGCAGGAACAACTATCGGAATGGAGCCCCAGACAGCTATTATGAGGAAGAGGACGGCGGA AGGGCTTGGTAGGGGTGATAGTTACGGGCTTGTGAATGGTTGTGGAGGAACTGTGGGAGGTACCCTTTGTGCTGCCCGCAACT ATGAGTTCTGTGTGAAGATAGTGCGCTGGTTGGAATGTGAAGGCTATCTAAAGGAAGACTTTAGAATGAAGTTTCTCACCTGG TTTAGCTTGAAGGCATCAGAGCATGAGAAAAGGGTCGTCAGTGTATTTATTGATACACTGCAAGACAATCCTGCAAGTTTGGC AGGGCAGTTGGTGGACACTTTCTCAGATATCGTTTCCTCAAAGAGACACCACATGGTTTCCAACGGCTTCCGTAATAAGCTGT GGCACTAGATTGAAAAAGTGCAACAAACAAGCTGAAACTAAGCAACCTAGTTTAGCTCTAAGTTTGTTTTTGGTGTTTTTATT TTTTTCATTTATATAGCTGCAAACAAAGCATTATTTGTGGGTTGATTGATGTATCATTTGTGATTGTCCTTCAGCTAGACAATT GCCTAGTTCTTAGAATGGCCAAACATTATCTCTTTACTCCTTGCAGATGCTTGGCCTTCAGTCCCCCCCCCCCCTTCTCCTGGG CGCCCGCCGTGCCCCATCAAAACGAATCACTCAGTGCTCACCAGTTATTTCTGGTAAAGTAAGTATGTTATGCCAGCACATTT GAACCTCTGCTTTTGATATAGCCTGTCATTATTGAAGTTATTTTGTTGTGATATTTGCTCTTCCTGTCTTCCATTGGTTCTCTATT TTGCTGTCATGCGCTCAAGTTGTTTATGGGCCTTCGGTGTCGAATGACTGAGCGTAAGTCGTGTTTTTTACTCTTTTAGCCTAG CAAAATTTGCACTTTTGGTCAATTTACCCTGGTTCTATTGCCAGTATGGGACTT

**PROTEIN ARGININE METHYLTRANSFERASE 10 (*PRMT10*):**

GAGAGAGAGAGAGAGAGAGCAGGCAGCGAAACCATGAGCAGAGGAGCTGGTGGAGCTTCCATGGCTGCTCCTGCTGTC GACAAGGCCGCCGATTACGCCAATTACTTCTGCACCTACGCGTTCCTCTTCCATCAGAAGGAGATGCTCTCTGATTGCGTCCG CATGGATGCCTACCACGATTCCATCTTCAAGAACTCCAAGCACTTCAAGGACAAGATTATTCTGGATGTTGGAACAGGCAGTG GAATTCTTGCCATATGGGCTGCGCAAGCAGGTGCAAAGAAGGTTTATGCTGTAGAAGCCACAAAAATGGCTGAACATGCTCG CCGATTAGCTGCTGGCAATGGGGTGGAGAATATTGTGGAGGTAATCGAGGGATCTATCGAGGACATAGAGCTACCTGAAAAA GTTGATGTCATCATATCAGAGTGGATGGGCTACTTCCTCATACGCGAGTCTATGTTTGATTCAGTCATTCATGCACGAGATCG GTGGTTGAAACCAACAGGGATCATGTATCCAAGTCATGCAAGAATGTGGGTTGCTCCTATGAGATCAGGTCTTGGGAAGGTC AAGTTGCAAGATTATGAAAATGTGATGGTTGATTGGGATGCGTTCGTTCAGGATACCCAGGATTATTATGGTGTGGATATGAG TGTTCTTACTGATCCCTTCCAAGAAGAGCAAAAGAAATACTACCTACAGACAGTTTTATGGAATAATCTTCATCCGACTCAAA TTGTTGGAACTCCTGTGATTGTAAAAGAGTTTGATTGTCTTACTGCTACTCTCAAGGATGTGGCTAGTGTTCATTCCTCGTTTC

AAATGCAATTATGGGAAAGCCAGACGAAGGTCTCTGGCTTCGCTGGATGGTTTGATGTCCATTTTAAGGGGAATGCAGGTGA
GCCGGCTGATCAAATAGTGGAATTAACAACAGCCCCGAGTGCAAATAAATCAACTCATTGGGGCCAGCAGGTTTTTCTTTTAC
ACCCACCTTTAGATTTTGAATCTGGGGATGTAATTGAAGGCGAACTGGACATCACACGAGCCAAGGACAACCACAGATTGCT
GGATCTCTCATTCAAACACAGATTGATCAAGTCCAATAGTAGTGGTGCCACGGGACCATTCACCTGCTCAAATTTCTACCTTG
AATGAGGCTATGTATGCATTTTTGTTCCCGGTCTGGCCTATTTGCAGGTGCACATTTTGAGTGAATTTTGGGAAAAGGGATAT
ACCTTTAGCTATTATATTGGTGAAAGTAGTAACTAGCTACTCATCACTGGCATATCCAAGGGTCCCTGGCTTTTCACGTATTCT
TTATGGACCATCTGCTGTATTTTCGCTTGGGATTGCAGGTTGCTTCGTACCTTTCCATCGAGAATTGTAAAGAAGTATAACATT
GCCGCTGTTTCTCGCGGCTTGTTGCATCAACAGTTTTTGCCAAGTTTTACCAACAATTTTTGCCTTGCCAAGTCTTATGGACAA
AGTCTACCAGATTTTTACTTGTTGAGCGGCTCTCTAAGTTGATTCCGCTGAGAGTTTTGTAACTTGATTATATGGCACGTAGAC
CTAAAAGTAAATTGCAATTTTAGAACCCCAGTAGAGTTTTCCCCCCTGAAATCAAGTTTGGGTGAACGTATATGGTGATGCGA
AAATTGCTGCTTCATCAAGCAGCTAGCATAAACATTGCGAGCAACGAGAGATGCTACATGGGAGATAATATGTGGAGTGCAT
GTAATTTGTTAGGTTGTGTGATGGAAGGCATGACCTTCTCATGCAATTTATTAAAAAGGAGTGTTTTGTGAATCCTGCTCTTTG
CGTTTCATTCAAGGAAATATAGAAGACGCACGGGACAGGTGGTCGAGCG

**AT-HOOK MOTIF NUCLEAR-LOCALIZED PROTEIN 21 (*AHL21*):**

TACCTCGAAAGGATTGAAGACACAGCTCCAAGCACTAGCACTATTGGTGTCAAAAGGATGGCATCTCTATTGAAGCTT
GGCTTTCAAGGTAGTTTGTGGGTCCTAGAGCACATGGACAAGCACAACTTGTTCACGCATAGCTTGAGGGATGGATCTTGGCT
CTTAAAGACAAATGTAACTAGGAAGCTTGTGTAGTACATGGGTTTGGACCTTTTACCTTTATTTGTGTGATTAATCTTGCTAGG
GGATCTAGGGGCCCAATGTTAGAATAACATCAATACGGTGGTTTTTGTGTGACTGGTCCCAATTGCTCTTGCTAGGGGGTATC
GAGGGACCTAGTGCTAGAATAACATCAATGAGGTGGAGGGAGGGACACCTTGGGCCAAGGAAGAGTGACATGTGACAAAAT
AGAAGGATGTTAGTAAGTTAGAGAACGTATTGTAACAATGTTAAAGAAAGAGATAATTCTTGTTAAGGAGGAACTAGCTAGT
GAAATAAGACATTTTTGGAGTGACATGAACAAGGAAGGGGTACCATGGGCCAAGCAAGGATAGGCATGATGTGGGAAATGA
GAATCTTGGAGGAGGTTGGGAGGCAAGGTAAAAATTGCCATAATATTAGGTTAGGTTGGGGTGGATATGGTTTCGAAAAAAG
ACCTGCTAGGATCGGGGACGTGAAAGTTTGAGGCCATCAATATGGGGGGCGAGGAGGAGGCGCTATCGACCAGGCATAGGA
GTCATGAGGCAATTGAGAGTTTGGGAGAAGGTTAGGGGGCAAGTTAAAAATGGCCATATTATTAAGGTTTAAGTTAGGTTGG
GGAACAATAGCTGAGGAGGACTCACCAAGGCTAGGATGAGCATTATGATTAGTATTATTTGAGTCTTCGTTTTTATTTTGTAG
TTGAAAAGGCTCTTCATTATCTGTGACGGGAAGCTGGCGGTCCGAGTCCTCAAGGGGCAAGCGGTCGTAGGAGGCGTTTAGG
AAGGAGGCGGCAATTACGATGACGGGCCCAGCGGCAAGTAGGTTACCAACAACACTACCACCTATGACATTACCTTGGCCAC
CGGCTAAGTAGACGGTCAAGCCGGTAGCCCCAGGGGGGGCCGGATGTGGAAGAAATGCTCCCGAGAGAGAGAGAATTTCAA
AGCTCCCTTGTAGAGTTACCGTTGAACCGGGTGCAGCGGGTTGTCGTAGCGTCACGTTGTTCACGGAGCCACTGCCGCTTAGA
ATGCAAAGACCACGATGTCGAACACGCGCAAAATTTGCGACGCACTCAACCACGTCACACGTGCTTGCGATCTCGAGCACAT
GTGCGCGCAATGCGGTGGCACTGTCCTTGCTTATGATGATGGGCGGCTTGGGTTTGTTCTTGGAGCCTAGGGGTCGGCCGCGT
GGGCGCCTTCCGATTTGTCCGGCCATAAGCTCTTTGCCCTTGTTCCGTCTCTTTGATTTGTTGGTATCGCCACTCATTAAACCCC
CGCTGCTTGGGCCGCTGCTCTCGCCGTCGTTGTCGTTGGCGTCGTCGTCAATATTGTTGTGTGCTACGCCGGCTTGCCGTCTCG
CCTCTTCTTCGCTGGCAACCAAGGCCTGCCTTGCCTGCATCGCTGAGATCATATTGCCAACTGTAGTCGATGCCGTTTTCAATA
TCAATGTGGATACCGGCATGCTAGAGAGAGGGAGAGAGAGAGAGA

**KNOTTED-LIKE FROM *ARABIDOPSIS THALIANA* (*KNAT1*):**

TAAATGGGAGTATATCATGAACTAAAAGAAAAAGAATTCTATGGCTTAGGAATCAAAAGGCCAACCTGCACAACCTC
TAGCTGATTAATTTGCAAATTGCACAATTTTGTATTGAGAGCGCCACGGGATTGATGTGTAAACATGAATAGTAAAGAGCAAC
ATAAACTACAAAAGGGATAACCTTTTAATGCGAAAATTCCAAACTGACATAATGTTGAGAGGGGGAGAAAGAAAGGGAGAT
GTAGCTTGCTTTTATTGGGCCACTGAACACTCGGTTTGCTGCAAGAATGATCCTCAATTCAGTGTTGGGAGGATGATCCTTAAT
GCAGAGTTGGGAGGATCATAGCTGCAGCTGCGGCTGCTGCTCCCGCTGCAGCATCCATGGTCTTGACAACGGCATAGGAAGT
AAGCGAATGAGGAGCAAGATGTCCATCTGTTGCAGGTGCATGTGCATGCAGAAGATGATGGTGATGGGCTCTTGAGTTGTCC
ACCATTACATACTGCATATACTCTGACGGTTTCCAGTGTCGCCTGCGCTGATTTATGAACCAACTGTTTATTTGCGTCTGATCA
AGCCCTGTTGACTCAGCTAAGCCTGCTTTCTCTGCCTCAGATGGGTACGGCCATTTGTAGTGTTGGCTCCACCAGGCCAAGAG
CTTCTGCCTTGCATCCTTGGGCAGTTTTCCTTTCTTCTTCTTCTTCAGGAACTCCTGCTTCAGGCTGCAAATGGAGCCGCTGTAC
TTGCGCAACAGCTGTTCTTTTAGCTTCTGATCCTCGGCAAGTGGGTCTATCCTCTCCTTGTGAAATTCGACCTCCCTGCAGCTG
GCGTCTCCCACTTCTTCCTCCACAGACGACCCCCCAGTGTTGGCCCATTGTTGGGCATTGGCCTCCCCTTTCTCGTCACCTGAT
TGAGATAGGCGCAGGGTTCCCTTGCTCAGCGAATTCAACTGCAGCTCTATCATTCTGAAGAAGGCCATGGCCTCCTTATAGGG
CTGCATCAGCTCGTCGTGATACTTCTGCAGCACGTTGCAATACGTCTCCATGAAGTGGTCCAACTCAGGGTCTGCACCCAAGG
CTGCTGGAGAGGATGATCTAAAGGTCTGGTACTTCTTGCTAATCTCGTCCAAACTTGAAACAACTTCTGCGGGAGCTCCAATC
TGCAGACACATACATGTCAAAAATCAAGTAAGCGACAGTTGTAATCATTGAGTACCCAAACAGGCTCTTACACAGACAGTAG
CGTGTGAGATAATTCTAAGGAGTCTTGTAGTATTGCGACAGTTACATAATTTCCATGCTGTAGATTCGTTATTTTTGTCGATTA
TTTGAAAAAGGCAGGCAAATTCACAACCTTGTGGCAATTCACATACGCCATAACAAGGCATGGGTATGTTCCGTGGGACACA
ATTTTCGCTCTCATTAGGTCTGTACTGTGCTCTGCCGAGGAATGTGTATTCTCCACCTCCATGGAGACCCTCCCCACAGCTGCT
CTACTTTCATGCATCATTGAGTTCGCGCTCGGGCTCACAAGCAGCATTGGAACACTCGTGGTAATCACATAACTACAATCACG
TCCACCTACCAATTGGGCACCCTGCACGGCTTTGGGTGCCCCTACATCAGGCTTCTCTTCCTCAGTCCGGAAGATGACAGTAC
TAGGTTCGAGATGATCAAGCGCAGAGATAATGGGAGTGTGCGTTTCAACATGATGGGATTGGTTGAAAGCATGCGTTGCTTG
GAGGAGGCCAGACTGCAGGCAAGGTTCGACGATCGCCGCATTGTAGTATGTGAAAGAAGGTGGAATCTGCGAAGCAGCGGA
GACAGTATTGGAATGCAAAATACTGAGCTGCTGCCTCTGATCATTATGGTACTGTTGATGGGGAGATCCCTGAGTATTATGAT
GGGCAGGTTGATGCGGATCATGTTGGGGATGATGATCACGATGATAATCAGTTTCTGCACCTAGATCTGTGGTCATCAGCGCC
ATCATCGCCCCCACTTGGCCCTCGCCATAGCTCACCGCATTGCATAAGCCTGAATACTGCATAGCTGCCCTCATGTGGCTGTC
ATAGTCCATCTCGTCTACCATCCTATCTGTCCCAACTTAAGATCTGTGTCCGCACCTTTAATCGTTGCTTGTAGTCAAATCCTG
CATGCTGCATACCCGCTGGAGAGGGGAGAATGAGATGCACGAGGAAGCATGGGAAGAGAGACAAGGGAAAGCTGGTATTAC
ATTACTCTCTGCTCAAGGAAGCGACTGCGCATTTTCTCATCCGTTTCAGCAACATGCCTCCGTAATGTCCCCTCTTGCCCAAAA
AAGCACCGTGGTAAGTTGCTAGTAAACGAGGGGCGGAGGGCGCTGGAGCGCCTATGTGCGCGCAGGCAGAGAGAGAGAGAG
AGAGGAGTAGACAGCATATCGATAAGGCAAAAGTGTGCAGAGAGCAGCAGTAGCTCCTCATTTGAGAGAGAAAGGGCCTAC
ATAGAGAGGAAAGACGTCCCAGAGAGCATTCTTGAGAGAGTGGCTGGCGTTGTAGCAGAGAGAGAGAGAGAGAGAG

**GLYCINE-RICH PROTEIN 2B (*GRP2B*):**

AGCAGAGCCGTCCGATGACCATTATACTGTGTACCAGGTGAAAACCCTAAATCCCAATTGAAAAGCCCTAACCTGCCT
TCTCACAGCTCCCGTGCTTGCATTTCACGCGAGCCAGCCCCTCTTCTCTGCTCCATCTGAGCTTCGCGGCTATGGCCAAGAGGA
CAGGAAAGGTTAAGTGGTTCAACGTCACCAAGGGCTTCGGCTTCATCACCCCTGATGATGGCTCCGAAGATCTCTTCGTTCAT
CAGACCTCCATTCTCTCTGAGGGCTTCCGCTCCTTGGCTGATGGCGAGGAGGTCGAGTTTACCATCGAGCTTAGTGACGATGG
CCGCACCAAGGCTGTCGATGTAACTGGACCCAACGGAGAGCCTGTCCAAGGAGGCGATGGAGGTGGTGGCGGTGGAGGTGG
TGGCTATGGTGGCGGAGGAGGTGGAGGCCGTGGCGGGGGAACTAGGGGCCGTAGTAGAGGAGGGCGAGGAGGA

**MULTICOPY SUPPRESSOR OF IRA 4 (*MSI4*):**

AACTTGTTTCCCTTAAACCTACTTGGTACCTGTAAGTGGCAGTCTACTCGAGTTCCCGCCTGCGAAAGAGCTTCATAAT
CTGCACGAAGCTGAAGAGTTCATCGCGCTCTCGTCCTTCGCAACGAGAACTCTCCCCTGGTTAGGGTTCTTTCTGTTGATCTTG
GCTCACTGAGATGCGGAGCGGTGATGGTGGTCCTGTAGAAGATCGCTACACACAATGGAAGTCCCTTGTTCCAGTTCTCTACG
ACTGGCTTGCAAACCATAATTTAGTGTGGCCTTCCCTCTCTTGCAGATGGGGCTCACAGTTGGAACAAGCGACATACAAAAAT
CGTCAACGCCTATATTTATCTGAACAGACTGATGGAACTGTTCCCAACACTTTGGTAATTGCAAATTGTGAAGTAGTGAAGCC
AAGAGTTGCAGCTGCTGAACATATTTCACAGTTCAACGAAGAAGCCAGGTCCCCTTTTGTAAAGAAGCACAAGACAATCATT
CATCCAGGAGAGGTCAACAGGATCAGAGAATTGCCTCAGAAGAGACAGATAATTGCAACTCATACAGATGGTCCAGAGGTGT
TTGTCTGGAATGTTGATACTCAACCAAACTGCCAAGTTGCTGTGGGCGCAAGTGTTTCAAAACCGGACCTGACTTTGACGGGA
CACACAGACAACGCAGAGTTTGCCCTTGCACTGAGTCCAGCTGCACCGCATGTTCTCTCTGGAGGCAAAGACCAATGTGTAGT
ACTGTGGAGTATAGAAGACTTTACAACTTCAGTTAAGGACCCTTCACCTTCCAAGCCTGTGACACCAACAGGAGGCAAGCAA
AAAGGTAGTGGACATCTCAGTCCTCACTGGCCGATTCTACAAAGGTTGCACCTCGTGGAATTTTTAAAGGACACTCAGAAAC
TGTGGAGGACGTACAGTTTCATCCTTCAAGCGATCAAGAATTCTGCAGCGTAGGAGATGATTCATGCCTGATTTTATGGGATG
CTCGTGCTGGACACGAGCCTGCATTGAAGATTGAGAAAGCACATGATGCTGATCTTCATTGTGTTGATTGGAATTCCCTTGAG
GACAATCTGCTGTTGACTGGGTCTGCAGATAATTCGGTGAGGATGTTTGACCGTCGAAAGTTGTCGGGAAAGGGTCAAGGAA
CCCCAATTCACAAATTTCAGGGCCACTCAGCGGCAGTGCTGTGTGTCCAGTGGTGTCCTGAGCGGAAGTCAGTGTTTGGAAGC
TCTGCTGAAGATGGCTATGTGAATATATGGGACTATGAAAAGGTGTCTCAGAAAAAAGAGGAGACAAAGCCATCAAAGCGT
GCAAACTGGCCTCCTCCAGGGCTTTTCTTTCAACATGCAGGACATAGAGACAAGGTGGTGGATTTCCAGTGGAACCTTCTGGA
TCCGTGGACAGTTGTGAGTGTATCTGCTGATTGTGAGAAGACAGGAGGAGGGGGAACATTGCAGATTTGGAGAATGAACGAT
CTGATTTACAGACCAGAAGAGGAAGTGTTGGCAGAATTGGAGCAGTTCAAAAGCCACATACTATCTTGTGCAGATGAGAAAT
AAAGATTTGGTCTGTATGCTGAAGTGCTACTTATACCAGCTGAGTTTTGATCATTTTGGGAGCAGCTCTTTTCATGCTGTGTCC
AGGCCAAGAGTTGTTCCCTTCAGAATTCACAAGGATGCTTTGGTTTTATAAAGAGATGACATGATAATGTTGTAAAGGCTTGC
TCTCATTTGGGTCTCACGTCATTAGCATTCTCGGTCTACGGATTTCAATGACTTTGCAGTGGAAGTGTGTGCGTGCATGCATGA
GCCCAGTTCAGAGGCCTGACTGTGTGCTGTTAACGTATTTGTGTGGTCACACGTGGGTGCATTGACCTGTGCATGAGTGTGAG
TTGGTGTACGCATATATGCTTTCATGCATGTGCACATGAGTGGATGTCTGGGTGTGTGCATTGCGAAGGTAGATAATCTAACA
GTTGTGTATATGATCTAGACATTAAGGTTCCTGCTCATGACAATGATTTAATTAAAAAGATCTCGAGTGTGCATGGTGTACAT
ATGCCGATATGGAAAGATACTCTCACACTGGGGTTGTTTCATGTGGGTTACTGATAGCTTTTTTACGTTTTGGCAAATTTTGGT
ACAAGTTGAGCAACTGGATGATCTTCACAAGTATGTCGATGACAAATCACTGTGAAGCCAGATTACGTGGTGGTAATTTTCTC
TTGATATGCTGTGAAGTGGTTTTCTATCAGTGACTTGGGCACCCTTTGCAAGGCTACTATCATGGAGGGCTCTTGCCTGATAGA
TCTGTGAGGCTCTTGCCCAAATGCATTTGTTTCTACGATTGGCCAATATTTGGAAAGGGTGATACCTCCTCGTGGTATTTGGGT
GTCTGTAATGGCTGTTCGGGAAGGACTCACTCTCCAAAAATAGAAGTGATTTCCAACACTTGTTTCATAGAAGATCCTAACTG
CTGTTAAATGCAATGTTTGTATTTTTTGAAAAGTGAAGCTATGCGATTGATGACACTGGCTTTCTTGAATTAGGAAGCCCATG
ATTCCTCATACTCGGTTTCAACTTGGGAGCTATCAGTGCTATCTAACTGATGTAAGAATTTTAGGCTAGAAGCATATCATTGCA
AGAGTTACATAAAAATGCTAAGGTATTTTCAAGAGAAAAA

**ENDO-BETA-MANNASE 6 (*MAN6*):**

TCTCTCTCTCTCTCTGTGTCTGTGGGGAGAGGCCATGGCTTCTTCTACTGGCAGCTTCCCTACCATGTGCATTGCGTTCT
TGGCTATATTCGCTCTCTTTTCTTGGTCTTGCAAGGCTTCTTACTACCCAGGCTCCAACTTCAATAGCTTTGTGCAAACAAGAG
CATCCCAGTTTGTTGTGAATGGTCACCCTTTCTATGTGAATGGCTTCAATGCCTACTACCTCGCCTACGTTGCTGTCGATTCAC
GCGACGAGGCTTCCGGCATCCTGCAGCAGGCTGCCGGCATGGGCCTCACCTTGTGCCGGACATGGGCATTTAATGATGGAGC
CTACCGTGCCTTGCAGGTCTCTCCTGGAGTCTACGATGAATCCTCTTTCAAGGCTTTAGATTTTGTGTTAAGTGAAGCCAAGAA
GAATGGGATGAGGGTGATATTGAGTTTGGTGGACAACTACCCAAACATGGGGGGAAGAGCTCAATATGTGCAATGGGCAAG
AAATGCCGGGGTCTCCCTTTATGACGACGACGACTTCTACACTAACCCCACCATCAAGGCCTTCTACAAGAACTATGTCAAGT
ATGTGATCACAAGAGTGAACACTTTGACTGGAGTGGCATACAAGGATGATCCTACAATTTTTGCATGGGAGCTCATGAATGA
GCCCAGGTGCCCGAGCGATCCATCTGGAGACACATTATATGGTTGGATTCGGGAGATGTCGTCTTATGTGAAATCCATTGATT
GCAATCATCTAGTCGAGGCGGGGCTTGAAGGCTTCTATGGGACTTGGAGTGAGGATAGGCAGCATCTAAATCCCGCAGGGAT
GTCCCTCTCAACAGGGACTGACTACATTAGGTTCAGTCAGATCTCTTCCATTGATTTCACGACGGTGCACTCCTATCCGGACCA
ATGGTGCGTAAAGTCCGTCTCTTGACTCAAATTCTGCATGTTGGGAGTATAAAAGCGCCGAACTCGCATGGGCAGTCGCCTGA
GTGCGTATTTGGTTGTACTCATTGTCTGATTCAACTGAAGATATGTGTATCCTGTACTAATGTTTGTGATGTGACTAAATGTGT
CCCTAATTGTGTGGTTGGTATAAACAGGCTACCCGGGCGAAGCTTCTCTGAACAATTGGCATTCTTCACAGCATGGGTAAGTG
CCCATGTGAGTGATGGGTTCGCCGTGAAGAAACCTCTTTTATTTGCAGAGTTTGGACTATCCAAGAACAGCCAGGGCAGCTAC
TCGGACGAGAACCGTGTCACTCTCTTCTCCGCCCTCTATGCCTCCATCTACAGCTCTGCCATGACGGGTGGACCGGCAGCAGG
GGCGCTTGTTTGGCAGATTTGCACTCAAGGTGTAGAGAACACCATGGCCACAGACGGTTACGCTGTTCTTCTCTCTCCTGATTC
CCCTATCACCTCCCTAATCTCTCTCCAATCTCAAAAGCTCACTGCTCTAAACCAATGAGCCACACACAAACAAGCACACATAG
AGACGCACTCTTTTTGCGTGCGCGCACACGCGGAGCATTGACAATCACTTAGACACAGTAAACACTCCCACACACACACATG
GAATCAAAGACACACCCATAAACACACACACCATCAACATTCACATAGATGCACATATCAAATTCAAAAGTTATCCATGATC
CATATGCAAATGTATACACAGAGGTGATGGCTGCAGTGAATGTGGAGTACTATAGAGAAGTACACGGCCTTACACACATACA
CACACATGTCACACACACACACAAGAGAGTCCTTTTGATGCATCCATCCAAAGCAAACATCACTATCAACAAGGCCCTAGAT
TGTGTCGTTGATAGCTCCGACCATCTAGATTACTTTCTATACACATTTCCATTAGGCTTTTAAGCTCTCCTTTTTCCTAGCACCA
TTGTACACACACAATATAATAGTGGCTACTTCAATTTTTTGAAGAATAACTTAAGCATATAACCTTTTTTTGTAACTTTTTCCTC
ACGTGAAATTTAATTATGTACATGCTTCCTTTACACCAAAGAAGGCAAGTTCCACAAATTAAGTGTGTAAACAAG

**JUMONJI DOMAIN-CONTAINING PROTEIN 22 (*JMJ22*):**

TTGCTCGACATTAAATAGTACTTAAAGCCTTTTCTTAAAAATCAATAAAGAGCTGTTGAGGAAGATTTTGGTGAAAAG
GAACCTCCGTATGAAAATCTTAGCAGTTGCAAATATTGGCAAGCTCAGCATACTTGCGTTTGAGAGTGTCATCTAGACCAAAA
TGAACTGATAGCATTCTTTTCAATTACTGTTAGAATTGATTGAGAAAAATGAGGCACTACAACTCTTTCCCAAGAGAAACAGA
AAAGGAGGCAAGCCGTATGGCAAACGCAGCATACATGCATTTGAGAGTGTCCTCTTGACAAAACGGAACTTCCCCAGAAATA
TGGAAAAGAGAGTCAGGGCATACTTGCAGGGCAAGCCCATGATTATAGCTTGAACCCATCAACAAGTACAATGGTCATTTGA
TTCTATAAAAGTCTGGAGTGAGATGGACATTTGAATTGGTGCCTGCAACACAAACACACACCGACTTGATGCCTTCAATTCCT
TAGGGTTCCGATTGAAGTGGATATGAGGAACCTCCTTGAAAGATGTCATCTTCCATTTTGCTGTAAGAAAGCAAAGGCTACAT
TACTTAGCGTGCACATACTCCACTGTTATCAGCAGAGTTAATGGAGCTGATGAGGCTGAAATTGAATGATTTTTGCCTTCGGT
GATGTTCCTTTTGAGTTTGAAAACCTTCAGGAAATACCTGCCAGCACCAGCTATATCAGCAGGAAGGGTAGCTAAAATGCCA
AAAAAATGGGTATGATAAGACCCTGCACACCAGCAGAGATACAATAATGTGTGAAGGCATCAGAAAACGCATGAATTTCTCA
GCTGATTTTTCAGGAAAGCCTGGAAGTAGTAAGGAAACTCTCAAAGCACTGTCATTATAACCTGTTAATCAGAAGCCGAATTT
AAACCCACCAACTTTAACGTCTGTTACCATCTCCCAAAACGAGACAGAACTTCTTTGCTTAGCTTTCTCTGTAGCTTTTCTTTC
CAGTTCTTCAATGGAACCAGGAAATAAATTGTCATACATGCTCTTGAAATTATCATACAAGTTGACTCGGTCTCTTGTACCCG

ATACCAGCTGAGCCGAATTTGGCCGTTTCAGAAACTCCAAGACATTGAGAAGATTGCTCCTGCTCACATAATTTTGTGTAATC
GCAACAGAATCCTCCAAATTGATAACCAAATGCCACCATCCATTTGGGACAAACATCACTTCACCTGCTCTGCATATGCACTC
TACAGGTTTTTCTTCCACTGTTTTGTCTGCTCATAGAAGTTCATAAACCATTCCATTATTGACACAGGAGCAGCCACCTCGGC
ACCATCTGGACTAGGATGCACACCTGGAGGATTAACATCCGGAGGGAAAAGTATCCACTTTTTGGCACCCCTTACCACAGCAT
TCCATGCAGATGTGCTGTTGGGATCAATGTGAAAAGAAGAACCTGATCTAGCAGGCCCAATTATAAGCCAACGAAAATTAGG
TCTGTCCTTCCCCAAAATAGAGAATAAATCCTCCCTGAAGTAACTTGGGACCACATAATCTTCATTTAACACTGGCATTTTCTC
TGCAAACTGAGGATCAAAAAGGTACAATGGGCGCTCTTCACAGATAGATTCTGAATATGAAAAATAATCATCCATTGTCATTT
CCACGGGACCTACGGCAAGCTTGGAATCTCCTGCTGCTTTCACCAGATACTCCTTGTTCCACTTCGTCAATGCTGGCCAGTTAG
ACAATGCATCTGTTATTATCACTGGTTTATTTGGCTCTTCATAACTGTCGACAAACTCCTGCACAGAAAGGTTGGCCCGCCTAT
CAATGTTGTCCACCAAAAGCCATTCTTCTTTCATTTTCAAGCTGCTGCAGAGCCAGCTCTGAAAGAGGTAGTCTGAGAAAAAA
CCTGTCACTTGTAGACGAGGCTCCGGATAAAGGGAAGAGGATGACGTGGCGTCGGCGTAGGTGCATCTCCAGTCAACTTTAA
ACCTAAAATCGCCTCCGAACTTATCCAACACAAGATTTCTCCATAGGAAATCCTGGTGAGAAAAGACATACAAAGCTTTACTC
ACAAGAGAAAGGTTAACCAGATCGATGGCTTGCAGAAACCCTAGAATTTCACACAGGGTTTCATCTCCGAGGACATGGAGGG
AGCCGAGGCCTGGGTCTCTGATGTTGTGCGCGAAAGCGGAGAAGAGAAGGTTTCCAAGAGGTTGGACCCCATAGCTGCAAAG
TGCTCTTTCCAGCTGCAGGCCTTCACGCTCATCGTCATGGGTCGCCATTGATGAAGTTTTGCTCCTCTTTAGACTGCAGTCCTC
CATATTGGGTCTCTCTTTATTCTTCTTCTTCTTCTTAATCTTCTTCTTCTTCTGATGCCTCTCGCCTGTGCTGTCATCTCCATTGG
AGCCCCATTTCGCCATGTCTTTGACCCTTCTCTTAGAGAGAGAGAGAGAGA

**WUSCHEL RELATED HOMEOBOX 13 (*WOX13*):**

GATTCACAATCCTGTAGTACATGCTCACCAAAGATAATTATACTACTGCCAACACATCATTAACCGCATTAACTACTT
ATCACATCAACCCTTGAGCCATGTTACAGGCAGTAATCACAGCAACATCTTTCATTTCCCCTCTTACTTGTATTCTGGCAAGAT
TTGGGCTGTGGCCTTCCTTTCTTCCACTTTAGTGCCCCGCTGACGATAGAAATTCAACTGATTTGGAAGTCCCATTAGAGACAG
ACTGACTTTCCTTCAAGCCAGCCGTAGGCGGTCGTAAGCATTGCCACTCGGAATTATCCTTCGACTGTTGCACACCATGGACA
TGCTCTGTAATCTAAAGAGTGGCAAACTTCTTTATTAACAGTCTGTCTGTTTCTCTTCATCTAATCTGTTTTCCTTGAGTAAAGT
ACACACCCATGTGTGATAGAATCGCTTTCAGATAATGATAACTATACAATGAATGTTTAAATTTAACTATAAATCCTTCCAGT
TAGAAGCTGGTTACAACACCACCTGAAGGGCATCACTGTAAAACTTGTCAAACTACAAAAAATTAGCACTTCCAGATCTAGC
AGGTTGTCTCAACCAAGGATTTGTTCGAACTTCCAAGATCACCACCAGATTATAGAAGCTTCCTGGATATTTAGAATTGACTG
CCTTGATCCGGAAAGCCTTCCGGACTACTATCTGCAGGCTTGGAAAGATTGAAAATCTTAAAACCAGCATACAAGTGCCATTG
AAACTAACTTTTGTACATGGAAAAATGTCGGATCTTAATTGAAGTTATTGCTTAGCATTTGGTTCCTGAAAAGCCTCATCGCAT
GACAATGTTAATCTCTGGAGCTTCAATGGTGGCGACTCTCTGCCCCAGTAATGATCATCTGCCGTCCTTGAAATTCTTGCAACA
CTGAAAATCCAGAAAAATCTGCTTTGCTTTCAGGCTCAGTCATGGGAATGTCCCACAGTTTACTGTCGACAAAACGGCTGGCT
TGCATTTCACCATGCCCATTGAATGCCGATTTGTTGAAAGAAACAGCAGGGCTACTAGCGTGTGCATGCCTTTGAGATAATTC
AGAAAATCCCTGGTAATGCTCAATGTTATCAGCAGCCCCGCTAGTCCCTGCAGCATCCAAACCGTCATTTGTGAAGTCTCCCT
CGGTCTTACCCTTCTTTTCTTCTGGAAAATCAACATCCGTGTCAAGCTCAGAATCTCCATTGTTAGGCCCACCAAGCTGCTGTT
TTCTCTTGGTTCGGGCTCTTCGGTTCTGAAACCAATTGTATACATTGGTTTCCGAAATTTGGCCATGCTGACTAAGCTCTATAG
TTATCTCCTTTATTCTTTGCTTATTGGGTGTGCCACTCTCCTCCTCAAAGAGCCTCTCGAGAATGTGAAGCTGAGTTTGACTCG
GTGTCCACCGCTGCCGGGATGTCAGCTTGTGTCCTATGGAATGCAGAGAAGAGTCGAAGGGGATAGATGGGCCTAGCCATAA
CCCTGGTGTGGAGCCTTGCTGCGCCATGATCGCCTTGTGCATTTCCACGAGCTGTTGGCAAATGGTTGCATATACAGAGATCT
GTCTTCTTAGGGTTTCCAGCTGTTCATCAGTCATAACTTGGCCATGCGGGGTTATGGGGCAAACTGGGTGATGTTGGGTTGTG
AGCGCATGACTATCAGCGGATGATGAGTGCTGTTGGTTTGCAAAAGCATGGAATCGCATGTCATAGTGCTGCTGATTGGCAA
ATGCTTGGATGTGCATATCAGGAGACGCATGCACCCCATGAAAGGGGGTCGCGTATACACCGGTATTCGCATGTGGCAAGTT
AGCCTTGGCAGACAAGTCCTCTCCTGCTCTATTTTCGAGCTCCATGTGATTCAACCCTCTTCAGTCCCCCTCTGATTCTAGGGT
TCGCCTGTCTTCATAGGAATGGAGACCATTAAGCAGCTCTCAGAGCAAGAGGCGTGTTTTGGAGGGGGACTCAAGTGGTGGG
TGGTTAGCGAGGAAGTGACATTTGCGGCAGCGCATGTGGGAGAGATTTGGACAATGCGCAGTGAGTTCTTTTGTGTGTTTGCC
GTGTGGGTGTGGAGAGCTTA

**FASCIATED STEM 4 (*FAS4*):**

AATGGAAAGCTTCATAATAGAGAGAGAGAGAGAGAGTGTGATTTCCAGAATCTTTAAACACCTTGCTTTTTCTTCCTC
TTTCCATAAAGGCTCTTTCCATCTTGTTGAGCTTCAAGTTTCAACTTGGTCCAAAGCCTTTCCAACTGTGAATGTGAACTCTTTT
GTACCCATAACAGAATTTCCTGGTACAGAAAGTCACTATGAGTATCCCACCTACAGTTGAGCTTGAAGCGGCTATCCACTGAT
GCAGCTTCTAAAGCATGCAGCAAATCACCGGAACGTTTGTGGGCATTTCCAGAAGGAGAAACTAAAATTAATGGGTCTGCGG
CTAGGTGGGGATGCAAGGCCTTCAAGGCAGGAAGTACTTTACCTTGAAGAAAGGCACATGCAAAAACAGCAATACGATGTTT
TTTGCTTTTCAAGGGAAACTTGTGTAATGGGAGTTGCCACAGATGAGGACCAAAGCATGGGGACACCCAACAGGAAACATCA
TCAGATAACTGATCATACCATGGTGGGGGATCTGAAAGAGGCTTTGAGAAAGTACATAAAGCATTTGCATGCATTACCAGCC
AATCTGCATTGACACTCGTCAGAACTTTCATATAAGGCCTAGATGTTCTCACAAGCTCATTGAACACCACAAAATTCGGTGCA
TTCTTCCTCAAGGAGGATGACGGATGCACATACACTTTCTCTTCTGTATTGCAAGTTTGATACTGAATTGTCTTGTGTTTTTTCT
GTTCACCTCCCTCTTTGCTTTTCAATATTTCCTGCAAACTCACCTTATGCGCGACTCTATCTGCCCAACCTGCACAAATTGCTTG
CCTCAACACATTCTCATGCTTAACACTCAATTCATGGTTCACTCCAGCCTTCCACATGGTTTCTACATAGGCTACGGAAAGTTC
ATCGCAAAAAGAGATGCAATTAGTTCCTCTTTCTTTACCGAACACAAGATAAATCTGTACAAGTTGTTTTCGCAACTTCGACA
TCTCATATAAAATCCTTGCGTGCAAGTGGTTGTTCCTGCAAAATTCCTCTGGATCATTGGAGACTTCATATGCACGCAAAGCC
CTAACCACACCAAGAGCATCACTAAGGGGATGGCTATAGCTTGACCTGACCTTTTTGACAGATGTTCTTAACTCTGTTGTTTTC
AATGCTGCATCTTTTCTCTCTGACTTACCATCAGCATACATTCTCTGTGACTTCTTTGATTTACTAACTGATTTTTCTATAGTTTC
TTCATTTGGATCCCCTGGACGAGCTAAGTCCGTCACAAAAGGACTATCCAAGCTTAGAGCTGCTACTGTTGCCCCAGCATAAG
CAAGCAAGATGCTTGTATCTTTAATATCATCTGAGCATCCAATCTGCATTGCAGTAAGCAACATTCGAGAATGACGAGGACTG
ATGGGATACAATGAGATCGCTTTCCCCAGAGGAGTCAAAAGGCCAGTGCTTGAGTCAAGTGCTGATAATGCATGCAGACAAT
GCTCAGCCTCTGCAAGATCTGTTTTATCAGGTTCAGACAAGAAAGGGAAATGGCTCACCTTGACAATATTCATGCTCTTTAAC
AAAAGAACAAGGCTCTCAATGGGAGCCCGATATATCTCTGGCTGAGAGAAGTCCCGGAACGTGTCGTTGAATATAGCAGATG
AATATAGGCGATAACAATGCCCAGGGCCTGTCCGACCCGACCTACCAGCACGCTGGGCTGCAGATGCTTTGCTTATCCATCCG
ACCTCATACCTTGTAATACCGCTTGCACGTTCAAAAATTTTATCCTTGACACGGCCACAATCCACCACATATCTGATGCCCGG
AATAGTGATTGAAGTTTCAGCGACATTTGTGGCTACAACCACTAGTCGAGATCCCTCAGGAACAGCACCAAAAATCTTCAGCT
GTGAGGCTGCTGGTAGAAGTGCATAGAGGGGCAGAACATACATCGGGCCAGCTACTTTTGCTGGTACCTGCTTTTCGACCACA
AATGGCTTCTCCAACTCTTCCCTCCTTCCCGTGCCAGCAAAAGAGTCAAATGCATCTTTGATAGCTTCCAAGCAGATAGAATTT
TCTGACCTCCACTCTGAATCACGCTTTGGCCTCCGTTCCTCCACTTCAATACCTTCTGTTTCTGAGAACTCCAGATCACTTTCTG
AACTAGAATCAAGGGGTGATTCATTGATTTCCCAATTTTCCACCATCTCTGCTTCATCATAGAATTCATCCTCTGCGTAACTGC
CATCAGCTGCTTCTGAGATATTCTTAAGATCATCATTATCTCCATTATTAGGTTGCTTTTCGTGCTTCTTTACACCTTCCTGTTTT
GGAGGAAAAGCTTTCTGAAGCCTCTTGCACAAATAATGAACTTCTGCTTGCCCTGTAAGAAACACAAGAACACCTCCAGGAG

GAAGAAATTTATGTATAGAGCAAACTTTTTTGTATGCTTTGCCCACATAGTCCACAAGCTCTGTTTTAGCAGAGAAGTGGACT
GTAACGGGAAACTGCCGTGCAGGAACTTCAACCACAGGGGGTGCACAAGCAAACAACTTCCTGTTTGCCACAAACTCATCTA
CAAGCAAAGTAGCACTCATTATAACAAGCTTCAAGGGGTATATGGAAGCTTTGCCTGCCCTACATTGCTCTTCATATAAGCTC
TGGCGAAGTGGCAGGACTCGAGACAGCATACCTATAAGAATATCTGTATTTAAACTCCTTTCATGTGCTTCATCTAGCACAAT
AACTGAATACTTTCTCAGCAAAAAATCTGACTGTACTTCTCTGAGAAGAATGCCATCTGTCATAAATTTGATGTGGGTATTTTT
CCCAGCTTTTCTATCGTGTCGCACTTGAAACCCCACTTCTTCACCTAATCTAAGGTTCATCTCATATGACACTCTCTTTGCTGTT
GCGAGAACCGCAACCCGGCGGGGCTGTGTCACACCAATCATTCCTGCTCTCTCTAGACAGGCACTTGAAGAGAACCCAGCCT
CGTATAAAAACTGAGGAACTTGAGTAGTTTTACCACAGCCAGTCTCTCCACATATAATTGTGATTGAATTTTCAGCAATAGCC
TCCATAATCTCTTGCTCCATCATTATAATGGGAAGCTTTTCTCTCGCACTTTTTACCTCCTCTGGTCTCCAAACATGTACAACTA
CTTTAGAGCCATTATCAGGAATGCCTCCACAGGCCTTAAGAGCATCACTATCCTTGGTGATAGTTCCCCCTTGGACCGTCTCA
ACTTTAAACACATCATCTCGCATGATGTCCGTATTATTAAATTCTGCTAGCTTTTTGCCCTTTTTCTTTTTCGTTTGTTTGGATGC
AACTCTTCTTTCCCTTGAATTATCATCATTAAAATTTTCATGCTCTTTAAAATATTTTCTGGCTTTCCTTGATCTTTCTTCAATTG
AAAAATTATTATCTATGCCAATACTGAGATCCGACAAAACAAAACCAAGTGTTTCTGTGCCACCTTCAGTTTGATCAATAATT
GCTGGCTGACTTGTGGTGTGCTGCACTTTCAAAGCATGTTTTGTATTACCCCATAACTCTCCATCGCTTTCACAACTTTTCATG
GGAAAACTATTGCCTGCTTGAAATTCTGTTTGTTTCT

**SWITCH1 (*SWI1*):**

GAGTCAAGTGATACAATGCTACAATATATCACGGGCTTCATTCATGAAATGCAAATAAAAGCAACCGTCTTCAACAAT
GCCAAACCAGAAGCAAACATTCTGAACAAGTTAAGACCCAGTAATTCTTATCGTAAAAAACATAAAAGAAAAAGAGGACCG
CCACTGCTTGTAAGTTCGCCCATCTCCGAAAAGGAAGCTTTGCAAGGTAGTGTATCTTGACTGAGTACTTGGCTTTTCATATTA
GAATGTGTACAATAGATTTCAAATTTTCTGCGCATGTGAACATAAAATCCGCGTTTGTGAATGCTTTGCAGACTCTTTCATATG
CAAAAGCATCTATGAAATTGAGCAGAAAAGCCTGCCACCAGATGCATCTCAGCAGCTGAAAAATCTTCGTGCTGCCCTGGTA
AAAGTTGGGTGGTGTACATCTATCCTCCTGAAAAAATGATGTTTTAACTGTTTTACAACAAGGAGGATGCACGTTTTTGCAAA
CGTTTATGCTTCACTGCAAAAAGTTTTTATTTGCGCGGTTGCCTAGCTTTGAAGATTCTAATAAGCTGGATTCAACTGATTAGT
CCAAACAGCTCCTCATAGCCGTTGTTGAAATATCATTTCCGATCGTTGTGGGGCATTGAGTGCAGGTCAGTGAAACTACAAAA
GAATATGTCACCATTGCGTATCCAAGCATGGCATCTTTGGCAGAACATTTTGAGGAATCGAAGGTGTTAGATGGAGCTTCCCA
AGAAAGAATAATTACAAAATGCCTTTTTAATCAGCTTTCCACATCCACTTGCATGCCAAGGTTTGATGAAGAGTTTTCAATGA
CACACGAAACTGCTCAAAGAGTGCTCAAGGGCCTTATTTCTTCCACACAGTGTCTCCGACAGGGCCATCTTGAATCATTCTGG
TTAATACCAGGCCTAGACACCACCTGCAGAACGACAAACACCCAAGCAAATAAGGACGAGAACTGTTGCAATGCCAAGGAA
ATTGCTTCATTATCCCCAAACCACGATACCATAAAGAAACATCGACCCTCTGAAAAGAACTCCCCAAAAGTGCTCAAAGTCTA
CACACACAAGAAATTTTGGAGGGGAAAAACAAAAGTCAAAAAAGAAAAACTATCTGAGAGCGAATCTTCTGGCAAACAAAA
TGAGCATAATAACAGCAGCACCCAAGAAAGGGAGGTGCCTACGTCATTAGATGATGCTGATCGTGAGGAACCCTTGCTGCAT
ACAGTTGATAGCAACCCTGGGTGTGTTTTCACAGAAGATCATGAGAAAAAATCACAGAACCATGATGCCTCAGAAGAAAACC
AGGGCATTGTCAGATGGGGGGTCCGGAAAAAGATTACCTCTAGGTGCCGCAGTAACACCTTTAGTGATGATTCCAAAAGTAA
AGGACTCTATTGTAAAGAGGATATGCACAAACAGAATCAGCACAACGTAGCAAAAACAGGTCACGGGTCAAATGCATATAA
GCATGTTGCGGTGACACGAATACTTACAGGCATTGAAAAAGGAAAGATGCTCAAAGTTCCCAAATTAATGGAAGGAAGATGG
TCTACTGAAAGGTTTAATCTCGATCCCTTGGCAAATTGAGTTACCCCTCATGAGCAGAGAAAAATTAACTCATCTGGTGACAG
CGAAATTGAACTCCTGAATGTTGTTGCAGATATAAAGCAGCTCAACTCAAGCTTTTTGAAATTATGAAAGACAAGGGTGCAGT
TCCTGGGAAACCCCTACTAAGGCCTGCACTGCGTGTAGAAGCCAGAAAACATATTGGAGATACAGGACTTCTTGATCATCTTC
TAAAGCATATGACAGACACTGTGATCAATGATGGTGAGCGCTTTCGTCGTCGGCACAACTCTGAAGGTGCAATGGAATACTG
GCTGGAGGATGCACGTCTACAGGAGTTGAGGAAACAAGCAGGTGTTGACCAGTACTGGATACCCCCTCCAGGATGGAAAATT
GGAGACATGATAGCTGACTTGTCTAAGCCACATGTAAACTCTTAAAAAACTGTCAGCTTTTTTTGAAGAGAAATTTTTGTGAT
CCTTCTTGATAATGCCTCAATTTGTGAATTTCCAGTCAACTACAATATGTGTGTCTGCATGTCGGTGCCAGCCTTTTAGAATCA
AGAAGTACATCTGTTTTCAATCGCGAGTCTGACTTCAAATCCTTTTTCTACAAGGCAGCAGCTCGACCGGGGAGCCTTGGAAA
ACCAGGAGAGGTTATATTCACGAAGGCTAAGCAGGAGATTTCAGAGAAGAAGAGCCCTGTTTCTATTGTCTTAAGGAGGAGC
CCTCGTTCAAAGGAAGCTGAGAATAAGGAAAATGTGTCTCCAATGGGGCAGCAGGGCGAGCTCATCAAGGCCTTTCAAGGTT
GGAAGAAAACACAGGGCTTCTCTCAAAGAGAAGTTCTCTCTGCAAAATCCGTGAATATGGCGAGCTCACTGTAGTGCAATTC
TCGGACAAAATAATCTTCGGAAGATTAAAAGAATTTAAGATTGTCGGCTTGTACTAGTACTTGTGTTGCATTTTGAGTCATCA
CTTTGATTTAGTAGAGAAGAAGAAGAAGTTC

**SLOW WALKER 1 (*SWA1*):**

TGTTGATAGTGTAGACGAATTTTCTAGCGAGTGTCGGGGCTCGAGGTATAGTGATAGTAGTGTTGCAAAAGACTATCA
TTGAATTTAAAGCAACATTTGGTGTCATCGTTGGATGATTTGCCTCAATGTGGCCTTGCTCAAGAATCATAACAAGAATCCTTC
ATCTAGGTAGTCCACAACACAATTAAGTGGGCTAAATCCCTCTCTAAGAAAACAGTGATATCGTCCAAACATCATGTGACATT
AACGAGTTGATATTGGATCATCCACGAGAATATGAGGAGCAATGGCTTCGCAATACGCCAACAATCCACTTGAAAATAAGTT
TGGTGCCCAGAACACGATAGCATTCATCACTGAGGGTTAACAAGAGTGTCAAAAAGCCTTCGTACAAGGGTACTAGAATTGA
AATGCATGACGTCTCTCTCAAAGGGCCAGTGTTGCATATCCCTTAGTCAGCTACCATGCTAGTCCACTATCACCTTGACATGC
AGCGAGTTTTGCACCTCCACCTACTTGGTCAGTAAAACCAGAGATCTCATCGATATAAATTGCACAATGTGTAGAAGATGAGG
GGCATCCTTGTCTTAGCCCAATGGTGTTGATAATCTCATATGATAACCATTGAGCACTGCATTCTCCCCACTCCTAGCTCATAC
AGACTAAAGCAAAGTAGATAGTTCCATGGTGCTTCAAATTACTGCTAATGTTTATTGAACAGATGCCTGAATCAAAGGTTGAA
CCAAGCCTTGTAGCACTTGTAAAGACTCCTGCAGACGAACTTCAGTTTCAACGTTCTCCCTCAGTACTGCCACCTGATGTAGA
ATTGATGGAGAAAGAACAAAGTTGCCAGCACATTTGTGAAGCACCTTGTGAGCAAAAGGAATTAATATCCTTGAAAATTTGG
GCAATGTCACATATCTCCTTAGAAAATCTAATAGCAATTCCAGCGACGTGCTATCCAGATTAGACACAGCGGATATCAAACCC
CGGCGAAGAATCAATTCTTCCATAACAGCCAAAACAACTGTTGGGTTCGACATCTTTAGGGATGAAACAAGAGCTTCTTTATA
CTGGAATTTGCGGAGATATATATCATGTTCAGCAAGCCGAGGCCTTCTTTGTTGAGCTACAAAGTAATCTCCTTCCGTAGCCTT
CTCTGCACGACCTCGCAAAAAATAGCGGTAATTGCTGGGTCGCAAGACAGTTTCTAATTTTGGTTGCTCTTGGAATATGCCGG
CAGCATTTTCTGTACTCCCTTCTTCATCTAAAGCCACCTTTTTCTTCTGGCGAATAAATAATTTCCCATCTGACATCCCTACAAC
CATAGTTGCCATTGATGGGGAAAGGTCCATTGACATGATCGGAGAATCGTACTTTGATGCATGAACCACCTTGAATTCATTGA
TATCAAAAACCTTGACATGGCCATCCAGGGACCCAGTCAAAAGCCGCCGGGAAGTAGCAATGTCACTATGTCGGGCTTTGAT
GGGCGGGGTAAGGCATAGGCATGTTACTGTTTTTTGATGGCTGGCAACTGCATGCAAAAGCTTTCCGCCTCCCAAAACATCCC
AAATCTTCACAACATTACCACCAGCAGTTGCTATGAGCCCTCCAGATGGAAAAAACAAAACGTCTTCTAGAGGCTTCCCATGC
TGATGTGCCATGACCCAAAACATATACCTGCAGCTGTAGTACTGATTTTGCCGAGCGAAGATCCCACAAATTTACTGTATGAT
CATAGGATCCTGTAGCCCAAAGGTCTGCACTAGAAGGGTTCGCACATCCACTTCGTACATAGTCGGAATGACCTTCTAACTTC
AATATTTCAGTCTGAGTTGCAACGTCCCACCACCGAACTGTATTGTCATCGCTTCCTGAAAGTACATGTAGCTTATCTAAAGG
GGAGTACGTAACCCAATGTACTGCCCTTGAATGTCCCTTTAGTTGGCGAAGCACAAGGCGACTATTGATATCAAAAACCTGGA
TGATCCCCATTTCACCCCCAGCAACCACTAGTTGCCCATCAGATCTAAAAACGCCAGAATATGCCACATCAGAGAAGCGTGA

GATCGTCTTTTTAACTTTACATGTTTGCCCATCATAAATCGAAACCCTAGTGGAAGAAGTAACAGCAAAGTCGTAAGGGGGTT
CGGGGCAGAAGTGAACGCAGGAGACGGCAGCGATCTGTTGGAGAATTTGCTTGGAGGAGAAGGACTTCCAATACTTGGACTC
CAAAGCAGAGGGTTTCCTGGAGGCCGCAAACCTCTTCGCCCTCACTGGCTTGTAGCTGCTGCTTTGCACTTCCATTCTCACGCG
CGCACGCACACAATAACCCCCTCCCGCTGTGTGCAGCGTTTAGAGCTAGCTCTTCTAAAACCCTCTTCCAAATTCGAAA

**BRI1-ASSOCIATED RECEPTOR KINASE (*BAK1*):**

ACACACACACACACACACACACTTACGCCCACAGCCACAAGGAGAATTGTTCAAGAAGAGGAAGCGCGTTCTCCTTC
ATCATGTTTCTATAAGCAGATCCTCATTCAGAGAAACCTCGCAGGTCTCTCTCTCACACACACACACTCACACACACAAACAC
ACACACGCACGCGCGTGCAGAGAGAGAGAGAGAGGGTACCTTTCTTTCTCTCTTTAGTCCTGTTAGCTCCACCCGCAGAGAGA
GAGAGAGAGAAAGTAACTGTGTAGCCTTATCAGCTCCAAGCTCCACGGAGAGAGAGAGAGAAAGAGTGCCCATTTTCTTGAA
AAGCCCCTTTATCCCATGGCTCCCTCTTCTGCTTGAATTCTTCCCCGCTCTCCTGCAAGCCCATGCCTGCCCCACAGTCCGCCCT
TGTTGCTGCCCATCTACCGTAGCATTTTCAAAAGGCTTCTCGACATGCGGGCTTCCTTCTTCCTGCTGCTGCTTCTCCAGCTCCA
GCTTCTCCCTCGCATTTACTGCAATTCTGAAGGTGATGCCCTCCATAATTTGCGGCTTGCTCTTACTGATCCAAGTAATGTGCT
TCAGAGCTGGGACCCTACTCTTGTGAACCCATGCACTTGGTTTCATGTCACGTGCAATCCAAGCAATAATGTTATCAGGGTTG
ATCTGGGGAATGCTCAGTTGTCTGGCCTGTTGGTTCCTGATCTTGGTGAACTTCAATCACTTCAGTATTTGGAGCTCTACAGCA
ACAATATTTCTGGAGACATTCCCAAAGAGCTTGGAAGTTTGGGTCAGCTAGTAAGTCTTGATTTATATCAGAATAAATTTACG
GGATCAATTCCTGACACATTGGGACAGCTGAGTCATCTTCGTTTCCTACGTCTAAACAACAATACTTTAGATGGCAGCATTCC
GTATTCCTTGACTAGTGTCAACGCACTTCAAGTACTTGATCTGTCAAACAATAATTTGTCAGGGAAAGTTCCTACCAATGGTTC
CTTTTCGCTTTTCACTCCCATCAGTTTTAATGGTAACAGCCAGCTTTGTGGCTCTGTTGTTAGTAAACCTTGTCCAGGAGAGCC
TCCGTTCCCTCCCCCACCGCCTTATCAGGCACCTCCGAGCCCTTCTTCAACTGGAAATAATGGAGGAAGGGCATCAACATCCA
GTACAGGTGCTATAGCAGGTGGAGTTGCTGCAGGTGCTGCATTGATTTTTGCAGTTCCTGCAATTGGGTTTGCATGGTATAGA
AGGCGAAGACCACAAGAGCATTTCTTTGATGTTCCAGCTGAGGAAGATCCGGAGGTGCACTTGGGGCAACTCAAACGTTTCT
CCTTACGTGAGCTTCAAGTGGCTACAGATAACTTCAACAACAAAAACATACTGGGAAGAGGTGGGTTTGGCAAAGTATACAA
AGGACGTCTTGCAGATGGCACACTAGTGGCAGTAAAACGTCTGAAGGAGGAAAGGAGTCCAGGTGGGGAGCTGCAGTTTCA
GACAGAGGTCGAGATGATAAGCATGGCCGTGCATCGCAATCTTCTACGCCTGAGAGGGTTCTGCATGACCCCAACAGAGCGG
CTTCTTGTTTATCCCTACATGTCTAATGGAAGTGTCTCCTCATGCCTGCGAGATAGAAGTCCTGGAGATGCACCGTTGGATTGG
CCCAGACGAAAGAGTATAGCCTTAGGTTCTGCCAGGGGTCTTTCTTATCTTCATGATCATTGTGATCCCAAAATTATTCACCGA
GATGTTAAGGCTGCCAACATTCTGCTGGACGAGGAGTTCGAAGCAGTAGTTGGGGACTTCGGGCTTGCTAAACTTATGGATTA
CAAAGATACACACGTAACGACAGCAGTGCGGGGCACTATTGGGCATATTGCACCGGAGTATCTGTCAACAGGGAAATCTTCC
GAGAAGACTGATGTGTTCGGGTTTGGAATCATGCTGCTGGAGTTAATAACTGGGCAACGAGCTTTTGATCTTGCTCGTCTTGC
CAACGACGATGATGTCATGTTACTGGATTGGGTAAGCTGTTAAGAACATGGGTTCATAATCCTATATTCATTTTTGTAGCAGT
GTTATTTGAAGGCAAATTTAAGAGAAGGTGGCTAAAGTTTGGAACTGTCTCTGATTTTCAGGAAATTTCTGACTATTTTATTGC
ATAAAAAGTACGCCAAAAATTGTGTACTCTTGGTGAAAATTAGGAAGAATATGAAACACATGTTTTTTGGATAAAAATATCA
CTTGGAATATTTTCCAATGGATAGGCTACTTATGTAGTTCAATCCGGGCATGTTCTCCCAGCGTGGCTACAAAATTTTAAAGG
GGTCCTCCCAAAATAAACACTCTTTCGAAGTGAGAGAGATGGTTTGATTTTTGATTTTTGATTTTGTTAGGAGTGTTGGACTGA
TGTGTTTGGATTCTAAATCCTGCAGGTCAAGGGTTTACTTCGAGAGCGCAGGATTGACATGCTTGTTGATCCCGACTTGAAGG
CCAACTATGAAACCCATGAAGTAGAGCAGCTAATCCAAGTTGCGTTGCTTTGCACGCAGAGTTCGCCCATGGACAGACCCAA
GATGTCAGAAGTGGTTCGGATGCTAGAAGGGGATGGGTTAGCAGAACGTTGGGAAGAGTGGCAAAGAGTTGAGGTGGTCCG
TAGCCAAGAGGTTGAGCTTTTACCTCATAGAAGTGATTGGATTGTGGATTCCGTCGACAATTTGCATGCAGTGGAACTTTCTG
GTCCAAGATAGACATAACAACCCTTCATTCTCCCAATGCAACTTCCAAATCGGTGGGGCCTCTATATAAAGCTGAGTCTTGGG
TTCTTGTGTATATTGAGTTCAGAGTAAGGAAATATATTATCCCAGCCTCTCTCGTCGTAGTTGTGAAATATCACGAGACATCTT
CGACAGCATATGCATGCCCTGACAGCTGCAAAAACTTGTAACAACAATTCTTGGCAAGCTTTAGCTTTAGTGAAGGAGAAATT
GTTTCCAAAGAAACACCACTATCAAGTGGGGGTTGAATCAATCTACCTTGTGAAGGCACCTGTTTTTGGCTTCTTGTAATTTGT
CTATTGCAAAACAATCACTGCTCCACTGACTGTGTGCGTCGTTTATCTTTGGATCTTTAAAGAAGAGAGTTGTATTTTCAATTC
GATGTGAGAAAACAAGTTGGCTTTTGTCAATTATGGTGAGTTGGAGTGGTCTGCATTTGCACTGGTTAGACTTATGGGTAAGG
ATGACGATAGGGAAATGACAATGTATAGACACGGAGAAAG

**FERTLIZATION-INDEPENDENT ENDOSPERM (*FIE*):**

TTTTTTGAGAGGTTCAACCGCATCCACATTGAAACATTTAGACGTGCTCATACAAATACAACAAAAGTTTAGGGGCTT
TCTATCATACAAAGTTGCACTTCTTCAATGTTCTTCCATGGAGGACATTGATTTAAGCTTTCTATCCAATTGTAAGAACATGTA
TATCATTGATTTAAGCTTGAGAGCTATAAAAAACCACAAGAGACTACTGGGCACAAAAAGAATTACTTTGTGCCAAAAAAAC
TGTATGGAGTGAGAATTTTGATTCCCCACAGCCATAAATACTGCCCTCTTAGTACTGAAATCTTTTAAAAGCGTAATGGAACT
TTAGATTAGTCCTAGCACTAGCATTGCTTTGTTTGGATTGGTCAGTCTTGTAAGCTGCAAAAATTCTTCTTGTGAAGCTCGAGT
GCTGTGCATACAAAATTTCATCCATCCAAGATCTTTAGAGGAAAACCCAACATGTTCAGTGGTCATCTTTACGGCACAGCATC
CCATCTCCAAAAACTACCATCCTCGCACGAGCACAAAATAGTACTGCACCATATGAAAATACAGAACTTAAGAGCACACATG
AGATTATACACCTTCCTGTTTTTTTGTCTCCTCATGCTTCCTACTGAAACTTCTTTTCGGGGGAGATACTATGAGAGCTACTAAT
TTCTTACACTCACACCCTCTTAATATTAAGATGCCAAGGATTGAGCCAACTCCTCCCCATTGAGAAGCACTCACTCTACCAATT
CAGCGAATGCTCAGCAGCCTAAGGAAAGTACTTTTAAGAAAAGCTCAGTATTTTATTGCTATAACATGCACAAACCTCCCATC
GAAAGACATAGCAGTTTGTCGTATTGGAGATTTGCACTGAGCATAAGATAATTTGGCCAACAATGTGGGGGGACTTGTTTGA
AGCTCCCACACAAAAACCTTCCCTTCTCTGTTCCCAATTGCTAGAGAATTGAAGTGGAAATCACATGAGAACTTGATGAACCA
GATGTCACAATCAGGGACAGGATACTTTTGTAACACGTCAACGGCACCATCATTGCTTCCACCGTGCTCTTTCATCAAAGCTT
CCCATAGAACTATTTCATTGTCCACGCTCTTTGATAGAATGAAATCTCCAAGCCAGCGGGTGCAGTCCACATAGTTCGAATGA
ACATTTGCTGTGAAAATAGGAAATTGCACATACTTTGTTGAGAACTTGGATGACTCATCTTTGCAATCGAGAGATTGTTCAAC
ATAAACCCAAAAATCTTTTAAACTCCAAATCTTGACAGTGTTGTCCATACCACAACTAGCAATCAGGTCCTTGTTAGACGGGT
GGAAGTCCACACTGAGAACTTCATTGCGATGTCCATTTGATCCTGCAAATATGAGAATACACGCACCTGAATTTACATTCCAC
AATCGCACAGACTCATCCTTGCTTGCAGAAGCTACCAAACAAGGCTTAAGGGCTTGTGTTCTCAACTCGTTGATTGAATCACC
ATGGCCAACGCAACTTTTGTACAGCTTTTCTTTGCCCACATTTATGACGCGGATGACACCGTTGCTCCCACTTGCAATTAACAA
AGGAGAATTTGCATCATCACAAGCCCAACTTACAGTGTAGAAGGCTTCATCTTTATCATCATCAACATAAGTTTGCAACACTG
CAATTCCGTTGTTCTCGAGCAAGTGATAAATTGTTACCCGGTTACCGCCTGCAGTGGCGAAGACGTTGTAGAAGCGGGAGTCA
ATGAAGTTGAAGCAGACAGCGTAAATGGGGCGTTTTCCCTCCTGGAGTCTGTTGGAGACCTTGTATTCCTTCCTCTTTTTAGCG
CCAGCCCCCGTGAGACCCACCACAGGGTCGGTCCCCAGCCCGCTCTTCACAGACATGTTCACAACGCACTCCTCACTAGCGAC
CTCAGATCAAAA

**MULTICOPY SUPPRESSOR OF IRA 1 (*MSI1*):**

AGAGAGAGAGAGAGAGAGTAGTAGTAGTTTATTTTGTAGGTGTGTGTGTGATGTCGAAAGAAGAGGTTGAGTTTCGG
GATGAGATGGAGGAACGCCTCATAAACGAAGAATACAGGATATGGAAGAAGAACACGCCATTTTTGTATGATCTGGTGATTA
CCCATGCGCTAGAATGGCCCTCTCTTACAGTGCAGTGGCTTCCTGATCGCAAGGAGCCGCCTGGCAAAGATTACTCTGTGCAG

AAGCTCATTTTGGGTACCCATACCTCAGACAATGAGCCCAACTATCTCATGCTGGCTGAGGTTCACCTCCCTTTGGAAGACTC
GGAGAGCGATGCCCGCCAGTATGATGAAGAGCGAGGAGAGGTTGGTGGCTTCGGTTGTGCTAATGGAAAGGTGCAAATAACT
CAGCAAATTAACCACGATGGAGAAGTGAATAGGGCTCGGTACATGCCTCAGAATCCATTTATTATTGCAACAAAGACAGTGA
GCGCAGAGGTTTATGTTTTTGATTATAGCAAGCACCCTTCCAAGCCACCTCAAGAAGGGCGTTGCAACCCAGATTTGAGACTG
CAGGGTCACAAGACAGAGGGGTACGGCTTGACTTGGAGCTCTTTGAAAGAGGGTCATTTGCTTAGTGGGTCAGATGATGCCC
GAATATGTTTATGGGATTTGAGTGCAAATCCCAAGGGCCAGAACATGGTTGACGCAAAACAAATTTATGAGTACCACAGTGG
CGTTGTGGAAGATGTTGCATGGCACTTGAAGCACGAGCATTTGTTTGGATCAGTCGGAGATGATTGTAGGTTGCTGATTTGGG
ACACCCGCAAAAGTGGTGATAAGCCCTTAAATTCAGTTGAGGAGGCTCATCTTGCTGAGGTCAACTGTCTAGCCTTCAATCCT
TTCAATGAGTGGGTTCTGGCCACAGGATCCGCTGATAAAACTGTTGCTCTTTATGATCTACGAAAGCTGTCCAAACCTTTACA
TACCTTTAGCAATCACACGGAAGAAGTGTTTCAGATAGGTTGGAGTCCTAACAACGAGACTATTCTTGCATCTTGTGGAGCAG
ATAGAAGAATGATGATTTGGGACCTTAGCAAAATTGGGGAGGAACAGTCTCCTGAAGATGCAGAAGATGGCCCACCAGAGCT
ACTCTTCATACATGGTGGTCACACAAGCAAGATTTCTGATTTTAGCTGGAATCCGAAGGATGATTGGGTGATCTCAAGTGTGG
CTGAAGACAATATCTTGCAAATTTGGCAGATGGCAGAAAACATTTATCATGATGATGATTTCGGACCCATGGATGATGGGAA
AGCTGTTTAATGAAGCAGCTTATTTCTTACAACAAGATTTGTATCACACGTGTGCCTGAGACGAGACTTGCTCATACAGGACA
TCTTGCCACCTGAGCAAAATGGAACATGGTGGCGTTGTTGCCTATCACCTTCATGTGAACTTGCATCGTTGTAAACGTTGTTTT
TCTTTGAGGTGAGCTATGTTTTAGGAGCAAGCTGCCAGGTGTCTTTGGTAGACACCAACCCAGGTTAGGTGCATGAGATTTTA
AAGTTTTATACCATGTGTGCTTTTGATTTTTAATCTTGTGACTTCCTTGCCAATGCAGTAGCCTAGAAAAGTCGCATTCTTGTTG
CATTCAAGTTGGCAGGATGATGACATTTGGGGGCAACTTTTGTTTGGTTTACTTAATTTTTATTTTGCAAGATGTTTTTCAATA
CTGAATGGTTGTCTAGACTGTACTGTGTTATTTGCTTGGGCTGCGAAAGACTAGAGAAGGCTTTGGT

**BABY BOOM (*BBM*):**

AGAGAGAGAGAGAGAGAAACTAGATTGTTGAAATTAGTGGGAGTTTTATACAAGAAAATTAAGTATCAGTCCAAGCG
GCAAAAAGGGGAGTGGGGCAAACGCCCAAGTGGTTAGCTTTCAAAGGAGAAGAAGCGGAGGCAAAGGGGTTAGGCCAATG
AGCTGCATGCAAGTTGATAGCGCTTCTTGCGTGGAGGCTCTCGTGGTAGCTGCTGCCATAGTTGGACCTCACAAGGCTAGGCG
TCGTATACGTGCAGGTAGCTGCTGCTTCCGGCCTCGCATGCATCATACCTTCTTCGCTTGTCGCGTGCCCCATACCCATATTAT
TGCTGTAGCTGCCTGCGTTCACCAGCTGCTCAAATGCGCAGCTCTTGCTCCCCCCTCCTCCTTCAGCACCACCATGGAAGGGTA
ACTTGGAGCTGAATTCAAAAGCTGAACGATTTTGACTGGAAGATGGGTGCTGATCCAAGCTGCTCAGCAAGTTGTGCATTTGA
AGGTCATGTACTTGCACATGGGGATCTTGGTATGTTTTCAGGCTACTTGTAGTATGGCAATCCTTCTGCTCTTCACTTTCTTCCA
TATTCTTCTGATGCACATTATAGCATGACTCATAGCTTTGATCTGGCTTAGAGCTGCTCCAAGTTGTAGTTGCCTGCTCATCAC
TAGTACAATTATTGATGCTTTGTGGATACCCTAATAAGAGCTGCCAATCTTGAGCACTTCTTGCTATACTCTCACTTACATGAT
TATGCACATTAATATTATTAGTGCTAGCATTGATATTATTGGAAAGATTGCTTGTATTGCTATTAAATGAGCTTCCTTCCATTA
AGCCAATAATGAGATTATTTTCCCCCGAACCTCCATTAAAATAGTGATGGTGCCCATTATGAAGAGGGTTGAAGAGTGAGGCT
TCATTGATTGGGGATATGTCTTTGAGCATGCGCTTGGCGAGTGTGGTGATAGGAAGGCTTGTGCTGCTGCATATCTTGGCGAT
GTCGTACCTGCTCATGTCAAAATTGGTCACCGCATTTATGCCTCGGAACTTTATGGCCGCTATATCATATGCCTCCGCAGCCTC
CTCTTGTGTGCCGAATGTTCCTAGGTACAAATCCTTGTTTCCTGCCACTCTTCCAATTCGAGCCTGCCAGCGTCCATGTTGATG
ATGTCTTGTGACTCCACGATAGATCGATGCTCCTCTAGAGAAGCCACTACTTTTTCTTCTCAAGGAAGCTACAAACTCTTGCCT
TGTCATGTTCTTCATCTCCTCCAATTCTTTTTCATAGTCATCAAGCTGGAAATTGATGGTAGTTGTAGGACCCCAGTACTTCAA
TGCTGCAAGATCATAGGCTCTTGCTGCTTTCTCCTCTTTGTCATATCCACCTAAGTATACTTGTCTTCCTTTGCGGGTTTGCCCC
TCCCGTCTGCAGCTGTTGTCCCAAAGGTGTGCTTCGAACCTTCCGGTCCAACGATGCTTTGTAACCCCGCGGTAAATGGAGGT
CCGCTGCCCAAAGGATTCAATGGACTTTCTAGGGGTAGACTCTGCCACTACATTAGCTGGTGCTGGATCTGGGCTGTGCCTCT
TGGAGTCCGTAGTGGTGGAAGAACTCTCTTGCACAGCTAATGTTCGATCCGTTGGGTTTGTAGCGTTTACATGCATAATGCTG
GGTGTCATTGATAGAGTTAACTGTGATGCTTCAAAAAGAGGCAATATTGGAAAGCTGCCCTCTTTCATGGCCTCCATATTACT
TTCCTTTAGCATGCCATTCTCTGTGTTAGATAAACTCTGGGACCTCATCCAAGAGTTCATCAATGAGAAGCCCAGCAAGTTTGT
GCTGTTTTGCAAGCCTAATTCACTGGTAAGGCTGGTGTTAACATTGTGTGTATTGCTAGAGGAGGTTACGTTCGCATTTCTAGC
TGAAGAGGTGATATGGGTGTTGCTTGTTGTAGTGTAAGGTAGCATTGAGATTAAGTGCTCATTAGTAAAAGATGATGAGTGAT
TAGAGCTAGGTAACACTGGATCTATGGGATTTTGTAGATGCATTGTGCTTGCAGGTTGGAAGCTATTATCTGCATTGTAGCCT
CCAGAAAATATATCGTAGCTTTTGTGAGGTTGTGGATTCGCTGAGGGGCTGCTGAGAGGTTGCTCTAGTGTGGTATTGCTGCA
AAACAGTTTGCCTTCAGCTTGTGGGATTAAGGGATTTTCCTCCTTAATGTAGTGGGATAGGAAGTTCTCCAGCTTTGCAGAGC
GCGGCTGCGTGTTGTTATTGTCCGCAGTGTTCAAGTGTTGGGCCAGTGATATGCCGGTAGCAGCAGCTGCATCTTCGTGATCA
TCATCAGCTCTGATGTTGTGAGTGTAGTGGCCTTCAAAGTCCCTCATGTTTGACCAGTCAGATAGCCCGGAGTCGAGATGTAA
GTGGGAGGCGGAGGCGGAGGCGAGCGCGGAGTGGACGATGCCACAGAGGGAGTGATGAGGAGGGAGTAGAATGTGCTCCA
TAAAATTTGGGAATGAATGCGCGCGTCGGCTTTGCTGAAAATATTGTAAAAGCGAGGTGGCGATCACTGGCGCAGACACACA
CACTGTGAAATACACAGTAGTCACTCAACACAGGGCACACAACACACTGTGACACAAACACACACTGTATTCACTCAACACA
GGGCTCAGTAAAATTACACACAGGCCTCTGGACTTCCACACGACAGTCAAGCACAAGATGAAGGAGTGCTGCTTCTGCTGCC
CCATTTCTGGTATTTATACTTTATATTTATCCCAATCTATTTTATTACTACTCTCCCTATCTCTCTCATTTAATAGTGTACCATTC
TCTATTTTCAAATAATAGTTATAACTTATTGTTATAATTTCTTCACCCGAAACCTTTTTTTATATTTAATTTTCTTCATCGGTTTT
AGTCCCCTAATAAAAATTTAGCTCTCCATGTACCGAAACATATGACGAATTTCATTTTCAT

**MATERNAL EFFECT EMBRYO ARREST 29 (*MEE29*):**

TCAAACCCCATCCACCCTACAGTCAAAACTCAATATCGTCGTCATTATTATACTGACGCGCTCTCGATGGTTTAGGGTC
TGAACGTCGGCAAGAAGATAGAGATTTTGGGTTTGCAGGGAAAATAATTTTGAGACTACCCATGGCGCAAGAAAAGCAGTTA
AGGGTATGGATCTCAGACAATCTTCATGCCCTGTTGGGTTTCTCAGAACCCAACCTGGTGCACTACTTGTCCGGGCTTGGTAA
ACGCACAACCTCCGCCCAAGAGCTTGCCTCAGAGTTAGTCAAATTGGAGCTTCCCTCTTCTTCTGAAACTGTCAGATTCGCTG
AAGAGCTCTTTGGACGCTTTCCGCGCAAATCCACAGGACCCAACAGCTACAAGCAGGCGGAGCGTGAGGCTGCTGCATTTGC
CCGCAAGCAAAATCATTATCAGTTGTTAGATGATGATAATGATGATGCCCTTGACAACGTGCAGGAAAACCAGGTAAAGAAG
AAAGATGATAATGTTGGGTCCAATGGTTCCGTGCCCAGAAAGGGTCCTAAACACATTCGGAAGAGGCCTGCAAATGTTTATG
ATGATGATGATGTGAGTACTGGCAGATTATCGAAAGAGAGAAGAGTACAGAACAATATTGACGACGATGATGATGATGAAG
ATGAGGATGATGAGGCTCAAAGAGAGCGTGAGAGGGAAGAAGACCAAAGAGAAAGAGAAGCTCTTGAGAAAAGGCTTAGA
GAAAAGGATGAAGCCGTCACAAAGAGACTTATGGAGCCTAAAATGACCTTCAGAGAGGAGGAAGAGGCAAAGCGACGTGCT
GAAGCTGAGGACAGAAAGGAGCTGCTACCCACTTTAAGAGAAGCTAGTAGACAAGTGTACTTGAAGTCAAGGGAGCAGAAA
AAGATTGATGAACTTCAGGATCAAATCGAAGATGAGCAATACCTGTTTGAGGGAGTGCAGCTTACAGCAAAGGAGAAGGAG
GAACATAGGTATCGAAAAACTGTGCTTGAGCTGGCCAAACAAGGAAATGAGGACATTGATGGCATCGAAGAGTATAGGATG
CCTGAAGCATATGACAAGGAAGGCAGAGTCAAGCAGGATAGGAGATTTGCTGTGGCTCTGGAGCGGTACAAAGATGTGGAT
GGTGAAAATGATAAGCTCAATCCTTTTGCAGAGCAAGAAGCTTGGGAGGAACATCAGATTGGTAAAGCAAGGCTCAAGTTTG
GTGCAGCAAATCAAAAGCAGAAAAGTGGTTATGATTATGTTTTTGAGGATCAAATTGAGTTCATCAAAGCCGATCTGCTTGAG
GGGTATAAAGACCAGGATATGGATATTGAAGAAGCTGAGGAACAAGAGAAATCAGCAGCAAAGTCAGCTCTGTTGCACCTG

CAGGAGGAGCGCAAAACTTTACCAATTTTTCGTTATCGAGAGGAGCTCTTGCAAGCTGTGAATGATCATCAGGTTCTTGTCAT
TGTTGGAGAAACTGGTTCAGGAAAAACAACCCAAATCCCACAGTACCTTCATGAGGCAGGGTATACAAAGAAAGGAAGGAT
TGGTTGTACACAGCCTAGGAGAGTTGCTGCCATGAGTGTAGCTGCACGCGTCGCACAAGAAGTGGGGGTGAAGCTTGGCCAT
GAGGTAGGATACTCAATTCGGTTTGAAGATTGCACCTCAGACAAAACAATAATGAAGTATATGACAGATGGTATGCTTCTTCG
TGAGTTCTTGGGTGAGCCAGATTTGGCAAGCTATAGTGTTATGATGGTAGATGAGGCTCATGAGCGAACTATTGCCACAGATG
TTTTGTTTGGCCTAGTGAAGGATATTATCCGTTTCAGGCCAGACATTAAGCTTCTCATTTCAAGTGCCACATTGGATGCCAAGA
AGTTTTCTGATTATTTTGACCGTGCTCCTATTTTTAGCATCCCTGGAAGGAGGTATCCAGTTGACATTATGTATACAAAAGCTC
CAGAGGCAGACTATCTAGAGGCAGCTGTTGTAACTATCTTGCAGATTCATGTGACACAACCATCAGGAGATGTGTTAGTTTTC
TTTACGGGGCAAGAGGAGATAGAGGCTGCAGAAGAGATACTAAAGGCACGCACCAGAGGCCTGGGTACCCGAATTTCTGAG
ATGATTATTTGCCCAATTTATGCAAACCTGCCATCTGACTTACAGGCCAAGATTTTTGAGGAGACCCCTGAAGGAGCCCGGAA
AGTAGTTCTGGCAACCAACATTGCAGAAACATCTCTCACAATTGATGGCATAAAATATGTGATAGATCCTGGTTTTTGTAAGC
AAAAATCTTACAACCCACGTACAGGCATGGAGTCGCTTATAGTGACACCCATTTCAAAAGCAGCAGCTCAACAGAGGGCTGG
CAGGGCTGGGAGGACGTCGCCTGGAAAGTGTTTTAGGTTGTATACCGCATGGTCTCATCAGCATGAGATGGAGGATAATACT
GTACCAGAGATCCAGAGGACAAATTTAGGGAATGTTGTTTTAATGCTGAAAAGTTTAGGCATTGACGACCTCATAAATTTTGA
CTTTCTGGATCCACCGCCTGCAGAAACCTTGCTTAGGGCTCTAGAACAACTGTATGCACTGGGAGCTCTAAATCACCGTGGAG
AGCTGACAAAACTGGGACGCAGAATGGCAGAATTCCCTCTTGATCCGATGCTTTCAAAGATGATTGTGGCGTCTGATAAGCAT
AAGTGCTCAGAGGAGATAGTGACGATAGCTGCAATGCTGTCTGTCGGAAATTCTATCTTTTATCGGCCCAAGGACAAACAGG
TTCATGCTGACAATGCAAGAATGAACTTCCATGCAGGCAACGTGGGCGATCAGATAGCCTTGCAGAAGGTTTATGATACCTG
GAAGGAGACAAACTTTTCAACACAGTGGTGCTATGAGAACTACATACAGGTACGGAGTATGAAGCGTGCTCGTGACATCAGA
GATCAGCTTGAAGGGCTCCTTGAACGAGTTGAGATTGAGCCTAGCAGCAGCCCAAATGACATAGAAGTAATAAAGAAGGCCA
TAACATCAGGCTATTTCTACCACACAGCAAAATTGCAAAAGAATGGTTCGTATCGAACAGTGAAGAACCCTCAAACTGTGCA
TATTCATCCAAGCTCAGGCCTTGCTGAGGTTTTACCTCGCTGGCTGGTATATTTTGAGCTGGTGAATACAACCAAGGAGTATA
TGCGACAGGTTATTGAGATAAAGCCAGAGTGGTTGGTGGAGATAGCTCCCCATTATTACAAAAAGCAAGATGTTGAAGATGC
TGGCTCGCAGAAGCTGCCCAAAGGCATGGGAAGGGCCAACATGGATTGATCACTGTGACGACAAGTATATGCATGAGAGGA
GAGCTTCAGATTATGGAGGTAAATGATGTCCAGAACTAGCAGTGTTGGTTCTCTCTTTTTGTGATGCCTGATGAATATCAAAT
CAAGCAGAGCGCTTATAGATTTTCCAAGATTAAAATCAACATGGATATTTTGGAGTGTGTATGGAAGTATTTGGGACCCTTGG
GCCTTGCTGCTGCCTGTGTCGACATGTTGCAAGCTTTGTTTGAACTTTGATCATAATGGTGGTTCTTGGAGGCTGGAGCGGGTT
TTCAATAGCATGGGGATTGCAGAGATTTATTCTATGCTCTCAACAAGGTCTAATGGATGCACCACCTCTTGCAGAAATTGCAT
TTTGACAGTTTTCACTAGTGAGACAGCACCAAATACAGATGATTTCTGATCTACTTCTGTATTTGCTGCTTTGAAGCAGGTGGG
GTGTGGAAAGTACGTCTTTCCTTGCTCTTGCAGCTCATTCAGGTAAGCAAGCAGTAATGAGACGGGCGCATGGATATAGATTT
GTCTGAGCGATGTTGGCTGTCCACTAATCCAACTCTACAGTTGTTGTATGGGTCTGCGTCATGTCATCATGCATAGACATTCAG
GTTCCTTAAACCAAAGGTCATGAAACACAAAGCTTTGAGCATGGACAGCAGAGATCATAGATACAAAGATCTGTGTACTTTTT
CTTCACAAGCTTTTGCTCTTTACTTTTGTGCAATGTTATTGAGGTATTTAAGCTCCAACAATGTATGCCCCGACCAGTGTAAAT
TATTGCCATGTTTGCTATTTGTTTTTTGATGGTAGCCTCACAAAAATTTCAGTCCATCAGGAACCAATCTTTGAAAGCTGTTGG
CAATGAACGGTATGAGAGTAGATTTCTATGTACCAGGCTTGTGG

**AGAMOUS-LIKE 62 (*AGL62*):**

GTGTGTTTACGTGTGCGCGCGCGAGAGTGGGCTGCTTAGACGTGGCAAAACTTGCCAAATTAAACCCTTTACAAACGC
CCTTTTACATCCTGAATTTCTGCGTCGCTTGGTAGCTACAATGTGCATTGCTTCTCTTGATCGCCCAGACAAACTTAAAATGCT
GATTAGCAATGTTAATTATCCTGTTTAATCATTGCTAATTACTATCAGAAATTGCGTATATCAACAATACTTAATTAGTTTCTA
ATTAATCTCTATTAGTCTTAATCATATATACATATAATGGGCTCTTTAGATGCCTGTGGATTTCTTTATTTTTCATTCCAGAGAT
TTACTGTTTTGTACTATCTATCTCTATAAATCTTAATAACTTCTGATAGAAATTGCTCCAAAAAAAAAGAAAGTACTACGAAAA
AATGGTTAGGTCTTTGCCCTAATTTCTGCTGAGGAGCTGAATGATACTATCTGAAACAGTGATCAAGGTATTTAATTGAGAAA
TCAAATGAGATTTTCTGGAATTCAGATGATTTGCAACAATGGGTCGTGCGAAAATAGAGATCAAGAAAATCGAAAACTCAAG
CGCGCGTCAGGTGTGCTTCTCAAAGCGGAGAGTAGGACTCATCAAGAAGGCAAGCGAGCTCTCAATTCTATGTGGGTCAGAG
GTGGGCATCATTGTGTTCTCTCAAGCTGGCAAGGCCTTCTCATTCGGGCACCCATGCATCGACTACGTGATCGACAAGACGCT
CAAGAGACCTGTTTCCGTAGACTCCGACAAAATCGAGGTCATACGGAGGCTCGAGAGCGAGTACAACACACTCCTGCAAGAG
CTTGAAGTCGAAAAAGAGAGACATATCGCCTTGCAAAAGCAGCTTCATGTGGATTACTACAACCATAAGCATTGCTGGAACC
AAAATTGGTGGCAAGAGCCCATTCATGCCATGGGTTTGCTAGAGTTGAAGCAGCATGCGGAGCGGTTAGAGGCATTCTACAG
GCTTATTGTCGAGCGGGCTCGATACTTGCAGTACGTGTCTAGCTTGGATCTGCCCTTAATGCAGCAGTATCAAGTGATGCTTA
ACAGACAACAAGCATCAGCGGGTGCTTTCATGCTTAAGCAACAGTTCTCGCCGGACCATCTTCATCACCATCAGCATGCCCCT
CAGCAACTTCCCCAGTATATGGACATGAATAGTATGCCATACAAAACTGTGCGCTCTTCACGGGCTCTTCACAAGGAGTTTAC
ACAAGAAGTGGATACAAGGGATGATACCCAAGGAGAGAGAACTATAATCTGCTATGAACACAGATGACGATCAGTATTCTTG
AAAGGGGTGCTTCGCTCTGGTGCATGTCATAAAGCATTTTTCCACACGGCATGCGCCCATACATGTCAAGCCGGAGATCAGAT
GCACAGTTGAATAACATTTGTTTACTGACCACTTGTGACATGCATGCATGTCTCCTGATTGATGTTATATGTAGATTTAACATA
AGTATGCATGCAATAGAAAATGGCAAATTTTGGTCTGCTCCTGTACGATGCCTACTGGCGAGCTGTTGGGGATGGTTGACCAG
TACTGACTTGGCCACCAGTTACGCCTGACGGACACGTAATGAAGAAAGCCGATAGAGTTTTAGGTACTTCAATTATTAAGGTA
CATTTTTTAAAGAGAGACAAGATATATTATTATGACAGTGTTTAAGGTCTAATGATATGGTATGAAAGTACAGCAGCCTCCAA
AATAAGATCCTGAAACAATCCTGCATGCCTTCATTCTACGCACTATTTATGATGTATGTATACCGAATACGTCTAGCAGCCTCT
GAACTGAGTACCTTAAACAATTAAACAAACTTTAGTTTAGAAAATTAGGCATAATTAGTTAAACAAAAACTTAAATTGCCAA
ATGATTTTGCAGGCATGCATGTGTTCTACATGATATCCGTCAATATGATGTTTCTTATGTCCTCAACCACATGTACGTGTGTGC
ATATTACGTGTCTGTTGATAAGATGTGTCTTTGTGTACTGTTCTACATGTATGTGCATGCACCTTGAGATAAAAATGTGTCTTA
TGTATGTACTACCACATGTATGTGCGTACCACAGAGCTGAATGTTGTTAATTGTTAACTTTTTGGCAATGAAGGTAGCAATGG
AGAGAGACATGGGTTCGATTGCTGGAACTTCAAATCTGGAGCGCTGTTTTTCAGTACAAGGTTTGGCGAATTCGATACCAAAG
GAAGAACCCAATGCTTCTTTCAGCAATATGCTGCTTCTAACAGCAGCAGATGAGGATCAAATACACATGAGTCAAGAGCGTG
TGAGGTGGGCAAGCCTAGAGCAGTCGTGTGTGCTCGAGCAACCAGAGCAACTGCCAGATGAACAAGCGTTGATGGGAGAGG
AAGGGCAACAGGAACCGCCTGATGAGAGCGAATTGGAGGCTCCCATGCCAATGGATCAAGCATTATATCTAGCAGATGATTA
TACCACCGATAACGAAAATTTTGACACCGACCTCTCAGACAATAATGAATACATGGATAATATGCTTGGAAGCTTGGCAGGTT
CCCTCTCTGCTGATCATCATGAGGATCACAGTAGTGTGCACGCTTCAGTTGACCTCTCTGCTGATGAAGAACATCAAGGGGAC
AATGTAATTAACACCGCTAATGACGCACATCGTAGTGATATGGTCGTCAACTTCTCTGCTAATGTGGATCATGAGGGTGATAT
GAATATCAGCCTGTCAGCTTATGATCTTCATGGTAGTATGGCTACAAATGTTGGTGGTAATTCTGAAAGTGGTTATAACATCA
ATGTACCAATTGAAGATATGGATGGTAAGTGTCCTGATATGGCTCGCATTCGCACAAATATGGACAAAGTGTCTTTTGATGAT
CCAGATAACCATGTCAGCCTGTCTTCGGAGGATCACGGAGTGCTTGTAGCAGGTCTCTCTGCTGATGATGATGCTAATGCTGC
TTGCCTCTCCGCTAATGATGACGATGACGGCAGTAAACTAGATAGCCTTGATGCCACTGGTACTAGCTTTGGGAAGTTCTATG
CTGATGGTGGTGAAACTTCTATTATGGCCAACCTTTCGGTCATTAACGAAGTGGAGAATGTTGCATCCAGCCCTCGTGCAGCT

CAAGAGAACATGTGTGATGGAAATTCTGGAGCCCTTTGGCCCTCTTGCGATGCTACTTATCTTACCGAGAATTCGTCCATACA
AGAGGATTCAGATCAGCTTATAGTTAGCATTTCAAATCAGGATTAGCCACTGCTACCAATCGTTAATGGAGTTGATCATCGCA
ATATTGTGAGGAACAATGTGCAAGGTAACAGTAATTAGCCATCTAGCAGCGCGATGTTTCAGCAGGAATGTCTTCATATCACT
AAGTCGAAAAGATGGTCTTAGACCAGTGTTCACAGTTTCGGCTAGAAACTCGAGAAAATCACATAAAATTCTACAAATACAC
GACGTATGTACAGACCTCTGCCCACTTAACGTTGCTCAAGTGTAGCACACAGAAGGGCTACCATAGGGTCCTGCACATAGTGT
GCATCGACTTAACAGTCATCATAATCAGTCATCGTCATCATGATTTAAGGAGTCCTATGCTTTACGGAGGCAGCACAAGCTGG
TATTATTATGCTAAATATGTATATACATATATATATATATATA

**AGAMOUS-LIKE 6 (*AGL6*):**

GACGTTCTCAAAGCGGCGGGGAGGCCTGTTGAAGAAAGCCCATGATCTCTCTGTCCTCTGCGACGCTGAGGTGGCAGT
CATTATCTTCTCGAGTAAAGGAAAACTCTTCGAGTTCGCCAGTCCCAACATGGCAAGCATATTGGACCGCTATTTGAAGTGTA
GTGAAGATGTAGGATGTGCCGATAACAACAATCTTTCGGATAAAAGAGCATTGACCCACTGCACAGAAGATCTTAAAACTTT
GCAAAGAAACTTTATTGGTGAGGATTTAGAGCGGCTATCTTTACAAGATCTTATCAATTTGGAGCATCAAATTCACGACACTT
TAGGTCGAGTTCGCACAAAAAAGGGGGAGCTACTCATTGAACAGCTCGAAGATATCAAGGAAAAGGTAAGGACAACAAGTG
CGAGTTCGAGCTTCTTGGCTAAACTAGTGGACTCATTCCCTTTGGATGCGGCGGGATCACAGTATTCAATGAATAACAGCATT
TGCACGATGCAGAACATTCCAGA

**ARGONAUTE 4 (*AGO4*):**

TTTTTTTGGAACAAAACACTTTTCTTTCATTTGTAGTGCATCAAGCTAACACTGCTTTTACATCAAAAAGTTAGAAGGC
CTCTCCTTTTCTGGAGTGGCAACCATTGTCTATACAGCAGCTCACAAAAGGCAAAACACATTCTATGGTGACCTTCATCAGGA
GTACAAAAATATTGCCTCTGTAATTACAGCTGGTGCTTAAGACCCTAGATGTACTGTAGAAAACAGCTCAGTACATGTTCGAG
CATGAAGACCCTAGCAGTAGAACATTTTAGTCTTCATTTGCGGATGCAGCTCCGGTAAATGGATAGGAGTTGACCGCTCAGAT
CTCCCGGTCATTGACGAAGTTTCTGAGCCACCCTCAGAGTCCAAGAAATTCCGGGCGTGCGTGGCCGCAAGGTGAGCATATG
CAATCGGAGCAACGGTTGAAACTGCTGTGGTGCTTCGAGCATATGTGTAACACAATGCATGAGTCATCTGTTGCAACTCATCC
ACTGTGAAGTTGTTCTCGTCAACCAAAACATGGTAATGAGTGGGGCGAGTTGTACCAATCAGGCCTGCGTGTGCGCACAGGT
AGAAGTCATAGTCCCTTGGATGGCAGAGGGTTGCATCCACAATGGTACCTGGAGCTACATTTCCACCACCGGTTGCTGGAAA
GAACCGAGTATGGTGCCTCTTCTGTACCACAATCAAAGTTATTCTCGGATCATAGCTGCCAGTGTTGTCTATTTTTTTGAAAGC
CTACACAATTAAAACAGAAATCATTATATAATAACTATACTAAACAACCGCATGTGTTTAAAACATCAAAATCAACAGGCTG
ACTGTATGTAACAGCCATATATAGCCATGTGACTTCATTGTAAAGGATTGTAAGCATTTCTTGCCCTCTTGAATGCCAGATATT
CATCTCTTAAAACTTGCTCAAACTGGCTTTCACTAACACCATCCCTAAAAATTATCACTTGCTTAGGTCGAGGGTCAGTGATTC
CAGGAACTTTCTTGCAAGAAATGTAAAAATCCTTCAACAACTCAACAATAATTCCACCACTGGGGTTGTCTTCTTCATAGAGC
CCTGCTATCATTTCAACTTTCCGAGACTGTGCTCTCATACGAACACCGTACTTGGAAAAGTACGGCCATTCTCGTGATGCCACC
ACTGCAGCTATTGAAGGAGAGTCTGAATATCCAGGAGGCCCATGAGAGACGTCCATCCCAAATATGATTGTAGGAACACTGG
ATATCTTGGGAAGTTTAGAGTTCAATTCCAATGCCAGCAAAGAATTATACCCTCCCACCTTTGCATTAATCTTCAAAGCAACA
TTGGTCAAATATTGGTCCTTCACTAGGTTATAAGGAGGAGCTGGGGAGGAGCTATGCATTGTGTGACAATGCACAGATTAGTT
TGCAACATCTTTTTTAGAGGCCCATATAAATCACTCTTGCTTTCCGGAAGAATACACAACATAAAAGCAGGCGGTGATCGCAT
TTTTGATTTCAAATCCTGTATTATTCGGTTAACCCTTTCTGCAGCAGATAAGCCTTGATCACTGAGTTGCTCGCATAGTATGAG
TTCGGGGCGTGAGATTTTCATTCCCTTGGTGCTGCAGACTATCTCCAGATCATTTGCAATCTTTTCCACAATATAGCGAGGCTT
TTTGTTATCAAAACTTATGACTACCCAGGATTCAATGTCGACAGCAGATGCCATTGTTTTATTGTTGAAGTTCCACCTCCCATT
CCGTGGTGTCTCTGTTCTAGACTCCCCAAACATCAAAGTGGGTGCGTCTAGTAGCCTTGCAGGGATCCTCCGCATTCTAGAAT
CAATTTTTATGTTAAAATCCTTTATGAGCTCATTTGAAGAATAATCACTAACATCCATAGCCTTTTGCACAACATCCCTTCTTT
CATCTGGTCCCTGGCGAGCTTGATCAATCATTTTCTGTCTCTGGGCACTTGACAGGGCTTTGGTGTATCTTTGTCCCGCCACAA
TGTTGCAAAGCTCTACTGGGATAAATGTGGGCCTCTTGGTTCTACCCGCTGCTATGCATGGCAAATTTGGAAACCGAAGATTT
ATGTTGTATTGCATTTCGTAATATTGTAGTATCGTCAACTCAGTCTCTACGAATTGCCCAGATTCATCCTTCTTCCTTTTCGTGA
ACTTCTGCTGCGAGCAAGAAGCATCACTGAAGCCTATAATTCTATGCACCATCTGGGTGTGGGGAGTCTCAACCTTGACCCCT
TTCAAAACTCTCTTCACCTTGTCCCAATCAATGCAATTCGGACTAGGGGGATTAAAGCGTTCCTTAATGAAATCCAAGACGCT
AGAAGCCTTGATGATAGTTGTTGTAGCAACGTCAAGGTTTAGTGACAGCCCCCTAAGAGAAGGCCGAAAGCTAACGTGATAC
CCTCGACAGGTTTGAACTCCTTCCCCAATATCAGCAACTCCAAACAATGGGCTGAAGTAGTTGTCCTTGATTAGGAGATAGCC
CTTGGAGGATGCATGCTGCTTCAGAACAATATCCAGCACCCGCAGAGCATCTTGGGCCCTGTCATTCTTTTGTCCATTCAAAA
TTGCATTGATGGCTCCCATCTTCACAGTAGCTGCATACTCAATTTTAACCTTAAATTCTCTAGTGTTCCTGGATGACATTTTGC
GCCGCTTTGTTGAATCATCATCTGCTGAGTTTGCTTTGTTCATCGCAGACTCCTCCAACAGGACTGTAAACTCCTGGGTTTGGA
ACTCGAGGGCACCTGGCGTGAATAGACTCTTCTCTCCATCATAGGCAAACCGCTTGCCTCCTAGTTCTTTTGCTCCGTATACTT
GTTGAAGCTTGCTGATAATGGACCGACAAAGGTTTTTATTTGCGATACCATCGTTTCCACCAGGCGCTGAGCCCCCATCTGAG
ATACTCACATCGTAGTGGCAAACGTCCTGCGATCCGGAGTAGTTGACCATAAAGTGATTGCAGAGCAGCTCAATGGGCCTGC
CCGCTTTTCCTAATCCGGGTCTTTGCATTTGTGAAGGCGGCTTCATTGCAGGTACGTCCGCCACATCTGTTGGGTTCGTTGATG
GCTCTCCTGTGAGTGCCGCATCTCCTGCTGCTTCTGCCACCGCTGCGGGGTCCCCCATGGCTCCAAGCTTGAGAATGGCACTTG
TGCAGAAGAACTGAAGAACTAAGGGCTGCGCAGTGCGAATTAAGAGAAAGGGCACAACTGAGAAAGGGAAGGTAAAAAGG
GGGGGAAAAGAGAGAGATAGAGAGAGAGAGAGGGAGAGAGA

**ARGONAUTE 9 (*AGO9*):**

TTTTTTTTGGAACAAAACACTTTTCTTTCATTTGTAGTGCATCAAGCTAACACTGCTTTTACATCAAAAAGTTAGAAGG
CCTCTCTCTCCTTTTCTGGAGTGGCAACCATTGTCTATACAGCAGCTCACAAAAGGCAAAACACATTCTATGGTGACCTTCATC
AGGAGTACAAAAATATTGCCTCTGTAATTACAGCTGGTGCTTAAGACCCTAGATGTACTGTAGAAAACAGCTCAGTACATGTT
CGAGCATGAAGACCCTAGCAGTAGAACATTTTAGTCTTCATTTGCGGATGCAGCTCCGGTAAATGGATAGGAGTTGACCGCTC
AGATCTCCCGGTCATTGACGAAGTTTCTGAGCCACCCTCAGAGTCCAAGAAATTCCGGGCGTGCGTGGCCGCAAGGTGAGCA
TATGCAATCGGAGCAACGGTTGAAACTGCTGTGGTGCTTCGAGCATATGTGTAACACAATGCATGAGTCATCTGTTGCAACTC
ATCCACTGTGAAGTTGTTCTCGTCAACCAAAACATGGTAATGAGTGGGGCGAGTTGTACCAATCAGGCCTGCGTGTGCGCACA
GGTAGAAGTCATAGTCCCTTGGATGGCAGAGGGTTGCATCCACAATGGTACCTGGAGCTACATTTCCACCACCGGTTGCTGGA
AAGAACCGAGTATGGTGCCTCTTCTGTACCACAATCAAAGTTATTCTCGGATCATAGCTGCCAGTGTTGTCTATTTTTTTGAAA
GCCCTCTTGAATGCCAGATATTCATCTCTTAAAACTTGCTCAAACTGGCTTTCACTAACACCATCCCTAAAAATTATCACTTGC
TTAGGTCGAGGGTCAGTGATTCCAGGAACTTTCTTGCAAGAAATGTAAAAATCCTTCAACAACTCAACAATAATTCCACCACT
GGGGTTGTCTTCTTCATAGAGCCCTGCTATCATTTCAACTTTCCGAGACTGTGCTCTCATACGAACACCGTACTTGGAAAAGTA
CGGCCATTCTCGTGATGCCACCACTGCAGCTATTGAAGGAGAGTCTGAATATCCAGGAGGCCCATGAGAGACGTCCATCCCA
AATATGATTGTAGGAACACTGGATATCTTGGGAAGTTTAGAGTTCAATTCCAATGCCAGCAAAGAATTATACCCTCCCACCTT
TGCATTAATCTTCAAAGCAACATTGGTCAAATATTGGTCCTTCACTAGGTTATAAGGAGGAGCTGGGGAGGAGCTATGCATTG
TGTGACAATGCACAGATTAGTTTGCAACATCTTTTTTAGAGGCCCATATAAATCACTCTTGCTTTCCGGAAGAATACACAACA
TAAAAGCAGGCGGTGATCGCATTTTTGATTTCAAATCCTGTATTATTCGGTTAACCCTTTCTGCAGCAGATAAGCCTTGATCAC

TGAGTTGCTCGCATAGTATGAGTTCGGGGCGTGAGATTTTCATTCCCTTGGTGCTGCAGACTATCTCCAGATCATTTGCAATCT
TTTCCACAATATAGCGAGGCTTTTTGTTATCAAAACTTATGACTACCCAGGATTCAATGTCGACAGCAGATGCCATTGTTTTAT
TGTTGAAGTTCCACCTCCCATTCCGTGGTGTCTCTGTTCTAGACTCCCCAAACATCAAAGTGGGTGCGTCTAGTAGCCTTGCAG
GGATCCTCCGCATTCTAGAATCAATTTTTATGTTAAAATCCTTTATGAGCTCATTTGAAGAATAATCACTAACATCCATAGCCT
GGCATAAATCTTTTGCACAACATCCCTTCTTTTCTCTGGGGCTTGGCGTCCTTGGTCAATCATTTTCCGTCTCTGGCCACTTGAT
AATGCTTTTGTGTATCTCTGCCCTCCTTTTATATTGCAAAGCTCTACTGGAATAAAAGTGGGCTTCCTGGGTCTTCCAGCTGCT
ATGCATGGCCAGTTTGGATACTTAAGAGTTATATTGTACTGCATTTTATAGTACTGCTGTATCGTCAATTCCGTCTCTACCAAT
TGCCCGGATTCGTCCTTTTGCTTTTTTATGAACTTCTGCTGGGAGCAAGGAACATCACTAAAGCCGATAATTCTATGCACCATC
TGCGTATGTGGAGTCTCAACCTTAACTCCTTTCAAAACTCTCTTCACCTTGTCCCAATCAATGCAATTAGGATTACGGGGCTGA
AAACGTTCCCTAATAAAATCTTTGACGCTGGAATACTTGATGACTGTCGTTGTAGCAATGTCAAGGTTTAGTGACAACCCCTG
AAGAGAAGGCCGGAAACTAACATGATACCCTTGACATGTCTGAACTCCTTCCCCAATATCAGCAACTCCAAACAATGGGCTG
AAGTAGTTGTCCTTGATTAGGAGATAGCCCTTGGAGGATGCATGCTGCTTCAGAACAATATCCAGCACCCGCAGAGCATCTTG
GGCCCTGTCATTCTTTTGTCCATTCAAAATTGCATTGATGGCTCCCATCTTCACAGTAGCTGCATACTCAATTTTAACCTTAAA
TTCTCTAGTGTTCCTGGATGACATTTTGCGCCGCTTTGTTGAATCATCATCTGCTGAGTTTGCTTTGTTCATCGCAGACTCCTCC
AACAGGACTGTAAACTCCTGGGTTTGGAACTCGAGGGCACCTGGCGTGAATAGACTCTTCTCTCCATCATAGGCAAACCGCTT
GCCTCCTAGTTCTTTTGCTCCGTATACTTGTTGAAGCTTGCTGATAATGGACCGACAAAGGTTTTTATTTGCGATACCATCGTT
TCCACCAGGCGCTGAGCCCCCATCTGAGATACTCACATCGTAGTGGCAAACGTCCTGCGATCCGGAGTAGTTGACCATAAAGT
GATTGCAGAGCAGCTCAATGGGCCTGCCCGCTTTTCCTAATCCGGGTCTTTGCATTTGTGAAGGCGGCTTCATTGCAGGTACG
TCCGCCACATCTGTTGGGTTCGTTGATGGCTCTCCTGTGAGTGCCGCATCTCCTGCTGCTTCTGCCACCGCTGCGGGGTCCCCC
ATGGCTCCAAGCTTGAGAATGGCACTTGTGCAGAAGAACTGAAGAACTAAGGGCTGCGCAGTGCGAATTAAGAGAAAGGGC
ACAACTGAGAAAGGGAGGGTGAAAAGAGAGAAGGTAAAAGAGGGGGGGAAAAGAGCGAGAGAGAGAGAGAACAAGAAGC
TGCTTAGAGAGAGAGAGAGAGAG

**ARGONAUTE 10 (*AGO10*):**

CATAATTTTTGGAGCTGATGTCACACATCCACATCCAGGGGAGTTCATCAATTGCTTCTGTGAGTTTTGTAGGAATGTG
AGTTACTTTATAGAGTTTTATGATTAGTATATGCATATATTGGTGAACTAGCTTATGGCTTTCTCTAACTCAATCAAATGTCTC
TTTGCTACAAGGTTGTTGCATCACAGGATTGGCCTAAAGTGACCAAATATCCGGAATTGTTCTGTGCACAAGAACATCGACAA
GAGCTCATCCAAGATTTGTTCAGAAGCTGGGTAGATCCTGTGAGAGGGCCTCAGACGGGTGGTATGAGCAAGTTTATATCCTG
CTTTTGGTTCTAATTAATGATCTTTTAAGAATGATATTGGGTGTCATTAAGTGTCATTTATTTGTAGGATGTAATTGTCAGGCA
GCTTTTAATCTCTTTTCGACGAGCAACAGGTCAAAAGCCTTTGGGAATTATATTCTACAGGTAAGGCATGAGGCTGTAGTTAA
TTAAATATGGCTTTTCTTAATTAAGACGGCAGCGCTGGACATATTTGTGGGAGGGATGGTGCAAGTGAGAGCCAGTTCTCTCA
AGTGCTGCTTTATGAAGTGGATGCTATTCGCAAGGCTTGTGCATCTCTGGAACCGGACTATCAGCCTCCTATCACATTTGTAGT
TGTCCAGAAGC

**SERRATE (*SE*):**

GTTTAGGGTTGTCTAATGGTCCCCGTTTCTGCTTCCCGTACCTTTGGGCTTACACACACACACGATATGATCGTGTGCT
TCAATGGTTTTCACCTGAGACAGGTTGCAGTTTAGATAGGGCTCTTACGCAGCTGCATATTCATCAGCGTCTTGGTGGATTAG
GGTTAGGGTTTCGCTGGCTTGACAGGAAATTGCCCAACAGAATTTCTCCATTTGGGTCGTCAGAATACCTAGGGTTTGAGTTT
GTCTGTCCATTTTGCCGTGCTTTTAGCTCTTGGTATGGCAGATGTATTGGATATTCCCCCGGAGCCTGCCCTTTTGCGCGACGA
TAAGCGCAGAGACAGGGTACCTGACCGCAAAGAAAGATCCGTTGATGATCGCAGACGCGATAGGGATCCCAGAGACCGGCG
GGATGAGGGGAATGAACGCGATGAGCGGCTGTCACGGCGTGATCATTTTGACAGGCGAAGTGACGATGAAAGACCGGGGAG
TGCCATGACTCGAGATAGGGACCGGGATTACAAGAGGCGCAGAACTCCGAGTCCTTCACCCCCTCCTCTCTATGGCCATAGA
GATCGAGAGCGCAGGATTTCTCCATTACGCCGCTCCCCTTCTTCTTACAAGCGGTCCAGGAGGGAGGATGACTATGATGGAGG
CCGGCGAGGTTCTCCTAGAATGGGCATGGATGACAGACGGGATAGAAGGATGGGAGGTGGTGGTGGTCGAAGTAATAGCTA
TAGCGGTGATGAAAGAAGCTATGGCAGACATCATGGGTTTCGTCCTGAGTTTGGTAGGGGTGGCTTTGCTGATGGACCTTTCT
CCTATGATGTTGGTCCTCGTAGAGAGGGACTTATGACGTACAAGCAGTTCATCACGGAGTTAGAGGATGATATCATTCCCACA
GAGGCTGAACGGAGGTACACCGAGTACAGAAACGAATTTATCTCGACACAAAAGAAGGCTTATTTTGAGCAGAACAAGCAG
GAGGATTGGTTGCGGGATAAATACGATCCATCACGTCTGGAAGCAGTCATTCAGAGGAGAAATGAAGCATGCAAAACTGCCG
CTAAGGAGTTCATTTTGGAGTTGGAATCTGGCTCTCTGGACATAGGCCCTAATGTTGTAGGACAGAATCCACAGGGTGCGCAA
GAAGTTTCCGAGGAAGAGACGGATGACAGGAGGAGGAATGGGCGTGGTTCTGCCAAAGAGCAGGAGTTTGATGCTCCCAAA
GCTCCTGCCATCTTTTGCGAACCAAGAAGGATAGAGAAAGATGTTGAACAGGCACGTGCTCTTGTCAGGAAACTTGATGGTG
AAAAGGGCATTGAAAGGAATATACTTTCAACCTCAGAAATGGATAAATCTGATGGGGAGAGAAGTGGTGGTAAGATGAATA
TTGTCGTTGTGAGAGGTGCTAATCATGTTCAGGGCTATGAAGGTGTGGAGCTCTTAGATGTGGTGATAACCTATCTATGGCGT
GTTCATTATGTTGATTATTATGGTTTCAAAGAGTACAAAGAACAGCCCAAGGTAATGCGGCATGTTAGAGGAGACGGCAAAG
TGAATGAAGATATGGCCTCATCTGTTGAATGGGAGAAGAAGGTTGATGGCACTTGGCAAGGAAGAATACAGGGACAGGATCT
TTTAGAGCTCATGTTGGGTAAGGAAAGAATGGAGTCTGCCACATTGCTAGCTTTGGATCCCCTCATTCGAAAGATAAAGGATG
AGAAATATGGGTGGAAGTATGGTTGTGGAGCAAAGAATTGCACTAAGCTTTTCCATGGGCCTGAATTTGTTCAAAAGCACCT
GAAGTTGAAGCACCCTGAGCTTATTCAAGATGTGGCAGTGAAAGTTTATGAAGAGCTATATTTTGAGAACTATATGAGTGATG
CTGATGCTCCTGGATCCACTCCAGTTATGGCTTCCCAGAAGGATAGACCTCGTAGGCCTCCTAGACCTAGTGCGCCCGATGAG
CCTGCACGGATAAGTGCTGGTTTGCCTTTGCCAGCTCCCAGCAGGGGAGGTTCTCGGGAGGCGGATAGAGGGGGCCGAGGTG
CCAGAGAGAGTGATAAAGTTGATAAACCGGAGAAGGTTCAAGAGGATGAGCAGTTTGAGCAACGGAACAATGATCAATCCC
CTTCGCAGGATTACCAGCAGTCTGGTGGGCCTTATGATTCTGCTGGGCCCTTCGAAGGAGGAAGAGGTGACACACAGATGTTC
GACCCATTCAGTGGACCTGGTGGTATGCGAGGGCCTCCTTTCGGTGCAGATATGGGCATGCCTCCCGTGTTGATGCCTGTTCC
AGGCGCTGGACCTCTTGGGCCTTTCGTTCCTGCTCCTCCAGAGGTTGCCATGCGTTTGTGGAGAGAGCAAGGTGGTGCAGGAC
CATTTCATCCAGCAGGAGTTTATGATGGCCCATTTGATAGTGAAGGAGGCAACTCTCGTGGCAACAGGAAACGGGCTGGTCC
ATCAGGAGGTGGTAGAATGGGGGCAGGTTTAATTGACACCCCACCTTTGCCCTTACCTTTACCAAACATGCGACCTGATGCCC
GTCGTCCTCTTCGCAGTTATCGGGATCTGGATGCACCGGAAGACGAAGTTACAGTTATTGACTACAGAAGTCTATAAAAGGCT
CTGTATGCCATGTGTATTCTTGTCTAAATCAGCAATGACGACGTGGAGTGTAGTGAAAGACCTTCACGTTGTCAACGCACTTT
TGTCGTTTTTAAATTGCAGCCATGTAAGATTGTGATCAAGGGCCAATTTGAAAGCCCACTCAGCTGTGCCTAAGTAGGATACA
TGCATTGAGGTACCCACGTAAGCAATCAAAGTGAAAACTTGAAGACAATTTATGGATTGTCCTCAACTTCTGTTTGTGATGCC
AGTCTTCGTGATGTTGGATGCTTCCGCTCATACTAATTTTGCTACTTTTGGGATGGGAATATTGAATCAGTGAAAAATTGTTTC
ATGTTAAA

**HIGH EXPRESSION OF OSMOTICALLY RESPONSIVE GENES 1 (*HOS1*):**

AAAAAAAACTCGTTTTTTGCTCCACATATTTTACAACCTACGAAATTTTGTTTAGTACTAAGGTCTTATATTCGAAATC
TGACTGTTTTTGTCCTCTTAATCAAACCACCAGGCAAAGAGCCCAAACATTGCAGTGCAAGTTTGATTCTTGCTACAGATAAG
TTACCCACTCAATGACTACTTCATCAGAATCCGAGAATTCCTGCATGTCCCAATTGCACTCTAGTACAACTCATGTAGCATATA

CACATGCAATTATCCAGTTTTATTAACCTTGCAACAAAAACCTTGCTTCAAGCATTTCACTACTCAAGTAGTCCTCACTGTGCT
TGATCATTTGGCTTGCAGAGGCTTAAGTTCCAGTTTACTTTTTTGAAAGCAAATGTGATCCAAGTGTCTCTGACTTGGTACAAT
GTCCCCAGAAAAAAACCAATTTAAAGCAAGCGTACAACGCAATACAAGACAAGGCCTTCAATAATAAGCTCTCTTTGTAGTCC
TTTGAGTTTCAAAACCAGTGATACTGCACCAAGGCTTAAACTCCAGAACCCTGAAAAATAGCTGATCATCCCTAGGATGTATT
AGTTTTAAGGCGGCCGGTACAAAGCCTTTGCAAAGAGTGCTAATAGTTAACAACTGAACAGTGTCAGCCTAAATGCAGCAAA
ATTGCTTCCTAGCTTTTCTAAACTCGAATCTTCCTGGAGCAGGAAGGATCCAAAACATCTGTATCACTGAAGTACAAACAAAA
CTGAACAGTTATGTCTAGGGGTGTAACAGCCACTCTTCATGCAGCTGCGAAAAGGTAATCTTCGTCACCAAGCCGTAGATCCC
TGGATCTTTTCAATGGGAGTAAGCACAATGGCAAGGCAGCTTTTCAGAACCAGATTGATCAGAAGGGTTCTGCTCTACATGAT
TAACCCCGGCACCAGATTCAAGTGTTCAGAATTAATGCTCTGTTAGTTTAGAACAGCTCTTGCTCATGACCTGATGGCAGCTG
ACTGCAACAGCCAACCAAAATGTGCAGGCGTACCATTGCACCACTGTGCCAAGTTTTCATCCAACCTGGCAGAACAGTGAAA
GACCTTGCAGTGCATCACTGCATTGACCTGTCACAGTTAAAAGAAAATTAACTTATCATCAAACCCACAATCGAGTAAAGCCT
TCAGCACAGGTGGCATAACAACACTGCAAGAACCAGTAGAACCTTCCAGTTCAAGCAACTTGAACAGAAGCGCAAACCGATG
GTCAAGCCCTGGCATCTGCCGCAGACGTAGCTTCTTTCAACCCCAAGAAGCCCATCAAACTTGTTTTGACAGGTCGGGTTGCC
GTCAACTCCAAAATAAAATTAAGAGGCGTTGCTACAATGCATGGCTTCTGAACGATCTACTCCCACTACTTGGCCTCCCAATA
GCAGCTGAGGCTTTCGCCCTCCCACCAGATACAGAGGGAGGGGGTAAATCCACGCCATCGTCTTCTGTTCGCCATCTCAGAAC
ATTTTTCCTCCTCGAATATGAGTTCTGGATATCGGCTCCTGATGGCAACAATCCATTTTGCTGCTCACTTCCTGACATTTTCAA
GCTGTTTGCAAATACAACATCCCCTGCAGTTGAATCTTGCAAGCAGACATCCAGTTGATCATGGGCTTTTCTCTTTCCATTCGG
GCTTGAATGTATCATCTGTCGCCTGGGCCTATATTTGGCCTCTCTAATTGTCAAATCAAATGTCTTATCATTTCTTGGTGACTCT
TTCTGTTCCAAGGCTCCATTTTCAGAACACCTGAACAAGGGAGCAGCTGTTCCTGGACCATTATCAGTAAAAGCGGACGCCCC
ATTCTGCAATCTCTCATTCAAACAGGTCCTACCATCAGTTTCGTGGACATATGATTTTGATTTGTTGAAAGATGGAACAGGTTT
AGCAGAACCATCACGCTCCTGAATATGAGCATTTGAAGTGGAAGAAAAAATATTCGGGCTCTCCTCATGAATATAGCCCTTCA
TATCTTGAGCTAATACTGCTCGCAGATTCCTGGTTGCTAAGGAAGTGCCCTCGCTCTCTGCATTGTAAATCAGTTTCTTGCCTA
GAATAGGACTCAAGAGTCCTTCTGAATACTCTATTTGAGGTGTGATGCCACTGGTGGTGGCAAGATCCAGCTGAGACAAGCC
ATCCGGTACCGCAGATGTAGTAATGAATGGCTTTCCGTGAAGTATAGATGGTTGACGGTAGTTGCCCCATGCTAGTGGATGGC
TCATCAAATCAGGTGAAGTACTGTGGGATAAGGGCAGAGAGTCGCTCTTCACATCATAAGAGATAAATGCTTGGTTTGTTGCA
CCGCGTAACTCAGCTTGTACAGCTGGGAACATTAACTTTAAATTGGGTGCTTCAGAAGCAGCTTCTTTTTTCCTTTCTGTTGAA
GGAAGCAATAGATGATCAAGGCTTCCAGTTCGTAACTGATCCTGTTGAATCTTTGGTAGCAACTCAACACCCTTCTCCACCAA
AGTAGTTCTTTGTGCGCATACAGTCTTCACAATTTCACTTATTTCCACATCTCTACATTGATCCAAAAATGACCTTTCTATTGCG
GAGAGCCTCCTATGCACCACATACGCCTCGCGATTGCGGCACCTCATAATGTAAAACATCACCATAAGACTGCCTGCTGTAGA
ACATGGGTTTATGGTTGCTTGCTCCAAAAGACATGTCAAAAGAATCTTCTCCTCATCCACGTGCCAAGGAAGCTCTATCATCT
TAGCTACCAAGTTTTTCCTTATGCTTAACCAGCAAATTTCTCCCACCAACAGTTCCATTTCTGATGACCAATCTCTAGAATTGT
CTTCACTGCCAAATTTTGATTTGGAAATTACATCCCTTTGAGTTTGTCCTTTAACTCTCGATATATATGAGCTCTGATATAGAT
ATGCATCTGTCAAAAGCCCACATTCTAGCCTCACCCTCACAGCAGTTACAGCATCTTGAAGGGCAGGAATTTCATCTCGGTCT
ACAAATCCTGAAAGCCTTAGGCTCCCATCTCCATTGCTGCAATGCAAAACATCCAAAGCCATATGTGGTTTTTGCCTCTCCAG
TAGGACACGAGCAACTTTAGGATGTATATTACAAGAAACGACCTCAGGAAGAAGCTGACATGCCTCCTCCAAAGCCAAGTCA
TCCTTCTCATCCAGCAAAAAGAAAACCCATGATTCTAGCATTGCATGCCTTGATATGTCAAATGTAAGAATATAATCGTCCAA
TACCTGCTTCCAATCCTCACTGTGGTTTGGCCAATGTCGGTCAAAAAGATAGTACAAAAAAATTGCTTTTTTGGCCAGAAAGA
GGTCTGCATTTGCTTCCAAAAAGAACAAGTCTACAGCTCCGCGCACACTTTCTGGAGGGTATACTGCTGCTGAAACGCGTATG
GCAGACTGCCCTCCAATCCCCTCTCTCCTTCTAAAAGGTGTAGAAGCTGTCGTCTGCCTCAAGGACCCTAATTCTAGGTTTTCC
CAGAGATATGCATTCTCATTACCTTTTCCGAACCCTACGTTTGACATTGCATCCTCAATAAACAAAGTTGCTGAAGTGTAAGA
AGTCACGTCTGATGCATTCCTTGTATGATCAGGCCACACTCTGCTGTTGGCTGCCAACTTACGTCCTTTTACAGCAATTCTCCA
ATCACTCGGTGAGGGATGAGATGACTGTAGATCCTCAAGAAACTGATGCCTGGCACACCAGCACATCACATCTAGATGCTGA
GATACTCTCCGGGTACCCTCTATAAGTTGTTGAAGCTCAAAAATTGGTGTGGAAGTTGCATCTGATATGGGACCTTCCAGAGA
TTCCAAAATCTGAGACAACCCAAGAAGCTTTGAGGAAAATCTCTCCATATCACCCTCTCTGCTTCTCATCTGTGTGGTTCCTAA
ACAATATATGTCATGTAAGCTCCGGGTGATGTGTAGAAATGTTTTTTTACACCAGTCTTTGACCACTGTTCCATCCAGGAGCAT
GGAGACAATTGCATGGCTAGAGACAGCCCTGTCATCCATACAGACATCTGAAACATAATGACAGATCACTGAGACCAATTTG
TACTCTAAAGCAATATCAAAAAAGAACCAGAGGCGCTGAACATCAGCATTTAGGTGTCCATCCCTCTCCCTAATATCTTCTGT
CCTAGAGCTTATGAGACCAGCCTCACAGCATTCTTCATAAAGGCGTGCTCTCATTGAAGGTCCATGCCTCCTGATTGGTGCTCT
GCAAATTGGGCAAACATCACATCTCTGGCCACATTCTGCACATAATGAGGCATGGTTACAGGAAGTCAAAGAGTGCTCAACA
ATTCTTCCACAGCTTCTCAGGTCCCTTGTTGCTCTGCATTTTTCAACGATCGCTTCACTGCAAAGCTCCCTGGGCTCAAGGCTG
ACAAGCCGTGACAAAACATCCTGGCATTTGTCTGTGATTCTAGGGTAGAGACGATCTCCATCGTTTTTCTGAAGCGACCACGC
GGCTCCCGCCATGCAAGCTTTCCTGCTCTCACTGCAGAAGCTGTAGGCGTATATGCACGAGGGCGTCCGTCTATATGTGCGCG
TGTGTTCGTACGTGCGCGTGTGTGTGCGCGCGTTCCAAATGCGAAGCACGTAAAAGCTTCTCGTTGCCCCCTTCAGTATACAA
AACACAGTGCGGTGGTGAACCAAAAGACGCCTATGTCAATCTCCCTGTAGTCTCCAAACACTTTAGACTCG

**ABA DEFICIENT 3 (*ABA3*):**

AGAAGTTTTCCCAAGCTATGGGAGCTTCGTGTCTGCTCGAAACCCTTAAAAGCATTCTGCTAACAGTAGCTTTCTACCT
GATGAGCTCCTTCTCCTCCTCCTTCGCAGAAGATTTAAGTGCAGAGGAACATGAAGTCCACTATGGAGACCATGGAGGTCCTT
CTGCAAGACAGAAAGGCAAAGATGAAGCCAGCATAGAAGTTCCGTGCCATCATGGAAAGCTTTCTGAAGAAGCTGCAGATC
ATGAAAGCTTCATGGAGCATGGAGCCAGGTGCATTACAGAACATGGAGCCACTTGCCTTACAGAAGAAGATTCACATACAGG
CGTTGCAGAACACGGAGAAGAATTTTTTTTCGATGCAGATGATGGAGAGCTGTCTTCAATACATGGAAACTTTCTTTGTAAAG
AGAGTGTAGGCACAAAACCTACCTTTGCAGATGGAGGCGTCTTGTCAGAAGACCAAGCAAGCCTTGGAAAACTCTCCTTTGC
GGAAGATGATGCAGCGCATAATGAAGACCCTCGCAAGCAGGAGTTCTTTTCACTGTATAAGGAGCAATACGGCTACGCTAAA
AGTGCCTTATCTATTGACAAACTGCGCTCCGATGAGTTTCCTCACCTTAAAGGTGATATATATATTTGGATCATGCTGGGGCAACT
CTTTATTCAAGATGGCAAATCCGCTCCAATTTGGAGGACTTATGTTCAAATTTATATGGGAATCCCCACAGCCAAAGTGGGTG
CAGCAACATTTCGTCAAACATGGTTGCAAATGCTCGTGAAGAGGTTCTGAAGTTCTTCAATGTGACAAGAACAGAGTACAAG
TGTGTGTTCACTGCTGGAGCAACAGCAGCTCTTAAACTAGTTGGAGAATGTTTTCCTTGGACCACCAAGAGCAGCTTTTGGTA
CACCATGGAAAACCATAATAGTGTGCTTGGAATAAGAGAATATGCTCTTGCAGCAGGATCCTCAGCCTATGCCATAGAGGTA
GAAGATTTTGGTCAGCAACAGAGTTTCAAATTAAGGGCACGCCAATCGCGCACAATAAGCTCTGCTTGCAAGTTAGAGGAAG
GCAAGGAGGACAAGAACACCTTGCTCATAGTTAGTTTGACGAGGCAGAGAATTGTTACTGACTTTTGAGCCATGTGTTCTGCA
ACAGGACTCACATTTAATTTGTTTGCATTTCCTTCGGAGTGCAACTTTTCTGGGAAAAAGTTTAACCTTGATTCAGTCAAGCTC
ATACAAGATGGAGCTCATGCAAAGGATTCTGTGAACAGGTGGATGGTGTTAATAGATGCTGCAAAGGGTTGTGCAACCTCTC
GGCTTGATCTTTCAAAGTACCCAGCAGATTTTGTAGCCATTTCTTTCTACAAAATTTTTGGGTATCCTACCGGTTTAGGGGCGC
TTCTCTTACGCAATGAATCTTCTAGGATATTAGAGAAGAGATATTTTGGGGGAGGTACTGTTGCAGTTTCTGTAGCAGATGTA
GATTTTGTTCAGAAGAGAGAGAATATTGAGCAGTGGTTGGAGGATGGAACTGTTCCTTTTCTTGGGATTGCTGCTTTACATAG

AGGTTTTTCGATTATTAATAGGCTTGGCATCTCTAACATTGGAAGGCATGTTGGAAGCCTTGCGAAATTCACAGCTGCTCAAC
TATCAGGGCTTAAGCACAAGAATGGGAGCAACGTCTGTGTGCTATATGGCAATCATGATTCTGTGCATTATTGGGGGGATAAC
TGCAGTCAGGGCCCAATTGTGACATTCAATTTAAAGCGTTTGGATGGGGCATGGGTCGGCCATCGAGAGGTTGAAGAACTTG
CATCTTTGAATGGAATCCATCTCCGGACAGGCTGTTTTTGCAATCCTGGTGCATGTTCAAAGTACTTAGACTTATCAGAGTTAG
AGATTCGAGCTAACCATGAGGCTGGCCATGTGTGTTGGGATGATCACGACGTTATTGATGGTAAGCCAACTGGAGCAGTTCG
AGTGTCTTTTGGCTATATGTCTACCATTGAGGACTCTTTGGCTTTACTTGATTTCATATGCAACTACTTTATGGAAAATCATAA
CTCTTATCTTGAGAATCAACCAACCCTCCGTGAGCTTGCGAGCTTGGCCTCAAATGATTATTGCAAATCAGATTGCAGTATGT
CCTTGGAATCAATTACTATTTATCCTATCAAATCTTGTGGTGGTTTTGCTGTGGATTCGTGGCCACTCGCGGATTCTGGCCTTCT
TTACGATCGAGAATGGTTGGTGATGAGCTCTGCAGGTTTTGTGTTAACACAAAAGAAGTGCCCCATGATGTGCCTGTTAGGGA
CTTACATAGAACAATCAACAAATACACTTCAGATAACGTCTCCAAACATGAAAACAAGACTAGAAATTCCTTTAGTATCTGCA
CCTCACCAGGAAGCTGTAGTGAGATTTGATTTGTGTGGAGAAGGGTCCATTGGTAGATCATATGAAAAGGAGGTTGCAGACT
GGTTCACAGAAGCTTTGGGTACGGCCTGCACACTGGTTAAAAAGCAGCCAAGAAGCAGGCATCTGCGGTTAAGAGGGGGGAT
ATGTGAATCCCAGGCATGCAAGGATGCAAGAGAACTCAGCTATGCAAATGAAGGTCAGTTTCTGCTAGTTTTACGGGCAAGC
GTTGAGGATATAAATCGGCGAGCAATCTCATCTTCACAGCGCATCCAATCCAAACAGAAAAAGAAAGCTCACCATTCATCAA
TGCAGGTAGACGCCATGCGTTTTAGGCCAAACTTCGTGGTATCTGGAGGGTCTGCTTTTGAAGAAGACAATTGGCAATCCCTT
GACATTGGAGGTCATAAGTTTATGGTTTTAGGGGGTTGCAACAGATGCCAAATGGTCAATATAGATCAATCAACAGGAAGGT
CGCAGGAAGGAAGCAACCCATTACTGACACTTGCCACCTACAGGCGGTCCAAGGGCAAAATACTGTTTGGGCTTCTTTTGGC
GCACAATCCTTCACAGCAGCATGAAATAGAACAAATGGAAAGAAATGGTGCCAAAAGTCGACATGAAAGAATCATCAAAAC
AGGTTGCAGAGTTCATGCAGAAAGGAATCGTACGGCCTAATTTAAAGAAGCTCATTAGAAGCTCGGACTGAACAGCTTCAAC
GACTTAAGGCCACAGGCATACACTACTGTTTTGGTAGTTTTGCAGGATTTTGGGCTTCATCACTCCTGCAAATTTGTGGCTGGA
GGCTTGCTTTGAAGCAATGACATGTAATTAGATCACTTCAAATTTCAGGGATCAGGAACTGGAGAGCTGGCATTGAATCTTAT
TTGATCACTTCAAAAGTTCAAGGATCAGGAACAGGAGAGCTGCGTTGTTCACCGTCATGAAATCCCAACATGCCAGATGAGT
GAATGTCAGCCATCCCATTTCAGTACTACAGTGACATCATCATGTGAACTTACACAGAATAAAGATCCTTTTTTTTTCATCTTC
CATTCTAGCGGTTCGACGAGGTTCATTATTGGTTCTCTTGAAGCAGACAGAAACTATCAGTGTGTATAATATGTATATACATTC
GCACATAGTCATGTATAAGCAATGGGTTTTCACGTACCATCAGACATTGACAGATCAAATAGTGGCCTGTTAGATAGTCCAAC
ATGAACAATTTTGAAAGAGAAATGCTCAGTGCTGGTTACGTATGAAGTTAATTTTGAAATTCATCCTGTTGTATTCATTTTGC
CAGCATTGTGTTTATGCTATTTGAAGTTGTTAGTCCTCACCATACTAATGACATAGAAG

**PYR1-LIKE 5 (*PYL5*):**

TGAGGACAACTGTTCTCTTATCCTCACAAAGGGCCCCATAGCGAGTCGTCTTCTCCATTCGGGGGGGATGGATGTGAA
GCTATCGATGATGCATCTGTGTGTGGCCGCATGGTCACTATGGGGTAGCTTCTCTGCTCTTCTTTCCTGAAGCCCCTCTCCCTG
CCTCTCTCCCTGCCTTCCATTCCACAATTTTGCCCCTCTTTGCCTCATCATTTCGTTCGCCCATCTCCTCCTCCTCCTCTGTCTGT
TCTTCATTTGGTCGCCCAGATCAATGGCAGTGCATCAGCTGTGGAGGTTCCCAAGGGGAGCAGATGATCTGGAATCTATGCTA
AATGAAAGGGAGGCAGTGACTCCGGGTTTGGAGCAGTTAAACAGAGATGAGCAAGCCAACCTACGAGATCTGATTTTCAGGT
ATCACACCCACTTCATAGCTTCAGGGCAGAGCTGCTCCATGGTGGCACAGAGGATAAAGGCCCCCGTGGAGGCGGTCTGGTC
TGTCGTGCGCCGCTTCGACAACCCGCAAGCATATAAGCACTTCATTAGCAGCTGCTTCATGCAAGGAGATGGCCAAGTGGGC
AGCACAAGAGAGGTGAGGGTGGTTTCGGGTCTCCCGGCTGCTAGTAGTACTGAGAGGCTCGAAGTCCTGGACGAGGAAAGGC
ATGTTTTAAGCTTCAAAGTTCTCGGAGGGCAACATCGCCTCAGAAACTATCACTCCACTACTACCCTCCATAGGTATGTGATT
GATGGGTGCCCTTCCACTATTGTGATAGAATCTTATGTAGTTGATGTGCCAGAAGGAAATACCAGAGAAGAGACCAGGGTGT
TTGCGGACACCATTGTGAAGTCCAACTTGCAATCCCTCGCCCGAACCTCAGAGCTCCTTGTTCGTGACGCACGGAGGCTGGCC
TTTCTGGCGTAGATTGGCTCAATTATGGTCTCTTTTGGCGAGCAAAATGGGAGGACCATGAAGGCCACAACACCTTTGCCTAG
TTTTATCTCATGCAAATGTGTGAATTCATGGTGGAAAGATTACCGATCTGATTTTTTTTAGTTCCAGATGGCAATGTTCGACTC
CCTCTTTTGCCAAAGCTTGGTGATCTTATGAAGGCAGAGCAAATTCGATCACAAGCTAATCACCAAAATTTACACTTGCAGAA
AGCATGTCTGATGTCTCCAGTGCACGCTACTTACCATATCGTCACACCATCATTGGAGTGATGCTTTTACAGTGTTCAGAAGTT
CTCTTCCAGCCGCCCCTTTTTCATTGGTTTATTGTATGTCACCATTCCAGCAGTGAATGATTGCGTGTTAGTGTATACAGTACC
AACCTGTACAGCTGCGTGGGTTTTAGTCTTAGCATCTTTGATGTGTATTATGTCTGGGTATTGTCTACTATGGATAAAGCATGG
ATGCTATGGTCAGGAACAATGGTCTTCCATAAGGTGAGTGTGTACTTTAAATTTGCAAGCTTTGTCTTCTTTGCAGTGGAATGG
ATTCTTGTGGTAAAGTGGTACTTCGCTGCATTGTCCTTTCTTATCAGCAAGTTGCATATACTGTTTCGCTGCTTTGTCCTCAGAA
AATGCTGCGTAAATGGCCCATCAGTATTGCCTGGAGTTGCTTAGTTAAGTCATGTAAATTGTCAATGACATGATACCCTCTGG
GCGAGTTGTTTTTGGAGCCTCCTATGTCCGGGACATTTGATTGTCGATGTCAGTGGTTTTTCATGAGCTAAGAGCTATTTTAAA
ATTGTAATATGAGATAATTCAGAGGTAGTTTTCA

**ULTRAVIOLET HYPERSENSITIVE 1 (*UVH1*):**

TCTCTCTCTCTCTCTCTCCTTGCTTGCCTGCATTGTTTGAATGACACGGCAACGAGCGGGGGAGCGCGATGCTGCCAAT
ACACGAACACATAGTAGGGGAGCTGGTGCAGGAGGATGGCATAGTGGTGATAGGGGCGGGGCTTGGTTTAGCCAAGGTCCTC
GCCTCCGTGCTGCGCCTGCACTCCCCTTCTGAAGGTGTTGTCCTCATTCTCTCTGCCTCTGATGGACAGCGTGCCGCCATCCAA
GAAGAGCTTCTTGAGCAAGACCCATCTGTTCCCATCATCTCAGAAATCAATAATGAGACATACTCTGTCTCTGATCGTGTTGC
TTTCTACACGCATGGCGGCACTGCTTTCATCACCTCCCGCATCCTAATCGTGGACCTCCTCAATGAACGTATTCCCCTTGCCCG
CCTCTCTGGCCTCATCTTCCTCAATGCCCATCGCCTCTCCGACGCATGTACAGAGTCTTTCATTGCCCGCCTCTATCGTGCAGG
CAACCGAAAGGGCTTTGTGCGTGCATTTTCTGACAAGCCCCAAGCTGTGATTAGTGGATTCTCGAAAACAGAGCGAATTATGA
GGAGTTTGTATGTGCGCAGATTGCATCTCTGGCCTCGATTTCAGATGTCAATTGCGTCGGCGTTGGAAGAACATCCCCCACAA
GTTATCGATTTGAGGGTTCCTCTGACATCTTCTATGCATGGCATAAATGCAGCAATTCTTGAGGTCATGGATGCATGCCTGAA
AGAGCTCAGAAAAACCAATAAGCTTGATGTTGATGATCTTACTCTTGAAAATGGGCTCTTCAAATCATTTGATGAAATTGTCA
GGCGGCAGCTTGACCCTATATGGCATACTCTTAGCAGAAAAATAAAGCAACTTGTCTCAGATTTGAAGACTCTGAGGAAACTT
GCTGATTACCTTGTGCGGTATGATGCGATCACATTCTTGAAGTATTTGGATACTCTACGTGCTTCTGAGGATGTGAGATCTGTT
TGGATTTTTGCTAACCCAGCACACAAGATATTTGAACTTGCCAAGCGTAGAGTTTTTCAAGTCATTAGGACAGATACAGGCAA
TCCGATTGTGTCAGAGAAAGGCAACAAGTTTTTTCGTGGCCGGGGTAGACAAAGCAATGGGTCAGATATGGGTGCAAAAAGG
AGAAAAAACAGTGAAGGAACTGCATCAGAAGCAACCAATTCAGATGCCAATGCGAGCGTGGACCAAGCGGGAGGCGTGCAA
GATGTTGAAGTGGTTGTAATGGCTGAGGAGATGCCGAAATGGAAGGTGTTGCGTGATGTATTGGATGAGATCCAAGAGGAAC
TACAGCTTATTGATTCAAGTAAGCCAGAGTCCCCTACTTATTCAGATGAAGGGCTTGGAACTGTATTAATTGCATGCAAGGAT
GAACGCACCTGCCTCCAGTTGCAAGATTGCATTAACAAAGGACCCCAGAAGCTTATGAGTGAGGCCTGGGAAAAATATCTGT
TGGGGAAGGCTGAGTTACATGGTATGCGTACCCGGAAAAAAAGAAGCAATTTGGGTTCTAGAGGTGTTGGTGTTTTGAATGG
TAAGCTCTCTTCGAGAAATGTAAGCAATGAGGAAGGAAACACAGCAAGGCAGGAAGAGGTAGCTCTTCTAGCAGCAGCTGC
AGAGGTGTCATCTCGCGAGGAGGAAAGTGGGACAAGCAAAAGGGGGGGCAGAGGTCGAGGCAGGGGCAG

**ELECTRON-TRANSFER FLAVOPROTEIN:UBIQUINONE OXIDOREDUCTASE (*ETFQO*):**

TGGCAGAGCTGGTACCTAATTGTGCCTGACAAGAACATAAAAGTATGAAGCACGAAAGTAGGTCTAAAGCCTTTCT

TTTTGTAAAAAGAAACTTATTGCTTGTTCTTCCTATACTCATGTTGCTTGAGAGCACAAGGACTCTACGTTTATAAACAATCTA
CAAGAGGTATTCTTGGTTGACATCACACTTTACATGTAGAGTATACATTCTATACATATTTACAATGTAAATTGTTGAGGGAG
CAAACTCCATTGTATATGAGACACATATAAGTGTAAGCTATCCTTGACGAAAGGCCTTAAATATGTATGAACTCTGTCATGTT
ACAGTCCAAGATAATAAGCTATCCTATCCTTCAGGACAGGTCTCATTAAACATGACTACACTTTGTGAAGTTACATGACCGAA
TATCCAGGACCGCCACCACCTTCGGGCACAGTCCATTCAATGTTTTGGGAAGGATCCTTTATATCACATGCCTTGCAATGAAG
ACAGTTTTGAGCGCTGATGTGTAGCTTTAGTTCCCCATCATCATTGCTTACATATTCATAAACACGTGCAGGACAATAGCGTG
ACTCGAGCCCTGCAAATTTTCGAGCATTCAAGCTTTCTGGAATACATGGATCCTTCAAATGTAGATGTGCAGGCTGGTCATGA
TCGTGGTTGGTGTTGCTTCTGTAGAGAGAAGTCAGCAAATCAAATGAGATCTTTCCATCAGGCTTTGAATACATCTTCTCTTTA
TGGATGTATGCAACTTCTGTAGCTTCATGGTCAGGCTTCCCATGCTTCAAAGTCCAGGGCAATCGGCCTTGAGAGATATACCG
CTCAAACGCCGAAATTGCGAGTCCTGGTAACATGCCATACCGAAATGCAGGTCGAAAATTTCGTGCTTTGTACAGCTCCTCCC
ATACCCATGAGTTCTTCAAAGCACTCCAATATCCCTCCATATTTGGCTCAGACTCTGATTCTTCCAAAAATGTAGTAAAAGCA
GCTTCAGCGGCAAGCATTCCTGATTTCATTGCAGTATGTGTGCCTTTAATCTTTGGGACATTTAGAAAACCAGCAGAGCATCC
TATTAGTGCACCCCCTGGAAAAATGGGTTTCGGTATGGACTGGAATCCCCCTTCATTTAAAGTACGAGCTCCATATTGTATGG
CAGTACCACCTTCCAACAAAGATCTTATTGATGGATGTGTTTTGAATTTCTGGAATTCCTCGTAAATATTAAGAAATGGATTCT
TGTAATCAAGAGCCACAACAAGTCCCATTGCAACCTTACAATTATCCATGTGATACACAAAAGATCCACCATAGGTTTTGCTA
TCCAATGGCCACCCAATTGTGTGCAAAACTAAGCCAGGTTGGTGCTTAGTTTGCTCCACTTCCCAGACTTCCTTTACACCCAGT
GCATAAGTCTGATGTTGAGCCTGTGCAAGCTCCCTTAAACAGAACTTTTTGATAAGCTTCTGCGAGAGAGAACCTCGACAACC
TTCCCCAAATAATGTAAGCCGTCCTCTCAACTCAACACCTCGCTGAAAAGTTGACTTTCTGTCCCCATCTTTGGAAATGCCCAT
ATCGTTTGTTGCAATGCCAACAACTTTGCTTCCTCCTTCATCAAAAAGTATCTCACTACCAGCAAACCCTGGATAAATTTCAAC
TCCTAGGTCTTCTGCTTTTTTTGCGAGCCATCGCACAAGCTCACTCAAGCTTATTACAAAATTGCCATCGTTTTTAAAGGGAGA
TGGTAATGGCACAGCACGGTTAGCAGTAAGCCACCAAAACTTGTCTGTTGAAACAGGTACGCTTATTGGAGCTCCTTGATGTT
TCCAATCTGGTAGAAGCTCATCGAGTGCACGTGTTTCAAGGACATTGCCGGAAATAATGTGCGCTCCTATTTCGGCTCCTTTTT
CGATGATGCAGACAGAGACGTCTTTGCCATGCTGTTTGGATAGCTGTTTGAGTTTGATGGCTGCAGAGAGGCCGGAAGGGCCT
GCGCCCACAATCACCACATCAAAATCCATGCGCTCTCGTTCTCCCGGGACTGCCTGCGAGCAAAATCCTAGTGCCTGCAGGGG
CTGCCATGGCGACCTCTTTGAAAGCAAACCGCCGCGACCTCTTGTGGCAGAGGCAAGCCATCGTAGCTGCTGCATTGTTGTGG
GTGCGCGTCTGCGTGTGCGTGTGTGTGAGGTGTGTGTGTGAGAGTGTGTGTGTAGTGAGGCGGAAAGCTCAATCAATGTGCA
GGAGAGGGAATTGCAGCATCTTATGAGGAACGAGAAGAAGGCGCGCCATAGCCGACACAGAGCTGCAGGATCTCAAGCTTG
CTTAGGGGAGGGGTAAAGCGAAAAGAGAAGAGAAAATCTGAGAAAGTAGCACGCATTACAAAACCCAAAAGAAAAGGAGA
AGGAAGCTTTCTG

**Y-FAMILY DNA POLYMERASE H (*POLH*):**

GTTTTTTTGGAAGAAAAAAGGCCATGGAGAAAGAAACGCCTTCTTCTCCTTCTGTAATACTGCATGTGGACCTTGACT
GCTTCTACGCAGCGGTGGAGCAGGTGAGGCTTGGGATCCCTTCAGAAAGGCCTCTCGCTGTGCAGCAATGGGAGGGTCTCAT
TGCAGTCAATTATGCTGCCCGAGCTGCTGGAATTGTCCGCCATGACCGAGTCACAGACGCTTTGCACAAGTGTCCAGATTTGC
AGCTTGTGCATGTTGAAACTATTGGGAATGATACATGGGATGGTGCAGCTGCTAGACGTGACACCGCAAAGGTGTCGCTGGA
AAGGTACCGACAGGCGTCAAGGTGCGTATTTGATATATTTCATGAACATGCTGATATCTGTGAGCGGGCAAGCATTGATGAA
GCCTATTTGGATGTTACCAAGAAGGTTGAACTTTTCATGAAGAAAAATTGCAATTGGGACCAAGAATTAGAACGTTTGGCTGG
TAGTCAACGTCTGCTAGGGGGCTCGGTTGAAAAAGCTGGCATGGTTGTAGAGAATGGAGCAGTTGAGATTTGCGATCCTGTT
AGTAGGAGGCTACTGGCTGGAGCAATTATGGCTGAGCAGATACGTACTGATGTCAGATCAAAGTTGGGGTACACTTGCTCTG
TGGGAATTGCAACCAATAAACTTCTGGCCAAGATAGCATCAGCTAGGAATAAGCCAGATCGACAAACATTGATTCTACCAGG
GGCAGTTGCCGATTTGATGGATGCGCTTCCTTTAAAGAAAGTCAAACTTCTGGGCGGTAAAAACGGTGAGGAACTTTCTGAG
AAATGGAGTTGCAAGACAGCTGGGGATGCTCAGACAATACCTTTAAAAAGTCTATTAGCGTCCTTTGGCGACCGTTTGGGAA
ATTATGTGTATAGAGCAGTGCGAGGAATTCATGAGGACAAAGTACAAGAAAAGCAGACAACTAAGTCCATGCTCGCAGCCA
AGTCTTTTCAGGCTACAAACGATCTTTCTGTTATCAGGATGTGGCTTGGTGTCCTAGCTGAGGAGTTATCAATCAGGATGCTGC
GTGATGCTACAGAAAATCATCGTCAACCTCACAATTTGCAGCTATACTACCGATCAGGATTATCTAGGCAAAGCGGAGACCA
TTCAAAGTCATGTTCTATGCCGCATGCAGTGATACAACTACTCATGTCATCATGCTTCAGAACTCCAAGAGCATCTGCCAGTA
GGCAGCATGCTAGTGAAGCAACGGGATTATCTATTAAGTGTGACAAAGAATTCAGTGGTGAAGGAGTGGCAAAGTTTAAAGA
GGAGACCAAGTGTGATAAGTCGGAAGTACCTCAATGCATTGAAGGTGAGGCTGAAAGAGAAGTACTTCTATTCAAGGAAGA
ATCTGGCTTTCTCGACCCCGAGGATAAAAGATTATTCACGGAAGAGTGTGGGAGAAATATGAAATACAATACAGTTGAAGAT
CAAACAGCCAAGGAGTTGTCCAAATTGTTGCAGCAAGTTTCCCTTCAGCTATTTGAGCGCATTGAAGATGCGTTCCCATGCAC
TCGTTTAGCCATTGCTGCCAGCAGTTTTCATGATGCACCAGCACAGGGTTCTCAATCTATCCAGCATTTTTTTGGTAGTGCCTT
TCAAGGTTCCAAGGTGCCTTCTGTCAAGTCTTCGATGGAGGGACCTGTAGGGGAAAATAAACTTTTAAGTAAGCGTAAAGGTT
TGCTAAGATATATGGATAATGGCACTGGTCTCAAGAAGCCTTGTGCAACTTCTTTGAATGAGTTGAAAGGGCTATACTGTGTG
ACGGAAGGTATTGAGGAACTCTCGCACAATGAAAATCCGAGGGACAGCTCTTCAGTAGAGAAAGATAGCTGTAGAAGTTGG
GTTGTTGAACAGGCAAAAGAGTTATTAGAGGATGAGGTGGTTTTGCCGGCATTGGAGGTGCTGGATTCTGGTCGGTGCTGTGA
AGACCCAATGGATCAACAAGAAGTAAAGCCAAGGGATGGTTTGTTTGTTCAATGTGCGCATCCGTCATCAGAAGTGCCCGGT
TATGGTGGGAGCTGTGAAGATCCGAAGGATCAACAAGGCGCACCTTCCGATGTAGGGTGCGGTATCAATTTGAAGGACATAG
ATGTGGATGAACAATGCCGCATTCTTGAGGAAATATGTAAGGCTAGGTCAGCTTCTCAAGTCAGCAGCATGTCTTCACGTGCT
AAATGTGTGCAGTCTCAAGAGACACGAGCTATAAAGCGCAGAAAGCCCAAGAGCTCTGATTTCAAGTCTGGCCAATTGTCCT
TGTCTAAATATTTTCACTCTTCTGATGCTCAAGGATGAAATGATCCAATTGTACAACCAATGGAGATTGTAAAATGGCCACCA
AAGGTTTATAGAGAATTTTTCACTCACATAAGGTTTTGTTGACATGCAATATTTATGCTGAGGTTTAGTGTATAGGCTACCTTG
TGGTGGTGCATTCTTAAATTGTAATCATGATGTCCTACTATATAATTACACCCAATGCTAAGTTTTGCAACCTTATGCATTCTT
AAATTGTAATCATGATGTCCTACTATATAATTACACCCAATGCTAAGTTTTGCAACCTTA

**DAMAGED DNA BINDING 2 (*DDB2*):**

GAGAGAGAGAGAGAGAGAAGCTTGCAAGCGTTTTGCACAGTTGTGGGCTATGGCGGGAAGGCCTCAGAGGGCTGTG
CGCAAGGTTGTGGTGCAGAGGGATAGCGAGTCTGAGGAAAGCTCCGATGAAGAAGTGGAGAACGACCCTCTTGACGAGGAG
GAAGTCGCCGCCGATGGTGAAGAACTCAATGATGGTGAAGAAGATGATGCAGAGGACAATGTAGCAACTCCATCTTCCTCCA
CGCTGAATGAGAAGAGAAAGCAAAAGAAACCCATCACCATTAGCCTTAAAGCAGCTAAATTGTGCAAGGTATGTAAGGCCA
AGGATCATCAAGCTGGTTTTGTGGGTTCTATTTACTTAGACTGCCCTAACAAGCCTTGCTTTCTCTGCAAACAGCCTGGACACA
CAACTGCAACATGTCCCCATCGCATTGCATCCGAGCATGGAGTCACACCTGCTCCAAGAAGACAATCTTTGGGGATGATCGAT
TTTGTGCACGAAAGGCAATTGCGAAGTCGAGTCCTGAAGATGGTGCCACCCCCTGTCATTCCAAATCGTGTGGATTGTGCCAT
AGTTAAGCTACACAGCCGGCGGGTCACAAATTTAGAATTTCATCCGACTAAAGATAACATACTTATATCTGGAGATAAGAAA
GGTCAAATTGGTATTTGGGACTTTGAGAAAGTTTACGAAAAGACTGTATACAGCTCAATTCATACTTGTATAGTAGGTAGCAT
AAGGTTCCATCCGGCAAATTCTGAGATGATATATAGCTGTGGTGCTGATGGTTTGGTCAGCTGCACTGACCTGGAAACAGGGC
TTCCTAATAAAGCAATTGATCTAAACCCGGATGGATGGAATGGCCCTTCGACATGGCGGATGATTTATGGAATGGATATCAAT

CCAACTAGGAATGTTGTACTTGCAAGTGATAACACTGGGTTTTTGCACCAGGTTGACATAAGGACAAACAGTACAATAGGAA
AGCCATTATTAATACACAAAAAGGGATCCAAGGTGGTTGGCTTGCATTGCAACCCAGTTGATTCAGACCTCTTCATCAGCAGT
GGCAATGACCACATGGCACGAATTTGGGATATGCGCTTTTTGGATTCCCAAAGTCCTCTGGCAGTACTGCCACATCCAAGAGT
GGTGAATTCAGCGTACTTTTCGCCTGCAACTGGAAACAAGATCTTAACAACTTGTCAGGATAATCGACTACGAGTTTATGATT
GCATTTTTAGTAACTTGGCAGAGCCTAGCAGAGAGATTGTACACAGTCATGACTTCAACCGCTATCTTACATGCTTTCGAGCT
GAATGGGACCCAAAGGATACCAGTGAAAATCTGGCAGTCATTGGCCGTTACATAAGTGACAACTTTGATGGTGTTGCATTGC
ATCCCATTGACTTCATTGATACTAGCACAGGTCAGCTTGTAGCTGAGGCTGTAGACAAGAACATCACGACCATTACTCCTGTA
AATAAGCTGCACCCTCGATTAGATGTCCTCGCTTCTGGCAGTTCAAGATCGTTGTATATCTGGCGGCCAAAAGATGAGATGGA
GGATGATGACAAGGCAAGGGAAGCTGAAGATGAAGTTAAGAAGAAGATCACCATTTTCACTGTGCCGGATGATGCTGCTGCA
AAGAAAGGGAAGGATAAGAGGAAGTTTGATGGAGATGATGATGAAGATGATGACATGCTTTGCAATGGGAAGAAAGGGAAA
GGCAAGGCTGTGCGCCCGTCGCCTACAGCGAAACAAAAAGCGCGCTGAGGGCTCTTCCATTTTTCCATCTCATAGATTTAGTC
TCACATCCCCGGGTGGACATTTCTGTACATTATAGGCTAGGATTTTTGTTTCTGTAAACCATTTAGTATATGGTGCGTAAGCAA
CTATCTTCTAATTTGAATATGATGTTTATGGTTTGTCATGCAAACTCTCTTTTCTCTTTTGCCCAAGGGAACATCAAGGTGAGG
TGTATACTTGTGACATCCAACGACAAATGTGATGCCACGAC

**STRESS ENHANCED PROTEIN 2 (*SEP2*):**

ACACACACACACACACACACCAACTCATGCTCTGTTTGCTGCTGTCTTTGCCCCTTCGAGGGCGCCATGGTGTCGTCGT
CGTGTGCGAGGAAGCATTGTCGGCGAAGAAAGCATCTTTATCATCATCATCATCATCATACTCACCCCATGTATCCAGCTACC
CCTTGCACCAGTCAAAACAGGGGGTCTACTTACCTCCCAGACTCTGCAGCTTGAATGAGTTCGGTACGCGCATGGCTTCCGAT
CGCTCTCATGACAAAACCCACCTCGATTTCTTCACATCGCTTTCTGCGATCATCGAGCTCTCTGCCCAAGTCCAGAATTGGGAA
GTACTCTCTGGTCGGCTTGCCATGATGGTGTTTGCATCAGCTCTAATGCTAGAGGCTTTGACTGGCAATAGTGTGTTTGAGAA
GATGGAAACGCAAAGACTTCTTGAATATTGTGGTGCTACTTTTGTTTCCATTCTCGTTGCAGCGGGATTTGCGGTTGCGTTGCA
AGCAAAGGCTCATGTGGCATACACAGTATCTAAAGGTTATGAGAAGTTAATAAATGGGCTTGTCGATAATGTCATCGATAAT
CTTCTCTTTGATGATGAAGTATGCTAGCTCTCTGTCTTGAGAAGCAGCAGCAAACTTCTCTTATGTCACCTTTCTAGAGGTATT
ACTGCGGACCCAAGCAAGCTTTAGCAATATGAAGCTTGTTTCTTGTAAATATGGAGGTATGTGGCACAAAGAACTGCCACTTC
CAGATGTTCTATATTTAGTGCAAGCCAATATTTGTTGGGGCATGACTTGTTGGGCTTTAATCATGGCACCACCCAGCTATAGA
GTTGCATGACCTTTGATTTGGTTAGATCGGTCGAAACTCTTGTTTTCAAATCACCGCAGTGGAACCGTGTGTAAAGATGCTGC
CCTTTTATGATCTGAATGCAAGGTCATGTTAAGTGTTCATAATCAATAGTTTTTGTAATTTTCATTTGTGTTGATTTGTAAATTT
TAGATCTCAAGAGGCTTGTCAAATGAGGGG

**SALT OVERLY SENSITIVE 1 (*SOS1*):**

CCGCCAAGAGCAACCCTGGACAGGTAAGCCACAACAGTTGCTGCATTGAAAGTCTGACCCACCACCAGCTCCAAGAA
CAACCGAAAGATAACGATTGCAGTTCCATCATTTATCAATGACTCTCCCTCAATCAAAGTATTCAGTTTTTTTGCCTGCACCAA
GCTCCTTGAGAAGTGCAACAACTGCAACCGGATCTGTTGCACTTAACAACCCTCCTAATAATAACGACGTGCTCCAGGTCCAT
TGGTATGGAAAGGC

**HISTIDINE KINASE 1 (*HK1*):**

TTTTTTTGAAAAAATTGATTACTTGTATTCAACCTTGCACAACCATTGCAATGATATAGAGGTAGCTCCAACTAGACA
AACTGGGAAAAAGAAGAGATTACAAGGGCATGAAGACCCCTGGGAAAATACACATGCCTTTTACATGAAAGGAGTTTGATG
GCAACGCACTTACACTGGCGCACACATTTCTCATATACGCAGTGCCGTGTGTTGGTCTCCACTAAGATAATGCCAAATATGTG
AAAGGAATTTTGAGAGAACAAGAGCATATGGCCATTGGTAATTTGCCTACTTCTCTGCACATCATCCTCACGAGACAACACAA
TATTTTTACAGTTGTACACGCTTATGGTGTCCATAACCATGACCTCTCAAAATTCTACATCTCTTTGCGACGTCTTGCCAAGCT
ATAGATTTTCCATCAGCAATGGCATTTTACCCTTTTTGTAAGGTTGATATTTATTTCAGTACTCATGGAGAACGGTGAACCAAT
TCGCACATTGTACGGCCCCGGACACTCATGCCACGATTCTCGGATATGCATTGCACACTCCATATAATGCACTATTGCCGGCG
CTGTACACATTGTGTCAACATACATACTCCTCCAAGAAATTTAGCATTTGAAACATATTCCTTAACTGCCAACTTGATCAACAT
ACAGTAGCACCTGTCATGTGACTTCCGTACTTCTCCCCAATAATTTGGTGACCACATCAAGTAAGGCATCGATTGATAGTGGC
TTTGTTAGGTAGAAATCCATACCCGCCTCTATACATTTCTTCTCATCTTGAGCCATCGCATGAGCTGTCAAAGCAATGACAGGT
GTGCGAATACCATACCTGCTTTCTTCACTACGGATACGTTGTGTTGCTTCATATCCATTCAAAATAGGCATCTCACAGTCCATT
AGAACCAGATTGAAAGGTTGATTTTTCTCAGCACTTCTCAATTCTCCTGCCAGCTTTACATCAGTGCCTTCTGGAGGAGAAACT
TTGCTCCTATACAAGTTCTCTAGAACTAGCCGAGCAGCTTCAGCACCATTATCAACGCATTCAACAGTGGCCCCAAGGCGAAT
TAGTTGTGTTTTTGTTAACCGCTGGAGGATTGCGTTATCCTCGGCCACAAGTATGTGCATGTCAGACAGGGCATTCTGGAATCT
GACTCTAGAAGTTGAAAGATTCTCTGAGCTTTTATGAACTAAGGAATCAGGAGCTGATGACAATGGCTGATCTTTTCTTTTTG
ATTTAGTATTGTCAGGAGAATCCATCAGCTGACCTACTTGGGAATCCAAGGTCGCCAAATTTGGTAAGCTATTCATCATATTA
GAAGCATTCCTTGTCACAAGCGCAACTGAATGAGCTTTAAGGTTTCTCTTTGCATCCAAGCTTGATTGGAGACTTCCTGTATCT
TGCTGGAAAGTGCTCATACAAGAGAGAGATGTATACAATGAGTTTGCAGCACTTAGGTCTACCTTTTCACGGTTCCTTGTAGA
CAAATGTGGAGTTTCCGCTTCAAAAGAGTCCCGCACTCCATTGCCGACCAACTCTTGTAGTAGATCCCAGACACACTTTAAGC
GAGACGCATACAGTGGCTTATGCAGGATCAGATTGCAAACTCTCGTTTGCCTTCTTAAAAACTGAATGTCAATGCTGGGAGTA
TTGGCATTCACTAACCATGCAACCCTGAACGGGAGAGATTGGTCACTCTCCACTAGCATTTCTTCCAAATCTGAGCACAAATT
ATCAAGAGATGTTATGGCAAAGGTAGCATCTACAATTAATAACAGATGTGCCTGTGCGCCATAGAGGTTAAGCTTTTGAGCA
GATCCATTGCCATCAGATGTGGGCAATCCTTGCAAGCAAACATCAAAGCTAGTTTCATTCCTTGCAAAGTTTTTCGCATCGAG
TTCATCCACCCAGGAGTCATATATGTGACTTTCAGGTGACCTTTGGCCCGGCCTTTCAGACAAACTGAAGCTTGCAGAAAAAA
CTTCATGTTTAATTCTTTCGATTGCAGGTGCAAGCTCATCCCAGTTTGCAATTGTCCATACCTGTAATCCCCTCTTCTCCATCCA
CTTCTTCACTACCCTTTTCCCTGCCTGGCCCTGCATGGCAAGTAGAACATGGACACCTTCCATTATAGGAGGGGAGATGAAGC
CAGTTTCCTCCAAAGTATTAGAAAATACAAAGGCTCCAGTAGGGTCTAGCTGCCAACCAGGAGAAACAGAATCACCACTAAC
ACCAAATGTTGCTTGTAAGGAAGCCACATTTTGCAGTTGGTTGTCCGGTGTGCTGTTGTCTTCAGGCACATGTTTAGCGTGCTG
CTTTCCACACTTGAAAATCAGATTAAATCGAAATCGAGTTCCAGGTTCACCAGGTTCATCTTTGTCAATGATGTTGATGTCTCC
TCCCATTAACCTTACCAGTGATCTCACTATGCCAAGGCCAAGGCCTGTTCCGTCATAGTTCCTTGGTCCAGATGACTCAACTTG
TACAAAATTTTCAAATACAGATTTCCATCGTTCTTTTGGTATGCCCCTTCCTGTGTCATCCACCTCAAACTCCACATGCATGTA
GTTGTCATCCTGATCTTGCTCAGATAAACCCTGAATTAGCTTACGTATTTCCTTTTCCCTTGACGGGGACCTTATAAGTTGATT
CCATGGCCATGACAAAAGCCTTCCCCGACAACTAAACTGTTGTCTCGTGCCCGAAATGCTGAGTGGTTTTACCCGTCCTCGCA
GGACAACATGCCCTTCAGAAGTGAACTTTACTCCGTTGCTTAGAAGGTTGGCCATGATCTGCTTCACTCGTCCAACATCCCCT
ATCACCCACGAAGCCTTCTCCACAAAACCATCTGATAGGTCCAAGGCAATTTCTATCCCCTTCTTCGAGCCCACAACAGAAAA
TGTGTCTATCACTTCTTCAAGGGCACTCACCAGCTCAAACTCCGACTCCTGCAACAAGAGCTTTCCTGCTTCTATCTTGCTCAT
GTCCAGGATTGAGTTCAAAATGCCCAGTAGGTTGGTGGCACAGGATCTCATTTGGATCAAATTGGACCCCAGTTCCGAAAATT
CAGAAGCATCACAAAGGCAAAGATCAATTAGCCCAAGAATAGCCGCCAAAGGAGTCCGAAGATCATGACTCATGTTGGCAA
ACACTAGACTCTTGTGGCCACTCTTTGTCTCAGCCCTCTTTGTCTCCTCCAACTGCTGTATTAATGCAGCCCTTAACCGCATCTC
ATTGCTAACACCTTTGGTCAGAAGCGTGACAAAGAAACACCCTACAATCCCCATACTGGTAGCCAATGCAACCAATACAGCT

AGAGTAATGTGGCTCCACTTCTCTGTCAAGCCCCAAACACTGGCGTAAGGGAGTACCACAGTACACACCTGCATGTACGTTGC
ATACAGATGTCAAAACTTATATGCCGACATCCATTGCCATGACAAACCTGTTTATTCTTGGAGGAACTGAATTCTTTTTTTGTT
TCTTTGAGCAAGTAAAACAACGCTAGAGGCATACCAATGTTGCTCCAGAAATGTGCAGAGGTGCACTGTCGATGATGTAACG
CGCTCCATGTATCTCCAATTTCATTACAGTGAAGGATTGGCTTTCCATGGTTGATATGTAAAAGAGGTGCGGCATTAAATATT
CTGCAGCACCTGCAACAAGAGCATTATCGGACTGAGTTGCCATTGTCTCTTTTTGGGAGGTGTTATCCATCTTCGTGGTTGAAA
AGCCAGCTCCCGTCTCGGCCACTAAATGACCATCTGTAGTGGCAAGAAAAATGGCACCCCCTTGAAAGTCAAATTGATTGAG
GAGGTTACTGATGTCTGGTGTAGGTACACCAACCGAGACAGCAGCCACTGGAATTGCAGATTGGCTTTGAAGGACTGGAGAT
AAAAAGGAGAATGAAACCTCCCCGGAATCTATCGGTAAAGAACCCCAGCTTGCCGTTCCATTTTTACCCTGCAATGAGCTTTT
CACTACTTCATTGTCCCAGTATGCTATAGGGGGTACTTGCAATGCTTCTCCAGTTGGGTAACCTGTAAGGTCATTGACTTTCTG
ATGGTACCACTGATACCAAGGGCGTGTGTCCGCCATATCCTCTCCAGTCTTCCATGGGATTGCAGGTGCATTTGAGGACTTAG
AATAGTCGCGAGAGAGATAAGATGTGTTAGCATAAGTGAGATATGTTCCTGTAACATCTGTTGCATAAGATAGCAGTACCCC
ACTTGCTCCAAAGAACGTGACTCGTGCTTTCTTGGCAAGAAACGAGAAGACTAGAAATAGAACAGGCCTCACCTGGTTGTGT
AGAATTGTTACAGAGGATAGGCTGTCTTGTGTAAGTGAACTGATGAATGTGGCCAGAGAGGCCGCAGATGCATCATTCCCGT
GAAGCACAGCATTCACATTGTCCTTGCTGTGAAGAAGTAGCTCACGGCGGAACTTTGAGGCTAGAGAACCCACAGAATGCTT
CACATTGGTGGACGTGAGCACCCACGTGAGCACCGTCATGATAATTAGCAATATGGCCAGCAGCCCCATCAAAGCGAGCCTG
GAAGTGAGAAGTGGGTTGCACATGACTCCACAGCGAGACCGTGTTGAAGCATTGCCTTTCCAACAATTCAGGAGCCGCCAGA
TACGGCAGTCCTGCATCCCTTCATGAGAATCATCGTCTTTGTCATCATCAATGTCCTTGTCCTTTTGTCTGTGAGCTTCTTCTC
GCTGTTGTGTCCCTTCTCCATATCCTCAGACATGCTTGTGGCCCATGGCATGTCAAGAACAGCTTGTTCCTGGCATAAAAGCCC
AATATCATAATTGAAGAAGGTGTCTGCAGCACGTGCCTTTGCTGGAGAGGGAGAGGAACCCAAAAAAGAGACCGGGCAGCA
GGAGTTCGCAGAAGCAAAATGGCGTCTTTCGAGCAAGTCTTTGTGCATCTTGGTAATTGTTAAATATACTAATGCTAATAATC
AGAGAAGCATCTGCATGTCAGTGGAGCAGAACTAAAGCATTTGAAGATGCTCGAATACAGAATCCATTGCGTCCTGTCGAAA
GCCAAGTCAGGTCGAGTTTGTAGTACATAGTTCAAAGACAAGCTAGGCCTTTATCTTCTTCTGCAATGTACTGTGAATGTGAG
CAAGACAAATGCATACCAAGATGAGCATGTGGAGTACAAGCCAGTGTCTAGCGCACACACAATTCTGTTCAATCAACTGGAA
AAGAACAGGAAGGGACTCCGAGGGAGGCAAGATTGGGAGCAGAAAAAGAAAGAGAGAAGACACTGTGGGAAAAGAGAGC
AAAGCAGAGGACGGGGTAGATCAAACACCGAGTATGAAAGGGACACTCTTGGAGGAGGCACACCAGTGATCAGTGTGCTGG
CAAGGCAAGGGGGGGGGGGGAAGAGGGGGAGAGGGGGAGAGA

**IRON REGULATED 2 (*IREG2*):**

CGAAGTATCCAAGTTGGTAAGTTGGTAAGAGCAATAAGTGATAAACTTGAGGTTGTATACAAAGAGCTAAGGTACCC
TTAATTGGCCCCTTTACGAGACTCAAAGTTCTAAGCAATTGAAGAAAAAAGCTCTTATAAATATTGTATCATTAATATTTACA
TATATACATAAACATGGCCTTGGGATACTTCAGCAATAGGTTTTAAAAAAACCATCAATCAAATTGATTCTACAGCCTACCCT
TAAACCGCTCACCTCCACTAAAAAGATACCGCAATGATTGAGGTCTTCTCGTGTGGAAGCTCTACCGTGACTTGAAGATGCGA
TCCAAATGCAGCAAATGGCCTCGAACACGATATACTTGAAGAGAGTAGAGTATGAGGGACATTGTCACGGCCCCATAAGACA
AGTAGACCAGCTTATCAAAGTCCTTGGGGTTGGAGATCAGCATGCCAGCAACAAAAGAGAGCATCTCTAAGAAAGATTGCAT
CGAATTTTGAACTCCTCCAACTATTCCACGTTCATATTCAGGGGTAGATTCCTGCATAAGTTGCAACACTGCCAAATCAAACA
TCCAAAGACCAAAACGTGAAGTCACGACACCCCCCATAAGAAGAGCAGAGGATATGGTTGAGTTAGATATCCAATTTGAAGT
CACACAAAATGAAAGGAAACTCCACTGCATCACAATGGACCAAAGCCCTGTTCTTATTGTCTGTAGACGGCCATGAAATTTTG
GATAAGCGAAAGTTGCACAAATACCAACTAACGCAGAAAATCCTCGTACAAGCCCTAAGATGTAAGAGGAAGTGCCTCTCCA
ACTCAACGCAGCTGTCATTAAAGACCCAAAGCTTAAAACTGTAAAATACAACATGGAAAGTGCCACAGCTGCTAGTAGCATT
TCCTGTGCTGCATACACTTTAAAACCTTCGACAAAATGTAGTTGTAAAACTCTCTGGTGAAGAATTGAGATCCATTGACACGC
TATTTCCCACAATCTTTGAGACGAATATGTTGTATGAGATCTATCTCCACTTTGAAAATTGTTGACTGATTCATTTGTCCTCCC
AATCTCCTCAGAAACACAACTACTCACAAGCTTATCAGTCGCAGGATCAATGCTCTCTCTGGCAATGTTTGGTACATGCATTT
CTATGTCATTACTGTTATCAATTACAATCCTTTTTTGCTGAAGTTCTAGAGATTCGTTGTACAAGCACCTCAGCAAATAGTACT
CGAACCCAATAGACACAACATTCCATACACATAATAAAATGGCAGCTGCCAGAACTGATACAAAGCTCATTATCATTCCCAC
AAGCACAGGAGCCATAAGCTTGCACGTAAGATCAATGCCTCGCATTGCTGAGTTTAGTTCTGTGAGTGTTGCTATGTGCTGTC
CTTCTGCTACAGCAAGCACCCAATCTCTCTCAACAGCAATGTTTGTAGCAAGGGCTGATAGGGCCGAGAGTCCACCAAATAA
ATTTACAAGGACAATGAGACTAGCAAATGTTTTATAACGTCCAAATGCAAAGTCCCGATCGAGAAGCAATATTGCTAAGGCC
AATGCTGCAGCCACAACTGACCCATTCCTTGTTCCCAAAGAAATTTGCAACACCTTGAGGCGTGGGGTTCTATCCACCCATTT
TCCAACAGTGATGCCCATGGAAACCACAGCCAAGTTTTCTACAAGCCCGTACAAAGAGGTGAGAAGTAATGAGTTTGGCCAA
ATCGCAATCATGAATAACCCAATCCCAAACTCCCACATTCTGGTACCCCAGCGTGTGAGGAAGTGGCTGACGTAGAGGTAAG
CAAGAAGACGGAATCTGGAAGGATACAAGCCTTTGAAGCTTCCATGACGGTCCTGCGTAGTTAGAGGTTCTTCCTCCTCCCGC
TGCTCCTCCATTTTTTCTCGTTGCTTCTCTCACTCTCTCTCACTCTCTCTCTCT

**PHOSPHATE TRANSPORTER TRAFFIC FACILITATOR1 (*PHF1*):**

TTATTATCGAGATCGCCGTCGACAGCAGCTGCAGAGAGTATATTAGCAGAGTATATTTCTGGCATCTTCTTTCGATATT
TTGCACGAACGCACGCACGCACGCACACAAAGCTCTTCGCATCTACAGGACCTCTCTAAGCGCAAGGATCTACACAGGCTTC
CATAATTTTGGCAAGGAAATCTGTGTTGTCAATTAGGGTTTCTGCAATGCCACGCACTCGCTACATGCGGGTCGTTAAGAAGT
GCGACTCCCCGCTTCTGTGTGCTTCTTGGACCGCTGGCAGCCCCAACTCTCAGAAAGAGGACATTGAATACATTGTCTGTGGG
AAGACTTCAAGTGAAACAGCTCCTCATGGCAGCTTAGAGCTCACCCGCTATGACTTCAAGGAGGACTGTATATCAGAGGCGG
TTCACAATCTAGTGGAAGATGATGTACCTCAGACTCTAGCAGTTCATCCACGAGGGGATGGTGTTATATGCTCGTTTTCAAAT
TCTTGCAAGCTGTTTGGGTTGGAGGGGCTAAAACTTAAAGCTTTGGATCAAGGTCTTGTCCCTTTGCAGGGCATTGGAACACA
AGGAAATCTGGTTTTTAGTGCAGATGGGTCTAGACTAGCAGCCGGTGGAAAGGATGGCACCTGCGAGTCTTTGAGTGGCCA
AGTTTAAATCTTTTGTTTGATCAGCCTGATGGCTGCAAGTTTCTTAGCAGTCTTGACATCAGCTTAGATGGTGCTTTTCTAGCT
GCCATACCGGATGGTGGGAACTACTGCCGTGTCTGGGAAATTGAGAAGTCAACAATTGTAACAACAATTCAGAGCAGCAACA
AGAATGAAATGTTTGGATCGTGTAAGTTTTCAAGGGATGGAACAAAGCCATTCTTGTTTATAACCATCTCTAAAGGTGGAAAA
GGTGTTATTGGAGTCTGGGAGATGGAAGCATGGAACAAAGTGGGCGTAAAGGCGCTCTCACAAGATCCCATTACATCATTAT
CCATCAGTCACAATGGCAAGTGGCTCGCAATTGGATGTGCTGCTGGGGATGTTCATGTTATTGAAGTGAAAAAAATGGAGAC
CTCTGAGCTTGTAAAAGATGCACATTTATCTGCAGTCACATTATTAGAATTCTCTCCAAATGGCAGAGCATTTTTGTCATTGTC
TTCTGATTCTACGATAAGGGTTACTAAACTCAACGTAGCCAAGCAGTGGAAAGAGTGGCAGATTTATTTGCTGTTGATAGGCC
TGATTTTGGCTTCTGGAGTTATTTTCTACATCTTTTTTGAGCACTCCGATTCTTTCTGGAATTTCCCTTTGGGCCGAGATCAGCC
TGCCCGGCCTCCTCGTGAGGCAATATATGGGATAATGCCTGATTTTGATGGAGGAGAATTTATGTAAAGCCATAAAAATTTAC
CAAACAGACTTAGATTTGTCCACTAGTGGATTCGCCCCGGCGAATGATGATGTGTTGTTGAGCAGCCCAAGGTGCATTACCCG
CATGGCATTACTACTCTGAATTTTGACTACACTCATTTGGTCTAAGATTTTATAGATAGAAACACACAGAATGCAGACGGGAT
TGGGAGAATAGCTCCCATGCTTTGTGTCAATGATAGTAATAGAATATTTGATGTGCTTTCTTAAAATGTTTGCCTCTGTCGTGC
ATTATTTG

**PYRIMIDINE 2 (*PYD2*):**

GACACAAAAATTGAGAGCATCCCAATCATGAAGATCTACAAGAGATATTGCCAATACATGAATTAGTGACATTGCTA
TTACATTATTATGCAAATAAACTCTAGGAGCTTGGTAGAACCTAAATACTCAAGATCCACCAATCATCGCGACTGTATTGGGT
CTTCATACTTGACTTGCATGATTGGGCTATTTGTAAAGGCAATGTATTCACACATGAATTAATTGATGATGCGGTTTGGTAGG
GCAAAATATGTCATACAAGAAAAATTTGTCCCCTATGATCTTGTCATTTGTGTAATCTGATAAGTCAAACCAATCTTAAACTT
ATTTATAAATTGAAATTGTACAGATTGGTGCACTCTTTAACAAAAAACCCTTGAGAAACTACTTCGCATCTTGCTTGCGGGCT
CTGAGAGATGAAATAAAGGCTGAATCAATTTTGTCAAGCCCATTAAATAAATAGCCAAAGGGAGGCAGAGATATGTATCTTC
CTGATCCTGGGCTCACATTCAACTTATCATTTTCCCAAACAACTCTTCCTCTGCTTATTGTCATTTCAACCTTTCCCGTCATGGT
CCATCCTTCATAAATATTTGTATCAATCTTTGAATGATGAGTCTTAGAACTGATAGTGAAGGTTGCATTGGGATTCAATAGAA
TAATATCTGCATCAGACCCAGGGATGATAGCTCCTTTTCTTGGGTAGATATTAAAAATTTGGGCACAGGCGGTACTTGTTACT
CGCACGTAGTCCATAGGTGTGATTTTTCCAGAGTTCACCATAATATCCCATAAAATATGCATTCGTTCCTCGATACCATTAACG
CCATTTGGGATCTTCCTGAAATCACCTTTCCCTGAAGCCTTCTGTGTGGAATTAAAAACACAATGATCCGTTCCTACGAGATCT
TTCTTCTCGCGATCTCAGAGACGCTCTAATAGTAGATACCACTGTCAAAACCCCAGATCTCCTTGAGATGCAAACTCGCCTCC
CAACCATGCTGATCATGACTTGATCAGCCCCACCGAGCCCGAGCTAGAGACCCTCGTGGAAATTGCTAGCCGCATTGCAATCC
ATGCCATACCCTTTGGCGTCCCATACGGTTGGATGATCGCTTACGTTGCTAGCTCCACGCTCCACATCTGCCTGGTGTGTCCCT
CTATGGCGGTGGTACTCCCCTTTATAGCCTCTGGGAAAAGCGCCTGCCAACAAAATCACAGAAGCCAAAAGTGATGAAGCAC
AGAAGCCAAAAGTGATGTTGTTGACCCATATTCCCACGGATAGCACTCTAGCCCGCCTATCCCTAGCGTGTACATGACATACT
CCCCTTCCTTTGAAGAGCTCACTCTTGAGCTCCAGAAAAAGAACTCAAGGGGAAGTAGGCACACTTGCAACACTTAGAGAAC
ACCCTCTATTGGTCAAAACCAATAGTCTCAAGAACAATCCGACCAAGAAAAATAATTAACCTACTGAAATTGATGACATGAG
CATCAAAACCTAGTCACTATGTGCCACATGAATCTTCTGGATAAGTCAAGTCTAACAGTGCTACTAGTAGACGAGGTAAGCA
ACCAG

**ACTIN-RELATED PROTEIN C3 (*ARPC3*):**

TTTTTTCCCATCAAACAAACAAAGAACATGCATTGCACTCCAATACATATTACAATGACTCCTTTCTAAAATCACAAA
TGGCAACAGCCTCTGACCAAGACCACATTGTAAGTGGGGTGTGCATTGAGGGCACTGCATTCATACCAAGACATCTCTTTCCC
ACAATCCTCAACACTTTTACCCTTCTGCATGAAAATTCACTATGACATATTTTGAAAACTTTAAAAGGATTTATTGATTGTTTT
GACACACTTATTCCCATTGATTCAGCCATTATCTCATACTTACACTTGGCACTCCCACTGATTCACCAATCTTATACTTACACTT
GGCACTCCCATGTTTGTAACCTTGTCAAGGTTACAAGGTTGAAGTATTGACATCCACCAGCAGAGTAAACACTTTTTCGATTTT
GTAGCCCATGGACAGTCCGGAAACATTCCAGAAATGTTTCCTAAGCAATGGCCATGTTCATGAATCGCCGTTTCGAGAAAGC
AACCCACCATTTATTCTGTGTACCATTTGGGTGGTAGGCTCGCTCCATAAGTCTGCCACTTGTTTCTTCCCTCAACTGCTTCAA
GTAGCTTCTGAGCAATTCGCCTTCTTGTTGAGACACATGAGGTGTAAACAAGCCCCCAAGAGGAAAGCCAGGTTCTCCAGGT
ATTGGAAACCTTTCAAGACCCAGAGTAATGATAGCTTTTGTTCCTTCCGCTTCAGTCCTGCAATTTTCTACCCTTTTGAGGGCC
ATATTAACATAGAGTGTCAAGTAGATCAAGAGTCTGTCAGCAGAGCTCTTGACATCAAACTGGCTCAGCTGCTGCTCTAATGT
GGCTCTTCAAACGAAGAAGAGGGTATCCCTTCTGTATTCACAAAAGAAGAGTGATAGACCTGCAACAAAATATGGAAACAAG
ATCGTATAATGCGAGTGAGTTTCCCCTCTCCCAACCTCATTTAATGGCTCGCATTTTTAGGGTTGCAGAAGGGTCAAGCTCATT
ACTGAGAGCTCCGGCATGCCATTTCTCTTCTTGCTCCGCCGCACTGCTGCTTTGTGCCGTCCACCAAAAGAAAATACCGTGTTC
GCATCATCGGGCAGTAGCCCAGTTGG

**ACYL-COA-BINDING PROTEIN 2 (*ACBP2*):**

CTCGTATCAAAGCTCTCTGTCTCTCTCTCTCTGTAAAGAAAGAAACCTTACATTGCACAAATTCTCGAGACATCTTTGA
CGTTTAGTATGTACAACACTAATCACCTAAGAAACTAAGGTTCAATATACATTAACCAACATATTACTGGATAACGAGTGTTT
ATTCAATCCCATGTTGACTGTCATGTGAATAGTTCCATGTAAAAGGTGCTCATTATATGTACTTGGTTGAGCAACAAAATTAC
AAGGCTGCAAAAGCTTGTTACTTAAACCAGACATTTCTTCAGTAAGTTCTTCAAACTATTTGATATCTTCACCTCCAGAGTTCT
CCTTACAAGGTGATAAGGAATGTTCATATGGTGTACAGCCATCATTGTCTACCAAGGATGGGTCAGCACCATTTTCCAATAGG
AAAGCAGCTATTTCATCCCATTCACAAGTGATAGCATAATGTAGAGCTGTTTGTCCGTCTGAATCCTTATCATTAATCTCTGCT
CCCCGAGAAATCAGTAATTTGACAATATCAAGGTGACCCTGGTCAGCTGCCCAATGTAATGCAGTCCGATTATCGCTATCTTT
CTGATCAACTGAAGTATACGATTGCAGCAAAGTTTCCAAACCTTTCAAATCGCCTTCCTTAGCACATTTGTGGATACCCTCCA
GACCCTCCTGCTCAATGGCATCGCTTAACATTGTGCTAAAAACAGGACCCATCCTTGGAGCCTTCTTACTGCTTCCTGTAGAA
AGGCCATGAGCTAACTCTTCATCTTTATTCCAATCTGGATACAGCTCTGAAAGCAGAGTGATGTAACGTTGCATTCCCTCTTCT
GGCGAAACATCTCTTAAACTTTTCCACGCATTCCATTTGGCTCGTGCAGAAAGTCGATAAGCTGAGGGTTGCGGAGAAGAGC
ATGGTCCCTCGGTTGCTTGTTTGTAAAGGCCATAGAATTGAAGTTTCCTATCATTTGATATTTTGGAAGAAGGATCTAGTACCA
GCTTTTCAAGAAAGGCAGCTGCTTTCCCAAAAGCTTTATCCAAGTCAGAACCATCAATGCCTTCCCAATCCTCGTCGTCTGAA
AGAAGGCTTTCCTCTTTGTCGGATCCTTCTTCAAGACTTGCTACATTCTTAGTGCCATCTAAGGGAGGGGCGGCATCTGCGAC
AAAGTCGTCCTCTGTGTGTGGCCCTTCATGTGCTACATCCGTGCCTTCACTAGATGCACCTTCTGAAGGCATAGTTCGTGTATT
GTCTAGCTTATCCACATTGTGAAGGGTATGCTCACCTTTTGAGAGCGTAAACCCCAAGGTGTCTCTAATCGTTGCAGACGGCT
GTGCTACATCAAGGGTTTCCACACTTGCAACCTGTGTAAACCCTGGGTCTCTCTCTTCCATTACCTCGGCAGAATCGGGGCTTT
CCGCATGCTCCCTCTCAACTCGAAGGTTCTCACTTCGAAAGGAATGCACAATCGAGATGAGCTTTACAAGCATAAAGGCAAA
GACAAGGGACATGAAGCACGAAGTAACCCATTCCTGCCAATCTGCCATGGCTAATGATGATCAATGCGAGCGCGACAAGGAG
GGCGGTGCTCGCTGGCTTTCTGGCTACGTTGTGTTGATGATTGTGATGTTGCACATGGAGGATCGCGGCTATCATGAGTAATA
TAAGTGAGGAAAAA

**HEAVY METAL ATPASE 2 (*HMA2*):**

CGACGGTGTTTCTTATAGTTTGTACGTGCAACTGGTACTATGTAGTAGGTCCCTATTTAGTCTCCCGTCTCCTTGCTCTT
CCTCAGAAGTGTCAACGCAGGCAGAGCTCCAATGGCGACCTTACCATCGCTCAAGTCTCATCACCACAAAGAAGGAAACCAT
TTGCTGCTACGCTACGGCAACAAGCTCTCATACTTCCTTGTCCCTCCTCACCAAGAGGTGGAGCTCTCTGAACTATGCTTTACC
GATCACAACCGCGGCATGCAGGATCCAAGAATCGAGCAATTTTGCGCTCGAGAGGGTGGTGGCCTTCTTGCTCATCTTCACCA
TGGCGAAGTGATTGAAGACATTTTCTTGGAGGAGTGCATCGAGGGGGCGGGATGTGGCCTTGGCTCTTTGCCATGGGAATTTT
CAGACTCATGTTTGAATGTGCTGTCGGGCGGGTACCAAGACCTTGAGCAGCACAAGGCAGGGAAAGAGGAGCATGTGCACAT
GGAGGGTTGTGGTCATGAGCAGATCAGACATGGGGATCATTTGGACTGGCTAGTACCTTTGAGTGATGGCTTGTTTGCGTTGA
GCCATGGTCTTGGAAATGACGATGGAGGGTTGGAGCATGGCCGGCTTGTAAAGATCGGTGATTCACATGGAAAAATGAGAGA
ACGTGAAAAGAGCCCTTACGATTTGTTCAAGTTTGAGGTTCCGCAAACTGGTGATTGGGATACTAGAACAAGAAGTTCTCCTC
AGCTAGGCTTTGAGCATTCCAGAGCTGGGAAATCTTCTCCAATAGAGCAAGTATCTGATGCCATGCAACCTTTGTTAAAAGAG
CAAGAGGTTAAGATAAGCATGGGAGTAGATATGGAACCAGTGATATTGCAAAGGACTACCATTGATGTGCTTGGAATCTGTT
GTCCAGCTGAAACTCCCTTGGTCAAAAAGATTCTTGAACCTGCCCCGGGAGTACAAGAAGTCTCTGTGAATGTTGCAGCTAGA
TTAGTAACCGTACACCATGATCCTTTCGCTACCCCCCCTGCCAGGCTAGTCAAATTGCTGAATGCCGCACATCTAGAAGCCAA
TCTTCACGTAACAGGGGAGTGGAAAGCAGCACGCAAATGGCCCTCTCCATACACATTCGCTTCTGGCCTCTTTGTGGCAATTG
CAGTTTTCCAGTATGTGTTTGATCCTTTGAAATGGGTGGCTCTGGGGGCTGTGGCAGTGGGCACACCACCTATCATTTTGAAG
AGCTTCACAGCCTTGCGACGTCGTATACTTGACATCAATTTACTGATGCTTATAGCAGTTGCTGGTTCTGTTGCTCTTGGAGAC

TATCTAGAAGCGGGCTTGATTGTGTTTCTCTTCACTCTCGCAGAGTGGCTAGAGTCAAGGTCCACTGACAGGGCACGACTCGC
AATATCTTCTGTGGCAAGTTTGGCTCCCCAGAATGCTGTTCTTGCAGACACTGGTGTAAGAGTTCCCGTCGCAGATATCAAGG
TTGGGACTAGGCTGGCTGTGAAGGCAGGAGAGTCCATTCCTATTGATGGCGTTGTGGCGTCTGGTAGGAGTGTTGTAGACGA
GAGCAGTTTGACTGGAGAGTCAATGCCGGTCGAGAAGGAGGTTGGCATACGCGTGTGGGCTGGCACCATCAATATGACCGGC
TACCTCTGTGTTGAAACATCTGCCCTAGCTGAAGATTCTGCTGTAGCCAGGATGATAAGGCTAGTAGAGGATGCTCAGAATCA
GCGCTCTCACGTAGAGCAGTTAATGGAGAGATTTGCCAAATACTACACTCCACTTGTGATAGTTGGTGCACTAGCGTTTGCTA
TAATCCCAGTAAGTGTGCATGCACATAACACTCGACACTGGCTGTACTTGGCGCTGGTTCTCCTTGTGGTCGCTTGCCCTTGTG
CGCTAGTGATTTCAACACCTATTACTACAACATGTGCAATAGCCCAAGCTGCTCGCACAGGCCTCCTTGTGAAGGGTGGAAA
CACTTGGAAACTCTTGGAAAACTGAAAGTGATTGCGATGGATAAAACCGGAACGCTTACTGAGGGCTGTTTTCAAGTAGTAC
AAATGCATTCCTTGGGCAAGGATGCCAATCTACAGAAAGTTCTATTTTGGTTATCATGTATAGAGAGCAAAGCAAGCCATCCA
ATAGCTTCAGCTGTTGTGGAATATGCAAGGCTTCATGGTGCAGAATCACCCAATGCTAACACGCTTGTGGATGACTTTAGAAT
ACTCGTTGGAGAAGGAGTCAGTGGTATTGTTGATGGTCATGAAATTTCTGTTGGGAATGAACGCCTAGCAAATCGGCTGCATT
GGATGGAAGCTGCGGCTGCTGAATCTTTTATTTTGGAGAGTTGGAAAAGCCAGGGGTTGACAATCTGCTGGGTTGGTGTGGAT
GGAAAGCTTGTGTTGATTTCAGCGCCGGTGACCAGCTTCGTACAGAAGCACGGGAGGCTGTTAAAGACATGAGGGACTTGG
GTCTTCAAGTAGCTATGCTTACTGGAGATAGTCTGGCAGTAGCCAGCACTGTACAAAGAAAGCTTGGACAAATTGATGTTCAT
GCTCAATTATTTCCCGAGGACAAAGTGGAGCTCCTCAAGCAATTGAAAACAGTTGGCCTCACTGGCATGGTTGGTGATGGGAT
CAATGATGCTCCCGCTTTAGCGGCTGCTGATGTTGGCATAGCTATGGGTGTTGCCGGAACTGCGATTGCTATGGAGACTGCTG
ACATTGCCCTCATGACTAATGATCTCAGAAAGCTGGCAACTGCTGTAAAACTTGGGCGAAAAGCTCGCAGAAAGATTCAGCA
AAATATCTTCTTATCCATCATAACCAAAATACTGGTGATTGTGCTTGCAGCAGTAGGGTATGCATCTTTGTGGGGTGCTGTTCT
TGCAGATGTTGGCACTTGTCTACTTGTCATCTTCAACAGCATGTTATTACTAGAAAGAAAGAAAGATGGATCAGGTTGTTTGG
GATTATTTACATTTGGGCGCAAGCCCAGAAGTAAAGTGTGCCAAAAAGATGTCTTACTAGTTAGTGAAAAGGATGCCAAAAC
GGAAGCGGAACCCTGTTGTCTAAAGGTATCAAGCCCTTCAGATGGTTCGAACACTTGCTGCAAGAAAAGGGAGGGTGATGGG
AGTTCTGTTTGCAATAGCCATCAGGAAAACCAATCAATCAAGTCCTGCTGTGCAGCAAAGTCGTGCTGTGATGCAAAGAACG
CAAACAGTGGAATGAGACGTAGAGTGAGCAATCGAGACCCTTGCGATAAGGATTCACAAGAAATTGCTGAGGGACCGTGCA
CTGCCATTAGTATTGCTGATCTTGAAGGATTATGCTTGTCATCTCAGATTGTGAACAACAGCAAAAAGAATTCAAGCGGGTCC
TCTTCAAAGGAGAATGCAAGTTGTTGCTCTAGTGGTAAATGTAATGGTTCCAAAAATGAGCAATTCAGCAGCAACGTAATAC
AGTCAGGAGATGATGGATCTTGGGGCACCGCCATTAAAAATGACATTGAAAACAATTCTCACAGCTCTACGCGAGTCGTGAA
TTCATGTAGCTCTAGAACCAGGATTGACCAGTGCTGCACGAATGATACTTTGCTGCTGGCTGCTATGCCTGCATGTTGCAGCT
CAAAATGCCAGGCCCCTCCAAGCCTAGAAGCTCAATCCACTAAGGGCTGTGAACAAGGGTGCTGTGAAGGGAAAAGTCGCA
AGGCAGAGGAGTCAGCAGCAGTACAGCCAAGACCAGTGTTGCAAGAGAAAGTTGAAGGAGTAGCTTCTGAGGGAAATGGTA
AGATCGGCTTGAAAGACAATTGCTGCATGTCTAGGCAAGTTGTGGGTTCATGTGGCTCTAGGAACAGTGCTGACCAGTGCTGC
ACGAATGATTCTCTAGTCGCTACATGTTGCAGCTTGGAAAGCCAAGCCCTTTCAAGCTTAGGGATTAGATCCGCAAAGAGTTG
TGAAGAAAGATGCTGTGTAGAGAGGAACTGCAAGGCAGAGGATTCAGCAGCAGATCAAGGGCCAAGGCCAGATTGCCAAGA
GCAACCTAGGCAAGATGTGAATTTGTGTAGTTCTAGGACAAGTGTTGGGCAGTGCTGCATTCATGATACTTTAGTGGCCACCA
CCCCTGCATGCTGCAGCTCGGAAAGACAAGCCATTCCCATCTTGGAGTGTAGATCTAATAAGGGTTGTGACCAAGGATGCTGT
GAAGGGAGAAGTCAGGAGCCAGCAGCCTGGAAGCCAAAACCAAACTGCCAAGAGAAAGCCGCAGGTGCATCTCTTGTGGGT
GGGGACAGGAATTGTAAGTGCGACATGAAAGAACACCCTTGCATCTCTAGGCAAGATGTGAATTTGTTTAGCCCTAGGACAA
GCATTGACCAATGCTGCATGAGAGACACTCTGATTGCTGCTTGTTGCAGCTCCAAATGCCAAGCTCCTCCAAGCTTAGAAATT
CAATCTACTAAGGATTGTTGTGCAGGATGCTGTGAAGAGAGAATCTGCGAAGCAGTGAAGGAAGCAGCAGAGGAGCCAAGG
TCAGAGTGCCATGGAAAAGATGGTGCAGCTTCTCTGGGTGTGGAGGGAAATGAGAAGCAGTGCTGTCAAGCCGGGCCCACTT
CGTATGATTGTGTGGGAGGCAATAAATGCTGTGCAACTGCTTGGAAAGATAAAGATGCATGCTGTGGTGGTGGTTCCACTAGT
AGTCATGGTAGTAAAGATGTGGGCTGTGCTAGTAATAATGATACCTGTCGTGGTGATAGCTGTGAAAAGGGTGAGGCAGCAA
CCGATGATAGTGCTTGTGTTGGCGGTGGTTGTTGTCGTAGTAGTAACATAAGACATGAGCTAGAAGCTAAAGATGGAAGCTA
TGATAGTTCATGCTTTGAGGGTAACACTTCTAAAGAAGAGATTGATTGTTGTCGGAGTGGTGGATGCTGTGGCTCAATTAGCC
AAGACATAAAGGCTGGGTCCTGTGGAGCTGTTGCCATGCCATCGATGGAAAAGCCAGGCCAAGCTGAAGGACCTGCCATGTC
ATCAATCGAAAAGCCAGGCCAAGCTAGAGGATGCCAAGGTAAAGATTGTTGCGCTGTCACTCAGTAAACTCACGAAAAGAGT
CTCAGCTGAGGTCTGTAGAAATCTACTTGTCTGCCTACGCTGAAACAGAGTAATTCCAGAGTAATCAATTTCCCCAGTGCTTT
ATGTCATTTTATATCCCCATTGCCGTCTTACAAAGCTGATGAGAAACAATATGGTCAGGGAGCTGGGCAAGCAGAAGGCATT
GAACCTTGGGTCCAACACCATAACATGTGACTTCGCTACTTCCTGGATCTCTTGTCTATCAATAGGACTTTGCTGTGGTAGACC
TTTTTCTGCACTATGATGTCTTGTATTGTAGTAGAAAACTTCTCTCCTTCATACCGGTAAGCTTGTTATCGGCCTCGCGCATACC
TTTCTTTGTACCATCCACTATCTTCTTGTGTAAAAATATTGGCCTCCATGATCCTTGTAGAGATAGTAATTATTAAATGTTTGTC
CAAGTGGTATGACCTAGACCAATCTAAAGTCCCACCAATCACCTATATCTTGGCCCCCCACACATTTCCCATGTCAACTTGCA
TTTGGGGGCTTGACATTTGCTGCCCCTGTAGGGCAGGATTGCGTGTGCATCAATACTCCTAGCAATAGCCGACTACGTGTGCC
AGCTCAAACACGAGAAGTTTTAGATAATTTCCCACCGTAATTCCCACTTTGCCGATTACGA

**PLANT CADMIUM RESISTANCE 2 (*PCR2*):**

GTAAAGGTGGTCTAAAATTTGACAAATAGACTATTAAACAAGCATAACACTACATTACACATAATGACACAAATCAT
GGCACTGCATAACGTTAGCATGCAGGAGCATTATATGGTACATCACTTTATACTAAGGGGCCAATGATAAATCCATATTGAAT
GATTGCAAATTATTTTCTCAAACAACACCTAGGATTAATATTACTAAAATAAGAGGGAAAAGTTTGGGATAAAAAAAATTTG
AGACAACATTCTTCGAATTCCTTATTTTTGCATGACATGAGGTTGAGGAGCAACATGTGCATATTTCTCAGAGTTAGCGGCCC
ATCCAAGGCAAGGATCAATGCCCCTAGCTTTGAGCTCACGATATTCTTGGCAGATAGCACATGCATCACAGAAGCAATGCAC
CACAAAGTCATGGCATGGTGTTGCTGGAAGGCCAAACTTTTGTCGAAGCTTGGTCCTATAATGGGTAGTGTACATACAAGGGC
ACCCTACACATGTTGCCCCCAAGTAGAATATAGCATTCATAGGACATGATTTGGTGCCCGTATCCAAAATCTCTCCAATTTGC
CCAAACGTCACGCACGGCAACCACCATGTAAAACAACAGTTTTTGAAGTCCTTGAAGCAGTCGAAGAGGCCCGAATGCCAGC
CTTGTTGGGAGGATGAGCTCGCCATGGCCTCTTTTAGAGAGAGAGAGGGAGTCTCAATAGGTGGTAGTTA

**HYPERSENSITIVE TO EXCESS BORON 2 (*HEB2*):**

GTTGGCTATGAAGTCCAATGCTTGCAACACCAGGCCATAGAGTTATTCTACCTTCCGGCTATATATTTGGATGGAGCC
CTGTGTTAAGAGGGCAACATTTTTTCAAAAATTGAGTGATTCCCCCAGAGCCATACACACACACCCAAGGTGCCAAGATTCGA
CCTTGGCCCATAAATTTCGTGAAAATCATTTGAAAGCTGATCTGAATCACTCAAAGCCCCCTTGTTTGCCAGATTTTCCATGTT
TAAGATATTGATAGTGGATATGCTGCTTCACAAAATTATCTGCAGCCAAATGTAGCATATTCAGCATGCAGGAGGTCTTATTG
CTGGTGTAATGCAGGAAGATTCTTTCTGCTCATCCACGATGTCAATGTTCATTTCCTCAGGAACATCGTCGAGATTTAGAAAA
TCTTCCTCTTGATCTTCAATGATGTCTGAATCTTTCCCATTGGCTTGTATGGAGCTGTTTTCCCGTTGGTCTTGTTTTTATTGGC
TATGAAGTCCAATGCCTGCAATACCAGGCCATAAAGGTATTCTACCTTCCGGCTATAGATTTGGATGGAGCCCTGTATTAAGA
GGGCAGCTTCTGCAAAATTGAAGGAATCATGGCCGTCTTCATCCGGAAGCGACACCAAAGAGAGCCCTGCTAGGTAGCTTTC
AAGCTCCTGCGCCACATCCACAGCCCAATTTGCTGCCAGGTCCCGATTAGGTTGCAGCAGATGCAAATATCGAGGCTCCTCTT

CCTTCTCCATATATCCTTTCAATTTGCACTGAATCTCAAGCTCTGATGGTGCACTGAGCTTTCCGTATCCAAAACGAGCAAGGT
GGGCATGCTTCTAAAAGACTCTTCATAATCCTTGATCATTCCTGTTACAATTTTCTGCAAAACCCTAGTGTTTCAACCCAGAAC
CCTGCTTTGTCCTCTCCAAAAAGCCCTTCAGATATCTGCAAATCCCTAGCATGCTCTATAACTTAAAGCTGCAGGGAAATCCTT
CTCGTCCGGGATCTCCCTAAACCCTAATTTATCAGCTAAAATTTACAGGGTAAAAGAAACCCCTAATCCCTTGTTGTAGCAGA
AACTTGCCGCAAGAACTTGAGAAACTACACAATGCGCTTCTGGTTCCTCCTCTCTGCGTCTGCTTGAAGTAAAATTCAAACAG
CTACCTAGTTCCCGCTCGCATCAACGACACAAAACTCTTTGGGTGCTTTCATACTTG

**ELONGATOR PROTEIN 6 (*ELP6*):**

CTACATACTTGCATAAAGGCATCATAAAAACCCCAACCAAGACAGAAAAACATTAAGGGCAAAACTCTATGACCAAA
CTTCTCTAATGGAAATCTTAATTTCTTGCAAATTCCCATGTTTGGTTGCTGTATACCCACTAAAATGAGTTGAGCACTTGCCCT
GCCCACGGCCCGAATAAGGGGCGTAAACCAAGTTTGTTGGAGGGTCAGCTTTAAAGGTGCAAGAGGATGTTCAAGTACAAAA
ATTTACTTCCTGCTTTCAGATTTGCTTTCCAGGAGCAAAATAAAGGACTGTGTTCTCCATCAGCTTGTAGTGAAGACCATAGCC
ATTCAAGTCTTGACGCACATTTGAGTGACCACTGGAGTCTGTATTGCAAGGCATCCTATGCTCAACAATCAGCTGCCCATGAA
CATCATTGGCTTGCCCTGTTGCGAGCGGTTCAACATTGATGACTGTGTCTGCCAAGTGCTCAAGCTGGTAAGCCATCACAGTC
TCATTGGCAGTTTCATAAACATCTCGATGAGTAAGAAGTAGCAAAGAACAAACCTCCTCATTGGAAGCAAGTGTCCTGCAAT
AGTGGAGAAAATCTCGAATATGCGTACTATTGCCTTCAGCCACCACTTCTAGCAATGATATGTCATCAATTATTATCCACACA
TTCTTTCCTGAAGTCTGCTTCAAAATCTTCACAAAACTCTCGAAGAGCGGAAACAGTTTGTTTTCAACACTTGCAGCAGCCAC
ATTTGGGGACAAGCTGGATAGCTCAAGGATGGATTTTGACAGCAAATCAATGTAGACAAGAGAGCCACGCTCTCGATGCACC
ACCAAATTGCATCCCTGTTTTCGGGCAATTCTGGTGTAGTGAGAAAAAGGCTCCTTAAGGGCAACGAAGATGACAAGGGAGT
CGGGAACAGGCAAGATGTAGGAGAGGAAGGAGAGGAGGAGGAAGGAGGCAGGAGAAGTGAGCGAGTCGTTGATCAACACC
ACCTTGTTTCGTGGGGGGGCTCCTCCACTCCACCCCAATGCCTCGTGCAGCATGGACATGCTCTAAGGCCCCATTCAACGCTG
ATTGGTTGAGAGAGCACGTTGTTTTGTTAACTTTGAGTTGGGCGTGCTTGGAAGCCGGAAGGTTTTTTCTCCAAAAGCTGTCGT
GACTCAGCAGCCAGATCAGCTTGGATGCTGCCTTGTGTGAACTGGCAAGAAACAGGCAGCTCGGTTGATTGGTCCACGTCAG
CCAGATCAGCTTGGCTACCGCCTTGTGTGAACAGCCAATAATCAGGCGTCTCCCTTGATTGGTCCACATCATCATTTTCACCTT
GGCTGCCGCCCTGTGTGAACCAGGCCTTATATTGAATGTCACTATTGAAAGGACGACCTTTTTTAGGTTGGATGTATGCATTGT
GTAGGATTTCAATTTGTTTATATGTGGTTTTTGCTTTTTTTCTTTGTTGCGAGGGGGGGCAAATAAAAACACAGCTGCTAAAAC
ACGTCACTCCATTACCGCTCCATTAAAAAGTGCATTACAGGAAGTAGTAAATAGGAATAGGTGTGTGGAGACTTTACATTCAC
TCTGCTTGTATTGTCACTTTGAGAGTTTAAAAGATTATGTTGGTTGCCTCAGCCTTGTCATGAATATCCCCTATCAGATGGAGG
GGTAACCCATTGAGTGATGCACTCCATCAAAAAGGCAGGTTATCAATTTTGTGGATTATGGATTATGGGGTGACAAAGTTTGT
ACACAGTTTGGTGCGTCTTGTACAAATTGTAGACTATAATTATGCAAATCAGCCTGTGTATAGCTTTATCCCTATGGGAGGGA
ATGACCGTTTTTTCTTAGACCCCCAGTTTTCCAGTTTTTTATGCAGCAAATTAGGAAAAGCCCTTAAAAGTTTACTTAAATCTC
AAAAATAGCCTTGAAAAAATACACTAATGGTCGGTTTTTCATGAGCACTAAGTATTTTTAGTGGTCACACTATTATTTCATGGT
CACAATATCATTTCAGAAAATGACCGTTGCTGCTAAATTTACATGCATGAGGGAAAAATCCGTAGGCAGCCAACATGTGGAA
AGGCCAAAACTGCTATTAGATTTCAATATGTTATTGCCACTGGAAGGACCATTATACTGGGGTAAATTCCAGCATTAAGAGTA
TTTCAAATTCAAGGAGGTTGTCTTGTTAAAAGCACCAAGCTCGGAAATTGTGTGTGTGACTGGAGTATAGTACTAGCGGCACT
CAGCTCCTACGTGATGCTTCTAATCAATCCTATTCTTGTCTATAAATACCGAGAGAGTATTTTGTG

**NUCLEOSIDE DIPHOSPHATE KINASE 1 (*NDPK1*):**

TCCAAAAAGCTTATCATACCTCATTTCACACAATAGCACCATTAAACTACAGAACCAGGTTCTTCGGAAAATATGAAT
ATTTATAGCATTTAGAGCCTAAGACAGAGATGCATGGGCGGCACTGGATGATCTATGAATGCCATCAACATTCAGTCAATAA
CTAGACCCAAAACTCATACACCGAGTGCACAAGAGAAAAGAGAATCACTGGAACAACCTGGGCAGCTAATGATCAAACCCA
CATTGCATGATTAGGGTGATACAGAAGCATCGTTGTCATGCCCTAACTAATCGCAGATGGAAATACGAATACTATCTGCTCAT
TGTCACACATCATGCATCCTCTTAACAACAACTCTGTAGGAATACCTTATGCCATGCTGCTGCAGTTTTCAATAGTTAAATTCC
CAAAACAGTGAAACACTCTGCTCCATTTTTGGACAAATCTAATCCTTGAGCAGTCTTTACACATGAGTTTGACATAACCCCTG
TTGAAATCAGAGACTTCCTTTTGTTGCATCTCTATCCAGATGTATATGGCAAGTTTCTATGCACGCAGGCATGAGACAAGTGC
TTTCATTGCGCCTCTATCCAGATGTCTATGGCAAGTTTCTGTGCGTGCAGTCACGAAGCTAGTAGGCCAATACAAGAAAATTA
AACAACTCACTTTCTCGACACCTCCAGGCATGTACACGCCATAAAATATCAGTGGCCAACTTATAGTAAACCTCAGACAACAA
TCACTATGCGGTGCAATGACATCTTCGCTTATTCTCAAGCTCTTCAACTTGATAAGATCAAATATCATGGTGCCTTCTGAGCAT
CAAAATTGTGCATCTCAACGTATGGGACACTGATAGCACTGAAGGCACCAAGATTTCCACAAAAAATCCAAAAATGGAGCTC
ATATGATAACATTTTTTTAAAGAAATCTCACTAAAAGGTTACGTTATTCCCGTAGCCATGGAGTCAACGTTGGCTCCCAATCA
CATAATTCACTCTCTTTGAACCACAAACCAATTTCACGCTCCCCATTGTCGGGGCTGTCACTTCCATGAATAACATTCCTTCCA
GTTTCAATACCCAAATCCCCACGGATTGTTCCCGGCTCAGCTTGTAAGGGATTTGTAGAACCTATGAGTTTCCTAGCAGAAGC
CACCACACCTGGTCCTTCCAATGCCAGACACACAACGGGTCCTGATGTGATGTACTTCACTAGCTTTGGAAAAAAAGGTTTAG
AGCTCAGGTCTTTATAGTGTTCCTCTGCAAGCTCCGTTGAACAACTGAAAAGCTTGAGTCCTTTTATGACAAAACCTTTTCTTT
CGAATCTTGAGATAATCTCTCCCACCAGGCCACGTTGTACTCCATCTGGCTTTATCATGACATATGACCTTTCGGTTGAACCCT
GAGATGCGCGGACAATTGGGGAGAGGCGCAAGCAATGGTGGTGGAGAGAGGCAGAATGGGAGAAGCGAAGCTGCGAGCGG
AAGAGGCTGTGGGAGCTGGAGAGGGTGACACGAGCAGATCGGGCGAGGCTGGAGCCAAATGGCTGGAGGCCTGAAGAAGAT
GATGCTGAAGAAGACGTTGAAGAGGAGGAGAGGAGGGCAACAGAGCCTGCCATAGCCACAGAAGTGCAGGCCATGCCTCCT
GCCCCTCGACTATTTGCCAGCGAACTCCCTGGGAGGCGGTGCAAGGAGAGCCAGGGCAAGTGTGGTGCCTTGTCAGCAAGCT
AATTGGGGGCAAAGGCACCTG

**GLYOXYLATE REDUCTASE 2 (*GLYR2*):**

AAGGGCGAGAGCAACGAGGAGCGAGCCACCTCCACACCTCTGTACGATGGCGTCCCTCCATGTGAAGCTTCCGCTCC
TCGTTGCTCTCGCCCTTTGCTGCTCTCATTTGATAGCCTCTCCCACTCTGGCTGGCTACTACGGCGCCGATAGCGCCGGATGGA
GCAACGCCCACGCCACTTTCTACGGCGGCAGCGATGCTTCTGGCACCATGGGTGGTGCCTGTGGCTACGGCGACCTCTACTCA
TCGGGGTATGGCACGAACACGGCAGCGCTGAGCACGGCGCTGTTTAACGCAGGGCTGAGCTGCGGAGCATGCTTTGAGCTGA
GGTGCGACGAAGCAGCAGATGGCCAGTGGTGCGTGAAGGGGACATCCGTGATAATAACGGCAACGAATTTCTGTCCGCCAAA
CAACGCGCAGCCAAGCGATGATGGGGGGTGGTGCAATCCGCCGCGAGAGCATTTCGACATGGCGCAGCCGGCGTTCGAGAA
GATAGGGGTGTACAGAGGGGGGATAGTGCCGGTGCAGTATAGGAGAGTGGCGTGCGAGAAGAAGGGGGGAGTGCGGTTTAC
AATGAACGGGCACGCGTATTTCAATTTGTTGCTGGTGACGAATGTGGGCGGCGCAGGGGACGTGACGGCGGTGTCAGTGAAG
GGAGCGGGGACGGGGTGGCAGCAGCTCAGCCGCAATTGGGGCCAAAATTGGCAGAGCAACGCCCATCTGACAGGCCAAAGC
CTGTCGTTTCAGATCACAACCAGTGATGGCCGCCTTCTCACCTCCTACAACGTCGCCCCCGCCGACTGGCGCTACGGCCAAAC
CTTCAGTGGCCTCCAATTCTAGCTGCTCTCCTCCACGTTTCCGCCCTCATTACTCACATACAGAAGTGGCTGCCTGTAGCTCAA
CCCTAATTCTACTTCAGCCCGCTGCACCTTAATCTCCTTGGCCATATCTATATATATTTTTTATATTTCTCTATTGCTGGATGTA
ATCTGTAGTGTAAGTTGAAAACAAGTTAAAATGTTTAAATTGCCTTCCTTTTACAACCATTAACCTTAACCAACTGCGCTACTG
CCCGTTGATAATTGCTTGCATTCATTGAGTTAAGAGCACGTCTCTTTTGTAAGAGTAACTACCACGGCACAAGCCAATTCCAC
ATAGGCAGATATTTAGAAACAATTATCCCTCTTGGTGTTCTAAATATGGTCATTGTAGCATCAGAACAAAAGGGCTACTCCAG

CAAGACCTTCAAAAACCTCAAATAATTGTGTGGCGGTGTGGGTGTGTATAAGTGGACACCCTCAAGGCAAAGAATGCAGCCA
GTTCAACAGCAAAGGTGCAGTTTGTGCCTTTACAATCAAGCTATACCCCATTGCATGACTAGGTTTGTTTGATATCATTTTAGT
GTAATTGATCACTTCTGCCAAAAGAAGCTTTTAAAGCAAGCTCAAGAAACTGAGCAAAAATGTCAATCCACTAACACTTGTTA
CTAAAAGCAGCTTTAAAAGAAAACATTTTTGTGTAGGTGGTAGCCGCATTGCAACCTGTTGCGAAACTAATCTTCTTTGATTG
CTTCAATTACCGCAGAGAAGTCCTGATCGCCAAAGCCTTTAGACTTGGCCACTTTGTACATTTCATTTACAGATGCTGCAACC
GGAACTGGTTGGGACACAGACTCAGCCAGTTGCAATGCGAGTCGTAAATCCTTTTGTTGGTGTTTAAGAGGAAATGACGGGG
AGAAGCTCTTCTTGATCATTGAGGGACCCTTCATTGCAAACATTGGCGAGCTTATGGCACCTTGAGATATCACCTCAATTATA
GTGCTTGGATCCAATCCAATCTTGTCACCCAAAACAAGACCCTCTGAAAAAGATGCCATCATGCTTCCCATGACCATGTTGAC
AATCAACTTCATGGCTGCACCATTCCCAACTTCACCAAGGTAAAACTTTGACTTGCCCATTACATCCAGCAGAGGCCCAGCTT
GTTCATACAAAACCTTATCTCCGGCAGTAAGGAATATAAGAGTTCCATTTTCAGCAGGTTTCTTCGATCCTGAAACCGGTGCC
TCGAGAAATGACGCGCCGGTAGCTTTTACAAGGCCACAAATCACTTTTGAAGTTCCACCATCTACTGTTGACACATCCACGTA
CCCTTTACCAGGGCCCAAGCCATGAATGACACCATGCTTTCCTGAAGCTACTTCCAACGCAGCAACAGGATCGGCGAGCATG
GCAAATGTGATATTGCACTGTGAGGCAACCTCTGCCGGAGAACTACCACAACTGGCTCCTTGGTTGATCAAAGGCTCGCATTT
TTTTGCATTCCTATTCCATACTGATACATCATACCCGGCCTTGATGAGATTGGTTGCCATGGCAGTGCCCATAATGCCAAGGCC
AAGGAATCCAACGGTGGGTTTTGCAGTATCATCAGAAGAGGGGGAGGAGCTGCAACAGCGCGCCGATGAGGGCTTGCGTGC
CAGGGAGAAGAAGGGCGGGCACAATAGATGACGGCGAGAGGGTGCGAAGAAGAGTGAGCTCGAACACGCACACGCTATCT
GGTTGAGCTGAGCCATCGCTCGTTTTG

**GLUTAMATE RECEPTOR 3.4 (*GLR3.4*):**

TCTCTCTCTCTCTCTCTAAGCTCTTTTGTTCACAAAAATCTATGGAGTTTGATTGTCTATTCTCACTCCCATCATTTTGAT
TCTGCTGGCCTTAGTAGTAATTTTATTCAGAGTTATCTTTTTTTTTTTTGGCATCTTTGAATTTCCTTGTTTCAATACTTTACACTT
GCACTCCTACATCCATCAGCTTCTTAAGTTCTCTAAATTTCTGCAATTCGGCATGTTTCGTGCTACCGGGCAAGCCCACATCGG
CTTTTTGATCTTAATAATGAATTATGCTTTAGCTCTCTATAGAAATTTTTAATTCAATCCAATTTTCATTAATTAATCTTCAATT
TTGGCAGCTTCTTGTCATAAACCATGATTCAAGCTTCTTTGCATAAGCCCAGGATTCAAGCTTCTTAAGCCTTTGTCCTTCTGT
AAGGGCCGGATTTCAAATTGTGGGTTCAAATCCCTATGTTATTCACTTGGTTTCTTTCACAAAAATTGTGTGTTTTGCAGCTTT
TGATGCTCTGGTTATACACCTTACTCTGTTAGGAGTAAGTCATGCAATTTTTTCAGATCATTCTGATTTTGCACGGGTTAAGCC
GCAATTTGTACTCCCCACGCATGACTGCGCATTGTGCGCATTTGCAATAGGTCATTACATTACTCCAATCATGCTTCAAATTCT
CCACTTATATAGCCACTTTTGTATATAGATAGCAGGGTCTATAGGCTATGATGCAATATGTTAAACACTCTGTTAATGATGAG
TGCCATGCCCCTTCTACATTTAGCAGCCAGAGTTCTTTTTTTCCTATTCAGTTGATTTTCATTCTACAGCATGCTTTCTTCAAGA
TGACAAAATGAATCAAGATTGATGTCGCCGTGCGCACTCTCGTTTGCATATGGTTCACATCGAGAACATTGAGAGTATTCCTG
TGGGCAAAGCCATCATTGGCATGTAGTAGTAGCCTGCATGATCCACTTAGAGCTGCCAAGCTAATAGCTCAATTTCCTTAGAC
TAGATTACAGCTTAGTTTATTCACAACTTTTCACGAAAACTTCGTTTGCGACATCGCTACTTTGCGAGACAATCAAGCAGTCA
AGAATGCAATATCCTTTCTCTTTGTGTGTATTGAGTTGTATTCTGCTTACACTCACATGGTGTTGTATGAATGCTTTGGGTGCTT
CCAACATTACAAATGGGCCTCCTGCAAGTGTGAAGATAGGGGCTCTGCTTGCTTTCAACTCGACAATTGAACAGTTTGCACGG
CGGGCTATACTGCTCGCAGTCCAGGATGTGAACAAAGCAGAAGATGTTCTAAATGGAACACAGCTTATCTTGGAGATGTTTG
ATAGCGGCTACAACAATCCAATACAAGGAGCTGCTTCTGCTGTGGAGCTTGTGAAGAAAGGCGTTGTGGCTGTTGTAGGTCCT
GAATTTACAGCAGTTTCGCAGTTTGCAGCTCATATAGGACAAGAAACAACTATCCCCTTTGTCTCATTTGGTGCAACAGATCC
AAACCTCTCAGAGATGCAATACCCTTATTTCATGCGTGTTGCACCTAGTGATGTGCTACAGATGGAAGCAGTTGCAGCCTTCA
TTGCCAATTATGGATGGAGAGAAGTGGTGGTGTTTCACATGGATGATGACTACGGTACAAATGGAGCCTCTGCCTTAAGCGAT
TCACTGCAAGTTAGAGGTAGCAAAATTGTGGACAAAGTAGCTTTTGTTCCAGGGATAGATAAGCCTGGAATTGACAAGGAGC
TTGCCCGGTTAGCTAATAAGCAGGCACGAATATTTGTCGTGCATACTCATCAAGATGTGGGTTTTCAGATCCTCCTAGAAGCG
TACTATTTAAACATGATGGTTTCTGGATATGTATGGATTGTGAATGATTTGATCACCACCCATCTAGGAGATCTGCACCTTGAT
GTGCGAATACCGCGATATACCAAGGGGGTCATTGGAGTACGGCGGCATGTAATGGACTCTCCACAGTTGGATAGCTTTTTGGT
GGAATGGAGGAGGACTTATCCTAATCAGTCACTTGATTTGGAGGCAAGCCAGATCAATGCCTACAGATTATATGCTTATGATG
CTGTCTGGGTAGTTGCTCGAGCCATTTCATCTTATTTAAATGACAAACATGAGATTAGCTTCCAGAACCCCCCTAAACTGCCC
ATTTATTCAGGGGGCGAGTCTGAACTAGCTCAAATGAGGAGATTTGTAGGAGGGGCAGTAATGAGGAATTATGTCTTGACTA
CAAAGTTTTTGGGTGCGTCTGGACTCGTAAAGTTTGATAAAAAAGGAGACCTTGTAGGTGCCGCTTTTGAGTACATAAACATG
GTGAGTAGAAGCCCCCATGTGGTGGGATACTGGTGGGCAAACAACTCTGGTGTCTCTTTAAATCCCCCTCCCAATCATGAGAT
TTCAAGTCTGAGTAGTGACACTTCTGATGATGAATTTTTCTTATCTCATGATGGCAAACCAGCAAATGCTACCAGAATGGCTA
TCATCTGGCCAGGCATGGCCACCACGGCACCACGGGGTTGGGTGCTCCCTAAGAATGGCAATCCTCTTAAAATTGGTGTCCCT
CGAAAAGCAGGTTACAACGAGCTGGTTAGCATTACTATTGGTGCTGACAATGTGACTGCGTATAGCGGGTTCTGCATACAAGT
GTTTGAAGCTGCCCTCAAGAACCTTCGGTACGCTGTGCCTTACAACTATGAAATGCTTGGTGATGGGATTACAACACCAGAAT
ACAACAAGTTGATTCTGAAGCTAGCTGACCAGGAATATGATGCTGTAGTTGGAGATGTAGCAGTCCTCGCGGATCGTCTTAGA
ATAGTTGATTTTACTCAACCATTCATAGAGTCTGGGCTAGTGGTGCTGGTACCTGTCAAAAACAACAGGGAAAGTAGTCCTTG
GGCTTTTCTCAGACCATTTACACTATACTTGTGGTTGACTGTGCTGTTATCATTTGTATTTACTGGCGCAGTCATTTGGATACTT
GAGCACAAAGTAAATGAAGATTTTCGAGGTCCTCCAAGGGCACAATTAGTTACTATGCTGACGTTCATCTTCTCCACCTTGTT
CTTTGCACATAGGGAAGACACTCGAAGCGTTTTAGGTCGATTTGTTTTGATTGTCTGGCTATTCGTGATCCTAATCATCAATTC
CAGCTACACAGCAAACCTTACCTCTATTTTAACAATTGAACAGCTAGCTCCAACCATTCAAGGGCTTGATAGCTTAATCCAAA
CAAATCTGCCCATAGGGTATCAAACTGGATCATTCGTTCGAGATTACTTAATAGGTCTCCATGTGAATCCAGGAAGACTGAAG
GAGCTAAGTTCTCGTGAAATGTATCAGAAAGCATTGGAGGCAGGGCCTTATAATGGGGGTGTTGCGGCAATCGTGGATGAGC
TTCCCTATGTGCAGCTGTTTCAAGAGTCTGATTGCAAACAATATATCATTGCTGGCCAGGTGTTCACAAAAAGCGGATGGGGA
TTTGCGTTTCCACGGGGATCCGATATAACAGCGGACTTGTCGCAAGCCATACTGGAGATGTCACAGTCGGGGGAGCTGCAGC
ACATAAGGGATTATTGGTTCCGGGATGTAGTATGCGAGGAGGTAGCGAGGAATGCCGCAACGGGATCAGGCAAGCTTGATTT
GGGAAGCTTCTGGGGTCTCTTTCTTATATCAGGGGTGGCATCAACAATATGTGTGTGTGTACATTTGCTGGTTTTATTGAAGAA
GTACAAGGAGCATAATAGGCTACAACGGAAGGAGGAGAGCATCCTCGTGAAGAATTCAGGTAGATGGGAGCATGTGAGACA
ATTCATGTCATTTGCAGACAAATCTTCTCATGGGTGCAAGCATGTGAGCAAGTATGATGCCTCCTTCAAAAACCATGCAGCTA
GTAGCTCTACATCTTCTTCCACTTGCATGACTAACTCTTGCAGTTAACTATATATATATATATCAAGTTGTGCATTCAAAA

**PPHB SUSCEPTIBLE 2 (*PBS2*):**

CAACATTATATCCAACATTACAAAGCCCGATTCGAACTCACACCATGAGGCTATGATTCCTTTTGGTATCTGTGTGTAC
TCTAGGTTAGTACGTGCGAGGAGATGACAGAGCGATGCCAGAGAATCGGTTGCAACGCCAGTTTCTCTGCAGATAACAACGC
CGACGATAGCTGCCGCTATCACCCCGGGCCTCCTTTGTTTCATGATGGTGGAAAGGAATGGAAATGCTGCAAGCAACGGAGC
CATGACTTTTCTCTGTTCATGGATATTCCTGGATGCACAACTGGCAGGCATACTTCTGAAAAACCTCAAACAACATTTACCTCT
TCAGCTAAGAAACCTGTCATGGCACCGGTTGGGCACAGAAGTTTTACCACCGTAAGCAGGGAAGCTTGTCCTAGGTGTAGAC
AGGGGTTCTTCTGCTCTGACCATGGTGCACAACCAGCAGTGAATACGTCTGTGCGAACCCAAGCATCAGCAACAATTGTTGAC
CCAGTTGTAGAGAGTAGCGAAAATTTACAAGAAAAAGTTAATGAAGAACCGAGAGTGGTGGACTTCAGTGTAGAGCAAACTT

GTAAGAACTGCAAGAAAACTTTCACAGAGAAGGAAAACTACGACACAGCATGTGATTATCACCCAGGGCCTCCTGTTTTCCA
TGACCGATCTCGGGGTTGGGCATGCTGCAACGTTCATGTAAAGGAATTCGAAGAGTTTTTGGAAATTCCTCCCTGCAAAAAAG
GATGGCACCGTGGAAACTTGTGAAGCATATTGAGCCTTCCAAGATTCTTCTCTTTTTTACAATGAATTGACTGTACACCATAA
AGACAACTGCTTAGCATTAGGAACTTTTTCAAGGTCTTTCTTGCTTTGGAGTTTCCACATGCCTACCAATGATCATCATTGTAA
GAACAATTTTGAAGCTGTTGGGCATGATATAGGAGAGCTTAGTTGGAGCTTGTCTCTGTTTTTCTTTTGTGACTATGTCGAAAC
CTACGGAATTTCTCTCAAGCT

**GLUTAMATE-CYSTEINE LIGASE (*GSH1*):**

GAGGTTAAATGTACACATTTAAGGGAATTTACCAAGAAAGGATTCCAAGGAACAAACCTTTATTTTTATTTGATTGTG
AAATTTTTATTTTATCCCAAAAGCCTTGTAGTCCTCTGAACAAAGATTGGTCTACAAGCCTTGGGGTCAATATGTGATCTAGA
ACCTTAGTAGTTCAAAGGCAGGGTCTACACTTTCCTTCCAACTACCTCTGTACAGCTCCAAAAGTCTCTCTGCTTGAGTCACAC
CTGTATTGGCAATTTCAACAACTTCGTCAAGAAAATGGGCCTCCCCATACCCTCTTCTTTGCAAACCCTCCTGTGCCAAGTTGA
CCACATCTTGTGCCACATGTTTTAGAAGAGAGTCTCTAAAAGGAGTCTTCAATCCCAATGTTGGTACCTGGTTTCTCAACAGC
AGGTGCTCATCCAAGGTCCAATCCCGTATTAAATCCAATGCTGCTTGAAGTGATATGTCATCATATAGTAGACCAACCTGCAC
AAACAAGTTTTTCAAAAGGAGATCCCAAAGGATTACAAGTAACAAGTTCATCTATAACTTTTTCTTTTTTATTCATGATTTTAG
GTAAGGGCCTAGAAAGTGGATCCCTGAGATTTCCATCCACACACACCCAACCCACGACAGGTTTCAAGCTCAGGACCTCGTC
CTTTCCAGCTCTACAACCAAGTCATCACGTGGTGCTCCCATGGACAGGTTCACCTGGAGCTTAAATATGTACCCAAAAAGCTG
GCACTGCACATAGTAGACTCCTTGGACCTGCATCAGCTCCTCTAAATTCCAGAAACTTTTTGAGCCTCACCTCTGTATAAATTG
TAGCAATATGAGATTCCCAGTCTTTGAAAGTAGCATTCTCACCAGGCAAATTAGGTAAATTTCCCACAATAAAGTCCTTGAAA
GACATGCCACAACAGTTGAACAAGTGATTATTGCGATACACCCAATACAAAGGAACATTCAAAGCATATTCAACATATCTCT
CGAATCTGTATACTAACAAAAGAAAACACTTAGACCCCATTGACCAACCCCTACTGTGGCTTTTTACATTGTCCTTAAACAGG
TATCATGAGGTACTCTTTCAATAACGACGCATCGGCAAACCCTGATTGCAGACGTACTCCTACTTCCCTTGAATATCTTTCCTA
CTGGTGTTACAAGGGCACATTAATTAAATGGTTAAAATGCCGGTTTGAGAAGCACATAATGTCATTAATCTTTAAGTAAAACC
CTGTACTGGCCTAGATACCCTGTACTGGCCTAGATATACAAATTGTGAGAGACCTGATTTCAGCAGAATATGAAAGAAAGGT
CAGTGCTGAGCCTTGTACCCCGTCTTTTTAGCTTTAGTATTGTGTTAATAGTAATGCTCTGAAGCTTTTCTGCATTAGCCTGGA
AGACGCTATGAAGAGTCTAGCATCTGCTTGAGCTGGCCAATAGAGCAAACACAAGGTTGAAGATTGGAAAATATTTAAATTA
TCTCTCTTCCATATTTAAAGCAATTTATACATTAAGAACATGTCATTCCTAGTTAGAACACCGCTTGGGCATACTATGGACACA
AAGGTTTCGAACCTTGTGTTTGCAGACCAGAAAATAAATCCAAGTAAATACAAGTTCGCGTATGTAAAAGTTAGAACACTTGT
GTAAGCATCATAAGACAAATAAATCAATCTTAGAGACTGGAATTCTGGCTGTAGATCAAACTCATCACAAAGCTGTCAATAT
GAACCAAAGCAAGTCTGCTGTAAGTCATCAACATATAGTAATAGCATGCAAATCAGCTTGAAAGTTATGCTCCTCAAATCAC
GTTTCTTTTTTTCCGAAAACTCTCACACCCCGGCGCCACATTTGTGGCTTTCTAAAAATCAAGAAAAATAACGCAAATGCACA
AAGGATATCACAAAGAATACACCTAAGAAAATGAAGCTCCAGATTTTACCCAAAGTTGTCATCGAAAACAAATGGTAGTATG
CCAGTCCGATTGTTATCCAGGTCCTTATATAGTTCACTTCTGTGGCTTAAAAGTCCACTAGGCTTCCCATCTGCGAAAGGAGA
GTTTGCAAAAAGAGCTATTGCAATAGGTTGTAGTGCAAGTCCAAGTCGAAATTTGTTCATCATATCTTTCTCAGAACTGAAGT
CCAAGTTCACCTGAGTAGAGCATGCTCGAAACAGCACTTCATTTCCAGTAGTACCAACTTTGGGAAAATATTCATGCATCATT
CTGCATTTTTTCTTAGGCATCAACGGAAATGCATCAACTGGCCACTTGGGATTAAAACCCATCCCACATAAGCCAAGCCCCAG
CTCATCTGCTACAGATTTCACCTCGAAAATATAGGAGTTTATTTCTGCCTGTGATTGATGTAAGGTTTCAAAAGGGGCACCAC
TAAGTTCAACTTGGCCGCCACCCTCCAAAGTCAAACTCTTTCTGTCCTTTAGTAGTCCAACTAATAATCCATCCTCCTTGATTC
TGTCCCAGTTGAAACGCCCGGCCATGCCT

**DICER-LIKE 4 (*DCL4*):**

GAGTAAATTATGAGGAAGAGGAACTAAGGTGATAATAATTATGTACTATGATAAGGACAGCTTTCAGGCTCTATTCAT
TAATTGGGTTTAAACAGGCATGCCAACTAAGGTGCATGTCTTTTGGATGGCAAAGTGAGATAGGCTTCACAAAAAGCCACAA
GAAGCCAGCCAAGTCAGGAGTTCTGCTGCTGCTGAATCCTTGGCTGCCTTAGCGGATTTCCTTGGATCTCCACGACATTCGAC
GAGTCCCACATCTGGCAGCTGCAGTACGGCATCGTATGTGAAACACTTGTCATGGGGTGGACCCTCCTCGGAGGAGCAAGTG
AATGATGGGGGCTCCCATCTTTTTCTTGAGCAAGCTTCATTCAGGGATGCTCTTGCAAGGCCCCTTGATGCTGCTTGCGTTGTC
GTGTCGATTGTACATGCAGGGACAGATGCTAAATTGGCAAACTGAAAGTTAGTTTCATTTCTGATACCAACACCATTAGTTTT
CATTAGCCATGAATTCTCCATAGACATATTTGAGCGCTGAAAGTAACTTCCTGGCTCCTCTGCTATTAATCTGGATGTACTTTT
TTGCTGATCTTTGATTTCATAAAGAAGTCCTTTTCCATTATCTTCAATAAATTTTTTGGCCTCATGATCCATGATTGATGCTGCA
CATTTCTCTTCCCTAGTCATTCCTACTCTAAATTGTGGCTCATTTACATTGCTTACTCCCAATATTTTTTCAAATCTGTTACTAAT
AGCACTTATACTGGCTATATGGTCCTGTATATGGTGAGTGCTACTGCCTGCACCAGTTGCTAAAAAGCCAGAGCTCAAAGCAG
TTGATGACAATGTGGCAAAGGAGCTCACTCTAAATTGTGGCTCATCTACATTGCTAGCTCCCAATATTGTTTCTGATGGCTCCT
TCGCCAATGTGGCAAAGGAGCCCACTCCAGATTGTGGCTCATTTACATTGCTAAATTTCCGTATTTCATCAAATTTCGTGCTGC
TGCCGGCACCAGTTGCTAAATATTCCATGTTCAAATCAGTTGGCGACAATGGTGCGCTGGGGATCATGTTTTGGGGGGTGTCA
CTTATTTTGAACTTGTCTGACGGCTCCTTTGTGGCATCCCTGCTCTTTGCTTTCTGAGAGTCTAGAGTCTCACCCTTGCATGAGT
GAACATAACCTAGAACCTTCAGTTCCCTAAGAGCGTTAATTGCAGCTCGTTTCTTTGCAGACTTCTTATCTTTAGATACAGAAG
TTCCCTTGACAATATTATTGCTCACATTGATTTCGTAAGTGTATTGGCATTGCCGGCCACCAAGAGACGATGAACTTTTAGTCC
ACGTCAAGTGATTCTTTTGGCATACTTCTTGCAGCTCACGAACAGGATGAAGACGTACAAGGCGACTTGAGATCAGACGCCCC
AATATTGGCTCAAACACTTTCCAGACCAGGCTCAGGTTAAACTCTCCATCCACACATATAGCGCCACTCAAGGACTCCAGAAT
GTCAGCAAGCACCTTCGAGCACTTGTCTCCTTCCCGGTCTCCAATAAGTTCCTGCTCTGAGGAAGCACAGATGTAGGAAATAA
ATTCATTTATGCCTTTCCGAAGTTCTGTAGAGTTCTCTATAAGATATGCATAAAGTTTATGCTGAATTGCAATCCTGGAAAAGC
TTTCGTTGCTCACTATTGTCGACCTCAAGTCAGTAAGCTCTCCCGGTTTCGAATCCTTGAATTCTCTGTAAAGGTGCTTTGTGA
TAAGGAAATCCAAAACAGAGTCACCAAGAAACTCCAAGCGCTGATAGCATTTCCCGAGATGATTGGTAAACGAAGCATGTGT
GAAGGCTTCTATTAGAAGCCCTTTGTGCATAAACCTATAGGACAGAAGCTTCTCGATTGCTTCAATATCAATTCTTCTGATCAA
CGAAAGATTGTTCTTGCTTGATGACTGAATCTTGACAATTTGTGAGATGTCTGTGGCTACTTCCAGTCCAATAAATTGCATGAA
TGATACCGCAGCCTGCTCACCACCATCTTCTAGGTAAGCACCAATCAATGCTTCAATAACATCTGCAACAGTTTTCCTCTGCAT
CCATCTGTGTCGCTTGTTACATGTCACAGAAGCCTTCCATTCTTTGCTTTCACTGGAGTCTCCGTGTAGGTCCCCAAGTAAACT
TTCACCACACACAGCTTTGCTGGGATGGCAAGGTGCTACCCAATGCTTAGGATCAAATAAGGTATCACGAATATATCCAGCA
AGACCAAGTTTGCTTCCAAGTGTGAACAGTGAAGAATTGCAGATTCTCTGATTACGTTGCAAACTCAGAAAACCTTCATCCAC
TTCCTCATGCTCAAGGAATAGACGACGACTGATGGCATACTTCAAGAAAGAGTCTCCAAGCAGCTCCAATCGTTCTAAAGAA
AATGAATCTAGGCATTTCTCTGTCGTTAGTGCTTCCAGAATCTTTTTTATGGAGACCTGAGACCCTTGCGGAAACTGTGAGCTG
AGAAAATTGCGGAATTGAGTCGCAACAAGAATGCTTTCAAGTTTGTGCAGCAGGGATGGCAGAATAACTGCAGCATTAACCA
AATCATCATAAAACCCAGAAAATTCGATTCTGCAGACCTCCGGAGGTAGCTCTACTAATGACTTTTCTGGCTCCACTGACACT
TTTTCTGCTTCCTTCAACTTTTCTGGTGTAGCCTTGCTGCCCTGATCTGGTGACAGATTCAAAACCTTTTCTCTTGACCTAGCCA
GTAAGAAATTGTGAGCGTCTCTTAGAAACCTAGACTTTACCAAGTGTTGTCCTTTAAGAACTATGTCATGGTCATACATGGTC
TTAAAATACTCCACATAAGTTTTGTATCGTGCATTTGGAAAAGAACTCTCTGCTCTCAGATCTGTGAACATCTCAATGACGCA

GTACAGATGTCTGCCATGAAGAGTCTTTACTAACAAACCAGGCACATCCTCTATACGTATAACACCGTTGGAGAAATATAAA
GCAGCACCATCGACTTTGGGGCAAACATTCTCTGTCACCCCAAAGGTGTAATTTGAATCTTCAAAAGGTTTTAGTTTGGAAAT
TCTAGTCCAATCAATTTCTGGCTCCTTGTTGGAATTGATCGGTTTGAGTGGCAATAAAAGATACCAGGAACCATCTGATATAG
GAGGCTCTTTGGTTTTTGAATGCTCCAAATCACGATCCAGAAGAACAGAGAACAATATTGATTGAAATTGCTCTGCTTGTGCT
ACCTTTTGTAAGAAAAAGATTCGTCAACTTATCCCTACTTTCATCATAGGATACGCTCCTTATGTTTTTGCAACTGTTTGTCTA
GCATATCTCTGTGAAGTCCGTCACCCTCTCGCATATCATTAGCGTACGTCTGTGGAGCCCGTCACCTTGTGCGACCGTGCTGTG
GAGCCGCCCTAAGCTCTGTCAGCAGCGCGCGCACTACCTGCAAAGCCTTAGGCTGCTGGTGGTCGCATCCTTGCCTGGTTCAC
AGTCTGAACAGTGGGCCTTTTTTTTGCCTCGCATTGCTCTCAAGGCCCACCCCCATCTAGATGAAGTGACCTCGCCACAAGCT
GCCGCAGCTATCACGCGACCTTGTCACCTGAAAGGGATCAAGAAAGACATTTTTGACAGGTTCAAGCTTGCATAAAACAGCC
CTTCCATGACGCAATTGGAGGGTTTCATGGAAGTTAGCAGCCCCTTCGGGTAACGGTGATTGAAGCAGCAAAACAAAGCTTA
CGTACTCCCTGTCACTAGGAATAGCATCGAAAGATATTCTGTAAGCCTGCAAGCACATTGCACTAGAACTACTTGTGGGTTGC
CGAAGCCACACGTCTGGTATCACAGTCTCCTGTAGCTCCTCAGTTGTTGCTTTGACAATGCTTGAATGATGATCTGTTTCAGCA
TCTTCTTCTACTGTGGTTGTTTGTGGTAAGAGATAATCTGTTAAGGCTCCCTTTGAATGTAGTAATCTGCATGCTTTCAAGCAG
GCTGCCTTTTTGGCTGCAGCTTCTGAGTCACAAATTTCTCCTTCAACCAGTCTTAGGCAGGCGTTTGAAGGAAGTGTAATGCTG
CATTTGATTCCATCTTGTTCTTTTAAAAAATGAAATGATGGCTTTGGCTGATAAAATTCATCACTTGGAAGCTTGGAGCAATAG
CGATGAAGTAATTGAACACTGGAATGAGTGTTGACAATAGCTCCCGTGCTCTCCACTTCATATAAATCAAAATTTTCTCGCTG
CCTCTGATTTGTAGATGGTAAATGCCCTTGTGAAATTTTCTCTTTGACAAAGCTTTCGCTGTGGACCAAGGTATCTAATAACGT
TGCTTCCGAGATGTTTGTTCTATCCTTAAGAATAACATAATGAGA

**CHITINASE A (*CHIA*):**

GTTTGCTTGAGATGGCGCGGTTCAACACAATTATAAGCTGCACCATCCTGCTCTTGACATTTTGCAGCCTCACCGGCTT
CACGGTGGCGGCCGGCAACCTGGTGACCTATTGGGGCCAGGGGGGGGGCACGGATGGCATCGAGGGGACCTTAGCAGAGGC
GTGCCAGTCCAACCTCTACAATACCCTCATCATCTCCTTCCTCGACGTCTTCGGCCAGGGCCAGCAGCCAAAGCTCGACCTGG
CAAACCACTGCGACCCTGAATCAGGCACCTGCACCAGCCTCTCCAACAACATTTCGACTTGCCAAAGCCTGAGCATAACAGT
ACTCCTCTCTATCGGCGGCGCGAACGGCACCTACGGCCTCTCTTCTGCTGATGACGCCGCCACTGTTGCTAGCTACATCTGGA
ACAATTACCTGGGCGGTCAGCAGTCCAGCAGCCGCCCTCTCGGACCGGCTGTCCTGGATGGTGTTGACTTCGACATTGAGACA
GGCACCGGTGCCTCTTACTACGGCAGCCT

**NECROTIC SPOTTED LESIONS 1 (*NSL1*):**

CGGGCCGCGAGCTCTCTCTGATTAGTACGGGCGGGCCGGCTTTTTGTCTGCTCCCGCTCCTCACCCTCTTTCACTTTCA
AAGTTAAGCATCTTCTCCTCTTCTCTCTTAACAATTCTGTGTATGCTCCTTCTGCACGTCGCCTCACGCTGGCTCAAGCTTAACG
AAACGGCTTTGACCCCCTCATGTTCATTTTTTGCAGATTGCACATATTCATATTGGGTTGTTTAGTTTGAGCTTTCGCAGCCCTT
CTCCTTGCCTCTTCACATTCCTTGTGTCATGGTGCCATAATTGGCGATTGTACCCCCCCAAAAAGACAATCCTGAATGAATACA
GACTGATTTGAAGATCTGTGTAGGGAAAGGATGTGCTAAATGGCTTCCAAGCTTGACATCCCAAATGCAGCTAAAGTCGCTA
ACACTGCCATCCAGTCTCTAGGTCGAGGTTATGATCTCACATGTGACTTAAGGTTTTCTACTTGCAAAGATATTGATGGCACAT
CCCTAATTGAGCTGAGAAACAATGAGACGGTTGAGCTTTCGCTTCCGGGGGGCGTGGTTGTCGCAAATGTGCCTGCAATCGTC
AAGTGTGACAAGGGTGAACGGACTCGTTTCCGGTCAGACATCCTCACTTTTGATGAGATGTCTCAACAGTTTAATCAGGGCCT
CTCACTATCTGGTAAAATTCCATGTGGTCTCTTCAATTACATGTTCAATTTCACCGGGTCATGGCAGAAGGATGCATCAACAA
CAAGGCACCTTGCTTTAGATGGTTGGTTTTACACATTGTATACTGTGGAGATGCCAAGATCTCAACTTGTCTTGAAGGAAGAC
ATAAAAGCATCAGTTCCAACATCCTGGGAACCAGCTGCTTTGGCCAGGTTTATTGAGACTTTTGGGACCCACATAATTGTAGG
AGTCAAAATCGGAGGGAAAGATGTTGTTTACATGAAACAGCATCAGTCATCACCGTCAACATCTGTTGACTTTCAGAAATTAC
TTGCTGAGGTGTCAGATGAACGATTTTTGCAGACTGAAGGTCGTACCAGTGTGGGCTCGAAGGACAGCCGCATCAACAAAAA
GCGTAGCGTAGAGTTTCAGTCATGGTCAGCTCATTTGGACACCTTCAGTCAGATAATCCACAATGATAAACATCATGTGACCA
TTATACCAAGAAGGAAGGGCGGTTTTGACCATGGTCAAAGCCATTCCGAATGGATTCATACTGTCCCTCTTGCCCCTGACGTG
ATTTCCGTGAGTTTGATACCCATTACCTTCCTACTCAATGGCGTTGCTGGGAGTGGTTTTTTGAGTCATGCTGTAAACCTCTAC
TTACGCTATAAACCGCCTATTGAAGAGCTGCGTCAGTTTTTGGAGTTTCAGCTCCCCCGTGAGTGGGCCCCAGTTTTCAGTGA
ATTGCCTTTAACTTTGTCTAGGAGAGAGCACAGCCCTTCAACTTTGCAGTTCACTTTGATGGGTCCTAAGCTTAAAGTTAGTAA
ATTGCAGGTGACAGTTGGGAAAAAACCGGTAACAGGAATGCGCTTATTCCTGGAGGGCAAGAGGTGCGACAAACTTGCAATT
CATCTTCAGCACCTGTCGGCTATTCCCAGATTTCTACAACCTCTGTGGGAAGATCATCCATTTGCAAGAGAGGTCACATGGCA
AGATCCAGATGACTACAGTACAAAATATTTTGAGCCAGTACAGTGGAAGAATTTTTCGCACGTCTGCACAGCACCTGTGGAG
AACACGGAGACATGGATAGGGGACCATGCAGGTGCATCTGTTGTTGTTGGAGCGCAGCTTGCAGTGAAGAGCTTTGGGCTGA
GAAATGTGCTGTATTTGAGGTTGCTTTACTCAAAGATACCAGAAGCTAAGATTCGGAGGTCCGAGTGGGATCACATGCCAGCT
TCTACACAAAAATCTGGAATGTTCTCGAGCTTTTTAAGTACCACTTTCTCATCCACACCGATGCCTACAAAGTATCCAGCTGTG
GTTATAAACTCCGGCATATTTCCAGAAGGCCCACCTAAACCTGTACACAATCATAAGCTTTTGAAATTTGTTGATACGACTGA
AATGACCAAGGGCCCTCAGGATATGCCTGGGCATTGGCTAGTGACTGGGGCCAAGCTCTACCTGGATAAGAGAAAAATCTCT
TTACGAGTCAAATACTCTCTTCTGACTGTTAGCTCAGATTGACTCTAAATCACAGCCTTGTAATTTTCGACTCTATGGGAGTCC
TAAGAGGAAGGACTCCCAAATCATAGAAGTGAGGTCAACTAGTCGTTACACATCGGCGCTGCTCTGTACAGTTTTTGCAGAAT
GCTCAAATAGAAAGTTACAGTGGAGGTTGGGAGCTGAGTAAGCACTTGTGCAGAAGCACCATAAAGTGCCCAACAACAATCA
GTGTTGTGACCTAATACTCATCAACGCAGTGGCGGTGAGTGAGGCTTAGAAACATTATCCAATCTTCTGTAAATTTTGTAAAA
CGCATGTATATGGATGCACATAGTTTTTTACAGTTGTGTGGCTTTTAGAAGTTCAATGTGTGTTAGTCCCAGCTTTGAAGCATC
CAGCTTACATGGTGAGTAATGTTCAGGGCAGGATTGATGCTCCGCTCATTTGTACTTAAAAGGCTGGCTACCCTTCAGAGTGT
GTGCTTGACGATTCGGGTGGACCAAGAAGTTCAAAGCTTCACAAAATTTCTGATGCATAGTAATGCTGGGGAAGACTAGACC
TTAAGCAAGGTGAAATATTGTAGCACAGAGAATTTGAATGCATATCATCATTTTGTTCAGTTCATTGGAGGAGCTTGGGCTCA
TCTCTTGACATCAGACAAGCGCTGGTCCGAGGTCGTTGAGGCGATGGTCTGAATTCATATCCTCTAGACCTTAACAACTAAGC
AAAGTGGCCAGAGAACTACATATTTTTGTGTGAAAATGGTAACCTTTTTTATTTGGCATCGAAAAAACAGTGTACTGCAAAAG
AGGAGTACAGAGCATCCAAGCCAACTGCGGCGGAAGAAACACAATTTCTCAGCACGAGCTGGTATGACAAGAAAGGACTAG
AGTGTGCTGTGCATCATCCAAAGAAAAACAAAACAAAACAAAACAATCCTTGCCGTTTTATGTAGTTATTTTGCATATTGCAA
CACATGCCTTTGCAAGGTTGTTCAAGGTGCTCAGTATAGTGGAGGTATTGAAATGTGCAAATGGCTCAAATGAAGCACAAGA
AGTTTCAGAAAATTTCCTTTGTGGATTTTGACTACTAGGGAAAAGCAAAGGGAAAGGGGCCTCTTTCTCACAAGGAAAAAAT
CCAAAACTTCATGTGTATATACCAAATTTACAAGGGGGTGTCATGAAGAGAGAGACACAGAGGACCTCAAGTGATGAGAGCT
TAGAAGTTAAGGAAGGAGATGGAGTCCAAGGCTTGAGCGAAGAGTTTGATATGGAATGAGATGATGAGCATGATGGAGGAG
GTGGTGAGGATTGAGTTTCAATTTTGATATAGAAGGAGATGAGGGAAAAAAGTATTGCAAGATGCCTGAGGGTCAAACAAGC
CCCTACATAAAGAAAAACGAGTTTAAAATGATAAAAAACAAGATGATATTCCACACTCAACTTGCCATTGCGGTCAAATCGT
CTTGCAACCAACCGTACGTTACATGGACAAGCTACAACGTGTCTAAAGAACAGTATCTTTGTGGCAACCATCACAAGAAGTTT
TGGACATTTTTC

**FATTY ACID AMIDE HYDROLASE (*FAAH*):**

TCTCTCTCTCTCTCTCTCTCTTTGTGGCCAAGATAAATGTAGCTGGTATAGAAGGGCAAACAAAGGTTTGCTTTCTTTGAT
AGGCACACGTGCGTGCACATGCATGCGCCTGTGTGCACGCGTGCGCTTGCGCAGCTCTTAAAGGATGCACGCACGTGCGTGC
GCGCGCACCCACACCCACACCCACACTCACACAGACAAAAGCATAGCTTCTTCCTTAACCACACAGATTGCCGGGCTTTGAAC
AAGATTACTATCAAGATCAATTTAGAGCACGGATGGATCGTATCAGATGTGTGTTCATTACAAAGGGTTACTACCAAGGATTG
TAGAGGAGCATACAAGGCACTTGGCATGAATGTCTGAAGAGCTAAGCAATGGGTCAGAAGTCTATGCCCCTTTTGGAGGATG
TTAAAATTGATGCTGTGAAGTATCACCGTCATCAGTTAGAAGCTCCTCACCTTACTGGTTATGGCCTCAAGATTTCTACGTGGT
TGCTGGAGAGTTATGTTGTTGGTGGCTTGATTATTTCAAACACATTACGTGCAAATAAGTACACAGAGAAACTATCCCAAATT
GTGATACCTGATGCGCCATTGTTTGCACCTCAATTTCCTGAGCAAGATCTGGAGCCCGCTGTCAAGGCAGTTGAAGAGACAAG
TTCACCTGTGAAAAGAGTTGAATTAGCTCTGGAATGTCTCCCTCCTGGCTTCAATAAAAGATCTGATGTTCCATTTCAATATTG
GACTATCCGGGATTATGCTCATGCTTATCGCAATGGATCTACTACCCCTTCAGATGTTGCAAGGCGTCTTATTGCGGCCATAG
AGGATTCTCAAACTAGAGACCCTCCCATGTCCTTATTTATTTCATTTGATGCCAAAGACATTGAGAAGCAAGCAGCAGAGTCA
ACACACCGTTTTTCTGAGGGGCGTCCTCTTTCTATATTAGATGGAATTTTTATGGCAGTCAAAGATGAGGTTGATTGCCTTCCG
TATGGCTCCCGGGGTGGCACAACTTGGTTTCACAAAGTAAGGGAGGTAAAAAACGATGCCGTTTGTGTTGCAAGGCTGCGTG
AATGTGGAGTAATTTTTGTTGGCAAAACCAATCAGCATGAGCTTGGTTCAGGGACAACAGGCAACAATCCTCATTATGGGAC
GGCTAGAAATCCCCATGATCCAAAAAGATATACAGGAGGCTCTTCTGCTGGATCAGCGGCATTGGTTGCTTGTGGACTGTGTC
CAGCAGCTTTAGGAACAGATGGTGGAGGGTCTGTTCGCATTCCCTCTGCATTATGTGGCATTGTTGGCTTGAAAACAACGCAT
GGGCGAACTTCGACTGAAGGTGTCATCGAGCTTGGATGGACAGTGGAGGTGGTAACACCAATGGCAAGCACTGTTGAAGACT
TACTGCTTGTGTATGCAGCCATGCTAGGATCACGGCCTAATGATGTTACAATTTCTAAGCCAGCTTTGCCAAGTTTTCCCATAA
TGGAAGGGTCTCAACACGATTTGGATGTTACTAAAGTGATAGGCTCCATTACATTTGGGAAATATACTGAGTGGTTCGAAGAT
GTTGCTAGCTCGGAAGTATTGGATGCCTGCGAAAAGGTTTTGCAGCTTCTTGTAGAGAACTATGGAATCAAGATTAAGGATGT
AATTTTGCCAGAGATGGATGAGATGCGCATAGGTCACGTTGTAACCATTGGATCAGAAATTTCTGCTGCTACTAATACTTACT
ATCTGAACGGGTTGGTTATTGACTCTGGTGCTCTTGCTGTGTCATCTGAGTTTTCTAAGCAATTTTACTTGAATTTACGATCTGT
TAATGTGTGAGAGTGCTGCCTGCTTTGTTACGAGTTCACTTTCACTTTATATGCATGTGGCTTGAGTACAGGAGAAGAAAAGA
TATTTCTTATGAAATCAGAAATACATTAGCTTTATTACGGTCATTTACGTCACGTGATTATGTGGCTGCTCAGCGCCTGCGGAG
AAGATCCATGTTCTATCATATGGAAGCATTCAAATCGGTGGACATTATTGTGACACCAACAACACCCATGACTGCACCTGTCA
TTCCCAGAAGTGCTCTCAAATTTGGAGAGAGCAATCTCGTTGTTGGAGGAGACCTGATGCGATTTGCAGTGGCACCAAATTTT
TTGGGTTTTCCAGCCATCACAGTTCCTGTTGGATTCGATAAAGATGGGCTTCCGATTGGTCTACAACTAATAGGACGACCTTG
GTCAGAAGCAACTCTACTACGGTTAGCAGCTGTAATTGAGAAGTTGTGCGCTTCAAAAAGAAAACAGCCAGCCGTGTTTTAT
GATCTTCTTGAAGGCGTGTAATGAAAGATGCCTTACCAATTTTAGATAAGAAAGGTCTTGTGATTGTAGGCAGCCTTCAAATT
TTTCATTTTTTCGAAGTTTAATCCAACAAATAATTTTGATGTAGTTGAGTTTATTCAGTGTGTTTCATGCATCAAAAAACTGAG
TTATGCAACATGAAGATGGTGGAACCTGTGCACTACACTATTTTTAAAGAGAGTAATATGCAACTTCCACCTACTTCCACTTG
TATACTTGACTACCTTGTAGACTTGTCTTTCCAAATTAACTACATGCAGCTTTCACTGTCTTTTTCAAAA

**PRIORITY IN SWEET LIFE 4 (*PSL4*):**

TTTTTTTGAAGGTGGTAAGATGCCTACATATTGAACCTTTAAATTTGCATTCCTTCGGTATACCAGAGTATTCTGCTTG
CCATGTCTGTATTAGACTTGGCTGACATAATTAAAGATTGTTAAACAGTACAGATGTCCGTCCTACAAAAAGAATCAGCTTTC
GCTTTGCTTGCCATAAAGCGTAGCTTTGGGGATGCCAGTTTCCTCGGCCAAATACTTTCATAGTATTCAGCACTATTTTGAAGA
GGAAAAAATACTCCCTGCAAAACTTTTGGGTGCCCTTCTCAAGCACATTAACCACTCATCATCGATATATTGACATGAAAATA
AAACATTACAATAGGGGGTGAATGTTGTGCATTGAATTACAGTTCATCGTGGGCAGGAACATCGCCTTGATTAAGCAGCCGCT
GTTCCAAAGTCGCTAATCTTTCTTGCAAACACACCGCAGGTGTGGAGAGCTCAGCAACATACTCGCAACGACTTGGCTCATCT
ACATCCCTCAGCTCTATTTTAGAACCACACCTCAACTTCACCTTCAAGCTCCGATCCGGACCATTCCAGCACTTGTCACCCCGT
ACAAAATGCATGCTTTTATACTCATCTTTGAAGCCATCCCAATTCCCTAAGGTAGTGGTTGAATGGGCCTCCACCTGCACCGC
ATGTCTAAAAGGACAAACCTTGTACGTATACTTCTGTAGCCGTAACTCGAAGCAATGGCCATAAAGGCTATAGAACTCTCCAT
CCTTTCCAAAATCTTGCTCCAGTTTCTTCTCAAGATCCGAAATCTTGGTCTGTATGTCGCTCAACTTTGAGGTTGAATCCTTGTA
TTCTTTGCGAATGTGCTCAGCTTCAGATTTATCCACTGGTGCTTTCACAAAAATGCCATCTGAAAATTCCTTGATCTTTTCCTTC
AACTTAGTCCACCAAGGACCTTCCTTCGAAGTCGCATGACCTGAGGAACCAGATGCCTCATCTGTCTCCTCTGCCTCATCCTG
ATCATCTATAACATCCTTCTCATCCTCTTCATCATCTTCATCATCTGTAGAGGCCTCATCTTCATGGTCTGCATCATCATCAAAC
GTGCCTTCGGACTCTGCATCATGATCATCTTCCACTCTTCCTGCCTCTTCTTTCTCAATGTTTGGGTTCTCTCCAGTCCAGCGAG
ATGCTACAAGACGACCTAACTCTTCTTTTGACAAACTCTCCTGGTTATGTTCTGATGAAATGATGTCTTTATCATCATCCAGTG
TCACTTTCTCTTCTTCAAGACCTTCCAGGTGGTCTTCCAAAATCTGATCATCTTTTGCGGGTTCTGTTTTTTCTTCTTCGGTAGCT
TCTTCTTTCGCATCTTTCATCTCTGTTTCTTTTCTTGCCTTCTCTTCTTTCTCTCGCTTCAGCTTCTCGTGTTGCTCAGCCTTTTCG
ATTTCTTCTTTCTTTGCTTTTAACTTCTGAACCTGGTCTTTCAACTTCTTCTCTTCTTGCTTCAACGTTGAGAGATCCACTTTCTC
CTTTGCAATCATGTGTTTGGCATCCTCAACATCTCGTTTCCTAACCTCCAACCCCCCTTTGAATGTCGCAATCTTCCTCTCCAAC
TTCTCCTTCATTACTTTTCCTACTTGAGAGCATGTATTGGAGCACTTTGTCTTCCCATCATACTCATCACTTCCATCACAACAAT
CACAAATGCCATCATTGACTCTTGAAGAAAATATTTCCACAGGTGTATGCCCTTTATTTCTACAGTGGAACTTCCCATTCGGAC
AAGCAGATGTCCCAGGCTCATCAGTCCCATCGACGCAATCACAAAAGTCGTCGTTTAATCGATTCGATTGAAAGCTCTTGGAC
CCATCTTTACACCGGATAGTCGCTGTTTTGTAGTAGGCTTCATCTTGAAGGGCGATGCCTCTGAAGTTGGGGGAGGGCGCAGG
AGCTGCAAGAGTGTGAGAGAGAAGAGAAGCTACCAGACAAGCTTGAAAGAGGAGGGCGAGGCCGCCCATCGCTGCTCTGAG
ATCGGCTCTCCTCGAGCAATGCTCGTAAGCGAACACACAAATCTCACTGTTCCTTAGAGAGAGACAAATAACATGGAGAGAG
ATGGCATGCATTTTTTTTCTTAAGTGAAAATAAAGGACCCAAAGCCGG

**SIRTUIN 2 (*SRT2*):**

GTCAAACCCGAGGTCGCGGGTTCGAACACTGCTGGCTTCCCCCAAAGAGAGAAAGTCTTTAGAAGCTTTAATACAAA
TGTAACTCCCAACCAAGTGACAAAGAGAGATGCGACATACATGCATCTCGTTGCTAATTTTTCATAGGATGGTAATTTTCTGA
TGAATATATAATTTGCTCCCTTATGTTGAAGACAGTCCTTAATGGCAGGAAGGGTTTCTCTCAGGATGATGGCACCATCTTTA
GTTTCAGCTTACAAGCTTTGCAGAACTTTTATGCTGTTTGAGGATCAAAAGCATCATCAGAAGGACCTTATCTGTAGCACAGA
TGTTGCATTGCAGTTGGATTTGCAAAAGAAGCCGATTGTTCCTGAGGCGCCCCCACCGAAACCTGATGAGATTGCTTCTCTTA
ATCAGTTTATTAAAAACAGCCACAAGCTTGTGGTACTAACTGGAGCAGGCATAAGCACGGAATGTGGAATCCCAGATTATAG
AAGCCCACACGGAGCTTATAGTACAGGTTTCAAGCCAATAACTCATCAGGAGTTTATTAATTCTGAACAAAGTCAGAAACGG
TATTGGGCGAGAAGCTATGCAGGCTGGAAAAGATTTATCTCTGCTGAGCCAGGAATAACACACATATCCATTGCGAAGATTG
AGTCAAGGGGATATATTCAGCATATTATTACACAAAATGGTGACAGGCTAGTCACTACAATACTTGGTTGCACCATTGTGCAG
GCAGCAATGCTCTGGAATTGCATGGCAGTACGCACGAAGTTATCTGTTTGAACTGCAGTAATATAACGAGTAGGCATGAGTTC
CAAAAGCAAGTGAAAGAGTTAAATCCAGAGTGGGCATCAGCATTGGAGGCTTTGGAAAATGGCAAGGCAGTCTTAAGTTCTG
GGGTGCAACAACGCCCAGATGGGGATATAGAAATAGATGAGAGATTTTGGGAGGGGAGGTTCCAAATTCCCACATGTCAAG
AGTGTGGGGGCGTCTTAAAACCCAATGTGGTGCTCTTTGGTGCCAATCTTCCAAAATCAAGAACAGAGGAAACACTAGCCAT
GATACGGAATGCAGATGCAGTTCTTGTCTTGGGATCTTCGCTGATGGTTTTATCTGCATTTCGACTTGCCAAAACAGCGCATG

AAATGGGAACACCTTTGGCCATTGTCAACATTGGGGTTACCAGAGCTGATAATTTGGCTTCTTTGAAGATTGAATGCCACACT
GGGGAGATTATGTCAAGAATTGTGGACAATGGGTCTTTAGGACAACGTGCGGCCTGATCGAGGAGAGGGATATTTAAACACA
CAAATACAGATTAAATTAAGCCCAAATAATCATGGGATTATACGCAACTAATCAATGGACTAAATTCAACAAGCTGGATGTT
TGGAAGGCTGTGGTCGATAATTCTCATTATGGATGGTTGTTCTACTGCTCTCATTGCATTTAAGCCTCTTGAATGGCATTCCAC
ATAGAGGTGTTTTGTAAATGTATGGGACCTAAGACAACATTCGTTCTCACATAAAATGAAGAGTTTGAAATTAAGCATACCCA
TTGTCAATTTATTTTTCTTCTTTGCATCAAAATTAAAAAGTGGGCCCAACTGTTTTTGGTTAAACACTCACT

**LAZARUS 1 (*LAZ1*):**

CATAAAAATACTAAAAAGGAATTGTCCATCAAATGACAATTCTTGATTAGAAATGTTCATGATTTTTTTAATGATTAA
AGGTAACCTCAAAAACTACTTGTCAATGGAAATCCCAAATTCTTAAGAGACCCAACAGATTTTAGCATGGAATGCCATACGA
TTAACTATTCTCTGATGCATTTGCAAGGCCAGATTTGTGCACCCACATGTCATGCATTTTTTTGTCTCTTAACTCTGAGGTGTAC
AAGGCTGCAAAAAGCCTGCACACTTTTTCTCATGATTGGAGGCTGATGCATTTGAGGCATGAGAATCACTTTTGGATTAGTAA
ACTTCATTTACACGAAGCAACCAAAATGGATGTATATACATCAAGACTCGTTACAAAAAAGATTTTGCTGTTGTCATTGCACA
GTCCAAACACCTTTGTTTGGAAGCACGTGTATAAAAGAAGATAAAATACATTGTAAGAGGAAGGCGGGAAGACCCATCTTTT
CTTGCTGTTTGTCAACTTGCTCCCATCCCAATTCCAGTCCCTTAATTGTAGTTGGAATTCGAGTACAACTCCATTTCTCTTATGC
TACTTATTCTCATCCTGTGTCTTTGAATAATTGTTCTCTGACTTGAATCCTTAAAACTGCAAGAGTACACCCAATTGACGAGGT
GACTTGCACCCTAGATAAGATCTGGCTCCATCTGGCTTACAAAAAAGCGCTAAAAAGGAGTGGCAGACAATCATTTTGAGTA
TCTTTCATTTACTTGAATGTCCAACGGTGACCCGAAGCATGGTAGTCACCCAACCCCTGATCACTACTCTCCCCACTGCTATCT
GCTGTCTTATAGGTGTAGGTGTATTTCTTTGTTTTACGCTTCTCAGCTCCACTATCACTCACACTTCCAACCAGCGAAGGGTCA
TCAATCCCTCTTATGGTTGGAATCCAGCTGTCATCTTTGACTTGCTTCTTGTGGTCCCTATGCATGGTTGCCTTCAAGCGACCA
AATCCTTGCCTAACTGGCTCCTGGACTTTTCCTACAGTAAATTTCACATCGTTTATAACATATTCTCCACCTTTGAATACAAGG
TCATGCATGCTTTCCTTCAGACTTGTTCCTATTTCTGCAACCTCCGAGTACGGGAGCCGTTTGACAGTAGGCCTTTCAATATCT
TTGACCTCTTCGGGATCAGGAGGCACACCAATAGCAGCATAATCTGACATAACAGAAAGGCTTCCACGCTTTTGTGAGCCCAT
TAGTTGATAGGGTTTTGCTGGCAATACATAGACATGAGCAACAGCAGCAAATCCCATCTCTATGCAAATAAAGAAATCTTGTA
TTCGAGCTTCTAAAGATTCTTCATTTTCGAACCTGGCTGATTGCATAGCACAGAAAATGGCTATTAAGAGACCCTGCCACCAT
GTGAGAAAAACAATGGACTTAAAACATAAGAACTTTGATAATGGTTGGATTGGTGCAAGTTCTTCTTTTGTGACGTTGTAGAA
CTGCATCAAGCAATATAGTGCCCAAGATTGACTCAAATTCATAACCACAGCAACGTAGAGGTACCCCCTGTTCCATTTAAATT
CTCCTTCTCCGTAAAGATCCCAGAATTGTAAAATCAATGCAACCATAGCACTGACAGTCTTCATAATCATAAACTGAACAATG
GCGAATTTGAATGCTTGATAAAACTGCCTCCCAAGATCCCAAGGCTGCAAAATATAACTGAGTGGAAAGCTATGTTTCACAG
CGGCTCTTCCGGCAGTCTGGACAAGAAGTGGAGTTCGAGGGCCTGCAACTGCTTGCCTTTCCATTGATTCCATTGTCTTCTCTT
CCCCACCTAGACAGGCAATGAGATAGCGCCCAAAGGAATACAATGCAAAAGCTTCGTAGCAGTCTCGTACGATCTCGAACAC
GAGAGCTGCATGTGAGCTGCATAACGACACATAAGATTCTATGGCATACAATGGAACTATCAGAACTACTCCTATGATCCATT
TTTGCTCCTCTGGATTGTTGTACCAAGAAAGATGTTCGAGTATCAACAAGGTTGAGAGTGCAACGGCAAAGAGTACAAAAAT
GCCAGTGAGTTTCCTGGCATTAACTGAAATATAGATGCTTTATATCTCTTCCCTGCCAAAAAAAATAGCAGAAGGTCAATCTT
ATTACTGTTGGAGCACAATCTTCTCAATTAGTCAAGCTACCCATCAATCATAAATGTTTCTACTGTCACAAACACCTTACTTA
TAAACTCCGTCTTCTCTTCAGTAGTCAACACAGATAGAACAAAAATACGAAAAACCCTTGAATAATGCGCACGACGGGCCAT
GTAAAATGGCTTTCTGTGCACGTGTTTACAAATGGACGCATGTGCAATACGTCTGTACAGCAATTCCATGAAACATGTTTGTG
AAGTGCGCAATAGTAATGAGGCAGAAACTGAAAAATTTTACCTCCAGTAACTGTAGACAAATTTCTTCTACTAAAAAACCCA
ACCTGAGAGAGAGAGAGAGAGA

**STROMAL CELL-DERIVED FACTOR 2-LIKE PROTEIN PRECURSOR (*SDF2*):**

ATGCCCTCTGCAGAATCTCATGATGCTATCAGAGAACAAAGGGAAAGGGGCCCCTTTTTCATAGGGAAAGTGTCCAT
AACTTCTTGGGTGGATTGTAGAGCTTATCTAGCCGCCACAAAGTTGGTAATTAATAGATGATAGACATTGTTTTACAAACCTT
TTGCCACTTTAAAAGGTATAAACATGATTGAAACGACTATTGTATTGTATTAATAGAGAGCACACTTCATATTGTTTGAGCTG
TTTAAGAAAAGTTTTATGGCCTCACGGTGAATCACCCAAATAATGGACATAGTGGGATGATTCAACCCAAATCTAAGTGGAT
GTGGAAATGGGCTCAAGATTTTACTTTGTCAGTTTTCATGGACATAGTGGGATGATTTAACCCTTCCGCCGCTTGGACCCCTCC
CTGAAGTGGGACCTTAAAACGTTCTACCCAAGATACAATTGCAAGGAGCCCACCGTCACCTCCTTCCACTGCTTTAATGATTC
TATGATTGTGTTTTATGATTATGTATCAATGATTCTATGATTATGTTTTATGATTGCACTTGGTCACAAGCTTGTGTGTATATTC
ACTCAAACTTACATGATGTCAGCACGATGATGTTCAGGTAACCTATGGAAGTGTGTTGAAGTTGATGCATGGGACCACTAAG
AGAGGTTTGCATTCCATTGACATATGATACAAAGCAGGGGGCCCATATCATGTTGTTACTGGTTTCAAGGAGTTTGTAACTCC
AATAGCTATTGGGTATGGAGCTCTTCCCCTTTCAATCCTAGTTTCACCTAGAATCACTTTATTGTCAAGTAGATCAGCTTTGTT
AGGTGGGTTTCTTGTTTTACATCCTCAAATTTTATTTCTTATTTGAGTTACTTGAAGTTATTGACTTGCAATAAAATGGCTTATT
GTACACACTATAGCCGGGAGCTTGTGAGTGTAGGAGGTTTACATGCATGTAGCAGCTTAGGAGATGTAGAAAAGCACAACAT
GTGTCGTAGCCTTGAAAACACACCCTCTCATTGCATTCTGTTGGTAAATTAGACAATCAAGAAGATGGGCCATGTCATTAGAT
TATTCTTTAGACACCCAAATTTTCTATTCAGTATGTGTCTTTTATGTACCCTTCTCTTTAAATTATTGGTATAAAATACACATGT
TGCACCTACATAAGTGGAGTTTCATGCTGCTACCTTTGTTATACATTAGAAAAGTGTTGATGACAATCTGCCGCTTAGCCCTTA
CAAGAATGAAGGACCTCCCAATGCTTTGCAGGCTGCCCCAATAAAGCATCAAAACTATGATGAACCACTACTTCAGCTTGGA
GGAAAATGCAAGAGGAAGGTATTTGCCAAACATGGTAACAATTGTCAGCATTTTCAATGCATGTGCAATCTATTTAGCTCTTC
TTTAAGGTGATTAGATACATGCCAAATTGCGATGTTAAAATTTTATTTAGTCCTGCAATCAAATTTATTTTCCCTTCAACACTA
GTACTCAAACATAGCTCCTCTCCACTTAATCATATTCCATATGACTTCTAATTCTATAACCATTTTACCTATGTATTGTCTCTTG
CAATAGGTTGGCGATGCACATGCTGATCATCAATCCCAAAAGCCTGATAACACCATGCACAACTGTATAGATCAACACTTCTC
GGCTCAAAACCAAAGTTGCTAGTGAGACTACTTTTGCTACCCTCGCTTCTTCAAGGGAATCAACATTGATTACTTTGCACTTCT
GCTCAAAAGGGCCGTCATGGTACTCATGGCGTCAACAAAGAGAAGGGCAAGGGATGCATCCAGCGGGAAGTTTGCTGGCCCC
CAACAGGATCCGCCAAAACACTCTCAAGGACGCCCAAAAGTAAAAAAAATCCCGAGGACAGCAACC

**CELL DIVISION CYCLE 5 (*CDC5*):**

GACCACATTGGTTGTGTGTGTCAAGGACACTGCAGTCATACTAAGATACCTTAAAAAATTTACAAGGATTTATTGATT
TTTTTAACACGCTCATTCCCATTGATTTAGCCATTATCGCATACTAATCTTATACTTACATGTGGCACTCTCATGTTTGATTCCC
TTATGTATGTGTCGAGGTTACAAGGTTTAAGTATTGAATTACACCATCAAAGTAAACACTTTTTCAAATTTTTAACCCATTGAG
ATTTCCGAAGCAATGACCATTGAGAGGACCATTTGGGTGGTACTATCTGAACCAAGAAGGCTACCACCTCCCATAGAGCAAG
AGAGAAAGGACATCTAGGAAGCTTGGAGCCTGAGGGTATAGAAGAAGAAGCAATAAGGCTTCAGATCCAATTGCAAAATAT
GATTCCAGTGAGCCAGAATGATGGTGGAGAAAATTAAAGGAAATCCTTAGCATGTAAGTTTCAAGACCCTATGGTGTTGCCT
CAACCTCTGCTGCTGAGCGTGCTGCCAGCTCAGCCTCTGCCATCACCTCTGCCTCTGCATCTGTACACGCCACCACTTCCACCT
CCTCAACCTCTTGTGCTGCCACCTCCGCCACTGCAGCTGCACTCGTCATCTCTGCCTCTGCCTCTGCAACTGTGCACACCGTCA
CTTCTGCGGCTCGTGCTGCCACCTCTGCCTCTGCAGCTGCACTCGTCGCCTCTGTGTCTGCAACTGTGCACGCCACCACTTCCG
CCTCCTCAACCTCCGCTGCTGCATGTGCTGCCACCTCAGCCTCTGCAGCTGCACACGTCGCGTCTGCATCTGCAGGTGTACACG
CCGCCATTTCCACCTCAAACTCCAGTGCGGCTGTCTCAGTCTCTGCAGCTGCACACGTCGCCTCTGCCTCTGCAGCTGTACACA

CCTCCATTTCCACCTCCGCTGCTGCAAGTGCCACAGCCTCTGCCTCCGCTGCTGCACGTGCCACAGCCTCTGCCTCAGCTGCTG
CCTCCGCTTCAGCAGCAGCACGTGCCACTGCAATATCATGTTCTTGAATAAGTTTCCTCCAATTCTCCCGTTCAACAAGCAAGT
TTTCATATCTTAGCTGAAGCTTGTGCTCATTCTCATTCTGCTGCTTCACTGCTGCTTGCAAACTCTCCAAGCGAGACGGTGCTG
CAAGCTGCTCCTGCTTAAACAGCAAGCGGAAGGACTCCAGTTCTGTTGCTAGAGTGTCAGCATCTTTAAACAATGCCTCAATT
TGCCCCCACAAGGTTTCTGCCCTGACCTGATAACCTTGAGTGAGTACTTTCAGCTTTTGCTCTGACCGGACCGCTTTCTTGGTT
TCCCCTTCCATGTGTTTCTTAATGATCTCAAATTCATATTGTAAAGCAGATAATCTTTCGTTGTTGCTTGCCACACTTGCAGAGT
TATATGCTTGTCGTGAAGGGAAGTACATCATATCCTCAGCATGAGCATCCCTTGCCTCTGAGTAATCATCAATTGTCGCAACA
TCATGCCCCATTGCCAATCGCAAGAAGTCCGACTCTTCTTCAATCATCATAGCTGCATCTTTCAGCTCCTCATCTTCAAAATCG
TCGAGTACAGGTATTTCAACTTGCTTTTTGCCATTGGCCACGTTCTTACTGGCCTTCTTCTTCTCCTTTCCTGATAATTCCTCCA
AAGGATATTTCGCATTATCGTGCTCCAAAAGAGCTGCCAATTCCATCTTTATTAACTTATCTGCCTGTTCCGTCAATGTTGTTG
CTACACTCTCTTCAGCCTGAGCCAAAGAACTCTTCAACCATTCAACTGACACTTGCGGAGGTCGTGGAAGGTCCCTTTGAAGG
ACTTTGGACCGCTTTAAAAGAAGGGCAGCTTGCCTTGCTACTTCTTCTGCTCGATCCCTTGCAAGCTTATCAGACATATCTTCT
TCCACAGCATCCACAGCCTCTGGTTCGTCTGGTAACAAATCACGCACAACAACTTGGTACTCATATTTAGGCACAGGAAGATC
ACCCAGGCCAGCCCTAAGGTTCCTTCGTATCTCTGCCTGCCTTGCTTTCTCAGCCTTGACACTTTCATAAGGGGTTTCCAATCC
TTCATTGATGTGAAGTTCGTCCCTGATTGGAGTACCCTTGGGAGTAGCATAGATGGAATCCTTGATGGGTGTCATTCTTGCTAA
AGGAGTTAAGCCTACCCCACCAGAAGGGGTTTGAGCTGGAGTAGCAATCGGATTTGGAGTTTGTATTTCCCTCTTTTTGGGTG
TCACTCCAGAAAAATCCCATGGATGCAAGTCCGGGTTCTCACCTCCAAGAAGAGGAGTCTGCACTTGCCTAAGCCGAGCTAA
GTTCTCAGCCTCCATCATGATAGCGTCACCTTTCCCTCCTGGAGTTCTTTGAGGAGTTCTGAGTGGTGTCATGCCAGTTCTAGG
AGTTTGCCCATAGTTTGCAAGCAGCATACGAGTAGCCCCACTTCCATCCCCAAGCTCATCATCGCCTGTAAGTAGATCATTAG
AATAACCCATCTTTGCTATCTCCTCCAGCTCACGATCTGAAATCTGAGGAGGAGGAAGCATAAGCTTCGACCTCTTACGAACA
GCCTCTGGGTCATTTAACCTGTTGATTTGCATTACAGAGGCTGGAGCATCCCTGCGCTCCGCAATCTTGTTGTGAGCGACATCC
TGCTTCCTCAACTGCGCCTCTACATCAACTCGCCTCTTCCCTTCCAACTCTTCAATGGTGGTTGGAAATTGAGGCTGTTCTACA
ACCATATCTTCCTGAGATATGTCAAAGAAACCTGGGGGAGGCTTCTTTTCAAAAGGAATTTCAGCATTGTAGTCAATGCCTCT
TTGTTTTCTCCTTCTATGCCTCCCCTCGATGCCTGCTGCTTTAAGCTCGCGGCGCTTTTGCAGAGAAGCGAGTCTCCTAGCTTCT
TCAAGCTGCTTCTCCCGAGCTTTCCTCTTGGCCTTCTTTCCGCGAGTATTCGCAAGCCTGGCCCTTGCTTCAGAAAGCATTTCTT
TTTCATCTTCATCCATGTCGACAGGGTCAGGCCTAGCGGGTTTGGACTCAGGGTTTGGATCAATTTCACCAGGCCGAAGTTTC
CTTGGATCATCTGAAGGCTCGTAATTCTCATCCCGAGCACAAGCTGCATCCAGCAATTTCTCGTACCGTTCTAAGCATTGTGCA
GGCGTGCGACCGACAATGGGAGCTATAGTTCGCCACTGAGTAGGCATGAGCTTTGCAAGATGTAAAAGCTTCTCATCTTCTTC
CCGAGTCCATTCCGTCTTTTTGATGGATGGATCCAGCCATTCATACCATCGTGCTTTGCACTGCTTTGCCGATTTTCGGACAAG
CAGAGATGATATACGAGCCCACTGATTCTTGCCGTACTTCATGACGGCAGCCTTCAGAATCTCATCTTCTGTGTTCTTCCAAAC
ACCGCCCTTTATCATGATCCTCATCTTGCCCTGAAAACCCTGAAAAGGAGCTGTTGTTTAAGAAGCCTCTGTGATACACAGGA
AAGATGAGCCTCGCAATGCAGGCAAGACTGCGTAGTTACACGGTTGACCTAAAGAAACTGAAGGCACAAGATAAACTACAG
GCACGCCGTTGCTTCGCATACGGCTTCGCAGGTAGTGAGACATTATGAAACCCCCTAAAACCCCCTAATTGAG

**APETALA 2 FAMILY PROTEIN INVOLVED IN SA MEDIATED DISEASE DEFENSE 1 (*APD1*):**

CGAGATTTGACAATTAAAAATGTATGCACATGCATTGAAAAGCTAGACATCCGCCAAAGCGAAACATGAAGTAGACA
AGCATGGCACATTAAAGTGAGCTTCTCAATTGCTCTCATACCACAACACACCACCCCTCATATCAATGTGAAAACAAATGAAC
AGATCTTGCGAGCTTCTTGCTTTCTTCATCTAAGACGCGGACAATGTCCTAATGAACAATAGTATATACTTTTGAAGAACTTGC
AATCCTGCTACCTCATTACTACCATACTTTTTACAGCATTTCATACTGCGCCCCTCCGAAATCTGGTCATACCTCGTGTTCCAA
ATTAAAAGGTGGCATAAGAAATTGCTCCATCTCCTCCTAGAAGCAAAGCAAACAATTAGAGGACAATGAAATCCCAAATCCT
TGCTTGCTTTTGGACACTCCCTAGCAGTTTGTTAAGTGATCCATGTGAATGCTCGACCCTCCTTTCTAATAAGGGGGTCTTGAG
CTAATAATGGTTTCCAAAACTTGGCAACTTTGTTTGCTAAGAGCTGTGGAACATCTGCTCATCTCTGGAAAACTAGTAGACTTT
ATAGCTTGCTTCGATTGGGAGTACGCCAATCAATGAACACGTGATATCACTGTAAGGATAATTAAGGATAAGGACTGTTGCCT
GCAAAGGCAGAATGCTGGTGAAGAGGAGGCGGCAAATAAGATTCTGACAAGTAGAAAGGTGCTGTCTGGTGAAAAGGCTCA
AAGGGTTGTCCTTGCTGCTGCATTAGCTTTAAAGCTGCTGATTGCTCATAAACAGAAGTGGGTGGTGAATGAGCCTGCTGAAA
GGATGCCAAAGTTGCGGCTTGGTGAAGAACAGATGCAGTTTCATCAAAAGTTGATGTACGCTGATACGCAGATGAGGGTTCA
TCAACAGATGCCATTTGATGAAAGGGTGGTACTTGATGGTACACCGATGCAGGTTTGTCATAACATTGAGACTGATGAAAAG
ACGATAAATCAATTCTTCCATCTTCAAGTAGCTCTGAAGAGCGCCGCCTCCGTACACAACTCTCTACCCTCTTCTGCTTCTTTT
TGTTTGCAATAGCTTCTTTTGTTATTTGAAGAAAATCCTCCCACTGAAGTCCCTGTAGCTCTTGCTTCTCCTCCTCAGAGAGTTC
AAAATTTGGTACTCTTCCGCAAAGGTAGGCAGCCCTGTCATACAAATGAGCTGCTTCTTCCATAGAAGCAACTGTGCCCAAAT
GTATTTGCTTTTTGTCCACCTTTATTGCTGCTTGCCACTTCATATTTTTGAAGTACACACCTCGTAGCAAACAAGGGTCCTGATT
CTCTGATTGCTTCCTTCTGGAAGCTTTCCTCAGCTTCAGTGCATGATCTTGAAATGCATTGTTTAAGTCCTTCATTGAGTTATCA
CATACAGAAAAGGAAGGGCTGGGTTCCGCTGCAGGAACCAGGTTTTTCTCCTGTCGTGTCTCCAAGATGCTCGATTCACATTT
CTGGTAGGGAATCTCTTCAACCACCTCCTTGCGCTCCATTGCACGTTTTTGCTCATTGAGTCTTGTAAGTTGATTGAGTATTTTA
TACCTTTCTTGTGCACCGAGGTGGCGCCATTTTCTGATGCTCACCATTTCACAAATGTATTGGCAATTCCAGAAACAGTAGTTC
TTTGGAGCCTCCAATTCTCAAGCAACTCTCACAGCTCGGAGTAATGAATGTAGTTACATCTATATATGAGGTGTTTGAGAACT
CTTTAGCAACCCCTTTACAAGCTAGTTCTTCCAAATGTTCTGGCTTCCAAGTCACAACCTGCTTGCTAATCTGTGAGAGCTGGT
CGGCGTCAAGGCTGCCTACCTTCTGGAGCTGCCGATCTTCGTCCCCCCCGCCAAACAGAAAATGCTCGCCACCTAGAGAGAGT
GTACAACGCAAATTTTG
